# Genomic insights into southern white rhinoceros (*Ceratotherium simum simum*) reproduction: revealing granulosa cell gene expression

**DOI:** 10.1101/2023.06.27.546732

**Authors:** Elena Ruggeri, Kristin Klohonatz, Marc-André Sirard, Barbara Durrant, Stephen Coleman

**Affiliations:** Reproductive Sciences, Conservation Science Wildlife Health, San Diego Zoo Wildlife Alliance, Escondido, California, United States of America; Select Breeders Services, Chesapeake City, Maryland, United States of America; Département des Sciences Animales, Université Laval, Québec City, Québec, Canada; Department of Animal Sciences, Colorado State University, Fort Collins, Colorado, United States

## Abstract

*In vivo*-collected granulosa cells (GC) from the southern white rhinoceros (SWR) provide a non-invasive assessment of the developmental status of oocytes prior to *in vitro* culture, which could aid in the development of assisted reproductive technologies (ART). Our study aimed to investigate gene expression in SWR granulosa cells, collected *in vivo* and gain preliminary insight into the transcriptional activity occurring within the cells during various stages of oocyte development. It was hypothesized there would be similarities between the SWR GC transcriptome and cattle and humans, two species for which well-annotated genomes are available and ART are commonly used. GC were collected from SWR following ovum pickup (OPU) and pooled from all aspirated follicles. Total RNA was isolated, libraries prepared, and sequencing performed using an Illumina NextSeq 500. Reads were aligned and annotated to CerSimCot1.0. Databases for cattle and human were acquired for comparison. This study identified 37,407 transcripts present in GC of SWR. It was determined that cattle and human transcriptomes are valuable resources with a homology of 45% with the SWR. In conclusion, these data provide preliminary, novel insights into the transcriptional activity of GC in the SWR that can be used to enhance ART in this species.

## Introduction

The southern white rhinoceros (*Ceratotherium simum simum*; SWR) is a useful reproductive model to study its subspecies (1), the functionally extinct northern white rhinoceros (*Ceratotherium sinum cottoni*; NWR). Substantial work has been done to evaluate the reproductive physiology and estrous cycles of four rhinoceros species (white rhinoceros *Ceratotherium simum*, black rhinoceros *Diceros bicornis*, Indian or greater one horned rhinoceros *Rhinoceros unicornis* and Sumatran rhinoceros *Dicerorhinus sumatrensis*), but there is no similar cycle length among them (2). Acyclic and irregularly cycling SWR have been reported in captivity (2-5), with cyclic females exhibiting estrous cycles from 20 to 80 days in duration (6) frequently categorized into two cycle lengths, long (clustered around 70 days) and short (clustered around 30 days), but to date there is no mechanistic explanation provided for the differences (3, 5, 7-11). This unusual pattern of irregular estrous cycles potentially complicates the development of ART in the SWR. However, regardless of length, the SWR cycle has been manipulated with GnRH to induce ovulation when follicles reach 30-35 mm (5, 6, 12). GnRH treatment is used to obtain high quality oocytes and achieve higher pregnancy rates in cattle, and this treatment may be successfully applied to the SWR (13, 14) to improve *in vitro* embryo development. To understand SWR oocyte development, it is imperative to describe the *in vivo* follicular environment.

Recently ovum pick-up (OPU) has been successfully performed in the SWR (15, 16), but the mechanisms of follicle growth, oocyte maturation, ovulation, and embryo development are only partly understood. *In vitro* maturation and blastocyst rates in this species are suboptimal compared to other domestic species and humans (17-21). Ovarian ultrasonography has been done routinely in very few SWR (12). The technical challenges of repeated ovarian ultrasound to closely follow the estrous cycle and limited opportunities to perform OPU result in incomplete information on the species’ reproductive physiology. Reliable data on follicle maturation and atresia, ovarian stimulation, oocyte maturation, and ovarian aging are needed in this species to improve *in vitro* techniques.

Genomics are a useful tool to elucidate cellular function and physiology (22-25) and to address reproductive questions, from oocyte quality and embryonic development to postnatal testing (26-28). In numerous species, including cattle (*Bos taurus*, (29-31), human (*Homo sapiens*, (32, 33), horse (*Equus caballus*, (34), pig) *Sus scrofa domesticus*, (35, 36), and mouse (*Mus musculus*, (37), the transcriptomes of granulosa cells (GC) have been evaluated to address questions about ovulation and oocyte developmental competence. Granulosa cells undergo dynamic changes during folliculogenesis, culminating in releasing a competent oocyte and forming the corpus luteum (29). Distinct gene expression profiles during ovulation and luteinization processes can be detected, allowing insight into the path of oocyte developmental competence. Therefore, it is valuable to study granulosa cell transcripts and generate their transcriptomic signature to elucidate unknown aspects of ovarian physiology in the SWR. Novel insight into granulosa cell gene expression and function during follicle development has the potential to improve reproductive management of the SWR and develop successful *in vitro* technologies by highlighting the processes and pathways being utilized by the GC during follicle development.

There is limited availability of publicly accessible transcriptomic analyses in the domestic horse, which is considered the closest model for the rhinoceros. *Equus caballus* (EquCab3) genome has been recently upgraded in annotation (38, 39), but only one microarray study has been published on the granulosa cell signature (34). In addition, a complete sequenced database is not available for the mare to elucidate granulosa cell expression profiles. Thus, data publicly available in established species such as cattle and humans (30, 40), are better resources for investigating granulosa cell expression profiles in the SWR. In addition, the incomplete and partial annotation of the SWR genome negatively impacts the assembly of a reference data set to compare expression data. Therefore for the most accurate reference, a newly assembled NWR genome and annotation, can be used (41), but manual curation is still needed and an intensive comparative analysis will help profile GC expression.

The current work is the first transcriptome-wide analysis of granulosa cells in the southern white rhinoceros. The aim of this study was to investigate *in vivo* GC to provide a preliminary insight into the transcriptional activity of these cells during follicle development. It was hypothesized there was a similarity between the SWR GC transcriptome and two species with well-developed assisted reproductive technologies, cattle and humans. Acknowledging limitations in the number of individuals and genomic platforms for the analysis, this preliminary report described expression profiles in GC in the SWR and compared that profile to established GC data from cattle (30) and humans (40). As previously shown in other species (26-28), these data can improve the understanding of ovarian dynamics and oocyte competence acquisition in the SWR, as a template to describe the transcriptomic signature of GC cells. The purpose of this study was to provide the first insight into the SWR granulosa cell transcriptome, to begin to understand the unique aspects of rhinoceros reproductive physiology and to generate a tool to optimize assisted reproductive technologies for this species.

Defining the reproductive physiology of the white rhinoceros and developing assisted reproductive technologies (ARTs) is an essential strategy to rescue the nearly extinct NWR (42). These techniques could be applied to other rhinoceros species and also have a wider impact for other endangered species (16). The use of ARTs (gamete collection and cryopreservation, artificial insemination, *in vitro* oocyte maturation, fertilization and embryo production, and embryo transfer) will aid in increasing genetic diversity and maintenance of *ex situ* populations.

## Results

Across all samples (n = 6), 37,407 transcripts were detected, which corresponded to 11,258 genes, with some genes expressing multiple transcripts (Supplemental List 1). For the remainder of the data analyses, genes, instead of transcripts, were utilized for expression information to avoid overrepresenting genes that had multiple transcripts. The biological pathways with the most gene representation in these samples were determined for the genes that were expressed in GC. These pathways included: the centrosome cycle, mitochondrial translation, DNA geometric change, nuclear-transcribed mRNA catabolic process, and mRNA splicing via spliceosome. The top 20 most represented biological pathways and the pathways associated with reproduction were also identified (Table 1).

**Table 1.** The top 20 biological pathways and reproduction-related pathways associated with transcripts identified in southern white rhinoceros granulosa cells.

| Top Rhinoceros Biological Pathways | Reproduction Related Pathways |
| --- | --- |
| Centrosome cycle | General transcription regulation |
| Mitochondrial translation | p53 pathway |
| DNA geometric change | Apoptosis signaling pathway |
| Nuclear-transcribed mRNA catabolic process | Wnt signaling pathway |
| mRNA splicing via spliceosome | DNA replication |
| rRNA processing | CCKR signaling map |
| DNA-templated DNA replication | General transcription by RNA polymerase I |
| Spindle organization | Parkinson disease |
| Negative regulation of translation | Transcription regulation by bZIP transcription factor |
| Regulation of double-strand break repair | Gonadotropin releasing hormone receptor pathway |
| Ribonucleoprotein complex assembly | Ornithine degradation |
| Microtubule cytoskeleton organization involved in mitosis | Hedgehog signaling pathway |
| Vacuolar transport |  |
| Phosphatidylinositol biosynthetic process |  |
| Macroautophagy |  |
| Double-strand break repair |  |
| Positive regulation of autophagy |  |
| Cell cycle checkpoint signaling |  |
| Vacuole organization |  |
| Protein stabilization |  |

The transcriptional profile of SWR granulosa cells was compared to the GC transcriptome of other species (cattle and human; Figure 1) across all follicle categories. Humans were the most similar, with 48% of the transcriptome in common with the SWR (Supplemental List 2). The pathways with the most representation associated with these genes were: mitochondrial translation, centrosome cycle, mitotic cytokinesis, nuclear-transcribed mRNA catabolic process, and mRNA splicing via spliceosome. The top 20 biological pathways from the genes in common are listed in Table 2.

**Table 2.** The top 20 biological pathways associated with the genes in common between the southern white rhinoceros and the human or cattle. Pathways in bold are in common between both comparisons.

| Top Rhinoceros and Human Biological Pathways | Top Rhinoceros and Bovine Biological Pathways |
| --- | --- |
| **Mitochondrial translation** | Positive regulation of transcription elongation from RNA polymerase II promoter |
| Centrosome cycle | **Mitotic cytokinesis** |
| Ribosomal small subunit biogenesis | Histone ubiquitination |
| **Regulation of transcription elongation from RNA polymerase II promoter** | **Mitochondrial translation** |
| **Mitotic cytokinesis** | Regulation of ubiquitin-protein transferase activity |
| Nuclear-transcribed mRNA catabolic process | Protein localization to endoplasmic reticulum |
| mRNA splicing via spliceosome | Nuclear envelope organization |
| rRNA processing | **Post-Golgi vesicle-mediated transport** |
| **Protein monoubiquitination** | Neuron death |
| DNA-templated DNA replication | **Regulation of transcription initiation from RNA polymerase II promoter** |
| **DNA duplex unwinding** | Positive regulation of DNA-templated transcription, initiation |
| **Post-Golgi vesicle-mediated transport** | **DNA duplex unwinding** |
| **Endosome organization** | **Protein monoubiquitination** |
| Spindle assembly | Regulation of translational initiation |
| Mitotic spindle organization | Lysosome organization |
| Endoplasmic reticulum organization | Chaperone-mediated protein folding |
| tRNA modification | Phosphatidylinositol biosynthetic process |
| Telomere maintenance | Negative regulation of chromosome organization |
| Double-strand break repair via homologous recombination | **Endosome organization** |
| Regulation of double-strand break repair | Vesicle budding from membrane |

**Figure 1.**
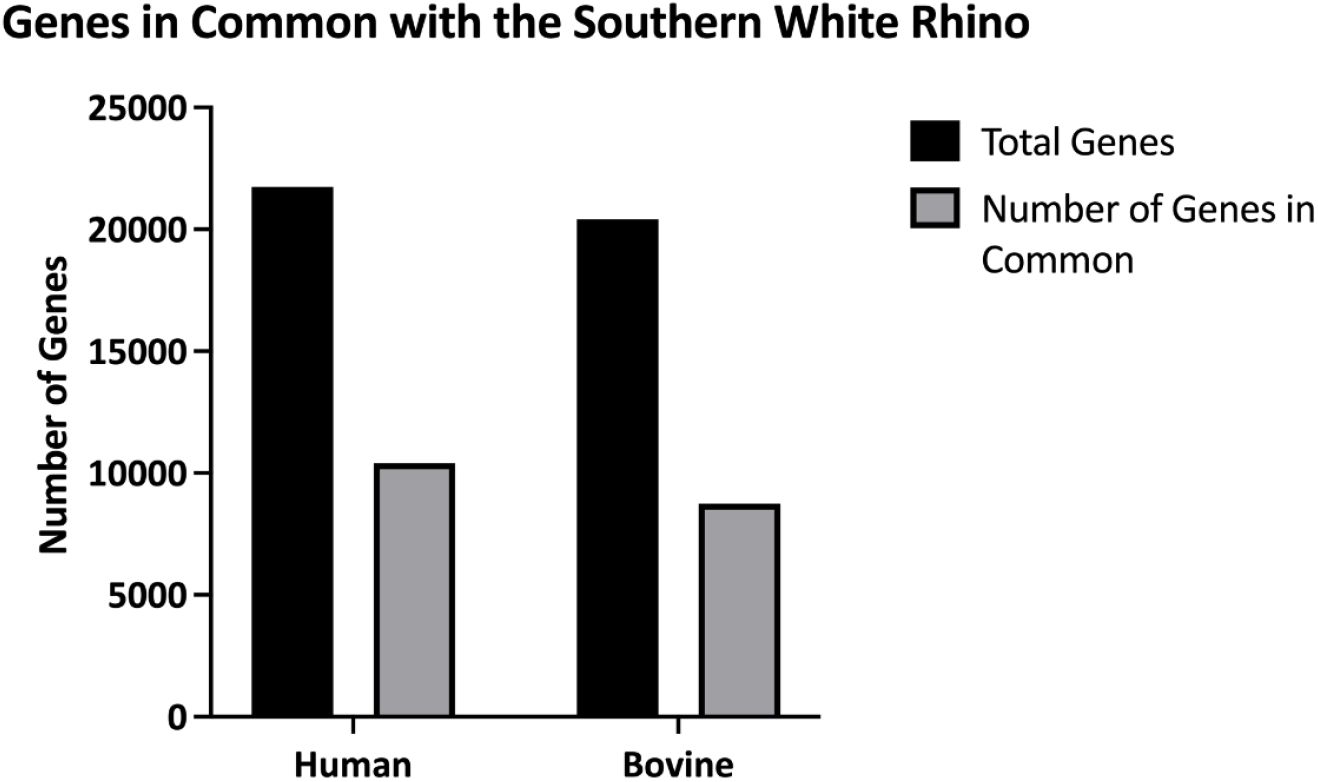
Expressed Genes in Common Between southern white rhinoceros, human, and cattle granulosa cells.

The cattle genome has 43% of the transcriptome in common with the SWR (Supplemental List 3). The top pathways associated with these genes were: positive regulation of transcription elongation from RNA polymerase II promoter, mitotic cytokinesis, histone ubiquitination, mitochondrial translation, and regulation of ubiquitin-protein transferase activity. The top 20 biological pathways from the genes in common can be found in Table 2.

Across all three species, there were 6,935 expressed genes in common (Supplemental List 4). The top biological processes for the genes in common differed from analyses when the species were individually compared. The top pathways for the genes in common between all three species were: Golgi to plasma membrane transport, nuclear-transcribed mRNA catabolic process, deadenylation-dependent decay, protein localization to the endoplasmic reticulum, and fatty acid catabolic process. The top 20 biological pathways from the genes in common are listed in Table 3. In addition, further evaluation of the pathways specifically involved in reproduction from the list of shared genes revealed 11 pathways (Table 3).

**Table 3.** The top 20 biological pathways and reproduction related pathways associated with the genes in common between granulosa cells of the southern white rhinoceros, cattle, and human.

| Top Overall Biological Pathways | Reproduction Related Pathways |
| --- | --- |
| Golgi to plasma membrane transport | General transcription regulation |
| Nuclear-transcribed mRNA catabolic process, deadenylation-dependent decay | p53 pathway |
| Protein localization to endoplasmic reticulum | Apoptosis signaling pathway |
| Fatty acid catabolic process | Wnt signaling pathway |
| Negative regulation of protein phosphorylation | CCKR signaling map |
| Hexose metabolic process | General transcription by RNA polymerase I |
| Microtubule organizing center organization | Parkinson disease |
| Phosphatidylinositol biosynthetic process | Transcription regulation by bZIP transcription factor |
| Regulation of protein serine/threonine kinase activity | Gonadotropin releasing hormone receptor pathway |
| Ribonucleoprotein complex assembly | Ornithine degradation |
| Cellular carbohydrate metabolic process | Hedgehog signaling pathway |
| Ras protein signal transduction |  |
| mRNA splicing via spliceosome |  |
| Protein glycosylation |  |
| rRNA processing |  |
| Translational elongation |  |
| Protein folding |  |
| Small molecule biosynthetic process |  |
| Vesicle fusion to plasma membrane |  |
| Cellular component disassembly |  |
| DNA repair |  |

Using the cattle database (30), the SWR transcriptome data were compared to cattle GC data from follicles in different stages (growing, plateau, and atretic based on follicle size in cattle). From the cattle data set, there were 5,722 cattle genes specific to this group of follicular stages. Of these 5,722 cattle genes, 3,388 were present in the SWR. The SWR dataset had the most genes (1,570) in common with cattle GC from follicles in the growing phase, representing 14% of the genes in the SWR transcriptome. 1,239 genes in the SWR were in common with the transcriptomic signature of GC cells from atretic follicles in cattle, representing 11% of the genes identified in the SWR. Only 579 SWR genes were common with cattle GC cells from follicles in the plateau phase. These 579 genes represent only 5% of the genes identified in the SWR (Figure 2; Supplemental List 5).

**Figure 2.**
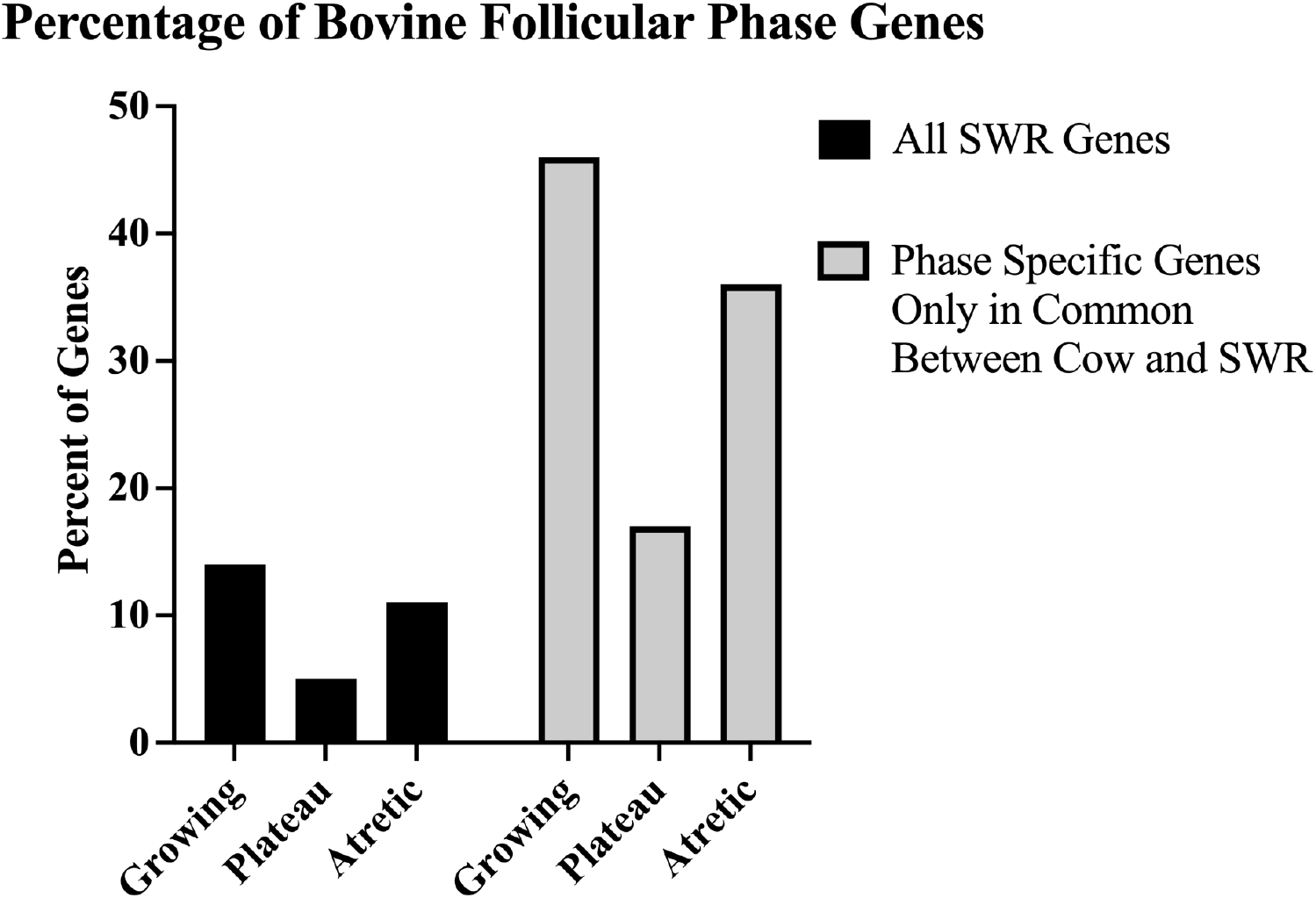
Percentage of SWR granulosa cell genes corresponding to bovine granulosa cell genes specific to the growth, plateau, and atretic follicle phases.

## Discussion

The aim of this study was to generate the first transcriptomic analysis of granulosa cells (GC) in the southern white rhinoceros (SWR). Although this is a preliminary study, this dataset provides new information useful for future studies in the field of reproductive physiology in the SWR. In addition, this dataset will continue to improve as more OPUs are performed, adding to the pool of granulosa cells used for sequencing. Due to the low number of individuals included in this study, the overall goal was to evaluate the transcriptomic profile of granulosa cells in this species. While an SWR genome is publicly available (CerSimSim1.0; GCA_000283155.1), there are gaps in the sequence and partial reference annotation. Because of these limitations, the new northern white rhinoceros (NWR) genome, CerSimCot1.0, was used to align and annotate of these data. The NWR is the closest relative to the SWR, allowing us to use its genome for this analysis. Due to the nature of a new genome assembly, additional manual curation was required to identify all annotated genes and transcripts. This additional step and analysis resulted in an overall improvement in the number of transcripts identified in our dataset. The large number of unidentified genes could be due to natural divergences in the genome and different mRNA splicing, pointing to the low-depth annotation of the genome.

Based upon PANTHER enrichment analysis, GC have an over-representation of multiple biological processes related to the cell cycle, DNA replication, RNA splicing, and translation (Table 1). This indicates the cells are actively going through replication and reorganization, which would be expected as follicle development and oocyte maturation are very dynamic processes requiring substantial transcriptional activity. Moreover, the data highlighted the Wnt signaling pathway as one of the most enriched pathways related to reproduction. The Wnt signaling pathway must be activated for GC to support oocyte maturation (43). In addition, studies *in vitro* have found that supplementing maturation medium with a Wnt activator results in GC layer thickening and increased oocyte maturation in mice (43). Therefore, these findings could help identify potential biomarkers for successful oocyte maturation that could be applied to SWR ARTs to increase *in vitro* oocyte maturation.

Apoptosis and p53 signaling pathways were also highly enriched in this dataset. p53 is a gene associated with apoptotic events in the follicle during peri-ovulatory periods (44) and apoptosis occurs throughout folliculogenesis (45). In the present study, apoptotic/p53 pathway over-representation may reflect the diversity of follicle sizes from which GC were collected, as it is unknown if the expression of the genes associated with these pathways is related to ovulation or follicular decline (44). Future studies will differentiate the GC based upon follicle type and size in order to define the specific role of these pathways.

As expected, due to the high transcriptional activity of developing follicles and GC, general transcription activity was the most over-represented pathway. This transcriptional activity is required for follicles to progress through folliculogenesis, culminating in ovulation (46). A deeper analysis into the genes expressed in these pathways could identify potential biomarkers of granulosa cell-expressed genes involved in oocyte competence acquisition and embryo development potential. By exploring the biological pathways identified and the genes associated with those pathways, this study aims to improve the understanding of the reproductive function of granulosa cells in this species and to aid *in vitro* culture systems by identifying candidate markers specific to the SWR.

The next focus of this study compared the SWR GC transcriptomic dataset to those of cattle and humans. Although a preferable model would have been the more closely related domestic horse, adequate sequencing data has yet to be published evaluating the entire transcriptome of GC, and only microarray data are available (34). As microarrays are made with known sequences bound to a microarray chip, it does not reveal the detailed sequences used in the sample, which is not comparable to this study (34). Therefore, the SWR transcriptome was compared to publicly available transcriptome datasets for cattle and human GC. There was 45% homology across the SWR, cattle, and human. Moreover, as the SWR/NWR genome annotation improves, this homology is expected to increase. 48% of the GC transcriptome of the SWR is shared with the transcriptome of human GC. This is key information because it indicates that human GC could be used as a potential model for some aspects of assisted reproductive technology in the SWR.

The cattle genome was also compared to the SWR dataset because large cattle datasets are publicly available (30). Surprisingly, there were fewer genes in common (43%) between the cattle and SWR GC (8,708 genes) compared to the human and SWR GC (10,408 genes). It is possible that this is due to the fact the human genome is more completely annotated than either the cattle or the SWR genome. One of the limitations of the SWR dataset was that the GC collected were derived from a pool of follicles from various developmental stages. Therefore, to extrapolate information about the follicular origin of these cells, the SWR data were compared to a well-described cattle GC dataset (30), that includes transcriptomic signatures of follicles in the growing, atretic, or plateau follicular stages. Although rhinoceros follicular dynamics have not traditionally been described as growing, plateau, and atretic, these are classic categories for follicle development and can aid in understanding the physiological status of the follicle and developing oocyte. (31, 47, 48). When comparing the data in this study to each of the follicular signatures in cattle, the highest percentage (14%) of all the SWR transcripts identified fell exclusively into the growing follicular phase (a total of 1571 genes). The second most abundant set of transcripts found in the SWR GC were specific to the atretic follicular phase of the cattle dataset. The ability to correlate follicle stage-specific genes to individual follicle categories (growing, atretic, or plateau) will allow future studies to evaluate the effect of different treatments and conditions (i.e. hormonal stimulation, OPU collection time, maternal age, reproductive health, etc.) on the quality of the oocyte collected. If the follicles are immature or atretic, the capacity to move further with IVF becomes limited.

As an example of a gene expressed in both the SWR and cattle GC transcriptome, in growing and plateau follicles, was StAR (steroidogenic acute regulatory protein)(49). StAR is a critical gene responsible for the initiation of steroid production in granulosa cells (45). Based upon percentages of genes in common between the different follicle categories in cattle, it is suggested that most of the SWR follicles aspirated at OPU were at the growing or atretic stages. It was not surprising to see a large number of genes associated with atresia, as it is a physiological event that occurs naturally in all follicles that are not recruited to ovulate, and is the final destiny of most growing follicles (50).

In conclusion, the focus of this study was to describe, for the first time, granulosa cell transcriptomic profiles in the SWR. In this threatened species, numerous aspects of ovarian physiology and follicular activity, particularly at the sub-cellular level, remain unknown and are of considerable importance in developing assisted reproductive technologies both for this species and the nearly extinct NWR. Molecular pathways involved in oocyte competence acquisition, reduced ovarian reserve, infertility, and maternal aging are some of the missing information for the reproductive physiology of this species. Studying the transcriptome of the GC provides an efficient and indirect way to understand oocyte developmental competence, due to the close interaction of the oocyte to its surrounding GC, without harming the valuable oocytes. As oocyte collection in this species is technically challenging (16), the ability to study aspects of oocyte maturation and follicle development is limited and relies on the GC surrounding the oocyte.

Previous studies have shown that sequencing of human oocytes and their granulosa cells across different follicular stages revealed unique transcriptional machinery, transcription factor networks, and interactions between the oocyte and GC that were specific to developmental stages (51). While the current study did not evaluate GS across follicular stages, it did provide baseline transcriptional information that can then be applied to future studies that will evaluate the different follicular stages. These data generated pathways that were over-represented, helping to identify candidate genes expressed in GC that could be expected as biomarkers of oocyte and embryo quality. Furthermore, future studies will be focused on gene expression dynamics throughout folliculogenesis by exploring the transcriptomes of rhinoceros GC at key stages of follicular development. Aspects of reproductive health may also be addressed by studying GC to further understand infertility and reproductive challenges in this species.

## Materials and Methods

### Animal management and ovum pick up (OPU)

All procedures, experiments, and methods were reviewed and approved by San Diego Zoo Wildlife Alliance’s Institutional Animal Care and Use Committee (IACUC; protocol number 18-018, United States Department of Agriculture Certificate #93-R-0151). In addition, guidelines set forth by ARRIVE (Animal Research: Reporting of *In Vivo* Experiments, https://arriveguidelines.org/arrive-guidelines) were followed. All methods for this study were performed in accordance with all relevant guidelines and regulations according to IACUC and ARRIVE.

This study used two female rhinoceroses from the San Diego Zoo Wildlife Alliance’s Safari Park in Escondido, CA. One female received a single injection of 1.8 mg of gonadotropin stimulation (15) 24 hr prior to OPU; the other female was naturally cyclic, providing GC from follicles of all maturation statuses at an unknown point in their estrous cycle. Both individuals were anesthetized with a combination of etorphine (1.5 - 1.9 mcg/kg), butorphanol (14 - 39 mcg/kg), and medetomidine (20 - 31 mcg/kg); one rhinoceros` also received azaperone (23 - 27 mcg/kg) administered intramuscularly. Both rhinoceroses required propofol administered i.v. during initial positioning for intubation (300 - 658 mcg/kg) with additional supplementation (821 mcg/kg) necessary one hour after initial anesthesia for the rhinoceros not given azaperone during induction. Anesthesia was reversed with atipamezole delivered i.m. at a 5:1 ratio to medetomidine (100 - 136 mcg/kg) and naltrexone delivered i.m. at a target 50:1 ratio to etorphine (80 - 110 mcg/kg) or atipamezole i.m. at a 7.5:1 ratio to medetomidine (231 mcg/kg) and naltrexone i.v. at 50:1 ratio to etorphine (77 mcg/kg) (15).

During OPU (15) each visible follicle was aspirated and rinsed with a 37°C flushing solution (Vigro) containing 12.5 I.U./ml of heparin. The follicle aspirate was maintained at 37°C and immediately transported to the laboratory, where it was filtered through a 0.22μm embryo filter (Professional Embryo Transfer Supply Inc.). The filter was rinsed with a complete flush medium (ABT complete flush; ABT360) to isolate cumulus oocyte complexes (COCs).

### Granulosa cell collection and RNA isolation/quantification

Free-floating mural granulosa cells were collected from both rhinoceroses during OPU. Non-atretic granulosa cells were selected before IVM (mural), pooled together, and stored in multiple aliquots at -80°C until RNA isolation. Total RNA was isolated from granulosa cells using the Arcturus PicoPure RNA Isolation Kit (Thermo Fisher Scientific, Waltham, MA) per the manufacturer’s instructions. There were three aliquots (technical replicates) from each rhinoceros resulting in a total of six samples. Granulosa cells were incubated with extraction buffer for 30 min before centrifugation to remove debris and extracellular material. The cell extract was incubated with ethanol, bound to, and washed on pre-conditioned purification columns. Total RNA was recovered into elution buffer and quantified using a Nanodrop (Nanodrop One Spectrophotometer; Fischer Scientific).

### RNA Sequencing

Following the manufacturer’s protocol, RNA-sequencing (cDNA) libraries were prepared using the NEBNext® Ultra™ II RNA Library Prep Kit for Illumina. Three technical replicate libraries were prepared for each animal using ten ng of total RNA (6 libraries total, 30 ng total RNA per animal). Briefly, RNA was fragmented, first- and second-strand cDNA synthesis was performed, and adapters were ligated. Each sample was amplified with a specific barcoded PCR primer for sample identification purposes. Prepared libraries were sent to the Sanford Burnham Prebys for sequencing. Single-end reads of 75 basepairs were generated for each sample on an Illumina NextSeq 500 (Illumina, San Diego, CA). Raw data files were uploaded to Gene Expression Omnibus with accession number GSE216035.

### Bioinformatic Analysis

Bioinformatic analysis was performed on the Galaxy web platform and used the public server at usegalaxy.org (52). The sequence quality of each sample was assessed by FastQC, and the results were aggregated for comparison using MultiQC (53, 54). Trimmomatic was utilized to remove the adapter sequences and low-quality bases (55). Bases were removed if their quality score was below a threshold of 25, and reads below 20 basepairs were removed from the analysis. Reads were aligned to the NWR genome CerSimCot1.0 (GCA_021442165.1) (41) using HISAT2 (56). The SWR genome (CerSimSim1.0; GCA_000283155.1) was not used for this analysis due to gaps in the sequencing and limited annotation. Stringtie was used for transcript assembly and quantification of both annotated and unannotated transcripts (57). Each sample was analyzed individually to assemble transcripts. Those resulting transcript annotations were combined with the CerSimCot1.0 gene annotation to create a final transcript annotation file representing the intersection and union of all inputs (57). Manual curation of gene names was required to ensure HGNC/VGNC naming conventions were maintained.

### Species Comparisons

Publicly available transcriptome datasets from cattle and human granulosa cells were compared to the SWR. Human transcripts were obtained from the Expression Atlas from the European Bioinformatics Institute (40). Bovine transcripts were obtained from Girard et al. (2015). The results from these transcriptomic studies were compared to the results obtained in the present study (29, 30). The comparisons between SWR/human genome and SWR/cattle genome were evaluated individually and across all three species. PANTHER was used to determine the biological pathways associated with the transcripts in common between species (58).

## Supporting information

All Supplementary Material

## Acknowledgements

We thank the Conservation Genetics group and Dr. Marisa Korody at the San Diego Zoo Wildlife Alliance for providing us with the newly sequenced NWR genome.

## Author Contributions

E.R. and B.D. collected samples. E.R. processed and prepared samples for RNA sequencing.

E.R. and K.K. performed bioinformatic analyses and wrote the initial draft of the manuscript.

M.A.S. aided in comparative data analysis and interpretation. B.D. aided in writing and revisions of the manuscript. S.C. supervised bioinformatic analysis and revision of the manuscript.

## Competing Interests

The authors have no competing interests to declare.

## Data Availability Statement

The datasets generated and analyzed during the current study are available in the Gene Expression Omnibus (GEO), accession number GSE216035.

## Notes

### Competing Interest Statement

The authors have declared no competing interest.

