## Supplementary material for "Genomic insights into southern white rhinoceros (*Ceratotherium simum simum*) reproduction: revealing granulosa cell gene expression": All Supplementary Material

### Supplemental List Legends

Supplemental List 1. The complete list of genes identified in southern white rhinoceros granulosa cells.

Supplemental List 2. The complete list of genes in common between southern white rhinoceros and human granulosa cells.

Supplemental List 3. The complete list of genes in common between southern white rhinoceros and cattle granulosa cells.

Supplemental List 4. The complete list of genes in common between southern white rhinoceros, cattle, and human granulosa cells.

Supplemental List 5. The complete list of genes in common between the southern white rhinoceros and cattle granulosa cells in each of three follicular stages (growing, plateau, and atretic).

**Supplemental List 1**

| |
|---|
| A2GL |
| A2M |
| A3GALT2 |
| A4GALT |
| AAAS |
| AACS |
| AADAC |
| AAGAB |
| AAK1 |
| AAMP |
| AAR2 |
| AARS1 |
| AARS2 |
| AARSD1 |
| AASDH |
| AASDHPPT |
| AASS |
| AATF |
| AATK |
| ABCA1 |
| ABCA12 |
| ABCA17 |
| ABCA2 |
| ABCA3 |
| ABCA4 |
| ABCA5 |
| ABCA6 |
| ABCA7 |
| ABCA9 |
| ABCB1 |
| ABCB10 |
| ABCB19 |
| ABCB6 |
| ABCB7 |
| ABCB8 |
| ABCB9 |
| ABCC1 |
| ABCC10 |
| ABCC11 |
| ABCC3 |
| ABCC4 |
| ABCC9 |

| |
|---|
| ABCD1 |
| ABCD3 |
| ABCD4 |
| ABCE1 |
| ABCF1 |
| ABCF2 |
| ABCF3 |
| ABCG1 |
| ABCG2 |
| ABHD1 |
| ABHD10 |
| ABHD11 |
| ABHD12 |
| ABHD13 |
| ABHD15 |
| ABHD16A |
| ABHD17A |
| ABHD18 |
| ABHD2 |
| ABHD3 |
| ABHD4 |
| ABHD5 |
| ABHD6 |
| ABHD8 |
| ABI1 |
| ABI3BP |
| ABITRAM |
| ABL2 |
| ABLM1 |
| ABR |
| ABRACL |
| ABRAXAS1 |
| ABRAXAS2 |
| ABT1 |
| ABTB1 |
| ABTB2 |
| ACAA1 |
| ACAA2 |
| ACACA |
| ACACB |
| ACAD10 |
| ACAD11 |
| ACAD8 |

| |
|---|
| ACADS |
| ACADSB |
| ACADVL |
| ACAN |
| ACAP1 |
| ACAP2 |
| ACAP3 |
| ACAT1 |
| ACAT2 |
| ACBD3 |
| ACBD4 |
| ACBD5 |
| ACBD6 |
| ACCS |
| ACD |
| ACER3 |
| ACHE |
| ACIN1 |
| ACKR2 |
| ACKR3 |
| ACKR4 |
| ACLY |
| ACOD1 |
| ACOT11 |
| ACOT12 |
| ACOT13 |
| ACOT2 |
| ACOT4 |
| ACOT6 |
| ACOT7 |
| ACOT8 |
| ACOT9 |
| ACOX1 |
| ACOX3 |
| ACP1 |
| ACP2 |
| ACP6 |
| ACRBP |
| ACSBG1 |
| ACSF2 |
| ACSF3 |
| ACSL1 |
| ACSL3 |

| |
|---|
| ACSL6 |
| ACSM3 |
| ACSS1 |
| ACSS2 |
| ACSS3 |
| ACTA1 |
| ACTA2 |
| ACTB |
| ACTBL2 |
| ACTG1 |
| ACTL6A |
| ACTL7A |
| ACTL7B |
| ACTN1 |
| ACTN4 |
| ACTR10 |
| ACTR1B |
| ACTR2 |
| ACTR3 |
| ACTR5 |
| ACTR6 |
| ACTR8 |
| ACTZ |
| ACVR1 |
| ACVR1B |
| ACVR2A |
| ACVR2B |
| ACY1 |
| ACYP1 |
| ACYP2 |
| ADA |
| ADAL |
| ADAM10 |
| ADAM12 |
| ADAM15 |
| ADAM17 |
| ADAM19 |
| ADAM20 |
| ADAM21 |
| ADAM22 |
| ADAM32 |
| ADAM8 |
| ADAM9 |

| |
|---|
| ADAMTS1 |
| ADAMTS12 |
| ADAMTS13 |
| ADAMTS18 |
| ADAMTS2 |
| ADAMTS3 |
| ADAMTS5 |
| ADAMTS6 |
| ADAMTS7 |
| ADAMTS8 |
| ADAMTS9 |
| ADAMTSL4 |
| ADAP1 |
| ADAP2 |
| ADARB1 |
| ADAT1 |
| ADAT2 |
| ADAT3 |
| ADCK1 |
| ADCK2 |
| ADCK5 |
| ADCY1 |
| ADCY10 |
| ADCY3 |
| ADCY4 |
| ADCY6 |
| ADCY7 |
| ADCY9 |
| ADDA |
| ADDG |
| ADGB |
| ADGRA3 |
| ADGRB1 |
| ADGRB2 |
| ADGRD1 |
| ADGRE4P |
| ADGRE5 |
| ADGRF1 |
| ADGRF2 |
| ADGRG1 |
| ADGRG6 |
| ADGRL1 |
| ADGRL2 |

| |
|---|
| ADGRL3 |
| ADGRV1 |
| ADH1A |
| ADH4 |
| ADH5 |
| ADHFE1 |
| ADI1 |
| ADIPOR1 |
| ADIPOR2 |
| ADM |
| ADNP2 |
| ADO |
| ADORA1 |
| ADORA2A |
| ADORA2B |
| ADPGK |
| ADPRHL1 |
| ADPRM |
| ADPRS |
| ADRA1D |
| ADRA2A |
| ADRB2 |
| ADRM1 |
| ADSL |
| ADTRP |
| AEBP1 |
| AEBP2 |
| AEN |
| AFAP1 |
| AFF2 |
| AFF3 |
| AFF4 |
| AFG1L |
| AFG3L2 |
| AFMID |
| AFTPH |
| AGAP3 |
| AGBL2 |
| AGBL3 |
| AGBL5 |
| AGFG1 |
| AGFG2 |
| AGGF1 |

| |
|---|
| AGK |
| AGL |
| AGMAT |
| AGO1 |
| AGO2 |
| AGO3 |
| AGO4 |
| AGPAT1 |
| AGPAT2 |
| AGPAT3 |
| AGPAT5 |
| AGPS |
| AGRN |
| AGT |
| AGTPBP1 |
| AGTR2 |
| AHCTF1 |
| AHCY |
| AHCYL1 |
| AHCYL2 |
| AHDC1 |
| AHI1 |
| AHNAK |
| AHNAK2 |
| AHR |
| AHSA1 |
| AHSA2P |
| AIDA |
| AIF1L |
| AIFM1 |
| AIG1 |
| AIMP1 |
| AIMP2 |
| AIP |
| AIRE |
| AJM1 |
| AJUBA |
| AK1 |
| AK2 |
| AK3 |
| AK4 |
| AK6 |
| AK9 |

| |
|---|
| AKAP1 |
| AKAP11 |
| AKAP12 |
| AKAP13 |
| AKAP17A |
| AKAP5 |
| AKAP6 |
| AKAP7 |
| AKAP8 |
| AKAP8L |
| AKAP9 |
| AKIP1 |
| AKIRIN1 |
| AKIRIN2 |
| AKNA |
| AKNRD54 |
| AKR1A1 |
| AKR1B1 |
| AKR7L |
| AKT1 |
| AKT1S1 |
| AKT2 |
| AKT3 |
| AKTIP |
| ALAS1 |
| ALB1 |
| ALCAM |
| ALDH16A1 |
| ALDH18A1 |
| ALDH1A1 |
| ALDH1B1 |
| ALDH1L1 |
| ALDH1L2 |
| ALDH2 |
| ALDH3B2 |
| ALDH4A1 |
| ALDH5A1 |
| ALDH6A1 |
| ALDH7A1 |
| ALDH9A1 |
| ALDOA |
| ALG1 |
| ALG11 |

| |
|---|
| ALG12 |
| ALG13 |
| ALG14 |
| ALG2 |
| ALG3 |
| ALG5 |
| ALG6 |
| ALG8 |
| ALKBH1 |
| ALKBH2 |
| ALKBH3 |
| ALKBH4 |
| ALKBH5 |
| ALKBH7 |
| ALMS1 |
| ALOX15 |
| ALOX5 |
| ALOX5AP |
| ALPI |
| ALPK1 |
| ALPK3 |
| ALPL |
| ALS |
| ALS2 |
| ALYREF |
| AMACR |
| AMBRA1 |
| AMD1 |
| AMDHD1 |
| AMER1 |
| AMER3 |
| AMFR |
| AMH |
| AMHR2 |
| AMIGO1 |
| AMIGO2 |
| AMIGO3 |
| AMMECR1 |
| AMN |
| AMN1 |
| AMOT |
| AMOTL1 |
| AMOTL2 |

| |
|---|
| AMPD2 |
| AMPD3 |
| AMY2A |
| AMZ2 |
| ANAPC1 |
| ANAPC10 |
| ANAPC11 |
| ANAPC13 |
| ANAPC15 |
| ANAPC16 |
| ANAPC2 |
| ANAPC7 |
| ANG |
| ANGEL1 |
| ANGEL2 |
| ANGPT2 |
| ANGPTL1 |
| ANGPTL2 |
| ANGPTL5 |
| ANGPTL7 |
| ANK1 |
| ANK2 |
| ANK3 |
| ANKD1B |
| ANKDD1B |
| ANKEF1 |
| ANKFY1 |
| ANKH |
| ANKHD1 |
| ANKIB1 |
| ANKLE1 |
| ANKLE2 |
| ANKMY2 |
| ANKRA2 |
| ANKRD10 |
| ANKRD11 |
| ANKRD12 |
| ANKRD13A |
| ANKRD13C |
| ANKRD13D |
| ANKRD16 |
| ANKRD17 |
| ANKRD2 |

| |
|---|
| ANKRD22 |
| ANKRD26 |
| ANKRD27 |
| ANKRD28 |
| ANKRD29 |
| ANKRD31 |
| ANKRD33B |
| ANKRD34A |
| ANKRD35 |
| ANKRD37 |
| ANKRD39 |
| ANKRD40 |
| ANKRD46 |
| ANKRD49 |
| ANKRD50 |
| ANKRD52 |
| ANKRD55 |
| ANKRD9 |
| ANKS3 |
| ANKS4B |
| ANKS6 |
| ANKZF1 |
| ANLN |
| ANO10 |
| ANO2 |
| ANO6 |
| ANO7 |
| ANO8 |
| ANO9 |
| ANOS1 |
| ANP32A |
| ANP32B |
| ANP32E |
| ANTKMT |
| ANTXR1 |
| ANTXR2 |
| ANXA1 |
| ANXA11 |
| ANXA13 |
| ANXA2 |
| ANXA2R |
| ANXA3 |
| ANXA5 |

| |
|---|
| ANXA6 |
| AOAH |
| AOC2 |
| AOC3 |
| AOPEP |
| AOX1 |
| AP1AR |
| AP1B1 |
| AP1G1 |
| AP1G2 |
| AP1M1 |
| AP1S1 |
| AP1S2 |
| AP2A1 |
| AP2A2 |
| AP2M1 |
| AP2S1 |
| AP3B1 |
| AP3D1 |
| AP3M1 |
| AP3S1 |
| AP3S2 |
| AP4B1 |
| AP4E1 |
| AP4M1 |
| AP4S1 |
| AP5B1 |
| AP5M1 |
| AP5S1 |
| AP5Z1 |
| APAF1 |
| APBB1 |
| APBB1IP |
| APBB2 |
| APBB3 |
| APC |
| APEH |
| APEX1 |
| APEX2 |
| APH1A |
| APH1B |
| API5 |
| APIP |

| |
|---|
| APLF |
| APLP2 |
| APMAP |
| APOA1 |
| APOB |
| APOBEC3CA |
| APOBR |
| APOD |
| APOE |
| APOL3 |
| APOLD1 |
| APOM |
| APOO |
| APOOL |
| APP |
| APPBP2 |
| APPL1 |
| APPL2 |
| APRT |
| APTX |
| AQP11 |
| AQP7 |
| AQP8 |
| AQR |
| AR |
| ARAF |
| ARAP1 |
| ARAP2 |
| ARAP3 |
| ARC |
| AREL1 |
| ARF1 |
| ARF2 |
| ARF4 |
| ARF5 |
| ARF6 |
| ARFGAP1 |
| ARFGAP2 |
| ARFGAP3 |
| ARFGEF1 |
| ARFGEF2 |
| ARFGEF3 |
| ARFIP1 |

| |
|---|
| ARFIP2 |
| ARFRP1 |
| ARG2 |
| ARGLU1 |
| ARHGAP1 |
| ARHGAP10 |
| ARHGAP11A |
| ARHGAP17 |
| ARHGAP18 |
| ARHGAP19 |
| ARHGAP20 |
| ARHGAP21 |
| ARHGAP22 |
| ARHGAP23 |
| ARHGAP25 |
| ARHGAP26 |
| ARHGAP29 |
| ARHGAP30 |
| ARHGAP31 |
| ARHGAP32 |
| ARHGAP33 |
| ARHGAP35 |
| ARHGAP39 |
| ARHGAP4 |
| ARHGAP40 |
| ARHGAP42 |
| ARHGAP44 |
| ARHGAP45 |
| ARHGAP5 |
| ARHGAP6 |
| ARHGAP9 |
| ARHGDIA |
| ARHGDIB |
| ARHGDIG |
| ARHGEF1 |
| ARHGEF10 |
| ARHGEF10L |
| ARHGEF11 |
| ARHGEF12 |
| ARHGEF16 |
| ARHGEF17 |
| ARHGEF18 |
| ARHGEF19 |

| |
|---|
| ARHGEF2 |
| ARHGEF25 |
| ARHGEF28 |
| ARHGEF3 |
| ARHGEF39 |
| ARHGEF4 |
| ARHGEF40 |
| ARHGEF6 |
| ARHGEF7 |
| ARHGEF9 |
| ARI1 |
| ARID1A |
| ARID1B |
| ARID2 |
| ARID3A |
| ARID3B |
| ARID4A |
| ARID5A |
| ARID5B |
| ARIH2 |
| ARL1 |
| ARL10 |
| ARL11 |
| ARL13B |
| ARL14 |
| ARL14EP |
| ARL15 |
| ARL2 |
| ARL2BP |
| ARL3 |
| ARL4A |
| ARL4C |
| ARL4D |
| ARL5A |
| ARL5B |
| ARL6 |
| ARL6IP1 |
| ARL6IP5 |
| ARL6IP6 |
| ARL8A |
| ARL8B |
| ARMC1 |
| ARMC3 |

| |
|---|
| ARMC5 |
| ARMC6 |
| ARMC7 |
| ARMC8 |
| ARMC9 |
| ARMCX1 |
| ARMCX2 |
| ARMCX3 |
| ARMCX6 |
| ARNT2 |
| ARNTL |
| ARPC1B |
| ARPC3 |
| ARPC4 |
| ARPC5 |
| ARPC5L |
| ARPIN |
| ARPP21 |
| ARRB1 |
| ARRDC1 |
| ARRDC2 |
| ARRDC3 |
| ARRDC4 |
| ARSA |
| ARSB |
| ARSG |
| ARSK |
| ART3 |
| ART4 |
| AS3MT |
| ASAH1 |
| ASAH2 |
| ASAP1 |
| ASAP2 |
| ASAP3 |
| ASB1 |
| ASB13 |
| ASB15 |
| ASB16 |
| ASB3 |
| ASB4 |
| ASB5 |
| ASB6 |

| |
|---|
| ASB7 |
| ASB8 |
| ASCC1 |
| ASCC2 |
| ASCC3 |
| ASCL1 |
| ASCL4 |
| ASF1A |
| ASF1B |
| ASH1L |
| ASL |
| ASMTL |
| ASNS |
| ASNSD1 |
| ASPG |
| ASPH |
| ASPHD2 |
| ASPM |
| ASPRV1 |
| ASPSCR1 |
| ASRGL1 |
| ASTE1 |
| ASXL1 |
| ASXL2 |
| ASXL3 |
| ATAD1 |
| ATAD2 |
| ATAD5 |
| ATE1 |
| ATF2 |
| ATF3 |
| ATF4 |
| ATF5 |
| ATF6 |
| ATF7 |
| ATF7IP |
| ATF7IP2 |
| ATG10 |
| ATG101 |
| ATG12 |
| ATG13 |
| ATG14 |
| ATG16L1 |

| |
|---|
| ATG16L2 |
| ATG2A |
| ATG2B |
| ATG3 |
| ATG4A |
| ATG4B |
| ATG4C |
| ATG5 |
| ATG7 |
| ATG9A |
| ATIC |
| ATL2 |
| ATL3 |
| ATM |
| ATMIN |
| ATN1 |
| ATOH1 |
| ATOH7 |
| ATOX1 |
| ATP10D |
| ATP11A |
| ATP11B |
| ATP11C |
| ATP12A |
| ATP13A1 |
| ATP13A2 |
| ATP1A1 |
| ATP1A3 |
| ATP1B1 |
| ATP1B2 |
| ATP1B3 |
| ATP23 |
| ATP2A1 |
| ATP2A2 |
| ATP2A3 |
| ATP2B2 |
| ATP2B4 |
| ATP2C1 |
| ATP4A |
| ATP4B |
| ATP5F1A |
| ATP5F1B |
| ATP5F1C |

| |
|---|
| ATP5F1D |
| ATP5F1E |
| ATP5IF1 |
| ATP5MC1 |
| ATP5MC3 |
| ATP5ME |
| ATP5MF |
| ATP5MG |
| ATP5MJ |
| ATP5PB |
| ATP5PD |
| ATP5PO |
| ATP6AP1 |
| ATP6AP2 |
| ATP6V0A1 |
| ATP6V0A2 |
| ATP6V0A4 |
| ATP6V0B |
| ATP6V0C |
| ATP6V0D1 |
| ATP6V0D2 |
| ATP6V0E1 |
| ATP6V1A |
| ATP6V1B2 |
| ATP6V1D |
| ATP6V1E1 |
| ATP6V1F |
| ATP6V1G1 |
| ATP6V1G2 |
| ATP6V1H |
| ATP7A |
| ATP7B |
| ATP8A1 |
| ATP8A2 |
| ATP8B1 |
| ATP8B2 |
| ATP8B3 |
| ATP8B4 |
| ATP9A |
| ATP9B |
| ATPAF1 |
| ATPAF2 |
| ATPSCKMT |

| |
|---|
| ATR |
| ATRAID |
| ATRIP |
| ATRN |
| ATRNL1 |
| ATRX |
| ATX10 |
| ATXN1 |
| ATXN1L |
| ATXN2 |
| ATXN2L |
| ATXN3 |
| ATXN7 |
| ATXN7L1 |
| ATXN7L2 |
| ATXN7L3 |
| ATXN7L3B |
| AUH |
| AUNIP |
| AUP1 |
| AURKA |
| AURKAIP1 |
| AURKC |
| AUTS2 |
| AVEN |
| AVL9 |
| AVPI1 |
| AXIN1 |
| AXIN2 |
| AXL |
| AZI2 |
| AZIN1 |
| B2M |
| B3GALNT1 |
| B3GALNT2 |
| B3GALT2 |
| B3GALT4 |
| B3GALT5 |
| B3GALT6 |
| B3GAT2 |
| B3GAT3 |
| B3GLCT |
| B3GNT2 |

| |
|---|
| B3GNT3 |
| B3GNT4 |
| B3GNT5 |
| B3GNT7 |
| B3GNT8 |
| B3GNT9 |
| B3GNTL1 |
| B4GALNT1 |
| B4GALNT2 |
| B4GALNT3 |
| B4GALT1 |
| B4GALT2 |
| B4GALT3 |
| B4GALT4 |
| B4GALT5 |
| B4GALT7 |
| B4GAT1 |
| B9D1 |
| B9D2 |
| BAALC |
| BAAT |
| BABAM1 |
| BABAM2 |
| BACE1 |
| BACE2 |
| BACH2 |
| BAD |
| BAG1 |
| BAG2 |
| BAG3 |
| BAG4 |
| BAG5 |
| BAG6 |
| BAHCC1 |
| BAHD1 |
| BAIAP2 |
| BAIAP2L1 |
| BAIAP2L2 |
| BAIAP3 |
| BAIP2 |
| BAK1 |
| BAMBI |
| BANF1 |

| |
|---|
| BANK1 |
| BANP |
| BAP1 |
| BARD1 |
| BASP1 |
| BATF2 |
| BAX |
| BAZ1A |
| BAZ1B |
| BAZ2A |
| BAZ2B |
| BBLN |
| BBOF1 |
| BBS1 |
| BBS12 |
| BBS2 |
| BBS4 |
| BBS5 |
| BBS7 |
| BBS9 |
| BBX |
| BCAM |
| BCAP29 |
| BCAP31 |
| BCAR1 |
| BCAR3 |
| BCAS1 |
| BCAS2 |
| BCAS3 |
| BCAS4 |
| BCAT2 |
| BCCIP |
| BCDIN3D |
| BCHE |
| BCKDHA |
| BCKDHB |
| BCKDK |
| BCL10 |
| BCL11A |
| BCL2 |
| BCL2L1 |
| BCL2L11 |
| BCL2L12 |

| |
|---|
| BCL2L15 |
| BCL3 |
| BCL6 |
| BCL6B |
| BCL7B |
| BCL7C |
| BCL9 |
| BCLAF1 |
| BCLAF3 |
| BCO1 |
| BCOR |
| BCORL1 |
| BCR |
| BCS1L |
| BDH2 |
| BDP1 |
| BECN1 |
| BEND3 |
| BEND5 |
| BEND6 |
| BEND7 |
| BEST1 |
| BEST2 |
| BEST4 |
| BET1L |
| BEX2 |
| BEX3 |
| BEX4 |
| BFAR |
| BFSP1 |
| BGN |
| BHLHA15 |
| BHLHB9 |
| BHLHE40 |
| BHLHE41 |
| BICD1 |
| BICD2 |
| BICDL1 |
| BICDL2 |
| BICRA |
| BICRAL |
| BID |
| BIK |

| |
|---|
| BIN1 |
| BIN2 |
| BIN3 |
| BIRC2 |
| BIRC3 |
| BIRC5 |
| BIRC6 |
| BIVM |
| BLCAP |
| BLM |
| BLMH |
| BLNK |
| BLOC1S1 |
| BLOC1S2 |
| BLOC1S3 |
| BLOC1S4 |
| BLOC1S5 |
| BLOC1S6 |
| BLVRA |
| BLVRB |
| BLZF1 |
| BMERB1 |
| BMF |
| BMP1 |
| BMP2 |
| BMP2K |
| BMP3 |
| BMP6 |
| BMPR1A |
| BMPR2 |
| BMS1 |
| BMT2 |
| BNC2 |
| BNIP1 |
| BNIP3 |
| BNIP3L |
| BOD1 |
| BOD1L1 |
| BOK |
| BOLA1 |
| BOLA2 |
| BOLA3 |
| BOP |

| |
|---|
| BOP1 |
| BORA |
| BORCS5 |
| BORCS6 |
| BORCS8 |
| BPGM |
| BPHL |
| BPNT1 |
| BPNT2 |
| BPTF |
| BRAF |
| BRAP |
| BRAT1 |
| BRCA1 |
| BRCC3 |
| BRD1 |
| BRD2 |
| BRD3 |
| BRD3OS |
| BRD4 |
| BRD7 |
| BRD8 |
| BRD9 |
| BRDT |
| BRF1 |
| BRF2 |
| BRI3 |
| BRI3BP |
| BRINP3 |
| BRIP1 |
| BRIX1 |
| BRK1 |
| BRME1 |
| BRMS1 |
| BRMS1L |
| BROX |
| BRPF1 |
| BRPF3 |
| BRSK1 |
| BRWD1 |
| BRWD3 |
| BSCL2 |
| BSDC1 |

| |
|---|
| BSG |
| BSN |
| BSPRY |
| BST1 |
| BST2 |
| BTAF1 |
| BTBD1 |
| BTBD10 |
| BTBD2 |
| BTBD3 |
| BTBD6 |
| BTBD7 |
| BTBD9 |
| BTC |
| BTD |
| BTF3 |
| BTF3L4 |
| BTG1 |
| BTG2 |
| BTG3 |
| BTK |
| BTN1A1 |
| BTN3A3 |
| BTNL9 |
| BTRC |
| BUB1 |
| BUB1B |
| BUB3 |
| BUD13 |
| BUD23 |
| BUD31 |
| BYSL |
| BZW1 |
| BZW2 |
| C10orf143 |
| C10orf71 |
| C11orf1 |
| C11orf24 |
| C11orf42 |
| C11orf54 |
| C11orf71 |
| C11orf98 |
| C12orf4 |

| |
|---|
| C12orf43 |
| C12orf56 |
| C12orf57 |
| C12orf60 |
| C12orf73 |
| C14ORF119 |
| C14orf93 |
| C15orf39 |
| C15orf40 |
| C15orf61 |
| C15orf62 |
| C16orf46 |
| C16orf72 |
| C16orf90 |
| C16orf91 |
| C17orf113 |
| C17orf49 |
| C17orf64 |
| C17orf75 |
| C17orf80 |
| C18orf21 |
| C18orf25 |
| C18orf32 |
| C18orf54 |
| C18orf63 |
| C19orf12 |
| C19orf44 |
| C19ORF53 |
| C19orf54 |
| C1D |
| C1GALT1 |
| C1GALT1C1 |
| C1orf109 |
| C1orf112 |
| C1orf115 |
| C1orf131 |
| C1orf159 |
| C1orf162 |
| C1orf174 |
| C1orf185 |
| C1orf198 |
| C1orf21 |
| C1orf210 |

| |
|---|
| C1orf216 |
| C1orf35 |
| C1orf43 |
| C1orf50 |
| C1orf52 |
| C1orf53 |
| C1orf74 |
| C1QA |
| C1QB |
| C1QBP |
| C1QC |
| C1QTNF1 |
| C1QTNF12 |
| C1QTNF2 |
| C1QTNF4 |
| C1QTNF6 |
| C1QTNF8 |
| C1R |
| C1RL |
| C1S |
| C2 |
| C20orf204 |
| C20orf27 |
| C20orf96 |
| C22orf31 |
| C22orf39 |
| C2CD2 |
| C2CD2L |
| C2CD3 |
| C2CD4A |
| C2CD4B |
| C2CD5 |
| C2orf42 |
| C2orf49 |
| C2orf69 |
| C2orf74 |
| C2orf76 |
| C2orf81 |
| C2ORF88 |
| C3AR1 |
| C3orf14 |
| C3orf18 |
| C3orf20 |

| |
|---|
| C3orf38 |
| C3orf62 |
| C3orf80 |
| C4A |
| C4orf19 |
| C4orf33 |
| C4orf46 |
| C4orf48 |
| C5AR1 |
| C5AR2 |
| C5ORF15 |
| C5orf22 |
| C5orf24 |
| C5orf34 |
| C5orf63 |
| C6orf120 |
| C6orf132 |
| C6orf136 |
| C6orf141 |
| C6orf163 |
| C6orf226 |
| C6ORF23 |
| C6orf47 |
| C6ORF58 |
| C6orf62 |
| C6orf89 |
| C7orf25 |
| C7orf31 |
| C7orf61 |
| C8G |
| C8orf33 |
| C8orf48 |
| C8orf76 |
| C8orf82 |
| C9orf131 |
| C9orf152 |
| C9orf40 |
| C9orf43 |
| C9orf64 |
| C9orf72 |
| C9orf78 |
| C9orf85 |
| CA12 |

| |
|---|
| CA2 |
| CA4 |
| CA5B |
| CA7 |
| CA9 |
| CAAP1 |
| CAB39L |
| CABIN1 |
| CABLES2 |
| CABP5 |
| CABYR |
| CACFD1 |
| CACHD1 |
| CACNA1B |
| CACNA1F |
| CACNA1G |
| CACNA1H |
| CACNA1S |
| CACNA2D1 |
| CACNB1 |
| CACNB2 |
| CACNB3 |
| CACTIN |
| CACUL2 |
| CACYBP |
| CAD |
| CADM1 |
| CADM4 |
| CALCOCO1 |
| CALCOCO2 |
| CALD1 |
| CALHM2 |
| CALHM5 |
| CALHM6 |
| CALM1 |
| CALM2 |
| CALR |
| CALU |
| CAMK2D |
| CAMK2G |
| CAMK2N1 |
| CAMKK1 |
| CAMKMT |

| |
|---|
| CAMKV |
| CAMLG |
| CAMSAP1 |
| CAMSAP2 |
| CAMSAP3 |
| CAMTA1 |
| CAND1 |
| CANT1 |
| CANX |
| CAP1 |
| CAPG |
| CAPN1 |
| CAPN10 |
| CAPN15 |
| CAPN2 |
| CAPN6 |
| CAPN7 |
| CAPN8 |
| CAPN9 |
| CAPNS1 |
| CAPNS2 |
| CAPRIN2 |
| CAPS2 |
| CAPZA1 |
| CAPZA2 |
| CAPZB |
| CARD19 |
| CARD6 |
| CARD8 |
| CARF |
| CARHSP1 |
| CARM1 |
| CARMIL1 |
| CARNMT1 |
| CARNS1 |
| CARS1 |
| CARS2 |
| CASC3 |
| CASD1 |
| CASKIN1 |
| CASKIN2 |
| CASP1 |
| CASP10 |

| |
|---|
| CASP14 |
| CASP2 |
| CASP3 |
| CASP7 |
| CASP8 |
| CASP8AP2 |
| CASP9 |
| CASQ1 |
| CASQ2 |
| CASTOR1 |
| CASTOR2 |
| CASZ1 |
| CATSPER2 |
| CATSPERE |
| CATSPERG |
| CAV2 |
| CAVIN1 |
| CAVIN2 |
| CAVIN3 |
| CBARP |
| CBFA2T2 |
| CBFB |
| CBL |
| CBLB |
| CBLC |
| CBLL1 |
| CBR1 |
| CBR3 |
| CBR4 |
| CBS |
| CBWD2 |
| CBX1 |
| CBX2 |
| CBX3 |
| CBX4 |
| CBX5 |
| CBX6 |
| CBX7 |
| CBX8 |
| CBY1 |
| CBY2 |
| CC2D1A |
| CC2D1B |

| |
|---|
| CC2D2A |
| CC2D2B |
| CCAR1 |
| CCAR2 |
| CCBE1 |
| CCDC102A |
| CCDC103 |
| CCDC106 |
| CCDC107 |
| CCDC112 |
| CCDC115 |
| CCDC117 |
| CCDC12 |
| CCDC121 |
| CCDC122 |
| CCDC124 |
| CCDC125 |
| CCDC126 |
| CCDC127 |
| CCDC137 |
| CCDC138 |
| CCDC14 |
| CCDC142 |
| CCDC149 |
| CCDC15 |
| CCDC157 |
| CCDC158 |
| CCDC159 |
| CCDC160 |
| CCDC163 |
| CCDC166 |
| CCDC167 |
| CCDC168 |
| CCDC170 |
| CCDC171 |
| CCDC174 |
| CCDC175 |
| CCDC177 |
| CCDC18 |
| CCDC180 |
| CCDC181 |
| CCDC183 |
| CCDC184 |

| |
|---|
| CCDC185 |
| CCDC186 |
| CCDC190 |
| CCDC191 |
| CCDC194 |
| CCDC22 |
| CCDC24 |
| CCDC25 |
| CCDC28A |
| CCDC28B |
| CCDC3 |
| CCDC30 |
| CCDC32 |
| CCDC34 |
| CCDC43 |
| CCDC47 |
| CCDC50 |
| CCDC51 |
| CCDC57 |
| CCDC59 |
| CCDC6 |
| CCDC61 |
| CCDC62 |
| CCDC65 |
| CCDC66 |
| CCDC69 |
| CCDC7 |
| CCDC71 |
| CCDC71L |
| CCDC73 |
| CCDC74B |
| CCDC77 |
| CCDC8 |
| CCDC80 |
| CCDC82 |
| CCDC85A |
| CCDC85B |
| CCDC85C |
| CCDC86 |
| CCDC87 |
| CCDC88A |
| CCDC88B |
| CCDC88C |

| |
|---|
| CCDC9 |
| CCDC90B |
| CCDC91 |
| CCDC93 |
| CCDC96 |
| CCDC97 |
| CCHCR1 |
| CCIN |
| CCKBR |
| CCL14 |
| CCL15 |
| CCL2 |
| CCL20 |
| CCL27 |
| CCL28 |
| CCL4 |
| CCL5 |
| CCL8 |
| CCM2 |
| CCN1 |
| CCN2 |
| CCNA2 |
| CCNB2 |
| CCNB3 |
| CCNC |
| CCND2 |
| CCND3 |
| CCNE1 |
| CCNE2 |
| CCNG1 |
| CCNG2 |
| CCNH |
| CCNI |
| CCNJ |
| CCNJL |
| CCNK |
| CCNL1 |
| CCNL2 |
| CCNQ |
| CCNT1 |
| CCNT2 |
| CCNY |
| CCNYL1 |

| |
|---|
| CCP110 |
| CCPG1 |
| CCR1 |
| CCR2 |
| CCR3 |
| CCR5 |
| CCR7 |
| CCRL2 |
| CCS |
| CCSAP |
| CCSER1 |
| CCSER2 |
| CCT2 |
| CCT3 |
| CCT4 |
| CCT5 |
| CCT6A |
| CCT7 |
| CCT8 |
| CCT8L2 |
| CCZ1 |
| CD101 |
| CD109 |
| CD14 |
| CD151 |
| CD163L1 |
| CD164 |
| CD180 |
| CD19 |
| CD1D |
| CD2 |
| CD200 |
| CD244 |
| CD247 |
| CD2AP |
| CD2BP2 |
| CD300A |
| CD300LB |
| CD300LF |
| CD302 |
| CD36 |
| CD37 |
| CD3E |

| |
|---|
| CD40 |
| CD44 |
| CD46 |
| CD47 |
| CD48 |
| CD5 |
| CD53 |
| CD55 |
| CD58 |
| CD6 |
| CD63 |
| CD68 |
| CD69 |
| CD7 |
| CD70 |
| CD72 |
| CD74 |
| CD79B |
| CD81 |
| CD82 |
| CD84 |
| CD86 |
| CD8A |
| CD9 |
| CD93 |
| CD96 |
| CD99L2 |
| CDADC1 |
| CDAN1 |
| CDC14A |
| CDC14B |
| CDC16 |
| CDC25B |
| CDC25C |
| CDC26 |
| CDC27 |
| CDC34 |
| CDC37 |
| CDC37L1 |
| CDC40 |
| CDC42 |
| CDC42BPA |
| CDC42BPB |

| |
|---|
| CDC42BPG |
| CDC42EP2 |
| CDC42EP3 |
| CDC42EP4 |
| CDC42EP5 |
| CDC42SE2 |
| CDC45 |
| CDC5L |
| CDC6 |
| CDC7 |
| CDC73 |
| CDCA2 |
| CDCA3 |
| CDCA4 |
| CDCA5 |
| CDCA7 |
| CDCA7L |
| CDCA8 |
| CDCC189 |
| CDCP1 |
| CDH10 |
| CDH12 |
| CDH17 |
| CDH18 |
| CDH2 |
| CDH24 |
| CDH3 |
| CDH6 |
| CDH8 |
| CDHR2 |
| CDHR5 |
| CDIN1 |
| CDIP1 |
| CDIPT |
| CDK1 |
| CDK10 |
| CDK12 |
| CDK13 |
| CDK16 |
| CDK17 |
| CDK18 |
| CDK19 |
| CDK20 |

| |
|---|
| CDK2AP1 |
| CDK2AP2 |
| CDK3 |
| CDK4 |
| CDK5 |
| CDK5R1 |
| CDK5RAP1 |
| CDK5RAP2 |
| CDK6 |
| CDK7 |
| CDK8 |
| CDK9 |
| CDKAL1 |
| CDKL1 |
| CDKL3 |
| CDKL5 |
| CDKN1A |
| CDKN1B |
| CDKN1C |
| CDKN2A |
| CDKN2AIPNL |
| CDKN2B |
| CDKN2C |
| CDKN2D |
| CDKN3 |
| CDO1 |
| CDON |
| CDPF1 |
| CDR2 |
| CDR2L |
| CDS1 |
| CDS2 |
| CDV3 |
| CDX1 |
| CDX2 |
| CDYL |
| CDYL2 |
| CE12 |
| CEACAM1 |
| CEACAM16 |
| CEACAM21 |
| CEBPA |
| CEBPB |

| |
|---|
| CEBPD |
| CEBPG |
| CEBPZ |
| CEBPZOS |
| CECR2 |
| CELA3B |
| CELF1 |
| CELF2 |
| CELSR1 |
| CELSR2 |
| CELSR3 |
| CEMIP |
| CEMIP2 |
| CENATAC |
| CENPA |
| CENPB |
| CENPC |
| CENPE |
| CENPF |
| CENPH |
| CENPI |
| CENPJ |
| CENPK |
| CENPM |
| CENPN |
| CENPO |
| CENPP |
| CENPQ |
| CENPT |
| CENPU |
| CENPV |
| CENPW |
| CENPX |
| CEP104 |
| CEP112 |
| CEP126 |
| CEP128 |
| CEP129 |
| CEP131 |
| CEP135 |
| CEP152 |
| CEP162 |
| CEP164 |

| |
|---|
| CEP170B |
| CEP19 |
| CEP192 |
| CEP250 |
| CEP295 |
| CEP295NL |
| CEP350 |
| CEP41 |
| CEP43 |
| CEP44 |
| CEP55 |
| CEP57 |
| CEP57L1 |
| CEP63 |
| CEP68 |
| CEP70 |
| CEP72 |
| CEP76 |
| CEP78 |
| CEP83 |
| CEP85L |
| CEP89 |
| CEP95 |
| CEP97 |
| CERCAM |
| CERK |
| CERS1 |
| CERS2 |
| CERS4 |
| CERT1 |
| CES1 |
| CETN2 |
| CETN3 |
| CFAP100 |
| CFAP20 |
| CFAP20DC |
| CFAP251 |
| CFAP298 |
| CFAP299 |
| CFAP300 |
| CFAP36 |
| CFAP410 |
| CFAP45 |

| |
|---|
| CFAP46 |
| CFAP54 |
| CFAP65 |
| CFAP69 |
| CFAP70 |
| CFAP91 |
| CFB |
| CFD |
| CFDP1 |
| CFDP2 |
| CFH |
| CFL1 |
| CFL2 |
| CGAS |
| CGGBP1 |
| CGN |
| CGNL1 |
| CGREF1 |
| CGRRF1 |
| CHAC1 |
| CHAC2 |
| CHADL |
| CHAF1A |
| CHAF1B |
| CHAMP1? |
| CHCHD2 |
| CHCHD3 |
| CHCHD4 |
| CHCHD6 |
| CHCHD7 |
| CHD1L |
| CHD2 |
| CHD3 |
| CHD4 |
| CHD6 |
| CHD8 |
| CHD9 |
| CHEK1 |
| CHEK2 |
| CHERP |
| CHFR |
| CHGB |
| CHI3L1 |

| |
|---|
| CHIC1 |
| CHIC2 |
| CHKA |
| CHL1 |
| CHML |
| CHMP1A |
| CHMP1B |
| CHMP2A |
| CHMP2B |
| CHMP4A |
| CHMP4B |
| CHMP4C |
| CHMP5 |
| CHMP6 |
| CHMP7 |
| CHORDC1 |
| CHP1 |
| CHP2 |
| CHPF2 |
| CHPT1 |
| CHRAC1 |
| CHRD |
| CHRDL1 |
| CHRM3 |
| CHRM5 |
| CHRNA3 |
| CHST11 |
| CHST12 |
| CHST13 |
| CHST14 |
| CHST15 |
| CHST3 |
| CHST4 |
| CHST7 |
| CHSY1 |
| CHSY3 |
| CHTOP |
| CHUK |
| CIAO1 |
| CIAO2A |
| CIAO2B |
| CIAO3 |
| CIAPIN1 |

| |
|---|
| CIART |
| CIB1 |
| CIB2 |
| CIBAR1 |
| CIC |
| CIDEC |
| CIITA |
| CILK1 |
| CILP1 |
| CINP |
| CIP2A |
| CIPC |
| CIR1 |
| CIRBP |
| CISD1 |
| CISD2 |
| CISD3 |
| CISH |
| CIT |
| CITED2 |
| CITED4 |
| CIZ1 |
| CKAP2 |
| CKAP2L |
| CKAP4 |
| CKAP5 |
| CKB |
| CKLF |
| CKMT1A |
| CKMT2 |
| CKS1B |
| CLASP1 |
| CLASP2 |
| CLASRP |
| CLBA1 |
| CLCA2 |
| CLCN2 |
| CLCN3 |
| CLCN4 |
| CLCN5 |
| CLCN6 |
| CLCN7 |
| CLDN1 |

| |
|---|
| CLDN11 |
| CLDN12 |
| CLDN15 |
| CLDN20 |
| CLDN23 |
| CLDN3 |
| CLDN4 |
| CLDN7 |
| CLDN8 |
| CLEC11A |
| CLEC12A |
| CLEC14A |
| CLEC16A |
| CLEC1A |
| CLEC3A |
| CLEC3B |
| CLGN |
| CLHC1 |
| CLIC1 |
| CLIC4 |
| CLINT1 |
| CLIP1 |
| CLIP2 |
| CLIP3 |
| CLK1 |
| CLK2 |
| CLK3 |
| CLK4 |
| CLMN |
| CLMP |
| CLN3 |
| CLN5 |
| CLN6 |
| CLN8 |
| CLNS1A |
| CLOCK |
| CLPB |
| CLPP |
| CLPS1 |
| CLPTM1 |
| CLPTM1L |
| CLPX |
| CLRN3 |

| |
|---|
| CLSPN |
| CLSTN1 |
| CLSTN2 |
| CLSTN3 |
| CLTA |
| CLTB |
| CLTC |
| CLU |
| CLUAP1 |
| CLYBL |
| CMAHP |
| CMAS |
| CMBL |
| CMC1 |
| CMC2 |
| CMC4 |
| CMKLR2 |
| CMPK1 |
| CMPK2 |
| CMTA2 |
| CMTM3 |
| CMTM6 |
| CMTM7 |
| CMTR1 |
| CMTR2 |
| CMYA5 |
| CNBP |
| CNDP2 |
| CNEP1R1 |
| CNIH1 |
| CNIH4 |
| CNKSR3 |
| CNN1 |
| CNN2 |
| CNN3 |
| CNNM2 |
| CNNM3 |
| CNNM4 |
| CNOT1 |
| CNOT10 |
| CNOT11 |
| CNOT3 |
| CNOT4 |

| |
|---|
| CNOT6 |
| CNOT6L |
| CNOT7 |
| CNOT8 |
| CNP |
| CNPPD1 |
| CNPY2 |
| CNPY3 |
| CNPY4 |
| CNR1 |
| CNR2 |
| CNTLN |
| CNTN1 |
| CNTN3 |
| CNTN6 |
| CNTNAP1 |
| CNTNAP2 |
| CNTRL |
| CNTROB |
| COA1 |
| COA3 |
| COA4 |
| COA5 |
| COA7 |
| COA8 |
| COASY |
| COBL |
| COBLL1 |
| COCH |
| COG1 |
| COG2 |
| COG3 |
| COG4 |
| COG5 |
| COG6 |
| COG7 |
| COG8 |
| COIL |
| COL13A1 |
| COL15A1 |
| COL17A1 |
| COL1A1 |
| COL21A1 |

| |
|---|
| COL24A1 |
| COL3A1 |
| COL4A1 |
| COL4A2 |
| COL4A3 |
| COL4A4 |
| COL4A5 |
| COL4A6 |
| COL6A1 |
| COL6A2 |
| COL6A5 |
| COL7A1 |
| COL8A2 |
| COL9A2 |
| COLCA2 |
| COLEC11 |
| COLEC12 |
| COLGALT1 |
| COMMD1 |
| COMMD10 |
| COMMD2 |
| COMMD4 |
| COMMD5 |
| COMMD6 |
| COMMD7 |
| COMMD8 |
| COMMD9 |
| COMT |
| COMTD1 |
| COP1 |
| COPA |
| COPB1 |
| COPB2 |
| COPE |
| COPG1 |
| COPG2 |
| COPRS |
| COPS3 |
| COPS5 |
| COPS6 |
| COPS7A |
| COPS8 |
| COPZ1 |

| |
|---|
| COPZ2 |
| COQ10A |
| COQ10B |
| COQ2 |
| COQ3 |
| COQ5 |
| COQ6 |
| COQ7 |
| COQ8A |
| COQ8B |
| COQ9 |
| CORIN |
| CORO1A |
| CORO1B |
| CORO1C |
| CORO2A |
| CORO6 |
| CORO7 |
| COTL1 |
| COX10 |
| COX11 |
| COX15 |
| COX16 |
| COX18 |
| COX19 |
| COX20 |
| COX4I1 |
| COX4I2 |
| COX5A |
| COX5B |
| COX6A1 |
| COX6B1 |
| COX6C |
| COX7A1 |
| COX7A2 |
| COX7A2L |
| COX8A |
| CPD |
| CPEB1 |
| CPEB2 |
| CPEB4 |
| CPED1 |
| CPLANE1 |

| |
|---|
| CPLANE2 |
| CPN2 |
| CPNE2 |
| CPNE3 |
| CPOX |
| CPPED |
| CPPED1 |
| CPSF1 |
| CPSF2 |
| CPSF3 |
| CPSF4 |
| CPT1A |
| CPT1B |
| CPT1C |
| CPT2 |
| CPTP |
| CPVL |
| CPXM1 |
| CPZ |
| CR1 |
| CRACD |
| CRACDL |
| CRACR2A |
| CRADD |
| CRAMP1 |
| CRAT |
| CRBN |
| CRCP |
| CREB1 |
| CREB3 |
| CREB3L1 |
| CREB3L2 |
| CREB3L3 |
| CREBL2 |
| CREBRF |
| CREBZF |
| CREG1 |
| CRELD1 |
| CRELD2 |
| CREM |
| CREPBP |
| CRIM1 |
| CRIP1 |

| |
|---|
| CRIP3 |
| CRIPT |
| CRK |
| CRKL |
| CRLF3 |
| CRLS1 |
| CRMA |
| CRNKL1 |
| CROCC |
| CRP |
| CRPPA |
| CRTAP |
| CRTC3 |
| CRY2 |
| CRYAB |
| CRYBG1 |
| CRYBG2 |
| CRYBG3 |
| CRYGC |
| CRYM |
| CRYZ |
| CRYZL1 |
| CS |
| CSAD |
| CSDE1 |
| CSF1 |
| CSF1R |
| CSF2RB |
| CSF3R |
| CSGALNACT1 |
| CSGALNACT2 |
| CSK |
| CSKMT |
| CSMD3 |
| CSN2 |
| CSN4 |
| CSNK1A1 |
| CSNK1D |
| CSNK1E |
| CSNK1G1 |
| CSNK1G2 |
| CSNK1G3 |
| CSNK2B |

| |
|---|
| CSPG4 |
| CSPG5 |
| CSPP1 |
| CSRNP1 |
| CSRNP2 |
| CSRP1 |
| CSRP2 |
| CST11 |
| CST3 |
| CST6 |
| CST7 |
| CSTF1 |
| CSTF2 |
| CSTF2T |
| CSTF3 |
| CSTL1 |
| CSTPP1 |
| CTAGE9 |
| CTBP1 |
| CTBP2 |
| CTC1 |
| CTCF |
| CTDNEP1 |
| CTDP1 |
| CTDSP1 |
| CTDSP2 |
| CTF1 |
| CTF18 |
| CTF2 |
| CTF8 |
| CTGF |
| CTH |
| CTHRC1 |
| CTIF |
| CTNNA1 |
| CTNNA2 |
| CTNNAL1 |
| CTNNB1 |
| CTNNBL1 |
| CTNND1 |
| CTPS1 |
| CTR9 |
| CTRL |

| |
|---|
| CTSA |
| CTSB |
| CTSC |
| CTSD |
| CTSH |
| CTSK |
| CTSL |
| CTSO |
| CTSV |
| CTSZ |
| CTTN |
| CTTNBP2NL |
| CTU1 |
| CTU2 |
| CTXN1 |
| CUBN |
| CUEDC2 |
| CUL1 |
| CUL2 |
| CUL3 |
| CUL4B |
| CUL5 |
| CUL7 |
| CUL9 |
| CUTA |
| CUTC |
| CUX1 |
| CUX2 |
| CUZD1 |
| CWC15 |
| CWC22 |
| CWC25 |
| CWC27 |
| CWF19L1 |
| CWF19L2 |
| CWH43 |
| CX3CL1 |
| CX3CR1 |
| CXADR |
| CXCL10 |
| CXCL16 |
| CXCL2 |
| CXCL8 |

| |
|---|
| CXCR2 |
| CXCR4 |
| CXCR5 |
| CXorf38 |
| CXorf58 |
| CXXC4 |
| CYB5 |
| CYB561A3 |
| CYB561D2 |
| CYB5B |
| CYB5D2 |
| CYB5R1 |
| CYB5R2 |
| CYB5R3 |
| CYB5R4 |
| CYBB |
| CYBC1 |
| CYBRD1 |
| CYC1 |
| CYCS |
| CYFIP2 |
| CYHR1 |
| CYLD |
| CYP11A1 |
| CYP19A1 |
| CYP1B1 |
| CYP20A1 |
| CYP27A1 |
| CYP27B1 |
| CYP2D6 |
| CYP2J2 |
| CYP2S1 |
| CYP2U1 |
| CYP4B1 |
| CYP4F3 |
| CYP4V2 |
| CYP8B1 |
| CYREN |
| CYRIB |
| CYSLTR1 |
| CYSLTR2 |
| CYSTM1 |
| CYTB |

| |
|---|
| CYTH2 |
| CYTH3 |
| CYTH4 |
| CYTIP |
| CZIB |
| D2HGDH |
| DAAM2 |
| DAB2 |
| DAB2IP |
| DACH1 |
| DACH2 |
| DACT2 |
| DAD1 |
| DAG1 |
| DAGLA |
| DAGLB |
| DALRD3 |
| DAP |
| DAPK1 |
| DAPK3 |
| DARS1 |
| DAXX |
| DAZAP1 |
| DAZAP2 |
| DBF4 |
| DBF4B |
| DBI |
| DBN1 |
| DBNDD1 |
| DBNL |
| DBP |
| DBR1 |
| DBT |
| DCAF1 |
| DCAF10 |
| DCAF11 |
| DCAF12 |
| DCAF12L2 |
| DCAF13 |
| DCAF15 |
| DCAF16 |
| DCAF17 |
| DCAF4 |

| |
|---|
| DCAF5 |
| DCAF6 |
| DCAF7 |
| DCAF8 |
| DCAKD |
| DCBLD1 |
| DCBLD2 |
| DCDC2 |
| DCHS1 |
| DCK |
| DCLRE1A |
| DCLRE1C |
| DCP1A |
| DCP1B |
| DCP2 |
| DCPS |
| DCT |
| DCTD |
| DCTN1 |
| DCTN2 |
| DCTN3 |
| DCTN5 |
| DCTN6 |
| DCTPP1 |
| DCUN1D1 |
| DCUN1D2 |
| DCUN1D3 |
| DCUN1D5 |
| DCXR |
| DDA1 |
| DDAH2 |
| DDB1 |
| DDB2 |
| DDHD1 |
| DDHD2 |
| DDI1 |
| DDI2 |
| DDIAS |
| DDIT3 |
| DDIT4 |
| DDN |
| DDO |
| DDOST |

| |
|---|
| DDR1 |
| DDR2 |
| DDRGK1 |
| DDT |
| DDX1 |
| DDX10 |
| DDX11L8 |
| DDX17 |
| DDX18 |
| DDX19A |
| DDX19B |
| DDX20 |
| DDX21 |
| DDX23 |
| DDX24 |
| DDX27 |
| DDX28 |
| DDX31 |
| DDX39A |
| DDX39B |
| DDX3X |
| DDX41 |
| DDX42 |
| DDX43 |
| DDX46 |
| DDX47 |
| DDX49 |
| DDX5 |
| DDX50 |
| DDX52 |
| DDX54 |
| DDX55 |
| DDX56 |
| DDX58 |
| DDX59 |
| DDX6 |
| DDX60 |
| DDX60L |
| DEAF1 |
| DECR1 |
| DECR2 |
| DEDD |
| DEDD2 |

| |
|---|
| DEF8 |
| DEFA5 |
| DEGS1 |
| DEGS2 |
| DENND10 |
| DENND1A |
| DENND1B |
| DENND1C |
| DENND2A |
| DENND2B |
| DENND2C |
| DENND2D |
| DENND3 |
| DENND4A |
| DENND4B |
| DENND4C |
| DENND5A |
| DENND5B |
| DENND6A |
| DENND6B |
| DENR |
| DEOC |
| DEPDC1 |
| DEPDC1B |
| DEPDC4 |
| DEPDC5 |
| DEPDC7 |
| DEPP1 |
| DEPTOR |
| DERL1 |
| DERL2 |
| DERL3 |
| DESI1 |
| DESI2 |
| DET1 |
| DEUP1 |
| DEXI |
| DFFA |
| DFFB |
| DGAT1 |
| DGAT2 |
| DGCR2 |
| DGCR6 |

| |
|---|
| DGCR8 |
| DGKA |
| DGKD |
| DGKE |
| DGKQ |
| DGKZ |
| DGUOK |
| DHCR24 |
| DHCR7 |
| DHDDS |
| DHDH |
| DHFR |
| DHH |
| DHODH |
| DHPS |
| DHRS1 |
| DHRS11 |
| DHRS13 |
| DHRS3 |
| DHRS4 |
| DHRS7 |
| DHRS7B |
| DHRS9 |
| DHRSX |
| DHTKD1 |
| DHX15 |
| DHX16 |
| DHX29 |
| DHX30 |
| DHX32 |
| DHX33 |
| DHX34 |
| DHX35 |
| DHX36 |
| DHX37 |
| DHX38 |
| DHX40 |
| DHX57 |
| DHX9 |
| DIABLO |
| DIAPH2 |
| DIAPH3 |
| DICER1 |

| |
|---|
| DIDO1 |
| DIMT1 |
| DIP2A |
| DIP2B |
| DIP2C |
| DIPK1A |
| DIPK1B |
| DIPK2A |
| DIRAS1 |
| DIRAS2 |
| DIRAS3 |
| DIS3 |
| DIS3L |
| DIS3L2 |
| DIXDC1 |
| DKC1 |
| DKK3 |
| DKKL1 |
| DLAT |
| DLC1 |
| DLD |
| DLEC1 |
| DLEU7 |
| DLG1 |
| DLG3 |
| DLG4 |
| DLG5 |
| DLGAP1 |
| DLGAP5 |
| DLL1 |
| DLL3 |
| DLST |
| DLX5 |
| DMAC2L |
| DMAP1 |
| DMD |
| DMKN |
| DMPK |
| DMTF1 |
| DMWD |
| DMXL1 |
| DMXL2 |
| DNA2 |

| |
|---|
| DNAAF11 |
| DNAAF2 |
| DNAAF3 |
| DNAAF5 |
| DNAAF9 |
| DNAH1 |
| DNAH2 |
| DNAH3 |
| DNAH7 |
| DNAH8 |
| DNAH9 |
| DNAI2 |
| DNAJA1 |
| DNAJA2 |
| DNAJA3 |
| DNAJA4 |
| DNAJB1 |
| DNAJB12 |
| DNAJB14 |
| DNAJB2 |
| DNAJB3 |
| DNAJB4 |
| DNAJB6 |
| DNAJB7 |
| DNAJB9 |
| DNAJC1 |
| DNAJC10 |
| DNAJC11 |
| DNAJC12 |
| DNAJC13 |
| DNAJC14 |
| DNAJC15 |
| DNAJC16 |
| DNAJC17 |
| DNAJC18 |
| DNAJC19 |
| DNAJC2 |
| DNAJC21 |
| DNAJC24 |
| DNAJC25 |
| DNAJC27 |
| DNAJC28 |
| DNAJC3 |

| |
|---|
| DNAJC30 |
| DNAJC4 |
| DNAJC5 |
| DNAJC7 |
| DNAJC8 |
| DNAJC9 |
| DNAL1 |
| DNAL4 |
| DNASE1L1 |
| DNASE2 |
| DND1 |
| DNHD1 |
| DNLZ |
| DNM1 |
| DNM1L |
| DNM2 |
| DNM3 |
| DNMBP |
| DNMT1 |
| DNMT3A |
| DNMT3B |
| DNPEP |
| DNPH1 |
| DNTTIP1 |
| DNTTIP2 |
| DOC2B |
| DOCK1 |
| DOCK11 |
| DOCK3 |
| DOCK4 |
| DOCK5 |
| DOCK6 |
| DOCK7 |
| DOCK8 |
| DOCK9 |
| DOHH |
| DOK1 |
| DOK2 |
| DOK3 |
| DOK4 |
| DOK7 |
| DOLK |
| DOLPP1 |

| |
|---|
| DOP1A |
| DOP1B |
| DOT1L |
| DPCD |
| DPF2 |
| DPF3 |
| DPH2 |
| DPH3 |
| DPH5 |
| DPH6 |
| DPH7 |
| DPM1 |
| DPM2 |
| DPM3 |
| DPP3 |
| DPP7 |
| DPY19L1 |
| DPY19L3 |
| DPY19L4 |
| DPY30 |
| DPYD |
| DPYS |
| DPYSL2 |
| DPYSL3 |
| DPYSL5 |
| DQX1 |
| DR1 |
| DRAM2 |
| DRAP1 |
| DRC3 |
| DRG1 |
| DRG2 |
| DROSHA |
| DRP2 |
| DSC2 |
| DSC3 |
| DSCC1 |
| DSE |
| DSEL |
| DSG2 |
| DSP |
| DST |
| DSTN |

| |
|---|
| DSTYK |
| DTD1 |
| DTD2 |
| DTL |
| DTNB |
| DTNBP1 |
| DTWD1 |
| DTWD2 |
| DTX1 |
| DTX2 |
| DTX3 |
| DTX3L |
| DTYMK |
| DUOX1 |
| DUS1L |
| DUS2 |
| DUS3L |
| DUS4L |
| DUSP1 |
| DUSP10 |
| DUSP11 |
| DUSP12 |
| DUSP14 |
| DUSP15 |
| DUSP16 |
| DUSP18 |
| DUSP19 |
| DUSP23 |
| DUSP28 |
| DUSP3 |
| DUSP5 |
| DUSP6 |
| DUSP7 |
| DUSP8 |
| DUSP9 |
| DUT |
| DVL1 |
| DVL2 |
| DXO |
| DYM |
| DYNC1H1 |
| DYNC1I2 |
| DYNC1LI1 |

| |
|---|
| DYNC1LI2 |
| DYNC2H1 |
| DYNC2I1 |
| DYNC2I2 |
| DYNC2LI1 |
| DYNLL1 |
| DYNLL2 |
| DYNLRB1 |
| DYNLRB2 |
| DYNLT1 |
| DYNLT2 |
| DYNLT2B |
| DYNLT3 |
| DYNLT4 |
| DYRK1A |
| DYRK1B |
| DYRK2 |
| DYRK3 |
| DZANK1 |
| DZIP1 |
| DZIP3 |
| E2F1 |
| E2F2 |
| E2F3 |
| E2F4 |
| E2F5 |
| E2F6 |
| E2F7 |
| E4F1 |
| EAF1 |
| EAF2 |
| EAPP |
| EARS2 |
| EBAG9 |
| EBF4 |
| EBNA1BP2 |
| EBNABP2 |
| EBP |
| EBPL |
| ECD |
| ECE1 |
| ECH1 |
| ECHDC1 |

| |
|---|
| ECHDC3 |
| ECHS1 |
| ECI1 |
| ECI2 |
| ECM2 |
| ECPAS |
| ECRG4 |
| ECSIT |
| ECT2 |
| EDC3 |
| EDC4 |
| EDEM1 |
| EDEM2 |
| EDEM3 |
| EDF1 |
| EDIL3 |
| EDN1 |
| EDNRA |
| EDRF1 |
| EEA1 |
| EED |
| EEF1A1 |
| EEF1A1P5 |
| EEF1AKMT1 |
| EEF1AKMT2 |
| EEF1AKMT4 |
| EEF1B2 |
| EEF1D |
| EEF1E1 |
| EEF1G |
| EEF2 |
| EEF2K |
| EEF2KMT |
| EEFSEC |
| EEPD1 |
| EFCAB11 |
| EFCAB14 |
| EFCAB2 |
| EFCAB6 |
| EFCAB7 |
| EFEMP2 |
| EFHC2 |
| EFHD2 |

| |
|---|
| EFL1 |
| EFNA1 |
| EFNA2 |
| EFNA5 |
| EFNB1 |
| EFNB2 |
| EFR3A |
| EFTUD2 |
| EGFL6 |
| EGFL7 |
| EGFL8 |
| EGLN1 |
| EGLN2 |
| EGLN3 |
| EGR1 |
| EGR2 |
| EGR3 |
| EHBP1 |
| EHBP1L1 |
| EHD1 |
| EHD2 |
| EHD4 |
| EHF |
| EHHADH |
| EHMT1 |
| EHMT2 |
| EI24 |
| EID1 |
| EID2 |
| EIF1 |
| EIF1AX |
| EIF1B |
| EIF2AK1 |
| EIF2AK2 |
| EIF2AK3 |
| EIF2AK4 |
| EIF2B1 |
| EIF2B2 |
| EIF2B4 |
| EIF2B5 |
| EIF2D |
| EIF2S1 |
| EIF2S2 |

| |
|---|
| EIF2S3 |
| EIF3A |
| EIF3B |
| EIF3D |
| EIF3E |
| EIF3F |
| EIF3G |
| EIF3I |
| EIF3J |
| EIF3K |
| EIF3L |
| EIF3M |
| EIF4A1 |
| EIF4A2 |
| EIF4A3 |
| EIF4B |
| EIF4E |
| EIF4E2 |
| EIF4E3 |
| EIF4EBP1 |
| EIF4EBP2 |
| EIF4ENIF1 |
| EIF4G1 |
| EIF4G2 |
| EIF4G3 |
| EIF4H |
| EIF5 |
| EIF5A2 |
| EIF5B |
| EIPR1 |
| ELAC1 |
| ELAC2 |
| ELAPOR2 |
| ELAV1 |
| ELAVL2 |
| ELF1 |
| ELF2 |
| ELF3 |
| ELF4 |
| ELK1 |
| ELK3 |
| ELK4 |
| ELL |

| |
|---|
| ELL2 |
| ELMO1 |
| ELMO2 |
| ELMO3 |
| ELMOD1 |
| ELMOD2 |
| ELMOD3 |
| ELOA |
| ELOB |
| ELOC |
| ELOVL1 |
| ELOVL3 |
| ELOVL5 |
| ELOVL6 |
| ELP1 |
| ELP2 |
| ELP3 |
| ELP4 |
| ELP5 |
| ELP6 |
| EMB |
| EMC1 |
| EMC10 |
| EMC2 |
| EMC3 |
| EMC4 |
| EMC6 |
| EMC7 |
| EMC8 |
| EMC9 |
| EMCN |
| EMD |
| EME1 |
| EME2 |
| EMG1 |
| EMID1 |
| EMILIN1 |
| EMILIN2 |
| EMILIN3 |
| EML1 |
| EML2 |
| EML3 |
| EML4 |

| |
|---|
| EML6 |
| EMP1 |
| EMP2 |
| EMP3 |
| EMSY |
| EMX2 |
| ENAH |
| ENC1 |
| ENDOD1 |
| ENDOG |
| ENDOV |
| ENGASE |
| ENHO |
| ENKD1 |
| ENO1A |
| ENO3 |
| ENOPH1 |
| ENOSF1 |
| ENOX1 |
| ENOX2 |
| ENPEP |
| ENPP1 |
| ENPP3 |
| ENPP4 |
| ENPP5 |
| ENPP6 |
| ENPP7 |
| ENSA |
| ENTPD1 |
| ENTPD2 |
| ENTPD3 |
| ENTPD4 |
| ENTPD5 |
| ENTPD6 |
| ENTPD7 |
| ENTPD8 |
| ENY2 |
| EOLA1 |
| EP400 |
| EPAS1 |
| EPB41 |
| EPB41L2 |
| EPB41L3 |

| |
|---|
| EPB41L4B |
| EPB41L5 |
| EPC1 |
| EPC2 |
| EPCAM |
| EPDR1 |
| EPG5 |
| EPHA1 |
| EPHA2 |
| EPHA4 |
| EPHA6 |
| EPHB1 |
| EPHB4 |
| EPHB6 |
| EPHX1 |
| EPHX2 |
| EPM2A |
| EPM2AIP1 |
| EPN1 |
| EPN2 |
| EPN3 |
| EPOP |
| EPRS1 |
| EPS15 |
| EPS15L1 |
| EPS8L2 |
| EPS8L3 |
| EPSTI1 |
| ERAL1 |
| ERAP1 |
| ERAS |
| ERBB2 |
| ERBB3 |
| ERBB4 |
| ERBIN |
| ERCC1 |
| ERCC2 |
| ERCC3 |
| ERCC4 |
| ERCC5 |
| ERCC6 |
| ERCC6L |
| ERCC6L2 |

| |
|---|
| ERCC8 |
| EREG |
| ERF |
| ERG28 |
| ERGIC1 |
| ERGIC2 |
| ERGIC3 |
| ERH |
| ERI1 |
| ERI2 |
| ERI3 |
| ERICH4 |
| ERLEC1 |
| ERLIN1 |
| ERLIN2 |
| ERMARD |
| ERMP1 |
| ERN1 |
| ERN2 |
| ERO1A |
| ERO1B |
| ERP27 |
| ERP29 |
| ERP44 |
| ERRFI1 |
| ERVFC1 |
| ERVW-4 |
| ESD |
| ESF1 |
| ESPL1 |
| ESPN |
| ESPNL |
| ESR1 |
| ESR2 |
| ESRP1 |
| ESRP2 |
| ESRRA |
| ESRRG |
| ESS2 |
| ESYT1 |
| ESYT2 |
| ETAA1 |
| ETF1 |

| |
|---|
| ETFA |
| ETFB |
| ETFBKMT |
| ETFDH |
| ETFRF1 |
| ETHE1 |
| ETNK1 |
| ETNK2 |
| ETS1 |
| ETS2 |
| ETV2 |
| ETV3 |
| ETV6 |
| EURL |
| EVA1B |
| EVC |
| EVC2 |
| EVI5 |
| EVI5L |
| EVL |
| EVPL |
| EVX1 |
| EWSR1 |
| EXD2 |
| EXD3 |
| EXO1 |
| EXO5 |
| EXOC1 |
| EXOC2 |
| EXOC3 |
| EXOC3L1 |
| EXOC3L2 |
| EXOC4 |
| EXOC5 |
| EXOC6 |
| EXOC6B |
| EXOC7 |
| EXOC8 |
| EXOG |
| EXOSC1 |
| EXOSC10 |
| EXOSC2 |
| EXOSC3 |

| |
|---|
| EXOSC4 |
| EXOSC5 |
| EXOSC6 |
| EXOSC7 |
| EXOSC8 |
| EXOSC9 |
| EXPH5 |
| EXT1 |
| EXT2 |
| EXTL1 |
| EXTL2 |
| EXTL3 |
| EYA3 |
| EZH1 |
| EZH2 |
| F11R |
| F12 |
| F2RL1 |
| F2RL3 |
| F3 |
| F5 |
| F8A1 |
| FA2H |
| FAAH |
| FAAP24 |
| FABP3 |
| FABP4 |
| FABP5 |
| FABP7 |
| FABPL |
| FADD |
| FADS1 |
| FADS2 |
| FADS3 |
| FAF1 |
| FAF2 |
| FAH |
| FAHD1 |
| FAHD2A |
| FAIM |
| FAM104A |
| FAM110A |
| FAM110B |

| |
|---|
| FAM110C |
| FAM111A |
| FAM111B |
| FAM114A1 |
| FAM114A2 |
| FAM117A |
| FAM117B |
| FAM118A |
| FAM118B |
| FAM120A |
| FAM120B |
| FAM120C |
| FAM126B |
| FAM131A |
| FAM131C |
| FAM133A |
| FAM133B |
| FAM135A |
| FAM13A |
| FAM13B |
| FAM13C |
| FAM149A |
| FAM149B1 |
| FAM151B |
| FAM155A |
| FAM156A |
| FAM161A |
| FAM161B |
| FAM162A |
| FAM168A |
| FAM168B |
| FAM169A |
| FAM170A |
| FAM171A1 |
| FAM171A2 |
| FAM171B |
| FAM172BP |
| FAM174A |
| FAM174C |
| FAM177A1 |
| FAM177B |
| FAM178B |
| FAM181B |

| |
|---|
| FAM184B |
| FAM185A |
| FAM189A1 |
| FAM189B |
| FAM193A |
| FAM193B |
| FAM199X |
| FAM200A |
| FAM200B |
| FAM20A |
| FAM20B |
| FAM20C |
| FAM210A |
| FAM210B |
| FAM213B |
| FAM214A |
| FAM214B |
| FAM216B |
| FAM217A |
| FAM217B |
| FAM219A |
| FAM219B |
| FAM220A |
| FAM222B |
| FAM229A |
| FAM229B |
| FAM234B |
| FAM241A |
| FAM241B |
| FAM24A |
| FAM26E |
| FAM32A |
| FAM3B |
| FAM3D |
| FAM43A |
| FAM50A |
| FAM50B |
| FAM53B |
| FAM53C |
| FAM71A |
| FAM71E1 |
| FAM72A |
| FAM76A |

| |
|---|
| FAM76B |
| FAM78A |
| FAM81A |
| FAM83C |
| FAM83D |
| FAM83E |
| FAM83F |
| FAM83G |
| FAM83H |
| FAM89B |
| FAM8A1 |
| FAM91A1 |
| FAM92A1 |
| FAM98A |
| FAM98B |
| FAM98C |
| FAN1 |
| FANCA |
| FANCB |
| FANCC |
| FANCD2 |
| FANCE |
| FANCF |
| FANCG |
| FANCI |
| FANCL |
| FANCM |
| FAR1 |
| FAR2 |
| FARP2 |
| FARS2 |
| FARSA |
| FARSB |
| FAS |
| FASTK |
| FASTKD1 |
| FASTKD2 |
| FASTKD3 |
| FASTKD5 |
| FAT1 |
| FAT2 |
| FAT4 |
| FAU |

| |
|---|
| FAXDC2 |
| FBF1 |
| FBH1 |
| FBLIM1 |
| FBLN1 |
| FBLN2 |
| FBLN5 |
| FBN1 |
| FBP1 |
| FBP2 |
| FBRSL1 |
| FBX30 |
| FBXL12 |
| FBXL14 |
| FBXL15 |
| FBXL17 |
| FBXL18 |
| FBXL19 |
| FBXL2 |
| FBXL20 |
| FBXL22 |
| FBXL4 |
| FBXL5 |
| FBXL7 |
| FBXL8 |
| FBXO10 |
| FBXO11 |
| FBXO15 |
| FBXO17 |
| FBXO21 |
| FBXO22 |
| FBXO24 |
| FBXO25 |
| FBXO28 |
| FBXO3 |
| FBXO31 |
| FBXO32 |
| FBXO33 |
| FBXO34 |
| FBXO38 |
| FBXO4 |
| FBXO42 |
| FBXO43 |

| |
|---|
| FBXO45 |
| FBXO46 |
| FBXO5 |
| FBXO6 |
| FBXO7 |
| FBXO8 |
| FBXO9 |
| FBXW11 |
| FBXW12 |
| FBXW2 |
| FBXW4 |
| FBXW5 |
| FBXW8 |
| FBXW9 |
| FCF1 |
| FCGBP |
| FCGR1B |
| FCGR1B?? |
| FCGR2A |
| FCGRT |
| FCHO1 |
| FCHO2 |
| FCHSD1 |
| FCHSD2 |
| FCRL3 |
| FCRL5 |
| FCSK |
| FDFT1 |
| FDPS |
| FDX1 |
| FDX2 |
| FDXACB1 |
| FDXR |
| FECH |
| FEM1A |
| FEM1B |
| FEM1C |
| FEN1 |
| FER |
| FER1L6 |
| FERMT2 |
| FES |
| FEZ2 |

| |
|---|
| FFAR4 |
| FGD1 |
| FGD2 |
| FGD3 |
| FGD4 |
| FGD5 |
| FGD6 |
| FGF1 |
| FGF13 |
| FGF17 |
| FGF2 |
| FGFBP1 |
| FGFR1OP2 |
| FGFR2 |
| FGFR3 |
| FGGY |
| FGL2 |
| FGR |
| FH |
| FHDC1 |
| FHIP1A |
| FHIP1B |
| FHIP2A |
| FHIP2B |
| FHL2 |
| FHL3 |
| FHOD1 |
| FHOD3 |
| FIBP |
| FICD |
| FIG4 |
| FIGN |
| FIGNL1 |
| FILIP1 |
| FILIP1L |
| FIP1L1 |
| FITM2 |
| FIZ1 |
| FJX1 |
| FKBP10 |
| FKBP11 |
| FKBP15 |
| FKBP3 |

| |
|---|
| FKBP5 |
| FKBP7 |
| FKBP8 |
| FKBP9 |
| FKBPL |
| FKRP |
| FKTN |
| FLAD1 |
| FLCN |
| FLI1 |
| FLII |
| FLNA |
| FLNB |
| FLOT1 |
| FLOT2 |
| FLRT1 |
| FLT3 |
| FLVCR1 |
| FLVCR2 |
| FLYWCH1 |
| FLYWCH2 |
| FMN1 |
| FMN2 |
| FMNL1 |
| FMNL2 |
| FMNL3 |
| FMO5 |
| FMR1 |
| FN3K |
| FN3KRP |
| FNBP1 |
| FNBP1L |
| FNBP4 |
| FNDC1 |
| FNDC10 |
| FNDC3A |
| FNDC3B |
| FNDC4 |
| FNDC9 |
| FNIP1 |
| FNIP2 |
| FNTA |
| FNTB |

| |
|---|
| FOCAD |
| FOLR1 |
| FOLR2 |
| FOS |
| FOSB |
| FOSL2 |
| FOXA1 |
| FOXA3 |
| FOXC1 |
| FOXC2 |
| FOXD2 |
| FOXD4L1 |
| FOXG1 |
| FOXH1 |
| FOXJ3 |
| FOXK1 |
| FOXK2 |
| FOXL2 |
| FOXM1 |
| FOXN2 |
| FOXO1 |
| FOXO3 |
| FOXO4 |
| FOXP1 |
| FOXP4 |
| FOXRED1 |
| FOXRED2 |
| FOXS1 |
| FPGS |
| FPGT |
| FPR2 |
| FRA10AC1 |
| FRAS1 |
| FRAT1 |
| FRAT2 |
| FREM2 |
| FRG1 |
| FRK |
| FRMD4B |
| FRMD6 |
| FRMD8 |
| FRMPD1 |
| FRRS1 |

| |
|---|
| FRRS1L |
| FRS2 |
| FRS3 |
| FRY |
| FRYL |
| FSCN1 |
| FSD1L |
| FSD2 |
| FSIP2 |
| FST |
| FSTL1 |
| FSTL3 |
| FTCD |
| FTH1 |
| FTL |
| FTO |
| FTSJ1 |
| FTSJ3 |
| FUBP1 |
| FUBP3 |
| FUCA1 |
| FUCA2 |
| FUNDC2 |
| FUOM |
| FURIN |
| FUS |
| FUT1 |
| FUT2 |
| FUT4 |
| FUZ |
| FXR1 |
| FXR2 |
| FXYD3 |
| FXYD4 |
| FXYD5 |
| FXYD6 |
| FYB1 |
| FYB2 |
| FYCO1 |
| FYTTD1 |
| FZD1 |
| FZD2 |
| FZD3 |

| |
|---|
| FZD4 |
| FZD5 |
| FZD6 |
| FZD7 |
| FZD8 |
| FZD9 |
| FZR1 |
| G0S2 |
| G2E3 |
| G3BP1 |
| G3BP2 |
| G6PC3 |
| G6PD |
| GAA |
| GAB1 |
| GAB2 |
| GABARAP |
| GABBR1 |
| GABPA |
| GABPB1 |
| GABPB2 |
| GABRA2 |
| GABRA3 |
| GABRG1 |
| GABRR2 |
| GADD45A |
| GADD45B |
| GADD45G |
| GADD45GIP1 |
| GADL1 |
| GAK |
| GAL3ST1 |
| GAL3ST2 |
| GAL3ST3 |
| GAL3ST4 |
| GALC |
| GALE |
| GALK1 |
| GALK2 |
| GALM |
| GALNS |
| GALNT1 |
| GALNT10 |

| |
|---|
| GALNT11 |
| GALNT12 |
| GALNT13 |
| GALNT16 |
| GALNT18 |
| GALNT2 |
| GALNT3 |
| GALNT4 |
| GALNT5 |
| GALNT8 |
| GALR3 |
| GALT |
| GAMT |
| GAN |
| GANAB |
| GANC |
| GAPDH |
| GAPDHS |
| GAPT |
| GAR1 |
| GAREM1 |
| GARIN5B |
| GARNL3 |
| GARRE1 |
| GARS1 |
| GAS1 |
| GAS2L1 |
| GAS6 |
| GAS7 |
| GAS8 |
| GASK1B |
| GASP2 |
| GATA4 |
| GATA6 |
| GATAD2A |
| GATAD2B |
| GATB |
| GATC |
| GATD1 |
| GATD3B |
| GBA |
| GBA2 |
| GBA3 |

| |
|---|
| GBAP1 |
| GBE1 |
| GBGT1 |
| GBP6 |
| GCA |
| GCAT |
| GCC1 |
| GCC2 |
| GCDH |
| GCFC2 |
| GCH1 |
| GCHFR |
| GCLM |
| GCN1 |
| GCNA |
| GCNT1 |
| GCNT2 |
| GCNT3 |
| GCSH |
| GCSP |
| GDA |
| GDAP2 |
| GDE1 |
| GDF1 |
| GDF9 |
| GDI1 |
| GDI2 |
| GDPD3 |
| GDPD5 |
| GDPGP1 |
| GEM |
| GEMIN2 |
| GEMIN4 |
| GEMIN5 |
| GEMIN6 |
| GEMIN7 |
| GEMIN8 |
| GEN1 |
| GET1 |
| GET3 |
| GET4 |
| GFER |
| GFM1 |

| |
|---|
| GFM2 |
| GFOD2 |
| GFPT1 |
| GFPT2 |
| GFUS |
| GGA1 |
| GGA2 |
| GGA3 |
| GGCT |
| GGCX |
| GGH |
| GGNBP2 |
| GGPS1 |
| GGT1 |
| GGT7 |
| GGTA1 |
| GHDC |
| GHITM |
| GHR |
| GID4 |
| GID8 |
| GIGYF1 |
| GIGYF2 |
| GIMAP1 |
| GIMAP4 |
| GIMAP6 |
| GIMAP7 |
| GIMAP8 |
| GIN1 |
| GINM1 |
| GINS1 |
| GINS2 |
| GINS3 |
| GINS4 |
| GIPC1 |
| GIPC2 |
| GIT1 |
| GIT2 |
| GJA1 |
| GJA5 |
| GJA8 |
| GJA9 |
| GJB3 |

| |
|---|
| GJC1 |
| GK |
| GK2 |
| GK5 |
| GKAP1 |
| GLA |
| GLB1 |
| GLB1L |
| GLCCI1 |
| GLDC |
| GLDN |
| GLG1 |
| GLI3 |
| GLI4 |
| GLIPR1L2 |
| GLIPR2 |
| GLIS2 |
| GLIS3 |
| GLMN |
| GLO1 |
| GLOD4 |
| GLRX |
| GLRX2 |
| GLRX3 |
| GLRX5 |
| GLS |
| GLS2 |
| GLT1D1 |
| GLT8D1 |
| GLT8D2 |
| GLUD1 |
| GLUL |
| GLYCTK |
| GLYR1 |
| GM2A |
| GMCL1 |
| GMDS |
| GMEB1 |
| GMEB2 |
| GMFB |
| GMFG |
| GMIP |
| GMNN |

| |
|---|
| GMPPA |
| GMPR |
| GMPR2 |
| GMPS |
| GNA11 |
| GNA12 |
| GNA13 |
| GNA14 |
| GNAI1 |
| GNAI2 |
| GNAL |
| GNAQ |
| GNAS |
| GNAZ |
| GNB1 |
| GNB1L |
| GNB2 |
| GNB4 |
| GNB5 |
| GNE |
| GNG10 |
| GNG11 |
| GNG12 |
| GNG3 |
| GNG5 |
| GNG7 |
| GNL1 |
| GNL2 |
| GNL3 |
| GNPAT |
| GNPDA1 |
| GNPDA2 |
| GNPNAT1 |
| GNPTAB |
| GNPTG |
| GNS |
| GOLGA2 |
| GOLGA3 |
| GOLGA5 |
| GOLGA7 |
| GOLGB1 |
| GOLIM4 |
| GOLM1 |

| |
|---|
| GOLPH3 |
| GOLPH3L |
| GOLT1B |
| GON4L |
| GON7 |
| GOPC |
| GORAB |
| GORASP2 |
| GOSR1 |
| GOSR2 |
| GOT1 |
| GOT2 |
| GP1BA |
| GP5 |
| GPA33 |
| GPAA1 |
| GPALPP1 |
| GPAM |
| GPANK1 |
| GPAT4 |
| GPATCH1 |
| GPATCH11 |
| GPATCH2 |
| GPATCH2L |
| GPATCH3 |
| GPATCH4 |
| GPATCH8 |
| GPBAR1 |
| GPBP1 |
| GPBP1L1 |
| GPC1 |
| GPC2 |
| GPC4 |
| GPC6 |
| GPCPD1 |
| GPD1 |
| GPD1L |
| GPD2 |
| GPER1 |
| GPHN |
| GPI |
| GPKOW |
| GPM6A |

| |
|---|
| GPN1 |
| GPN2 |
| GPN3 |
| GPNMB |
| GPR101 |
| GPR108 |
| GPR132 |
| GPR135 |
| GPR137 |
| GPR137B |
| GPR141 |
| GPR143 |
| GPR146 |
| GPR15 |
| GPR153 |
| GPR155 |
| GPR157 |
| GPR162 |
| GPR173 |
| GPR176 |
| GPR179 |
| GPR18 |
| GPR183 |
| GPR27 |
| GPR3 |
| GPR31 |
| GPR34 |
| GPR35 |
| GPR39 |
| GPR4 |
| GPR52 |
| GPR63 |
| GPR65 |
| GPR68 |
| GPR75 |
| GPR82 |
| GPR84 |
| GPR85 |
| GPR88 |
| GPR89A |
| GPRASP1 |
| GPRC5A |
| GPRC5B |

| |
|---|
| GPRC5C |
| GPRC5D |
| GPRIN3 |
| GPS1 |
| GPSM1 |
| GPSM2 |
| GPT |
| GPT2 |
| GPX1 |
| GPX3 |
| GPX7 |
| GPX8 |
| GRAMD1A |
| GRAMD1B |
| GRAMD1C |
| GRAMD2B |
| GRB2 |
| GRB7 |
| GREB1 |
| GREM1 |
| GREM2 |
| GRHL1 |
| GRHPR |
| GRID1 |
| GRID2 |
| GRIK5 |
| GRIN2 |
| GRIN2C |
| GRIN3A |
| GRIN3B |
| GRINA |
| GRIP1 |
| GRIPAP1 |
| GRK1 |
| GRK2 |
| GRK3 |
| GRK4 |
| GRK5 |
| GRK6 |
| GRM2 |
| GRM3 |
| GRM8 |
| GRN |

| |
|---|
| GRPEL1 |
| GRPEL2 |
| GRSF1 |
| GRTP1 |
| GRWD1 |
| GSAP |
| GSDMB |
| GSE1 |
| GSG1 |
| GSK3A |
| GSK3B |
| GSKIP |
| GSPT1 |
| GSS |
| GSTA1 |
| GSTA4 |
| GSTCD |
| GSTK1 |
| GSTM2 |
| GSTM3 |
| GSTM5 |
| GSTO1 |
| GSTP1 |
| GSTZ1 |
| GSX1 |
| GTDC1 |
| GTF2A1 |
| GTF2A2 |
| GTF2B |
| GTF2E1 |
| GTF2E2 |
| GTF2F1 |
| GTF2F2 |
| GTF2H1 |
| GTF2H2 |
| GTF2H3 |
| GTF2I |
| GTF2IRD1 |
| GTF3A |
| GTF3C1 |
| GTF3C2 |
| GTF3C3 |
| GTF3C4 |

| |
|---|
| GTF3C5 |
| GTPBP1 |
| GTPBP10 |
| GTPBP10 |
| GTPBP2 |
| GTPBP3 |
| GTPBP4 |
| GTPBP6 |
| GTPBP8 |
| GTSF1L |
| GUCA2B |
| GUCD1 |
| GUCY1A1 |
| GUCY1B1 |
| GUCY2C |
| GUCY2D |
| GUF1 |
| GUK1 |
| GULP1 |
| GUSB |
| GVINP1 |
| GXYLT1 |
| GYG1 |
| GYG2 |
| GYS1 |
| GZF1 |
| GZMA |
| H1-0 |
| H1-1 |
| H1-10 |
| H1-4 |
| H2AC1 |
| H2AC11 |
| H2AC12 |
| H2AC19 |
| H2AC20 |
| H2AC21 |
| H2AC4 |
| H2AC6 |
| H2AC7 |
| H2AC8 |
| H2AJ |
| H2AP |

| |
|---|
| H2AX |
| H2AZ1 |
| H2AZ2 |
| H2BC1 |
| H2BC11 |
| H2BC12 |
| H2BC13 |
| H2BC14 |
| H2BC15 |
| H2BC18 |
| H2BC21 |
| H2BC3 |
| H2BC9 |
| H2BE1 |
| H3-3A |
| H3C1 |
| H4C1 |
| H6PD |
| HABP4 |
| HACD1 |
| HACD2 |
| HACD3 |
| HACD4 |
| HACE1 |
| HACL1 |
| HADH |
| HADHA |
| HAGH |
| HAGHL |
| HAPLN1 |
| HAPLN3 |
| HARBI1 |
| HARS1 |
| HARS2 |
| HAS3 |
| HAT1 |
| HAUS1 |
| HAUS2 |
| HAUS3 |
| HAUS4 |
| HAUS5 |
| HAUS6 |
| HAUS7 |

| |
|---|
| HAUS8 |
| HAVCR2 |
| HAX1 |
| HBB |
| HBE1 |
| HBEGF |
| HBP1 |
| HBS1L |
| HCAR2 |
| HCCS |
| HCFC1 |
| HCFC1R1 |
| HCFC2 |
| HCK |
| HCN1 |
| HCRT |
| HDAC1 |
| HDAC10 |
| HDAC11 |
| HDAC2 |
| HDAC3 |
| HDAC4 |
| HDAC5 |
| HDAC6 |
| HDAC7 |
| HDAC8 |
| HDAC9 |
| HDC |
| HDDC2 |
| HDDC3 |
| HDGF |
| HDGFL2 |
| HDGFL3 |
| HDHD2 |
| HDHD3 |
| HDHD5 |
| HDLBP |
| HEATR1 |
| HEATR3 |
| HEATR4 |
| HEATR5B |
| HEATR6 |
| HEBP1 |

| |
|---|
| HEBP2 |
| HECTD1 |
| HECTD2 |
| HECTD3 |
| HECTD4 |
| HEG1 |
| HELB |
| HELLS |
| HELQ |
| HELZ |
| HELZ2 |
| HEMK1 |
| HEPACAM2 |
| HEPH |
| HERC1 |
| HERC2 |
| HERC4 |
| HERC5 |
| HERPUD2 |
| HES4 |
| HES6 |
| HEXA |
| HEXB |
| HEXD |
| HEXIM1 |
| HEXIM2 |
| HEY2 |
| HFE |
| HFM1 |
| HGF |
| HGH1 |
| HGNAT |
| HGS |
| HHAT |
| HHEX |
| HHIPL1 |
| HHIPL2 |
| HIBADH |
| HIC1 |
| HIC2 |
| HID1 |
| HIF1A |
| HIF1AN |

| |
|---|
| HIGD1A |
| HIGD2A |
| HIKESHI |
| HILPDA |
| HINFP |
| HINT1 |
| HINT2 |
| HINT3 |
| HIP1R |
| HIPK1 |
| HIPK3 |
| HIRA |
| HIRIP3 |
| HIVEP1 |
| HIVEP2 |
| HIVEP3 |
| HJURP |
| HK1 |
| HK2 |
| HK3 |
| HKDC1 |
| HLA-A |
| HLA-DMB |
| HLA-DQB1 |
| HLCS |
| HLX |
| HM13 |
| HMBS |
| HMCN1 |
| HMCN2 |
| HMG20A |
| HMG20B |
| HMGA1 |
| HMGB1 |
| HMGB2 |
| HMGB3 |
| HMGCL |
| HMGCR |
| HMGCS1 |
| HMGN1 |
| HMGN2 |
| HMGN3 |
| HMGN4 |

| |
|---|
| HMGN5 |
| HMGXB4 |
| HMMR |
| HMOX1 |
| HMOX2 |
| HNF4A |
| HNF4G |
| HNMT |
| HNRNPA0 |
| HNRNPA1 |
| HNRNPA2 |
| HNRNPA3 |
| HNRNPAB |
| HNRNPC |
| HNRNPD |
| HNRNPDL |
| HNRNPF |
| HNRNPH1 |
| HNRNPH2 |
| HNRNPH3 |
| HNRNPK |
| HNRNPL |
| HNRNPM |
| HNRNPR |
| HNRNPU |
| HNRNPUL1 |
| HNRNPUL2 |
| HOMER1 |
| HOMER2 |
| HOMER3 |
| HOMEZ |
| HOOK1 |
| HOOK2 |
| HOOK3 |
| HOXA10 |
| HOXA13 |
| HOXB13 |
| HOXC10 |
| HOXC11 |
| HOXC4 |
| HOXC5 |
| HOXC6 |
| HOXC8 |

| |
|---|
| HOXC9 |
| HOXD11 |
| HOXD3 |
| HOXD4 |
| HOXD8 |
| HP1BP3 |
| HPDL |
| HPF1 |
| HPPD |
| HPRT |
| HPS1 |
| HPS3 |
| HPS4 |
| HPS6 |
| HPSE |
| HRAS |
| HRCT1 |
| HRH1 |
| HRH2 |
| HROB |
| HS1BP3 |
| HS2ST1 |
| HS3ST1 |
| HS3ST2 |
| HS3ST3A1 |
| HS6ST1 |
| HS6ST3 |
| HSC20 |
| HSD11B1L |
| HSD11B2 |
| HSD17B1 |
| HSD17B10 |
| HSD17B12 |
| HSD17B14 |
| HSD17B2 |
| HSD17B3 |
| HSD17B4 |
| HSD17B7 |
| HSD17B8 |
| HSD3B2 |
| HSD3B7 |
| HSDL1 |
| HSDL2 |

| |
|---|
| HSF1 |
| HSF2 |
| HSF2BP |
| HSF4 |
| HSFX1 |
| HSFX3 |
| HSH2D |
| HSP90AA1 |
| HSP90AB1 |
| HSP90B1 |
| HSPA12B |
| HSPA13 |
| HSPA14 |
| HSPA1A |
| HSPA1L |
| HSPA4L |
| HSPA5 |
| HSPA6 |
| HSPA8 |
| HSPA9 |
| HSPB1 |
| HSPB11 |
| HSPB6 |
| HSPBAP1 |
| HSPBP1 |
| HSPD1 |
| HSPE1 |
| HSPG2 |
| HTATIP2 |
| HTATSF1 |
| HTN3 |
| HTR1F |
| HTR3A |
| HTR5A |
| HTR7 |
| HTRA1 |
| HTRA2 |
| HUS1 |
| HUWE1 |
| HVCN1 |
| HYAL1 |
| HYAL2 |
| HYI |

| |
|---|
| HYKK |
| HYLS1 |
| HYOU1 |
| HYPK |
| IAH1 |
| IARS1 |
| IARS2 |
| IBA57 |
| IBTK |
| ICA1L |
| ICA69 |
| ICAM1 |
| ICAM2 |
| ICAM3 |
| ICE1 |
| ICE2 |
| ICMT |
| ICOSLG |
| ID1 |
| ID2 |
| ID3 |
| IDE |
| IDH1 |
| IDH2 |
| IDH3A |
| IDH3B |
| IDI1 |
| IDNK |
| IDO1 |
| IDS |
| IDUA |
| IER2 |
| IER3 |
| IER3IP1 |
| IER5 |
| IER5L |
| IFFO2 |
| IFI16 |
| IFI27 |
| IFI27L2 |
| IFI30 |
| IFI35 |
| IFI44L |

| |
|---|
| IFI6 |
| IFIH1 |
| IFIT1 |
| IFIT2 |
| IFIT5 |
| IFITM3 |
| IFNA4 |
| IFNAR1 |
| IFNAR2 |
| IFND1 |
| IFNGR1 |
| IFNGR2 |
| IFNLR1 |
| IFNW1 |
| IFRD1 |
| IFRD2 |
| IFT122 |
| IFT172 |
| IFT20 |
| IFT22 |
| IFT25 |
| IFT27 |
| IFT43 |
| IFT46 |
| IFT52 |
| IFT57 |
| IFT74 |
| IFT80 |
| IFT81 |
| IFT88 |
| IFTAP |
| IGBP1 |
| IGDCC4 |
| IGF1R |
| IGF2 |
| IGF2BP2 |
| IGFBP2 |
| IGFBP3 |
| IGFBP4 |
| IGFBP5 |
| IGFBP6 |
| IGFBP7 |
| IGHMBP2 |

| |
|---|
| IGSF11 |
| IGSF6 |
| IGSF8 |
| IGSF9 |
| IHH |
| IK |
| IKBIP |
| IKBKB |
| IKBKE |
| IKBKG |
| IKZF1 |
| IKZF2 |
| IKZF3 |
| IKZF4 |
| IKZF5 |
| IL10 |
| IL10RA |
| IL10RB |
| IL11RA |
| IL13RA1 |
| IL15 |
| IL16 |
| IL17D |
| IL17RA |
| IL17RC |
| IL17RD |
| IL18 |
| IL18BP |
| IL18RAP |
| IL1A |
| IL1RAP |
| IL1RN |
| IL20RA |
| IL27RA |
| IL2RB |
| IL2RG |
| IL32 |
| IL34 |
| IL3RA |
| IL4I1 |
| IL4R |
| IL6 |
| IL6ST |

| |
|---|
| ILDR1 |
| ILF2 |
| ILF3 |
| ILK |
| ILKAP |
| ILRUN |
| ILVBL |
| IMMT |
| IMP3 |
| IMP4 |
| IMPA1 |
| IMPA2 |
| IMPACT |
| IMPDH1 |
| IMPDH2 |
| IMUP |
| INAFM2 |
| INAVA |
| INCENP |
| ING1 |
| ING2 |
| ING3 |
| ING4 |
| ING5 |
| INHA |
| INHBA |
| INHBB |
| INIP |
| INKA1 |
| INKA2 |
| INO80 |
| INO80B |
| INO80C |
| INO80D |
| INO80E |
| INPP1 |
| INPP4A |
| INPP5A |
| INPP5B |
| INPP5D |
| INPP5E |
| INPP5F |
| INPP5K |

| |
|---|
| INPPL1 |
| INSIG1 |
| INSIG2 |
| INSL6 |
| INSR |
| INSYN2A |
| INTS10 |
| INTS11 |
| INTS12 |
| INTS13 |
| INTS14 |
| INTS15 |
| INTS2 |
| INTS3 |
| INTS5 |
| INTS6 |
| INTS6L |
| INTS7 |
| INTS8 |
| INTS9 |
| INTU |
| IP6K1 |
| IPMK |
| IPO13 |
| IPO4 |
| IPO5 |
| IPO7 |
| IPO8 |
| IPP |
| IPPK |
| IQCB1 |
| IQCD |
| IQCE |
| IQCG |
| IQGAP1 |
| IQGAP2 |
| IQGAP3 |
| IQSEC1 |
| IQSEC2 |
| IRAG1 |
| IRAG2 |
| IRAK1 |
| IRAK1BP1 |

| |
|---|
| IRAK2 |
| IRAK3 |
| IRAK4 |
| IREB2 |
| IRF1 |
| IRF2 |
| IRF2BP1 |
| IRF2BP2 |
| IRF2BPL |
| IRF3 |
| IRF6 |
| IRF7 |
| IRF8 |
| IRS1 |
| IRS2 |
| IRX3 |
| IRX5 |
| ISCU |
| ISG20 |
| ISG20L2 |
| ISLR2 |
| ISOC1 |
| IST1 |
| ISX |
| ISY1 |
| ISYNA1 |
| ITFG1 |
| ITFG2 |
| ITGA3 |
| ITGA4 |
| ITGA6 |
| ITGAE |
| ITGAL |
| ITGAM |
| ITGAV |
| ITGAX |
| ITGB1 |
| ITGB1BP1 |
| ITGB1BP2 |
| ITGB3 |
| ITGB3BP |
| ITGB5 |
| ITGB7 |

| |
|---|
| ITGB8 |
| ITGBL1 |
| ITIH1 |
| ITM2A |
| ITM2B |
| ITM2C |
| ITPA |
| ITPK1 |
| ITPKA |
| ITPKB |
| ITPKC |
| ITPR2 |
| ITPR3 |
| ITPRID2 |
| ITPRIP |
| ITPRIPL1 |
| ITPRIPL2 |
| ITSN1 |
| ITSN2 |
| IVD |
| IVNS1ABP |
| IWS1 |
| JADE1 |
| JADE2 |
| JADE3 |
| JAG1 |
| JAGN1 |
| JAK1 |
| JAK3 |
| JAKMIP2 |
| JAM3 |
| JARID2 |
| JAZF1 |
| JCAD |
| JCHAIN |
| JDP2 |
| JHY |
| JKAMP |
| JMJD1C |
| JMJD4 |
| JMJD6 |
| JMJD7 |
| JMJD8 |

| |
|---|
| JMY |
| JOSD1 |
| JOSD2 |
| JPT1 |
| JPT2 |
| JRK |
| JRKL |
| JSRP1 |
| JTB |
| JUN |
| JUNB |
| JUND |
| JUP |
| KALRN |
| KANK1 |
| KANK2 |
| KANK3 |
| KANSL1 |
| KANSL1L |
| KANSL3 |
| KARS1 |
| KASH5 |
| KAT2A |
| KAT2B |
| KAT5 |
| KAT6A |
| KAT6B |
| KAT8 |
| KATNA1 |
| KATNAL1 |
| KATNB1 |
| KATNBL1 |
| KATNIP |
| KAZALD1 |
| KAZN |
| KBTBD11 |
| KBTBD2 |
| KBTBD3 |
| KBTBD6 |
| KBTBD7 |
| KCMF1 |
| KCNA2 |
| KCNA3 |

| |
|---|
| KCNA4 |
| KCNA5 |
| KCNA6 |
| KCNAB1 |
| KCNAB2 |
| KCNC2 |
| KCNC3 |
| KCND2 |
| KCNE2 |
| KCNE3 |
| KCNE4 |
| KCNE5 |
| KCNG2 |
| KCNH2 |
| KCNIP4 |
| KCNJ10 |
| KCNJ14 |
| KCNJ15 |
| KCNJ16 |
| KCNJ2 |
| KCNJ5 |
| KCNK1 |
| KCNK3 |
| KCNK5 |
| KCNN3 |
| KCNQ1 |
| KCNQ4 |
| KCNQ5 |
| KCNRG |
| KCNS3 |
| KCNT2 |
| KCTD1 |
| KCTD10 |
| KCTD11 |
| KCTD12 |
| KCTD14 |
| KCTD15 |
| KCTD18 |
| KCTD20 |
| KCTD21 |
| KCTD3 |
| KCTD5 |
| KCTD7 |

| |
|---|
| KCTD9 |
| KDELC1 |
| KDELR1 |
| KDELR2 |
| KDELR3 |
| KDF1 |
| KDM1A |
| KDM1B |
| KDM2A |
| KDM2B |
| KDM3A |
| KDM3B |
| KDM4A |
| KDM4B |
| KDM4C |
| KDM5A |
| KDM5B |
| KDM5C |
| KDM6A |
| KDM7A |
| KDM8 |
| KDSR |
| KEAP1 |
| KHDC4 |
| KHDRBS1 |
| KHNYN |
| KHSRP |
| KIAA0040 |
| KIAA0100 |
| KIAA0319 |
| KIAA0319L |
| KIAA0513 |
| KIAA0586 |
| KIAA0753 |
| KIAA0825 |
| KIAA0895 |
| KIAA0930 |
| KIAA1107 |
| KIAA1109 |
| KIAA1143 |
| KIAA1191 |
| KIAA1210 |
| KIAA1217 |

| |
|---|
| KIAA1328 |
| KIAA1522 |
| KIAA1549 |
| KIAA1586 |
| KIAA1614 |
| KIAA1671 |
| KIAA1958 |
| KIAA2013 |
| KIAA2026 |
| KICS2 |
| KIDINS220 |
| KIF11 |
| KIF13A |
| KIF13B |
| KIF14 |
| KIF15 |
| KIF16B |
| KIF18A |
| KIF18B |
| KIF19 |
| KIF1B |
| KIF1C |
| KIF20A |
| KIF20B |
| KIF22 |
| KIF23 |
| KIF24 |
| KIF26B |
| KIF2A |
| KIF2C |
| KIF3A |
| KIF3B |
| KIF3C |
| KIF4A |
| KIF7 |
| KIFAP3 |
| KIFBP |
| KIFC1 |
| KIFC2 |
| KIFC3 |
| KIN |
| KIRREL3 |
| KIT |

| |
|---|
| KITLG |
| KLC1 |
| KLC2 |
| KLC3 |
| KLC4 |
| KLDC9 |
| KLF11 |
| KLF12 |
| KLF13 |
| KLF15 |
| KLF16 |
| KLF2 |
| KLF4 |
| KLF5 |
| KLF6 |
| KLF7 |
| KLF8 |
| KLHDC1 |
| KLHDC10 |
| KLHDC2 |
| KLHDC4 |
| KLHDC7A |
| KLHDC8B |
| KLHL11 |
| KLHL12 |
| KLHL15 |
| KLHL17 |
| KLHL18 |
| KLHL2 |
| KLHL20 |
| KLHL21 |
| KLHL23 |
| KLHL24 |
| KLHL25 |
| KLHL26 |
| KLHL28 |
| KLHL3 |
| KLHL32 |
| KLHL36 |
| KLHL38 |
| KLHL41 |
| KLHL42 |
| KLHL5 |

| |
|---|
| KLHL7 |
| KLHL8 |
| KLHL9 |
| KLK14 |
| KLK15 |
| KMT2A |
| KMT2B |
| KMT2C |
| KMT2D |
| KMT2E |
| KMT5A |
| KMT5B |
| KMT5C |
| KNL1 |
| KNOP1 |
| KNSTRN |
| KPCB |
| KPNA3 |
| KPNA4 |
| KPNA5 |
| KPNA6 |
| KPNB1 |
| KRAS |
| KRBA1 |
| KRBA2 |
| KRBOX4 |
| KRCC1 |
| KREMEN1 |
| KRI1 |
| KRIT1 |
| KRR1 |
| KRT18 |
| KRT19 |
| KRT20 |
| KRT72 |
| KRT8 |
| KRTAP7-1 |
| KRTCAP2 |
| KSR1 |
| KTI12 |
| KTN1 |
| KXD1 |
| KY |

| |
|---|
| KYAT1 |
| KYAT3 |
| L1CAM |
| L2HGDH |
| L3HYPDH |
| L3MBTL1 |
| L3MBTL2 |
| L3MBTL3 |
| LACC1 |
| LACTB |
| LACTB2 |
| LAD1 |
| LAG3 |
| LAGE3 |
| LAMA1 |
| LAMA2 |
| LAMA3 |
| LAMA5 |
| LAMB1 |
| LAMB2 |
| LAMB3 |
| LAMC1 |
| LAMC3 |
| LAMP1 |
| LAMP2 |
| LAMTOR1 |
| LAMTOR2 |
| LAMTOR5 |
| LANCL2 |
| LAP3 |
| LAPTM4A |
| LAPTM4B |
| LAPTM5 |
| LARGE1 |
| LARGE2 |
| LARP1 |
| LARP1B |
| LARP4 |
| LARP4B |
| LARP6 |
| LARP7 |
| LARS1 |
| LARS2 |

| |
|---|
| LAS1L |
| LASP1 |
| LAT |
| LATS2 |
| LBH |
| LBR |
| LBX2 |
| LCA5 |
| LCK |
| LCLAT1 |
| LCMT1 |
| LCMT2 |
| LCN2 |
| LCOR |
| LCORL |
| LCP1 |
| LCP2 |
| LDAF1 |
| LDAH |
| LDB1 |
| LDHA |
| LDHAL6B |
| LDHB |
| LDHD |
| LDLR |
| LDLRAD2 |
| LDLRAD3 |
| LDLRAP1 |
| LDOC1 |
| LEF1 |
| LEMD2 |
| LEMD3 |
| LENG1 |
| LENG8 |
| LENG9 |
| LEO1 |
| LEPROT |
| LEPROTL1 |
| LETM1 |
| LETM2 |
| LETMD1 |
| LEXM |
| LFNG |

| |
|---|
| LGALS3 |
| LGALS3BP |
| LGALS8 |
| LGALS9 |
| LGALSL |
| LGI3 |
| LGMN |
| LGR4 |
| LGSN |
| LHCGR |
| LHFPL2 |
| LHPP |
| LIAS |
| LIF |
| LIFR |
| LIG1 |
| LIG3 |
| LIG4 |
| LILRA5 |
| LILRA6 |
| LIM2 |
| LIMA1 |
| LIMD1 |
| LIMD2 |
| LIMK1 |
| LIMK2 |
| LIMS1 |
| LIN37 |
| LIN52 |
| LIN54 |
| LIN7B |
| LIN7C |
| LIN9 |
| LINC02693 |
| LINGO2 |
| LINS1 |
| LIPA |
| LIPC |
| LIPE |
| LIPH |
| LIPK |
| LIPT1 |
| LIPT2 |

| |
|---|
| LITAF |
| LIX1L |
| LLGL2 |
| LLPH |
| LMAN1 |
| LMAN2 |
| LMAN2L |
| LMBRD1 |
| LMBRD2 |
| LMBRL |
| LMCD1 |
| LMF1 |
| LMF2 |
| LMLN |
| LMNA |
| LMNB1 |
| LMNB2 |
| LMNTD1 |
| LMNTD2 |
| LMO2 |
| LMO4 |
| LMO7 |
| LMTK2 |
| LNPEP |
| LNPK |
| LNX2 |
| LONP1 |
| LONP2 |
| LONRF1 |
| LONRF3 |
| LOXL3 |
| LOXL4 |
| LPAR2 |
| LPAR5 |
| LPAR6 |
| LPCAT3 |
| LPCAT4 |
| LPGAT1 |
| LPIN1 |
| LPIN3 |
| LPP |
| LPXN |
| LRATD1 |

| |
|---|
| LRATD2 |
| LRBA |
| LRCH1 |
| LRCH2 |
| LRCH3 |
| LRCH4 |
| LRFN1 |
| LRFN3 |
| LRFN4 |
| LRIF1 |
| LRIG1 |
| LRIG2 |
| LRIG3 |
| LRMDA |
| LRP1 |
| LRP10 |
| LRP12 |
| LRP4 |
| LRP5 |
| LRP6 |
| LRP8 |
| LRPAP1 |
| LRPPRC |
| LRR1 |
| LRRC1 |
| LRRC10B |
| LRRC14 |
| LRRC17 |
| LRRC20 |
| LRRC24 |
| LRRC25 |
| LRRC26 |
| LRRC27 |
| LRRC28 |
| LRRC31 |
| LRRC37A2 |
| LRRC4 |
| LRRC40 |
| LRRC41 |
| LRRC42 |
| LRRC45 |
| LRRC46 |
| LRRC47 |

| |
|---|
| LRRC49 |
| LRRC4B |
| LRRC4C |
| LRRC51 |
| LRRC57 |
| LRRC58 |
| LRRC59 |
| LRRC61 |
| LRRC66 |
| LRRC70 |
| LRRC74B |
| LRRC75A |
| LRRC8A |
| LRRC8D |
| LRRC9 |
| LRRCC1 |
| LRRFIP1 |
| LRRFIP2 |
| LRRK1 |
| LRRK2 |
| LRRN1 |
| LRRN2 |
| LRRN3 |
| LRRTM2 |
| LRSAM1 |
| LRTM2 |
| LRWD1 |
| LSG1 |
| LSM1 |
| LSM10 |
| LSM11 |
| LSM12 |
| LSM14A |
| LSM14B |
| LSM3 |
| LSM4 |
| LSM5 |
| LSM6 |
| LSM7 |
| LSM8 |
| LSMEM2 |
| LSR |
| LSS |

| |
|---|
| LTA4H |
| LTB |
| LTB4R |
| LTBP1 |
| LTBP3 |
| LTBP4 |
| LTBR |
| LTN1 |
| LTO1 |
| LTOR4 |
| LTV1 |
| LUC7L |
| LUC7L2 |
| LUC7L3 |
| LURAP1 |
| LURAP1L |
| LUZP1 |
| LUZP2 |
| LXN |
| LY6G5B |
| LY75 |
| LY9 |
| LYAR |
| LYN |
| LYNX1 |
| LYPLA1 |
| LYPLA2 |
| LYRM1 |
| LYRM2 |
| LYRM4 |
| LYRM7 |
| LYRM9 |
| LYSMD1 |
| LYSMD2 |
| LYSMD3 |
| LYST |
| LYZ |
| LZIC |
| LZTFL1 |
| LZTR1 |
| LZTS1 |
| LZTS2 |
| LZTS3 |

| |
|---|
| M6PR |
| MACIR |
| MACO1 |
| MACROD1 |
| MACROH2A1 |
| MAD1L1 |
| MAD2L1 |
| MAD2L1BP |
| MAD2L2 |
| MADD |
| MAEA |
| MAF |
| MAF1 |
| MAFB |
| MAFK |
| MAGEA10 |
| MAGEA13P |
| MAGED1 |
| MAGED2 |
| MAGED4B |
| MAGEE2 |
| MAGEL2 |
| MAGI1 |
| MAGI3 |
| MAGIX |
| MAGOH |
| MAGOHB |
| MAGT1 |
| MAIP1 |
| MAK16 |
| MAL2 |
| MALSU1 |
| MAMDC4 |
| MAML1 |
| MAML2 |
| MAML3 |
| MAMLD1 |
| MAMSTR |
| MAN1A1 |
| MAN1A2 |
| MAN1B1 |
| MAN1C1 |
| MAN2A1 |

| |
|---|
| MAN2A2 |
| MAN2B1 |
| MAN2C1 |
| MANBA |
| MANBAL |
| MANEA |
| MANEAL |
| MANF |
| MAOA |
| MAOB |
| MAP10 |
| MAP1B |
| MAP1LC3B |
| MAP2 |
| MAP2K1 |
| MAP2K2 |
| MAP2K3 |
| MAP2K5 |
| MAP2K6 |
| MAP2K7 |
| MAP3K1 |
| MAP3K10 |
| MAP3K11 |
| MAP3K12 |
| MAP3K14 |
| MAP3K15 |
| MAP3K2 |
| MAP3K21 |
| MAP3K3 |
| MAP3K4 |
| MAP3K5 |
| MAP3K6 |
| MAP3K7 |
| MAP3K8 |
| MAP3K9 |
| MAP4 |
| MAP4K1 |
| MAP4K2 |
| MAP4K3 |
| MAP4K4 |
| MAP4K5 |
| MAP6 |
| MAP7 |

| |
|---|
| MAP7D1 |
| MAP7D3 |
| MAP9 |
| MAPK11 |
| MAPK12 |
| MAPK13 |
| MAPK14 |
| MAPK15 |
| MAPK4 |
| MAPK6 |
| MAPK7 |
| MAPK8 |
| MAPK8IP1 |
| MAPK8IP3 |
| MAPKAP1 |
| MAPKAPK3 |
| MAPKBP1 |
| MAPRE2 |
| MARCHF1 |
| MARCHF2 |
| MARCHF5 |
| MARCHF6 |
| MARCHF7 |
| MARCHF8 |
| MARCHF9 |
| MARCKSL1 |
| MARF1 |
| MARK1 |
| MARK2 |
| MARK3 |
| MARK4 |
| MARS1 |
| MARS2 |
| MARVELD1 |
| MARVELD2 |
| MARVELD3 |
| MASP2 |
| MAST1 |
| MAST2 |
| MAST3 |
| MAST4 |
| MASTL |
| MAT2B |

| |
|---|
| MATK |
| MATN4 |
| MATR3 |
| MAU2 |
| MAVS |
| MAX |
| MAZ |
| MB21D2 |
| MBD1 |
| MBD2 |
| MBD3 |
| MBD4 |
| MBD5 |
| MBD6 |
| MBIP |
| MBLAC1 |
| MBLAC2 |
| MBNL2 |
| MBNL3 |
| MBOAT2 |
| MBP |
| MBTD1 |
| MBTPS1 |
| MBTPS2 |
| MCAM |
| MCAT |
| MCC |
| MCCC1 |
| MCCC2 |
| MCEE |
| MCF2L |
| MCF2L2 |
| MCFD2 |
| MCL1 |
| MCM10 |
| MCM2 |
| MCM3 |
| MCM3AP |
| MCM4 |
| MCM5 |
| MCM6 |
| MCM7 |
| MCM8 |

| |
|---|
| MCM9 |
| MCMBP |
| MCMDC2 |
| MCOLN1 |
| MCPH1 |
| MCRIP1 |
| MCRIP2 |
| MCRS1 |
| MCTP1 |
| MCTP2 |
| MCTS1 |
| MCU |
| MCUB |
| MDC1 |
| MDFIC |
| MDH2 |
| MDK |
| MDM1 |
| MDM2 |
| MDM4 |
| MDN1 |
| MDP1 |
| ME1 |
| ME2 |
| ME3 |
| MEA1 |
| MEAF6 |
| MEAK7 |
| MECOM |
| MECP2 |
| MECR |
| MED1 |
| MED10 |
| MED11 |
| MED12 |
| MED12L |
| MED13 |
| MED13L |
| MED14 |
| MED15 |
| MED16 |
| MED17 |
| MED18 |

| |
|---|
| MED19 |
| MED21 |
| MED22 |
| MED23 |
| MED24 |
| MED25 |
| MED26 |
| MED27 |
| MED28 |
| MED29 |
| MED30 |
| MED31 |
| MED4 |
| MED7 |
| MED9 |
| MEF2A |
| MEF2C |
| MEF2D |
| MEFV |
| MEGF8 |
| MEGF9 |
| MEI1 |
| MEI4 |
| MEIOB |
| MEIS1 |
| MEIS2 |
| MELK |
| MEN1 |
| MEP1A |
| MEP1B |
| MEPCE |
| MEPE |
| MERTK |
| MESD |
| MESP1 |
| MESP2 |
| MEST |
| METAP1D |
| METRN |
| METRNL |
| METTL1 |
| METTL13 |
| METTL14 |

| |
|---|
| METTL15 |
| METTL16 |
| METTL17 |
| METTL18 |
| METTL21A |
| METTL23 |
| METTL24 |
| METTL26 |
| METTL27 |
| METTL2A |
| METTL3 |
| METTL4 |
| METTL5 |
| METTL6 |
| METTL7A |
| METTL7B |
| METTL8 |
| METTL9 |
| MEX3A |
| MEX3B |
| MEX3C |
| MEX3D |
| MFAP1 |
| MFAP2 |
| MFAP3 |
| MFAP3L |
| MFGE8 |
| MFHAS1 |
| MFN1 |
| MFRP |
| MFSD1 |
| MFSD10 |
| MFSD11 |
| MFSD12 |
| MFSD13A |
| MFSD14A |
| MFSD14B |
| MFSD3 |
| MFSD4B |
| MFSD6 |
| MFSD6L |
| MFSD8 |
| MFSD9 |

| |
|---|
| MFT2 |
| MGA |
| MGAM |
| MGARP |
| MGAT1 |
| MGAT2 |
| MGAT4A |
| MGAT4B |
| MGAT5 |
| MGME1 |
| MGMT |
| MGRN1 |
| MGST1 |
| MGST2 |
| MGST3 |
| MIA2 |
| MIA3 |
| MIB1 |
| MIB2 |
| MICA |
| MICAL1 |
| MICAL3 |
| MICALL1 |
| MICALL2 |
| MICB |
| MICOS10 |
| MICOS13 |
| MICU1 |
| MICU2 |
| MICU3 |
| MID1 |
| MID1IP1 |
| MID1IPI |
| MID2 |
| MIDEAS |
| MIDN |
| MIEF1 |
| MIEF2 |
| MIEN1 |
| MIER1 |
| MIER2 |
| MIER3 |
| MIF |

| |
|---|
| MIF4GD |
| MIGA1 |
| MIGA2 |
| MIIP |
| MINAR1 |
| MINDY1 |
| MINDY2 |
| MINDY3 |
| MINDY4 |
| MINK1 |
| MINPP1 |
| MIOS |
| MIPEP |
| MIPOL1 |
| MIS12 |
| MIS18A |
| MIS18BP1 |
| MITD1 |
| MITF |
| MIX23 |
| MK167 |
| MKI67 |
| MKKS |
| MKLN1 |
| MKNK1 |
| MKNK2 |
| MKRN2 |
| MKS1 |
| MKX |
| MLEC |
| MLF1 |
| MLH1 |
| MLH3 |
| MLIP |
| MLKL |
| MLLT1 |
| MLLT3 |
| MLLT6 |
| MLPH |
| MLST8 |
| MLXIP |
| MLYCD |
| MMAA |

| |
|---|
| MMAB |
| MMACHC |
| MMADHC |
| MMD |
| MMGT1 |
| MMP14 |
| MMP15 |
| MMP17 |
| MMP19 |
| MMP2 |
| MMP25 |
| MMP9 |
| MMS19 |
| MMS22L |
| MMUT |
| MN1 |
| MNAT1 |
| MND1 |
| MNDA |
| MNS1 |
| MNT |
| MOAP1 |
| MOB1A |
| MOB1B |
| MOB3A |
| MOB3C |
| MOB4 |
| MOCOS |
| MOCS1 |
| MOCS2 |
| MOCS3 |
| MOGS |
| MON1A |
| MON1B |
| MORC2 |
| MORC3 |
| MORF4L1 |
| MORF4L2 |
| MORN4 |
| MOSMO |
| MOSPD1 |
| MOSPD3 |
| MOV10 |

| |
|---|
| MPC1 |
| MPDU1 |
| MPDZ |
| MPEG1 |
| MPG |
| MPHOSPH10 |
| MPHOSPH6 |
| MPHOSPH8 |
| MPHOSPH9 |
| MPI |
| MPL |
| MPLKIP |
| MPND |
| MPP1 |
| MPP2 |
| MPP7 |
| MPPE1 |
| MPPED2 |
| MPRIP |
| MPST |
| MPV17L2 |
| MPZL3 |
| MR1 |
| MRAP |
| MRAP2 |
| MRAS |
| MRC1 |
| MRC2 |
| MRFAP1 |
| MRFAP1L1 |
| MRGBP |
| MRGPRX3 |
| MRGRF |
| MRI1 |
| MRM1 |
| MRM2 |
| MRM3 |
| MRNIP |
| MRO |
| MROH1 |
| MRPL1 |
| MRPL10 |
| MRPL11 |

| |
|---|
| MRPL12 |
| MRPL13 |
| MRPL14 |
| MRPL15 |
| MRPL16 |
| MRPL17 |
| MRPL18 |
| MRPL19 |
| MRPL20 |
| MRPL21 |
| MRPL22 |
| MRPL23 |
| MRPL24 |
| MRPL27 |
| MRPL28 |
| MRPL3 |
| MRPL30 |
| MRPL32 |
| MRPL33 |
| MRPL34 |
| MRPL35 |
| MRPL36 |
| MRPL37 |
| MRPL38 |
| MRPL39 |
| MRPL4 |
| MRPL40 |
| MRPL41 |
| MRPL42 |
| MRPL43 |
| MRPL44 |
| MRPL45 |
| MRPL46 |
| MRPL47 |
| MRPL48 |
| MRPL49 |
| MRPL51 |
| MRPL52 |
| MRPL53 |
| MRPL54 |
| MRPL55 |
| MRPL57 |
| MRPL58 |

| |
|---|
| MRPL9 |
| MRPS10 |
| MRPS11 |
| MRPS15 |
| MRPS16 |
| MRPS17 |
| MRPS18B |
| MRPS18C |
| MRPS2 |
| MRPS21 |
| MRPS22 |
| MRPS23 |
| MRPS24 |
| MRPS25 |
| MRPS26 |
| MRPS27 |
| MRPS28 |
| MRPS30 |
| MRPS31 |
| MRPS33 |
| MRPS34 |
| MRPS35 |
| MRPS5 |
| MRPS6 |
| MRPS7 |
| MRPS9 |
| MRRF |
| MRS2 |
| MRTFA |
| MRTFB |
| MS4A12 |
| MS4A3 |
| MS4A4A |
| MS4A8 |
| MSANTD2 |
| MSANTD4 |
| MSH2 |
| MSH3 |
| MSH4 |
| MSH5 |
| MSH6 |
| MSI1 |
| MSI2 |

| |
|---|
| MSL1 |
| MSL2 |
| MSL3 |
| MSMO1 |
| MSN |
| MSR1 |
| MSRA |
| MSRB2 |
| MSRB3 |
| MST1 |
| MST1R |
| MSTO1 |
| MT-ND4L |
| MT1A |
| MT2A |
| MT3 |
| MTA1 |
| MTA2 |
| MTA3 |
| MTAP |
| MTARC1 |
| MTARC2 |
| MTBP |
| MTCH1 |
| MTCH2 |
| MTCL1 |
| MTCP1 |
| MTDH |
| MTERF1 |
| MTERF2 |
| MTERF3 |
| MTF1 |
| MTF2 |
| MTFMT |
| MTFP1 |
| MTFR1 |
| MTFR1L |
| MTFR2 |
| MTG1 |
| MTG2 |
| MTHFD1L |
| MTHFD2 |
| MTHFD2L |

| |
|---|
| MTHFSD |
| MTIF2 |
| MTIF3 |
| MTLN |
| MTM1 |
| MTMR1 |
| MTMR10 |
| MTMR11 |
| MTMR12 |
| MTMR14 |
| MTMR2 |
| MTMR3 |
| MTMR4 |
| MTMR6 |
| MTMR7 |
| MTMR9 |
| MTO1 |
| MTOR |
| MTPAP |
| MTPN |
| MTR |
| MTRES1 |
| MTREX |
| MTRF1 |
| MTRF1L |
| MTRR |
| MTSS1 |
| MTSS2 |
| MTTP |
| MTURN |
| MTUS1 |
| MTUS2 |
| MTX1 |
| MTX2 |
| MTX3 |
| MUC1 |
| MUC13 |
| MUC16 |
| MUC17 |
| MUC19 |
| MUC2 |
| MUC20 |
| MUC5B |

| |
|---|
| MUL1 |
| MUS81 |
| MVB12A |
| MVB12B |
| MVD |
| MVK |
| MVP |
| MX1 |
| MX2 |
| MXD3 |
| MXD4 |
| MXI1 |
| MXRA5 |
| MXRA7 |
| MXRA8 |
| MYADM |
| MYBBP1A |
| MYBL1 |
| MYBL2 |
| MYBPC1 |
| MYC |
| MYCBP |
| MYCBP2 |
| MYCBPAP |
| MYCN |
| MYD88 |
| MYDGF |
| MYEF2 |
| MYG1 |
| MYH10 |
| MYH11 |
| MYH14 |
| MYH7 |
| MYH9 |
| MYL12B |
| MYL2 |
| MYL6 |
| MYL6B |
| MYL9 |
| MYLIP |
| MYLK |
| MYLPF |
| MYO10 |

| |
|---|
| MYO15B |
| MYO18A |
| MYO19 |
| MYO1A |
| MYO1B |
| MYO1C |
| MYO1D |
| MYO1E |
| MYO1F |
| MYO1G |
| MYO3B |
| MYO5A |
| MYO5B |
| MYO5C |
| MYO6 |
| MYO7A |
| MYO7B |
| MYO9A |
| MYO9B |
| MYOC |
| MYOCD |
| MYOF |
| MYOM1 |
| MYORG |
| MYOZ1 |
| MYPN |
| MYPOP |
| MYRF |
| MYSM1 |
| MYZAP |
| MZF1 |
| MZT1 |
| N/A |
| N4BP1 |
| N4BP2 |
| N4BP2L1 |
| N4BP3 |
| N6AMT1 |
| NAA10 |
| NAA15 |
| NAA16 |
| NAA20 |
| NAA25 |

| |
|---|
| NAA30 |
| NAA35 |
| NAA38 |
| NAA40 |
| NAA60 |
| NAAA |
| NAALADL1 |
| NAALADL2 |
| NAB1 |
| NAB2 |
| NABP1 |
| NACA |
| NACAD |
| NACC1 |
| NACC2 |
| NADK |
| NADK2 |
| NAE1 |
| NAF1 |
| NAGA |
| NAGK |
| NAGLU |
| NAGPA |
| NAIF1 |
| NAIP |
| NALF1 |
| NALF2 |
| NAMPT |
| NANOS1 |
| NANP |
| NANS |
| NAP1L1 |
| NAP1L3 |
| NAP1L5 |
| NAPA |
| NAPB |
| NAPEPLD |
| NAPG |
| NAPRT |
| NAPSA |
| NARF |
| NARS1 |
| NASP |

| |
|---|
| NAT1 |
| NAT10 |
| NAT14 |
| NAT8 |
| NAT8B |
| NAT9 |
| NATD1 |
| NAV1 |
| NAV2 |
| NAV3 |
| NAXD |
| NAXE |
| NBAS |
| NBEA |
| NBEAL2 |
| NBL1 |
| NBN |
| NBPF4 |
| NBPF5P |
| NBR1 |
| NCAM1 |
| NCAM2 |
| NCAPD2 |
| NCAPD3 |
| NCAPG |
| NCAPG2 |
| NCAPH |
| NCAPH2 |
| NCBP1 |
| NCBP2 |
| NCBP2AS2 |
| NCDN |
| NCEH1 |
| NCF1 |
| NCF2 |
| NCF4 |
| NCK1 |
| NCK2 |
| NCKAP1 |
| NCKAP1L |
| NCKAP5 |
| NCKAP5L |
| NCKIPSD |

| |
|---|
| NCL |
| NCLN |
| NCOA1 |
| NCOA2 |
| NCOA3 |
| NCOA4 |
| NCOA5 |
| NCOA6 |
| NCOA7 |
| NCOR1 |
| NCOR2 |
| NCR3LG1 |
| ncRNA? |
| NCS1 |
| NCSTN |
| NDC1 |
| NDC80 |
| NDE1 |
| NDEL1 |
| NDFA11 |
| NDFIP1 |
| NDFIP2 |
| NDKM |
| NDOR1 |
| NDRG1 |
| NDRG2 |
| NDST2 |
| NDUFA10 |
| NDUFA12 |
| NDUFA13 |
| NDUFA2 |
| NDUFA3 |
| NDUFA4 |
| NDUFA6 |
| NDUFA8 |
| NDUFAB1 |
| NDUFAF1 |
| NDUFAF2 |
| NDUFAF3 |
| NDUFAF4 |
| NDUFAF5 |
| NDUFAF6 |
| NDUFAF7 |

| |
|---|
| NDUFAF8 |
| NDUFB1 |
| NDUFB10 |
| NDUFB11 |
| NDUFB2 |
| NDUFB4 |
| NDUFB5 |
| NDUFB6 |
| NDUFB7 |
| NDUFB8 |
| NDUFB9 |
| NDUFC1 |
| NDUFC2 |
| NDUFS2 |
| NDUFS3 |
| NDUFS5 |
| NDUFS6 |
| NDUFS7 |
| NDUFV1 |
| NDUFV2 |
| NDUFV3 |
| NEAT1 |
| NEBL |
| NECAP1 |
| NECAP2 |
| NECTIN1 |
| NECTIN2 |
| NEDD4 |
| NEDD8 |
| NEDD9 |
| NEFH |
| NEFL |
| NEFM |
| NEGR1 |
| NEIL1 |
| NEIL2 |
| NEIL3 |
| NEK1 |
| NEK10 |
| NEK2 |
| NEK3 |
| NEK4 |
| NEK6 |

| |
|---|
| NEK7 |
| NEK8 |
| NEK9 |
| NELFA |
| NELFB |
| NELFCD |
| NELFE |
| NELL1 |
| NELL2 |
| NEMF |
| NEMP1 |
| NEMP2 |
| NENF |
| NEO1 |
| NES |
| NET1 |
| NETR |
| NEU1 |
| NEU3 |
| NEUL1 |
| NEURL1B |
| NEURL2 |
| NEURL4 |
| NEXN |
| NF1 |
| NF2IP |
| NFAM1 |
| NFAT5 |
| NFATC1 |
| NFATC3 |
| NFE2 |
| NFE2L1 |
| NFE2L3 |
| NFIC |
| NFIL3 |
| NFIX |
| NFKB1 |
| NFKBIA |
| NFKBIB |
| NFKBID |
| NFKBIE |
| NFKBIL1 |
| NFKBIZ |

| |
|---|
| NFS1 |
| NFX1 |
| NFXL1 |
| NFYA |
| NFYB |
| NFYC |
| NGDN |
| NGLY1 |
| NGRN |
| NHEJ1 |
| NHLRC1 |
| NHLRC2 |
| NHLRC3 |
| NHP2 |
| NHS |
| NHSL2 |
| NIBAN1 |
| NIBAN2 |
| NICN1 |
| NID1 |
| NID2 |
| NIF3L1 |
| NIFK |
| NIN |
| NINJ1 |
| NINL |
| NIP7 |
| NIPA1 |
| NIPA2 |
| NIPAL1 |
| NIPAL3 |
| NIPBL |
| NIPSNAP2 |
| NIPSNAP3A |
| NISCH |
| NIT1 |
| NIT2 |
| NKAIN1 |
| NKAP |
| NKAPD1 |
| NKAPL |
| NKD1 |
| NKG7 |

| |
|---|
| NKIRAS1 |
| NKIRAS2 |
| NKPD1 |
| NKRF |
| NKTR |
| NKX1-2 |
| NLE1 |
| NLGN2 |
| NLGN3 |
| NLK |
| NLN |
| NLRC3 |
| NLRC4 |
| NLRP1 |
| NLRP3 |
| NLRP5 |
| NLRP6 |
| NMB |
| NMD3 |
| NME2 |
| NME3 |
| NME6 |
| NME7 |
| NMI |
| NMNAT1 |
| NMNAT3 |
| NMRAL1 |
| NMRK1 |
| NMT1 |
| NNAT |
| NNT |
| NOA1 |
| NOB1 |
| NOC2L |
| NOC3L |
| NOC4L |
| NOCT |
| NOD1 |
| NOD2 |
| NODAL |
| NOL10 |
| NOL11 |
| NOL12 |

| |
|---|
| NOL4L |
| NOL6 |
| NOL7 |
| NOL8 |
| NOL9 |
| NOM1 |
| NOMO1 |
| NONO |
| NOP14 |
| NOP16 |
| NOP2 |
| NOP53 |
| NOP56 |
| NOP58 |
| NOP9 |
| NOPCHAP1 |
| NOS1AP |
| NOS2 |
| NOSIP |
| NOTCH1 |
| NOTCH2 |
| NOTCH3 |
| NOTCH4 |
| nothing |
| NOTUM |
| NOVA1 |
| NOXA1 |
| NOXO1 |
| NOXRED1 |
| NPAS3 |
| NPAT |
| NPB |
| NPC2 |
| NPDC1 |
| NPEPL1 |
| NPEPPS |
| NPHP1 |
| NPHP3 |
| NPHP4 |
| NPL |
| NPLOC4 |
| NPM1 |
| NPM3 |

| |
|---|
| NPNT |
| NPR2 |
| NPR3 |
| NPRL2 |
| NPRL3 |
| NPTN |
| NPTX1 |
| NPTXR |
| NPY1R |
| NPY4R |
| NQO1 |
| NQO2 |
| NR1D2 |
| NR1H2 |
| NR1H4 |
| NR1I2 |
| NR2C2 |
| NR2C2AP |
| NR2E1 |
| NR2F1 |
| NR2F2 |
| NR2F6 |
| NR3C2 |
| NR4A1 |
| NR4A2 |
| NR4A3 |
| NR5A1 |
| NR5A2 |
| NR6A1 |
| NRAP |
| NRARP |
| NRBF2 |
| NRBP1 |
| NRBP2 |
| NRDC |
| NRDE2 |
| NREP |
| NRF1 |
| NRIP1 |
| NRIP3 |
| NRM |
| NRP1 |
| NRROS |

| |
|---|
| NRSN2 |
| NRXN1 |
| NSA2 |
| NSD1 |
| NSD2 |
| NSD3 |
| NSDHL |
| NSF |
| NSFL1C |
| NSL1 |
| NSMAF |
| NSMCE1 |
| NSMCE2 |
| NSMCE3 |
| NSMCE4A |
| NSMF |
| NSUN3 |
| NSUN4 |
| NSUN5 |
| NSUN6 |
| NSUN7 |
| NT5C |
| NT5C2 |
| NT5C3A |
| NT5C3B |
| NT5DC1 |
| NT5DC2 |
| NT5DC3 |
| NT5E |
| NT5M |
| NTAN1 |
| NTAQ1 |
| NTHL1 |
| NTMT1 |
| NTN3 |
| NTN4 |
| NTN5 |
| NTNG1 |
| NTRK1 |
| NUAK1 |
| NUAK2 |
| NUB1 |
| NUBP1 |

| |
|---|
| NUBP2 |
| NUBPL |
| NUCB1 |
| NUCB2 |
| NUCKS1 |
| NUDC |
| NUDCD1 |
| NUDCD2 |
| NUDCD3 |
| NUDT1 |
| NUDT12 |
| NUDT13 |
| NUDT14 |
| NUDT15 |
| NUDT16 |
| NUDT16L1 |
| NUDT17 |
| NUDT18 |
| NUDT2 |
| NUDT21 |
| NUDT22 |
| NUDT3 |
| NUDT4B |
| NUDT5 |
| NUDT6 |
| NUDT7 |
| NUDT8 |
| NUDT9 |
| NUF2 |
| NUFIP1 |
| NUFIP2 |
| NUGGC |
| NUMA1 |
| NUMB |
| NUMBL |
| NUP133 |
| NUP153 |
| NUP155 |
| NUP160 |
| NUP188 |
| NUP205 |
| NUP210 |
| NUP214 |

| |
|---|
| NUP35 |
| NUP42 |
| NUP43 |
| NUP50 |
| NUP54 |
| NUP58 |
| NUP62 |
| NUP85 |
| NUP88 |
| NUP93 |
| NUP98 |
| NUPR1 |
| NUPR2 |
| NUS1 |
| NUSAP1 |
| NVL |
| NXF1 |
| NXPE3 |
| NXPE4 |
| NXPH1 |
| NXT1 |
| NXT2 |
| NYAP1 |
| NYNRIN |
| OAF |
| OARD1 |
| OAS3 |
| OASL |
| OASL2 |
| OAT |
| OAZ1 |
| OAZ2 |
| OBI1 |
| OBSCN |
| OBSL1 |
| OCEL1 |
| OCIAD1 |
| OCIAD2 |
| OCLN |
| OCRL |
| ODAD3 |
| ODAD4 |
| ODC1 |

| |
|---|
| ODF2 |
| ODF2L |
| ODR4 |
| OE5AU1 |
| OFD1 |
| OGA |
| OGFOD1 |
| OGFOD3 |
| OGFR |
| OGFRL1 |
| OGG1 |
| OGT1 |
| OIP5 |
| OIT3 |
| OLA1 |
| OLFM1 |
| OLFM4 |
| OLFML1 |
| OLFML2A |
| OLFML2B |
| OLIG1 |
| OLR1 |
| OMA1 |
| OMD |
| ONECUT1 |
| OPA1 |
| OPA3 |
| OPCML |
| OPHN1 |
| OPRK1 |
| OPTN |
| OR10V1 |
| OR12D3 |
| OR13C2 |
| OR13C6P |
| OR13C7 |
| OR13C8 |
| OR1J4 |
| OR1N1 |
| OR2A1 |
| OR2A7 |
| OR2AG1 |
| OR2C3 |

| |
|---|
| OR2F1 |
| OR2G6 |
| OR2S2 |
| OR2T33 |
| OR4D9 |
| OR4K2 |
| OR4S2 |
| OR51B5 |
| OR51F2 |
| OR51Q1 |
| OR52A1 |
| OR52B4 |
| OR52E5 |
| OR52I1 |
| OR52M1 |
| OR52N4 |
| OR56A1 |
| OR5M10 |
| OR5M11 |
| OR6C68 |
| OR7D2 |
| OR7E24 |
| OR8B12 |
| OR8B3 |
| OR8K3 |
| OR8S1 |
| ORAI1 |
| ORAI3 |
| ORC1 |
| ORC2 |
| ORC3 |
| ORC4 |
| ORC6 |
| ORMDL2 |
| ORMDL3 |
| OS9 |
| OSBP |
| OSBP2 |
| OSBPL10 |
| OSBPL11 |
| OSBPL1A |
| OSBPL2 |
| OSBPL3 |

| |
|---|
| OSBPL5 |
| OSBPL8 |
| OSBPL9 |
| OSCAR |
| OSCP1 |
| OSER1 |
| OSGEP |
| OSGEPL1 |
| OSGIN1 |
| OSM |
| OSR2 |
| OSTC |
| OSTF1 |
| OSTM1 |
| OTOP2 |
| OTUB1 |
| OTUD1 |
| OTUD3 |
| OTUD4 |
| OTUD5 |
| OTUD6B |
| OTULIN |
| OTULINL |
| OVAAL |
| OVCA2 |
| OVGP1 |
| OVOL1 |
| OXA1L |
| OXCT1 |
| OXER1 |
| OXNAD1 |
| OXR1 |
| OXSM |
| OXSR1 |
| P2RX4 |
| P2RX6 |
| P2RY1 |
| P2RY10 |
| P2RY2 |
| P2RY4 |
| P3H1 |
| P3H2 |
| P3H3 |

| |
|---|
| P3H4 |
| P4HA1 |
| P4HA2 |
| P4HA3 |
| P4HB |
| P4HTM |
| PA2G4 |
| PAAF1 |
| PABIR1 |
| PABIR2 |
| PABPC1 |
| PABPC4 |
| PABPC4L |
| PABPC5 |
| PABPN1 |
| PACC1 |
| PACRGL |
| PACS2 |
| PACSIN2 |
| PACSIN3 |
| PADI1 |
| PADI2 |
| PAF1 |
| PAFAH1B1 |
| PAFAH1B2 |
| PAFAH1B3 |
| PAFAH2 |
| PAG1 |
| PAIP1 |
| PAIP2 |
| PAK1 |
| PAK1IP1 |
| PAK2 |
| PAK3 |
| PAK4 |
| PALB2 |
| PALD1 |
| PALLD |
| PALMD |
| PALS1 |
| PALS2 |
| PAM |
| PAN2 |

| |
|---|
| PAN3 |
| PANK2 |
| PANK3 |
| PANK4 |
| PANX1 |
| PANX2 |
| PAOX |
| PAPLN |
| PAPOLA |
| PAPOLB |
| PAPOLG |
| PAPSS1 |
| PAPSS2 |
| PAQR3 |
| PAQR4 |
| PAQR5 |
| PAQR6 |
| PAQR8 |
| PARD3 |
| PARD6A |
| PARD6B |
| PARD6G |
| PARG |
| PARK7 |
| PARL |
| PARN |
| PARP1 |
| PARP11 |
| PARP12 |
| PARP14 |
| PARP16 |
| PARP2 |
| PARP6 |
| PARP8 |
| PARP9 |
| PARPBP |
| PARS2 |
| PARVA |
| PARVG |
| PASK |
| PATJ |
| PATL1 |
| PATZ1 |

| |
|---|
| PAWR |
| PAXBP1 |
| PAXIP1 |
| PAXX |
| PBDC1 |
| PBK |
| PBLD |
| PBRM1 |
| PBX1 |
| PBX2 |
| PBX3 |
| PBX4 |
| PBXIP1 |
| PC |
| PCAT1 |
| PCAT2 |
| PCBD1 |
| PCBD2 |
| PCBP1 |
| PCBP2 |
| PCBP4 |
| PCCA |
| PCCB |
| PCDH1 |
| PCDH10 |
| PCDH18 |
| PCDH7 |
| PCDH9 |
| PCDHA1 |
| PCDHA3 |
| PCDHA4 |
| PCDHAC2 |
| PCDHB11 |
| PCDHB14 |
| PCDHB15 |
| PCDHB18 |
| PCDHB4 |
| PCDHB6 |
| PCDHB7 |
| PCDHGA11 |
| PCDHGA12 |
| PCDHGA3 |
| PCDHGA4 |

| |
|---|
| PCDHGA5 |
| PCDHGA6 |
| PCDHGA7 |
| PCDHGA8 |
| PCDHGA9 |
| PCDHGB1 |
| PCDHGB2 |
| PCDHGB6 |
| PCDHGC5 |
| PCED1A |
| PCED1B |
| PCF11 |
| PCGF1 |
| PCGF2 |
| PCGF3 |
| PCGF5 |
| PCGF6 |
| PCID2 |
| PCIF1 |
| PCK1 |
| PCK2 |
| PCLAF |
| PCLO |
| PCM1 |
| PCMT1 |
| PCMTD1 |
| PCMTD2 |
| PCNA |
| PCNP |
| PCNT |
| PCNX1 |
| PCNX3 |
| PCNX4 |
| PCOLCE |
| PCOLCE2 |
| PCP4L1 |
| PCSK1N |
| PCSK4 |
| PCSK5 |
| PCSK6 |
| PCSK7 |
| PCTP |
| PCYOX1 |

| |
|---|
| PCYOX1L |
| PCYT1A |
| PCYT1B |
| PCYT2 |
| PDAP1 |
| PDCD1 |
| PDCD10 |
| PDCD11 |
| PDCD1LG2 |
| PDCD2 |
| PDCD2L |
| PDCD4 |
| PDCD5 |
| PDCD6 |
| PDCD6IP |
| PDCD7 |
| PDCL |
| PDCL3 |
| PDE10A |
| PDE11A |
| PDE12 |
| PDE2A |
| PDE4B |
| PDE4C |
| PDE4D |
| PDE4DIP |
| PDE5A |
| PDE6D |
| PDE7A |
| PDE7B |
| PDE8A |
| PDE8B |
| PDE9A |
| PDF |
| PDGFA |
| PDGFB |
| PDGFC |
| PDGFD |
| PDGFRB |
| PDHA1 |
| PDHB |
| PDHX |
| PDIA2 |

| |
|---|
| PDIA3 |
| PDIA4 |
| PDIA5 |
| PDIA6 |
| PDIK1L |
| PDK1 |
| PDK3 |
| PDK4 |
| PDLIM1 |
| PDLIM2 |
| PDLIM4 |
| PDLIM5 |
| PDLIM7 |
| PDP2 |
| PDPK1 |
| PDRG1 |
| PDS5A |
| PDS5B |
| PDSS1 |
| PDXDC1 |
| PDXK |
| PDXP |
| PDZD11 |
| PDZD2 |
| PDZD4 |
| PDZD8 |
| PDZK1 |
| PDZRN3 |
| PDZRN4 |
| PEAK1 |
| PEBP1 |
| PECAM1 |
| PECR |
| PEDS1 |
| PEG10 |
| PEG3 |
| PELI2 |
| PELI3 |
| PELO |
| PELP1 |
| PEMT |
| PENK |
| PER1 |

| |
|---|
| PER2 |
| PER3 |
| PERP |
| PET100 |
| PET117 |
| PEX1 |
| PEX10 |
| PEX11A |
| PEX11B |
| PEX11G |
| PEX12 |
| PEX13 |
| PEX14 |
| PEX16 |
| PEX19 |
| PEX2 |
| PEX3 |
| PEX5 |
| PEX6 |
| PEX7 |
| PF4 |
| PFAS |
| PFDN1 |
| PFDN2 |
| PFDN4 |
| PFDN5 |
| PFDN6 |
| PFKFB2 |
| PFKFB3 |
| PFKFB4 |
| PFKL |
| PFKM |
| PFKP |
| PFN1 |
| PFN2 |
| PGAM1 |
| PGAM2 |
| PGAM5 |
| PGAP1 |
| PGAP2 |
| PGAP3 |
| PGAP4 |
| PGAP6 |

| |
|---|
| PGBD1 |
| PGBD2 |
| PGBD5 |
| PGBP |
| PGF |
| PGGT1B |
| PGK1 |
| PGLS |
| PGM1 |
| PGM2 |
| PGM2L1 |
| PGM3 |
| PGM5 |
| PGP |
| PGPEP1 |
| PGR |
| PGRMC1 |
| PGRMC2 |
| PGS1 |
| PHACTR1 |
| PHACTR2 |
| PHACTR4 |
| PHAF1 |
| PHAX |
| PHB1 |
| PHB2 |
| PHC1 |
| PHC2 |
| PHC3 |
| PHETA1 |
| PHETA2 |
| PHF1 |
| PHF11 |
| PHF12 |
| PHF13 |
| PHF14 |
| PHF19 |
| PHF2 |
| PHF20 |
| PHF21A |
| PHF23 |
| PHF24 |
| PHF3 |

| |
|---|
| PHF5A |
| PHF6 |
| PHF8 |
| PHGDH |
| PHGR1 |
| PHIP |
| PHKA1 |
| PHKA2 |
| PHKB |
| PHKG2 |
| PHLDA1 |
| PHLDA2 |
| PHLDA3 |
| PHLDB1 |
| PHLDB2 |
| PHLDB3 |
| PHLPP1 |
| PHLPP2 |
| PHOSPHO2 |
| PHRF1 |
| PHYH |
| PHYKPL |
| PI16 |
| PI4K2A |
| PI4K2B |
| PI4KA |
| PI4KB |
| PIAS1 |
| PIAS2 |
| PIAS3 |
| PIAS4 |
| PICALM |
| PICK1 |
| PIDD1 |
| PIEZO1 |
| PIF1 |
| PIGA |
| PIGBOS1 |
| PIGC |
| PIGF |
| PIGG |
| PIGH |
| PIGK |

| |
|---|
| PIGL |
| PIGM |
| PIGN |
| PIGO |
| PIGP |
| PIGR |
| PIGS |
| PIGT |
| PIGU |
| PIGV |
| PIGY |
| PIGZ |
| PIH1D1 |
| PIH1D2 |
| PIK3C2A |
| PIK3C2B |
| PIK3C3 |
| PIK3CA |
| PIK3CB |
| PIK3CD |
| PIK3CG |
| PIK3IP1 |
| PIK3R1 |
| PIK3R2 |
| PIK3R3 |
| PIK3R4 |
| PIK3R5 |
| PIK3R6 |
| PIKFYVE |
| PIM1 |
| PIM2 |
| PIM3 |
| PIMREG |
| PIN1 |
| PIN4 |
| PINK1 |
| PINX1 |
| PIP4K2A |
| PIP4K2B |
| PIP4P1 |
| PIP5K1A |
| PIP5K1B |
| PIP5K1C |

| |
|---|
| PIR |
| PIRT |
| PISD |
| PITHD1 |
| PITPNA |
| PITPNM1 |
| PITPNM2 |
| PITRM1 |
| PITX3 |
| PIWIL2 |
| PJA1 |
| PJVK |
| PKD1 |
| PKD2 |
| PKD2L2 |
| PKDCC |
| PKHD1 |
| PKIA |
| PKIG |
| PKMYT1 |
| PKN1 |
| PKN2 |
| PKN3 |
| PKNOX1 |
| PKP1 |
| PKP2 |
| PKP3 |
| PKP4 |
| PLA2G12A |
| PLA2G15 |
| PLA2G6 |
| PLA2G7 |
| PLAA |
| PLAAT2 |
| PLAAT3 |
| PLAC8 |
| PLAGL1 |
| PLAGL2 |
| PLAU |
| PLAUR |
| PLB1 |
| PLBD1 |
| PLBD2 |

| |
|---|
| PLCB2 |
| PLCB3 |
| PLCB4 |
| PLCD1 |
| PLCD3 |
| PLCE1 |
| PLCG1 |
| PLCG2 |
| PLCH2 |
| PLCL2 |
| PLCXD1 |
| PLCXD2 |
| PLD2 |
| PLD3 |
| PLEC |
| PLEK |
| PLEK2 |
| PLEKHA1 |
| PLEKHA2 |
| PLEKHA3 |
| PLEKHA4 |
| PLEKHA5 |
| PLEKHA6 |
| PLEKHA7 |
| PLEKHA8 |
| PLEKHB1 |
| PLEKHD1 |
| PLEKHF1 |
| PLEKHF2 |
| PLEKHG1 |
| PLEKHG2 |
| PLEKHG3 |
| PLEKHG4B |
| PLEKHG5 |
| PLEKHG6 |
| PLEKHG7 |
| PLEKHH1 |
| PLEKHH3 |
| PLEKHJ1 |
| PLEKHM1 |
| PLEKHM2 |
| PLEKHM3 |
| PLEKHN1 |

| |
|---|
| PLEKHO1 |
| PLEKHO2 |
| PLGRKT |
| PLIN1 |
| PLIN2 |
| PLIN3 |
| PLK1 |
| PLK2 |
| PLK3 |
| PLK4 |
| PLOD1 |
| PLOD2 |
| PLOD3 |
| PLP2 |
| PLPBP |
| PLPP1 |
| PLPP2 |
| PLPP3 |
| PLPP5 |
| PLPP6 |
| PLPP7 |
| PLPPR2 |
| PLPPR4 |
| PLRG1 |
| PLS3 |
| PLSCR4 |
| PLSCR5 |
| PLTP |
| PLVAP |
| PLXDC1 |
| PLXNA4 |
| PLXNB1 |
| PLXNB2 |
| PLXNB3 |
| PLXNC1 |
| PLXND1 |
| PM20D2 |
| PMEL |
| PMF1 |
| PML |
| PMM1 |
| PMM2 |
| PMP22 |

| |
|---|
| PMPCA |
| PMPCB |
| PMS1 |
| PMVK |
| PNISR |
| PNKD |
| PNKP |
| PNMA1 |
| PNMA6E |
| PNMA6F |
| PNMA8A |
| PNN |
| PNO1 |
| PNP |
| PNPLA2 |
| PNPLA3 |
| PNPLA4 |
| PNPLA6 |
| PNPLA7 |
| PNPLA8 |
| PNPO |
| PNPT1 |
| PNRC1 |
| PNRC2 |
| POC1A |
| POC1B |
| POC5 |
| POF1B |
| POFUT1 |
| POFUT2 |
| POGK |
| POGLUT1 |
| POGLUT2 |
| POGLUT3 |
| POGZ |
| POLA1 |
| POLA2 |
| POLB |
| POLD1 |
| POLD2 |
| POLD3 |
| POLD4 |
| POLDIP2 |

| |
|---|
| POLDIP3 |
| POLE |
| POLE2 |
| POLE3 |
| POLE4 |
| POLG |
| POLG2 |
| POLH |
| POLI |
| POLK |
| POLL |
| POLM |
| POLQ |
| POLR1C |
| POLR1D |
| POLR1E |
| POLR1F |
| POLR1G |
| POLR1H |
| POLR2A |
| POLR2B |
| POLR2C |
| POLR2D |
| POLR2E |
| POLR2F |
| POLR2G |
| POLR2H |
| POLR2I |
| POLR2J |
| POLR2L |
| POLR2M |
| POLR3A |
| POLR3B |
| POLR3C |
| POLR3E |
| POLR3F |
| POLR3GL |
| POLR3H |
| POLR3K |
| POLRMT |
| POM121 |
| POM121C |
| POM121L2 |

| |
|---|
| POMC |
| POMGNT1 |
| POMGNT2 |
| POMK |
| POMP |
| POMT1 |
| POMT2 |
| PON2 |
| PON3 |
| POP1 |
| POP5 |
| POP7 |
| POR |
| PORCN |
| POSTN |
| POT1 |
| POU2AF1 |
| POU2F1 |
| POU2F3 |
| POU5F1 |
| POU6F1 |
| PP4C |
| PPA1 |
| PPA2 |
| PPAN |
| PPARA |
| PPARD |
| PPARG |
| PPARGC1B |
| PPAT |
| PPCDC |
| PPCS |
| PPDPF |
| PPFIA1 |
| PPFIA4 |
| PPFIBP1 |
| PPFIBP2 |
| PPHLN1 |
| PPIA |
| PPIB |
| PPIC |
| PPID |
| PPIE |

| |
|---|
| PPIF |
| PPIG |
| PPIH |
| PPIL1 |
| PPIL2 |
| PPIL3 |
| PPIL4 |
| PPIL6 |
| PPIP5K1 |
| PPIP5K2 |
| PPL |
| PPM1A |
| PPM1B |
| PPM1D |
| PPM1F |
| PPM1G |
| PPM1J |
| PPM1K |
| PPM1M |
| PPOX |
| PPP1CA |
| PPP1CC |
| PPP1R10 |
| PPP1R11 |
| PPP1R12A |
| PPP1R12B |
| PPP1R12C |
| PPP1R13B |
| PPP1R13L |
| PPP1R14A |
| PPP1R14B |
| PPP1R14C |
| PPP1R14D |
| PPP1R15A |
| PPP1R15B |
| PPP1R18 |
| PPP1R2 |
| PPP1R21 |
| PPP1R26 |
| PPP1R32 |
| PPP1R35 |
| PPP1R37 |
| PPP1R3A |

| |
|---|
| PPP1R3B |
| PPP1R3D |
| PPP1R3E |
| PPP1R3F |
| PPP1R3G |
| PPP1R7 |
| PPP1R8 |
| PPP1R9A |
| PPP1R9B |
| PPP2CA |
| PPP2CB |
| PPP2R1A |
| PPP2R1B |
| PPP2R2A |
| PPP2R2D |
| PPP2R3A |
| PPP2R3B |
| PPP2R3C |
| PPP2R5A |
| PPP2R5B |
| PPP2R5C |
| PPP2R5D |
| PPP2R5E |
| PPP3CA |
| PPP3CB |
| PPP3R1 |
| PPP3R2 |
| PPP4C |
| PPP4R1 |
| PPP4R2 |
| PPP4R3A |
| PPP4R3B |
| PPP4R4 |
| PPP5C |
| PPP6C |
| PPP6R2 |
| PPP6R3 |
| PPR1A |
| PPRC1 |
| PPT1 |
| PPT2 |
| PPTC7 |
| PPWD1 |

| |
|---|
| PQBP1 |
| PQLC2 |
| PRADC1 |
| PRAF2 |
| PRAM1 |
| PRAME |
| PRC1 |
| PRCC |
| PRCP |
| PRD1B |
| PRDM1 |
| PRDM10 |
| PRDM11 |
| PRDM15 |
| PRDM2 |
| PRDM4 |
| PRDM5 |
| PRDM7 |
| PRDX2 |
| PRDX3 |
| PRDX4 |
| PRDX5 |
| PRDX6 |
| PREB |
| PRELID1 |
| PRELID3B |
| PREP |
| PREPL |
| PREX1 |
| PRICKLE1 |
| PRICKLE3 |
| PRICKLE4 |
| PRIM1 |
| PRIM2 |
| PRIMPOL |
| PRKAA1 |
| PRKAA2 |
| PRKAB1 |
| PRKACA |
| PRKACB |
| PRKAG1 |
| PRKAG2 |
| PRKAR1B |

| |
|---|
| PRKAR2A |
| PRKCD |
| PRKCE |
| PRKCI |
| PRKCSH |
| PRKCZ |
| PRKD1 |
| PRKD2 |
| PRKD3 |
| PRKDC |
| PRKRA |
| PRKRIP1 |
| PRMT1 |
| PRMT2 |
| PRMT3 |
| PRMT5 |
| PRMT6 |
| PRMT7 |
| PRMT9 |
| PROA |
| PROB1 |
| PRODH |
| PROM1 |
| PRORP |
| PRORSD1 |
| PROS1 |
| PROSER1 |
| PROSER2 |
| PROSER3 |
| PRPF18 |
| PRPF19 |
| PRPF3 |
| PRPF31 |
| PRPF38A |
| PRPF38B |
| PRPF39 |
| PRPF40A |
| PRPF40B |
| PRPF4B |
| PRPF6 |
| PRPF8 |
| PRPS1 |
| PRPS2 |

| |
|---|
| PRPSAP1 |
| PRPSAP2 |
| PRR11 |
| PRR12 |
| PRR14 |
| PRR14L |
| PRR15 |
| PRR15L |
| PRR23A |
| PRR5 |
| PRR7 |
| PRRC1 |
| PRRC2A |
| PRRC2B |
| PRRC2C |
| PRRG2 |
| PRRT1B |
| PRSS35 |
| PRSS36 |
| PRSS47P |
| PRUNE2 |
| PRX |
| PRXL2A |
| PRXL2B |
| PRXL2C |
| PSAP |
| PSAT1 |
| PSD3 |
| PSD4 |
| PSEN1 |
| PSEN2 |
| PSENEN |
| PSIP1 |
| PSKH1 |
| PSMA1 |
| PSMA2 |
| PSMA3 |
| PSMA5 |
| PSMA6 |
| PSMA7 |
| PSMB1 |
| PSMB10 |
| PSMB11 |

| |
|---|
| PSMB2 |
| PSMB3 |
| PSMB4 |
| PSMB5 |
| PSMB7 |
| PSMB9 |
| PSMC1 |
| PSMC2 |
| PSMC3 |
| PSMC3IP |
| PSMC5 |
| PSMC6 |
| PSMD1 |
| PSMD10 |
| PSMD11 |
| PSMD12 |
| PSMD13 |
| PSMD14 |
| PSMD2 |
| PSMD3 |
| PSMD4 |
| PSMD5 |
| PSMD6 |
| PSMD7 |
| PSMD8 |
| PSMD9 |
| PSME1 |
| PSME2 |
| PSME3 |
| PSME4 |
| PSMF1 |
| PSMG2 |
| PSMG3 |
| PSMG4 |
| PSPC1 |
| PSPH |
| PSRC1 |
| PSTK |
| PSTPIP1 |
| PTAFR |
| PTAR1 |
| PTBP1 |
| PTBP3 |

| |
|---|
| PTCD1 |
| PTCD2 |
| PTCD3 |
| PTCH1 |
| PTCH2 |
| PTCHD1 |
| PTCHD4 |
| PTCRA |
| PTDSS1 |
| PTDSS2 |
| PTEN |
| PTER |
| PTGDS |
| PTGER1 |
| PTGER2 |
| PTGER3 |
| PTGER4 |
| PTGES2 |
| PTGES3 |
| PTGES3L |
| PTGFR |
| PTGFRN |
| PTGS1 |
| PTGS2 |
| PTH |
| PTH2 |
| PTHLH |
| PTK2 |
| PTK2B |
| PTK6 |
| PTK7 |
| PTMA |
| PTN |
| PTN4 |
| PTOV1 |
| PTP4A1 |
| PTP4A2 |
| PTP4A3 |
| PTPA |
| PTPDC1 |
| PTPMT1 |
| PTPN1 |
| PTPN11 |

| |
|---|
| PTPN12 |
| PTPN18 |
| PTPN2 |
| PTPN21 |
| PTPN23 |
| PTPN3 |
| PTPN6 |
| PTPN9 |
| PTPRA |
| PTPRB |
| PTPRC |
| PTPRD |
| PTPRE |
| PTPRF |
| PTPRG |
| PTPRH |
| PTPRJ |
| PTPRK |
| PTPRM |
| PTPRN2 |
| PTPRQ |
| PTPRS |
| PTRHD1 |
| PTS |
| PTTG1 |
| PTX3 |
| PTX4 |
| PUDP |
| PUF60 |
| PUM1 |
| PUM2 |
| PUM3 |
| PUR2 |
| PURA |
| PURB |
| PURG |
| PUS1 |
| PUS3 |
| PUS7 |
| PUS7L |
| PUSL1 |
| PVR |
| PWP1 |

| |
|---|
| PWP2 |
| PWWP2A |
| PWWP2B |
| PWWP3B |
| PXDC1 |
| PXDN |
| PXK |
| PXMP2 |
| PXMP4 |
| PXN |
| PYCARD |
| PYCR1 |
| PYCR3 |
| PYGB |
| PYGL |
| PYGM |
| PYGO1 |
| PYM1 |
| PYROXD1 |
| PYURF |
| QARS1 |
| QDPR |
| QKI |
| QPCT |
| QPCTL |
| QRFPR |
| QRICH1 |
| QRSL1 |
| QSER1 |
| QSOX1 |
| QSOX2 |
| QTRT1 |
| QTRT2 |
| R3HCC1 |
| R3HCC1L |
| R3HCCIL |
| R3HDM2 |
| R3HDM4 |
| RAB10 |
| RAB11A |
| RAB11B |
| RAB11FIP1 |
| RAB11FIP2 |

| |
|---|
| RAB11FIP3 |
| RAB11FIP4 |
| RAB11FIP5 |
| RAB12 |
| RAB13 |
| RAB14 |
| RAB15 |
| RAB18 |
| RAB19 |
| RAB1A |
| RAB1B |
| RAB20 |
| RAB21 |
| RAB22A |
| RAB23 |
| RAB24 |
| RAB25 |
| RAB27A |
| RAB27B |
| RAB28 |
| RAB29 |
| RAB2A |
| RAB2B |
| RAB30 |
| RAB31 |
| RAB31P |
| RAB32 |
| RAB33B |
| RAB34 |
| RAB35 |
| RAB36 |
| RAB37 |
| RAB38 |
| RAB39A |
| RAB3B |
| RAB3D |
| RAB3GAP1 |
| RAB3GAP2 |
| RAB3IL1 |
| RAB40B |
| RAB40C |
| RAB43 |
| RAB4A |

| |
|---|
| RAB4B |
| RAB5A |
| RAB5C |
| RAB5IF |
| RAB6A |
| RAB8A |
| RAB8B |
| RAB9A |
| RAB9B |
| RABEP1 |
| RABEP2 |
| RABEPK |
| RABGAP1L |
| RABGGTA |
| RABIF |
| RABL2A |
| RABL3 |
| RABL6 |
| RAC1 |
| RAC2 |
| RAC3 |
| RACGAP1 |
| RACK1 |
| RAD1 |
| RAD17 |
| RAD18 |
| RAD21 |
| RAD23A |
| RAD23B |
| RAD50 |
| RAD51 |
| RAD51AP1 |
| RAD51B |
| RAD51C |
| RAD51D |
| RAD52 |
| RAD54B |
| RAD54L2 |
| RAD9A |
| RADX |
| RAE1 |
| RAF1 |
| RAI1 |

| |
|---|
| RAI2 |
| RALA |
| RALB |
| RALBP1 |
| RALGAPA1 |
| RALGAPA2 |
| RALGDS |
| RALGPS1 |
| RALGPS2 |
| RALY |
| RALYL |
| RAMAC |
| RAN |
| RANBP1 |
| RANBP10 |
| RANBP17 |
| RANBP3 |
| RANBP6 |
| RANBP9 |
| RANGAP1 |
| RANGRF |
| RAP1A |
| RAP1B |
| RAP1GAP |
| RAP1GAP2 |
| RAP2A |
| RAP2B |
| RAP2C |
| RAPGEF1 |
| RAPGEF2 |
| RAPGEF3 |
| RAPGEF4 |
| RAPGEF6 |
| RAPGEFL1 |
| RAPH1 |
| RAPSN |
| RARA |
| RARG |
| RARRES1 |
| RARRES2 |
| RARS1 |
| RARS2 |
| RASA1 |

| |
|---|
| RASA2 |
| RASA3 |
| RASAL2 |
| RASAL3 |
| RASD1 |
| RASEF |
| RASGEF1B |
| RASGEF1C |
| RASGRF1 |
| RASGRP2 |
| RASGRP3 |
| RASGRP4 |
| RASIP1 |
| RASL10B |
| RASL11A |
| RASL11B |
| RASSF1 |
| RASSF10 |
| RASSF2 |
| RASSF3 |
| RASSF5 |
| RASSF6 |
| RASSF7 |
| RASSF8 |
| RASSF9 |
| RAVER1 |
| RAVER2 |
| RB1 |
| RB1CC1 |
| RBBP4 |
| RBBP5 |
| RBBP6 |
| RBBP7 |
| RBBP8 |
| RBBP9 |
| RBCK1 |
| RBFA |
| RBFOX2 |
| RBIS |
| RBKS |
| RBL1 |
| RBL2 |
| RBM10 |

| |
|---|
| RBM12 |
| RBM12B |
| RBM14 |
| RBM15 |
| RBM15B |
| RBM17 |
| RBM18 |
| RBM22 |
| RBM25 |
| RBM27 |
| RBM28 |
| RBM3 |
| RBM33 |
| RBM34 |
| RBM41 |
| RBM42 |
| RBM43 |
| RBM45 |
| RBM47 |
| RBM48 |
| RBM4B |
| RBM5 |
| RBM6 |
| RBM7 |
| RBM8A |
| RBMS1 |
| RBMS3 |
| RBMX |
| RBMX2 |
| RBP2 |
| RBP7 |
| RBPMS2 |
| RBSN |
| RBX1 |
| RC3H1 |
| RCAN3 |
| RCBTB1 |
| RCBTB2 |
| RCC1 |
| RCC1L |
| RCC2 |
| RCCD1 |
| RCE1 |

| |
|---|
| RCHY1 |
| RCL1 |
| RCN1 |
| RCN2 |
| RCN3 |
| RCOR1 |
| RCOR2 |
| RCOR3 |
| RCSD1 |
| RDH10 |
| RDH11 |
| RDH13 |
| RDH14 |
| RDM1 |
| RECK |
| RECQL |
| RECQL4 |
| RECQL5 |
| REEP1 |
| REEP2 |
| REEP4 |
| REEP5 |
| REEP6 |
| REL |
| RELA |
| RELB |
| RELCH |
| RELL1 |
| RELL2 |
| RELN |
| RELT |
| REM1 |
| REM2 |
| REN |
| RENBP |
| REP15 |
| REPIN1 |
| REPS1 |
| REPS2 |
| RER1 |
| RERE |
| RERG |
| RESF1 |

| |
|---|
| REST |
| RETREG1 |
| RETREG2 |
| RETREG3 |
| RETSAT |
| REV1 |
| REV3L |
| REX1BD |
| REXO1 |
| REXO2 |
| REXO5 |
| RFC1 |
| RFC2 |
| RFC3 |
| RFC4 |
| RFC5 |
| RFESD |
| RFFL |
| RFK |
| RFLNB |
| RFNG |
| RFT1 |
| RFTN1 |
| RFTN2 |
| RFWD3 |
| RFX1 |
| RFX2 |
| RFX7 |
| RFXANK |
| RFXAP |
| RGCC |
| RGL1 |
| RGL2 |
| RGMA |
| RGMB |
| RGS1 |
| RGS10 |
| RGS11 |
| RGS12 |
| RGS18 |
| RGS19 |
| RGS2 |
| RGS3 |

| |
|---|
| RGS4 |
| RGS5 |
| RGS9BP |
| RHBDD1 |
| RHBDD2 |
| RHBDD3 |
| RHBDF2 |
| RHBDL1 |
| RHBDL2 |
| RHCE |
| RHEB |
| RHNO1 |
| RHOB |
| RHOBTB1 |
| RHOBTB2 |
| RHOBTB3 |
| RHOD |
| RHOF |
| RHOG |
| RHOH |
| RHOQ |
| RHOT1 |
| RHOT2 |
| RHOU |
| RHPN2 |
| RIBC1 |
| RIC1 |
| RIC3 |
| RIC8A |
| RIC8B |
| RICTOR |
| RIDA |
| RIF1 |
| RILPL1 |
| RILPL2 |
| RIMKLA |
| RIMKLB |
| RIMS1 |
| RIMS2 |
| RIN1 |
| RIN3 |
| RING1 |
| RINT1 |

| |
|---|
| RIOK1 |
| RIOK2 |
| RIOK3 |
| RIOX1 |
| RIOX2 |
| RIPK1 |
| RIPK2 |
| RIPOR2 |
| RIT1 |
| RIT2 |
| RITA1 |
| RL10 |
| RLF |
| RLIM |
| RMC1 |
| RMD1 |
| RMDN2 |
| RMDN3 |
| RMI1 |
| RMI2 |
| RMND5A |
| RMND5B |
| RNASE13 |
| RNASE4 |
| RNASE6 |
| RNASEH1 |
| RNASEH2A |
| RNASEH2B |
| RNASEH2C |
| RNASEK |
| RNASET2 |
| RND2 |
| RND3 |
| RNF10 |
| RNF103 |
| RNF11 |
| RNF111 |
| RNF112 |
| RNF113A |
| RNF114 |
| RNF115 |
| RNF123 |
| RNF126 |

| |
|---|
| RNF128 |
| RNF13 |
| RNF133 |
| RNF135 |
| RNF139 |
| RNF14 |
| RNF141 |
| RNF145 |
| RNF148 |
| RNF149 |
| RNF152 |
| RNF166 |
| RNF167 |
| RNF168 |
| RNF169 |
| RNF17 |
| RNF170 |
| RNF180 |
| RNF181 |
| RNF182 |
| RNF185 |
| RNF187 |
| RNF19B |
| RNF2 |
| RNF20 |
| RNF208 |
| RNF213 |
| RNF214 |
| RNF215 |
| RNF216 |
| RNF219 |
| RNF220 |
| RNF222 |
| RNF223 |
| RNF24 |
| RNF25 |
| RNF26 |
| RNF31 |
| RNF32 |
| RNF4 |
| RNF40 |
| RNF41 |
| RNF44 |

| |
|---|
| RNF5 |
| RNF6 |
| RNF7 |
| RNFT1 |
| RNFT2 |
| RNGTT |
| RNH1 |
| RNLS |
| RNMT |
| RNPC3 |
| RNPEP |
| RNPEPL1 |
| RNPS1 |
| RO60 |
| ROBO1 |
| ROCK1 |
| ROCK2 |
| ROGDI |
| ROM1 |
| ROMO1 |
| RP2 |
| RP9 |
| RPA1 |
| RPA2 |
| RPA3 |
| RPA4 |
| RPAIN |
| RPAP1 |
| RPAP3 |
| RPE |
| RPF1 |
| RPF2 |
| RPF3 |
| RPGR |
| RPGRIP1L |
| RPH3AL |
| RPIA |
| RPL10A |
| RPL11 |
| RPL12 |
| RPL13 |
| RPL13A |
| RPL14 |

| |
|---|
| RPL15 |
| RPL17 |
| RPL18 |
| RPL18A |
| RPL19 |
| RPL21 |
| RPL22 |
| RPL22L1 |
| RPL23 |
| RPL23A |
| RPL24 |
| RPL26 |
| RPL27 |
| RPL27A |
| RPL28 |
| RPL29 |
| RPL3 |
| RPL31 |
| RPL32 |
| RPL34 |
| RPL35 |
| RPL36 |
| RPL36A |
| RPL37 |
| RPL37P3 |
| RPL38 |
| RPL4 |
| RPL5 |
| RPL6 |
| RPL7 |
| RPL7A |
| RPL7L1 |
| RPL8 |
| RPL9 |
| RPLP1 |
| RPLP2 |
| RPN1 |
| RPN2 |
| RPP21 |
| RPP25 |
| RPP30 |
| RPP38 |
| RPP40 |

| |
|---|
| RPRD1A |
| RPRD2 |
| RPRM |
| RPS10 |
| RPS12 |
| RPS13 |
| RPS14 |
| RPS15 |
| RPS15A |
| RPS16 |
| RPS17 |
| RPS18 |
| RPS19BP1 |
| RPS2 |
| RPS20 |
| RPS21 |
| RPS23 |
| RPS24 |
| RPS25 |
| RPS26 |
| RPS27 |
| RPS27A |
| RPS27L |
| RPS3 |
| RPS3A |
| RPS5 |
| RPS6 |
| RPS6KA1 |
| RPS6KA2 |
| RPS6KA3 |
| RPS6KA4 |
| RPS6KA5 |
| RPS6KB1 |
| RPS6KB2 |
| RPS6KC1 |
| RPS7 |
| RPS8 |
| RPS9 |
| RPSA |
| RPTOR |
| RPUSD1 |
| RPUSD2 |
| RPUSD4 |

| |
|---|
| RRAD |
| RRAGA |
| RRAGB |
| RRAGC |
| RRAGD |
| RRAS |
| RRAS2 |
| RRBP1 |
| RREB1 |
| RRM1 |
| RRM2 |
| RRM2B |
| RRN3 |
| RRNAD1 |
| RRP1 |
| RRP12 |
| RRP15 |
| RRP1B |
| RRP36 |
| RRP7A |
| RRP8 |
| RRP9 |
| RRS1 |
| RSAD1 |
| RSAD2 |
| RSBN1 |
| RSBN1L |
| RSC1A1 |
| RSF1 |
| RSG1 |
| RSKR |
| RSL1D1 |
| RSPH3 |
| RSPH4A |
| RSPH6A |
| RSPRY1 |
| RSRC2 |
| RSRP1 |
| RT87P |
| RTCA |
| RTCB |
| RTF1 |
| RTF2 |

| |
|---|
| RTL4 |
| RTL5 |
| RTL6 |
| RTL8A |
| RTL8C |
| RTN1 |
| RTN2 |
| RTN3 |
| RTN4 |
| RTN4IP1 |
| RTN4R |
| RTRAF |
| RTTN |
| RUBCN |
| RUBCNL |
| RUFY1 |
| RUFY2 |
| RUFY3 |
| RUNDC1 |
| RUNX1 |
| RUNX2 |
| RUNX3 |
| RUSC1 |
| RUSC2 |
| RUSF1 |
| RUVBL1 |
| RUVBL2 |
| RWDD1 |
| RWDD2A |
| RWDD2B |
| RWDD3 |
| RWDD4 |
| RXFP3 |
| RXFP4 |
| RXRA |
| RXRB |
| RXYLT1 |
| RYBP |
| RYK |
| RYR1 |
| RYR2 |
| RYR3 |
| S100A10 |

| |
|---|
| S100A12 |
| S100A13 |
| S100A16 |
| S100A2 |
| S100A4 |
| S100A6 |
| S100A9 |
| S100P |
| S100PBP |
| S1PR1 |
| S1PR2 |
| S1PR4 |
| SAA2 |
| SAA4 |
| SAAL1 |
| SAC3D1 |
| SACM1L |
| SACS |
| SAE1 |
| SAFB |
| SAFB2 |
| SALL2 |
| SAMD1 |
| SAMD10 |
| SAMD11 |
| SAMD15 |
| SAMD4A |
| SAMD4B |
| SAMD5 |
| SAMD8 |
| SAMD9 |
| SAMD9L |
| SAMHD1 |
| SAMM50 |
| SANBR |
| SAP18 |
| SAP30 |
| SAP30BP |
| SAP30L |
| SAR1A |
| SAR1B |
| SARAF |
| SARDH |

| |
|---|
| SARG |
| SARM1 |
| SARNP |
| SARS1 |
| SARS2 |
| SART1 |
| SART3 |
| SASH1 |
| SASH3 |
| SASS6 |
| SAT1 |
| SAT2 |
| SAV1 |
| SAXO2 |
| SAYSD1 |
| SBDS |
| SBF1 |
| SBF2 |
| SBK1 |
| SBSN |
| SBSPON |
| SC5D |
| SCAF1 |
| SCAF11 |
| SCAF4 |
| SCAF8 |
| SCAI |
| SCAMP1 |
| SCAMP2 |
| SCAMP3 |
| SCAMP4 |
| SCAND1 |
| SCAPER |
| SCARA3 |
| SCARB2 |
| SCARF1 |
| SCARF2 |
| SCCPDH |
| SCD5 |
| SCFD1 |
| SCFD2 |
| SCG5 |
| SCGB1A1 |

| |
|---|
| SCHIP1 |
| SCIN |
| SCLT1 |
| SCLY |
| SCMH1 |
| SCML1 |
| SCML2 |
| SCN10A |
| SCN2A |
| SCN9A |
| SCNM1 |
| SCNN1A |
| SCNN1B |
| SCO1 |
| SCO2 |
| SCOC |
| SCP2 |
| SCPEP1 |
| SCRIB |
| SCRN1 |
| SCRN2 |
| SCRN3 |
| SCRT2 |
| SCUBE1 |
| SCUBE2 |
| SCUBE3 |
| SCX |
| SCYL1 |
| SCYL3 |
| SDAD1 |
| SDC1 |
| SDC2 |
| SDC3 |
| SDC4 |
| SDCBP |
| SDCBP2 |
| SDCCAG8 |
| SDE2 |
| SDF2 |
| SDF2L1 |
| SDF4 |
| SDHAF1 |
| SDHAF3 |

| |
|---|
| SDHB |
| SDHC |
| SDHD |
| SDK1 |
| SDR39U1 |
| SDR42E1 |
| SEC11A |
| SEC11C |
| SEC13 |
| SEC14L1 |
| SEC14L2 |
| SEC16A |
| SEC16B |
| SEC22A |
| SEC22B |
| SEC22C |
| SEC23A |
| SEC23B |
| SEC23IP |
| SEC24A |
| SEC24B |
| SEC24C |
| SEC24D |
| SEC31A |
| SEC31B |
| SEC61A2 |
| SEC61B |
| SEC61G |
| SEC62 |
| SEC63 |
| SECISBP2 |
| SECISBP2L |
| SECTM1 |
| SEH1L |
| SEL1L |
| SEL1L3 |
| SELE |
| SELENBP1 |
| SELENOI |
| SELENOM |
| SELENOO |
| SELENOS |
| SELENOT |

| |
|---|
| SELENOV |
| SELP |
| SELPLG |
| SEM1 |
| SEMA3A |
| SEMA3B |
| SEMA3F |
| SEMA3G |
| SEMA4B |
| SEMA4C |
| SEMA4D |
| SEMA4F |
| SEMA4G |
| SEMA5A |
| SEMA6A |
| SEMA6B |
| SEMA6C |
| SEMA6D |
| SEMA7A |
| SENP1 |
| SENP2 |
| SENP3 |
| SENP5 |
| SENP6 |
| SENP7 |
| SENP8 |
| SEPHS1 |
| SEPHS2 |
| SEPTIN1 |
| SEPTIN10 |
| SEPTIN11 |
| SEPTIN2 |
| SEPTIN5 |
| SEPTIN6 |
| SEPTIN7 |
| SEPTIN8 |
| SEPTIN9 |
| SERAC1 |
| SERBP1 |
| SERGEF |
| SERINC1 |
| SERINC2 |
| SERINC3 |

| |
|---|
| SERINC4 |
| SERINC5 |
| SERP1 |
| Serpina1b |
| SERPINA3 |
| SERPINA5 |
| SERPINB1 |
| SERPINB2 |
| SERPINB5 |
| SERPINB6 |
| SERPINB7 |
| SERPINB9 |
| SERPINE1 |
| SERPINE2 |
| SERPINH1 |
| SERPINI1 |
| SERTAD2 |
| SERTAD3 |
| SESN3 |
| SET |
| SETBP1 |
| SETD1A |
| SETD1B |
| SETD2 |
| SETD4 |
| SETD5 |
| SETD6 |
| SETD7 |
| SETD9 |
| SETMAR |
| SETX |
| SF1 |
| SF3A1 |
| SF3A2 |
| SF3A3 |
| SF3B1 |
| SF3B2 |
| SF3B3 |
| SF3B4 |
| SF3B5 |
| SF3B6 |
| SFI1 |
| SFN |

| |
|---|
| SFPQ |
| SFR1 |
| SFRP1 |
| SFRP4 |
| SFSWAP |
| SFT2D1 |
| SFT2D2 |
| SFT2D3 |
| SFXN1 |
| SFXN2 |
| SFXN4 |
| SFXN5 |
| SGCE |
| SGK1 |
| SGK3 |
| SGMS2 |
| SGO1 |
| SGO2 |
| SGPL1 |
| SGPP1 |
| SGPP2 |
| SGSM1 |
| SGSM2 |
| SGSM3 |
| SGTA |
| SH2B1 |
| SH2B2 |
| SH2B3 |
| SH2D3C |
| SH2D4A |
| SH2D4B |
| SH2D6 |
| SH3BGRL |
| SH3BGRL2 |
| SH3BGRL3 |
| SH3BP1 |
| SH3BP2 |
| SH3BP4 |
| SH3BP5 |
| SH3BP5L |
| SH3D19 |
| SH3D21 |
| SH3GL1 |

| |
|---|
| SH3GL2 |
| SH3GLB1 |
| SH3GLB2 |
| SH3KBP1 |
| SH3PXD2A |
| SH3PXD2B |
| SH3RF1 |
| SH3RF2 |
| SH3TC1 |
| SH3TC2 |
| SH3YL1 |
| SHARPIN |
| SHB |
| SHC1 |
| SHCBP1 |
| SHCBP1L |
| SHD |
| SHF |
| SHFL |
| SHISA4 |
| SHISA5 |
| SHISA8 |
| SHKBP1 |
| SHLD1 |
| SHLD2 |
| SHMT1 |
| SHMT2 |
| SHOC2 |
| SHPK |
| SHPRH |
| SHQ1 |
| SHROOM1 |
| SHROOM2 |
| SHROOM3 |
| SHROOM4 |
| SHTN1 |
| SIAE |
| SIAH1 |
| SIAH2 |
| SIDT2 |
| SIGLEC1 |
| SIGLEC10 |
| SIGLEC14 |

| |
|---|
| SIGLEC5 |
| SIGLEC8 |
| SIGMAR1 |
| SIK2 |
| SIL1 |
| SIMC1 |
| SIN3B |
| SINHCAF |
| SIPA1 |
| SIPA1L1 |
| SIPA1L2 |
| SIPA1L3 |
| SIR4 |
| SIRPA |
| SIRPB1 |
| SIRT1 |
| SIRT2 |
| SIRT3 |
| SIRT5 |
| SIRT6 |
| SIRT7 |
| SIT1 |
| SIVA1 |
| SIX5 |
| SKA2 |
| SKA3 |
| SKI |
| SKIDA1 |
| SKIL |
| SKIV2L |
| SKP1 |
| SKP2 |
| SLA |
| SLA2 |
| SLAIN1 |
| SLAIN2 |
| SLAMF7 |
| SLAMF8 |
| SLAMF9 |
| SLBP |
| SLC10A3 |
| SLC10A4 |
| SLC10A5 |

| |
|---|
| SLC11A1 |
| SLC11A2 |
| SLC12A2 |
| SLC12A4 |
| SLC12A5 |
| SLC12A7 |
| SLC12A9 |
| SLC13A2 |
| SLC13A4 |
| SLC15A1 |
| SLC15A3 |
| SLC15A4 |
| SLC16A1 |
| SLC16A11 |
| SLC16A12 |
| SLC16A14 |
| SLC16A3 |
| SLC16A5 |
| SLC16A6 |
| SLC16A7 |
| SLC17A5 |
| SLC19A1 |
| SLC19A2 |
| SLC19A3 |
| SLC1A1 |
| SLC1A4 |
| SLC1A5 |
| SLC1A6 |
| SLC1A7 |
| SLC20A1 |
| SLC20A2 |
| SLC22A15 |
| SLC22A18 |
| SLC22A23 |
| SLC22A4 |
| SLC22A5 |
| SLC22A7 |
| SLC23A1 |
| SLC23A2 |
| SLC25A1 |
| SLC25A10 |
| SLC25A11 |
| SLC25A12 |

| |
|---|
| SLC25A13 |
| SLC25A15 |
| SLC25A16 |
| SLC25A17 |
| SLC25A19 |
| SLC25A22 |
| SLC25A23 |
| SLC25A24 |
| SLC25A25 |
| SLC25A26 |
| SLC25A27 |
| SLC25A28 |
| SLC25A29 |
| SLC25A30 |
| SLC25A32 |
| SLC25A35 |
| SLC25A36 |
| SLC25A37 |
| SLC25A38 |
| SLC25A39 |
| SLC25A4 |
| SLC25A42 |
| SLC25A43 |
| SLC25A46 |
| SLC25A5 |
| SLC25A51 |
| SLC25A53 |
| SLC26A1 |
| SLC26A11 |
| SLC26A2 |
| SLC26A3 |
| SLC26A4 |
| SLC26A9 |
| SLC27A1 |
| SLC27A3 |
| SLC27A4 |
| SLC27A5 |
| SLC27A6 |
| SLC28A3 |
| SLC29A2 |
| SLC29A3 |
| SLC29A4 |
| SLC2A1 |

| |
|---|
| SLC2A10 |
| SLC2A13 |
| SLC2A2 |
| SLC2A3 |
| SLC2A4RG |
| SLC2A6 |
| SLC2A8 |
| SLC2A9 |
| SLC30A1 |
| SLC30A4 |
| SLC30A5 |
| SLC30A6 |
| SLC30A7 |
| SLC30A9 |
| SLC31A1 |
| SLC31A2 |
| SLC33A1 |
| SLC34A1 |
| SLC34A2 |
| SLC35A1 |
| SLC35A2 |
| SLC35A4 |
| SLC35B1 |
| SLC35B2 |
| SLC35B3 |
| SLC35B4 |
| SLC35C1 |
| SLC35C2 |
| SLC35D2 |
| SLC35E1 |
| SLC35E4 |
| SLC35F2 |
| SLC35F5 |
| SLC35F6 |
| SLC35G2 |
| SLC36A1 |
| SLC36A4 |
| SLC37A3 |
| SLC37A4 |
| SLC38A10 |
| SLC38A2 |
| SLC38A4 |
| SLC38A5 |

| |
|---|
| SLC38A6 |
| SLC38A7 |
| SLC39A1 |
| SLC39A10 |
| SLC39A11 |
| SLC39A14 |
| SLC39A3 |
| SLC39A4 |
| SLC39A5 |
| SLC39A7 |
| SLC39A8 |
| SLC39A9 |
| SLC3A2 |
| SLC40A1 |
| SLC41A1 |
| SLC41A2 |
| SLC41A3 |
| SLC43A1 |
| SLC43A2 |
| SLC44A1 |
| SLC44A2 |
| SLC44A3 |
| SLC44A4 |
| SLC45A1 |
| SLC45A2 |
| SLC45A4 |
| SLC46A1 |
| SLC47A1 |
| SLC47A2 |
| SLC48A1 |
| SLC49A3 |
| SLC49A4 |
| SLC4A10 |
| SLC4A1AP |
| SLC4A2 |
| SLC4A3 |
| SLC50A1 |
| SLC51B |
| SLC52A1 |
| SLC53A3 |
| SLC5A12 |
| SLC5A3 |
| SLC5A6 |

| |
|---|
| SLC66A1 |
| SLC66A2 |
| SLC66A3 |
| Slc6a12 |
| SLC6A15 |
| SLC6A16?? |
| SLC6A3 |
| SLC6A6 |
| SLC6A8 |
| SLC7A11 |
| SLC7A2 |
| SLC7A3 |
| SLC7A4 |
| SLC7A5 |
| SLC7A6 |
| SLC7A6OS |
| SLC7A7 |
| SLC7A8 |
| SLC8A1 |
| SLC8A2 |
| SLC8B1 |
| SLC9A1 |
| SLC9A3R1 |
| SLC9A3R2 |
| SLC9A6 |
| SLC9A7 |
| SLC9A8 |
| SLC9A9 |
| SLC9B1 |
| SLCO2B1 |
| SLCO3A1 |
| SLCO4C1 |
| SLF1 |
| SLF2 |
| SLFN11 |
| SLFN13 |
| SLFN5 |
| SLIRP |
| SLITRK1 |
| SLITRK2 |
| SLITRK6 |
| SLK |
| SLMAP |

| |
|---|
| SLTM |
| SLU7 |
| SLX1A |
| SMAD1 |
| SMAD2 |
| SMAD3 |
| SMAD4 |
| SMAD6 |
| SMAD7 |
| SMAGP |
| SMAP |
| SMAP1 |
| SMAP2 |
| SMARCA1 |
| SMARCA4 |
| SMARCA5 |
| SMARCC1 |
| SMARCC2 |
| SMARCD2 |
| SMARCD3 |
| SMARCE1 |
| SMC1A |
| SMC1B |
| SMC2 |
| SMC3 |
| SMC4 |
| SMC5 |
| SMC6 |
| SMCHD1 |
| SMCR8 |
| SMDT1 |
| SMG5 |
| SMG6 |
| SMG7 |
| SMG8 |
| SMG9 |
| SMIM1 |
| SMIM10L1 |
| SMIM10L2B |
| SMIM12 |
| SMIM13 |
| SMIM14 |
| SMIM15 |

| |
|---|
| SMIM19 |
| SMIM20 |
| SMIM22 |
| SMIM26 |
| SMIM29 |
| SMIM3 |
| SMIM4 |
| SMIM7 |
| SMN1 |
| SMNDC1 |
| SMO |
| SMOX |
| SMPD1 |
| SMPD2 |
| SMPD4 |
| SMPDL3A |
| SMPDL3B |
| SMS |
| SMTN |
| SMU1 |
| SMUG1 |
| SMURF1 |
| SMURF2 |
| SMYD2 |
| SMYD3 |
| SMYD4 |
| SMYD5 |
| SNAI1 |
| SNAP23 |
| SNAP29 |
| SNAP47 |
| SNAPC1 |
| SNAPC3 |
| SNAPC4 |
| SNAPC5 |
| SNAPIN |
| SND1 |
| SNF8 |
| SNIP1 |
| SNN |
| SNPH |
| SNRK |
| SNRNP200 |

| |
|---|
| SNRNP25 |
| SNRNP27 |
| SNRNP35 |
| SNRNP40 |
| SNRNP48 |
| SNRNP70 |
| SNRPA1 |
| SNRPB |
| SNRPB2 |
| SNRPC |
| SNRPD1 |
| SNRPD3 |
| SNRPE |
| SNRPF |
| SNRPG |
| SNRPN |
| SNTA1 |
| SNTB1 |
| SNTB2 |
| SNTG2 |
| SNU13 |
| SNUPN |
| SNURF |
| SNW1 |
| SNX1 |
| SNX10 |
| SNX11 |
| SNX13 |
| SNX14 |
| SNX15 |
| SNX16 |
| SNX17 |
| SNX18 |
| SNX19 |
| SNX2 |
| SNX20 |
| SNX21 |
| SNX24 |
| SNX25 |
| SNX27 |
| SNX29 |
| SNX3 |
| SNX30 |

| |
|---|
| SNX32 |
| SNX5 |
| SNX6 |
| SNX7 |
| SNX8 |
| SNX9 |
| SOAT1 |
| SOBP |
| SOCS1 |
| SOCS2 |
| SOCS3 |
| SOCS4 |
| SOCS5 |
| SOCS6 |
| SOD1 |
| SOD2 |
| SOD3 |
| SOGA1 |
| SOGA3 |
| SON |
| SORBS2 |
| SORBS3 |
| SORD |
| SORL1 |
| SOS1 |
| SOS2 |
| SOSTDC1 |
| SOWAHA |
| SOWAHB |
| SOWAHC |
| SOWAHD |
| SOX12 |
| SOX13 |
| SOX3 |
| SOX4 |
| SOX5 |
| SP100 |
| SP110 |
| SP130 |
| SP140L |
| SP3 |
| SP4 |
| SPA17 |

| |
|---|
| SPAAR |
| SPACA6 |
| SPAG1 |
| SPAG4 |
| SPAG5 |
| SPAG7 |
| SPAG8 |
| SPAG9 |
| SPARC |
| SPART |
| SPAST |
| SPATA13 |
| SPATA18 |
| SPATA2 |
| SPATA20 |
| SPATA24 |
| SPATA2L |
| SPATA31D1 |
| SPATA33 |
| SPATA4 |
| SPATA5 |
| SPATA5L1 |
| SPATA6 |
| SPATA6L |
| SPATA7 |
| SPC24 |
| SPC25 |
| SPCS1 |
| SPCS2 |
| SPCS3 |
| SPDEF |
| SPDL1 |
| SPDYA |
| SPECC1L |
| SPEF2 |
| SPEM3 |
| SPEN |
| SPG11 |
| SPG7 |
| SPHK1 |
| SPHK2 |
| SPHKAP |
| SPI1 |

| |
|---|
| SPICE1 |
| SPIDR |
| SPIN1 |
| SPIN2A |
| SPIN3 |
| SPIN4 |
| SPINDOC |
| SPINT1 |
| SPINT2 |
| SPIRE1 |
| SPNS1 |
| SPOCK1 |
| SPOCK2 |
| SPON1 |
| SPON2 |
| SPOP |
| SPOPL |
| SPOUT1 |
| SPPL2A |
| SPPL2B |
| SPPL3 |
| SPR |
| SPRED2 |
| SPRING1 |
| SPRR1B |
| SPRTN |
| SPRY2 |
| SPRY3 |
| SPRY4 |
| SPRYD7 |
| SPSB1 |
| SPSB3 |
| SPSB4 |
| SPTAN1 |
| SPTB |
| SPTBN1 |
| SPTLC1 |
| SPTLC2 |
| SPTLC3 |
| SPTSSA |
| SPTY2D1 |
| SQLE |
| SQOR |

| |
|---|
| SQSTM1 |
| SRA1 |
| SRBD1 |
| SRC |
| SRCAP |
| SRD5A3 |
| SREBF1 |
| SREBF2 |
| SREK1 |
| SREK1IP1 |
| SRF |
| SRFBP1 |
| SRGAP1 |
| SRGAP2 |
| SRGN |
| SRI |
| SRM |
| SRP14 |
| SRP19 |
| SRP54 |
| SRP68 |
| SRP72 |
| SRP9 |
| SRPK1 |
| SRPK2 |
| SRPK3 |
| SRPRA |
| SRPRB |
| SRR |
| SRRD |
| SRRM1 |
| SRRM2 |
| SRRT |
| SRSF1 |
| SRSF10 |
| SRSF11 |
| SRSF12 |
| SRSF2 |
| SRSF3 |
| SRSF4 |
| SRSF5 |
| SRSF6 |
| SRSF7 |

| |
|---|
| SRSF9 |
| SS18 |
| SS18L1 |
| SS18L2 |
| SSB |
| SSBP1 |
| SSBP2 |
| SSBP3 |
| SSBP4 |
| SSC5D |
| SSH1 |
| SSH2 |
| SSH3 |
| SSNA1 |
| SSPN |
| SSPOP |
| SSR1 |
| SSR2 |
| SSR3 |
| SSR4 |
| SSRP1 |
| SSU72 |
| SSX2IP |
| ST13 |
| ST14 |
| ST3GAL2 |
| ST3GAL3 |
| ST3GAL4 |
| ST3GAL5 |
| ST3GAL6 |
| ST6GAL1 |
| ST6GAL2 |
| ST6GALNAC1 |
| ST6GALNAC2 |
| ST6GALNAC3 |
| ST6GALNAC4 |
| ST6GALNAC6 |
| ST7 |
| ST8SIA4 |
| ST8SIA5 |
| STAB1 |
| STAC |
| STAG1 |

| |
|---|
| STAM |
| STAM2 |
| STAMBP |
| STAP2 |
| STAR |
| STARD10 |
| STARD13 |
| STARD3 |
| STARD3NL |
| STARD4 |
| STARD5 |
| STARD6 |
| STARD7 |
| STARD8 |
| STARD9 |
| STAT1 |
| STAT2 |
| STAT3 |
| STAT5A |
| STAU1 |
| STAU2 |
| STBD1 |
| STC1 |
| STEAP1 |
| STEAP3 |
| STEEP1 |
| STIL |
| STIM1 |
| STIM2 |
| STIMATE |
| STING1 |
| STIP1 |
| STK10 |
| STK11 |
| STK11IP |
| STK16 |
| STK17A |
| STK17B |
| STK19 |
| STK24 |
| STK25 |
| STK26 |
| STK3 |

| |
|---|
| STK32A |
| STK33 |
| STK35 |
| STK36 |
| STK38 |
| STK38L |
| STK39 |
| STMN1 |
| STN1 |
| STOM |
| STOML1 |
| STOML2 |
| STON1 |
| STON2 |
| STOX1 |
| STOX2 |
| STPG2 |
| STRA6 |
| STRADB |
| STRBP |
| STRIP1 |
| STRN |
| STRN3 |
| STRN4 |
| STS |
| STT3A |
| STUB1 |
| STX10 |
| STX11 |
| STX12 |
| STX16 |
| STX17 |
| STX18 |
| STX19 |
| STX1A |
| STX1B |
| STX2 |
| STX4 |
| STX5 |
| STX6 |
| STX7 |
| STX8 |
| STXBP3 |

| |
|---|
| STXBP4 |
| STXBP5 |
| STXBP5L |
| STYK1 |
| STYX |
| SUB1 |
| SUCLA2 |
| SUCLG1 |
| SUCLG2 |
| SUCNR1 |
| SUCO |
| SUFU |
| SUGCT |
| SUGP1 |
| SUGP2 |
| SUGT1 |
| SULT1A1 |
| SULT1A4 |
| SULT1B1 |
| SUMF1 |
| SUMF2 |
| SUMO1 |
| SUMO2 |
| SUMO3 |
| SUN1 |
| SUN2 |
| SUOX |
| SUPT16H |
| SUPT20H |
| SUPT3H |
| SUPT4H1 |
| SUPT5H |
| SUPT6H |
| SUPT7L |
| SUPV3L1 |
| SURF1 |
| SURF2 |
| SURF4 |
| SUSD1 |
| SUSD3 |
| SUV39H1 |
| SUV39H2 |
| SUZ12 |

| |
|---|
| SV2C |
| SVBP |
| SVIL |
| SWAP70 |
| SWI5 |
| SWSAP1 |
| SWT1 |
| SYAP1 |
| SYBU |
| SYCE2 |
| SYDE1 |
| SYDE2 |
| SYF2 |
| SYK |
| SYN1 |
| SYNC |
| SYNCRIP |
| SYNE1 |
| SYNE2 |
| SYNE3 |
| SYNE4 |
| SYNGAP1 |
| SYNGR1 |
| SYNGR3 |
| SYNJ1 |
| SYNJ2 |
| SYNJ2BP |
| SYNM |
| SYNPO2 |
| SYNPO2L |
| SYNRG |
| SYP |
| SYPL1 |
| SYS1 |
| SYT1 |
| SYT10 |
| SYT11 |
| SYT16 |
| SYT17 |
| SYT7 |
| SYT9 |
| SYTC |
| SYTL1 |

| |
|---|
| SYTL2 |
| SYTL3 |
| SYTL4 |
| SYTL5 |
| SYVN1 |
| SZRD1 |
| SZT2 |
| TAB2 |
| TAB3 |
| TAC1 |
| TAC4 |
| TACC2 |
| TACC3 |
| TACR1 |
| TACR3 |
| TADA1 |
| TADA2A |
| TADA2B |
| TADA3 |
| TAF1 |
| TAF10 |
| TAF11 |
| TAF12 |
| TAF13 |
| TAF15 |
| TAF1B |
| TAF1C |
| TAF1D |
| TAF3 |
| TAF4 |
| TAF4B |
| TAF5 |
| TAF5L |
| TAF6 |
| TAF6L |
| TAF7 |
| TAF8 |
| TAF9B |
| TAFA3 |
| TAFAZZIN |
| TAGAP |
| TAGLN2 |
| TAL2 |

| |
|---|
| TALDO1 |
| TALDO1P1 |
| TAMALIN |
| TAMM41 |
| TANGO2 |
| TANGO6 |
| TAOK1 |
| TAOK2 |
| TAP1 |
| TAP2 |
| TAPBPL |
| TAPT1 |
| TARBP1 |
| TARBP2 |
| TARDBP |
| TARM1 |
| TARS2 |
| TARS3 |
| TAS1R2 |
| TAS2R33 |
| TASL |
| TASOR |
| TASOR2 |
| TASP1 |
| TATDN1 |
| TATDN2 |
| TATDN3 |
| TAX1BP1 |
| TAX1BP3 |
| TBATA |
| TBC1D1 |
| TBC1D10A |
| TBC1D10B |
| TBC1D10C |
| TBC1D12 |
| TBC1D13 |
| TBC1D14 |
| TBC1D15 |
| TBC1D16 |
| TBC1D17 |
| TBC1D19 |
| TBC1D2 |
| TBC1D20 |

| |
|---|
| TBC1D22A |
| TBC1D22B |
| TBC1D24 |
| TBC1D2B |
| TBC1D30 |
| TBC1D31 |
| TBC1D32 |
| TBC1D4 |
| TBC1D7 |
| TBC1D8 |
| TBC1D8B |
| TBC1D9 |
| TBC1D9B |
| TBCA |
| TBCB |
| TBCC |
| TBCD |
| TBK1 |
| TBKBP1 |
| TBL1X |
| TBL1XR1 |
| TBL2 |
| TBL3 |
| TBP |
| TBPL1 |
| TBRG1 |
| TBRG4 |
| TBXAS1 |
| TCAIM |
| TCAM1P |
| TCAP |
| TCEA1 |
| TCEA2 |
| TCEA3 |
| TCEAL1 |
| TCEAL3 |
| TCEAL4 |
| TCEAL7 |
| TCEAL8 |
| TCEAL9 |
| TCEANC |
| TCEANC2 |
| TCERG1 |

| |
|---|
| TCF12 |
| TCF20 |
| TCF23 |
| TCF25 |
| TCF3 |
| TCF4 |
| TCF7 |
| TCF7L1 |
| TCF7L2 |
| TCHP |
| TCIM |
| TCN1 |
| TCN2 |
| TCOF1 |
| TCP1 |
| TCP11 |
| TCP11L2 |
| TCTA |
| TCTN1 |
| TCTN2 |
| TDG |
| TDO2 |
| TDP1 |
| TDP2 |
| TDRD12 |
| TDRD15 |
| TDRD3 |
| TDRD7 |
| TEAD1 |
| TEAD3 |
| TEC |
| TECPR1 |
| TECPR2 |
| TECR |
| TEFM |
| TEK |
| TELO2 |
| TEN1 |
| TENM2 |
| TENM3 |
| TENT2 |
| TENT4A |
| TENT4B |

| |
|---|
| TENT5A |
| TENT5C |
| TEP1 |
| TERF1 |
| TERF2 |
| TERF2IP |
| TERT |
| TES |
| TESC |
| TESK1 |
| TESK2 |
| TESPA1 |
| TET1 |
| TET2 |
| TET3 |
| TEX13D |
| TEX14 |
| TEX15 |
| TEX2 |
| TEX261 |
| TEX264 |
| TEX45 |
| TF |
| TF3C6 |
| TFAM |
| TFAP4 |
| TFB1M |
| TFB2M |
| TFCP2 |
| TFDP1 |
| TFE3 |
| TFEB |
| TFG |
| TFIP11 |
| TFPI2 |
| TFPT |
| TFRC |
| TGDS |
| TGFB1 |
| TGFB1I1 |
| TGFBI |
| TGFBR2 |
| TGFBRAP1 |

| |
|---|
| TGIF1 |
| TGIF2 |
| TGM2 |
| TGM3 |
| TGS1 |
| THAP1 |
| THAP11 |
| THAP12 |
| THAP3 |
| THAP4 |
| THAP5 |
| THAP6 |
| THAP7 |
| THAP9 |
| THBS1 |
| THBS2 |
| THBS3 |
| THBS4 |
| THEGL |
| THEM6 |
| THEMIS2 |
| THG1L |
| THNSL1 |
| THNSL2 |
| THOC1 |
| THOC2 |
| THOC3 |
| THOC5 |
| THOC6 |
| THOC7 |
| THOP1 |
| THRA |
| THRAP3 |
| THSD4 |
| THSD7A |
| THTPA |
| THUMPD1 |
| THUMPD2 |
| THUMPD3 |
| THYN1 |
| TIA1 |
| TIAL1 |
| TIAM1 |

| |
|---|
| TIAM2 |
| TICAM2 |
| TIFA |
| TIGAR |
| TIGD1 |
| TIGD2 |
| TIGD3 |
| TIGD4 |
| TIGD5 |
| TIGD6 |
| TIGD7 |
| TIMELESS |
| TIMM10 |
| TIMM10B |
| TIMM13 |
| TIMM17A |
| TIMM17B |
| TIMM21 |
| TIMM23 |
| TIMM29 |
| TIMM44 |
| TIMM50 |
| TIMM8B |
| TIMM9 |
| TIMMDC1 |
| TIMP1 |
| TIMP2 |
| TIMP4 |
| TINAGL1 |
| TINF2 |
| TIPARP |
| TIPIN |
| TIPRL |
| TJAP1 |
| TJP1 |
| TJP2 |
| TJP3 |
| TK1 |
| TK2 |
| TKT |
| TKTL2 |
| TLCD1 |
| TLCD3A |

| |
|---|
| TLCD4 |
| TLE1 |
| TLE2 |
| TLE3 |
| TLE4 |
| TLE5 |
| TLK1 |
| TLK2 |
| TLN1 |
| TLNRD1 |
| TLR1 |
| TLR10 |
| TLR2 |
| TLR3 |
| TLR4 |
| TLR5 |
| TLR6 |
| TLR7 |
| TLR8 |
| TM2D1 |
| TM2D2 |
| TM2D3 |
| TM4SF1 |
| TM4SF20 |
| TM6S1 |
| TM6SF2 |
| TM7SF2 |
| TM7SF3 |
| TM9S3 |
| TM9SF1 |
| TM9SF2 |
| TM9SF4 |
| TMA16 |
| TMBIM1 |
| TMBIM4 |
| TMBIM6 |
| TMC4 |
| TMC5 |
| TMC6 |
| TMC8 |
| TMCC1 |
| TMCC2 |
| TMCC3 |

| |
|---|
| TMCO1 |
| TMCO4 |
| TMCO6 |
| TMED1 |
| TMED10 |
| TMED2 |
| TMED3 |
| TMED4 |
| TMED5 |
| TMED7 |
| TMED9 |
| TMEM101 |
| TMEM104 |
| TMEM106A |
| TMEM106C |
| TMEM108 |
| TMEM109 |
| TMEM11 |
| TMEM115 |
| TMEM117 |
| TMEM120A |
| TMEM120B |
| TMEM121 |
| TMEM123 |
| TMEM125 |
| TMEM126B |
| TMEM127 |
| TMEM128 |
| TMEM129 |
| TMEM131 |
| TMEM131L |
| TMEM132A |
| TMEM134 |
| TMEM135 |
| TMEM138 |
| TMEM139 |
| TMEM140 |
| TMEM141 |
| TMEM143 |
| TMEM144 |
| TMEM147 |
| TMEM14A |
| TMEM14C |

| |
|---|
| TMEM150A |
| TMEM150B |
| TMEM151B |
| TMEM160 |
| TMEM161A |
| TMEM161B |
| TMEM164 |
| TMEM165 |
| TMEM167B |
| TMEM17 |
| TMEM170A |
| TMEM170B |
| TMEM171 |
| TMEM174 |
| TMEM175 |
| TMEM177 |
| TMEM178A |
| TMEM179B |
| TMEM18 |
| TMEM183A |
| TMEM184A |
| TMEM184C |
| TMEM185A |
| TMEM185B |
| TMEM186 |
| TMEM187 |
| TMEM19 |
| TMEM192 |
| TMEM198 |
| TMEM199 |
| TMEM200A |
| TMEM201 |
| TMEM203 |
| TMEM208 |
| TMEM209 |
| TMEM219 |
| TMEM220 |
| TMEM222 |
| TMEM223 |
| TMEM229B |
| TMEM230 |
| TMEM231 |
| TMEM234 |

| |
|---|
| TMEM236 |
| TMEM238 |
| TMEM240 |
| TMEM241 |
| TMEM242 |
| TMEM243 |
| TMEM245 |
| TMEM25 |
| TMEM250 |
| TMEM251 |
| TMEM254 |
| TMEM256 |
| TMEM259 |
| TMEM26 |
| TMEM260 |
| TMEM263 |
| TMEM266 |
| TMEM267 |
| TMEM268 |
| TMEM30A |
| TMEM30B |
| TMEM33 |
| TMEM35A |
| TMEM35B |
| TMEM37 |
| TMEM38A |
| TMEM38B |
| TMEM39A |
| TMEM39B |
| TMEM41A |
| TMEM41B |
| TMEM43 |
| TMEM45A |
| TMEM45B |
| TMEM47 |
| TMEM50A |
| TMEM50B |
| TMEM51 |
| TMEM52 |
| TMEM53 |
| TMEM54 |
| TMEM59 |
| TMEM59L |

| |
|---|
| TMEM60 |
| TMEM61 |
| TMEM62 |
| TMEM64 |
| TMEM65 |
| TMEM67 |
| TMEM68 |
| TMEM70 |
| TMEM81 |
| TMEM86A |
| TMEM87A |
| TMEM87B |
| TMEM88 |
| TMEM89 |
| TMEM8B |
| TMEM9 |
| TMEM92 |
| TMEM94 |
| TMEM97 |
| TMEM98 |
| TMEM9B |
| TMF1 |
| TMLHE |
| TMOD3 |
| TMPO |
| TMPPE |
| TMPRSS2 |
| TMPRSS9 |
| TMSB4X |
| TMTC1 |
| TMTC3 |
| TMTC4 |
| TMUB1 |
| TMUB2 |
| TMX1 |
| TMX2 |
| TMX4 |
| TNC |
| TNF |
| TNFAIP1 |
| TNFAIP2 |
| TNFAIP3 |
| TNFAIP8 |

| |
|---|
| TNFAIP8L2 |
| TNFRSF10A |
| TNFRSF11A |
| TNFRSF12A |
| TNFRSF14 |
| TNFRSF17 |
| TNFRSF1A |
| TNFRSF1B |
| TNFRSF21 |
| TNFRSF6B |
| TNFRSF8 |
| TNFSF12 |
| TNFSF9 |
| TNIK |
| TNIP2 |
| TNK1 |
| TNKS |
| TNKS1BP1 |
| TNNC1 |
| TNNT1 |
| TNPO2 |
| TNPO3 |
| TNRC18 |
| TNRC6A |
| TNRC6B |
| TNS2 |
| TNS3 |
| TNXB |
| TOB2 |
| TOE1 |
| TOGARAM1 |
| TOGARAM2 |
| TOLLIP |
| TOM1 |
| TOM1L1 |
| TOM1L2 |
| TOMM20 |
| TOMM22 |
| TOMM34 |
| TOMM40 |
| TOMM40L |
| TOMM6 |
| TOMM7 |

| |
|---|
| TOMM70 |
| TONSL |
| TOP1 |
| TOP2A |
| TOP2B |
| TOP3B |
| TOPBP1 |
| TOPORS |
| TOR1A |
| TOR1AIP1 |
| TOR1AIP2 |
| TOR1B |
| TOR2A |
| TOR3A |
| TOR4A |
| TOX2 |
| TOX4 |
| TP53 |
| TP53BP1 |
| TP53BP2 |
| TP53I13 |
| TP53I3 |
| TP53INP1 |
| TP53INP2 |
| TP53RK |
| TP53TG5 |
| TPBG |
| TPBGL |
| TPCN2 |
| TPD52 |
| TPD52L2 |
| TPGS1 |
| TPGS2 |
| TPH1 |
| TPK1 |
| TPM1 |
| TPM2 |
| TPM3 |
| TPM4 |
| TPMT |
| TPP1 |
| TPP2 |
| TPPP |

| |
|---|
| TPR |
| TPRA1 |
| TPRN |
| TPST1 |
| TPST2 |
| TPT1 |
| TPTE2 |
| TPX2 |
| TRA2A |
| TRA2B |
| TRADD |
| TRAF1 |
| TRAF2 |
| TRAF3 |
| TRAF3IP2 |
| TRAF3IP3 |
| TRAF4 |
| TRAF5 |
| TRAF6 |
| TRAF7 |
| TRAFD1 |
| TRAIP |
| TRAK1 |
| TRAK2 |
| TRAM1L1 |
| TRAM2 |
| TRANK1 |
| TRAP1 |
| TRAPPC1 |
| TRAPPC10 |
| TRAPPC11 |
| TRAPPC12 |
| TRAPPC13 |
| TRAPPC14 |
| TRAPPC3 |
| TRAPPC4 |
| TRAPPC5 |
| TRAPPC6B |
| TRAPPC8 |
| TRAPPC9 |
| TRAT1 |
| TRBM |
| TRDC |

| |
|---|
| TRDMT1 |
| TREH |
| TREM1 |
| TREM2 |
| TREX1 |
| TRGC1 |
| TRIAP1 |
| TRIB1 |
| TRIB2 |
| TRIB3 |
| TRIL |
| TRIM11 |
| TRIM14 |
| TRIM15 |
| TRIM16 |
| TRIM2 |
| TRIM21 |
| TRIM22 |
| TRIM23 |
| TRIM24 |
| TRIM25 |
| TRIM26 |
| TRIM27 |
| TRIM28 |
| TRIM3 |
| TRIM31 |
| TRIM32 |
| TRIM33 |
| TRIM34 |
| TRIM35 |
| TRIM36 |
| TRIM38 |
| TRIM39 |
| TRIM41 |
| TRIM45 |
| TRIM47 |
| TRIM5 |
| TRIM50 |
| TRIM56 |
| TRIM59 |
| TRIM6 |
| TRIM60 |
| TRIM62 |

| |
|---|
| TRIM65 |
| TRIM66 |
| TRIM68 |
| TRIM7 |
| TRIM8 |
| TRIM9 |
| TRIML1 |
| TRIO |
| TRIP10 |
| TRIP11 |
| TRIP12 |
| TRIP13 |
| TRIP4 |
| TRIQK |
| TRIR |
| TRIT1 |
| TRM11 |
| TRMO |
| TRMT1 |
| TRMT10A |
| TRMT10B |
| TRMT112 |
| TRMT12 |
| TRMT44 |
| TRMT5 |
| TRMT6 |
| TRMT61A |
| TRMT61B |
| TRNAU1AP |
| TRNP1 |
| TRNT1 |
| TROAP |
| TRPC1 |
| TRPC4 |
| TRPC4AP |
| TRPM2 |
| TRPM4 |
| TRPS1 |
| TRPT1 |
| TRPV1 |
| TRPV2 |
| TRRAP |
| TRUB1 |

| |
|---|
| TRUB2 |
| TS1R3 |
| TSBP1 |
| TSC2 |
| TSC22D1 |
| TSC22D3 |
| TSC22D4 |
| TSEN15 |
| TSFM |
| TSG101 |
| TSGA10 |
| TSHR |
| TSHZ1 |
| TSHZ3 |
| TSKS |
| TSKU |
| TSLP |
| TSN |
| TSNARE1 |
| TSNAX |
| TSPAN1 |
| TSPAN14 |
| TSPAN15 |
| TSPAN17 |
| TSPAN3 |
| TSPAN31 |
| TSPAN5 |
| TSPAN6 |
| TSPAN7 |
| TSPAN8 |
| TSPAN9 |
| TSPO |
| TSPOAP1 |
| TSPY26P |
| TSPYL1 |
| TSPYL2 |
| TSPYL4 |
| TSPYL5 |
| TSR1 |
| TSR2 |
| TSR3 |
| TSSC4 |
| TSSK1B |

| |
|---|
| TSSK2 |
| TSSK3 |
| TST |
| TSTD1 |
| TSTD2 |
| TSTD3 |
| TTBK2 |
| TTC1 |
| TTC12 |
| TTC13 |
| TTC14 |
| TTC17 |
| TTC19 |
| TTC21B |
| TTC22 |
| TTC23 |
| TTC26 |
| TTC27 |
| TTC28 |
| TTC30A |
| TTC30B |
| TTC31 |
| TTC32 |
| TTC33 |
| TTC38 |
| TTC39A |
| TTC39B |
| TTC39C |
| TTC4 |
| TTC5 |
| TTC7A |
| TTC7B |
| TTC8 |
| TTC9C |
| TTF1 |
| TTI1 |
| TTI2 |
| TTK |
| TTL |
| TTLL1 |
| TTLL11 |
| TTLL12 |
| TTLL3 |

| |
|---|
| TTLL5 |
| TTLL7 |
| TTN |
| TTPAL |
| TTYH2 |
| TTYH3 |
| TUB |
| TUBA1B |
| TUBA1C |
| TUBA4A |
| TUBA8 |
| TUBB2A |
| TUBB2B |
| TUBB3 |
| TUBD1 |
| TUBE1 |
| TUBG1 |
| TUBG2 |
| TUBGCP2 |
| TUBGCP3 |
| TUBGCP4 |
| TUBGCP5 |
| TUBGCP6 |
| TUFM |
| TULP3 |
| TULP4 |
| TUSC1 |
| TUSC2 |
| TUSC3 |
| TUT1 |
| TUT4 |
| TVP23B |
| TWF1 |
| TWF2 |
| TWIST1 |
| TWNK |
| TWSG1 |
| TXLNA |
| TXN |
| TXN2 |
| TXNDC11 |
| TXNDC15 |
| TXNDC16 |

| |
|---|
| TXNDC17 |
| TXNDC5 |
| TXNDC9 |
| TXNIP |
| TXNL1 |
| TXNL4A |
| TXNL4B |
| TXNRD1 |
| TXNRD3 |
| TYMP |
| TYMS |
| TYRO3 |
| TYSND1 |
| TYW1 |
| TYW3 |
| TYW5 |
| U2AF1L4 |
| U2AF2 |
| U2SURP |
| UACA |
| UAP1L1 |
| UBA1 |
| UBA2 |
| UBA3 |
| UBA5 |
| UBA52 |
| UBA7 |
| UBAC2 |
| UBALD1 |
| UBALD2 |
| UBAP2 |
| UBAP2L |
| UBASH3B |
| UBB |
| UBC |
| UBD |
| UBE2A |
| UBE2B |
| UBE2C |
| UBE2D1 |
| UBE2D2 |
| UBE2D3 |
| UBE2D4 |

| |
|---|
| UBE2E1 |
| UBE2E2 |
| UBE2E3 |
| UBE2F |
| UBE2G2 |
| UBE2H |
| UBE2J1 |
| UBE2J2 |
| UBE2K |
| UBE2L3 |
| UBE2L6 |
| UBE2M |
| UBE2N |
| UBE2O |
| UBE2Q1 |
| UBE2Q2 |
| UBE2R2 |
| UBE2S |
| UBE2T |
| UBE2V1 |
| UBE2V2 |
| UBE2W |
| UBE2Z |
| UBE3B |
| UBE3C |
| UBE3D |
| UBE4A |
| UBE4B |
| UBFD1 |
| UBIAD1 |
| UBL3 |
| UBL4A |
| UBL5 |
| UBL7 |
| UBLCP1 |
| UBN2 |
| UBP1 |
| UBQLN1 |
| UBQLN2 |
| UBQLN4 |
| UBR1 |
| UBR2 |
| UBR3 |

| |
|---|
| UBR4 |
| UBR5 |
| UBR7 |
| UBTD1 |
| UBTD2 |
| UBTF |
| UBXN1 |
| UBXN2A |
| UBXN2B |
| UBXN4 |
| UBXN6 |
| UBXN7 |
| UBXN8 |
| UCHL1 |
| UCHL3 |
| UCHL5 |
| UCK1 |
| UCK2 |
| UCKL1 |
| UCN2 |
| UCP1 |
| UCP2 |
| UCP5 |
| UEVLD |
| UFD1 |
| UFL1 |
| UFM1 |
| UFSP1 |
| UGCG |
| UGDH |
| UGGT2 |
| UGPA |
| UGT1A1 |
| UGT1A10 |
| UGT1A3 |
| UHRF1 |
| UHRF1BP1L |
| UHRF2 |
| UIMC1 |
| ULBP3 |
| ULK1 |
| ULK2 |
| ULK3 |

| |
|---|
| ULK4 |
| UMAD1 |
| UMPS |
| UNC119 |
| UNC119B |
| UNC13B |
| UNC13D |
| UNC45A |
| UNC45B |
| UNC50 |
| UNC5B |
| UNC5C |
| UNC5CL |
| UNC5D |
| UNC93B1 |
| UNG |
| UNK |
| UNKL |
| UPB1 |
| UPF1 |
| UPF2 |
| UPF3A |
| UPF3B |
| UPK1B |
| UPK3A |
| UPP1 |
| UPRT |
| UQCC1 |
| UQCC2 |
| UQCC3 |
| UQCR10 |
| UQCRC1 |
| UQCRC2 |
| UQCRFS1 |
| UQCRH |
| UQCRQ |
| URAH |
| URB1 |
| URB2 |
| URGCP |
| URI1 |
| URM1 |
| UROD |

| |
|---|
| UROS |
| USB1 |
| USE1 |
| USF1 |
| USF2 |
| USF3 |
| USH1C |
| USO1 |
| USP1 |
| USP10 |
| USP11 |
| USP12 |
| USP14 |
| USP15 |
| USP16 |
| USP18 |
| USP19 |
| USP2 |
| USP20 |
| USP21 |
| USP22 |
| USP24 |
| USP25 |
| USP26 |
| USP27X |
| USP28 |
| USP30 |
| USP31 |
| USP33 |
| USP34 |
| USP35 |
| USP37 |
| USP38 |
| USP39 |
| USP4 |
| USP40 |
| USP42 |
| USP44 |
| USP45 |
| USP46 |
| USP47 |
| USP48 |
| USP49 |

| |
|---|
| USP5 |
| USP53 |
| USP54 |
| USP7 |
| USP9X |
| USPL1 |
| UST |
| UTP11 |
| UTP14A |
| UTP15 |
| UTP18 |
| UTP20 |
| UTP23 |
| UTP25 |
| UTP3 |
| UTP4 |
| UTP6 |
| UVRAG |
| UVSSA |
| UXS1 |
| UXT |
| VAC14 |
| VAMP3 |
| VAMP4 |
| VAMP5 |
| VAMP7 |
| VAMP8 |
| VANGL1 |
| VANGL2 |
| VAPA |
| VAPB |
| VARS1 |
| VARS2 |
| VASH1 |
| VASN |
| VASP |
| VAT1 |
| VAV1 |
| VBP1 |
| VCAM1 |
| VCAN |
| VCPKMT |
| VDAC1 |

| |
|---|
| VDAC2 |
| VDAC3 |
| VDR |
| VEGFA |
| VEGFB |
| VEGFC |
| VEPH1 |
| VEZF1 |
| VEZT |
| VGF |
| VGLL4 |
| VHL |
| VIL1 |
| VILL |
| VIM |
| VIRMA |
| VKORC1 |
| VKORC1L1 |
| VMA21 |
| VMP1 |
| VN1R3 |
| VN1R4 |
| VP37B |
| VPS11 |
| VPS13A |
| VPS13B |
| VPS13C |
| VPS13D |
| VPS16 |
| VPS18 |
| VPS25 |
| VPS26B |
| VPS26C |
| VPS28 |
| VPS29 |
| VPS33A |
| VPS33B |
| VPS35 |
| VPS35L |
| VPS36 |
| VPS37A |
| VPS37D |
| VPS39 |

| |
|---|
| VPS41 |
| VPS45 |
| VPS4A |
| VPS4B |
| VPS50 |
| VPS51 |
| VPS52 |
| VPS53 |
| VPS54 |
| VPS72 |
| VPS8 |
| VPS9D1 |
| VRK1 |
| VSIG10 |
| VSIG10L |
| VSIR |
| VSTM4 |
| VSX2 |
| VTA1 |
| VTI1A |
| VTI1B |
| VV |
| VWA1 |
| VWA5A |
| VWA5B2 |
| VWA7 |
| VWF |
| WAC |
| WAPL |
| WARS1 |
| WARS2 |
| WASF1 |
| WASF2 |
| WASF3 |
| WASHC1 |
| WASHC3 |
| WASHC4 |
| WASHC5 |
| WASL |
| WBP1 |
| WBP11 |
| WBP1L |
| WBP2 |

| |
|---|
| WDCP |
| WDFY1 |
| WDFY2 |
| WDFY4 |
| WDHD1 |
| WDPCP |
| WDR1 |
| WDR11 |
| WDR12 |
| WDR13 |
| WDR18 |
| WDR20 |
| WDR24 |
| WDR25 |
| WDR26 |
| WDR27 |
| WDR3 |
| WDR35 |
| WDR36 |
| WDR37 |
| WDR4 |
| WDR41 |
| WDR44 |
| WDR45 |
| WDR45B |
| WDR46 |
| WDR48 |
| WDR5 |
| WDR53 |
| WDR54 |
| WDR55 |
| WDR59 |
| WDR5B |
| WDR6 |
| WDR61 |
| WDR62 |
| WDR7 |
| WDR70 |
| WDR72 |
| WDR73 |
| WDR74 |
| WDR75 |
| WDR76 |

| |
|---|
| WDR77 |
| WDR81 |
| WDR82 |
| WDR83 |
| WDR83OS |
| WDR86 |
| WDR90 |
| WDR91 |
| WDR97 |
| WDTC1 |
| WEE1 |
| WFDC8 |
| WFIKKN1 |
| WFS1 |
| WHAMM |
| WIF1 |
| WIPF3 |
| WIPI1 |
| WIZ |
| WLS |
| WNK1 |
| WNK2 |
| WNK3 |
| WNK4 |
| WNT10B |
| WNT5A |
| WNT6 |
| WNT9A |
| WRAP73 |
| WRNIP1 |
| WSB2 |
| WT1 |
| WTAP |
| WWC1 |
| WWC2 |
| WWC3 |
| WWOX |
| WWP1 |
| WWP2 |
| WWTR1 |
| XAB2 |
| XAF1 |
| XBP1 |

| |
|---|
| XDH |
| XIAP |
| XK |
| XKR2 |
| XKR4 |
| XKR5 |
| XKR8 |
| XPA |
| XPC |
| XPNPEP1 |
| XPNPEP3 |
| XPO1 |
| XPO4 |
| XPO5 |
| XPO6 |
| XPO7 |
| XPOT |
| XPR1 |
| XRCC1 |
| XRCC3 |
| XRCC4 |
| XRCC5 |
| XRCC6 |
| XRN1 |
| XRN2 |
| XXYLT1 |
| XYLT2 |
| YAE1 |
| YAF2 |
| YAP1 |
| YARS1 |
| YARS2 |
| YBEY |
| YBX1 |
| YBX2 |
| YBX3 |
| YEATS2 |
| YEATS4 |
| YES1 |
| YIF1A |
| YIPF1 |
| YIPF2 |
| YIPF3 |

| |
|---|
| YIPF4 |
| YIPF5 |
| YIPF6 |
| YJU2 |
| YJU2B |
| YKT6 |
| YLPM1 |
| YME1L1 |
| YPEL1 |
| YPEL3 |
| YRDC |
| YTHDC1 |
| YTHDC2 |
| YTHDF1 |
| YTHDF2 |
| YTHDF3 |
| YWHAB |
| YWHAE |
| YWHAG |
| YWHAH |
| YWHAQ |
| YWHAZ |
| YY1 |
| ZADH2 |
| ZAN |
| ZAP70 |
| ZAR1 |
| ZAR1L |
| ZBED1 |
| ZBED10P |
| ZBED2 |
| ZBED3 |
| ZBED4 |
| ZBED5 |
| ZBED6 |
| ZBED6CL |
| ZBED8 |
| ZBED9 |
| ZBF343 |
| ZBP1 |
| ZBTB1 |
| ZBTB11 |
| ZBTB12 |

| |
|---|
| ZBTB14 |
| ZBTB16 |
| ZBTB17 |
| ZBTB18 |
| ZBTB2 |
| ZBTB20 |
| ZBTB21 |
| ZBTB22 |
| ZBTB24 |
| ZBTB25 |
| ZBTB3 |
| ZBTB33 |
| ZBTB34 |
| ZBTB37 |
| ZBTB38 |
| ZBTB39 |
| ZBTB4 |
| ZBTB40 |
| ZBTB41 |
| ZBTB42 |
| ZBTB43 |
| ZBTB44 |
| ZBTB45 |
| ZBTB47 |
| ZBTB48 |
| ZBTB49 |
| ZBTB5 |
| ZBTB6 |
| ZBTB7A |
| ZBTB7B |
| ZBTB7C |
| ZBTB8A |
| ZBTB8OS |
| ZBTB9 |
| ZC2HC1C |
| ZC3H1 |
| ZC3H12A |
| ZC3H12C |
| ZC3H13 |
| ZC3H14 |
| ZC3H15 |
| ZC3H18 |
| ZC3H3 |

| |
|---|
| ZC3H4 |
| ZC3H6 |
| ZC3H7A |
| ZC3H7B |
| ZC3H8 |
| ZC3HAV1 |
| ZC3HC1 |
| ZC4H2 |
| ZCCHC10 |
| ZCCHC14 |
| ZCCHC17 |
| ZCCHC18 |
| ZCCHC2 |
| ZCCHC24 |
| ZCCHC3 |
| ZCCHC4 |
| ZCCHC7 |
| ZCCHC8 |
| ZCRB1 |
| ZCWPW1 |
| ZCWPW2 |
| ZDHHC1 |
| ZDHHC12 |
| ZDHHC13 |
| ZDHHC14 |
| ZDHHC15 |
| ZDHHC16 |
| ZDHHC17 |
| ZDHHC18 |
| ZDHHC2 |
| ZDHHC20 |
| ZDHHC23 |
| ZDHHC24 |
| ZDHHC4 |
| ZDHHC5 |
| ZDHHC7 |
| ZDHHC8 |
| ZDHHC9 |
| ZEB1 |
| ZEB2 |
| ZER1 |
| ZFAND1 |
| ZFAND2A |

| |
|---|
| ZFAND2B |
| ZFAND3 |
| ZFAND5 |
| ZFAND6 |
| ZFAT |
| ZFHX2 |
| ZFHX3 |
| ZFP1 |
| ZFP2 |
| ZFP28 |
| ZFP3 |
| ZFP36L1 |
| ZFP36L2 |
| ZFP41 |
| ZFP42 |
| ZFP64 |
| ZFP69 |
| ZFP69B |
| ZFP82 |
| ZFP91 |
| ZFPL1 |
| ZFPM1 |
| ZFR |
| ZFR2 |
| ZFTA |
| ZFX |
| ZFYVE1 |
| ZFYVE16 |
| ZFYVE19 |
| ZFYVE21 |
| ZFYVE26 |
| ZFYVE27 |
| ZFYVE9 |
| ZG16 |
| ZGFR1 |
| ZGPAT |
| ZGRF1 |
| ZHX2 |
| ZHX3 |
| ZKSCAN1 |
| ZKSCAN2 |
| ZKSCAN4 |
| ZKSCAN5 |

| |
|---|
| ZKSCAN7 |
| ZKSCAN8 |
| ZMAT1 |
| ZMAT2 |
| ZMAT3 |
| ZMAT5 |
| ZMIZ1 |
| ZMIZ2 |
| ZMPSTE24 |
| ZMYM1 |
| ZMYM2 |
| ZMYM3 |
| ZMYM4 |
| ZMYM5 |
| ZMYND10 |
| ZMYND11 |
| ZMYND12 |
| ZMYND8 |
| ZNF106 |
| ZNF112 |
| ZNF12 |
| ZNF124 |
| ZNF131 |
| ZNF132 |
| ZNF133 |
| ZNF134 |
| ZNF135 |
| ZNF136 |
| ZNF140 |
| ZNF142 |
| ZNF143 |
| ZNF146 |
| ZNF148 |
| ZNF157 |
| ZNF165 |
| ZNF169 |
| ZNF174 |
| ZNF175 |
| ZNF177 |
| ZNF18 |
| ZNF180 |
| ZNF182 |
| ZNF184 |

| |
|---|
| ZNF19 |
| ZNF197 |
| ZNF2 |
| ZNF205 |
| ZNF207 |
| ZNF208 |
| ZNF211 |
| ZNF212 |
| ZNF213 |
| ZNF214 |
| ZNF217 |
| ZNF219 |
| ZNF22 |
| ZNF227 |
| ZNF229 |
| ZNF236 |
| ZNF239 |
| ZNF24 |
| ZNF248 |
| ZNF25 |
| ZNF250 |
| ZNF251 |
| ZNF252 |
| ZNF256 |
| ZNF26 |
| ZNF260 |
| ZNF263 |
| ZNF268 |
| ZNF27 |
| ZNF271P |
| ZNF274 |
| ZNF275 |
| ZNF276 |
| ZNF277 |
| ZNF280A |
| ZNF280B |
| ZNF281 |
| ZNF282 |
| ZNF283 |
| ZNF286A |
| ZNF287 |
| ZNF292 |
| ZNF296 |

| |
|---|
| ZNF3 |
| ZNF300 |
| ZNF304 |
| ZNF311 |
| ZNF316 |
| ZNF317 |
| ZNF318 |
| ZNF319 |
| ZNF32 |
| ZNF322 |
| ZNF324 |
| ZNF326 |
| ZNF329 |
| ZNF330 |
| ZNF331 |
| ZNF333 |
| ZNF34 |
| ZNF341 |
| ZNF346 |
| ZNF347 |
| ZNF35 |
| ZNF350 |
| ZNF354A |
| ZNF358 |
| ZNF362 |
| ZNF365 |
| ZNF367 |
| ZNF37A |
| ZNF382 |
| ZNF383 |
| ZNF384 |
| ZNF385A |
| ZNF385B |
| ZNF385D |
| ZNF391 |
| ZNF394 |
| ZNF395 |
| ZNF397 |
| ZNF398 |
| ZNF404 |
| ZNF407 |
| ZNF408 |
| ZNF41 |

| |
|---|
| ZNF410 |
| ZNF414 |
| ZNF416 |
| ZNF418 |
| ZNF419 |
| ZNF420 |
| ZNF423 |
| ZNF426 |
| ZNF428 |
| ZNF432 |
| ZNF436 |
| ZNF438 |
| ZNF445 |
| ZNF446 |
| ZNF449 |
| ZNF451 |
| ZNF462 |
| ZNF467 |
| ZNF469 |
| ZNF470 |
| ZNF48 |
| ZNF483 |
| ZNF484 |
| ZNF485 |
| ZNF496 |
| ZNF501 |
| ZNF503 |
| ZNF507 |
| ZNF510 |
| ZNF511 |
| ZNF512 |
| ZNF512B |
| ZNF513 |
| ZNF516 |
| ZNF518A |
| ZNF518B |
| ZNF52 |
| ZNF521 |
| ZNF524 |
| ZNF526 |
| ZNF530 |
| ZNF536 |
| ZNF544 |

| |
|---|
| ZNF548 |
| ZNF549 |
| ZNF550 |
| ZNF551 |
| ZNF554 |
| ZNF555 |
| ZNF564 |
| ZNF565 |
| ZNF566 |
| ZNF567 |
| ZNF568 |
| ZNF57 |
| ZNF570 |
| ZNF574 |
| ZNF575 |
| ZNF576 |
| ZNF579 |
| ZNF581 |
| ZNF583 |
| ZNF584 |
| ZNF585A |
| ZNF586 |
| ZNF593 |
| ZNF594 |
| ZNF596 |
| ZNF597 |
| ZNF598 |
| ZNF599 |
| ZNF608 |
| ZNF609 |
| ZNF613 |
| ZNF614 |
| ZNF618 |
| ZNF619 |
| ZNF621 |
| ZNF622 |
| ZNF623 |
| ZNF628 |
| ZNF629 |
| ZNF638 |
| ZNF641 |
| ZNF644 |
| ZNF646 |

| |
|---|
| ZNF652 |
| ZNF653 |
| ZNF654 |
| ZNF655 |
| ZNF658 |
| ZNF660 |
| ZNF662 |
| ZNF664 |
| ZNF667 |
| ZNF668 |
| ZNF671 |
| ZNF672 |
| ZNF674 |
| ZNF677 |
| ZNF684 |
| ZNF687 |
| ZNF688 |
| ZNF689 |
| ZNF69 |
| ZNF691 |
| ZNF692 |
| ZNF696 |
| ZNF697 |
| ZNF703 |
| ZNF704 |
| ZNF706 |
| ZNF707 |
| ZNF709 |
| ZNF71 |
| ZNF710 |
| ZNF711 |
| ZNF713 |
| ZNF717 |
| ZNF74 |
| ZNF740 |
| ZNF746 |
| ZNF750 |
| ZNF75A |
| ZNF75D |
| ZNF76 |
| ZNF764 |
| ZNF768 |
| ZNF77 |

| |
|---|
| ZNF771 |
| ZNF772 |
| ZNF775 |
| ZNF777 |
| ZNF780B |
| ZNF782 |
| ZNF783 |
| ZNF784 |
| ZNF785 |
| ZNF786 |
| ZNF787 |
| ZNF789 |
| ZNF79 |
| ZNF790 |
| ZNF792 |
| ZNF793 |
| ZNF8 |
| ZNF800 |
| ZNF805 |
| ZNF81 |
| ZNF813 |
| ZNF814 |
| ZNF821 |
| ZNF827 |
| ZNF829 |
| ZNF831 |
| ZNF835 |
| ZNF839 |
| ZNF84 |
| ZNF846 |
| ZNF850 |
| ZNF853 |
| ZNF865 |
| ZNF875 |
| ZNF891 |
| ZNFX1 |
| ZNHIT1 |
| ZNHIT2 |
| ZNHIT3 |
| ZNP709 |
| ZNRD2 |
| ZNRF1 |
| ZNRF3 |

| |
|---|
| ZPR1 |
| ZRANB1 |
| ZRANB2 |
| ZRANB3 |
| ZRSR2 |
| ZSCAN12 |
| ZSCAN16 |
| ZSCAN18 |
| ZSCAN2 |
| ZSCAN20 |
| ZSCAN21 |
| ZSCAN22 |
| ZSCAN23 |
| ZSCAN26 |
| ZSCAN29 |
| ZSCAN30 |
| ZSCAN31 |
| ZSCAN32 |
| ZSCAN4 |
| ZSCAN9 |
| ZSWIM1 |
| ZSWIM3 |
| ZSWIM4 |
| ZSWIM7 |
| ZSWIM8 |
| ZSWIM9 |
| ZUP1 |
| ZW10 |
| ZWILCH |
| ZWINT |
| ZXDC |
| ZYG11B |
| ZYX |
| ZZEF1 |
| ZZZ3 |

**Supplemental List 2**

| Gene Name |
|---|
| A2M |
| A4GALT |
| AAAS |
| AACS |
| AADAC |
| AAGAB |
| AAK1 |
| AAMP |
| AAR2 |
| AARS1 |
| AARS2 |
| AARSD1 |
| AASDH |
| AASDHPPT |
| AASS |
| AATF |
| AATK |
| ABCA1 |
| ABCA12 |
| ABCA2 |
| ABCA3 |
| ABCA4 |
| ABCA5 |
| ABCA7 |
| ABCB1 |
| ABCB10 |
| ABCB6 |
| ABCB7 |
| ABCB8 |
| ABCB9 |
| ABCC1 |
| ABCC10 |
| ABCC11 |
| ABCC3 |
| ABCC4 |
| ABCC9 |
| ABCD1 |
| ABCD3 |
| ABCD4 |
| ABCE1 |
| ABCF1 |
| ABCF2 |
| ABCF3 |

| |
|---|
| ABCG1 |
| ABHD1 |
| ABHD10 |
| ABHD11 |
| ABHD12 |
| ABHD13 |
| ABHD15 |
| ABHD16A |
| ABHD17A |
| ABHD18 |
| ABHD2 |
| ABHD3 |
| ABHD4 |
| ABHD5 |
| ABHD6 |
| ABHD8 |
| ABI1 |
| ABI3BP |
| ABITRAM |
| ABL2 |
| ABR |
| ABRACL |
| ABRAXAS1 |
| ABRAXAS2 |
| ABT1 |
| ABTB1 |
| ABTB2 |
| ACAA1 |
| ACAA2 |
| ACACA |
| ACACB |
| ACAD10 |
| ACAD11 |
| ACAD8 |
| ACADS |
| ACADSB |
| ACADVL |
| ACAP1 |
| ACAP2 |
| ACAP3 |
| ACAT1 |
| ACAT2 |
| ACBD3 |
| ACBD4 |
| ACBD5 |

| |
|---|
| ACBD6 |
| ACCS |
| ACD |
| ACER3 |
| ACHE |
| ACIN1 |
| ACKR2 |
| ACKR4 |
| ACLY |
| ACOT11 |
| ACOT12 |
| ACOT13 |
| ACOT2 |
| ACOT4 |
| ACOT6 |
| ACOT7 |
| ACOT8 |
| ACOT9 |
| ACOX1 |
| ACOX3 |
| ACP1 |
| ACP2 |
| ACP6 |
| ACRBP |
| ACSBG1 |
| ACSF2 |
| ACSF3 |
| ACSL1 |
| ACSL3 |
| ACSL6 |
| ACSM3 |
| ACSS1 |
| ACSS2 |
| ACTA1 |
| ACTA2 |
| ACTB |
| ACTBL2 |
| ACTG1 |
| ACTL6A |
| ACTL7A |
| ACTN1 |
| ACTN4 |
| ACTR10 |
| ACTR1B |
| ACTR2 |

| |
|---|
| ACTR3 |
| ACTR5 |
| ACTR6 |
| ACTR8 |
| ACVR1 |
| ACVR1B |
| ACVR2A |
| ACVR2B |
| ACY1 |
| ACYP1 |
| ACYP2 |
| ADA |
| ADAL |
| ADAM10 |
| ADAM15 |
| ADAM17 |
| ADAM19 |
| ADAM20 |
| ADAM21 |
| ADAM22 |
| ADAM32 |
| ADAM8 |
| ADAM9 |
| ADAMTS1 |
| ADAMTS12 |
| ADAMTS13 |
| ADAMTS2 |
| ADAMTS3 |
| ADAMTS5 |
| ADAMTS6 |
| ADAMTS7 |
| ADAMTS8 |
| ADAMTS9 |
| ADAMTSL4 |
| ADAP1 |
| ADAP2 |
| ADARB1 |
| ADAT1 |
| ADAT2 |
| ADAT3 |
| ADCK1 |
| ADCK2 |
| ADCK5 |
| ADCY1 |
| ADCY3 |

| |
|---|
| ADCY4 |
| ADCY6 |
| ADCY7 |
| ADCY9 |
| ADGRA3 |
| ADGRB1 |
| ADGRB2 |
| ADGRE5 |
| ADGRF1 |
| ADGRG1 |
| ADGRG6 |
| ADGRL1 |
| ADGRL2 |
| ADGRL3 |
| ADGRV1 |
| ADH5 |
| ADHFE1 |
| ADI1 |
| ADIPOR1 |
| ADIPOR2 |
| ADM |
| ADNP2 |
| ADO |
| ADORA1 |
| ADORA2A |
| ADORA2B |
| ADPGK |
| ADPRHL1 |
| ADPRM |
| ADPRS |
| ADRA1D |
| ADRA2A |
| ADRM1 |
| ADSL |
| ADTRP |
| AEBP1 |
| AEBP2 |
| AEN |
| AFAP1 |
| AFF3 |
| AFF4 |
| AFG1L |
| AFG3L2 |
| AFMID |
| AFTPH |

| |
|---|
| AGAP3 |
| AGBL2 |
| AGBL3 |
| AGBL5 |
| AGFG1 |
| AGFG2 |
| AGGF1 |
| AGK |
| AGL |
| AGMAT |
| AGO1 |
| AGO2 |
| AGO3 |
| AGO4 |
| AGPAT1 |
| AGPAT2 |
| AGPAT3 |
| AGPAT5 |
| AGPS |
| AGRN |
| AGTPBP1 |
| AGTR2 |
| AHCTF1 |
| AHCY |
| AHCYL1 |
| AHCYL2 |
| AHDC1 |
| AHI1 |
| AHNAK |
| AHNAK2 |
| AHR |
| AHSA1 |
| AHSA2P |
| AIDA |
| AIF1L |
| AIFM1 |
| AIG1 |
| AIMP1 |
| AIMP2 |
| AIP |
| AJM1 |
| AJUBA |
| AK1 |
| AK2 |
| AK3 |

| |
|---|
| AK4 |
| AK6 |
| AK9 |
| AKAP1 |
| AKAP11 |
| AKAP12 |
| AKAP13 |
| AKAP17A |
| AKAP5 |
| AKAP6 |
| AKAP7 |
| AKAP8 |
| AKAP8L |
| AKAP9 |
| AKIP1 |
| AKIRIN1 |
| AKIRIN2 |
| AKNA |
| AKR1A1 |
| AKR1B1 |
| AKR7L |
| AKT1 |
| AKT1S1 |
| AKT2 |
| AKT3 |
| AKTIP |
| ALAS1 |
| ALCAM |
| ALDH16A1 |
| ALDH18A1 |
| ALDH1A1 |
| ALDH1B1 |
| ALDH1L2 |
| ALDH2 |
| ALDH3B2 |
| ALDH4A1 |
| ALDH5A1 |
| ALDH6A1 |
| ALDH7A1 |
| ALDH9A1 |
| ALDOA |
| ALG1 |
| ALG11 |
| ALG12 |
| ALG13 |

| |
|---|
| ALG14 |
| ALG2 |
| ALG3 |
| ALG5 |
| ALG6 |
| ALG8 |
| ALKBH1 |
| ALKBH2 |
| ALKBH3 |
| ALKBH4 |
| ALKBH5 |
| ALKBH7 |
| ALMS1 |
| ALOX15 |
| ALPK1 |
| ALPK3 |
| ALPL |
| ALS2 |
| ALYREF |
| AMACR |
| AMBRA1 |
| AMD1 |
| AMDHD1 |
| AMER1 |
| AMFR |
| AMH |
| AMIGO1 |
| AMIGO2 |
| AMIGO3 |
| AMMECR1 |
| AMN |
| AMN1 |
| AMOT |
| AMOTL1 |
| AMOTL2 |
| AMPD2 |
| AMPD3 |
| AMZ2 |
| ANAPC1 |
| ANAPC10 |
| ANAPC11 |
| ANAPC13 |
| ANAPC15 |
| ANAPC16 |
| ANAPC2 |

| |
|---|
| ANAPC7 |
| ANG |
| ANGEL1 |
| ANGEL2 |
| ANGPT2 |
| ANGPTL1 |
| ANGPTL5 |
| ANGPTL7 |
| ANK1 |
| ANK2 |
| ANK3 |
| ANKDD1B |
| ANKEF1 |
| ANKFY1 |
| ANKH |
| ANKHD1 |
| ANKIB1 |
| ANKLE1 |
| ANKLE2 |
| ANKMY2 |
| ANKRA2 |
| ANKRD10 |
| ANKRD11 |
| ANKRD12 |
| ANKRD13A |
| ANKRD13C |
| ANKRD13D |
| ANKRD16 |
| ANKRD17 |
| ANKRD2 |
| ANKRD22 |
| ANKRD26 |
| ANKRD27 |
| ANKRD28 |
| ANKRD29 |
| ANKRD31 |
| ANKRD33B |
| ANKRD34A |
| ANKRD37 |
| ANKRD39 |
| ANKRD40 |
| ANKRD46 |
| ANKRD49 |
| ANKRD50 |
| ANKRD52 |

| |
|---|
| ANKRD9 |
| ANKS3 |
| ANKS4B |
| ANKS6 |
| ANKZF1 |
| ANLN |
| ANO10 |
| ANO6 |
| ANO7 |
| ANO8 |
| ANO9 |
| ANOS1 |
| ANP32A |
| ANP32B |
| ANP32E |
| ANTKMT |
| ANTXR1 |
| ANTXR2 |
| ANXA1 |
| ANXA11 |
| ANXA13 |
| ANXA2 |
| ANXA2R |
| ANXA3 |
| ANXA5 |
| AOAH |
| AOC2 |
| AOC3 |
| AOPEP |
| AOX1 |
| AP1AR |
| AP1B1 |
| AP1G1 |
| AP1G2 |
| AP1M1 |
| AP1S1 |
| AP1S2 |
| AP2A1 |
| AP2A2 |
| AP2M1 |
| AP2S1 |
| AP3B1 |
| AP3D1 |
| AP3M1 |
| AP3S1 |

| |
|---|
| AP3S2 |
| AP4B1 |
| AP4E1 |
| AP4M1 |
| AP4S1 |
| AP5B1 |
| AP5M1 |
| AP5S1 |
| AP5Z1 |
| APAF1 |
| APBB1 |
| APBB2 |
| APBB3 |
| APC |
| APEH |
| APEX1 |
| APEX2 |
| APH1A |
| APH1B |
| API5 |
| APIP |
| APLF |
| APLP2 |
| APMAP |
| APOA1 |
| APOBR |
| APOD |
| APOE |
| APOL3 |
| APOLD1 |
| APOM |
| APOO |
| APOOL |
| APP |
| APPBP2 |
| APPL1 |
| APPL2 |
| APRT |
| APTX |
| AQP11 |
| AQP8 |
| AQR |
| AR |
| ARAF |
| ARAP1 |

| |
|---|
| ARAP2 |
| ARC |
| AREL1 |
| ARF1 |
| ARF4 |
| ARF5 |
| ARF6 |
| ARFGAP1 |
| ARFGAP2 |
| ARFGAP3 |
| ARFGEF1 |
| ARFGEF2 |
| ARFGEF3 |
| ARFIP1 |
| ARFIP2 |
| ARFRP1 |
| ARG2 |
| ARGLU1 |
| ARHGAP1 |
| ARHGAP10 |
| ARHGAP11A |
| ARHGAP17 |
| ARHGAP18 |
| ARHGAP19 |
| ARHGAP20 |
| ARHGAP21 |
| ARHGAP22 |
| ARHGAP23 |
| ARHGAP26 |
| ARHGAP29 |
| ARHGAP30 |
| ARHGAP31 |
| ARHGAP32 |
| ARHGAP33 |
| ARHGAP35 |
| ARHGAP39 |
| ARHGAP4 |
| ARHGAP40 |
| ARHGAP42 |
| ARHGAP44 |
| ARHGAP45 |
| ARHGAP5 |
| ARHGAP9 |
| ARHGDIA |
| ARHGDIB |

| |
|---|
| ARHGDIG |
| ARHGEF1 |
| ARHGEF10 |
| ARHGEF10L |
| ARHGEF11 |
| ARHGEF12 |
| ARHGEF16 |
| ARHGEF17 |
| ARHGEF18 |
| ARHGEF19 |
| ARHGEF2 |
| ARHGEF25 |
| ARHGEF3 |
| ARHGEF39 |
| ARHGEF4 |
| ARHGEF40 |
| ARHGEF6 |
| ARHGEF7 |
| ARHGEF9 |
| ARID1A |
| ARID1B |
| ARID2 |
| ARID3A |
| ARID3B |
| ARID4A |
| ARID5A |
| ARID5B |
| ARIH2 |
| ARL1 |
| ARL10 |
| ARL11 |
| ARL13B |
| ARL14EP |
| ARL15 |
| ARL2 |
| ARL2BP |
| ARL3 |
| ARL4A |
| ARL4C |
| ARL4D |
| ARL5A |
| ARL5B |
| ARL6 |
| ARL6IP1 |
| ARL6IP5 |

| |
|---|
| ARL6IP6 |
| ARL8A |
| ARL8B |
| ARMC1 |
| ARMC5 |
| ARMC6 |
| ARMC7 |
| ARMC8 |
| ARMC9 |
| ARMCX1 |
| ARNT2 |
| ARNTL |
| ARPC1B |
| ARPC3 |
| ARPC4 |
| ARPC5 |
| ARPC5L |
| ARPIN |
| ARPP21 |
| ARRB1 |
| ARRDC1 |
| ARRDC2 |
| ARRDC3 |
| ARRDC4 |
| ARSA |
| ARSB |
| ARSG |
| ARSK |
| ART4 |
| AS3MT |
| ASAH1 |
| ASAH2 |
| ASAP1 |
| ASAP2 |
| ASAP3 |
| ASB1 |
| ASB13 |
| ASB16 |
| ASB3 |
| ASB4 |
| ASB6 |
| ASB7 |
| ASB8 |
| ASCC1 |
| ASCC2 |

| |
|---|
| ASCC3 |
| ASCL1 |
| ASF1A |
| ASF1B |
| ASH1L |
| ASL |
| ASMTL |
| ASNS |
| ASNSD1 |
| ASPG |
| ASPH |
| ASPHD2 |
| ASPM |
| ASPRV1 |
| ASPSCR1 |
| ASRGL1 |
| ASTE1 |
| ASXL1 |
| ASXL2 |
| ASXL3 |
| ATAD1 |
| ATAD2 |
| ATAD5 |
| ATE1 |
| ATF2 |
| ATF3 |
| ATF4 |
| ATF5 |
| ATF6 |
| ATF7 |
| ATF7IP |
| ATF7IP2 |
| ATG10 |
| ATG101 |
| ATG12 |
| ATG13 |
| ATG14 |
| ATG16L1 |
| ATG16L2 |
| ATG2A |
| ATG2B |
| ATG3 |
| ATG4A |
| ATG4B |
| ATG4C |

| |
|---|
| ATG5 |
| ATG7 |
| ATG9A |
| ATIC |
| ATL2 |
| ATL3 |
| ATM |
| ATMIN |
| ATN1 |
| ATOH7 |
| ATOX1 |
| ATP10D |
| ATP11A |
| ATP11B |
| ATP11C |
| ATP12A |
| ATP13A1 |
| ATP13A2 |
| ATP1A1 |
| ATP1A3 |
| ATP1B1 |
| ATP1B2 |
| ATP1B3 |
| ATP23 |
| ATP2A1 |
| ATP2A2 |
| ATP2A3 |
| ATP2B4 |
| ATP2C1 |
| ATP4A |
| ATP4B |
| ATP5F1A |
| ATP5F1B |
| ATP5F1C |
| ATP5F1D |
| ATP5F1E |
| ATP5IF1 |
| ATP5MC1 |
| ATP5MC3 |
| ATP5ME |
| ATP5MF |
| ATP5MG |
| ATP5MJ |
| ATP5PB |
| ATP5PD |

| |
|---|
| ATP5PO |
| ATP6AP1 |
| ATP6AP2 |
| ATP6V0A1 |
| ATP6V0A2 |
| ATP6V0A4 |
| ATP6V0B |
| ATP6V0C |
| ATP6V0D1 |
| ATP6V0D2 |
| ATP6V0E1 |
| ATP6V1A |
| ATP6V1B2 |
| ATP6V1D |
| ATP6V1E1 |
| ATP6V1F |
| ATP6V1G1 |
| ATP6V1G2 |
| ATP6V1H |
| ATP7A |
| ATP7B |
| ATP8A1 |
| ATP8A2 |
| ATP8B1 |
| ATP8B2 |
| ATP8B3 |
| ATP9A |
| ATP9B |
| ATPAF1 |
| ATPAF2 |
| ATPSCKMT |
| ATR |
| ATRAID |
| ATRIP |
| ATRN |
| ATRNL1 |
| ATRX |
| ATXN1L |
| ATXN2 |
| ATXN2L |
| ATXN3 |
| ATXN7 |
| ATXN7L1 |
| ATXN7L2 |
| ATXN7L3 |

| |
|---|
| ATXN7L3B |
| AUH |
| AUNIP |
| AUP1 |
| AURKA |
| AURKAIP1 |
| AURKC |
| AUTS2 |
| AVEN |
| AVL9 |
| AVPI1 |
| AXIN1 |
| AXIN2 |
| AXL |
| AZI2 |
| AZIN1 |
| B2M |
| B3GALNT2 |
| B3GALT2 |
| B3GALT4 |
| B3GALT6 |
| B3GAT2 |
| B3GAT3 |
| B3GLCT |
| B3GNT2 |
| B3GNT3 |
| B3GNT4 |
| B3GNT5 |
| B3GNT7 |
| B3GNT8 |
| B3GNT9 |
| B3GNTL1 |
| B4GALNT1 |
| B4GALNT2 |
| B4GALNT3 |
| B4GALT1 |
| B4GALT2 |
| B4GALT3 |
| B4GALT4 |
| B4GALT5 |
| B4GALT7 |
| B4GAT1 |
| B9D1 |
| B9D2 |
| BAALC |

| |
|---|
| BAAT |
| BABAM1 |
| BABAM2 |
| BACE1 |
| BACE2 |
| BACH2 |
| BAD |
| BAG1 |
| BAG2 |
| BAG3 |
| BAG4 |
| BAG5 |
| BAG6 |
| BAHCC1 |
| BAHD1 |
| BAIAP2 |
| BAIAP2L1 |
| BAIAP2L2 |
| BAIAP3 |
| BAK1 |
| BAMBI |
| BANF1 |
| BANP |
| BAP1 |
| BARD1 |
| BASP1 |
| BATF2 |
| BAX |
| BAZ1A |
| BAZ1B |
| BAZ2A |
| BAZ2B |
| BBOF1 |
| BBS1 |
| BBS12 |
| BBS2 |
| BBS4 |
| BBS5 |
| BBS7 |
| BBS9 |
| BBX |
| BCAM |
| BCAP29 |
| BCAP31 |
| BCAR1 |

| |
|---|
| BCAR3 |
| BCAS1 |
| BCAS2 |
| BCAS3 |
| BCAS4 |
| BCAT2 |
| BCCIP |
| BCDIN3D |
| BCHE |
| BCKDHA |
| BCKDHB |
| BCKDK |
| BCL10 |
| BCL11A |
| BCL2 |
| BCL2L1 |
| BCL2L11 |
| BCL2L12 |
| BCL2L15 |
| BCL3 |
| BCL6 |
| BCL6B |
| BCL7B |
| BCL7C |
| BCL9 |
| BCLAF1 |
| BCLAF3 |
| BCO1 |
| BCOR |
| BCORL1 |
| BCR |
| BCS1L |
| BDH2 |
| BDP1 |
| BECN1 |
| BEND3 |
| BEND5 |
| BEND6 |
| BEND7 |
| BEST1 |
| BEST2 |
| BEST4 |
| BET1L |
| BEX2 |
| BEX3 |

| |
|---|
| BEX4 |
| BFAR |
| BFSP1 |
| BHLHA15 |
| BHLHB9 |
| BHLHE40 |
| BHLHE41 |
| BICD1 |
| BICD2 |
| BICDL1 |
| BICDL2 |
| BICRA |
| BICRAL |
| BID |
| BIK |
| BIN1 |
| BIN3 |
| BIRC2 |
| BIRC3 |
| BIRC5 |
| BIRC6 |
| BIVM |
| BLCAP |
| BLM |
| BLMH |
| BLOC1S1 |
| BLOC1S2 |
| BLOC1S3 |
| BLOC1S4 |
| BLOC1S5 |
| BLOC1S6 |
| BLVRA |
| BLVRB |
| BLZF1 |
| BMERB1 |
| BMF |
| BMP1 |
| BMP2 |
| BMP2K |
| BMP3 |
| BMP6 |
| BMPR1A |
| BMPR2 |
| BMS1 |
| BMT2 |

| |
|---|
| BNC2 |
| BNIP1 |
| BNIP3 |
| BNIP3L |
| BOD1 |
| BOD1L1 |
| BOK |
| BOLA1 |
| BOLA3 |
| BOP1 |
| BORA |
| BORCS5 |
| BORCS6 |
| BORCS8 |
| BPGM |
| BPHL |
| BPNT1 |
| BPNT2 |
| BPTF |
| BRAF |
| BRAP |
| BRAT1 |
| BRCA1 |
| BRCC3 |
| BRD1 |
| BRD2 |
| BRD3 |
| BRD3OS |
| BRD4 |
| BRD7 |
| BRD8 |
| BRD9 |
| BRDT |
| BRF1 |
| BRF2 |
| BRI3 |
| BRI3BP |
| BRINP3 |
| BRIP1 |
| BRIX1 |
| BRK1 |
| BRME1 |
| BRMS1 |
| BRMS1L |
| BROX |

| |
|---|
| BRPF1 |
| BRPF3 |
| BRSK1 |
| BRWD1 |
| BRWD3 |
| BSCL2 |
| BSDC1 |
| BSG |
| BSN |
| BSPRY |
| BST1 |
| BST2 |
| BTAF1 |
| BTBD1 |
| BTBD10 |
| BTBD2 |
| BTBD3 |
| BTBD6 |
| BTBD7 |
| BTBD9 |
| BTC |
| BTD |
| BTF3 |
| BTF3L4 |
| BTG1 |
| BTG2 |
| BTG3 |
| BTK |
| BTN1A1 |
| BTN3A3 |
| BTNL9 |
| BTRC |
| BUB1 |
| BUB1B |
| BUB3 |
| BUD13 |
| BUD23 |
| BUD31 |
| BYSL |
| BZW1 |
| BZW2 |
| C10orf143 |
| C11orf1 |
| C11orf24 |
| C11orf42 |

| |
|---|
| C11orf54 |
| C11orf71 |
| C11orf98 |
| C12orf4 |
| C12orf43 |
| C12orf56 |
| C12orf57 |
| C12orf60 |
| C12orf73 |
| C14orf119 |
| C14orf93 |
| C15orf39 |
| C15orf40 |
| C15orf61 |
| C15orf62 |
| C16orf46 |
| C16orf72 |
| C16orf91 |
| C17orf113 |
| C17orf49 |
| C17orf75 |
| C17orf80 |
| C18orf21 |
| C18orf25 |
| C18orf32 |
| C18orf54 |
| C19orf12 |
| C19orf44 |
| C19orf53 |
| C19orf54 |
| C1D |
| C1GALT1 |
| C1GALT1C1 |
| C1orf109 |
| C1orf112 |
| C1orf115 |
| C1orf131 |
| C1orf159 |
| C1orf162 |
| C1orf174 |
| C1orf198 |
| C1orf21 |
| C1orf210 |
| C1orf216 |
| C1orf35 |

| |
|---|
| C1orf43 |
| C1orf50 |
| C1orf52 |
| C1orf53 |
| C1orf74 |
| C1QBP |
| C1QC |
| C1QTNF12 |
| C1QTNF4 |
| C1QTNF6 |
| C1R |
| C1RL |
| C1S |
| C2 |
| C20orf204 |
| C20orf27 |
| C20orf96 |
| C22orf39 |
| C2CD2 |
| C2CD2L |
| C2CD3 |
| C2CD5 |
| C2orf42 |
| C2orf49 |
| C2orf69 |
| C2orf74 |
| C2orf76 |
| C2orf88 |
| C3orf14 |
| C3orf18 |
| C3orf20 |
| C3orf38 |
| C3orf62 |
| C3orf80 |
| C4A |
| C4orf19 |
| C4orf33 |
| C4orf46 |
| C4orf48 |
| C5AR1 |
| C5AR2 |
| C5orf15 |
| C5orf22 |
| C5orf24 |
| C5orf34 |

| |
|---|
| C6orf120 |
| C6orf132 |
| C6orf136 |
| C6orf141 |
| C6orf163 |
| C6orf226 |
| C6orf47 |
| C6orf62 |
| C6orf89 |
| C7orf25 |
| C7orf31 |
| C7orf61 |
| C8G |
| C8orf33 |
| C8orf48 |
| C8orf76 |
| C8orf82 |
| C9orf131 |
| C9orf152 |
| C9orf40 |
| C9orf43 |
| C9orf64 |
| C9orf72 |
| C9orf78 |
| C9orf85 |
| CA12 |
| CA2 |
| CA4 |
| CA5B |
| CA7 |
| CA9 |
| CAAP1 |
| CAB39L |
| CABIN1 |
| CABLES2 |
| CABYR |
| CACFD1 |
| CACHD1 |
| CACNA1B |
| CACNA1G |
| CACNA1H |
| CACNA2D1 |
| CACNB1 |
| CACNB2 |
| CACNB3 |

| |
|---|
| CACTIN |
| CACYBP |
| CAD |
| CADM1 |
| CADM4 |
| CALCOCO1 |
| CALCOCO2 |
| CALD1 |
| CALHM2 |
| CALHM5 |
| CALM1 |
| CALM2 |
| CALR |
| CALU |
| CAMK2D |
| CAMK2G |
| CAMK2N1 |
| CAMKK1 |
| CAMKMT |
| CAMKV |
| CAMLG |
| CAMSAP1 |
| CAMSAP2 |
| CAMSAP3 |
| CAMTA1 |
| CAND1 |
| CANT1 |
| CANX |
| CAP1 |
| CAPG |
| CAPN1 |
| CAPN10 |
| CAPN15 |
| CAPN2 |
| CAPN6 |
| CAPN7 |
| CAPNS1 |
| CAPRIN2 |
| CAPS2 |
| CAPZA1 |
| CAPZA2 |
| CAPZB |
| CARD19 |
| CARD6 |
| CARD8 |

| |
|---|
| CARF |
| CARHSP1 |
| CARM1 |
| CARMIL1 |
| CARNMT1 |
| CARNS1 |
| CARS1 |
| CARS2 |
| CASC3 |
| CASD1 |
| CASKIN1 |
| CASKIN2 |
| CASP10 |
| CASP2 |
| CASP3 |
| CASP7 |
| CASP8 |
| CASP8AP2 |
| CASP9 |
| CASQ1 |
| CASTOR1 |
| CASTOR2 |
| CASZ1 |
| CATSPER2 |
| CATSPERE |
| CATSPERG |
| CAV2 |
| CAVIN1 |
| CAVIN2 |
| CBARP |
| CBFA2T2 |
| CBFB |
| CBL |
| CBLB |
| CBLC |
| CBLL1 |
| CBR1 |
| CBR4 |
| CBS |
| CBWD2 |
| CBX1 |
| CBX2 |
| CBX3 |
| CBX4 |
| CBX5 |

| |
|---|
| CBX6 |
| CBX7 |
| CBX8 |
| CBY1 |
| CC2D1A |
| CC2D1B |
| CC2D2A |
| CC2D2B |
| CCAR1 |
| CCAR2 |
| CCBE1 |
| CCDC102A |
| CCDC106 |
| CCDC107 |
| CCDC112 |
| CCDC115 |
| CCDC117 |
| CCDC12 |
| CCDC121 |
| CCDC122 |
| CCDC124 |
| CCDC125 |
| CCDC126 |
| CCDC127 |
| CCDC137 |
| CCDC138 |
| CCDC14 |
| CCDC142 |
| CCDC149 |
| CCDC15 |
| CCDC157 |
| CCDC158 |
| CCDC159 |
| CCDC160 |
| CCDC163 |
| CCDC166 |
| CCDC167 |
| CCDC168 |
| CCDC171 |
| CCDC174 |
| CCDC177 |
| CCDC18 |
| CCDC180 |
| CCDC181 |
| CCDC183 |

| |
|---|
| CCDC184 |
| CCDC186 |
| CCDC191 |
| CCDC194 |
| CCDC22 |
| CCDC24 |
| CCDC25 |
| CCDC28A |
| CCDC28B |
| CCDC3 |
| CCDC30 |
| CCDC32 |
| CCDC34 |
| CCDC43 |
| CCDC47 |
| CCDC50 |
| CCDC51 |
| CCDC57 |
| CCDC59 |
| CCDC6 |
| CCDC61 |
| CCDC62 |
| CCDC65 |
| CCDC66 |
| CCDC69 |
| CCDC7 |
| CCDC71 |
| CCDC71L |
| CCDC73 |
| CCDC74B |
| CCDC77 |
| CCDC80 |
| CCDC82 |
| CCDC85A |
| CCDC85B |
| CCDC85C |
| CCDC86 |
| CCDC87 |
| CCDC88A |
| CCDC88B |
| CCDC88C |
| CCDC9 |
| CCDC90B |
| CCDC91 |
| CCDC93 |

| |
|---|
| CCDC96 |
| CCDC97 |
| CCHCR1 |
| CCIN |
| CCL2 |
| CCL20 |
| CCL28 |
| CCL5 |
| CCL8 |
| CCM2 |
| CCN1 |
| CCN2 |
| CCNA2 |
| CCNB2 |
| CCNB3 |
| CCNC |
| CCND3 |
| CCNE1 |
| CCNE2 |
| CCNG1 |
| CCNG2 |
| CCNH |
| CCNI |
| CCNJ |
| CCNJL |
| CCNK |
| CCNL1 |
| CCNL2 |
| CCNQ |
| CCNT1 |
| CCNT2 |
| CCNY |
| CCNYL1 |
| CCP110 |
| CCPG1 |
| CCR7 |
| CCS |
| CCSAP |
| CCSER1 |
| CCSER2 |
| CCT2 |
| CCT3 |
| CCT4 |
| CCT5 |
| CCT6A |

| |
|---|
| CCT7 |
| CCT8 |
| CCZ1 |
| CD109 |
| CD14 |
| CD151 |
| CD164 |
| CD180 |
| CD200 |
| CD2AP |
| CD2BP2 |
| CD302 |
| CD36 |
| CD37 |
| CD40 |
| CD44 |
| CD46 |
| CD47 |
| CD55 |
| CD58 |
| CD63 |
| CD68 |
| CD70 |
| CD72 |
| CD74 |
| CD81 |
| CD82 |
| CD84 |
| CD8A |
| CD9 |
| CD99L2 |
| CDADC1 |
| CDAN1 |
| CDC14A |
| CDC14B |
| CDC16 |
| CDC25B |
| CDC25C |
| CDC26 |
| CDC27 |
| CDC34 |
| CDC37 |
| CDC37L1 |
| CDC40 |
| CDC42 |

| |
|---|
| CDC42BPA |
| CDC42BPB |
| CDC42BPG |
| CDC42EP2 |
| CDC42EP3 |
| CDC42EP4 |
| CDC42EP5 |
| CDC42SE2 |
| CDC45 |
| CDC5L |
| CDC6 |
| CDC7 |
| CDC73 |
| CDCA2 |
| CDCA3 |
| CDCA4 |
| CDCA5 |
| CDCA7 |
| CDCA7L |
| CDCA8 |
| CDCP1 |
| CDH10 |
| CDH12 |
| CDH18 |
| CDH2 |
| CDH24 |
| CDH3 |
| CDH6 |
| CDH8 |
| CDHR2 |
| CDHR5 |
| CDIN1 |
| CDIP1 |
| CDIPT |
| CDK1 |
| CDK10 |
| CDK12 |
| CDK13 |
| CDK16 |
| CDK17 |
| CDK18 |
| CDK19 |
| CDK20 |
| CDK2AP1 |
| CDK2AP2 |

| |
|---|
| CDK3 |
| CDK4 |
| CDK5 |
| CDK5R1 |
| CDK5RAP1 |
| CDK5RAP2 |
| CDK6 |
| CDK7 |
| CDK8 |
| CDK9 |
| CDKAL1 |
| CDKL1 |
| CDKL3 |
| CDKL5 |
| CDKN1A |
| CDKN1B |
| CDKN1C |
| CDKN2A |
| CDKN2AIPNL |
| CDKN2B |
| CDKN2C |
| CDKN2D |
| CDKN3 |
| CDO1 |
| CDON |
| CDPF1 |
| CDR2 |
| CDR2L |
| CDS1 |
| CDS2 |
| CDV3 |
| CDYL |
| CDYL2 |
| CEACAM1 |
| CEACAM16 |
| CEACAM21 |
| CEBPA |
| CEBPB |
| CEBPD |
| CEBPG |
| CEBPZ |
| CEBPZOS |
| CECR2 |
| CELF1 |
| CELF2 |

| |
|---|
| CELSR1 |
| CELSR2 |
| CELSR3 |
| CEMIP |
| CEMIP2 |
| CENATAC |
| CENPA |
| CENPB |
| CENPC |
| CENPE |
| CENPF |
| CENPH |
| CENPI |
| CENPJ |
| CENPK |
| CENPM |
| CENPN |
| CENPO |
| CENPP |
| CENPQ |
| CENPT |
| CENPU |
| CENPV |
| CENPW |
| CENPX |
| CEP104 |
| CEP112 |
| CEP126 |
| CEP128 |
| CEP131 |
| CEP135 |
| CEP152 |
| CEP162 |
| CEP164 |
| CEP170B |
| CEP19 |
| CEP192 |
| CEP250 |
| CEP295 |
| CEP295NL |
| CEP350 |
| CEP41 |
| CEP43 |
| CEP44 |
| CEP55 |

| |
|---|
| CEP57 |
| CEP57L1 |
| CEP63 |
| CEP68 |
| CEP70 |
| CEP72 |
| CEP76 |
| CEP78 |
| CEP83 |
| CEP85L |
| CEP89 |
| CEP95 |
| CEP97 |
| CERCAM |
| CERK |
| CERS1 |
| CERS2 |
| CERS4 |
| CERT1 |
| CES1 |
| CETN2 |
| CETN3 |
| CFAP20 |
| CFAP20DC |
| CFAP251 |
| CFAP298 |
| CFAP299 |
| CFAP300 |
| CFAP36 |
| CFAP410 |
| CFAP45 |
| CFAP54 |
| CFAP69 |
| CFAP70 |
| CFAP91 |
| CFB |
| CFD |
| CFDP1 |
| CFH |
| CFL1 |
| CFL2 |
| CGAS |
| CGGBP1 |
| CGN |
| CGNL1 |

| |
|---|
| CGREF1 |
| CGRRF1 |
| CHAC1 |
| CHAC2 |
| CHADL |
| CHAF1A |
| CHAF1B |
| CHCHD2 |
| CHCHD3 |
| CHCHD4 |
| CHCHD6 |
| CHCHD7 |
| CHD1L |
| CHD2 |
| CHD3 |
| CHD4 |
| CHD6 |
| CHD8 |
| CHD9 |
| CHEK1 |
| CHEK2 |
| CHERP |
| CHFR |
| CHGB |
| CHI3L1 |
| CHIC1 |
| CHIC2 |
| CHKA |
| CHL1 |
| CHML |
| CHMP1A |
| CHMP1B |
| CHMP2A |
| CHMP2B |
| CHMP4A |
| CHMP4B |
| CHMP4C |
| CHMP5 |
| CHMP6 |
| CHMP7 |
| CHORDC1 |
| CHP1 |
| CHP2 |
| CHPF2 |
| CHPT1 |

| |
|---|
| CHRAC1 |
| CHRD |
| CHRDL1 |
| CHRM3 |
| CHRM5 |
| CHRNA3 |
| CHST11 |
| CHST12 |
| CHST13 |
| CHST14 |
| CHST15 |
| CHST3 |
| CHST7 |
| CHSY1 |
| CHSY3 |
| CHTOP |
| CHUK |
| CIAO1 |
| CIAO2A |
| CIAO2B |
| CIAO3 |
| CIAPIN1 |
| CIART |
| CIB1 |
| CIB2 |
| CIBAR1 |
| CIC |
| CIITA |
| CILK1 |
| CINP |
| CIP2A |
| CIPC |
| CIR1 |
| CIRBP |
| CISD1 |
| CISD2 |
| CISD3 |
| CISH |
| CIT |
| CITED2 |
| CITED4 |
| CIZ1 |
| CKAP2 |
| CKAP2L |
| CKAP4 |

| |
|---|
| CKAP5 |
| CKB |
| CKLF |
| CKMT1A |
| CKMT2 |
| CKS1B |
| CLASP1 |
| CLASP2 |
| CLASRP |
| CLBA1 |
| CLCA2 |
| CLCN2 |
| CLCN3 |
| CLCN5 |
| CLCN6 |
| CLCN7 |
| CLDN1 |
| CLDN11 |
| CLDN12 |
| CLDN15 |
| CLDN20 |
| CLDN23 |
| CLDN3 |
| CLDN4 |
| CLDN7 |
| CLDN8 |
| CLEC11A |
| CLEC16A |
| CLEC3A |
| CLEC3B |
| CLGN |
| CLHC1 |
| CLIC1 |
| CLIC4 |
| CLINT1 |
| CLIP1 |
| CLIP2 |
| CLIP3 |
| CLK1 |
| CLK2 |
| CLK3 |
| CLK4 |
| CLMN |
| CLMP |
| CLN3 |

| |
|---|
| CLN5 |
| CLN6 |
| CLN8 |
| CLNS1A |
| CLOCK |
| CLPB |
| CLPP |
| CLPTM1 |
| CLPTM1L |
| CLPX |
| CLSPN |
| CLSTN1 |
| CLSTN2 |
| CLSTN3 |
| CLTA |
| CLTB |
| CLTC |
| CLU |
| CLUAP1 |
| CLYBL |
| CMAS |
| CMBL |
| CMC1 |
| CMC2 |
| CMPK1 |
| CMPK2 |
| CMTM3 |
| CMTM6 |
| CMTM7 |
| CMTR1 |
| CMTR2 |
| CMYA5 |
| CNBP |
| CNDP2 |
| CNEP1R1 |
| CNIH1 |
| CNIH4 |
| CNKSR3 |
| CNN1 |
| CNN2 |
| CNN3 |
| CNNM2 |
| CNNM3 |
| CNNM4 |
| CNOT1 |

| |
|---|
| CNOT10 |
| CNOT11 |
| CNOT3 |
| CNOT4 |
| CNOT6 |
| CNOT6L |
| CNOT7 |
| CNOT8 |
| CNP |
| CNPPD1 |
| CNPY2 |
| CNPY3 |
| CNPY4 |
| CNR2 |
| CNTLN |
| CNTN1 |
| CNTN3 |
| CNTN6 |
| CNTNAP1 |
| CNTNAP2 |
| CNTRL |
| CNTROB |
| COA1 |
| COA3 |
| COA4 |
| COA5 |
| COA7 |
| COA8 |
| COASY |
| COBL |
| COBLL1 |
| COCH |
| COG1 |
| COG2 |
| COG3 |
| COG4 |
| COG5 |
| COG6 |
| COG7 |
| COG8 |
| COIL |
| COL15A1 |
| COL17A1 |
| COL1A1 |
| COL21A1 |

| |
|---|
| COL3A1 |
| COL4A1 |
| COL4A2 |
| COL4A3 |
| COL4A4 |
| COL4A5 |
| COL4A6 |
| COL6A1 |
| COL6A2 |
| COL7A1 |
| COL9A2 |
| COLEC12 |
| COLGALT1 |
| COMMD1 |
| COMMD10 |
| COMMD2 |
| COMMD4 |
| COMMD5 |
| COMMD6 |
| COMMD7 |
| COMMD8 |
| COMMD9 |
| COMT |
| COMTD1 |
| COP1 |
| COPA |
| COPB1 |
| COPB2 |
| COPE |
| COPG1 |
| COPG2 |
| COPRS |
| COPS3 |
| COPS5 |
| COPS6 |
| COPS7A |
| COPS8 |
| COPZ1 |
| COPZ2 |
| COQ10A |
| COQ10B |
| COQ2 |
| COQ3 |
| COQ5 |
| COQ6 |

| |
|---|
| COQ7 |
| COQ8A |
| COQ8B |
| COQ9 |
| CORIN |
| CORO1A |
| CORO1B |
| CORO1C |
| CORO2A |
| CORO6 |
| CORO7 |
| COTL1 |
| COX10 |
| COX11 |
| COX15 |
| COX16 |
| COX18 |
| COX19 |
| COX20 |
| COX4I1 |
| COX4I2 |
| COX5A |
| COX5B |
| COX6A1 |
| COX6B1 |
| COX6C |
| COX7A1 |
| COX7A2 |
| COX7A2L |
| COX8A |
| CPD |
| CPEB1 |
| CPEB2 |
| CPEB4 |
| CPED1 |
| CPLANE1 |
| CPLANE2 |
| CPNE2 |
| CPNE3 |
| CPOX |
| CPPED1 |
| CPSF1 |
| CPSF2 |
| CPSF3 |
| CPSF4 |

| |
|---|
| CPT1A |
| CPT1B |
| CPT1C |
| CPT2 |
| CPTP |
| CPVL |
| CPXM1 |
| CPZ |
| CRACD |
| CRACDL |
| CRADD |
| CRAMP1 |
| CRAT |
| CRBN |
| CRCP |
| CREB1 |
| CREB3 |
| CREB3L1 |
| CREB3L2 |
| CREBL2 |
| CREBRF |
| CREBZF |
| CREG1 |
| CRELD1 |
| CRELD2 |
| CREM |
| CRIM1 |
| CRIP1 |
| CRIP3 |
| CRIPT |
| CRK |
| CRKL |
| CRLF3 |
| CRLS1 |
| CRNKL1 |
| CROCC |
| CRPPA |
| CRTAP |
| CRTC3 |
| CRY2 |
| CRYAB |
| CRYBG1 |
| CRYBG2 |
| CRYBG3 |
| CRYM |

| |
|---|
| CRYZ |
| CRYZL1 |
| CS |
| CSAD |
| CSDE1 |
| CSF1 |
| CSF1R |
| CSGALNACT1 |
| CSGALNACT2 |
| CSK |
| CSKMT |
| CSMD3 |
| CSNK1A1 |
| CSNK1D |
| CSNK1E |
| CSNK1G1 |
| CSNK1G2 |
| CSNK1G3 |
| CSNK2B |
| CSPG4 |
| CSPG5 |
| CSPP1 |
| CSRNP1 |
| CSRNP2 |
| CSRP1 |
| CSRP2 |
| CST3 |
| CST6 |
| CST7 |
| CSTF1 |
| CSTF2 |
| CSTF2T |
| CSTF3 |
| CSTL1 |
| CTBP1 |
| CTBP2 |
| CTC1 |
| CTCF |
| CTDNEP1 |
| CTDP1 |
| CTDSP1 |
| CTDSP2 |
| CTF1 |
| CTH |
| CTHRC1 |

| |
|---|
| CTIF |
| CTNNA1 |
| CTNNA2 |
| CTNNAL1 |
| CTNNB1 |
| CTNNBL1 |
| CTNND1 |
| CTPS1 |
| CTR9 |
| CTRL |
| CTSA |
| CTSB |
| CTSC |
| CTSD |
| CTSH |
| CTSK |
| CTSL |
| CTSO |
| CTSV |
| CTSZ |
| CTTN |
| CTTNBP2NL |
| CTU1 |
| CTU2 |
| CTXN1 |
| CUEDC2 |
| CUL1 |
| CUL2 |
| CUL3 |
| CUL4B |
| CUL5 |
| CUL7 |
| CUL9 |
| CUTA |
| CUTC |
| CUX1 |
| CUZD1 |
| CWC15 |
| CWC22 |
| CWC25 |
| CWC27 |
| CWF19L1 |
| CWF19L2 |
| CX3CL1 |
| CX3CR1 |

| |
|---|
| CXADR |
| CXCL10 |
| CXCL16 |
| CXCL8 |
| CXCR4 |
| CXorf38 |
| CXXC4 |
| CYB561A3 |
| CYB561D2 |
| CYB5B |
| CYB5D2 |
| CYB5R1 |
| CYB5R2 |
| CYB5R3 |
| CYB5R4 |
| CYBC1 |
| CYBRD1 |
| CYC1 |
| CYCS |
| CYFIP2 |
| CYHR1 |
| CYLD |
| CYP19A1 |
| CYP1B1 |
| CYP20A1 |
| CYP27A1 |
| CYP27B1 |
| CYP2D6 |
| CYP2J2 |
| CYP2U1 |
| CYP4B1 |
| CYP4F3 |
| CYP4V2 |
| CYP8B1 |
| CYREN |
| CYRIB |
| CYSLTR1 |
| CYSTM1 |
| CYTH2 |
| CYTH3 |
| CZIB |
| D2HGDH |
| DAAM2 |
| DAB2 |
| DAB2IP |

| |
|---|
| DACH1 |
| DACH2 |
| DAD1 |
| DAG1 |
| DAGLA |
| DAGLB |
| DALRD3 |
| DAP |
| DAPK1 |
| DAPK3 |
| DARS1 |
| DAXX |
| DAZAP1 |
| DAZAP2 |
| DBF4 |
| DBF4B |
| DBI |
| DBN1 |
| DBNDD1 |
| DBNL |
| DBP |
| DBR1 |
| DBT |
| DCAF1 |
| DCAF10 |
| DCAF11 |
| DCAF12 |
| DCAF12L2 |
| DCAF13 |
| DCAF15 |
| DCAF16 |
| DCAF17 |
| DCAF4 |
| DCAF5 |
| DCAF6 |
| DCAF7 |
| DCAF8 |
| DCAKD |
| DCBLD1 |
| DCBLD2 |
| DCDC2 |
| DCHS1 |
| DCK |
| DCLRE1A |
| DCLRE1C |

| |
|---|
| DCP1A |
| DCP1B |
| DCP2 |
| DCPS |
| DCTD |
| DCTN1 |
| DCTN2 |
| DCTN3 |
| DCTN5 |
| DCTN6 |
| DCTPP1 |
| DCUN1D1 |
| DCUN1D2 |
| DCUN1D3 |
| DCUN1D5 |
| DCXR |
| DDA1 |
| DDAH2 |
| DDB1 |
| DDB2 |
| DDHD1 |
| DDHD2 |
| DDI2 |
| DDIAS |
| DDIT3 |
| DDIT4 |
| DDN |
| DDOST |
| DDR1 |
| DDR2 |
| DDRGK1 |
| DDT |
| DDX1 |
| DDX10 |
| DDX11L8 |
| DDX17 |
| DDX18 |
| DDX19A |
| DDX19B |
| DDX20 |
| DDX21 |
| DDX23 |
| DDX24 |
| DDX27 |
| DDX28 |

| |
|---|
| DDX31 |
| DDX39A |
| DDX39B |
| DDX3X |
| DDX41 |
| DDX42 |
| DDX46 |
| DDX47 |
| DDX49 |
| DDX5 |
| DDX50 |
| DDX52 |
| DDX54 |
| DDX55 |
| DDX56 |
| DDX58 |
| DDX59 |
| DDX6 |
| DDX60 |
| DDX60L |
| DEAF1 |
| DECR1 |
| DECR2 |
| DEDD |
| DEDD2 |
| DEF8 |
| DEFA5 |
| DEGS1 |
| DEGS2 |
| DENND10 |
| DENND1A |
| DENND1B |
| DENND1C |
| DENND2A |
| DENND2B |
| DENND2C |
| DENND2D |
| DENND3 |
| DENND4A |
| DENND4B |
| DENND4C |
| DENND5A |
| DENND5B |
| DENND6A |
| DENND6B |

| |
|---|
| DENR |
| DEPDC1 |
| DEPDC1B |
| DEPDC4 |
| DEPDC5 |
| DEPDC7 |
| DEPTOR |
| DERL1 |
| DERL2 |
| DERL3 |
| DESI1 |
| DESI2 |
| DET1 |
| DEXI |
| DFFA |
| DFFB |
| DGAT1 |
| DGAT2 |
| DGCR2 |
| DGCR6 |
| DGCR8 |
| DGKA |
| DGKD |
| DGKE |
| DGKQ |
| DGKZ |
| DGUOK |
| DHCR24 |
| DHCR7 |
| DHDDS |
| DHDH |
| DHFR |
| DHH |
| DHODH |
| DHPS |
| DHRS1 |
| DHRS11 |
| DHRS13 |
| DHRS3 |
| DHRS4 |
| DHRS7 |
| DHRS7B |
| DHRSX |
| DHTKD1 |
| DHX15 |

| |
|---|
| DHX16 |
| DHX29 |
| DHX30 |
| DHX32 |
| DHX33 |
| DHX34 |
| DHX35 |
| DHX36 |
| DHX37 |
| DHX38 |
| DHX40 |
| DHX57 |
| DHX9 |
| DIABLO |
| DIAPH2 |
| DIAPH3 |
| DICER1 |
| DIDO1 |
| DIMT1 |
| DIP2A |
| DIP2B |
| DIP2C |
| DIPK1A |
| DIPK1B |
| DIPK2A |
| DIRAS1 |
| DIRAS2 |
| DIRAS3 |
| DIS3 |
| DIS3L |
| DIS3L2 |
| DIXDC1 |
| DKC1 |
| DKK3 |
| DKKL1 |
| DLAT |
| DLC1 |
| DLD |
| DLEC1 |
| DLEU7 |
| DLG1 |
| DLG3 |
| DLG4 |
| DLG5 |
| DLGAP1 |

| |
|---|
| DLGAP5 |
| DLL1 |
| DLL3 |
| DLST |
| DLX5 |
| DMAC2L |
| DMAP1 |
| DMD |
| DMKN |
| DMPK |
| DMTF1 |
| DMWD |
| DMXL1 |
| DMXL2 |
| DNA2 |
| DNAAF2 |
| DNAAF3 |
| DNAAF5 |
| DNAH1 |
| DNAH2 |
| DNAH3 |
| DNAH7 |
| DNAJA1 |
| DNAJA2 |
| DNAJA3 |
| DNAJA4 |
| DNAJB1 |
| DNAJB12 |
| DNAJB14 |
| DNAJB2 |
| DNAJB4 |
| DNAJB6 |
| DNAJB7 |
| DNAJB9 |
| DNAJC1 |
| DNAJC10 |
| DNAJC11 |
| DNAJC12 |
| DNAJC13 |
| DNAJC14 |
| DNAJC16 |
| DNAJC17 |
| DNAJC18 |
| DNAJC19 |
| DNAJC2 |

| |
|---|
| DNAJC21 |
| DNAJC24 |
| DNAJC25 |
| DNAJC27 |
| DNAJC28 |
| DNAJC3 |
| DNAJC30 |
| DNAJC4 |
| DNAJC5 |
| DNAJC7 |
| DNAJC8 |
| DNAJC9 |
| DNAL1 |
| DNAL4 |
| DNASE1L1 |
| DNASE2 |
| DNHD1 |
| DNLZ |
| DNM1 |
| DNM1L |
| DNM2 |
| DNM3 |
| DNMBP |
| DNMT1 |
| DNMT3A |
| DNMT3B |
| DNPEP |
| DNPH1 |
| DNTTIP1 |
| DNTTIP2 |
| DOC2B |
| DOCK1 |
| DOCK11 |
| DOCK3 |
| DOCK4 |
| DOCK5 |
| DOCK6 |
| DOCK7 |
| DOCK8 |
| DOCK9 |
| DOHH |
| DOK1 |
| DOK3 |
| DOK4 |
| DOK7 |

| |
|---|
| DOLK |
| DOLPP1 |
| DOP1A |
| DOP1B |
| DOT1L |
| DPCD |
| DPF2 |
| DPF3 |
| DPH2 |
| DPH3 |
| DPH5 |
| DPH6 |
| DPH7 |
| DPM1 |
| DPM2 |
| DPM3 |
| DPP3 |
| DPP7 |
| DPY19L1 |
| DPY19L3 |
| DPY19L4 |
| DPY30 |
| DPYD |
| DPYSL2 |
| DPYSL3 |
| DPYSL5 |
| DQX1 |
| DR1 |
| DRAM2 |
| DRAP1 |
| DRC3 |
| DRG1 |
| DRG2 |
| DROSHA |
| DRP2 |
| DSC2 |
| DSC3 |
| DSCC1 |
| DSE |
| DSEL |
| DSG2 |
| DSP |
| DST |
| DSTN |
| DSTYK |

| |
|---|
| DTD1 |
| DTD2 |
| DTL |
| DTNB |
| DTNBP1 |
| DTWD1 |
| DTWD2 |
| DTX1 |
| DTX2 |
| DTX3 |
| DTX3L |
| DTYMK |
| DUOX1 |
| DUS1L |
| DUS2 |
| DUS3L |
| DUS4L |
| DUSP1 |
| DUSP10 |
| DUSP11 |
| DUSP12 |
| DUSP14 |
| DUSP16 |
| DUSP18 |
| DUSP19 |
| DUSP23 |
| DUSP28 |
| DUSP3 |
| DUSP5 |
| DUSP6 |
| DUSP7 |
| DUSP8 |
| DUSP9 |
| DUT |
| DVL1 |
| DVL2 |
| DXO |
| DYM |
| DYNC1H1 |
| DYNC1I2 |
| DYNC1LI1 |
| DYNC1LI2 |
| DYNC2H1 |
| DYNC2I1 |
| DYNC2I2 |

| |
|---|
| DYNC2LI1 |
| DYNLL1 |
| DYNLL2 |
| DYNLRB1 |
| DYNLT1 |
| DYNLT2 |
| DYNLT2B |
| DYNLT3 |
| DYNLT4 |
| DYRK1A |
| DYRK1B |
| DYRK2 |
| DYRK3 |
| DZANK1 |
| DZIP1 |
| DZIP3 |
| E2F1 |
| E2F2 |
| E2F3 |
| E2F4 |
| E2F5 |
| E2F6 |
| E2F7 |
| E4F1 |
| EAF1 |
| EAF2 |
| EAPP |
| EARS2 |
| EBAG9 |
| EBF4 |
| EBNA1BP2 |
| EBP |
| EBPL |
| ECD |
| ECE1 |
| ECH1 |
| ECHDC1 |
| ECHDC3 |
| ECHS1 |
| ECI1 |
| ECI2 |
| ECPAS |
| ECSIT |
| ECT2 |
| EDC3 |

| |
|---|
| EDC4 |
| EDEM1 |
| EDEM2 |
| EDEM3 |
| EDF1 |
| EDIL3 |
| EDN1 |
| EDNRA |
| EDRF1 |
| EEA1 |
| EED |
| EEF1A1 |
| EEF1A1P5 |
| EEF1AKMT1 |
| EEF1AKMT2 |
| EEF1B2 |
| EEF1D |
| EEF1E1 |
| EEF1G |
| EEF2 |
| EEF2K |
| EEF2KMT |
| EEFSEC |
| EEPD1 |
| EFCAB11 |
| EFCAB14 |
| EFCAB2 |
| EFCAB6 |
| EFCAB7 |
| EFEMP2 |
| EFHD2 |
| EFL1 |
| EFNA1 |
| EFNA2 |
| EFNA5 |
| EFNB1 |
| EFNB2 |
| EFR3A |
| EFTUD2 |
| EGFL7 |
| EGFL8 |
| EGLN1 |
| EGLN2 |
| EGLN3 |
| EGR1 |

| |
|---|
| EGR2 |
| EGR3 |
| EHBP1 |
| EHBP1L1 |
| EHD1 |
| EHD2 |
| EHD4 |
| EHF |
| EHHADH |
| EHMT1 |
| EHMT2 |
| EI24 |
| EID1 |
| EID2 |
| EIF1 |
| EIF1AX |
| EIF1B |
| EIF2AK1 |
| EIF2AK2 |
| EIF2AK3 |
| EIF2AK4 |
| EIF2B1 |
| EIF2B2 |
| EIF2B4 |
| EIF2B5 |
| EIF2D |
| EIF2S1 |
| EIF2S2 |
| EIF2S3 |
| EIF3A |
| EIF3B |
| EIF3D |
| EIF3E |
| EIF3F |
| EIF3G |
| EIF3I |
| EIF3J |
| EIF3K |
| EIF3L |
| EIF3M |
| EIF4A1 |
| EIF4A2 |
| EIF4A3 |
| EIF4B |
| EIF4E |

| |
|---|
| EIF4E2 |
| EIF4E3 |
| EIF4EBP1 |
| EIF4EBP2 |
| EIF4ENIF1 |
| EIF4G1 |
| EIF4G2 |
| EIF4G3 |
| EIF4H |
| EIF5 |
| EIF5A2 |
| EIF5B |
| EIPR1 |
| ELAC1 |
| ELAC2 |
| ELAPOR2 |
| ELAVL2 |
| ELF1 |
| ELF2 |
| ELF3 |
| ELF4 |
| ELK1 |
| ELK3 |
| ELK4 |
| ELL |
| ELL2 |
| ELMO1 |
| ELMO2 |
| ELMO3 |
| ELMOD1 |
| ELMOD2 |
| ELMOD3 |
| ELOA |
| ELOB |
| ELOC |
| ELOVL1 |
| ELOVL5 |
| ELOVL6 |
| ELP1 |
| ELP2 |
| ELP3 |
| ELP4 |
| ELP5 |
| ELP6 |
| EMB |

| |
|---|
| EMC1 |
| EMC10 |
| EMC2 |
| EMC3 |
| EMC4 |
| EMC6 |
| EMC7 |
| EMC8 |
| EMC9 |
| EMD |
| EME1 |
| EME2 |
| EMG1 |
| EMID1 |
| EMILIN1 |
| EMILIN2 |
| EMILIN3 |
| EML1 |
| EML2 |
| EML3 |
| EML4 |
| EML6 |
| EMP1 |
| EMP2 |
| EMP3 |
| EMSY |
| ENAH |
| ENC1 |
| ENDOD1 |
| ENDOG |
| ENDOV |
| ENGASE |
| ENHO |
| ENKD1 |
| ENO3 |
| ENOPH1 |
| ENOSF1 |
| ENOX1 |
| ENOX2 |
| ENPP1 |
| ENPP4 |
| ENSA |
| ENTPD1 |
| ENTPD2 |
| ENTPD4 |

| |
|---|
| ENTPD5 |
| ENTPD6 |
| ENTPD7 |
| ENTPD8 |
| ENY2 |
| EOLA1 |
| EP400 |
| EPAS1 |
| EPB41 |
| EPB41L2 |
| EPB41L3 |
| EPB41L4B |
| EPB41L5 |
| EPC1 |
| EPC2 |
| EPCAM |
| EPDR1 |
| EPG5 |
| EPHA1 |
| EPHA2 |
| EPHA4 |
| EPHA6 |
| EPHB1 |
| EPHB4 |
| EPHB6 |
| EPHX1 |
| EPHX2 |
| EPM2A |
| EPM2AIP1 |
| EPN1 |
| EPN2 |
| EPN3 |
| EPOP |
| EPRS1 |
| EPS15 |
| EPS15L1 |
| EPS8L2 |
| ERAL1 |
| ERAP1 |
| ERAS |
| ERBB2 |
| ERBB3 |
| ERBB4 |
| ERBIN |
| ERCC1 |

| |
|---|
| ERCC2 |
| ERCC3 |
| ERCC4 |
| ERCC5 |
| ERCC6 |
| ERCC6L |
| ERCC6L2 |
| ERCC8 |
| EREG |
| ERF |
| ERG28 |
| ERGIC1 |
| ERGIC2 |
| ERGIC3 |
| ERH |
| ERI1 |
| ERI2 |
| ERI3 |
| ERICH4 |
| ERLEC1 |
| ERLIN1 |
| ERLIN2 |
| ERMARD |
| ERMP1 |
| ERN1 |
| ERO1A |
| ERO1B |
| ERP29 |
| ERP44 |
| ERRFI1 |
| ESD |
| ESF1 |
| ESPL1 |
| ESPN |
| ESR2 |
| ESRP1 |
| ESRP2 |
| ESRRA |
| ESRRG |
| ESS2 |
| ESYT1 |
| ESYT2 |
| ETAA1 |
| ETF1 |
| ETFA |

| |
|---|
| ETFB |
| ETFBKMT |
| ETFDH |
| ETFRF1 |
| ETHE1 |
| ETNK1 |
| ETNK2 |
| ETS1 |
| ETS2 |
| ETV2 |
| ETV3 |
| ETV6 |
| EVA1B |
| EVC |
| EVI5 |
| EVI5L |
| EVL |
| EVPL |
| EWSR1 |
| EXD2 |
| EXD3 |
| EXO1 |
| EXO5 |
| EXOC1 |
| EXOC2 |
| EXOC3 |
| EXOC3L1 |
| EXOC4 |
| EXOC5 |
| EXOC6 |
| EXOC6B |
| EXOC7 |
| EXOC8 |
| EXOG |
| EXOSC1 |
| EXOSC10 |
| EXOSC2 |
| EXOSC3 |
| EXOSC4 |
| EXOSC5 |
| EXOSC6 |
| EXOSC7 |
| EXOSC8 |
| EXOSC9 |
| EXPH5 |

| |
|---|
| EXT1 |
| EXT2 |
| EXTL1 |
| EXTL2 |
| EXTL3 |
| EYA3 |
| EZH1 |
| EZH2 |
| F11R |
| F12 |
| F2RL1 |
| F3 |
| F5 |
| FA2H |
| FAAH |
| FAAP24 |
| FABP3 |
| FABP4 |
| FABP5 |
| FABP7 |
| FADD |
| FADS1 |
| FADS2 |
| FADS3 |
| FAF1 |
| FAF2 |
| FAH |
| FAHD1 |
| FAHD2A |
| FAIM |
| FAM104A |
| FAM110A |
| FAM110B |
| FAM110C |
| FAM111A |
| FAM111B |
| FAM114A1 |
| FAM114A2 |
| FAM117A |
| FAM117B |
| FAM118A |
| FAM118B |
| FAM120A |
| FAM120B |
| FAM120C |

| |
|---|
| FAM126B |
| FAM131A |
| FAM131C |
| FAM133A |
| FAM133B |
| FAM135A |
| FAM13A |
| FAM13B |
| FAM13C |
| FAM149B1 |
| FAM151B |
| FAM155A |
| FAM156A |
| FAM161A |
| FAM161B |
| FAM162A |
| FAM168A |
| FAM168B |
| FAM169A |
| FAM171A1 |
| FAM171A2 |
| FAM171B |
| FAM174A |
| FAM174C |
| FAM177A1 |
| FAM178B |
| FAM184B |
| FAM185A |
| FAM189A1 |
| FAM189B |
| FAM193A |
| FAM193B |
| FAM199X |
| FAM200A |
| FAM200B |
| FAM20A |
| FAM20B |
| FAM20C |
| FAM210A |
| FAM210B |
| FAM214A |
| FAM214B |
| FAM217A |
| FAM217B |
| FAM219A |

| |
|---|
| FAM219B |
| FAM220A |
| FAM222B |
| FAM229A |
| FAM229B |
| FAM234B |
| FAM241A |
| FAM241B |
| FAM32A |
| FAM43A |
| FAM50A |
| FAM50B |
| FAM53B |
| FAM53C |
| FAM71E1 |
| FAM72A |
| FAM76A |
| FAM76B |
| FAM78A |
| FAM81A |
| FAM83D |
| FAM83E |
| FAM83G |
| FAM83H |
| FAM89B |
| FAM8A1 |
| FAM91A1 |
| FAM98A |
| FAM98B |
| FAM98C |
| FAN1 |
| FANCA |
| FANCB |
| FANCC |
| FANCD2 |
| FANCE |
| FANCF |
| FANCG |
| FANCI |
| FANCL |
| FANCM |
| FAR1 |
| FAR2 |
| FARP2 |
| FARS2 |

| |
|---|
| FARSA |
| FARSB |
| FAS |
| FASTK |
| FASTKD1 |
| FASTKD2 |
| FASTKD3 |
| FASTKD5 |
| FAT1 |
| FAT2 |
| FAT4 |
| FAU |
| FAXDC2 |
| FBF1 |
| FBH1 |
| FBLIM1 |
| FBLN1 |
| FBLN2 |
| FBLN5 |
| FBN1 |
| FBP1 |
| FBP2 |
| FBRSL1 |
| FBXL12 |
| FBXL14 |
| FBXL15 |
| FBXL17 |
| FBXL18 |
| FBXL19 |
| FBXL2 |
| FBXL20 |
| FBXL22 |
| FBXL4 |
| FBXL5 |
| FBXL7 |
| FBXL8 |
| FBXO10 |
| FBXO11 |
| FBXO15 |
| FBXO17 |
| FBXO21 |
| FBXO22 |
| FBXO24 |
| FBXO25 |
| FBXO28 |

| |
|---|
| FBXO3 |
| FBXO31 |
| FBXO32 |
| FBXO33 |
| FBXO34 |
| FBXO38 |
| FBXO4 |
| FBXO42 |
| FBXO43 |
| FBXO45 |
| FBXO46 |
| FBXO5 |
| FBXO6 |
| FBXO7 |
| FBXO8 |
| FBXO9 |
| FBXW11 |
| FBXW12 |
| FBXW2 |
| FBXW4 |
| FBXW5 |
| FBXW8 |
| FBXW9 |
| FCF1 |
| FCGBP |
| FCGR1B |
| FCGRT |
| FCHO1 |
| FCHO2 |
| FCHSD1 |
| FCHSD2 |
| FCSK |
| FDFT1 |
| FDPS |
| FDX1 |
| FDX2 |
| FDXACB1 |
| FDXR |
| FECH |
| FEM1A |
| FEM1B |
| FEM1C |
| FEN1 |
| FER |
| FERMT2 |

| |
|---|
| FES |
| FEZ2 |
| FFAR4 |
| FGD1 |
| FGD2 |
| FGD3 |
| FGD4 |
| FGD5 |
| FGD6 |
| FGF1 |
| FGF13 |
| FGF17 |
| FGF2 |
| FGFBP1 |
| FGFR1OP2 |
| FGFR2 |
| FGFR3 |
| FGGY |
| FGL2 |
| FH |
| FHDC1 |
| FHL2 |
| FHL3 |
| FHOD1 |
| FHOD3 |
| FIBP |
| FICD |
| FIG4 |
| FIGN |
| FIGNL1 |
| FILIP1 |
| FILIP1L |
| FIP1L1 |
| FITM2 |
| FIZ1 |
| FJX1 |
| FKBP10 |
| FKBP11 |
| FKBP15 |
| FKBP3 |
| FKBP5 |
| FKBP7 |
| FKBP8 |
| FKBP9 |
| FKBPL |

| |
|---|
| FKRP |
| FKTN |
| FLAD1 |
| FLCN |
| FLII |
| FLNA |
| FLNB |
| FLOT1 |
| FLOT2 |
| FLRT1 |
| FLVCR1 |
| FLVCR2 |
| FLYWCH1 |
| FLYWCH2 |
| FMN1 |
| FMN2 |
| FMNL1 |
| FMNL2 |
| FMNL3 |
| FMO5 |
| FMR1 |
| FN3K |
| FN3KRP |
| FNBP1 |
| FNBP1L |
| FNBP4 |
| FNDC1 |
| FNDC10 |
| FNDC3A |
| FNDC3B |
| FNDC4 |
| FNDC9 |
| FNIP1 |
| FNIP2 |
| FNTA |
| FNTB |
| FOCAD |
| FOLR1 |
| FOS |
| FOSB |
| FOSL2 |
| FOXA1 |
| FOXA3 |
| FOXC1 |
| FOXD2 |

| |
|---|
| FOXD4L1 |
| FOXG1 |
| FOXH1 |
| FOXJ3 |
| FOXK1 |
| FOXK2 |
| FOXL2 |
| FOXM1 |
| FOXN2 |
| FOXO1 |
| FOXO3 |
| FOXO4 |
| FOXP1 |
| FOXP4 |
| FOXRED1 |
| FOXRED2 |
| FOXS1 |
| FPGS |
| FPGT |
| FPR2 |
| FRA10AC1 |
| FRAS1 |
| FRAT1 |
| FRAT2 |
| FREM2 |
| FRG1 |
| FRK |
| FRMD4B |
| FRMD6 |
| FRMD8 |
| FRMPD1 |
| FRRS1 |
| FRRS1L |
| FRS2 |
| FRS3 |
| FRY |
| FRYL |
| FSCN1 |
| FSD1L |
| FSD2 |
| FSIP2 |
| FST |
| FSTL1 |
| FSTL3 |
| FTCD |

| |
|---|
| FTH1 |
| FTL |
| FTO |
| FTSJ1 |
| FTSJ3 |
| FUBP1 |
| FUBP3 |
| FUCA1 |
| FUCA2 |
| FUNDC2 |
| FUOM |
| FURIN |
| FUS |
| FUT1 |
| FUT2 |
| FUZ |
| FXR1 |
| FXR2 |
| FXYD3 |
| FXYD5 |
| FYB1 |
| FYCO1 |
| FYTTD1 |
| FZD1 |
| FZD2 |
| FZD3 |
| FZD4 |
| FZD5 |
| FZD6 |
| FZD7 |
| FZD8 |
| FZD9 |
| FZR1 |
| G0S2 |
| G2E3 |
| G3BP1 |
| G3BP2 |
| G6PC3 |
| G6PD |
| GAA |
| GAB1 |
| GAB2 |
| GABARAP |
| GABBR1 |
| GABPA |

| |
|---|
| GABPB1 |
| GABPB2 |
| GABRA2 |
| GABRA3 |
| GABRG1 |
| GABRR2 |
| GADD45A |
| GADD45B |
| GADD45GIP1 |
| GADL1 |
| GAK |
| GAL3ST1 |
| GAL3ST2 |
| GAL3ST3 |
| GAL3ST4 |
| GALC |
| GALE |
| GALK1 |
| GALK2 |
| GALM |
| GALNS |
| GALNT1 |
| GALNT10 |
| GALNT11 |
| GALNT12 |
| GALNT13 |
| GALNT16 |
| GALNT18 |
| GALNT2 |
| GALNT4 |
| GALNT5 |
| GALR3 |
| GALT |
| GAMT |
| GAN |
| GANAB |
| GANC |
| GAPDH |
| GAPDHS |
| GAR1 |
| GAREM1 |
| GARNL3 |
| GARRE1 |
| GARS1 |
| GAS1 |

| |
|---|
| GAS2L1 |
| GAS6 |
| GAS7 |
| GAS8 |
| GASK1B |
| GATA6 |
| GATAD2A |
| GATAD2B |
| GATB |
| GATC |
| GATD1 |
| GATD3B |
| GBA |
| GBA2 |
| GBA3 |
| GBAP1 |
| GBE1 |
| GBP6 |
| GCA |
| GCAT |
| GCC1 |
| GCC2 |
| GCDH |
| GCFC2 |
| GCH1 |
| GCHFR |
| GCLM |
| GCN1 |
| GCNA |
| GCNT1 |
| GCNT3 |
| GCSH |
| GDA |
| GDAP2 |
| GDE1 |
| GDF9 |
| GDI1 |
| GDI2 |
| GDPD3 |
| GDPD5 |
| GDPGP1 |
| GEM |
| GEMIN2 |
| GEMIN4 |
| GEMIN5 |

| |
|---|
| GEMIN6 |
| GEMIN7 |
| GEMIN8 |
| GEN1 |
| GET1 |
| GET3 |
| GET4 |
| GFER |
| GFM1 |
| GFM2 |
| GFOD2 |
| GFPT1 |
| GFPT2 |
| GFUS |
| GGA1 |
| GGA2 |
| GGA3 |
| GGCT |
| GGCX |
| GGH |
| GGNBP2 |
| GGPS1 |
| GGT1 |
| GGT7 |
| GHDC |
| GHITM |
| GHR |
| GID4 |
| GID8 |
| GIGYF1 |
| GIGYF2 |
| GIMAP1 |
| GIN1 |
| GINM1 |
| GINS1 |
| GINS2 |
| GINS3 |
| GINS4 |
| GIPC1 |
| GIT1 |
| GIT2 |
| GJA1 |
| GJA5 |
| GJA9 |
| GJB3 |

| |
|---|
| GJC1 |
| GK |
| GK5 |
| GKAP1 |
| GLA |
| GLB1 |
| GLB1L |
| GLCCI1 |
| GLDC |
| GLDN |
| GLG1 |
| GLI4 |
| GLIPR1L2 |
| GLIPR2 |
| GLIS2 |
| GLIS3 |
| GLMN |
| GLO1 |
| GLOD4 |
| GLRX |
| GLRX2 |
| GLRX3 |
| GLRX5 |
| GLS |
| GLS2 |
| GLT8D1 |
| GLT8D2 |
| GLUD1 |
| GLUL |
| GLYCTK |
| GLYR1 |
| GM2A |
| GMCL1 |
| GMDS |
| GMEB1 |
| GMEB2 |
| GMFB |
| GMFG |
| GMIP |
| GMNN |
| GMPPA |
| GMPR |
| GMPR2 |
| GMPS |
| GNA11 |

| |
|---|
| GNA12 |
| GNA13 |
| GNA14 |
| GNAI1 |
| GNAI2 |
| GNAL |
| GNAQ |
| GNAS |
| GNAZ |
| GNB1 |
| GNB1L |
| GNB2 |
| GNB4 |
| GNB5 |
| GNE |
| GNG10 |
| GNG11 |
| GNG12 |
| GNG3 |
| GNG5 |
| GNG7 |
| GNL1 |
| GNL2 |
| GNL3 |
| GNPAT |
| GNPDA1 |
| GNPDA2 |
| GNPNAT1 |
| GNPTAB |
| GNPTG |
| GNS |
| GOLGA2 |
| GOLGA3 |
| GOLGA5 |
| GOLGA7 |
| GOLGB1 |
| GOLIM4 |
| GOLM1 |
| GOLPH3 |
| GOLPH3L |
| GOLT1B |
| GON4L |
| GON7 |
| GOPC |
| GORAB |

| |
|---|
| GORASP2 |
| GOSR1 |
| GOSR2 |
| GOT1 |
| GOT2 |
| GP1BA |
| GPAA1 |
| GPALPP1 |
| GPAM |
| GPANK1 |
| GPAT4 |
| GPATCH1 |
| GPATCH11 |
| GPATCH2 |
| GPATCH2L |
| GPATCH3 |
| GPATCH4 |
| GPATCH8 |
| GPBAR1 |
| GPBP1 |
| GPBP1L1 |
| GPC1 |
| GPC2 |
| GPC4 |
| GPC6 |
| GPCPD1 |
| GPD1 |
| GPD1L |
| GPD2 |
| GPER1 |
| GPHN |
| GPI |
| GPKOW |
| GPM6A |
| GPN1 |
| GPN2 |
| GPN3 |
| GPNMB |
| GPR101 |
| GPR108 |
| GPR135 |
| GPR137 |
| GPR137B |
| GPR141 |
| GPR143 |

| |
|---|
| GPR146 |
| GPR15 |
| GPR153 |
| GPR155 |
| GPR157 |
| GPR162 |
| GPR173 |
| GPR176 |
| GPR179 |
| GPR18 |
| GPR27 |
| GPR3 |
| GPR34 |
| GPR35 |
| GPR4 |
| GPR63 |
| GPR65 |
| GPR68 |
| GPR75 |
| GPR82 |
| GPR84 |
| GPR89A |
| GPRASP1 |
| GPRC5A |
| GPRC5B |
| GPRC5C |
| GPRC5D |
| GPRIN3 |
| GPS1 |
| GPSM1 |
| GPSM2 |
| GPT |
| GPT2 |
| GPX1 |
| GPX3 |
| GPX7 |
| GPX8 |
| GRAMD1A |
| GRAMD1C |
| GRAMD2B |
| GRB2 |
| GRB7 |
| GREB1 |
| GREM1 |
| GRHL1 |

| |
|---|
| GRHPR |
| GRID1 |
| GRID2 |
| GRIK5 |
| GRIN2C |
| GRIN3A |
| GRIN3B |
| GRINA |
| GRIP1 |
| GRIPAP1 |
| GRK2 |
| GRK3 |
| GRK4 |
| GRK5 |
| GRK6 |
| GRM3 |
| GRN |
| GRPEL1 |
| GRPEL2 |
| GRSF1 |
| GRTP1 |
| GRWD1 |
| GSAP |
| GSDMB |
| GSE1 |
| GSG1 |
| GSK3A |
| GSK3B |
| GSKIP |
| GSPT1 |
| GSS |
| GSTA4 |
| GSTCD |
| GSTK1 |
| GSTM2 |
| GSTM3 |
| GSTO1 |
| GSTP1 |
| GSTZ1 |
| GTDC1 |
| GTF2A1 |
| GTF2A2 |
| GTF2B |
| GTF2E1 |
| GTF2E2 |

| |
|---|
| GTF2F1 |
| GTF2F2 |
| GTF2H1 |
| GTF2H2 |
| GTF2H3 |
| GTF2I |
| GTF2IRD1 |
| GTF3A |
| GTF3C1 |
| GTF3C2 |
| GTF3C3 |
| GTF3C4 |
| GTF3C5 |
| GTPBP1 |
| GTPBP10 |
| GTPBP2 |
| GTPBP3 |
| GTPBP4 |
| GTPBP6 |
| GTPBP8 |
| GUCA2B |
| GUCD1 |
| GUCY1A1 |
| GUCY1B1 |
| GUF1 |
| GUK1 |
| GULP1 |
| GUSB |
| GXYLT1 |
| GYG1 |
| GYG2 |
| GYS1 |
| GZF1 |
| H1-0 |
| H1-1 |
| H1-10 |
| H1-4 |
| H2AC11 |
| H2AC12 |
| H2AC19 |
| H2AC20 |
| H2AC21 |
| H2AC4 |
| H2AC6 |
| H2AC7 |

| |
|---|
| H2AC8 |
| H2AJ |
| H2AP |
| H2AX |
| H2AZ1 |
| H2AZ2 |
| H2BC11 |
| H2BC12 |
| H2BC13 |
| H2BC14 |
| H2BC15 |
| H2BC18 |
| H2BC21 |
| H2BC3 |
| H2BC9 |
| H3-3A |
| H3C1 |
| H4C1 |
| H6PD |
| HABP4 |
| HACD1 |
| HACD2 |
| HACD3 |
| HACD4 |
| HACE1 |
| HACL1 |
| HADH |
| HADHA |
| HAGH |
| HAGHL |
| HAPLN1 |
| HAPLN3 |
| HARBI1 |
| HARS1 |
| HARS2 |
| HAS3 |
| HAT1 |
| HAUS1 |
| HAUS2 |
| HAUS3 |
| HAUS4 |
| HAUS5 |
| HAUS6 |
| HAUS7 |
| HAUS8 |

| |
|---|
| HAVCR2 |
| HAX1 |
| HBE1 |
| HBEGF |
| HBP1 |
| HBS1L |
| HCCS |
| HCFC1 |
| HCFC1R1 |
| HCFC2 |
| HCN1 |
| HCRT |
| HDAC1 |
| HDAC10 |
| HDAC11 |
| HDAC2 |
| HDAC3 |
| HDAC4 |
| HDAC5 |
| HDAC6 |
| HDAC7 |
| HDAC8 |
| HDAC9 |
| HDC |
| HDDC2 |
| HDDC3 |
| HDGF |
| HDGFL2 |
| HDGFL3 |
| HDHD2 |
| HDHD3 |
| HDHD5 |
| HDLBP |
| HEATR1 |
| HEATR3 |
| HEATR4 |
| HEATR5B |
| HEATR6 |
| HEBP1 |
| HEBP2 |
| HECTD1 |
| HECTD2 |
| HECTD3 |
| HECTD4 |
| HEG1 |

| |
|---|
| HELB |
| HELLS |
| HELQ |
| HELZ |
| HELZ2 |
| HEMK1 |
| HEPH |
| HERC1 |
| HERC2 |
| HERC4 |
| HERC5 |
| HERPUD2 |
| HES4 |
| HES6 |
| HEXA |
| HEXB |
| HEXD |
| HEXIM1 |
| HEXIM2 |
| HEY2 |
| HFE |
| HFM1 |
| HGF |
| HGH1 |
| HGS |
| HHAT |
| HHEX |
| HHIPL1 |
| HHIPL2 |
| HIBADH |
| HIC1 |
| HIC2 |
| HID1 |
| HIF1A |
| HIF1AN |
| HIGD1A |
| HIGD2A |
| HIKESHI |
| HILPDA |
| HINFP |
| HINT1 |
| HINT2 |
| HINT3 |
| HIP1R |
| HIPK1 |

| |
|---|
| HIPK3 |
| HIRA |
| HIRIP3 |
| HIVEP1 |
| HIVEP2 |
| HIVEP3 |
| HJURP |
| HK1 |
| HK2 |
| HK3 |
| HLA-A |
| HLA-DMB |
| HLA-DQB1 |
| HLCS |
| HLX |
| HM13 |
| HMBS |
| HMCN1 |
| HMG20A |
| HMG20B |
| HMGA1 |
| HMGB1 |
| HMGB2 |
| HMGB3 |
| HMGCL |
| HMGCR |
| HMGCS1 |
| HMGN1 |
| HMGN2 |
| HMGN3 |
| HMGN4 |
| HMGN5 |
| HMGXB4 |
| HMMR |
| HMOX1 |
| HMOX2 |
| HNRNPA0 |
| HNRNPA1 |
| HNRNPA3 |
| HNRNPAB |
| HNRNPC |
| HNRNPD |
| HNRNPDL |
| HNRNPF |
| HNRNPH1 |

| |
|---|
| HNRNPH2 |
| HNRNPH3 |
| HNRNPK |
| HNRNPL |
| HNRNPM |
| HNRNPR |
| HNRNPU |
| HNRNPUL1 |
| HNRNPUL2 |
| HOMER1 |
| HOMER2 |
| HOMER3 |
| HOMEZ |
| HOOK1 |
| HOOK2 |
| HOOK3 |
| HOXA10 |
| HOXA13 |
| HOXB13 |
| HOXC10 |
| HOXC4 |
| HOXC5 |
| HOXC6 |
| HOXC8 |
| HOXC9 |
| HOXD8 |
| HP1BP3 |
| HPDL |
| HPF1 |
| HPS1 |
| HPS3 |
| HPS4 |
| HPS6 |
| HPSE |
| HRAS |
| HRH1 |
| HROB |
| HS1BP3 |
| HS2ST1 |
| HS3ST1 |
| HS3ST2 |
| HS3ST3A1 |
| HS6ST1 |
| HS6ST3 |
| HSD11B1L |

| |
|---|
| HSD11B2 |
| HSD17B1 |
| HSD17B10 |
| HSD17B12 |
| HSD17B14 |
| HSD17B3 |
| HSD17B4 |
| HSD17B7 |
| HSD17B8 |
| HSD3B2 |
| HSD3B7 |
| HSDL1 |
| HSDL2 |
| HSF1 |
| HSF2 |
| HSF2BP |
| HSF4 |
| HSFX1 |
| HSH2D |
| HSP90AA1 |
| HSP90AB1 |
| HSP90B1 |
| HSPA12B |
| HSPA13 |
| HSPA14 |
| HSPA1A |
| HSPA1L |
| HSPA4L |
| HSPA5 |
| HSPA6 |
| HSPA8 |
| HSPA9 |
| HSPB1 |
| HSPB11 |
| HSPB6 |
| HSPBAP1 |
| HSPBP1 |
| HSPD1 |
| HSPE1 |
| HSPG2 |
| HTATIP2 |
| HTATSF1 |
| HTN3 |
| HTR1F |
| HTR7 |

| |
|---|
| HTRA1 |
| HTRA2 |
| HUS1 |
| HUWE1 |
| HVCN1 |
| HYAL1 |
| HYAL2 |
| HYI |
| HYKK |
| HYLS1 |
| HYOU1 |
| HYPK |
| IAH1 |
| IARS1 |
| IARS2 |
| IBA57 |
| IBTK |
| ICA1L |
| ICAM1 |
| ICE1 |
| ICE2 |
| ICMT |
| ICOSLG |
| ID1 |
| ID2 |
| ID3 |
| IDE |
| IDH1 |
| IDH2 |
| IDH3A |
| IDH3B |
| IDI1 |
| IDNK |
| IDO1 |
| IDS |
| IDUA |
| IER2 |
| IER3 |
| IER3IP1 |
| IER5 |
| IER5L |
| IFFO2 |
| IFI16 |
| IFI27 |
| IFI27L2 |

| |
|---|
| IFI30 |
| IFI35 |
| IFI6 |
| IFIH1 |
| IFIT1 |
| IFIT2 |
| IFIT5 |
| IFITM3 |
| IFNAR1 |
| IFNAR2 |
| IFNGR1 |
| IFNGR2 |
| IFNLR1 |
| IFRD1 |
| IFRD2 |
| IFT122 |
| IFT172 |
| IFT20 |
| IFT22 |
| IFT27 |
| IFT43 |
| IFT46 |
| IFT52 |
| IFT57 |
| IFT74 |
| IFT80 |
| IFT81 |
| IFT88 |
| IFTAP |
| IGBP1 |
| IGDCC4 |
| IGF1R |
| IGF2 |
| IGF2BP2 |
| IGFBP2 |
| IGFBP3 |
| IGFBP4 |
| IGFBP5 |
| IGFBP6 |
| IGFBP7 |
| IGHMBP2 |
| IGSF11 |
| IGSF6 |
| IGSF8 |
| IGSF9 |

| |
|---|
| IHH |
| IK |
| IKBIP |
| IKBKB |
| IKBKE |
| IKBKG |
| IKZF2 |
| IKZF3 |
| IKZF4 |
| IKZF5 |
| IL10 |
| IL10RA |
| IL10RB |
| IL11RA |
| IL13RA1 |
| IL15 |
| IL16 |
| IL17D |
| IL17RA |
| IL17RC |
| IL17RD |
| IL18 |
| IL18BP |
| IL1A |
| IL1RAP |
| IL1RN |
| IL27RA |
| IL2RB |
| IL32 |
| IL34 |
| IL3RA |
| IL4I1 |
| IL4R |
| IL6ST |
| ILF2 |
| ILF3 |
| ILK |
| ILKAP |
| ILRUN |
| ILVBL |
| IMMT |
| IMP3 |
| IMP4 |
| IMPA1 |
| IMPA2 |

| |
|---|
| IMPACT |
| IMPDH1 |
| IMPDH2 |
| INAFM2 |
| INAVA |
| INCENP |
| ING1 |
| ING2 |
| ING3 |
| ING4 |
| ING5 |
| INHA |
| INHBA |
| INHBB |
| INIP |
| INKA1 |
| INKA2 |
| INO80 |
| INO80B |
| INO80C |
| INO80D |
| INO80E |
| INPP1 |
| INPP4A |
| INPP5A |
| INPP5B |
| INPP5D |
| INPP5E |
| INPP5F |
| INPP5K |
| INPPL1 |
| INSIG1 |
| INSIG2 |
| INSR |
| INSYN2A |
| INTS10 |
| INTS11 |
| INTS12 |
| INTS13 |
| INTS14 |
| INTS2 |
| INTS3 |
| INTS5 |
| INTS6 |
| INTS6L |

| |
|---|
| INTS7 |
| INTS8 |
| INTS9 |
| INTU |
| IP6K1 |
| IPMK |
| IPO13 |
| IPO4 |
| IPO5 |
| IPO7 |
| IPO8 |
| IPP |
| IPPK |
| IQCB1 |
| IQCD |
| IQCE |
| IQCG |
| IQGAP1 |
| IQGAP2 |
| IQGAP3 |
| IQSEC1 |
| IQSEC2 |
| IRAK1 |
| IRAK1BP1 |
| IRAK2 |
| IRAK3 |
| IRAK4 |
| IREB2 |
| IRF1 |
| IRF2 |
| IRF2BP1 |
| IRF2BP2 |
| IRF2BPL |
| IRF3 |
| IRF7 |
| IRS1 |
| IRS2 |
| IRX3 |
| IRX5 |
| ISCU |
| ISG20 |
| ISG20L2 |
| ISOC1 |
| IST1 |
| ISY1 |

| |
|---|
| ISYNA1 |
| ITFG1 |
| ITFG2 |
| ITGA3 |
| ITGA4 |
| ITGA6 |
| ITGAE |
| ITGAL |
| ITGAV |
| ITGAX |
| ITGB1 |
| ITGB1BP1 |
| ITGB1BP2 |
| ITGB3 |
| ITGB3BP |
| ITGB5 |
| ITGB7 |
| ITGB8 |
| ITGBL1 |
| ITM2A |
| ITM2B |
| ITM2C |
| ITPA |
| ITPK1 |
| ITPKA |
| ITPKB |
| ITPKC |
| ITPR2 |
| ITPR3 |
| ITPRID2 |
| ITPRIP |
| ITPRIPL2 |
| ITSN1 |
| ITSN2 |
| IVD |
| IVNS1ABP |
| IWS1 |
| JADE1 |
| JADE2 |
| JADE3 |
| JAG1 |
| JAGN1 |
| JAK1 |
| JAK3 |
| JAKMIP2 |

| |
|---|
| JAM3 |
| JARID2 |
| JAZF1 |
| JCAD |
| JCHAIN |
| JDP2 |
| JHY |
| JKAMP |
| JMJD1C |
| JMJD4 |
| JMJD6 |
| JMJD7 |
| JMJD8 |
| JMY |
| JOSD1 |
| JOSD2 |
| JPT1 |
| JPT2 |
| JRK |
| JRKL |
| JSRP1 |
| JTB |
| JUN |
| JUNB |
| JUND |
| JUP |
| KALRN |
| KANK1 |
| KANK2 |
| KANK3 |
| KANSL1 |
| KANSL1L |
| KANSL3 |
| KARS1 |
| KASH5 |
| KAT2A |
| KAT2B |
| KAT5 |
| KAT6A |
| KAT6B |
| KAT8 |
| KATNA1 |
| KATNAL1 |
| KATNB1 |
| KATNBL1 |

| |
|---|
| KATNIP |
| KAZALD1 |
| KAZN |
| KBTBD11 |
| KBTBD2 |
| KBTBD3 |
| KBTBD6 |
| KBTBD7 |
| KCMF1 |
| KCNA4 |
| KCNAB1 |
| KCNAB2 |
| KCNC2 |
| KCNC3 |
| KCND2 |
| KCNE2 |
| KCNE3 |
| KCNE4 |
| KCNE5 |
| KCNG2 |
| KCNH2 |
| KCNIP4 |
| KCNJ10 |
| KCNJ14 |
| KCNJ15 |
| KCNJ2 |
| KCNJ5 |
| KCNK1 |
| KCNK3 |
| KCNK5 |
| KCNN3 |
| KCNQ1 |
| KCNQ4 |
| KCNQ5 |
| KCNRG |
| KCNS3 |
| KCNT2 |
| KCTD1 |
| KCTD10 |
| KCTD11 |
| KCTD12 |
| KCTD14 |
| KCTD15 |
| KCTD18 |
| KCTD20 |

| |
|---|
| KCTD21 |
| KCTD3 |
| KCTD5 |
| KCTD7 |
| KCTD9 |
| KDELR1 |
| KDELR2 |
| KDELR3 |
| KDF1 |
| KDM1A |
| KDM1B |
| KDM2A |
| KDM2B |
| KDM3A |
| KDM3B |
| KDM4A |
| KDM4B |
| KDM4C |
| KDM5A |
| KDM5B |
| KDM5C |
| KDM6A |
| KDM7A |
| KDM8 |
| KDSR |
| KEAP1 |
| KHDC4 |
| KHDRBS1 |
| KHNYN |
| KHSRP |
| KIAA0040 |
| KIAA0100 |
| KIAA0319 |
| KIAA0319L |
| KIAA0513 |
| KIAA0586 |
| KIAA0753 |
| KIAA0825 |
| KIAA0895 |
| KIAA0930 |
| KIAA1109 |
| KIAA1143 |
| KIAA1191 |
| KIAA1210 |
| KIAA1217 |

| |
|---|
| KIAA1328 |
| KIAA1522 |
| KIAA1549 |
| KIAA1586 |
| KIAA1614 |
| KIAA1671 |
| KIAA1958 |
| KIAA2013 |
| KIAA2026 |
| KIDINS220 |
| KIF11 |
| KIF13A |
| KIF13B |
| KIF14 |
| KIF15 |
| KIF16B |
| KIF18A |
| KIF18B |
| KIF1B |
| KIF1C |
| KIF20A |
| KIF20B |
| KIF22 |
| KIF23 |
| KIF24 |
| KIF26B |
| KIF2A |
| KIF2C |
| KIF3A |
| KIF3B |
| KIF3C |
| KIF4A |
| KIF7 |
| KIFAP3 |
| KIFBP |
| KIFC1 |
| KIFC2 |
| KIFC3 |
| KIN |
| KIRREL3 |
| KIT |
| KITLG |
| KLC1 |
| KLC2 |
| KLC3 |

| |
|---|
| KLC4 |
| KLF11 |
| KLF12 |
| KLF13 |
| KLF15 |
| KLF16 |
| KLF2 |
| KLF4 |
| KLF5 |
| KLF6 |
| KLF7 |
| KLF8 |
| KLHDC1 |
| KLHDC10 |
| KLHDC2 |
| KLHDC4 |
| KLHDC8B |
| KLHL11 |
| KLHL12 |
| KLHL15 |
| KLHL17 |
| KLHL18 |
| KLHL2 |
| KLHL20 |
| KLHL21 |
| KLHL23 |
| KLHL24 |
| KLHL25 |
| KLHL26 |
| KLHL28 |
| KLHL3 |
| KLHL36 |
| KLHL41 |
| KLHL42 |
| KLHL5 |
| KLHL7 |
| KLHL8 |
| KLHL9 |
| KLK14 |
| KLK15 |
| KMT2A |
| KMT2B |
| KMT2C |
| KMT2D |
| KMT2E |

| |
|---|
| KMT5A |
| KMT5B |
| KMT5C |
| KNL1 |
| KNOP1 |
| KNSTRN |
| KPNA3 |
| KPNA4 |
| KPNA5 |
| KPNA6 |
| KPNB1 |
| KRAS |
| KRBA1 |
| KRBA2 |
| KRBOX4 |
| KRCC1 |
| KRI1 |
| KRIT1 |
| KRR1 |
| KRT18 |
| KRT19 |
| KRT20 |
| KRT8 |
| KRTAP7-1 |
| KRTCAP2 |
| KSR1 |
| KTI12 |
| KTN1 |
| KXD1 |
| KYAT1 |
| KYAT3 |
| L1CAM |
| L2HGDH |
| L3HYPDH |
| L3MBTL1 |
| L3MBTL2 |
| L3MBTL3 |
| LACTB |
| LACTB2 |
| LAD1 |
| LAG3 |
| LAGE3 |
| LAMA1 |
| LAMA2 |
| LAMA3 |

| |
|---|
| LAMA5 |
| LAMB1 |
| LAMB2 |
| LAMB3 |
| LAMC1 |
| LAMC3 |
| LAMP1 |
| LAMP2 |
| LAMTOR1 |
| LAMTOR2 |
| LAMTOR5 |
| LANCL2 |
| LAP3 |
| LAPTM4A |
| LAPTM4B |
| LAPTM5 |
| LARGE1 |
| LARGE2 |
| LARP1 |
| LARP1B |
| LARP4 |
| LARP4B |
| LARP6 |
| LARP7 |
| LARS1 |
| LARS2 |
| LAS1L |
| LASP1 |
| LAT |
| LATS2 |
| LBH |
| LBR |
| LBX2 |
| LCA5 |
| LCLAT1 |
| LCMT1 |
| LCMT2 |
| LCN2 |
| LCOR |
| LCORL |
| LCP1 |
| LDAH |
| LDB1 |
| LDHA |
| LDHB |

| |
|---|
| LDHD |
| LDLR |
| LDLRAD2 |
| LDLRAD3 |
| LDLRAP1 |
| LDOC1 |
| LEMD2 |
| LEMD3 |
| LENG1 |
| LENG8 |
| LENG9 |
| LEO1 |
| LEPROT |
| LEPROTL1 |
| LETM1 |
| LETM2 |
| LETMD1 |
| LFNG |
| LGALS3 |
| LGALS3BP |
| LGALS8 |
| LGALS9 |
| LGALSL |
| LGI3 |
| LGMN |
| LGR4 |
| LGSN |
| LHFPL2 |
| LHPP |
| LIAS |
| LIF |
| LIFR |
| LIG1 |
| LIG3 |
| LIG4 |
| LILRA5 |
| LILRA6 |
| LIMA1 |
| LIMD1 |
| LIMD2 |
| LIMK1 |
| LIMK2 |
| LIMS1 |
| LIN37 |
| LIN52 |

| |
|---|
| LIN54 |
| LIN7B |
| LIN7C |
| LIN9 |
| LINC02693 |
| LINGO2 |
| LINS1 |
| LIPA |
| LIPC |
| LIPE |
| LIPH |
| LIPK |
| LIPT1 |
| LIPT2 |
| LITAF |
| LIX1L |
| LLGL2 |
| LLPH |
| LMAN1 |
| LMAN2 |
| LMAN2L |
| LMBRD1 |
| LMBRD2 |
| LMCD1 |
| LMF1 |
| LMF2 |
| LMLN |
| LMNA |
| LMNB1 |
| LMNB2 |
| LMNTD1 |
| LMNTD2 |
| LMO4 |
| LMO7 |
| LMTK2 |
| LNPEP |
| LNPK |
| LNX2 |
| LONP1 |
| LONP2 |
| LONRF1 |
| LOXL3 |
| LOXL4 |
| LPAR2 |
| LPAR6 |

| |
|---|
| LPCAT3 |
| LPCAT4 |
| LPGAT1 |
| LPIN1 |
| LPIN3 |
| LPP |
| LPXN |
| LRATD1 |
| LRATD2 |
| LRBA |
| LRCH2 |
| LRCH3 |
| LRCH4 |
| LRFN1 |
| LRFN3 |
| LRFN4 |
| LRIF1 |
| LRIG1 |
| LRIG2 |
| LRIG3 |
| LRMDA |
| LRP1 |
| LRP10 |
| LRP12 |
| LRP4 |
| LRP5 |
| LRP6 |
| LRP8 |
| LRPAP1 |
| LRPPRC |
| LRR1 |
| LRRC1 |
| LRRC10B |
| LRRC14 |
| LRRC17 |
| LRRC20 |
| LRRC24 |
| LRRC25 |
| LRRC26 |
| LRRC27 |
| LRRC28 |
| LRRC31 |
| LRRC37A2 |
| LRRC4 |
| LRRC40 |

| |
|---|
| LRRC41 |
| LRRC42 |
| LRRC45 |
| LRRC46 |
| LRRC47 |
| LRRC49 |
| LRRC4B |
| LRRC4C |
| LRRC57 |
| LRRC58 |
| LRRC59 |
| LRRC66 |
| LRRC74B |
| LRRC75A |
| LRRC8A |
| LRRC8D |
| LRRCC1 |
| LRRFIP1 |
| LRRFIP2 |
| LRRK1 |
| LRRK2 |
| LRRN1 |
| LRRN2 |
| LRRN3 |
| LRRTM2 |
| LRSAM1 |
| LRWD1 |
| LSG1 |
| LSM1 |
| LSM10 |
| LSM11 |
| LSM12 |
| LSM14A |
| LSM14B |
| LSM3 |
| LSM4 |
| LSM5 |
| LSM6 |
| LSM7 |
| LSM8 |
| LSMEM2 |
| LSR |
| LSS |
| LTA4H |
| LTB |

| |
|---|
| LTB4R |
| LTBP1 |
| LTBP3 |
| LTBP4 |
| LTBR |
| LTN1 |
| LTO1 |
| LTV1 |
| LUC7L |
| LUC7L2 |
| LUC7L3 |
| LURAP1 |
| LURAP1L |
| LUZP1 |
| LUZP2 |
| LXN |
| LY6G5B |
| LY75 |
| LY9 |
| LYAR |
| LYN |
| LYNX1 |
| LYPLA1 |
| LYPLA2 |
| LYRM1 |
| LYRM2 |
| LYRM4 |
| LYRM7 |
| LYRM9 |
| LYSMD1 |
| LYSMD2 |
| LYSMD3 |
| LYST |
| LYZ |
| LZIC |
| LZTFL1 |
| LZTR1 |
| LZTS1 |
| LZTS2 |
| LZTS3 |
| M6PR |
| MACIR |
| MACO1 |
| MACROD1 |
| MACROH2A1 |

| |
|---|
| MAD1L1 |
| MAD2L1 |
| MAD2L1BP |
| MAD2L2 |
| MADD |
| MAEA |
| MAF |
| MAF1 |
| MAFB |
| MAFK |
| MAGEA10 |
| MAGED1 |
| MAGED2 |
| MAGED4B |
| MAGEL2 |
| MAGI1 |
| MAGI3 |
| MAGIX |
| MAGOH |
| MAGOHB |
| MAGT1 |
| MAIP1 |
| MAK16 |
| MAL2 |
| MALSU1 |
| MAMDC4 |
| MAML1 |
| MAML2 |
| MAML3 |
| MAMLD1 |
| MAMSTR |
| MAN1A1 |
| MAN1A2 |
| MAN1B1 |
| MAN1C1 |
| MAN2A1 |
| MAN2A2 |
| MAN2B1 |
| MAN2C1 |
| MANBA |
| MANBAL |
| MANEA |
| MANEAL |
| MANF |
| MAOA |

| |
|---|
| MAP10 |
| MAP1B |
| MAP1LC3B |
| MAP2 |
| MAP2K1 |
| MAP2K2 |
| MAP2K3 |
| MAP2K5 |
| MAP2K6 |
| MAP2K7 |
| MAP3K1 |
| MAP3K10 |
| MAP3K11 |
| MAP3K12 |
| MAP3K14 |
| MAP3K15 |
| MAP3K2 |
| MAP3K21 |
| MAP3K3 |
| MAP3K4 |
| MAP3K5 |
| MAP3K6 |
| MAP3K7 |
| MAP3K8 |
| MAP3K9 |
| MAP4 |
| MAP4K1 |
| MAP4K2 |
| MAP4K3 |
| MAP4K4 |
| MAP4K5 |
| MAP7 |
| MAP7D1 |
| MAP7D3 |
| MAP9 |
| MAPK11 |
| MAPK12 |
| MAPK13 |
| MAPK14 |
| MAPK15 |
| MAPK4 |
| MAPK6 |
| MAPK7 |
| MAPK8 |
| MAPK8IP1 |

| |
|---|
| MAPK8IP3 |
| MAPKAP1 |
| MAPKAPK3 |
| MAPKBP1 |
| MAPRE2 |
| MARCHF1 |
| MARCHF2 |
| MARCHF5 |
| MARCHF6 |
| MARCHF7 |
| MARCHF8 |
| MARCHF9 |
| MARCKSL1 |
| MARF1 |
| MARK1 |
| MARK2 |
| MARK3 |
| MARK4 |
| MARS1 |
| MARS2 |
| MARVELD1 |
| MARVELD2 |
| MARVELD3 |
| MASP2 |
| MAST1 |
| MAST2 |
| MAST3 |
| MAST4 |
| MASTL |
| MAT2B |
| MATK |
| MATN4 |
| MATR3 |
| MAU2 |
| MAVS |
| MAX |
| MAZ |
| MB21D2 |
| MBD1 |
| MBD2 |
| MBD3 |
| MBD4 |
| MBD5 |
| MBD6 |
| MBIP |

| |
|---|
| MBLAC1 |
| MBLAC2 |
| MBNL2 |
| MBNL3 |
| MBOAT2 |
| MBTD1 |
| MBTPS1 |
| MBTPS2 |
| MCAM |
| MCAT |
| MCC |
| MCCC1 |
| MCCC2 |
| MCEE |
| MCF2L |
| MCF2L2 |
| MCFD2 |
| MCL1 |
| MCM10 |
| MCM2 |
| MCM3 |
| MCM3AP |
| MCM4 |
| MCM5 |
| MCM6 |
| MCM7 |
| MCM8 |
| MCM9 |
| MCMBP |
| MCMDC2 |
| MCOLN1 |
| MCPH1 |
| MCRIP1 |
| MCRIP2 |
| MCRS1 |
| MCTP1 |
| MCTS1 |
| MCU |
| MCUB |
| MDC1 |
| MDFIC |
| MDH2 |
| MDK |
| MDM1 |
| MDM2 |

| |
|---|
| MDM4 |
| MDN1 |
| MDP1 |
| ME1 |
| ME2 |
| ME3 |
| MEA1 |
| MEAF6 |
| MEAK7 |
| MECOM |
| MECP2 |
| MECR |
| MED1 |
| MED10 |
| MED11 |
| MED12 |
| MED12L |
| MED13 |
| MED13L |
| MED14 |
| MED15 |
| MED16 |
| MED17 |
| MED18 |
| MED19 |
| MED21 |
| MED22 |
| MED23 |
| MED24 |
| MED25 |
| MED26 |
| MED27 |
| MED28 |
| MED29 |
| MED30 |
| MED31 |
| MED4 |
| MED7 |
| MED9 |
| MEF2A |
| MEF2C |
| MEF2D |
| MEFV |
| MEGF8 |
| MEGF9 |

| |
|---|
| MEI1 |
| MEIOB |
| MEIS1 |
| MEIS2 |
| MELK |
| MEN1 |
| MEP1B |
| MEPCE |
| MERTK |
| MESD |
| MESP1 |
| MESP2 |
| MEST |
| METAP1D |
| METRN |
| METRNL |
| METTL1 |
| METTL14 |
| METTL15 |
| METTL16 |
| METTL17 |
| METTL18 |
| METTL21A |
| METTL23 |
| METTL26 |
| METTL27 |
| METTL2A |
| METTL3 |
| METTL4 |
| METTL5 |
| METTL6 |
| METTL7A |
| METTL7B |
| METTL8 |
| METTL9 |
| MEX3A |
| MEX3B |
| MEX3C |
| MEX3D |
| MFAP1 |
| MFAP2 |
| MFAP3 |
| MFAP3L |
| MFGE8 |
| MFHAS1 |

| |
|---|
| MFN1 |
| MFSD1 |
| MFSD10 |
| MFSD11 |
| MFSD12 |
| MFSD13A |
| MFSD14A |
| MFSD14B |
| MFSD3 |
| MFSD4B |
| MFSD6 |
| MFSD8 |
| MFSD9 |
| MGA |
| MGAM |
| MGARP |
| MGAT1 |
| MGAT2 |
| MGAT4A |
| MGAT4B |
| MGAT5 |
| MGME1 |
| MGMT |
| MGRN1 |
| MGST1 |
| MGST2 |
| MGST3 |
| MIA2 |
| MIA3 |
| MIB1 |
| MIB2 |
| MICA |
| MICAL1 |
| MICAL3 |
| MICALL1 |
| MICALL2 |
| MICB |
| MICOS10 |
| MICOS13 |
| MICU1 |
| MICU2 |
| MICU3 |
| MID1 |
| MID1IP1 |
| MIDEAS |

| |
|---|
| MIDN |
| MIEF1 |
| MIEF2 |
| MIEN1 |
| MIER1 |
| MIER2 |
| MIER3 |
| MIF |
| MIF4GD |
| MIGA1 |
| MIGA2 |
| MIIP |
| MINAR1 |
| MINDY1 |
| MINDY2 |
| MINDY3 |
| MINDY4 |
| MINK1 |
| MINPP1 |
| MIOS |
| MIPEP |
| MIPOL1 |
| MIS12 |
| MIS18A |
| MIS18BP1 |
| MITD1 |
| MITF |
| MIX23 |
| MKI67 |
| MKKS |
| MKLN1 |
| MKNK1 |
| MKNK2 |
| MKRN2 |
| MKS1 |
| MKX |
| MLEC |
| MLF1 |
| MLH1 |
| MLH3 |
| MLIP |
| MLKL |
| MLLT1 |
| MLLT3 |
| MLLT6 |

| |
|---|
| MLPH |
| MLST8 |
| MLXIP |
| MLYCD |
| MMAA |
| MMAB |
| MMACHC |
| MMADHC |
| MMD |
| MMGT1 |
| MMP14 |
| MMP15 |
| MMP17 |
| MMP19 |
| MMP2 |
| MMP25 |
| MMP9 |
| MMS19 |
| MMS22L |
| MMUT |
| MN1 |
| MNAT1 |
| MND1 |
| MNS1 |
| MNT |
| MOAP1 |
| MOB1A |
| MOB1B |
| MOB3A |
| MOB3C |
| MOB4 |
| MOCOS |
| MOCS1 |
| MOCS2 |
| MOCS3 |
| MOGS |
| MON1A |
| MON1B |
| MORC2 |
| MORC3 |
| MORF4L1 |
| MORF4L2 |
| MORN4 |
| MOSMO |
| MOSPD1 |

| |
|---|
| MOSPD3 |
| MOV10 |
| MPC1 |
| MPDU1 |
| MPDZ |
| MPG |
| MPHOSPH10 |
| MPHOSPH6 |
| MPHOSPH8 |
| MPHOSPH9 |
| MPI |
| MPL |
| MPLKIP |
| MPND |
| MPP1 |
| MPP2 |
| MPP7 |
| MPPE1 |
| MPPED2 |
| MPRIP |
| MPST |
| MPV17L2 |
| MPZL3 |
| MR1 |
| MRAP |
| MRAP2 |
| MRAS |
| MRC2 |
| MRFAP1 |
| MRFAP1L1 |
| MRGBP |
| MRGPRX3 |
| MRI1 |
| MRM1 |
| MRM2 |
| MRM3 |
| MRNIP |
| MRO |
| MROH1 |
| MRPL1 |
| MRPL10 |
| MRPL11 |
| MRPL12 |
| MRPL13 |
| MRPL14 |

| |
|---|
| MRPL15 |
| MRPL16 |
| MRPL17 |
| MRPL18 |
| MRPL19 |
| MRPL20 |
| MRPL21 |
| MRPL22 |
| MRPL23 |
| MRPL24 |
| MRPL27 |
| MRPL28 |
| MRPL3 |
| MRPL30 |
| MRPL32 |
| MRPL33 |
| MRPL34 |
| MRPL35 |
| MRPL36 |
| MRPL37 |
| MRPL38 |
| MRPL39 |
| MRPL4 |
| MRPL40 |
| MRPL41 |
| MRPL42 |
| MRPL43 |
| MRPL44 |
| MRPL45 |
| MRPL46 |
| MRPL47 |
| MRPL48 |
| MRPL49 |
| MRPL51 |
| MRPL52 |
| MRPL53 |
| MRPL54 |
| MRPL55 |
| MRPL57 |
| MRPL58 |
| MRPL9 |
| MRPS10 |
| MRPS11 |
| MRPS15 |
| MRPS16 |

| |
|---|
| MRPS17 |
| MRPS18B |
| MRPS18C |
| MRPS2 |
| MRPS21 |
| MRPS22 |
| MRPS23 |
| MRPS24 |
| MRPS25 |
| MRPS26 |
| MRPS27 |
| MRPS28 |
| MRPS30 |
| MRPS31 |
| MRPS33 |
| MRPS34 |
| MRPS35 |
| MRPS5 |
| MRPS6 |
| MRPS7 |
| MRPS9 |
| MRRF |
| MRS2 |
| MRTFA |
| MRTFB |
| MSANTD2 |
| MSANTD4 |
| MSH2 |
| MSH3 |
| MSH4 |
| MSH5 |
| MSH6 |
| MSI1 |
| MSI2 |
| MSL1 |
| MSL2 |
| MSL3 |
| MSMO1 |
| MSN |
| MSR1 |
| MSRA |
| MSRB2 |
| MSRB3 |
| MST1 |
| MST1R |

| |
|---|
| MSTO1 |
| MT-ND4L |
| MT1A |
| MT2A |
| MT3 |
| MTA1 |
| MTA2 |
| MTA3 |
| MTAP |
| MTARC1 |
| MTBP |
| MTCH1 |
| MTCH2 |
| MTCL1 |
| MTCP1 |
| MTDH |
| MTERF1 |
| MTERF2 |
| MTERF3 |
| MTF1 |
| MTF2 |
| MTFMT |
| MTFP1 |
| MTFR1 |
| MTFR1L |
| MTFR2 |
| MTG1 |
| MTG2 |
| MTHFD1L |
| MTHFD2 |
| MTHFD2L |
| MTHFSD |
| MTIF2 |
| MTIF3 |
| MTLN |
| MTM1 |
| MTMR1 |
| MTMR10 |
| MTMR11 |
| MTMR12 |
| MTMR14 |
| MTMR2 |
| MTMR3 |
| MTMR4 |
| MTMR6 |

| |
|---|
| MTMR7 |
| MTMR9 |
| MTO1 |
| MTOR |
| MTPAP |
| MTPN |
| MTR |
| MTRES1 |
| MTREX |
| MTRF1 |
| MTRF1L |
| MTRR |
| MTSS1 |
| MTSS2 |
| MTTP |
| MTURN |
| MTUS1 |
| MTX1 |
| MTX2 |
| MTX3 |
| MUC1 |
| MUC20 |
| MUL1 |
| MUS81 |
| MVB12A |
| MVB12B |
| MVD |
| MVK |
| MVP |
| MX1 |
| MXD3 |
| MXD4 |
| MXI1 |
| MXRA7 |
| MXRA8 |
| MYADM |
| MYBBP1A |
| MYBL1 |
| MYBL2 |
| MYBPC1 |
| MYC |
| MYCBP |
| MYCBP2 |
| MYCBPAP |
| MYCN |

| |
|---|
| MYD88 |
| MYDGF |
| MYEF2 |
| MYG1 |
| MYH10 |
| MYH11 |
| MYH14 |
| MYH9 |
| MYL12B |
| MYL6 |
| MYL6B |
| MYL9 |
| MYLIP |
| MYLK |
| MYLPF |
| MYO10 |
| MYO15B |
| MYO18A |
| MYO19 |
| MYO1A |
| MYO1B |
| MYO1C |
| MYO1D |
| MYO1E |
| MYO1F |
| MYO1G |
| MYO3B |
| MYO5A |
| MYO5B |
| MYO5C |
| MYO6 |
| MYO7A |
| MYO9A |
| MYO9B |
| MYOC |
| MYOCD |
| MYOF |
| MYOM1 |
| MYORG |
| MYOZ1 |
| MYPN |
| MYPOP |
| MYRF |
| MYSM1 |
| MZF1 |

| |
|---|
| MZT1 |
| N4BP1 |
| N4BP2 |
| N4BP2L1 |
| N4BP3 |
| N6AMT1 |
| NAA10 |
| NAA15 |
| NAA16 |
| NAA20 |
| NAA25 |
| NAA30 |
| NAA35 |
| NAA38 |
| NAA40 |
| NAA60 |
| NAAA |
| NAALADL1 |
| NAALADL2 |
| NAB1 |
| NAB2 |
| NABP1 |
| NACA |
| NACAD |
| NACC1 |
| NACC2 |
| NADK |
| NADK2 |
| NAE1 |
| NAF1 |
| NAGA |
| NAGK |
| NAGLU |
| NAGPA |
| NAIF1 |
| NAIP |
| NAMPT |
| NANOS1 |
| NANP |
| NANS |
| NAP1L1 |
| NAP1L3 |
| NAP1L5 |
| NAPA |
| NAPB |

| |
|---|
| NAPEPLD |
| NAPG |
| NAPRT |
| NAPSA |
| NARF |
| NARS1 |
| NASP |
| NAT1 |
| NAT10 |
| NAT14 |
| NAT9 |
| NATD1 |
| NAV1 |
| NAV2 |
| NAV3 |
| NAXD |
| NAXE |
| NBAS |
| NBEA |
| NBEAL2 |
| NBL1 |
| NBN |
| NBR1 |
| NCAM1 |
| NCAM2 |
| NCAPD2 |
| NCAPD3 |
| NCAPG |
| NCAPG2 |
| NCAPH |
| NCAPH2 |
| NCBP1 |
| NCBP2 |
| NCBP2AS2 |
| NCDN |
| NCEH1 |
| NCF2 |
| NCK1 |
| NCK2 |
| NCKAP1 |
| NCKAP1L |
| NCKAP5 |
| NCKAP5L |
| NCKIPSD |
| NCL |

| |
|---|
| NCLN |
| NCOA1 |
| NCOA2 |
| NCOA3 |
| NCOA4 |
| NCOA5 |
| NCOA6 |
| NCOA7 |
| NCOR1 |
| NCOR2 |
| NCR3LG1 |
| NCS1 |
| NCSTN |
| NDC1 |
| NDC80 |
| NDE1 |
| NDEL1 |
| NDFIP1 |
| NDFIP2 |
| NDOR1 |
| NDRG1 |
| NDRG2 |
| NDST2 |
| NDUFA10 |
| NDUFA12 |
| NDUFA13 |
| NDUFA2 |
| NDUFA3 |
| NDUFA4 |
| NDUFA6 |
| NDUFA8 |
| NDUFAB1 |
| NDUFAF1 |
| NDUFAF2 |
| NDUFAF3 |
| NDUFAF4 |
| NDUFAF5 |
| NDUFAF6 |
| NDUFAF7 |
| NDUFAF8 |
| NDUFB1 |
| NDUFB10 |
| NDUFB11 |
| NDUFB2 |
| NDUFB4 |

| |
|---|
| NDUFB5 |
| NDUFB6 |
| NDUFB7 |
| NDUFB8 |
| NDUFB9 |
| NDUFC1 |
| NDUFC2 |
| NDUFS2 |
| NDUFS3 |
| NDUFS5 |
| NDUFS6 |
| NDUFS7 |
| NDUFV1 |
| NDUFV2 |
| NDUFV3 |
| NEAT1 |
| NEBL |
| NECAP1 |
| NECAP2 |
| NECTIN1 |
| NECTIN2 |
| NEDD4 |
| NEDD8 |
| NEDD9 |
| NEFH |
| NEFL |
| NEFM |
| NEGR1 |
| NEIL1 |
| NEIL2 |
| NEIL3 |
| NEK1 |
| NEK10 |
| NEK2 |
| NEK3 |
| NEK4 |
| NEK6 |
| NEK7 |
| NEK8 |
| NEK9 |
| NELFA |
| NELFB |
| NELFCD |
| NELFE |
| NELL1 |

| |
|---|
| NELL2 |
| NEMF |
| NEMP1 |
| NEMP2 |
| NENF |
| NEO1 |
| NES |
| NET1 |
| NEU1 |
| NEU3 |
| NEURL1B |
| NEURL2 |
| NEURL4 |
| NEXN |
| NF1 |
| NFAM1 |
| NFAT5 |
| NFATC1 |
| NFATC3 |
| NFE2 |
| NFE2L1 |
| NFE2L3 |
| NFIC |
| NFIL3 |
| NFIX |
| NFKB1 |
| NFKBIA |
| NFKBIB |
| NFKBID |
| NFKBIE |
| NFKBIL1 |
| NFKBIZ |
| NFS1 |
| NFX1 |
| NFXL1 |
| NFYA |
| NFYB |
| NFYC |
| NGDN |
| NGLY1 |
| NGRN |
| NHEJ1 |
| NHLRC1 |
| NHLRC2 |
| NHLRC3 |

| |
|---|
| NHP2 |
| NHSL2 |
| NIBAN1 |
| NIBAN2 |
| NICN1 |
| NID1 |
| NID2 |
| NIF3L1 |
| NIFK |
| NIN |
| NINJ1 |
| NINL |
| NIP7 |
| NIPA1 |
| NIPA2 |
| NIPAL1 |
| NIPAL3 |
| NIPBL |
| NIPSNAP2 |
| NIPSNAP3A |
| NISCH |
| NIT1 |
| NIT2 |
| NKAIN1 |
| NKAP |
| NKAPD1 |
| NKD1 |
| NKG7 |
| NKIRAS1 |
| NKIRAS2 |
| NKPD1 |
| NKRF |
| NKTR |
| NKX1-2 |
| NLE1 |
| NLGN2 |
| NLGN3 |
| NLK |
| NLN |
| NLRC3 |
| NLRC4 |
| NLRP1 |
| NLRP6 |
| NMB |
| NMD3 |

| |
|---|
| NME2 |
| NME3 |
| NME6 |
| NME7 |
| NMI |
| NMNAT1 |
| NMNAT3 |
| NMRAL1 |
| NMRK1 |
| NMT1 |
| NNAT |
| NNT |
| NOA1 |
| NOB1 |
| NOC2L |
| NOC3L |
| NOC4L |
| NOCT |
| NOD1 |
| NODAL |
| NOL10 |
| NOL11 |
| NOL12 |
| NOL4L |
| NOL6 |
| NOL7 |
| NOL8 |
| NOL9 |
| NOM1 |
| NOMO1 |
| NONO |
| NOP14 |
| NOP16 |
| NOP2 |
| NOP53 |
| NOP56 |
| NOP58 |
| NOP9 |
| NOS1AP |
| NOSIP |
| NOTCH1 |
| NOTCH2 |
| NOTCH3 |
| NOTCH4 |
| NOTUM |

| |
|---|
| NOVA1 |
| NOXA1 |
| NOXRED1 |
| NPAS3 |
| NPAT |
| NPB |
| NPC2 |
| NPDC1 |
| NPEPL1 |
| NPEPPS |
| NPHP1 |
| NPHP3 |
| NPHP4 |
| NPL |
| NPLOC4 |
| NPM1 |
| NPM3 |
| NPNT |
| NPR2 |
| NPR3 |
| NPRL2 |
| NPRL3 |
| NPTN |
| NPTX1 |
| NPTXR |
| NPY1R |
| NQO1 |
| NQO2 |
| NR1D2 |
| NR1H2 |
| NR2C2 |
| NR2C2AP |
| NR2F1 |
| NR2F2 |
| NR2F6 |
| NR3C2 |
| NR4A1 |
| NR4A2 |
| NR4A3 |
| NR5A2 |
| NR6A1 |
| NRARP |
| NRBF2 |
| NRBP1 |
| NRBP2 |

| |
|---|
| NRDC |
| NRDE2 |
| NREP |
| NRF1 |
| NRIP1 |
| NRIP3 |
| NRM |
| NRP1 |
| NRSN2 |
| NRXN1 |
| NSA2 |
| NSD1 |
| NSD2 |
| NSD3 |
| NSDHL |
| NSF |
| NSFL1C |
| NSL1 |
| NSMAF |
| NSMCE1 |
| NSMCE2 |
| NSMCE3 |
| NSMCE4A |
| NSMF |
| NSUN3 |
| NSUN4 |
| NSUN5 |
| NSUN6 |
| NT5C |
| NT5C2 |
| NT5C3A |
| NT5C3B |
| NT5DC1 |
| NT5DC2 |
| NT5DC3 |
| NT5E |
| NT5M |
| NTAN1 |
| NTAQ1 |
| NTHL1 |
| NTMT1 |
| NTN3 |
| NTN4 |
| NTN5 |
| NTNG1 |

| |
|---|
| NTRK1 |
| NUAK1 |
| NUAK2 |
| NUB1 |
| NUBP1 |
| NUBP2 |
| NUBPL |
| NUCB1 |
| NUCB2 |
| NUCKS1 |
| NUDC |
| NUDCD1 |
| NUDCD2 |
| NUDCD3 |
| NUDT1 |
| NUDT12 |
| NUDT13 |
| NUDT14 |
| NUDT15 |
| NUDT16 |
| NUDT16L1 |
| NUDT17 |
| NUDT18 |
| NUDT2 |
| NUDT21 |
| NUDT22 |
| NUDT3 |
| NUDT5 |
| NUDT6 |
| NUDT7 |
| NUDT8 |
| NUDT9 |
| NUF2 |
| NUFIP1 |
| NUFIP2 |
| NUGGC |
| NUMA1 |
| NUMB |
| NUMBL |
| NUP133 |
| NUP153 |
| NUP155 |
| NUP160 |
| NUP188 |
| NUP205 |

| |
|---|
| NUP210 |
| NUP214 |
| NUP35 |
| NUP42 |
| NUP43 |
| NUP50 |
| NUP54 |
| NUP58 |
| NUP62 |
| NUP85 |
| NUP88 |
| NUP93 |
| NUP98 |
| NUPR1 |
| NUS1 |
| NUSAP1 |
| NVL |
| NXF1 |
| NXPE3 |
| NXPH1 |
| NXT1 |
| NXT2 |
| NYAP1 |
| NYNRIN |
| OAF |
| OARD1 |
| OAS3 |
| OASL |
| OAT |
| OAZ1 |
| OAZ2 |
| OBI1 |
| OBSCN |
| OBSL1 |
| OCEL1 |
| OCIAD1 |
| OCIAD2 |
| OCLN |
| OCRL |
| ODAD3 |
| ODAD4 |
| ODC1 |
| ODF2 |
| ODF2L |
| ODR4 |

| |
|---|
| OFD1 |
| OGA |
| OGFOD1 |
| OGFOD3 |
| OGFR |
| OGFRL1 |
| OGG1 |
| OIP5 |
| OIT3 |
| OLA1 |
| OLFM1 |
| OLFM4 |
| OLFML1 |
| OLFML2A |
| OLFML2B |
| OLR1 |
| OMA1 |
| ONECUT1 |
| OPA1 |
| OPA3 |
| OPHN1 |
| OPRK1 |
| OPTN |
| OR51B5 |
| OR7D2 |
| ORAI1 |
| ORAI3 |
| ORC1 |
| ORC2 |
| ORC3 |
| ORC4 |
| ORC6 |
| ORMDL2 |
| ORMDL3 |
| OS9 |
| OSBP |
| OSBP2 |
| OSBPL10 |
| OSBPL11 |
| OSBPL1A |
| OSBPL2 |
| OSBPL3 |
| OSBPL5 |
| OSBPL8 |
| OSBPL9 |

| |
|---|
| OSCAR |
| OSCP1 |
| OSER1 |
| OSGEP |
| OSGEPL1 |
| OSGIN1 |
| OSR2 |
| OSTC |
| OSTF1 |
| OSTM1 |
| OTUB1 |
| OTUD1 |
| OTUD3 |
| OTUD4 |
| OTUD5 |
| OTUD6B |
| OTULIN |
| OTULINL |
| OVAAL |
| OVGP1 |
| OVOL1 |
| OXA1L |
| OXCT1 |
| OXER1 |
| OXNAD1 |
| OXR1 |
| OXSM |
| OXSR1 |
| P2RX4 |
| P2RY1 |
| P2RY4 |
| P3H1 |
| P3H2 |
| P3H3 |
| P3H4 |
| P4HA1 |
| P4HA2 |
| P4HA3 |
| P4HB |
| P4HTM |
| PA2G4 |
| PAAF1 |
| PABIR1 |
| PABIR2 |
| PABPC1 |

| |
|---|
| PABPC4 |
| PABPC5 |
| PABPN1 |
| PACC1 |
| PACRGL |
| PACS2 |
| PACSIN2 |
| PACSIN3 |
| PADI1 |
| PADI2 |
| PAF1 |
| PAFAH1B1 |
| PAFAH1B2 |
| PAFAH1B3 |
| PAFAH2 |
| PAG1 |
| PAIP1 |
| PAIP2 |
| PAK1 |
| PAK1IP1 |
| PAK2 |
| PAK4 |
| PALB2 |
| PALD1 |
| PALLD |
| PALMD |
| PAM |
| PAN2 |
| PAN3 |
| PANK2 |
| PANK3 |
| PANK4 |
| PANX1 |
| PANX2 |
| PAOX |
| PAPLN |
| PAPOLA |
| PAPOLG |
| PAPSS1 |
| PAPSS2 |
| PAQR3 |
| PAQR4 |
| PAQR5 |
| PAQR6 |
| PAQR8 |

| |
|---|
| PARD3 |
| PARD6A |
| PARD6B |
| PARD6G |
| PARG |
| PARK7 |
| PARL |
| PARN |
| PARP1 |
| PARP11 |
| PARP12 |
| PARP14 |
| PARP16 |
| PARP2 |
| PARP6 |
| PARP9 |
| PARPBP |
| PARS2 |
| PARVA |
| PASK |
| PATJ |
| PATL1 |
| PATZ1 |
| PAWR |
| PAXBP1 |
| PAXIP1 |
| PAXX |
| PBDC1 |
| PBK |
| PBLD |
| PBRM1 |
| PBX1 |
| PBX2 |
| PBX3 |
| PBX4 |
| PBXIP1 |
| PC |
| PCAT1 |
| PCBD1 |
| PCBD2 |
| PCBP1 |
| PCBP2 |
| PCBP4 |
| PCCA |
| PCCB |

| |
|---|
| PCDH1 |
| PCDH10 |
| PCDH18 |
| PCDH7 |
| PCDH9 |
| PCDHA3 |
| PCDHA4 |
| PCDHB11 |
| PCDHB14 |
| PCDHB15 |
| PCDHB4 |
| PCDHB6 |
| PCDHB7 |
| PCDHGA11 |
| PCDHGA12 |
| PCDHGA3 |
| PCDHGA4 |
| PCDHGA5 |
| PCDHGA6 |
| PCDHGA7 |
| PCDHGA8 |
| PCDHGA9 |
| PCDHGB1 |
| PCDHGB2 |
| PCDHGB6 |
| PCDHGC5 |
| PCED1A |
| PCED1B |
| PCF11 |
| PCGF1 |
| PCGF2 |
| PCGF3 |
| PCGF5 |
| PCGF6 |
| PCID2 |
| PCIF1 |
| PCK2 |
| PCLAF |
| PCLO |
| PCM1 |
| PCMT1 |
| PCMTD1 |
| PCMTD2 |
| PCNA |
| PCNP |

| |
|---|
| PCNT |
| PCNX1 |
| PCNX3 |
| PCNX4 |
| PCOLCE |
| PCOLCE2 |
| PCP4L1 |
| PCSK1N |
| PCSK4 |
| PCSK5 |
| PCSK6 |
| PCSK7 |
| PCTP |
| PCYOX1 |
| PCYOX1L |
| PCYT1A |
| PCYT2 |
| PDAP1 |
| PDCD10 |
| PDCD11 |
| PDCD2 |
| PDCD2L |
| PDCD4 |
| PDCD5 |
| PDCD6 |
| PDCD6IP |
| PDCD7 |
| PDCL |
| PDCL3 |
| PDE12 |
| PDE2A |
| PDE4B |
| PDE4C |
| PDE4D |
| PDE4DIP |
| PDE5A |
| PDE6D |
| PDE7A |
| PDE7B |
| PDE8A |
| PDE8B |
| PDE9A |
| PDF |
| PDGFA |
| PDGFB |

| |
|---|
| PDGFC |
| PDGFD |
| PDGFRB |
| PDHA1 |
| PDHB |
| PDHX |
| PDIA2 |
| PDIA3 |
| PDIA4 |
| PDIA5 |
| PDIA6 |
| PDIK1L |
| PDK1 |
| PDK3 |
| PDK4 |
| PDLIM1 |
| PDLIM2 |
| PDLIM4 |
| PDLIM5 |
| PDLIM7 |
| PDP2 |
| PDPK1 |
| PDRG1 |
| PDS5A |
| PDS5B |
| PDSS1 |
| PDXDC1 |
| PDXK |
| PDXP |
| PDZD11 |
| PDZD2 |
| PDZD4 |
| PDZD8 |
| PDZK1 |
| PDZRN3 |
| PDZRN4 |
| PEAK1 |
| PEBP1 |
| PECR |
| PEDS1 |
| PEG10 |
| PEG3 |
| PELI2 |
| PELI3 |
| PELO |

| |
|---|
| PELP1 |
| PEMT |
| PENK |
| PER1 |
| PER2 |
| PER3 |
| PERP |
| PET100 |
| PET117 |
| PEX1 |
| PEX10 |
| PEX11A |
| PEX11B |
| PEX11G |
| PEX12 |
| PEX13 |
| PEX14 |
| PEX16 |
| PEX19 |
| PEX2 |
| PEX3 |
| PEX5 |
| PEX6 |
| PEX7 |
| PF4 |
| PFAS |
| PFDN1 |
| PFDN2 |
| PFDN4 |
| PFDN5 |
| PFDN6 |
| PFKFB2 |
| PFKFB3 |
| PFKFB4 |
| PFKL |
| PFKM |
| PFKP |
| PFN1 |
| PFN2 |
| PGAM1 |
| PGAM5 |
| PGAP1 |
| PGAP2 |
| PGAP3 |
| PGAP4 |

| |
|---|
| PGAP6 |
| PGBD1 |
| PGBD2 |
| PGBD5 |
| PGF |
| PGGT1B |
| PGK1 |
| PGLS |
| PGM1 |
| PGM2 |
| PGM2L1 |
| PGM3 |
| PGM5 |
| PGP |
| PGPEP1 |
| PGR |
| PGRMC1 |
| PGRMC2 |
| PGS1 |
| PHACTR1 |
| PHACTR2 |
| PHACTR4 |
| PHAX |
| PHB2 |
| PHC1 |
| PHC2 |
| PHC3 |
| PHETA1 |
| PHETA2 |
| PHF1 |
| PHF11 |
| PHF12 |
| PHF13 |
| PHF14 |
| PHF19 |
| PHF2 |
| PHF20 |
| PHF21A |
| PHF23 |
| PHF24 |
| PHF3 |
| PHF5A |
| PHF6 |
| PHF8 |
| PHGDH |

| |
|---|
| PHIP |
| PHKA1 |
| PHKA2 |
| PHKB |
| PHKG2 |
| PHLDA1 |
| PHLDA2 |
| PHLDA3 |
| PHLDB1 |
| PHLDB2 |
| PHLDB3 |
| PHLPP1 |
| PHLPP2 |
| PHOSPHO2 |
| PHRF1 |
| PHYH |
| PHYKPL |
| PI16 |
| PI4K2A |
| PI4K2B |
| PI4KA |
| PI4KB |
| PIAS1 |
| PIAS2 |
| PIAS3 |
| PIAS4 |
| PICALM |
| PICK1 |
| PIDD1 |
| PIEZO1 |
| PIF1 |
| PIGA |
| PIGBOS1 |
| PIGC |
| PIGF |
| PIGG |
| PIGH |
| PIGK |
| PIGL |
| PIGM |
| PIGN |
| PIGO |
| PIGP |
| PIGR |
| PIGS |

| |
|---|
| PIGT |
| PIGU |
| PIGV |
| PIGZ |
| PIH1D1 |
| PIH1D2 |
| PIK3C2A |
| PIK3C2B |
| PIK3C3 |
| PIK3CA |
| PIK3CB |
| PIK3CD |
| PIK3IP1 |
| PIK3R1 |
| PIK3R2 |
| PIK3R3 |
| PIK3R4 |
| PIK3R5 |
| PIKFYVE |
| PIM1 |
| PIM2 |
| PIM3 |
| PIMREG |
| PIN1 |
| PIN4 |
| PINK1 |
| PINX1 |
| PIP4K2A |
| PIP4K2B |
| PIP4P1 |
| PIP5K1A |
| PIP5K1B |
| PIP5K1C |
| PIR |
| PISD |
| PITHD1 |
| PITPNA |
| PITPNM1 |
| PITPNM2 |
| PITRM1 |
| PITX3 |
| PIWIL2 |
| PJA1 |
| PJVK |
| PKD1 |

| |
|---|
| PKD2 |
| PKD2L2 |
| PKDCC |
| PKIA |
| PKIG |
| PKMYT1 |
| PKN1 |
| PKN2 |
| PKN3 |
| PKNOX1 |
| PKP1 |
| PKP2 |
| PKP3 |
| PKP4 |
| PLA2G12A |
| PLA2G15 |
| PLA2G6 |
| PLA2G7 |
| PLAA |
| PLAAT2 |
| PLAAT3 |
| PLAC8 |
| PLAGL1 |
| PLAGL2 |
| PLAU |
| PLAUR |
| PLBD1 |
| PLBD2 |
| PLCB3 |
| PLCB4 |
| PLCD1 |
| PLCD3 |
| PLCE1 |
| PLCG1 |
| PLCG2 |
| PLCH2 |
| PLCL2 |
| PLCXD1 |
| PLCXD2 |
| PLD2 |
| PLD3 |
| PLEC |
| PLEK |
| PLEK2 |
| PLEKHA1 |

| |
|---|
| PLEKHA2 |
| PLEKHA3 |
| PLEKHA4 |
| PLEKHA5 |
| PLEKHA7 |
| PLEKHA8 |
| PLEKHB1 |
| PLEKHD1 |
| PLEKHF1 |
| PLEKHF2 |
| PLEKHG1 |
| PLEKHG2 |
| PLEKHG3 |
| PLEKHG4B |
| PLEKHG5 |
| PLEKHG6 |
| PLEKHH1 |
| PLEKHH3 |
| PLEKHJ1 |
| PLEKHM1 |
| PLEKHM2 |
| PLEKHM3 |
| PLEKHN1 |
| PLEKHO1 |
| PLEKHO2 |
| PLGRKT |
| PLIN1 |
| PLIN2 |
| PLIN3 |
| PLK1 |
| PLK2 |
| PLK3 |
| PLK4 |
| PLOD1 |
| PLOD2 |
| PLOD3 |
| PLP2 |
| PLPBP |
| PLPP1 |
| PLPP2 |
| PLPP3 |
| PLPP5 |
| PLPP6 |
| PLPP7 |
| PLPPR2 |

| |
|---|
| PLPPR4 |
| PLRG1 |
| PLS3 |
| PLSCR4 |
| PLSCR5 |
| PLTP |
| PLXDC1 |
| PLXNB1 |
| PLXNB2 |
| PLXNB3 |
| PLXNC1 |
| PLXND1 |
| PM20D2 |
| PMEL |
| PMF1 |
| PML |
| PMM1 |
| PMM2 |
| PMP22 |
| PMPCA |
| PMPCB |
| PMS1 |
| PMVK |
| PNISR |
| PNKD |
| PNKP |
| PNMA1 |
| PNMA8A |
| PNN |
| PNO1 |
| PNP |
| PNPLA2 |
| PNPLA3 |
| PNPLA4 |
| PNPLA6 |
| PNPLA7 |
| PNPLA8 |
| PNPO |
| PNPT1 |
| PNRC1 |
| PNRC2 |
| POC1A |
| POC1B |
| POC5 |
| POF1B |

| |
|---|
| POFUT1 |
| POFUT2 |
| POGK |
| POGLUT1 |
| POGLUT2 |
| POGLUT3 |
| POGZ |
| POLA1 |
| POLA2 |
| POLB |
| POLD1 |
| POLD2 |
| POLD3 |
| POLD4 |
| POLDIP2 |
| POLDIP3 |
| POLE |
| POLE2 |
| POLE3 |
| POLE4 |
| POLG |
| POLG2 |
| POLH |
| POLI |
| POLK |
| POLL |
| POLM |
| POLQ |
| POLR1C |
| POLR1D |
| POLR1E |
| POLR1F |
| POLR1G |
| POLR1H |
| POLR2A |
| POLR2B |
| POLR2C |
| POLR2D |
| POLR2E |
| POLR2F |
| POLR2G |
| POLR2H |
| POLR2I |
| POLR2J |
| POLR2L |

| |
|---|
| POLR2M |
| POLR3A |
| POLR3B |
| POLR3C |
| POLR3E |
| POLR3F |
| POLR3GL |
| POLR3H |
| POLR3K |
| POLRMT |
| POM121 |
| POM121C |
| POM121L2 |
| POMC |
| POMGNT1 |
| POMGNT2 |
| POMK |
| POMP |
| POMT1 |
| POMT2 |
| PON2 |
| POP1 |
| POP5 |
| POP7 |
| POR |
| PORCN |
| POSTN |
| POT1 |
| POU2AF1 |
| POU2F1 |
| POU5F1 |
| POU6F1 |
| PPA1 |
| PPA2 |
| PPAN |
| PPARA |
| PPARD |
| PPARG |
| PPARGC1B |
| PPAT |
| PPCDC |
| PPCS |
| PPDPF |
| PPFIA1 |
| PPFIA4 |

| |
|---|
| PPFIBP1 |
| PPFIBP2 |
| PPHLN1 |
| PPIA |
| PPIB |
| PPIC |
| PPID |
| PPIE |
| PPIF |
| PPIG |
| PPIH |
| PPIL1 |
| PPIL2 |
| PPIL3 |
| PPIL4 |
| PPIL6 |
| PPIP5K1 |
| PPIP5K2 |
| PPL |
| PPM1A |
| PPM1B |
| PPM1D |
| PPM1F |
| PPM1G |
| PPM1J |
| PPM1K |
| PPM1M |
| PPOX |
| PPP1CA |
| PPP1CC |
| PPP1R10 |
| PPP1R11 |
| PPP1R12A |
| PPP1R12B |
| PPP1R12C |
| PPP1R13B |
| PPP1R13L |
| PPP1R14A |
| PPP1R14B |
| PPP1R14C |
| PPP1R14D |
| PPP1R15A |
| PPP1R15B |
| PPP1R18 |
| PPP1R2 |

| |
|---|
| PPP1R21 |
| PPP1R26 |
| PPP1R32 |
| PPP1R35 |
| PPP1R37 |
| PPP1R3A |
| PPP1R3B |
| PPP1R3D |
| PPP1R3E |
| PPP1R3F |
| PPP1R3G |
| PPP1R7 |
| PPP1R8 |
| PPP1R9A |
| PPP1R9B |
| PPP2CA |
| PPP2CB |
| PPP2R1A |
| PPP2R1B |
| PPP2R2A |
| PPP2R2D |
| PPP2R3A |
| PPP2R3B |
| PPP2R3C |
| PPP2R5A |
| PPP2R5B |
| PPP2R5C |
| PPP2R5D |
| PPP2R5E |
| PPP3CA |
| PPP3CB |
| PPP3R1 |
| PPP4C |
| PPP4R1 |
| PPP4R2 |
| PPP4R3A |
| PPP4R3B |
| PPP4R4 |
| PPP5C |
| PPP6C |
| PPP6R2 |
| PPP6R3 |
| PPRC1 |
| PPT1 |
| PPT2 |

| |
|---|
| PPTC7 |
| PPWD1 |
| PQBP1 |
| PRADC1 |
| PRAF2 |
| PRAM1 |
| PRAME |
| PRC1 |
| PRCC |
| PRCP |
| PRDM1 |
| PRDM10 |
| PRDM11 |
| PRDM15 |
| PRDM2 |
| PRDM4 |
| PRDM5 |
| PRDM7 |
| PRDX2 |
| PRDX3 |
| PRDX4 |
| PRDX5 |
| PRDX6 |
| PREB |
| PRELID1 |
| PRELID3B |
| PREP |
| PREPL |
| PREX1 |
| PRICKLE1 |
| PRICKLE3 |
| PRICKLE4 |
| PRIM1 |
| PRIM2 |
| PRIMPOL |
| PRKAA1 |
| PRKAA2 |
| PRKAB1 |
| PRKACA |
| PRKACB |
| PRKAG1 |
| PRKAG2 |
| PRKAR1B |
| PRKAR2A |
| PRKCD |

| |
|---|
| PRKCE |
| PRKCI |
| PRKCSH |
| PRKCZ |
| PRKD1 |
| PRKD2 |
| PRKD3 |
| PRKDC |
| PRKRA |
| PRKRIP1 |
| PRMT1 |
| PRMT2 |
| PRMT3 |
| PRMT5 |
| PRMT6 |
| PRMT7 |
| PRMT9 |
| PROB1 |
| PRODH |
| PRORP |
| PROS1 |
| PROSER1 |
| PROSER2 |
| PROSER3 |
| PRPF18 |
| PRPF19 |
| PRPF3 |
| PRPF31 |
| PRPF38A |
| PRPF38B |
| PRPF39 |
| PRPF40A |
| PRPF40B |
| PRPF4B |
| PRPF6 |
| PRPF8 |
| PRPS1 |
| PRPS2 |
| PRPSAP1 |
| PRPSAP2 |
| PRR11 |
| PRR12 |
| PRR14 |
| PRR14L |
| PRR15 |

| |
|---|
| PRR15L |
| PRR23A |
| PRR5 |
| PRR7 |
| PRRC1 |
| PRRC2A |
| PRRC2C |
| PRRG2 |
| PRSS35 |
| PRSS36 |
| PRUNE2 |
| PRX |
| PRXL2A |
| PRXL2B |
| PRXL2C |
| PSAP |
| PSAT1 |
| PSD3 |
| PSD4 |
| PSEN1 |
| PSEN2 |
| PSENEN |
| PSIP1 |
| PSKH1 |
| PSMA1 |
| PSMA2 |
| PSMA3 |
| PSMA5 |
| PSMA6 |
| PSMA7 |
| PSMB1 |
| PSMB10 |
| PSMB2 |
| PSMB3 |
| PSMB4 |
| PSMB5 |
| PSMB7 |
| PSMB9 |
| PSMC1 |
| PSMC2 |
| PSMC3 |
| PSMC3IP |
| PSMC5 |
| PSMC6 |
| PSMD1 |

| |
|---|
| PSMD10 |
| PSMD11 |
| PSMD12 |
| PSMD13 |
| PSMD14 |
| PSMD2 |
| PSMD3 |
| PSMD4 |
| PSMD5 |
| PSMD6 |
| PSMD7 |
| PSMD8 |
| PSMD9 |
| PSME1 |
| PSME2 |
| PSME3 |
| PSME4 |
| PSMF1 |
| PSMG2 |
| PSMG3 |
| PSMG4 |
| PSPC1 |
| PSPH |
| PSRC1 |
| PSTK |
| PTAFR |
| PTAR1 |
| PTBP1 |
| PTBP3 |
| PTCD1 |
| PTCD2 |
| PTCD3 |
| PTCH1 |
| PTCH2 |
| PTCHD4 |
| PTDSS1 |
| PTDSS2 |
| PTEN |
| PTER |
| PTGDS |
| PTGER1 |
| PTGER2 |
| PTGER3 |
| PTGER4 |
| PTGES2 |

| |
|---|
| PTGES3 |
| PTGES3L |
| PTGFR |
| PTGFRN |
| PTGS2 |
| PTH2 |
| PTHLH |
| PTK2 |
| PTK2B |
| PTK6 |
| PTK7 |
| PTMA |
| PTN |
| PTOV1 |
| PTP4A1 |
| PTP4A2 |
| PTP4A3 |
| PTPA |
| PTPDC1 |
| PTPMT1 |
| PTPN1 |
| PTPN11 |
| PTPN12 |
| PTPN18 |
| PTPN2 |
| PTPN21 |
| PTPN23 |
| PTPN3 |
| PTPN6 |
| PTPN9 |
| PTPRA |
| PTPRB |
| PTPRC |
| PTPRD |
| PTPRE |
| PTPRF |
| PTPRG |
| PTPRH |
| PTPRJ |
| PTPRK |
| PTPRM |
| PTPRQ |
| PTPRS |
| PTRHD1 |
| PTS |

| |
|---|
| PTTG1 |
| PTX3 |
| PTX4 |
| PUDP |
| PUF60 |
| PUM1 |
| PUM2 |
| PUM3 |
| PURA |
| PURB |
| PURG |
| PUS1 |
| PUS3 |
| PUS7 |
| PUS7L |
| PUSL1 |
| PVR |
| PWP1 |
| PWP2 |
| PWWP2A |
| PWWP2B |
| PXDC1 |
| PXK |
| PXMP2 |
| PXMP4 |
| PXN |
| PYCR1 |
| PYCR3 |
| PYGB |
| PYGL |
| PYGM |
| PYGO1 |
| PYM1 |
| PYROXD1 |
| PYURF |
| QARS1 |
| QDPR |
| QKI |
| QPCTL |
| QRFPR |
| QRICH1 |
| QRSL1 |
| QSER1 |
| QSOX1 |
| QSOX2 |

| |
|---|
| QTRT1 |
| QTRT2 |
| R3HCC1 |
| R3HCC1L |
| R3HDM2 |
| R3HDM4 |
| RAB10 |
| RAB11A |
| RAB11B |
| RAB11FIP1 |
| RAB11FIP2 |
| RAB11FIP3 |
| RAB11FIP4 |
| RAB11FIP5 |
| RAB12 |
| RAB13 |
| RAB14 |
| RAB15 |
| RAB18 |
| RAB1A |
| RAB1B |
| RAB20 |
| RAB21 |
| RAB22A |
| RAB23 |
| RAB24 |
| RAB25 |
| RAB27A |
| RAB27B |
| RAB28 |
| RAB29 |
| RAB2A |
| RAB2B |
| RAB30 |
| RAB31 |
| RAB32 |
| RAB33B |
| RAB34 |
| RAB35 |
| RAB36 |
| RAB37 |
| RAB38 |
| RAB39A |
| RAB3B |
| RAB3D |

| |
|---|
| RAB3GAP1 |
| RAB3GAP2 |
| RAB3IL1 |
| RAB40B |
| RAB40C |
| RAB43 |
| RAB4A |
| RAB4B |
| RAB5A |
| RAB5C |
| RAB5IF |
| RAB6A |
| RAB8A |
| RAB8B |
| RAB9A |
| RABEP1 |
| RABEP2 |
| RABEPK |
| RABGAP1L |
| RABGGTA |
| RABIF |
| RABL2A |
| RABL3 |
| RABL6 |
| RAC1 |
| RAC2 |
| RAC3 |
| RACGAP1 |
| RACK1 |
| RAD1 |
| RAD17 |
| RAD18 |
| RAD21 |
| RAD23A |
| RAD23B |
| RAD50 |
| RAD51 |
| RAD51AP1 |
| RAD51B |
| RAD51C |
| RAD51D |
| RAD52 |
| RAD54B |
| RAD54L2 |
| RAD9A |

| |
|---|
| RAE1 |
| RAF1 |
| RAI1 |
| RAI2 |
| RALA |
| RALB |
| RALBP1 |
| RALGAPA1 |
| RALGAPA2 |
| RALGDS |
| RALGPS1 |
| RALGPS2 |
| RALY |
| RALYL |
| RAMAC |
| RAN |
| RANBP1 |
| RANBP10 |
| RANBP17 |
| RANBP3 |
| RANBP6 |
| RANBP9 |
| RANGAP1 |
| RANGRF |
| RAP1A |
| RAP1B |
| RAP1GAP |
| RAP1GAP2 |
| RAP2A |
| RAP2B |
| RAP2C |
| RAPGEF1 |
| RAPGEF2 |
| RAPGEF3 |
| RAPGEF4 |
| RAPGEF6 |
| RAPGEFL1 |
| RAPH1 |
| RAPSN |
| RARA |
| RARG |
| RARRES1 |
| RARS1 |
| RARS2 |
| RASA1 |

| |
|---|
| RASA2 |
| RASA3 |
| RASAL2 |
| RASAL3 |
| RASD1 |
| RASEF |
| RASGEF1B |
| RASGEF1C |
| RASGRP2 |
| RASGRP3 |
| RASGRP4 |
| RASIP1 |
| RASL10B |
| RASL11A |
| RASL11B |
| RASSF1 |
| RASSF2 |
| RASSF3 |
| RASSF5 |
| RASSF6 |
| RASSF7 |
| RASSF8 |
| RASSF9 |
| RAVER1 |
| RAVER2 |
| RB1 |
| RB1CC1 |
| RBBP4 |
| RBBP5 |
| RBBP6 |
| RBBP7 |
| RBBP8 |
| RBBP9 |
| RBCK1 |
| RBFA |
| RBFOX2 |
| RBIS |
| RBKS |
| RBL1 |
| RBL2 |
| RBM10 |
| RBM12 |
| RBM12B |
| RBM14 |
| RBM15 |

| |
|---|
| RBM15B |
| RBM17 |
| RBM18 |
| RBM22 |
| RBM25 |
| RBM27 |
| RBM28 |
| RBM3 |
| RBM33 |
| RBM34 |
| RBM41 |
| RBM42 |
| RBM43 |
| RBM45 |
| RBM47 |
| RBM48 |
| RBM4B |
| RBM5 |
| RBM6 |
| RBM7 |
| RBM8A |
| RBMS1 |
| RBMS3 |
| RBMX |
| RBMX2 |
| RBP2 |
| RBP7 |
| RBPMS2 |
| RBSN |
| RBX1 |
| RC3H1 |
| RCAN3 |
| RCBTB1 |
| RCBTB2 |
| RCC1 |
| RCC1L |
| RCC2 |
| RCCD1 |
| RCE1 |
| RCHY1 |
| RCL1 |
| RCN1 |
| RCN2 |
| RCN3 |
| RCOR1 |

| |
|---|
| RCOR2 |
| RCOR3 |
| RCSD1 |
| RDH10 |
| RDH11 |
| RDH13 |
| RDH14 |
| RDM1 |
| RECK |
| RECQL |
| RECQL4 |
| RECQL5 |
| REEP1 |
| REEP2 |
| REEP4 |
| REEP5 |
| REEP6 |
| REL |
| RELA |
| RELB |
| RELCH |
| RELL1 |
| RELL2 |
| RELN |
| RELT |
| REM2 |
| RENBP |
| REP15 |
| REPIN1 |
| REPS1 |
| REPS2 |
| RER1 |
| RERE |
| RERG |
| RESF1 |
| REST |
| RETREG1 |
| RETREG2 |
| RETREG3 |
| RETSAT |
| REV1 |
| REV3L |
| REX1BD |
| REXO1 |
| REXO2 |

| |
|---|
| REXO5 |
| RFC1 |
| RFC2 |
| RFC3 |
| RFC4 |
| RFC5 |
| RFESD |
| RFFL |
| RFK |
| RFLNB |
| RFNG |
| RFT1 |
| RFTN1 |
| RFTN2 |
| RFWD3 |
| RFX1 |
| RFX2 |
| RFX7 |
| RFXANK |
| RFXAP |
| RGCC |
| RGL1 |
| RGL2 |
| RGMA |
| RGMB |
| RGS10 |
| RGS11 |
| RGS12 |
| RGS18 |
| RGS19 |
| RGS2 |
| RGS4 |
| RGS5 |
| RGS9BP |
| RHBDD1 |
| RHBDD2 |
| RHBDD3 |
| RHBDF2 |
| RHBDL1 |
| RHBDL2 |
| RHCE |
| RHEB |
| RHNO1 |
| RHOB |
| RHOBTB1 |

| |
|---|
| RHOBTB2 |
| RHOBTB3 |
| RHOD |
| RHOF |
| RHOG |
| RHOQ |
| RHOT1 |
| RHOT2 |
| RHOU |
| RHPN2 |
| RIBC1 |
| RIC1 |
| RIC3 |
| RIC8A |
| RIC8B |
| RICTOR |
| RIDA |
| RIF1 |
| RILPL1 |
| RILPL2 |
| RIMKLA |
| RIMKLB |
| RIMS1 |
| RIMS2 |
| RIN1 |
| RING1 |
| RINT1 |
| RIOK1 |
| RIOK2 |
| RIOK3 |
| RIOX1 |
| RIOX2 |
| RIPK1 |
| RIPK2 |
| RIT1 |
| RITA1 |
| RLF |
| RLIM |
| RMC1 |
| RMDN2 |
| RMDN3 |
| RMI1 |
| RMI2 |
| RMND5A |
| RMND5B |

| |
|---|
| RNASE4 |
| RNASEH1 |
| RNASEH2A |
| RNASEH2B |
| RNASEH2C |
| RNASEK |
| RNASET2 |
| RND2 |
| RND3 |
| RNF10 |
| RNF103 |
| RNF11 |
| RNF111 |
| RNF112 |
| RNF113A |
| RNF114 |
| RNF115 |
| RNF123 |
| RNF126 |
| RNF13 |
| RNF133 |
| RNF135 |
| RNF139 |
| RNF14 |
| RNF141 |
| RNF145 |
| RNF148 |
| RNF149 |
| RNF152 |
| RNF166 |
| RNF167 |
| RNF168 |
| RNF169 |
| RNF170 |
| RNF181 |
| RNF182 |
| RNF185 |
| RNF187 |
| RNF19B |
| RNF2 |
| RNF20 |
| RNF208 |
| RNF213 |
| RNF214 |
| RNF215 |

| |
|---|
| RNF216 |
| RNF220 |
| RNF222 |
| RNF223 |
| RNF24 |
| RNF25 |
| RNF26 |
| RNF31 |
| RNF32 |
| RNF4 |
| RNF40 |
| RNF41 |
| RNF44 |
| RNF5 |
| RNF6 |
| RNF7 |
| RNFT1 |
| RNFT2 |
| RNGTT |
| RNH1 |
| RNLS |
| RNMT |
| RNPC3 |
| RNPEP |
| RNPEPL1 |
| RNPS1 |
| RO60 |
| ROBO1 |
| ROCK1 |
| ROCK2 |
| ROGDI |
| ROM1 |
| ROMO1 |
| RP2 |
| RP9 |
| RPA1 |
| RPA2 |
| RPA3 |
| RPA4 |
| RPAIN |
| RPAP1 |
| RPAP3 |
| RPE |
| RPF1 |
| RPF2 |

| |
|---|
| RPGR |
| RPGRIP1L |
| RPH3AL |
| RPIA |
| RPL10A |
| RPL11 |
| RPL12 |
| RPL13 |
| RPL13A |
| RPL14 |
| RPL15 |
| RPL17 |
| RPL18 |
| RPL18A |
| RPL19 |
| RPL21 |
| RPL22 |
| RPL22L1 |
| RPL23 |
| RPL23A |
| RPL24 |
| RPL26 |
| RPL27 |
| RPL27A |
| RPL28 |
| RPL29 |
| RPL3 |
| RPL31 |
| RPL32 |
| RPL34 |
| RPL35 |
| RPL36 |
| RPL36A |
| RPL37 |
| RPL37P3 |
| RPL38 |
| RPL4 |
| RPL5 |
| RPL6 |
| RPL7 |
| RPL7A |
| RPL7L1 |
| RPL8 |
| RPL9 |
| RPLP1 |

| |
|---|
| RPLP2 |
| RPN1 |
| RPN2 |
| RPP21 |
| RPP25 |
| RPP30 |
| RPP38 |
| RPP40 |
| RPRD1A |
| RPRD2 |
| RPRM |
| RPS10 |
| RPS12 |
| RPS13 |
| RPS14 |
| RPS15 |
| RPS15A |
| RPS16 |
| RPS17 |
| RPS18 |
| RPS19BP1 |
| RPS2 |
| RPS20 |
| RPS21 |
| RPS23 |
| RPS24 |
| RPS25 |
| RPS26 |
| RPS27 |
| RPS27A |
| RPS27L |
| RPS3 |
| RPS3A |
| RPS5 |
| RPS6 |
| RPS6KA1 |
| RPS6KA2 |
| RPS6KA3 |
| RPS6KA4 |
| RPS6KA5 |
| RPS6KB1 |
| RPS6KB2 |
| RPS6KC1 |
| RPS7 |
| RPS8 |

| |
|---|
| RPS9 |
| RPSA |
| RPTOR |
| RPUSD1 |
| RPUSD2 |
| RPUSD4 |
| RRAD |
| RRAGA |
| RRAGB |
| RRAGC |
| RRAGD |
| RRAS |
| RRAS2 |
| RRBP1 |
| RREB1 |
| RRM1 |
| RRM2 |
| RRM2B |
| RRN3 |
| RRNAD1 |
| RRP1 |
| RRP12 |
| RRP15 |
| RRP1B |
| RRP36 |
| RRP7A |
| RRP8 |
| RRP9 |
| RRS1 |
| RSAD1 |
| RSAD2 |
| RSBN1 |
| RSBN1L |
| RSF1 |
| RSKR |
| RSL1D1 |
| RSPH3 |
| RSPH4A |
| RSPRY1 |
| RSRC2 |
| RSRP1 |
| RTCA |
| RTCB |
| RTF1 |
| RTF2 |

| |
|---|
| RTL5 |
| RTL6 |
| RTL8A |
| RTL8C |
| RTN1 |
| RTN2 |
| RTN3 |
| RTN4 |
| RTN4IP1 |
| RTN4R |
| RTRAF |
| RTTN |
| RUBCN |
| RUFY1 |
| RUFY2 |
| RUFY3 |
| RUNDC1 |
| RUNX1 |
| RUNX3 |
| RUSC1 |
| RUSC2 |
| RUSF1 |
| RUVBL1 |
| RUVBL2 |
| RWDD1 |
| RWDD2A |
| RWDD2B |
| RWDD3 |
| RWDD4 |
| RXRA |
| RXRB |
| RXYLT1 |
| RYBP |
| RYK |
| RYR1 |
| RYR2 |
| RYR3 |
| S100A10 |
| S100A13 |
| S100A16 |
| S100A2 |
| S100A4 |
| S100A6 |
| S100A9 |
| S100P |

| |
|---|
| S100PBP |
| S1PR1 |
| S1PR2 |
| SAA2 |
| SAA4 |
| SAAL1 |
| SAC3D1 |
| SACM1L |
| SACS |
| SAE1 |
| SAFB |
| SAFB2 |
| SALL2 |
| SAMD1 |
| SAMD10 |
| SAMD11 |
| SAMD15 |
| SAMD4A |
| SAMD4B |
| SAMD5 |
| SAMD8 |
| SAMD9 |
| SAMHD1 |
| SAMM50 |
| SAP18 |
| SAP30 |
| SAP30BP |
| SAP30L |
| SAR1A |
| SAR1B |
| SARAF |
| SARDH |
| SARM1 |
| SARNP |
| SARS1 |
| SARS2 |
| SART1 |
| SART3 |
| SASH1 |
| SASS6 |
| SAT1 |
| SAT2 |
| SAV1 |
| SAXO2 |
| SAYSD1 |

| |
|---|
| SBDS |
| SBF1 |
| SBF2 |
| SBK1 |
| SBSN |
| SBSPON |
| SC5D |
| SCAF1 |
| SCAF11 |
| SCAF4 |
| SCAF8 |
| SCAI |
| SCAMP1 |
| SCAMP2 |
| SCAMP3 |
| SCAMP4 |
| SCAND1 |
| SCAPER |
| SCARA3 |
| SCARB2 |
| SCARF1 |
| SCARF2 |
| SCCPDH |
| SCD5 |
| SCFD1 |
| SCFD2 |
| SCG5 |
| SCIN |
| SCLT1 |
| SCLY |
| SCMH1 |
| SCML1 |
| SCML2 |
| SCN2A |
| SCN9A |
| SCNM1 |
| SCNN1A |
| SCNN1B |
| SCO1 |
| SCOC |
| SCP2 |
| SCPEP1 |
| SCRIB |
| SCRN1 |
| SCRN2 |

| |
|---|
| SCRN3 |
| SCUBE1 |
| SCUBE2 |
| SCUBE3 |
| SCX |
| SCYL1 |
| SCYL3 |
| SDAD1 |
| SDC1 |
| SDC2 |
| SDC3 |
| SDC4 |
| SDCBP |
| SDCBP2 |
| SDCCAG8 |
| SDE2 |
| SDF2 |
| SDF2L1 |
| SDF4 |
| SDHAF1 |
| SDHAF3 |
| SDHB |
| SDHC |
| SDHD |
| SDR39U1 |
| SEC11A |
| SEC11C |
| SEC13 |
| SEC14L1 |
| SEC14L2 |
| SEC16A |
| SEC22A |
| SEC22B |
| SEC22C |
| SEC23A |
| SEC23B |
| SEC23IP |
| SEC24A |
| SEC24B |
| SEC24C |
| SEC24D |
| SEC31A |
| SEC31B |
| SEC61A2 |
| SEC61B |

| |
|---|
| SEC61G |
| SEC62 |
| SEC63 |
| SECISBP2 |
| SECISBP2L |
| SECTM1 |
| SEH1L |
| SEL1L |
| SEL1L3 |
| SELE |
| SELENBP1 |
| SELENOI |
| SELENOM |
| SELENOO |
| SELENOS |
| SELENOT |
| SELENOV |
| SELPLG |
| SEM1 |
| SEMA3A |
| SEMA3B |
| SEMA3F |
| SEMA3G |
| SEMA4B |
| SEMA4C |
| SEMA4D |
| SEMA4F |
| SEMA4G |
| SEMA5A |
| SEMA6A |
| SEMA6B |
| SEMA6C |
| SEMA6D |
| SEMA7A |
| SENP1 |
| SENP2 |
| SENP3 |
| SENP5 |
| SENP6 |
| SENP7 |
| SENP8 |
| SEPHS1 |
| SEPHS2 |
| SEPTIN1 |
| SEPTIN10 |

| |
|---|
| SEPTIN11 |
| SEPTIN2 |
| SEPTIN5 |
| SEPTIN6 |
| SEPTIN7 |
| SEPTIN8 |
| SEPTIN9 |
| SERAC1 |
| SERBP1 |
| SERGEF |
| SERINC1 |
| SERINC2 |
| SERINC3 |
| SERINC5 |
| SERP1 |
| SERPINA5 |
| SERPINB1 |
| SERPINB2 |
| SERPINB6 |
| SERPINB7 |
| SERPINE1 |
| SERPINE2 |
| SERPINH1 |
| SERPINI1 |
| SERTAD2 |
| SERTAD3 |
| SESN3 |
| SET |
| SETBP1 |
| SETD1A |
| SETD1B |
| SETD2 |
| SETD4 |
| SETD5 |
| SETD6 |
| SETD7 |
| SETD9 |
| SETMAR |
| SETX |
| SF1 |
| SF3A1 |
| SF3A2 |
| SF3A3 |
| SF3B1 |
| SF3B2 |

| |
|---|
| SF3B3 |
| SF3B4 |
| SF3B5 |
| SF3B6 |
| SFI1 |
| SFN |
| SFPQ |
| SFR1 |
| SFRP1 |
| SFRP4 |
| SFSWAP |
| SFT2D1 |
| SFT2D2 |
| SFXN1 |
| SFXN2 |
| SFXN4 |
| SFXN5 |
| SGCE |
| SGK1 |
| SGK3 |
| SGMS2 |
| SGO1 |
| SGO2 |
| SGPL1 |
| SGPP1 |
| SGPP2 |
| SGSM1 |
| SGSM2 |
| SGSM3 |
| SGTA |
| SH2B1 |
| SH2B2 |
| SH2B3 |
| SH2D3C |
| SH2D4A |
| SH2D6 |
| SH3BGRL |
| SH3BGRL2 |
| SH3BGRL3 |
| SH3BP1 |
| SH3BP2 |
| SH3BP4 |
| SH3BP5 |
| SH3BP5L |
| SH3D19 |

| |
|---|
| SH3D21 |
| SH3GL1 |
| SH3GL2 |
| SH3GLB1 |
| SH3GLB2 |
| SH3KBP1 |
| SH3PXD2A |
| SH3PXD2B |
| SH3RF1 |
| SH3TC2 |
| SH3YL1 |
| SHARPIN |
| SHB |
| SHC1 |
| SHCBP1 |
| SHD |
| SHF |
| SHFL |
| SHISA4 |
| SHISA5 |
| SHISA8 |
| SHKBP1 |
| SHLD1 |
| SHLD2 |
| SHMT1 |
| SHMT2 |
| SHOC2 |
| SHPK |
| SHPRH |
| SHQ1 |
| SHROOM1 |
| SHROOM2 |
| SHROOM3 |
| SHTN1 |
| SIAE |
| SIAH1 |
| SIAH2 |
| SIDT2 |
| SIGLEC10 |
| SIGLEC8 |
| SIGMAR1 |
| SIK2 |
| SIL1 |
| SIMC1 |
| SIN3B |

| |
|---|
| SINHCAF |
| SIPA1 |
| SIPA1L1 |
| SIPA1L2 |
| SIPA1L3 |
| SIRPA |
| SIRT1 |
| SIRT2 |
| SIRT3 |
| SIRT5 |
| SIRT6 |
| SIRT7 |
| SIT1 |
| SIVA1 |
| SIX5 |
| SKA2 |
| SKA3 |
| SKI |
| SKIDA1 |
| SKIL |
| SKIV2L |
| SKP1 |
| SKP2 |
| SLA2 |
| SLAIN1 |
| SLAIN2 |
| SLAMF7 |
| SLBP |
| SLC10A3 |
| SLC10A4 |
| SLC10A5 |
| SLC11A1 |
| SLC11A2 |
| SLC12A2 |
| SLC12A4 |
| SLC12A5 |
| SLC12A7 |
| SLC12A9 |
| SLC13A4 |
| SLC15A1 |
| SLC15A3 |
| SLC15A4 |
| SLC16A1 |
| SLC16A11 |
| SLC16A12 |

| |
|---|
| SLC16A14 |
| SLC16A3 |
| SLC16A5 |
| SLC16A6 |
| SLC16A7 |
| SLC17A5 |
| SLC19A1 |
| SLC19A2 |
| SLC1A1 |
| SLC1A4 |
| SLC1A5 |
| SLC20A1 |
| SLC20A2 |
| SLC22A18 |
| SLC22A23 |
| SLC22A4 |
| SLC22A5 |
| SLC23A1 |
| SLC23A2 |
| SLC25A1 |
| SLC25A10 |
| SLC25A11 |
| SLC25A12 |
| SLC25A13 |
| SLC25A15 |
| SLC25A16 |
| SLC25A17 |
| SLC25A19 |
| SLC25A22 |
| SLC25A23 |
| SLC25A24 |
| SLC25A25 |
| SLC25A26 |
| SLC25A27 |
| SLC25A28 |
| SLC25A29 |
| SLC25A30 |
| SLC25A32 |
| SLC25A35 |
| SLC25A36 |
| SLC25A37 |
| SLC25A38 |
| SLC25A39 |
| SLC25A4 |
| SLC25A42 |

| |
|---|
| SLC25A43 |
| SLC25A46 |
| SLC25A5 |
| SLC25A51 |
| SLC25A53 |
| SLC26A1 |
| SLC26A11 |
| SLC26A2 |
| SLC26A3 |
| SLC26A4 |
| SLC27A1 |
| SLC27A3 |
| SLC27A4 |
| SLC27A5 |
| SLC27A6 |
| SLC29A2 |
| SLC29A3 |
| SLC29A4 |
| SLC2A1 |
| SLC2A13 |
| SLC2A3 |
| SLC2A4RG |
| SLC2A6 |
| SLC2A8 |
| SLC30A1 |
| SLC30A4 |
| SLC30A5 |
| SLC30A6 |
| SLC30A7 |
| SLC30A9 |
| SLC31A1 |
| SLC31A2 |
| SLC33A1 |
| SLC34A2 |
| SLC35A1 |
| SLC35A2 |
| SLC35A4 |
| SLC35B1 |
| SLC35B2 |
| SLC35B3 |
| SLC35B4 |
| SLC35C1 |
| SLC35C2 |
| SLC35D2 |
| SLC35E1 |

| |
|---|
| SLC35E4 |
| SLC35F2 |
| SLC35F5 |
| SLC35F6 |
| SLC35G2 |
| SLC36A1 |
| SLC36A4 |
| SLC37A3 |
| SLC37A4 |
| SLC38A10 |
| SLC38A2 |
| SLC38A4 |
| SLC38A5 |
| SLC38A6 |
| SLC38A7 |
| SLC39A1 |
| SLC39A10 |
| SLC39A11 |
| SLC39A14 |
| SLC39A3 |
| SLC39A4 |
| SLC39A5 |
| SLC39A7 |
| SLC39A8 |
| SLC39A9 |
| SLC3A2 |
| SLC40A1 |
| SLC41A1 |
| SLC41A2 |
| SLC41A3 |
| SLC43A1 |
| SLC43A2 |
| SLC44A1 |
| SLC44A2 |
| SLC44A3 |
| SLC44A4 |
| SLC45A1 |
| SLC45A4 |
| SLC46A1 |
| SLC47A1 |
| SLC47A2 |
| SLC48A1 |
| SLC49A3 |
| SLC49A4 |
| SLC4A1AP |

| |
|---|
| SLC4A2 |
| SLC4A3 |
| SLC50A1 |
| SLC51B |
| SLC52A1 |
| SLC5A12 |
| SLC5A3 |
| SLC5A6 |
| SLC66A1 |
| SLC66A2 |
| SLC66A3 |
| SLC6A12 |
| SLC6A15 |
| SLC6A6 |
| SLC6A8 |
| SLC7A11 |
| SLC7A5 |
| SLC7A6 |
| SLC7A6OS |
| SLC7A7 |
| SLC8A1 |
| SLC8A2 |
| SLC8B1 |
| SLC9A1 |
| SLC9A3R1 |
| SLC9A3R2 |
| SLC9A6 |
| SLC9A7 |
| SLC9A8 |
| SLC9B1 |
| SLCO2B1 |
| SLCO3A1 |
| SLCO4C1 |
| SLF1 |
| SLF2 |
| SLFN5 |
| SLIRP |
| SLITRK1 |
| SLITRK6 |
| SLK |
| SLMAP |
| SLTM |
| SLU7 |
| SLX1A |
| SMAD1 |

| |
|---|
| SMAD2 |
| SMAD3 |
| SMAD4 |
| SMAD6 |
| SMAD7 |
| SMAGP |
| SMAP1 |
| SMAP2 |
| SMARCA1 |
| SMARCA4 |
| SMARCA5 |
| SMARCC1 |
| SMARCC2 |
| SMARCD2 |
| SMARCD3 |
| SMARCE1 |
| SMC1A |
| SMC1B |
| SMC2 |
| SMC3 |
| SMC4 |
| SMC5 |
| SMC6 |
| SMCHD1 |
| SMCR8 |
| SMDT1 |
| SMG5 |
| SMG6 |
| SMG7 |
| SMG8 |
| SMG9 |
| SMIM1 |
| SMIM10L1 |
| SMIM10L2B |
| SMIM12 |
| SMIM13 |
| SMIM14 |
| SMIM15 |
| SMIM19 |
| SMIM20 |
| SMIM22 |
| SMIM26 |
| SMIM29 |
| SMIM3 |
| SMIM4 |

| |
|---|
| SMIM7 |
| SMN1 |
| SMNDC1 |
| SMO |
| SMOX |
| SMPD1 |
| SMPD2 |
| SMPD4 |
| SMPDL3A |
| SMPDL3B |
| SMS |
| SMTN |
| SMU1 |
| SMUG1 |
| SMURF1 |
| SMURF2 |
| SMYD2 |
| SMYD3 |
| SMYD4 |
| SMYD5 |
| SNAI1 |
| SNAP23 |
| SNAP29 |
| SNAP47 |
| SNAPC1 |
| SNAPC3 |
| SNAPC4 |
| SNAPC5 |
| SNAPIN |
| SND1 |
| SNF8 |
| SNIP1 |
| SNN |
| SNPH |
| SNRK |
| SNRNP200 |
| SNRNP25 |
| SNRNP27 |
| SNRNP35 |
| SNRNP40 |
| SNRNP48 |
| SNRNP70 |
| SNRPA1 |
| SNRPB |
| SNRPB2 |

| |
|---|
| SNRPC |
| SNRPD1 |
| SNRPD3 |
| SNRPE |
| SNRPF |
| SNRPG |
| SNRPN |
| SNTA1 |
| SNTB1 |
| SNTB2 |
| SNU13 |
| SNUPN |
| SNURF |
| SNW1 |
| SNX1 |
| SNX10 |
| SNX11 |
| SNX13 |
| SNX14 |
| SNX15 |
| SNX16 |
| SNX17 |
| SNX18 |
| SNX19 |
| SNX2 |
| SNX21 |
| SNX24 |
| SNX25 |
| SNX27 |
| SNX29 |
| SNX3 |
| SNX30 |
| SNX32 |
| SNX5 |
| SNX6 |
| SNX7 |
| SNX8 |
| SNX9 |
| SOAT1 |
| SOBP |
| SOCS1 |
| SOCS2 |
| SOCS3 |
| SOCS4 |
| SOCS5 |

| |
|---|
| SOCS6 |
| SOD1 |
| SOD2 |
| SOD3 |
| SOGA1 |
| SON |
| SORBS3 |
| SORD |
| SORL1 |
| SOS1 |
| SOS2 |
| SOSTDC1 |
| SOWAHA |
| SOWAHC |
| SOWAHD |
| SOX12 |
| SOX13 |
| SOX3 |
| SOX4 |
| SOX5 |
| SP100 |
| SP110 |
| SP140L |
| SP3 |
| SP4 |
| SPA17 |
| SPACA6 |
| SPAG1 |
| SPAG4 |
| SPAG5 |
| SPAG7 |
| SPAG8 |
| SPAG9 |
| SPARC |
| SPART |
| SPAST |
| SPATA13 |
| SPATA18 |
| SPATA2 |
| SPATA20 |
| SPATA24 |
| SPATA2L |
| SPATA33 |
| SPATA5 |
| SPATA5L1 |

| |
|---|
| SPATA6 |
| SPATA6L |
| SPATA7 |
| SPC24 |
| SPC25 |
| SPCS1 |
| SPCS2 |
| SPCS3 |
| SPDEF |
| SPDL1 |
| SPDYA |
| SPECC1L |
| SPEF2 |
| SPEN |
| SPG11 |
| SPG7 |
| SPHK1 |
| SPHK2 |
| SPI1 |
| SPICE1 |
| SPIDR |
| SPIN1 |
| SPIN2A |
| SPIN3 |
| SPIN4 |
| SPINDOC |
| SPINT1 |
| SPINT2 |
| SPIRE1 |
| SPNS1 |
| SPOCK1 |
| SPOCK2 |
| SPON1 |
| SPON2 |
| SPOP |
| SPOPL |
| SPOUT1 |
| SPPL2A |
| SPPL2B |
| SPPL3 |
| SPR |
| SPRED2 |
| SPRING1 |
| SPRR1B |
| SPRTN |

| |
|---|
| SPRY2 |
| SPRY3 |
| SPRY4 |
| SPRYD7 |
| SPSB1 |
| SPSB3 |
| SPSB4 |
| SPTAN1 |
| SPTB |
| SPTBN1 |
| SPTLC1 |
| SPTLC2 |
| SPTLC3 |
| SPTSSA |
| SPTY2D1 |
| SQLE |
| SQOR |
| SQSTM1 |
| SRA1 |
| SRBD1 |
| SRC |
| SRCAP |
| SRD5A3 |
| SREBF1 |
| SREBF2 |
| SREK1 |
| SREK1IP1 |
| SRF |
| SRFBP1 |
| SRGAP1 |
| SRGAP2 |
| SRGN |
| SRI |
| SRM |
| SRP14 |
| SRP19 |
| SRP54 |
| SRP68 |
| SRP72 |
| SRP9 |
| SRPK1 |
| SRPK2 |
| SRPK3 |
| SRPRA |
| SRPRB |

| |
|---|
| SRR |
| SRRD |
| SRRM1 |
| SRRM2 |
| SRRT |
| SRSF1 |
| SRSF10 |
| SRSF11 |
| SRSF12 |
| SRSF2 |
| SRSF3 |
| SRSF4 |
| SRSF5 |
| SRSF6 |
| SRSF7 |
| SRSF9 |
| SS18 |
| SS18L1 |
| SS18L2 |
| SSB |
| SSBP1 |
| SSBP2 |
| SSBP3 |
| SSBP4 |
| SSC5D |
| SSH1 |
| SSH2 |
| SSH3 |
| SSNA1 |
| SSPOP |
| SSR1 |
| SSR2 |
| SSR3 |
| SSR4 |
| SSRP1 |
| SSU72 |
| SSX2IP |
| ST13 |
| ST14 |
| ST3GAL2 |
| ST3GAL3 |
| ST3GAL4 |
| ST3GAL5 |
| ST3GAL6 |
| ST6GAL1 |

| |
|---|
| ST6GAL2 |
| ST6GALNAC2 |
| ST6GALNAC3 |
| ST6GALNAC4 |
| ST6GALNAC6 |
| ST7 |
| ST8SIA4 |
| ST8SIA5 |
| STAB1 |
| STAC |
| STAG1 |
| STAM |
| STAM2 |
| STAMBP |
| STAP2 |
| STAR |
| STARD10 |
| STARD13 |
| STARD3 |
| STARD3NL |
| STARD4 |
| STARD5 |
| STARD6 |
| STARD7 |
| STARD8 |
| STARD9 |
| STAT1 |
| STAT2 |
| STAT3 |
| STAT5A |
| STAU1 |
| STAU2 |
| STBD1 |
| STC1 |
| STEAP1 |
| STEAP3 |
| STEEP1 |
| STIL |
| STIM1 |
| STIM2 |
| STIMATE |
| STING1 |
| STIP1 |
| STK10 |
| STK11 |

| |
|---|
| STK11IP |
| STK16 |
| STK17A |
| STK17B |
| STK19 |
| STK24 |
| STK25 |
| STK26 |
| STK3 |
| STK33 |
| STK35 |
| STK36 |
| STK38 |
| STK38L |
| STK39 |
| STMN1 |
| STN1 |
| STOM |
| STOML1 |
| STOML2 |
| STON1 |
| STON2 |
| STOX1 |
| STRADB |
| STRBP |
| STRIP1 |
| STRN |
| STRN3 |
| STRN4 |
| STS |
| STT3A |
| STUB1 |
| STX10 |
| STX11 |
| STX12 |
| STX16 |
| STX17 |
| STX18 |
| STX19 |
| STX1A |
| STX1B |
| STX2 |
| STX4 |
| STX5 |
| STX6 |

| |
|---|
| STX7 |
| STX8 |
| STXBP3 |
| STXBP4 |
| STXBP5 |
| STXBP5L |
| STYX |
| SUB1 |
| SUCLA2 |
| SUCLG1 |
| SUCLG2 |
| SUCO |
| SUFU |
| SUGCT |
| SUGP1 |
| SUGP2 |
| SUGT1 |
| SULT1A1 |
| SULT1B1 |
| SUMF1 |
| SUMF2 |
| SUMO1 |
| SUMO2 |
| SUMO3 |
| SUN1 |
| SUN2 |
| SUOX |
| SUPT16H |
| SUPT20H |
| SUPT3H |
| SUPT4H1 |
| SUPT5H |
| SUPT6H |
| SUPT7L |
| SUPV3L1 |
| SURF1 |
| SURF2 |
| SURF4 |
| SUSD1 |
| SUSD3 |
| SUV39H1 |
| SUV39H2 |
| SUZ12 |
| SV2C |
| SVBP |

| |
|---|
| SVIL |
| SWAP70 |
| SWI5 |
| SWSAP1 |
| SWT1 |
| SYAP1 |
| SYBU |
| SYCE2 |
| SYDE1 |
| SYDE2 |
| SYF2 |
| SYK |
| SYN1 |
| SYNC |
| SYNCRIP |
| SYNE1 |
| SYNE2 |
| SYNE3 |
| SYNE4 |
| SYNGAP1 |
| SYNGR1 |
| SYNGR3 |
| SYNJ1 |
| SYNJ2 |
| SYNJ2BP |
| SYNM |
| SYNPO2 |
| SYNPO2L |
| SYNRG |
| SYP |
| SYPL1 |
| SYS1 |
| SYT1 |
| SYT10 |
| SYT11 |
| SYT17 |
| SYT7 |
| SYT9 |
| SYTL1 |
| SYTL2 |
| SYTL3 |
| SYTL4 |
| SYTL5 |
| SYVN1 |
| SZRD1 |

| |
|---|
| SZT2 |
| TAB2 |
| TAB3 |
| TAC1 |
| TAC4 |
| TACC2 |
| TACC3 |
| TACR1 |
| TACR3 |
| TADA1 |
| TADA2A |
| TADA2B |
| TADA3 |
| TAF1 |
| TAF10 |
| TAF11 |
| TAF12 |
| TAF13 |
| TAF15 |
| TAF1B |
| TAF1C |
| TAF1D |
| TAF3 |
| TAF4 |
| TAF4B |
| TAF5 |
| TAF5L |
| TAF6 |
| TAF6L |
| TAF7 |
| TAF8 |
| TAF9B |
| TAFA3 |
| TAGAP |
| TAGLN2 |
| TAL2 |
| TALDO1 |
| TAMALIN |
| TAMM41 |
| TANGO2 |
| TANGO6 |
| TAOK1 |
| TAOK2 |
| TAP1 |
| TAP2 |

| |
|---|
| TAPBPL |
| TAPT1 |
| TARBP1 |
| TARBP2 |
| TARDBP |
| TARM1 |
| TARS2 |
| TARS3 |
| TASOR |
| TASOR2 |
| TASP1 |
| TATDN1 |
| TATDN2 |
| TATDN3 |
| TAX1BP1 |
| TAX1BP3 |
| TBC1D1 |
| TBC1D10A |
| TBC1D10B |
| TBC1D10C |
| TBC1D12 |
| TBC1D13 |
| TBC1D14 |
| TBC1D15 |
| TBC1D16 |
| TBC1D17 |
| TBC1D19 |
| TBC1D2 |
| TBC1D20 |
| TBC1D22A |
| TBC1D22B |
| TBC1D24 |
| TBC1D2B |
| TBC1D30 |
| TBC1D31 |
| TBC1D32 |
| TBC1D4 |
| TBC1D7 |
| TBC1D8 |
| TBC1D8B |
| TBC1D9 |
| TBC1D9B |
| TBCA |
| TBCB |
| TBCC |

| |
|---|
| TBCD |
| TBK1 |
| TBKBP1 |
| TBL1X |
| TBL1XR1 |
| TBL2 |
| TBL3 |
| TBP |
| TBPL1 |
| TBRG1 |
| TBRG4 |
| TCAIM |
| TCAM1P |
| TCAP |
| TCEA1 |
| TCEA2 |
| TCEA3 |
| TCEAL1 |
| TCEAL3 |
| TCEAL4 |
| TCEAL8 |
| TCEAL9 |
| TCEANC |
| TCEANC2 |
| TCERG1 |
| TCF12 |
| TCF20 |
| TCF23 |
| TCF25 |
| TCF3 |
| TCF4 |
| TCF7 |
| TCF7L1 |
| TCF7L2 |
| TCHP |
| TCIM |
| TCN1 |
| TCOF1 |
| TCP1 |
| TCP11 |
| TCP11L2 |
| TCTA |
| TCTN1 |
| TCTN2 |
| TDG |

| |
|---|
| TDP1 |
| TDP2 |
| TDRD12 |
| TDRD3 |
| TDRD7 |
| TEAD1 |
| TEAD3 |
| TEC |
| TECPR1 |
| TECPR2 |
| TECR |
| TEFM |
| TEK |
| TELO2 |
| TEN1 |
| TENM2 |
| TENM3 |
| TENT2 |
| TENT4A |
| TENT4B |
| TENT5A |
| TENT5C |
| TEP1 |
| TERF1 |
| TERF2 |
| TERF2IP |
| TERT |
| TES |
| TESC |
| TESK1 |
| TESK2 |
| TESPA1 |
| TET1 |
| TET2 |
| TET3 |
| TEX14 |
| TEX15 |
| TEX2 |
| TEX261 |
| TEX264 |
| TEX45 |
| TF |
| TFAM |
| TFAP4 |
| TFB1M |

| |
|---|
| TFB2M |
| TFCP2 |
| TFDP1 |
| TFE3 |
| TFEB |
| TFG |
| TFIP11 |
| TFPI2 |
| TFPT |
| TFRC |
| TGDS |
| TGFB1 |
| TGFBI |
| TGFBR2 |
| TGFBRAP1 |
| TGIF1 |
| TGIF2 |
| TGM2 |
| TGS1 |
| THAP1 |
| THAP11 |
| THAP12 |
| THAP3 |
| THAP4 |
| THAP5 |
| THAP6 |
| THAP7 |
| THAP9 |
| THBS1 |
| THBS2 |
| THBS3 |
| THEM6 |
| THEMIS2 |
| THG1L |
| THNSL1 |
| THNSL2 |
| THOC1 |
| THOC2 |
| THOC3 |
| THOC5 |
| THOC6 |
| THOC7 |
| THOP1 |
| THRA |
| THRAP3 |

| |
|---|
| THSD4 |
| THSD7A |
| THTPA |
| THUMPD1 |
| THUMPD2 |
| THUMPD3 |
| THYN1 |
| TIA1 |
| TIAL1 |
| TIAM1 |
| TIAM2 |
| TICAM2 |
| TIFA |
| TIGAR |
| TIGD1 |
| TIGD2 |
| TIGD3 |
| TIGD4 |
| TIGD5 |
| TIGD6 |
| TIGD7 |
| TIMELESS |
| TIMM10 |
| TIMM10B |
| TIMM13 |
| TIMM17A |
| TIMM17B |
| TIMM21 |
| TIMM23 |
| TIMM29 |
| TIMM44 |
| TIMM50 |
| TIMM8B |
| TIMM9 |
| TIMMDC1 |
| TIMP1 |
| TIMP2 |
| TINF2 |
| TIPARP |
| TIPIN |
| TIPRL |
| TJAP1 |
| TJP1 |
| TJP2 |
| TJP3 |

| |
|---|
| TK1 |
| TK2 |
| TKT |
| TLCD1 |
| TLCD3A |
| TLCD4 |
| TLE1 |
| TLE2 |
| TLE3 |
| TLE4 |
| TLE5 |
| TLK1 |
| TLK2 |
| TLN1 |
| TLNRD1 |
| TLR10 |
| TLR3 |
| TLR4 |
| TLR6 |
| TLR7 |
| TLR8 |
| TM2D1 |
| TM2D2 |
| TM2D3 |
| TM4SF1 |
| TM4SF20 |
| TM7SF2 |
| TM7SF3 |
| TM9SF1 |
| TM9SF2 |
| TM9SF4 |
| TMA16 |
| TMBIM1 |
| TMBIM4 |
| TMBIM6 |
| TMC4 |
| TMC5 |
| TMC6 |
| TMC8 |
| TMCC1 |
| TMCC2 |
| TMCC3 |
| TMCO1 |
| TMCO4 |
| TMCO6 |

| |
|---|
| TMED1 |
| TMED10 |
| TMED2 |
| TMED3 |
| TMED4 |
| TMED5 |
| TMED7 |
| TMED9 |
| TMEM101 |
| TMEM104 |
| TMEM106A |
| TMEM106C |
| TMEM108 |
| TMEM109 |
| TMEM11 |
| TMEM115 |
| TMEM117 |
| TMEM120A |
| TMEM120B |
| TMEM121 |
| TMEM123 |
| TMEM125 |
| TMEM126B |
| TMEM127 |
| TMEM128 |
| TMEM129 |
| TMEM131 |
| TMEM131L |
| TMEM132A |
| TMEM134 |
| TMEM135 |
| TMEM138 |
| TMEM139 |
| TMEM140 |
| TMEM141 |
| TMEM143 |
| TMEM144 |
| TMEM147 |
| TMEM14A |
| TMEM14C |
| TMEM150A |
| TMEM151B |
| TMEM160 |
| TMEM161A |
| TMEM161B |

| |
|---|
| TMEM164 |
| TMEM165 |
| TMEM167B |
| TMEM17 |
| TMEM170A |
| TMEM170B |
| TMEM171 |
| TMEM175 |
| TMEM177 |
| TMEM178A |
| TMEM179B |
| TMEM18 |
| TMEM183A |
| TMEM184A |
| TMEM184C |
| TMEM185A |
| TMEM185B |
| TMEM186 |
| TMEM187 |
| TMEM19 |
| TMEM192 |
| TMEM198 |
| TMEM199 |
| TMEM201 |
| TMEM203 |
| TMEM208 |
| TMEM209 |
| TMEM219 |
| TMEM220 |
| TMEM222 |
| TMEM223 |
| TMEM229B |
| TMEM230 |
| TMEM231 |
| TMEM234 |
| TMEM236 |
| TMEM238 |
| TMEM240 |
| TMEM241 |
| TMEM242 |
| TMEM243 |
| TMEM245 |
| TMEM25 |
| TMEM250 |
| TMEM251 |

| |
|---|
| TMEM254 |
| TMEM256 |
| TMEM259 |
| TMEM26 |
| TMEM260 |
| TMEM263 |
| TMEM266 |
| TMEM267 |
| TMEM268 |
| TMEM30A |
| TMEM30B |
| TMEM33 |
| TMEM35A |
| TMEM35B |
| TMEM37 |
| TMEM38A |
| TMEM38B |
| TMEM39A |
| TMEM39B |
| TMEM41A |
| TMEM41B |
| TMEM43 |
| TMEM45A |
| TMEM45B |
| TMEM47 |
| TMEM50A |
| TMEM50B |
| TMEM51 |
| TMEM52 |
| TMEM53 |
| TMEM54 |
| TMEM59 |
| TMEM59L |
| TMEM60 |
| TMEM61 |
| TMEM62 |
| TMEM64 |
| TMEM65 |
| TMEM67 |
| TMEM68 |
| TMEM70 |
| TMEM81 |
| TMEM86A |
| TMEM87A |
| TMEM87B |

| |
|---|
| TMEM88 |
| TMEM89 |
| TMEM8B |
| TMEM9 |
| TMEM94 |
| TMEM97 |
| TMEM98 |
| TMEM9B |
| TMF1 |
| TMLHE |
| TMOD3 |
| TMPO |
| TMPPE |
| TMPRSS2 |
| TMPRSS9 |
| TMSB4X |
| TMTC1 |
| TMTC3 |
| TMTC4 |
| TMUB1 |
| TMUB2 |
| TMX1 |
| TMX2 |
| TMX4 |
| TNC |
| TNFAIP1 |
| TNFAIP2 |
| TNFAIP3 |
| TNFAIP8 |
| TNFAIP8L2 |
| TNFRSF10A |
| TNFRSF11A |
| TNFRSF12A |
| TNFRSF14 |
| TNFRSF1A |
| TNFRSF1B |
| TNFRSF21 |
| TNFSF12 |
| TNFSF9 |
| TNIK |
| TNIP2 |
| TNK1 |
| TNKS |
| TNKS1BP1 |
| TNNC1 |

| |
|---|
| TNNT1 |
| TNPO2 |
| TNPO3 |
| TNRC18 |
| TNRC6A |
| TNRC6B |
| TNS2 |
| TNS3 |
| TNXB |
| TOB2 |
| TOE1 |
| TOGARAM1 |
| TOGARAM2 |
| TOLLIP |
| TOM1 |
| TOM1L1 |
| TOM1L2 |
| TOMM20 |
| TOMM22 |
| TOMM34 |
| TOMM40 |
| TOMM40L |
| TOMM7 |
| TOMM70 |
| TONSL |
| TOP1 |
| TOP2A |
| TOP2B |
| TOP3B |
| TOPBP1 |
| TOPORS |
| TOR1A |
| TOR1AIP1 |
| TOR1AIP2 |
| TOR1B |
| TOR2A |
| TOR3A |
| TOR4A |
| TOX2 |
| TOX4 |
| TP53 |
| TP53BP1 |
| TP53BP2 |
| TP53I13 |
| TP53I3 |

| |
|---|
| TP53INP1 |
| TP53INP2 |
| TP53RK |
| TP53TG5 |
| TPBG |
| TPBGL |
| TPCN2 |
| TPD52 |
| TPD52L2 |
| TPGS1 |
| TPGS2 |
| TPH1 |
| TPK1 |
| TPM1 |
| TPM2 |
| TPM3 |
| TPM4 |
| TPMT |
| TPP1 |
| TPP2 |
| TPPP |
| TPR |
| TPRA1 |
| TPRN |
| TPST1 |
| TPST2 |
| TPT1 |
| TPX2 |
| TRA2A |
| TRA2B |
| TRADD |
| TRAF1 |
| TRAF2 |
| TRAF3 |
| TRAF3IP2 |
| TRAF4 |
| TRAF5 |
| TRAF6 |
| TRAF7 |
| TRAFD1 |
| TRAIP |
| TRAK1 |
| TRAK2 |
| TRAM1L1 |
| TRAM2 |

| |
|---|
| TRANK1 |
| TRAP1 |
| TRAPPC1 |
| TRAPPC10 |
| TRAPPC11 |
| TRAPPC12 |
| TRAPPC13 |
| TRAPPC14 |
| TRAPPC3 |
| TRAPPC4 |
| TRAPPC5 |
| TRAPPC6B |
| TRAPPC8 |
| TRAPPC9 |
| TRDMT1 |
| TRIAP1 |
| TRIB1 |
| TRIB2 |
| TRIB3 |
| TRIL |
| TRIM11 |
| TRIM14 |
| TRIM16 |
| TRIM2 |
| TRIM21 |
| TRIM22 |
| TRIM23 |
| TRIM24 |
| TRIM25 |
| TRIM26 |
| TRIM27 |
| TRIM28 |
| TRIM3 |
| TRIM32 |
| TRIM33 |
| TRIM34 |
| TRIM35 |
| TRIM36 |
| TRIM38 |
| TRIM39 |
| TRIM41 |
| TRIM45 |
| TRIM47 |
| TRIM5 |
| TRIM50 |

| |
|---|
| TRIM56 |
| TRIM59 |
| TRIM6 |
| TRIM62 |
| TRIM65 |
| TRIM66 |
| TRIM68 |
| TRIM7 |
| TRIM8 |
| TRIM9 |
| TRIO |
| TRIP10 |
| TRIP11 |
| TRIP12 |
| TRIP13 |
| TRIP4 |
| TRIQK |
| TRIR |
| TRIT1 |
| TRMO |
| TRMT1 |
| TRMT10A |
| TRMT10B |
| TRMT112 |
| TRMT12 |
| TRMT44 |
| TRMT5 |
| TRMT6 |
| TRMT61A |
| TRMT61B |
| TRNAU1AP |
| TRNP1 |
| TRNT1 |
| TROAP |
| TRPC1 |
| TRPC4 |
| TRPC4AP |
| TRPM4 |
| TRPS1 |
| TRPT1 |
| TRPV1 |
| TRPV2 |
| TRRAP |
| TRUB1 |
| TRUB2 |

| |
|---|
| TSC2 |
| TSC22D1 |
| TSC22D3 |
| TSC22D4 |
| TSEN15 |
| TSFM |
| TSG101 |
| TSGA10 |
| TSHZ1 |
| TSHZ3 |
| TSKS |
| TSKU |
| TSN |
| TSNARE1 |
| TSNAX |
| TSPAN1 |
| TSPAN14 |
| TSPAN15 |
| TSPAN17 |
| TSPAN3 |
| TSPAN31 |
| TSPAN5 |
| TSPAN6 |
| TSPAN7 |
| TSPAN8 |
| TSPAN9 |
| TSPO |
| TSPY26P |
| TSPYL1 |
| TSPYL2 |
| TSPYL4 |
| TSPYL5 |
| TSR1 |
| TSR2 |
| TSR3 |
| TSSC4 |
| TSSK3 |
| TST |
| TSTD1 |
| TSTD2 |
| TSTD3 |
| TTBK2 |
| TTC1 |
| TTC12 |
| TTC13 |

| |
|---|
| TTC14 |
| TTC17 |
| TTC19 |
| TTC21B |
| TTC23 |
| TTC26 |
| TTC27 |
| TTC28 |
| TTC30A |
| TTC30B |
| TTC31 |
| TTC32 |
| TTC33 |
| TTC38 |
| TTC39A |
| TTC39B |
| TTC39C |
| TTC4 |
| TTC5 |
| TTC7A |
| TTC7B |
| TTC8 |
| TTC9C |
| TTF1 |
| TTI1 |
| TTI2 |
| TTK |
| TTL |
| TTLL1 |
| TTLL11 |
| TTLL12 |
| TTLL3 |
| TTLL5 |
| TTLL7 |
| TTN |
| TTPAL |
| TTYH2 |
| TTYH3 |
| TUB |
| TUBA1B |
| TUBA1C |
| TUBA4A |
| TUBA8 |
| TUBB2A |
| TUBB2B |

| |
|---|
| TUBB3 |
| TUBD1 |
| TUBE1 |
| TUBG1 |
| TUBG2 |
| TUBGCP2 |
| TUBGCP3 |
| TUBGCP4 |
| TUBGCP5 |
| TUBGCP6 |
| TUFM |
| TULP3 |
| TULP4 |
| TUSC1 |
| TUSC2 |
| TUSC3 |
| TUT1 |
| TUT4 |
| TVP23B |
| TWF1 |
| TWF2 |
| TWIST1 |
| TWNK |
| TWSG1 |
| TXLNA |
| TXN |
| TXN2 |
| TXNDC11 |
| TXNDC15 |
| TXNDC16 |
| TXNDC17 |
| TXNDC5 |
| TXNDC9 |
| TXNIP |
| TXNL1 |
| TXNL4A |
| TXNL4B |
| TXNRD1 |
| TXNRD3 |
| TYMP |
| TYMS |
| TYRO3 |
| TYSND1 |
| TYW1 |
| TYW3 |

| |
|---|
| TYW5 |
| U2AF1L4 |
| U2AF2 |
| U2SURP |
| UACA |
| UAP1L1 |
| UBA1 |
| UBA2 |
| UBA3 |
| UBA5 |
| UBA52 |
| UBA7 |
| UBAC2 |
| UBALD1 |
| UBALD2 |
| UBAP2 |
| UBAP2L |
| UBASH3B |
| UBB |
| UBC |
| UBE2A |
| UBE2B |
| UBE2C |
| UBE2D1 |
| UBE2D2 |
| UBE2D3 |
| UBE2D4 |
| UBE2E1 |
| UBE2E2 |
| UBE2E3 |
| UBE2F |
| UBE2G2 |
| UBE2H |
| UBE2J1 |
| UBE2J2 |
| UBE2K |
| UBE2L3 |
| UBE2L6 |
| UBE2M |
| UBE2N |
| UBE2O |
| UBE2Q1 |
| UBE2Q2 |
| UBE2R2 |
| UBE2S |

| |
|---|
| UBE2T |
| UBE2V1 |
| UBE2V2 |
| UBE2W |
| UBE2Z |
| UBE3B |
| UBE3C |
| UBE3D |
| UBE4A |
| UBE4B |
| UBFD1 |
| UBIAD1 |
| UBL3 |
| UBL4A |
| UBL5 |
| UBL7 |
| UBLCP1 |
| UBN2 |
| UBP1 |
| UBQLN1 |
| UBQLN2 |
| UBQLN4 |
| UBR1 |
| UBR2 |
| UBR3 |
| UBR4 |
| UBR5 |
| UBR7 |
| UBTD1 |
| UBTD2 |
| UBTF |
| UBXN1 |
| UBXN2A |
| UBXN2B |
| UBXN4 |
| UBXN6 |
| UBXN7 |
| UBXN8 |
| UCHL1 |
| UCHL3 |
| UCHL5 |
| UCK1 |
| UCK2 |
| UCKL1 |
| UCN2 |

| |
|---|
| UCP2 |
| UEVLD |
| UFD1 |
| UFL1 |
| UFM1 |
| UFSP1 |
| UGCG |
| UGDH |
| UGGT2 |
| UHRF1 |
| UHRF1BP1L |
| UHRF2 |
| UIMC1 |
| ULBP3 |
| ULK1 |
| ULK2 |
| ULK3 |
| ULK4 |
| UMAD1 |
| UMPS |
| UNC119 |
| UNC119B |
| UNC13B |
| UNC13D |
| UNC45A |
| UNC50 |
| UNC5B |
| UNC5C |
| UNC5CL |
| UNC5D |
| UNC93B1 |
| UNG |
| UNK |
| UNKL |
| UPB1 |
| UPF1 |
| UPF2 |
| UPF3A |
| UPF3B |
| UPK1B |
| UPP1 |
| UPRT |
| UQCC1 |
| UQCC2 |
| UQCC3 |

| |
|---|
| UQCR10 |
| UQCRC1 |
| UQCRC2 |
| UQCRFS1 |
| UQCRH |
| UQCRQ |
| URB1 |
| URB2 |
| URGCP |
| URI1 |
| URM1 |
| UROD |
| UROS |
| USB1 |
| USE1 |
| USF1 |
| USF2 |
| USF3 |
| USO1 |
| USP1 |
| USP10 |
| USP11 |
| USP12 |
| USP14 |
| USP15 |
| USP16 |
| USP18 |
| USP19 |
| USP2 |
| USP20 |
| USP21 |
| USP22 |
| USP24 |
| USP25 |
| USP27X |
| USP28 |
| USP30 |
| USP31 |
| USP33 |
| USP34 |
| USP35 |
| USP37 |
| USP38 |
| USP39 |
| USP4 |

| |
|---|
| USP40 |
| USP42 |
| USP44 |
| USP45 |
| USP46 |
| USP47 |
| USP48 |
| USP49 |
| USP5 |
| USP53 |
| USP54 |
| USP7 |
| USP9X |
| USPL1 |
| UST |
| UTP11 |
| UTP14A |
| UTP15 |
| UTP18 |
| UTP20 |
| UTP23 |
| UTP25 |
| UTP3 |
| UTP4 |
| UTP6 |
| UVRAG |
| UVSSA |
| UXS1 |
| UXT |
| VAC14 |
| VAMP3 |
| VAMP4 |
| VAMP5 |
| VAMP7 |
| VAMP8 |
| VANGL1 |
| VANGL2 |
| VAPA |
| VAPB |
| VARS1 |
| VARS2 |
| VASH1 |
| VASN |
| VASP |
| VAT1 |

| |
|---|
| VAV1 |
| VBP1 |
| VCAM1 |
| VCAN |
| VCPKMT |
| VDAC1 |
| VDAC2 |
| VDAC3 |
| VDR |
| VEGFA |
| VEGFB |
| VEGFC |
| VEPH1 |
| VEZF1 |
| VEZT |
| VGF |
| VGLL4 |
| VHL |
| VIL1 |
| VILL |
| VIM |
| VIRMA |
| VKORC1 |
| VKORC1L1 |
| VMA21 |
| VMP1 |
| VPS11 |
| VPS13A |
| VPS13B |
| VPS13C |
| VPS13D |
| VPS16 |
| VPS18 |
| VPS25 |
| VPS26B |
| VPS26C |
| VPS28 |
| VPS29 |
| VPS33A |
| VPS33B |
| VPS35 |
| VPS35L |
| VPS36 |
| VPS37A |
| VPS37D |

| |
|---|
| VPS39 |
| VPS41 |
| VPS45 |
| VPS4A |
| VPS4B |
| VPS50 |
| VPS51 |
| VPS52 |
| VPS53 |
| VPS54 |
| VPS72 |
| VPS8 |
| VPS9D1 |
| VRK1 |
| VSIG10 |
| VSIG10L |
| VSIR |
| VSTM4 |
| VTA1 |
| VTI1A |
| VTI1B |
| VWA1 |
| VWA5A |
| VWA5B2 |
| VWA7 |
| VWF |
| WAC |
| WAPL |
| WARS1 |
| WARS2 |
| WASF1 |
| WASF2 |
| WASF3 |
| WASHC1 |
| WASHC3 |
| WASHC4 |
| WASHC5 |
| WASL |
| WBP1 |
| WBP11 |
| WBP1L |
| WBP2 |
| WDCP |
| WDFY1 |
| WDFY2 |

| |
|---|
| WDHD1 |
| WDPCP |
| WDR1 |
| WDR11 |
| WDR12 |
| WDR13 |
| WDR18 |
| WDR20 |
| WDR24 |
| WDR25 |
| WDR26 |
| WDR27 |
| WDR3 |
| WDR35 |
| WDR36 |
| WDR37 |
| WDR4 |
| WDR41 |
| WDR44 |
| WDR45 |
| WDR45B |
| WDR46 |
| WDR48 |
| WDR5 |
| WDR53 |
| WDR54 |
| WDR55 |
| WDR59 |
| WDR5B |
| WDR6 |
| WDR61 |
| WDR62 |
| WDR7 |
| WDR70 |
| WDR73 |
| WDR74 |
| WDR75 |
| WDR76 |
| WDR77 |
| WDR81 |
| WDR82 |
| WDR83 |
| WDR83OS |
| WDR86 |
| WDR90 |

| |
|---|
| WDR91 |
| WDR97 |
| WDTC1 |
| WEE1 |
| WFDC8 |
| WFIKKN1 |
| WFS1 |
| WHAMM |
| WIF1 |
| WIPI1 |
| WIZ |
| WLS |
| WNK1 |
| WNK2 |
| WNK3 |
| WNK4 |
| WNT10B |
| WNT5A |
| WNT6 |
| WNT9A |
| WRAP73 |
| WRNIP1 |
| WSB2 |
| WT1 |
| WTAP |
| WWC1 |
| WWC2 |
| WWC3 |
| WWOX |
| WWP1 |
| WWP2 |
| WWTR1 |
| XAB2 |
| XAF1 |
| XIAP |
| XK |
| XKR4 |
| XKR5 |
| XKR8 |
| XPA |
| XPC |
| XPNPEP1 |
| XPNPEP3 |
| XPO1 |
| XPO4 |

| |
|---|
| XPO5 |
| XPO6 |
| XPO7 |
| XPOT |
| XPR1 |
| XRCC1 |
| XRCC3 |
| XRCC4 |
| XRCC5 |
| XRCC6 |
| XRN1 |
| XRN2 |
| XXYLT1 |
| XYLT2 |
| YAE1 |
| YAF2 |
| YAP1 |
| YARS1 |
| YARS2 |
| YBEY |
| YBX1 |
| YBX2 |
| YBX3 |
| YEATS2 |
| YEATS4 |
| YES1 |
| YIF1A |
| YIPF1 |
| YIPF2 |
| YIPF3 |
| YIPF4 |
| YIPF5 |
| YIPF6 |
| YJU2 |
| YKT6 |
| YLPM1 |
| YME1L1 |
| YPEL1 |
| YPEL3 |
| YRDC |
| YTHDC1 |
| YTHDC2 |
| YTHDF1 |
| YTHDF2 |
| YTHDF3 |

| |
|---|
| YWHAB |
| YWHAE |
| YWHAG |
| YWHAH |
| YWHAQ |
| YWHAZ |
| YY1 |
| ZADH2 |
| ZAP70 |
| ZAR1L |
| ZBED1 |
| ZBED2 |
| ZBED3 |
| ZBED4 |
| ZBED5 |
| ZBED6 |
| ZBED8 |
| ZBED9 |
| ZBTB1 |
| ZBTB11 |
| ZBTB12 |
| ZBTB14 |
| ZBTB16 |
| ZBTB17 |
| ZBTB18 |
| ZBTB2 |
| ZBTB20 |
| ZBTB21 |
| ZBTB22 |
| ZBTB24 |
| ZBTB25 |
| ZBTB3 |
| ZBTB33 |
| ZBTB34 |
| ZBTB37 |
| ZBTB38 |
| ZBTB39 |
| ZBTB4 |
| ZBTB40 |
| ZBTB41 |
| ZBTB42 |
| ZBTB43 |
| ZBTB44 |
| ZBTB45 |
| ZBTB47 |

| |
|---|
| ZBTB48 |
| ZBTB49 |
| ZBTB5 |
| ZBTB6 |
| ZBTB7A |
| ZBTB7B |
| ZBTB8A |
| ZBTB8OS |
| ZC2HC1C |
| ZC3H12A |
| ZC3H12C |
| ZC3H13 |
| ZC3H14 |
| ZC3H15 |
| ZC3H18 |
| ZC3H3 |
| ZC3H4 |
| ZC3H6 |
| ZC3H7A |
| ZC3H7B |
| ZC3H8 |
| ZC3HAV1 |
| ZC3HC1 |
| ZC4H2 |
| ZCCHC10 |
| ZCCHC14 |
| ZCCHC17 |
| ZCCHC18 |
| ZCCHC2 |
| ZCCHC24 |
| ZCCHC3 |
| ZCCHC4 |
| ZCCHC7 |
| ZCCHC8 |
| ZCRB1 |
| ZCWPW1 |
| ZCWPW2 |
| ZDHHC1 |
| ZDHHC12 |
| ZDHHC13 |
| ZDHHC14 |
| ZDHHC15 |
| ZDHHC16 |
| ZDHHC17 |
| ZDHHC18 |

| |
|---|
| ZDHHC2 |
| ZDHHC20 |
| ZDHHC23 |
| ZDHHC24 |
| ZDHHC4 |
| ZDHHC5 |
| ZDHHC7 |
| ZDHHC8 |
| ZDHHC9 |
| ZEB1 |
| ZEB2 |
| ZER1 |
| ZFAND1 |
| ZFAND2A |
| ZFAND2B |
| ZFAND3 |
| ZFAND5 |
| ZFAND6 |
| ZFAT |
| ZFHX2 |
| ZFHX3 |
| ZFP1 |
| ZFP2 |
| ZFP28 |
| ZFP36L1 |
| ZFP36L2 |
| ZFP41 |
| ZFP42 |
| ZFP64 |
| ZFP69 |
| ZFP69B |
| ZFP82 |
| ZFP91 |
| ZFPL1 |
| ZFPM1 |
| ZFR |
| ZFR2 |
| ZFX |
| ZFYVE1 |
| ZFYVE16 |
| ZFYVE19 |
| ZFYVE21 |
| ZFYVE26 |
| ZFYVE27 |
| ZFYVE9 |

| |
|---|
| ZG16 |
| ZGPAT |
| ZGRF1 |
| ZHX3 |
| ZKSCAN1 |
| ZKSCAN2 |
| ZKSCAN4 |
| ZKSCAN5 |
| ZKSCAN7 |
| ZKSCAN8 |
| ZMAT1 |
| ZMAT2 |
| ZMAT3 |
| ZMAT5 |
| ZMIZ1 |
| ZMIZ2 |
| ZMPSTE24 |
| ZMYM1 |
| ZMYM2 |
| ZMYM3 |
| ZMYM4 |
| ZMYM5 |
| ZMYND10 |
| ZMYND11 |
| ZMYND12 |
| ZMYND8 |
| ZNF106 |
| ZNF112 |
| ZNF12 |
| ZNF124 |
| ZNF131 |
| ZNF132 |
| ZNF133 |
| ZNF134 |
| ZNF135 |
| ZNF136 |
| ZNF140 |
| ZNF142 |
| ZNF143 |
| ZNF146 |
| ZNF148 |
| ZNF157 |
| ZNF165 |
| ZNF169 |
| ZNF174 |

| |
|---|
| ZNF175 |
| ZNF18 |
| ZNF180 |
| ZNF182 |
| ZNF184 |
| ZNF19 |
| ZNF197 |
| ZNF2 |
| ZNF205 |
| ZNF207 |
| ZNF208 |
| ZNF211 |
| ZNF212 |
| ZNF213 |
| ZNF217 |
| ZNF219 |
| ZNF227 |
| ZNF229 |
| ZNF236 |
| ZNF239 |
| ZNF24 |
| ZNF248 |
| ZNF25 |
| ZNF250 |
| ZNF251 |
| ZNF256 |
| ZNF26 |
| ZNF260 |
| ZNF263 |
| ZNF268 |
| ZNF271P |
| ZNF274 |
| ZNF275 |
| ZNF276 |
| ZNF277 |
| ZNF280A |
| ZNF280B |
| ZNF281 |
| ZNF282 |
| ZNF283 |
| ZNF286A |
| ZNF287 |
| ZNF292 |
| ZNF296 |
| ZNF3 |

| |
|---|
| ZNF300 |
| ZNF304 |
| ZNF316 |
| ZNF317 |
| ZNF318 |
| ZNF319 |
| ZNF32 |
| ZNF322 |
| ZNF324 |
| ZNF326 |
| ZNF329 |
| ZNF330 |
| ZNF331 |
| ZNF333 |
| ZNF34 |
| ZNF341 |
| ZNF346 |
| ZNF347 |
| ZNF35 |
| ZNF350 |
| ZNF354A |
| ZNF358 |
| ZNF362 |
| ZNF365 |
| ZNF367 |
| ZNF37A |
| ZNF382 |
| ZNF383 |
| ZNF384 |
| ZNF385A |
| ZNF385B |
| ZNF385D |
| ZNF391 |
| ZNF394 |
| ZNF395 |
| ZNF397 |
| ZNF398 |
| ZNF404 |
| ZNF407 |
| ZNF408 |
| ZNF41 |
| ZNF410 |
| ZNF414 |
| ZNF416 |
| ZNF418 |

| |
|---|
| ZNF419 |
| ZNF420 |
| ZNF423 |
| ZNF426 |
| ZNF428 |
| ZNF432 |
| ZNF436 |
| ZNF438 |
| ZNF445 |
| ZNF446 |
| ZNF449 |
| ZNF451 |
| ZNF462 |
| ZNF467 |
| ZNF470 |
| ZNF48 |
| ZNF483 |
| ZNF484 |
| ZNF485 |
| ZNF496 |
| ZNF501 |
| ZNF503 |
| ZNF507 |
| ZNF510 |
| ZNF511 |
| ZNF512 |
| ZNF512B |
| ZNF513 |
| ZNF516 |
| ZNF518A |
| ZNF518B |
| ZNF524 |
| ZNF526 |
| ZNF530 |
| ZNF544 |
| ZNF548 |
| ZNF549 |
| ZNF550 |
| ZNF551 |
| ZNF554 |
| ZNF555 |
| ZNF564 |
| ZNF565 |
| ZNF566 |
| ZNF567 |

| |
|---|
| ZNF568 |
| ZNF57 |
| ZNF570 |
| ZNF574 |
| ZNF575 |
| ZNF576 |
| ZNF579 |
| ZNF581 |
| ZNF583 |
| ZNF584 |
| ZNF585A |
| ZNF586 |
| ZNF593 |
| ZNF594 |
| ZNF596 |
| ZNF597 |
| ZNF598 |
| ZNF599 |
| ZNF608 |
| ZNF609 |
| ZNF613 |
| ZNF614 |
| ZNF618 |
| ZNF619 |
| ZNF621 |
| ZNF622 |
| ZNF623 |
| ZNF628 |
| ZNF629 |
| ZNF638 |
| ZNF641 |
| ZNF644 |
| ZNF646 |
| ZNF652 |
| ZNF653 |
| ZNF654 |
| ZNF655 |
| ZNF658 |
| ZNF660 |
| ZNF664 |
| ZNF667 |
| ZNF668 |
| ZNF671 |
| ZNF672 |
| ZNF674 |

| |
|---|
| ZNF677 |
| ZNF684 |
| ZNF687 |
| ZNF688 |
| ZNF689 |
| ZNF691 |
| ZNF692 |
| ZNF696 |
| ZNF697 |
| ZNF703 |
| ZNF704 |
| ZNF706 |
| ZNF707 |
| ZNF709 |
| ZNF71 |
| ZNF710 |
| ZNF711 |
| ZNF713 |
| ZNF717 |
| ZNF74 |
| ZNF740 |
| ZNF746 |
| ZNF750 |
| ZNF75A |
| ZNF75D |
| ZNF76 |
| ZNF764 |
| ZNF768 |
| ZNF77 |
| ZNF771 |
| ZNF772 |
| ZNF775 |
| ZNF777 |
| ZNF780B |
| ZNF782 |
| ZNF783 |
| ZNF784 |
| ZNF785 |
| ZNF786 |
| ZNF787 |
| ZNF789 |
| ZNF79 |
| ZNF790 |
| ZNF792 |
| ZNF793 |

| |
|---|
| ZNF8 |
| ZNF800 |
| ZNF805 |
| ZNF81 |
| ZNF813 |
| ZNF814 |
| ZNF821 |
| ZNF827 |
| ZNF829 |
| ZNF835 |
| ZNF839 |
| ZNF84 |
| ZNF846 |
| ZNF850 |
| ZNF853 |
| ZNF865 |
| ZNF875 |
| ZNF891 |
| ZNFX1 |
| ZNHIT1 |
| ZNHIT2 |
| ZNHIT3 |
| ZNRD2 |
| ZNRF1 |
| ZNRF3 |
| ZPR1 |
| ZRANB1 |
| ZRANB2 |
| ZRANB3 |
| ZRSR2 |
| ZSCAN12 |
| ZSCAN16 |
| ZSCAN18 |
| ZSCAN2 |
| ZSCAN20 |
| ZSCAN21 |
| ZSCAN22 |
| ZSCAN23 |
| ZSCAN26 |
| ZSCAN29 |
| ZSCAN30 |
| ZSCAN31 |
| ZSCAN32 |
| ZSCAN4 |
| ZSCAN9 |

| |
|---|
| ZSWIM1 |
| ZSWIM3 |
| ZSWIM4 |
| ZSWIM7 |
| ZSWIM8 |
| ZSWIM9 |
| ZUP1 |
| ZW10 |
| ZWILCH |
| ZWINT |
| ZXDC |
| ZYG11B |
| ZYX |
| ZZEF1 |
| ZZZ3 |

**Supplemental List 3**

| Gene Name |
|---|
| A2M |
| AAAS |
| AACS |
| AADAC |
| AAK1 |
| AAMP |
| AARSD1 |
| AASDH |
| AASDHPPT |
| AASS |
| AATF |
| AATK |
| ABCA1 |
| ABCA12 |
| ABCA2 |
| ABCA3 |
| ABCA4 |
| ABCA5 |
| ABCA6 |
| ABCA7 |
| ABCA9 |
| ABCB1 |
| ABCB10 |
| ABCB6 |
| ABCB7 |
| ABCB8 |
| ABCB9 |
| ABCC1 |
| ABCC10 |
| ABCC11 |
| ABCC3 |
| ABCC4 |
| ABCD1 |
| ABCD3 |
| ABCD4 |
| ABCE1 |
| ABCF2 |
| ABCF3 |
| ABCG1 |
| ABCG2 |
| ABHD1 |
| ABHD10 |
| ABHD11 |

| |
|---|
| ABHD12 |
| ABHD13 |
| ABHD2 |
| ABHD3 |
| ABHD4 |
| ABHD5 |
| ABHD6 |
| ABHD8 |
| ABI1 |
| ABI3BP |
| ABL2 |
| ABR |
| ABT1 |
| ABTB1 |
| ABTB2 |
| ACAA1 |
| ACAA2 |
| ACACA |
| ACACB |
| ACAD10 |
| ACAD11 |
| ACAD8 |
| ACADS |
| ACADSB |
| ACADVL |
| ACAN |
| ACAP1 |
| ACAP2 |
| ACAP3 |
| ACAT1 |
| ACAT2 |
| ACBD3 |
| ACBD4 |
| ACBD5 |
| ACBD6 |
| ACCS |
| ACD |
| ACER3 |
| ACHE |
| ACIN1 |
| ACLY |
| ACOT11 |
| ACOT12 |
| ACOT13 |
| ACOT2 |

| |
|---|
| ACOT4 |
| ACOT7 |
| ACOT8 |
| ACOT9 |
| ACOX1 |
| ACOX3 |
| ACP1 |
| ACP2 |
| ACP6 |
| ACRBP |
| ACSBG1 |
| ACSF2 |
| ACSF3 |
| ACSL1 |
| ACSL3 |
| ACSL6 |
| ACSM3 |
| ACSS1 |
| ACSS2 |
| ACSS3 |
| ACTA1 |
| ACTA2 |
| ACTB |
| ACTBL2 |
| ACTG1 |
| ACTL6A |
| ACTL7A |
| ACTL7B |
| ACTN1 |
| ACTN4 |
| ACTR10 |
| ACTR1B |
| ACTR2 |
| ACTR3 |
| ACTR6 |
| ACTR8 |
| ACVR1 |
| ACVR1B |
| ACVR2A |
| ACVR2B |
| ACY1 |
| ACYP1 |
| ACYP2 |
| ADA |
| ADAL |

| |
|---|
| ADAM10 |
| ADAM12 |
| ADAM15 |
| ADAM17 |
| ADAM19 |
| ADAM20 |
| ADAM21 |
| ADAM22 |
| ADAM32 |
| ADAM8 |
| ADAM9 |
| ADAMTS1 |
| ADAMTS12 |
| ADAMTS13 |
| ADAMTS18 |
| ADAMTS2 |
| ADAMTS3 |
| ADAMTS5 |
| ADAMTS6 |
| ADAMTS8 |
| ADAMTS9 |
| ADAMTSL4 |
| ADAP1 |
| ADAP2 |
| ADARB1 |
| ADAT1 |
| ADAT2 |
| ADCK1 |
| ADCK2 |
| ADCK5 |
| ADCY1 |
| ADCY3 |
| ADCY4 |
| ADCY6 |
| ADCY7 |
| ADCY9 |
| ADH4 |
| ADH5 |
| ADHFE1 |
| ADI1 |
| ADIPOR1 |
| ADIPOR2 |
| ADM |
| ADNP2 |
| ADORA1 |

| |
|---|
| ADORA2A |
| ADORA2B |
| ADPGK |
| ADPRHL1 |
| ADRA1D |
| ADRA2A |
| ADRB2 |
| ADRM1 |
| ADSL |
| AEBP1 |
| AEBP2 |
| AFAP1 |
| AFF2 |
| AFF3 |
| AFF4 |
| AFG3L2 |
| AFMID |
| AFTPH |
| AGAP3 |
| AGBL3 |
| AGBL5 |
| AGFG1 |
| AGFG2 |
| AGGF1 |
| AGK |
| AGL |
| AGMAT |
| AGPAT1 |
| AGPAT2 |
| AGPAT3 |
| AGPAT5 |
| AGPS |
| AGRN |
| AGT |
| AGTPBP1 |
| AGTR2 |
| AHCTF1 |
| AHCY |
| AHCYL1 |
| AHCYL2 |
| AHDC1 |
| AHI1 |
| AHNAK |
| AHNAK2 |
| AHR |

| |
|---|
| AHSA1 |
| AIDA |
| AIF1L |
| AIFM1 |
| AIMP1 |
| AIMP2 |
| AIP |
| AIRE |
| AK1 |
| AK2 |
| AK6 |
| AKAP1 |
| AKAP11 |
| AKAP12 |
| AKAP13 |
| AKAP5 |
| AKAP6 |
| AKAP7 |
| AKAP8 |
| AKAP8L |
| AKIP1 |
| AKIRIN1 |
| AKIRIN2 |
| AKNA |
| AKR1A1 |
| AKR1B1 |
| AKT1 |
| AKT1S1 |
| AKT2 |
| AKT3 |
| AKTIP |
| ALAS1 |
| ALCAM |
| ALDH16A1 |
| ALDH18A1 |
| ALDH1A1 |
| ALDH1B1 |
| ALDH1L1 |
| ALDH1L2 |
| ALDH2 |
| ALDH4A1 |
| ALDH5A1 |
| ALDH6A1 |
| ALDH7A1 |
| ALDH9A1 |

| |
|---|
| ALDOA |
| ALG1 |
| ALG11 |
| ALG12 |
| ALG13 |
| ALG14 |
| ALG2 |
| ALG3 |
| ALG5 |
| ALG6 |
| ALG8 |
| ALKBH1 |
| ALKBH2 |
| ALKBH3 |
| ALKBH4 |
| ALKBH5 |
| ALKBH7 |
| ALMS1 |
| ALOX15 |
| ALOX5 |
| ALOX5AP |
| ALPI |
| ALPK1 |
| ALPL |
| ALS2 |
| AMACR |
| AMBRA1 |
| AMD1 |
| AMDHD1 |
| AMFR |
| AMH |
| AMHR2 |
| AMIGO1 |
| AMIGO2 |
| AMIGO3 |
| AMN |
| AMN1 |
| AMOT |
| AMOTL1 |
| AMOTL2 |
| AMPD2 |
| AMPD3 |
| AMY2A |
| AMZ2 |
| ANAPC1 |

| |
|---|
| ANAPC10 |
| ANAPC11 |
| ANAPC13 |
| ANAPC7 |
| ANG |
| ANGEL1 |
| ANGEL2 |
| ANGPT2 |
| ANGPTL1 |
| ANGPTL2 |
| ANGPTL5 |
| ANGPTL7 |
| ANK1 |
| ANK2 |
| ANK3 |
| ANKFY1 |
| ANKH |
| ANKIB1 |
| ANKLE1 |
| ANKLE2 |
| ANKMY2 |
| ANKRA2 |
| ANKRD10 |
| ANKRD11 |
| ANKRD12 |
| ANKRD13A |
| ANKRD13C |
| ANKRD13D |
| ANKRD16 |
| ANKRD17 |
| ANKRD22 |
| ANKRD26 |
| ANKRD27 |
| ANKRD28 |
| ANKRD29 |
| ANKRD31 |
| ANKRD33B |
| ANKRD34A |
| ANKRD35 |
| ANKRD37 |
| ANKRD39 |
| ANKRD40 |
| ANKRD46 |
| ANKRD49 |
| ANKRD50 |

| |
|---|
| ANKRD52 |
| ANKRD55 |
| ANKRD9 |
| ANKS3 |
| ANKS4B |
| ANKS6 |
| ANKZF1 |
| ANLN |
| ANO10 |
| ANO2 |
| ANO6 |
| ANO8 |
| ANO9 |
| ANP32A |
| ANP32B |
| ANP32E |
| ANTXR1 |
| ANTXR2 |
| ANXA1 |
| ANXA11 |
| ANXA13 |
| ANXA2 |
| ANXA3 |
| ANXA5 |
| ANXA6 |
| AOAH |
| AOC3 |
| AOX1 |
| AP1AR |
| AP1B1 |
| AP1G1 |
| AP1G2 |
| AP1M1 |
| AP1S1 |
| AP1S2 |
| AP2A1 |
| AP2A2 |
| AP2M1 |
| AP2S1 |
| AP3B1 |
| AP3D1 |
| AP3M1 |
| AP3S1 |
| AP3S2 |
| AP4B1 |

| |
|---|
| AP4E1 |
| AP4M1 |
| AP4S1 |
| APAF1 |
| APBB1 |
| APBB1IP |
| APBB2 |
| APBB3 |
| APC |
| APEH |
| APEX1 |
| APEX2 |
| APH1A |
| APH1B |
| API5 |
| APIP |
| APLF |
| APLP2 |
| APMAP |
| APOA1 |
| APOB |
| APOD |
| APOE |
| APOL3 |
| APOLD1 |
| APOM |
| APOO |
| APP |
| APPBP2 |
| APPL1 |
| APPL2 |
| APRT |
| APTX |
| AQP11 |
| AQP7 |
| AQP8 |
| AQR |
| AR |
| ARAF |
| ARAP1 |
| ARAP2 |
| ARC |
| ARF1 |
| ARF4 |
| ARF5 |

| |
|---|
| ARF6 |
| ARFGAP2 |
| ARFGAP3 |
| ARFGEF1 |
| ARFGEF2 |
| ARFIP1 |
| ARFIP2 |
| ARFRP1 |
| ARG2 |
| ARGLU1 |
| ARHGAP1 |
| ARHGAP10 |
| ARHGAP11A |
| ARHGAP17 |
| ARHGAP18 |
| ARHGAP19 |
| ARHGAP20 |
| ARHGAP21 |
| ARHGAP22 |
| ARHGAP23 |
| ARHGAP25 |
| ARHGAP26 |
| ARHGAP29 |
| ARHGAP30 |
| ARHGAP4 |
| ARHGAP5 |
| ARHGAP6 |
| ARHGAP9 |
| ARHGDIA |
| ARHGDIB |
| ARHGDIG |
| ARHGEF1 |
| ARHGEF10 |
| ARHGEF10L |
| ARHGEF11 |
| ARHGEF12 |
| ARHGEF16 |
| ARHGEF18 |
| ARHGEF19 |
| ARHGEF2 |
| ARHGEF3 |
| ARHGEF4 |
| ARHGEF6 |
| ARHGEF7 |
| ARHGEF9 |

| |
|---|
| ARID1A |
| ARID1B |
| ARID2 |
| ARID3A |
| ARID3B |
| ARID4A |
| ARID5A |
| ARID5B |
| ARIH2 |
| ARL1 |
| ARL10 |
| ARL11 |
| ARL13B |
| ARL14 |
| ARL15 |
| ARL2 |
| ARL2BP |
| ARL3 |
| ARL4A |
| ARL4C |
| ARL4D |
| ARL5A |
| ARL5B |
| ARL6 |
| ARL6IP1 |
| ARL6IP5 |
| ARL6IP6 |
| ARL8B |
| ARMC1 |
| ARMC3 |
| ARMC5 |
| ARMC6 |
| ARMC7 |
| ARMC8 |
| ARMC9 |
| ARMCX1 |
| ARMCX2 |
| ARMCX3 |
| ARNT2 |
| ARNTL |
| ARPC1B |
| ARPC3 |
| ARPC4 |
| ARPC5 |
| ARPC5L |

| |
|---|
| ARPP21 |
| ARRB1 |
| ARRDC1 |
| ARRDC2 |
| ARRDC3 |
| ARRDC4 |
| ARSA |
| ARSG |
| ARSK |
| ART3 |
| ART4 |
| AS3MT |
| ASAH1 |
| ASAP1 |
| ASAP2 |
| ASAP3 |
| ASB1 |
| ASB13 |
| ASB15 |
| ASB16 |
| ASB3 |
| ASB4 |
| ASB5 |
| ASB6 |
| ASB7 |
| ASB8 |
| ASCC1 |
| ASCC2 |
| ASCC3 |
| ASCL1 |
| ASCL4 |
| ASF1A |
| ASF1B |
| ASH1L |
| ASL |
| ASMTL |
| ASNS |
| ASNSD1 |
| ASPG |
| ASPH |
| ASPHD2 |
| ASPM |
| ASPSCR1 |
| ASRGL1 |
| ASTE1 |

| |
|---|
| ASXL1 |
| ASXL2 |
| ASXL3 |
| ATAD1 |
| ATAD2 |
| ATAD5 |
| ATE1 |
| ATF2 |
| ATF3 |
| ATF4 |
| ATF5 |
| ATF7 |
| ATF7IP |
| ATF7IP2 |
| ATG10 |
| ATG101 |
| ATG12 |
| ATG13 |
| ATG16L1 |
| ATG16L2 |
| ATG2A |
| ATG2B |
| ATG3 |
| ATG4A |
| ATG4B |
| ATG4C |
| ATG5 |
| ATG7 |
| ATG9A |
| ATIC |
| ATL3 |
| ATM |
| ATMIN |
| ATN1 |
| ATOH1 |
| ATOH7 |
| ATOX1 |
| ATP10D |
| ATP11A |
| ATP11B |
| ATP11C |
| ATP12A |
| ATP13A1 |
| ATP13A2 |
| ATP1A1 |

| |
|---|
| ATP1A3 |
| ATP1B1 |
| ATP1B2 |
| ATP1B3 |
| ATP2A1 |
| ATP2A2 |
| ATP2A3 |
| ATP2B4 |
| ATP2C1 |
| ATP4A |
| ATP4B |
| ATP6AP1 |
| ATP6AP2 |
| ATP6V0A1 |
| ATP6V0A2 |
| ATP6V0A4 |
| ATP6V0B |
| ATP6V0C |
| ATP6V0D1 |
| ATP6V0D2 |
| ATP6V0E1 |
| ATP6V1A |
| ATP6V1B2 |
| ATP6V1D |
| ATP6V1E1 |
| ATP6V1F |
| ATP6V1G1 |
| ATP6V1G2 |
| ATP6V1H |
| ATP7A |
| ATP7B |
| ATP8A1 |
| ATP8A2 |
| ATP8B1 |
| ATP8B2 |
| ATP8B3 |
| ATP8B4 |
| ATP9B |
| ATPAF1 |
| ATPAF2 |
| ATR |
| ATRIP |
| ATRN |
| ATRNL1 |
| ATRX |

| |
|---|
| ATXN1 |
| ATXN1L |
| ATXN2 |
| ATXN2L |
| ATXN3 |
| ATXN7 |
| ATXN7L1 |
| ATXN7L2 |
| ATXN7L3 |
| AUH |
| AUP1 |
| AURKA |
| AURKAIP1 |
| AURKC |
| AUTS2 |
| AVEN |
| AVL9 |
| AVPI1 |
| AXIN1 |
| AXIN2 |
| AXL |
| AZI2 |
| AZIN1 |
| B2M |
| B3GALNT1 |
| B3GALNT2 |
| B3GALT2 |
| B3GALT4 |
| B3GALT5 |
| B3GALT6 |
| B3GAT2 |
| B3GAT3 |
| B3GNT2 |
| B3GNT3 |
| B3GNT4 |
| B3GNT5 |
| B3GNT7 |
| B3GNT8 |
| B3GNT9 |
| B3GNTL1 |
| B4GALNT1 |
| B4GALNT2 |
| B4GALNT3 |
| B4GALT1 |
| B4GALT2 |

| |
|---|
| B4GALT3 |
| B4GALT4 |
| B4GALT5 |
| B4GALT7 |
| B9D1 |
| B9D2 |
| BAALC |
| BACE1 |
| BACE2 |
| BACH2 |
| BAD |
| BAG1 |
| BAG2 |
| BAG3 |
| BAG4 |
| BAG5 |
| BAHCC1 |
| BAHD1 |
| BAIAP2 |
| BAIAP2L2 |
| BAK1 |
| BAMBI |
| BANF1 |
| BANP |
| BARD1 |
| BASP1 |
| BATF2 |
| BAX |
| BAZ1B |
| BAZ2A |
| BAZ2B |
| BBS12 |
| BBS2 |
| BBS4 |
| BBS5 |
| BBS7 |
| BBS9 |
| BBX |
| BCAM |
| BCAP29 |
| BCAP31 |
| BCAR1 |
| BCAR3 |
| BCAS1 |
| BCAS2 |

| |
|---|
| BCAS3 |
| BCAS4 |
| BCAT2 |
| BCCIP |
| BCDIN3D |
| BCHE |
| BCKDHA |
| BCKDHB |
| BCKDK |
| BCL10 |
| BCL11A |
| BCL2 |
| BCL2L1 |
| BCL2L11 |
| BCL2L12 |
| BCL2L15 |
| BCL3 |
| BCL6 |
| BCL6B |
| BCL7B |
| BCL7C |
| BCL9 |
| BCLAF1 |
| BCOR |
| BCORL1 |
| BCR |
| BCS1L |
| BDH2 |
| BDP1 |
| BECN1 |
| BEND3 |
| BEND5 |
| BEND6 |
| BEND7 |
| BEST1 |
| BEST2 |
| BEST4 |
| BET1L |
| BEX2 |
| BEX4 |
| BFAR |
| BFSP1 |
| BGN |
| BHLHB9 |
| BHLHE41 |

| |
|---|
| BICD1 |
| BICD2 |
| BID |
| BIN1 |
| BIN2 |
| BIN3 |
| BIRC3 |
| BIRC5 |
| BIRC6 |
| BLCAP |
| BLM |
| BLMH |
| BLNK |
| BLOC1S1 |
| BLOC1S2 |
| BLVRA |
| BLVRB |
| BLZF1 |
| BMF |
| BMP1 |
| BMP2 |
| BMP3 |
| BMP6 |
| BMPR1A |
| BMPR2 |
| BMS1 |
| BNC2 |
| BNIP1 |
| BNIP3 |
| BNIP3L |
| BOK |
| BOLA1 |
| BOLA3 |
| BOP1 |
| BPGM |
| BPNT1 |
| BPTF |
| BRAF |
| BRCA1 |
| BRCC3 |
| BRD2 |
| BRD3 |
| BRD4 |
| BRD7 |
| BRD8 |

| |
|---|
| BRD9 |
| BRDT |
| BRI3 |
| BRI3BP |
| BRIP1 |
| BRIX1 |
| BRMS1L |
| BRPF1 |
| BRPF3 |
| BRSK1 |
| BRWD1 |
| BRWD3 |
| BSCL2 |
| BSDC1 |
| BSG |
| BSPRY |
| BST1 |
| BST2 |
| BTAF1 |
| BTBD1 |
| BTBD10 |
| BTBD3 |
| BTBD6 |
| BTBD7 |
| BTC |
| BTD |
| BTF3 |
| BTF3L4 |
| BTG1 |
| BTG2 |
| BTG3 |
| BTK |
| BTN1A1 |
| BTN3A3 |
| BTNL9 |
| BTRC |
| BUB1 |
| BUB1B |
| BUB3 |
| BUD13 |
| BUD31 |
| BYSL |
| BZW1 |
| BZW2 |
| C1D |

| |
|---|
| C1GALT1 |
| C1GALT1C1 |
| C1orf35 |
| C1orf52 |
| C1QA |
| C1QB |
| C1QBP |
| C1QC |
| C1QTNF1 |
| C1QTNF2 |
| C1QTNF6 |
| C1QTNF8 |
| C1R |
| C1RL |
| C1S |
| C2 |
| C2CD2 |
| C2CD2L |
| C3AR1 |
| C4A |
| C5AR1 |
| C6orf120 |
| C8G |
| C9orf64 |
| CA12 |
| CA2 |
| CA4 |
| CA5B |
| CA7 |
| CA9 |
| CAB39L |
| CABIN1 |
| CABLES2 |
| CABP5 |
| CABYR |
| CACHD1 |
| CACNA1F |
| CACNA1G |
| CACNA1H |
| CACNA2D1 |
| CACNB1 |
| CACNB2 |
| CACNB3 |
| CACYBP |
| CAD |

| |
|---|
| CADM1 |
| CADM4 |
| CALCOCO1 |
| CALCOCO2 |
| CALD1 |
| CALHM2 |
| CALM1 |
| CALR |
| CALU |
| CAMK2D |
| CAMK2G |
| CAMK2N1 |
| CAMKK1 |
| CAMKV |
| CAMLG |
| CAMSAP1 |
| CAMTA1 |
| CAND1 |
| CANT1 |
| CANX |
| CAP1 |
| CAPG |
| CAPN1 |
| CAPN10 |
| CAPN2 |
| CAPN6 |
| CAPN7 |
| CAPNS1 |
| CAPRIN2 |
| CAPZA1 |
| CAPZA2 |
| CAPZB |
| CARD6 |
| CARHSP1 |
| CARM1 |
| CARS2 |
| CASC3 |
| CASD1 |
| CASKIN1 |
| CASKIN2 |
| CASP1 |
| CASP14 |
| CASP2 |
| CASP3 |
| CASP7 |

| |
|---|
| CASP8 |
| CASP8AP2 |
| CASP9 |
| CASQ1 |
| CASQ2 |
| CASZ1 |
| CATSPER2 |
| CAV2 |
| CBFA2T2 |
| CBFB |
| CBL |
| CBLB |
| CBLC |
| CBLL1 |
| CBR1 |
| CBR3 |
| CBR4 |
| CBS |
| CBX1 |
| CBX2 |
| CBX3 |
| CBX5 |
| CBX6 |
| CBX7 |
| CBX8 |
| CBY1 |
| CC2D1A |
| CC2D1B |
| CC2D2A |
| CCAR1 |
| CCBE1 |
| CCDC102A |
| CCDC103 |
| CCDC106 |
| CCDC107 |
| CCDC112 |
| CCDC115 |
| CCDC117 |
| CCDC12 |
| CCDC121 |
| CCDC122 |
| CCDC124 |
| CCDC125 |
| CCDC126 |
| CCDC127 |

| |
|---|
| CCDC137 |
| CCDC138 |
| CCDC14 |
| CCDC142 |
| CCDC149 |
| CCDC15 |
| CCDC157 |
| CCDC158 |
| CCDC159 |
| CCDC160 |
| CCDC18 |
| CCDC22 |
| CCDC24 |
| CCDC25 |
| CCDC28A |
| CCDC28B |
| CCDC32 |
| CCDC34 |
| CCDC43 |
| CCDC47 |
| CCDC50 |
| CCDC51 |
| CCDC59 |
| CCDC61 |
| CCDC65 |
| CCDC66 |
| CCDC69 |
| CCDC7 |
| CCDC71 |
| CCDC73 |
| CCDC74B |
| CCDC77 |
| CCDC8 |
| CCDC80 |
| CCDC82 |
| CCDC85A |
| CCDC85B |
| CCDC85C |
| CCDC86 |
| CCDC88A |
| CCDC88B |
| CCDC88C |
| CCDC9 |
| CCDC90B |
| CCDC91 |

| |
|---|
| CCDC93 |
| CCDC96 |
| CCDC97 |
| CCHCR1 |
| CCIN |
| CCKBR |
| CCL14 |
| CCL2 |
| CCL20 |
| CCL27 |
| CCL28 |
| CCL4 |
| CCL5 |
| CCL8 |
| CCM2 |
| CCNA2 |
| CCNB2 |
| CCNC |
| CCND2 |
| CCND3 |
| CCNE1 |
| CCNE2 |
| CCNG1 |
| CCNG2 |
| CCNH |
| CCNI |
| CCNJ |
| CCNK |
| CCNL2 |
| CCNT1 |
| CCNT2 |
| CCNY |
| CCNYL1 |
| CCPG1 |
| CCR1 |
| CCR2 |
| CCR3 |
| CCR5 |
| CCR7 |
| CCRL2 |
| CCS |
| CCT2 |
| CCT3 |
| CCT4 |
| CCT5 |

| |
|---|
| CCT6A |
| CCT7 |
| CCT8 |
| CD101 |
| CD109 |
| CD14 |
| CD151 |
| CD164 |
| CD180 |
| CD1D |
| CD2 |
| CD200 |
| CD244 |
| CD247 |
| CD2AP |
| CD2BP2 |
| CD300A |
| CD300LB |
| CD300LF |
| CD302 |
| CD36 |
| CD37 |
| CD3E |
| CD40 |
| CD44 |
| CD46 |
| CD47 |
| CD48 |
| CD5 |
| CD53 |
| CD55 |
| CD58 |
| CD6 |
| CD63 |
| CD68 |
| CD69 |
| CD7 |
| CD70 |
| CD72 |
| CD74 |
| CD81 |
| CD82 |
| CD84 |
| CD86 |
| CD8A |

| |
|---|
| CD9 |
| CD96 |
| CD99L2 |
| CDADC1 |
| CDAN1 |
| CDC14A |
| CDC14B |
| CDC16 |
| CDC25B |
| CDC25C |
| CDC26 |
| CDC27 |
| CDC34 |
| CDC37 |
| CDC37L1 |
| CDC40 |
| CDC42 |
| CDC42BPA |
| CDC42BPG |
| CDC42EP2 |
| CDC42EP3 |
| CDC42EP4 |
| CDC42EP5 |
| CDC42SE2 |
| CDC5L |
| CDC6 |
| CDC7 |
| CDC73 |
| CDCA2 |
| CDCA3 |
| CDCA4 |
| CDCA5 |
| CDCA7 |
| CDCA7L |
| CDCA8 |
| CDCP1 |
| CDH10 |
| CDH12 |
| CDH17 |
| CDH18 |
| CDH2 |
| CDH24 |
| CDH3 |
| CDH6 |
| CDH8 |

| |
|---|
| CDHR2 |
| CDHR5 |
| CDIPT |
| CDK1 |
| CDK10 |
| CDK12 |
| CDK13 |
| CDK16 |
| CDK17 |
| CDK18 |
| CDK19 |
| CDK20 |
| CDK2AP1 |
| CDK2AP2 |
| CDK3 |
| CDK4 |
| CDK5 |
| CDK5R1 |
| CDK5RAP1 |
| CDK5RAP2 |
| CDK6 |
| CDK7 |
| CDK8 |
| CDK9 |
| CDKAL1 |
| CDKL1 |
| CDKL3 |
| CDKL5 |
| CDKN1A |
| CDKN1B |
| CDKN1C |
| CDKN2A |
| CDKN2AIPNL |
| CDKN2B |
| CDKN2C |
| CDKN2D |
| CDKN3 |
| CDO1 |
| CDON |
| CDR2 |
| CDR2L |
| CDS2 |
| CDV3 |
| CDX1 |
| CDX2 |

| |
|---|
| CDYL |
| CDYL2 |
| CEBPA |
| CEBPD |
| CEBPG |
| CEBPZ |
| CELA3B |
| CELSR1 |
| CELSR2 |
| CENPB |
| CENPE |
| CENPF |
| CENPH |
| CENPI |
| CENPJ |
| CENPK |
| CENPM |
| CENPN |
| CENPO |
| CENPP |
| CENPQ |
| CENPT |
| CENPV |
| CEP135 |
| CEP164 |
| CEP250 |
| CEP350 |
| CEP55 |
| CEP57 |
| CEP68 |
| CEP70 |
| CEP72 |
| CEP76 |
| CEP78 |
| CEP97 |
| CERCAM |
| CES1 |
| CETN2 |
| CETN3 |
| CFB |
| CFD |
| CFDP1 |
| CFDP2 |
| CFL1 |
| CFL2 |

| |
|---|
| CGN |
| CGREF1 |
| CGRRF1 |
| CHAC1 |
| CHAC2 |
| CHADL |
| CHAF1A |
| CHAF1B |
| CHCHD3 |
| CHCHD4 |
| CHCHD6 |
| CHCHD7 |
| CHD1L |
| CHD2 |
| CHD3 |
| CHD4 |
| CHD8 |
| CHD9 |
| CHEK1 |
| CHEK2 |
| CHERP |
| CHFR |
| CHGB |
| CHI3L1 |
| CHIC1 |
| CHIC2 |
| CHKA |
| CHL1 |
| CHML |
| CHMP1A |
| CHMP1B |
| CHMP2A |
| CHMP2B |
| CHMP4A |
| CHMP4B |
| CHMP4C |
| CHMP5 |
| CHMP6 |
| CHORDC1 |
| CHPT1 |
| CHRAC1 |
| CHRD |
| CHRDL1 |
| CHRM3 |
| CHRM5 |

| |
|---|
| CHST11 |
| CHST12 |
| CHST13 |
| CHST14 |
| CHST3 |
| CHST4 |
| CHST7 |
| CHSY1 |
| CHSY3 |
| CHUK |
| CIAO1 |
| CIAPIN1 |
| CIB1 |
| CIDEC |
| CINP |
| CIRBP |
| CISD1 |
| CISD2 |
| CISD3 |
| CISH |
| CIT |
| CITED2 |
| CITED4 |
| CIZ1 |
| CKAP2 |
| CKAP2L |
| CKAP4 |
| CKAP5 |
| CKB |
| CKMT2 |
| CKS1B |
| CLASP1 |
| CLASP2 |
| CLCA2 |
| CLCN2 |
| CLCN3 |
| CLCN4 |
| CLCN5 |
| CLCN6 |
| CLCN7 |
| CLDN1 |
| CLDN11 |
| CLDN12 |
| CLDN15 |
| CLDN20 |

| |
|---|
| CLDN23 |
| CLDN3 |
| CLDN4 |
| CLDN7 |
| CLDN8 |
| CLEC11A |
| CLEC12A |
| CLEC14A |
| CLEC16A |
| CLEC1A |
| CLEC3A |
| CLEC3B |
| CLGN |
| CLIC1 |
| CLIC4 |
| CLINT1 |
| CLIP1 |
| CLIP2 |
| CLIP3 |
| CLK1 |
| CLK3 |
| CLK4 |
| CLMN |
| CLN3 |
| CLN5 |
| CLN6 |
| CLN8 |
| CLNS1A |
| CLPB |
| CLPP |
| CLPTM1 |
| CLPTM1L |
| CLPX |
| CLRN3 |
| CLSPN |
| CLSTN1 |
| CLSTN2 |
| CLSTN3 |
| CLTA |
| CLTB |
| CLTC |
| CLU |
| CLUAP1 |
| CLYBL |
| CMBL |

| |
|---|
| CMC1 |
| CMPK1 |
| CMPK2 |
| CMTM3 |
| CMTM6 |
| CMTM7 |
| CMYA5 |
| CNBP |
| CNDP2 |
| CNIH4 |
| CNKSR3 |
| CNN1 |
| CNN2 |
| CNN3 |
| CNNM2 |
| CNNM3 |
| CNNM4 |
| CNOT1 |
| CNOT10 |
| CNOT3 |
| CNOT4 |
| CNOT6 |
| CNOT6L |
| CNOT7 |
| CNOT8 |
| CNP |
| CNPY2 |
| CNPY3 |
| CNPY4 |
| CNR2 |
| CNTLN |
| CNTN1 |
| CNTN3 |
| CNTN6 |
| CNTNAP1 |
| CNTROB |
| COASY |
| COBL |
| COBLL1 |
| COCH |
| COG1 |
| COG2 |
| COG3 |
| COG4 |
| COG5 |

| |
|---|
| COG6 |
| COG7 |
| COG8 |
| COIL |
| COL13A1 |
| COL15A1 |
| COL17A1 |
| COL1A1 |
| COL21A1 |
| COL24A1 |
| COL3A1 |
| COL4A1 |
| COL4A2 |
| COL4A3 |
| COL4A4 |
| COL4A5 |
| COL4A6 |
| COL6A1 |
| COL6A2 |
| COL7A1 |
| COL8A2 |
| COL9A2 |
| COLEC11 |
| COLEC12 |
| COMMD1 |
| COMMD10 |
| COMMD2 |
| COMMD4 |
| COMMD5 |
| COMMD6 |
| COMMD7 |
| COMMD8 |
| COMMD9 |
| COMT |
| COMTD1 |
| COPA |
| COPB1 |
| COPB2 |
| COPE |
| COPG2 |
| COPS3 |
| COPS5 |
| COPS6 |
| COPS7A |
| COPS8 |

| |
|---|
| COPZ1 |
| COQ10A |
| COQ10B |
| COQ2 |
| COQ3 |
| COQ5 |
| COQ6 |
| COQ7 |
| COQ9 |
| CORIN |
| CORO1A |
| CORO1B |
| CORO1C |
| CORO2A |
| CORO6 |
| CORO7 |
| COTL1 |
| COX10 |
| COX11 |
| COX15 |
| COX16 |
| COX18 |
| COX19 |
| COX5A |
| COX5B |
| COX6A1 |
| COX6B1 |
| COX6C |
| COX7A1 |
| COX7A2 |
| COX7A2L |
| COX8A |
| CPD |
| CPEB1 |
| CPEB2 |
| CPEB4 |
| CPN2 |
| CPNE3 |
| CPOX |
| CPPED1 |
| CPSF1 |
| CPSF2 |
| CPSF3 |
| CPSF4 |
| CPT1A |

| |
|---|
| CPT1B |
| CPT1C |
| CPT2 |
| CPVL |
| CPXM1 |
| CPZ |
| CRADD |
| CRAT |
| CRBN |
| CRCP |
| CREB1 |
| CREB3 |
| CREB3L1 |
| CREB3L2 |
| CREB3L3 |
| CREBL2 |
| CREBZF |
| CREG1 |
| CRELD1 |
| CRELD2 |
| CREM |
| CRIM1 |
| CRIP1 |
| CRIP3 |
| CRIPT |
| CRK |
| CRLF3 |
| CRLS1 |
| CRNKL1 |
| CROCC |
| CRP |
| CRTAP |
| CRTC3 |
| CRY2 |
| CRYAB |
| CRYBG3 |
| CRYGC |
| CRYM |
| CRYZ |
| CRYZL1 |
| CS |
| CSDE1 |
| CSF1 |
| CSF1R |
| CSF2RB |

| |
|---|
| CSGALNACT1 |
| CSGALNACT2 |
| CSK |
| CSMD3 |
| CSN2 |
| CSNK1A1 |
| CSNK1D |
| CSNK1E |
| CSNK1G1 |
| CSNK1G2 |
| CSNK1G3 |
| CSNK2B |
| CSPG4 |
| CSPG5 |
| CSPP1 |
| CSRNP1 |
| CSRNP2 |
| CSRP1 |
| CSRP2 |
| CST11 |
| CST3 |
| CST6 |
| CST7 |
| CSTF1 |
| CSTF2 |
| CSTF2T |
| CSTF3 |
| CSTL1 |
| CTBP1 |
| CTBP2 |
| CTCF |
| CTDSP1 |
| CTDSP2 |
| CTF1 |
| CTGF |
| CTH |
| CTHRC1 |
| CTNNA1 |
| CTNNA2 |
| CTNNAL1 |
| CTNNB1 |
| CTNNBL1 |
| CTNND1 |
| CTR9 |
| CTSA |

| |
|---|
| CTSB |
| CTSC |
| CTSD |
| CTSH |
| CTSK |
| CTSO |
| CTSZ |
| CTTN |
| CTTNBP2NL |
| CTU1 |
| CTU2 |
| CTXN1 |
| CUBN |
| CUEDC2 |
| CUL1 |
| CUL2 |
| CUL3 |
| CUL4B |
| CUL5 |
| CUL7 |
| CUTA |
| CUTC |
| CUX1 |
| CUX2 |
| CWC15 |
| CWC22 |
| CWC25 |
| CWC27 |
| CWF19L1 |
| CWF19L2 |
| CWH43 |
| CX3CR1 |
| CXADR |
| CXCL10 |
| CXCL16 |
| CXCL2 |
| CXCR4 |
| CXCR5 |
| CYB561D2 |
| CYB5B |
| CYB5R1 |
| CYB5R3 |
| CYBB |
| CYC1 |
| CYCS |

| |
|---|
| CYFIP2 |
| CYHR1 |
| CYLD |
| CYP11A1 |
| CYP19A1 |
| CYP1B1 |
| CYP20A1 |
| CYP27A1 |
| CYP27B1 |
| CYP2J2 |
| CYP2S1 |
| CYP2U1 |
| CYP4B1 |
| CYP4F3 |
| CYP4V2 |
| CYP8B1 |
| CYSLTR1 |
| CYSLTR2 |
| CYTH2 |
| CYTH3 |
| CYTH4 |
| CYTIP |
| D2HGDH |
| DAAM2 |
| DAB2 |
| DAB2IP |
| DACH1 |
| DAD1 |
| DAG1 |
| DAGLA |
| DAGLB |
| DALRD3 |
| DAP |
| DAPK1 |
| DAPK3 |
| DAXX |
| DAZAP1 |
| DAZAP2 |
| DBF4 |
| DBI |
| DBN1 |
| DBNDD1 |
| DBNL |
| DBP |
| DBR1 |

| |
|---|
| DBT |
| DCAF10 |
| DCAF11 |
| DCAF12 |
| DCAF15 |
| DCAF16 |
| DCAF17 |
| DCAF4 |
| DCAF5 |
| DCAF6 |
| DCAF7 |
| DCAF8 |
| DCAKD |
| DCBLD2 |
| DCHS1 |
| DCK |
| DCLRE1A |
| DCLRE1C |
| DCP1A |
| DCP1B |
| DCP2 |
| DCPS |
| DCT |
| DCTD |
| DCTN1 |
| DCTN2 |
| DCTN3 |
| DCTN5 |
| DCTN6 |
| DCTPP1 |
| DCUN1D1 |
| DCUN1D3 |
| DCUN1D5 |
| DCXR |
| DDA1 |
| DDAH2 |
| DDB1 |
| DDB2 |
| DDHD1 |
| DDHD2 |
| DDI1 |
| DDI2 |
| DDIT3 |
| DDIT4 |
| DDN |

| |
|---|
| DDO |
| DDOST |
| DDR1 |
| DDR2 |
| DDRGK1 |
| DDT |
| DDX1 |
| DDX10 |
| DDX17 |
| DDX18 |
| DDX19A |
| DDX19B |
| DDX20 |
| DDX21 |
| DDX23 |
| DDX24 |
| DDX27 |
| DDX28 |
| DDX31 |
| DDX3X |
| DDX41 |
| DDX46 |
| DDX47 |
| DDX49 |
| DDX5 |
| DDX50 |
| DDX52 |
| DDX54 |
| DDX55 |
| DDX56 |
| DDX58 |
| DDX59 |
| DDX6 |
| DEAF1 |
| DECR1 |
| DECR2 |
| DEDD |
| DEDD2 |
| DEF8 |
| DEGS1 |
| DEGS2 |
| DENND1A |
| DENND1B |
| DENND1C |
| DENND2A |

| |
|---|
| DENND2C |
| DENND2D |
| DENND3 |
| DENND4A |
| DENND4B |
| DENND4C |
| DENND5A |
| DENND5B |
| DEPDC1 |
| DEPDC1B |
| DEPDC5 |
| DEPDC7 |
| DERL1 |
| DERL2 |
| DERL3 |
| DET1 |
| DEXI |
| DFFA |
| DFFB |
| DGAT1 |
| DGAT2 |
| DGCR2 |
| DGCR8 |
| DGKA |
| DGKE |
| DGKQ |
| DGKZ |
| DGUOK |
| DHCR24 |
| DHCR7 |
| DHDDS |
| DHDH |
| DHFR |
| DHH |
| DHODH |
| DHPS |
| DHRS1 |
| DHRS11 |
| DHRS13 |
| DHRS4 |
| DHRS7 |
| DHRS7B |
| DHRS9 |
| DHRSX |
| DHTKD1 |

| |
|---|
| DHX15 |
| DHX16 |
| DHX29 |
| DHX30 |
| DHX32 |
| DHX33 |
| DHX34 |
| DHX35 |
| DHX36 |
| DHX37 |
| DHX38 |
| DHX40 |
| DHX57 |
| DHX9 |
| DIABLO |
| DIAPH2 |
| DIAPH3 |
| DICER1 |
| DIDO1 |
| DIP2A |
| DIP2B |
| DIP2C |
| DIRAS1 |
| DIRAS2 |
| DIRAS3 |
| DIS3 |
| DIS3L |
| DIS3L2 |
| DIXDC1 |
| DKC1 |
| DKK3 |
| DKKL1 |
| DLAT |
| DLC1 |
| DLD |
| DLEC1 |
| DLG1 |
| DLG4 |
| DLG5 |
| DLGAP1 |
| DLGAP5 |
| DLL1 |
| DLL3 |
| DLX5 |
| DMAP1 |

| |
|---|
| DMD |
| DMKN |
| DMPK |
| DMTF1 |
| DMWD |
| DMXL1 |
| DMXL2 |
| DNA2 |
| DNAH1 |
| DNAH2 |
| DNAH7 |
| DNAH9 |
| DNAI2 |
| DNAJA2 |
| DNAJA3 |
| DNAJA4 |
| DNAJB1 |
| DNAJB12 |
| DNAJB14 |
| DNAJB2 |
| DNAJB4 |
| DNAJB6 |
| DNAJB7 |
| DNAJB9 |
| DNAJC10 |
| DNAJC11 |
| DNAJC12 |
| DNAJC13 |
| DNAJC14 |
| DNAJC15 |
| DNAJC16 |
| DNAJC17 |
| DNAJC18 |
| DNAJC19 |
| DNAJC2 |
| DNAJC21 |
| DNAJC24 |
| DNAJC25 |
| DNAJC27 |
| DNAJC3 |
| DNAJC30 |
| DNAJC4 |
| DNAJC5 |
| DNAJC7 |
| DNAJC8 |

| |
|---|
| DNAJC9 |
| DNAL1 |
| DNAL4 |
| DNASE1L1 |
| DNASE2 |
| DND1 |
| DNHD1 |
| DNM1 |
| DNM1L |
| DNM2 |
| DNMBP |
| DNMT1 |
| DNMT3A |
| DNMT3B |
| DNPEP |
| DNTTIP1 |
| DNTTIP2 |
| DOC2B |
| DOCK1 |
| DOCK11 |
| DOCK3 |
| DOCK4 |
| DOCK5 |
| DOCK6 |
| DOCK7 |
| DOCK8 |
| DOCK9 |
| DOHH |
| DOK1 |
| DOK2 |
| DOK3 |
| DOK4 |
| DOK7 |
| DOLK |
| DOLPP1 |
| DPCD |
| DPF2 |
| DPF3 |
| DPH2 |
| DPH3 |
| DPH5 |
| DPM1 |
| DPM2 |
| DPM3 |
| DPP3 |

| |
|---|
| DPP7 |
| DPY19L1 |
| DPY19L3 |
| DPY19L4 |
| DPY30 |
| DPYD |
| DPYS |
| DPYSL2 |
| DPYSL3 |
| DPYSL5 |
| DQX1 |
| DR1 |
| DRAM2 |
| DRAP1 |
| DRG1 |
| DRG2 |
| DRP2 |
| DSC2 |
| DSCC1 |
| DSE |
| DSG2 |
| DSP |
| DST |
| DSTN |
| DSTYK |
| DTD1 |
| DTNB |
| DTNBP1 |
| DTWD1 |
| DTWD2 |
| DTX1 |
| DTX2 |
| DTX3 |
| DTX3L |
| DTYMK |
| DUOX1 |
| DUS1L |
| DUS3L |
| DUS4L |
| DUSP1 |
| DUSP10 |
| DUSP11 |
| DUSP12 |
| DUSP14 |
| DUSP15 |

| |
|---|
| DUSP16 |
| DUSP18 |
| DUSP19 |
| DUSP23 |
| DUSP3 |
| DUSP5 |
| DUSP6 |
| DUSP7 |
| DUSP9 |
| DUT |
| DVL1 |
| DVL2 |
| DYM |
| DYNC1I2 |
| DYNC1LI1 |
| DYNC2LI1 |
| DYNLL1 |
| DYNLL2 |
| DYNLRB1 |
| DYNLRB2 |
| DYNLT1 |
| DYNLT3 |
| DYRK1A |
| DYRK1B |
| DYRK3 |
| DZIP1 |
| DZIP3 |
| E2F1 |
| E2F2 |
| E2F3 |
| E2F4 |
| E2F5 |
| E2F6 |
| E4F1 |
| EAF1 |
| EAF2 |
| EAPP |
| EBAG9 |
| EBNA1BP2 |
| EBP |
| EBPL |
| ECD |
| ECE1 |
| ECH1 |
| ECHDC1 |

| |
|---|
| ECHDC3 |
| ECHS1 |
| ECM2 |
| ECRG4 |
| ECSIT |
| ECT2 |
| EDC3 |
| EDC4 |
| EDEM1 |
| EDEM2 |
| EDEM3 |
| EDF1 |
| EDIL3 |
| EDN1 |
| EDNRA |
| EEA1 |
| EED |
| EEF1A1 |
| EEF1D |
| EEF1E1 |
| EEF1G |
| EEF2 |
| EEF2K |
| EEFSEC |
| EEPD1 |
| EFCAB2 |
| EFCAB6 |
| EFCAB7 |
| EFEMP2 |
| EFHD2 |
| EFNA1 |
| EFNA2 |
| EFNA5 |
| EFNB1 |
| EFNB2 |
| EFR3A |
| EFTUD2 |
| EGFL6 |
| EGFL7 |
| EGFL8 |
| EGLN1 |
| EGLN2 |
| EGLN3 |
| EGR1 |
| EGR3 |

| |
|---|
| EHBP1L1 |
| EHD1 |
| EHD2 |
| EHD4 |
| EHF |
| EHHADH |
| EHMT1 |
| EHMT2 |
| EI24 |
| EID1 |
| EID2 |
| EIF1 |
| EIF1B |
| EIF2AK1 |
| EIF2AK2 |
| EIF2AK3 |
| EIF2AK4 |
| EIF2B1 |
| EIF2B2 |
| EIF2B4 |
| EIF2B5 |
| EIF2S1 |
| EIF2S2 |
| EIF2S3 |
| EIF3B |
| EIF3D |
| EIF3E |
| EIF3F |
| EIF3G |
| EIF3I |
| EIF3J |
| EIF3K |
| EIF3L |
| EIF3M |
| EIF4A1 |
| EIF4A2 |
| EIF4A3 |
| EIF4B |
| EIF4E |
| EIF4E2 |
| EIF4E3 |
| EIF4EBP1 |
| EIF4EBP2 |
| EIF4ENIF1 |
| EIF4G2 |

| |
|---|
| EIF4G3 |
| EIF4H |
| EIF5 |
| EIF5A2 |
| EIF5B |
| ELAC1 |
| ELAC2 |
| ELF1 |
| ELF2 |
| ELF3 |
| ELF4 |
| ELK1 |
| ELK3 |
| ELK4 |
| ELL |
| ELL2 |
| ELMO1 |
| ELMO2 |
| ELMO3 |
| ELMOD1 |
| ELMOD2 |
| ELMOD3 |
| ELOVL1 |
| ELOVL3 |
| ELOVL5 |
| ELOVL6 |
| ELP2 |
| ELP3 |
| ELP4 |
| EMCN |
| EMD |
| EME1 |
| EME2 |
| EMG1 |
| EMID1 |
| EMILIN1 |
| EMILIN2 |
| EML1 |
| EML2 |
| EML3 |
| EML4 |
| EML6 |
| EMP2 |
| EMP3 |
| EMX2 |

| |
|---|
| ENAH |
| ENC1 |
| ENDOD1 |
| ENDOG |
| ENHO |
| ENO3 |
| ENOPH1 |
| ENOSF1 |
| ENOX1 |
| ENOX2 |
| ENPEP |
| ENPP1 |
| ENPP3 |
| ENPP4 |
| ENPP5 |
| ENPP6 |
| ENSA |
| ENTPD1 |
| ENTPD2 |
| ENTPD3 |
| ENTPD4 |
| ENTPD5 |
| ENTPD6 |
| ENTPD7 |
| ENTPD8 |
| ENY2 |
| EP400 |
| EPAS1 |
| EPB41 |
| EPB41L2 |
| EPB41L3 |
| EPB41L4B |
| EPB41L5 |
| EPC1 |
| EPC2 |
| EPCAM |
| EPDR1 |
| EPHA1 |
| EPHA2 |
| EPHA4 |
| EPHB1 |
| EPHB4 |
| EPHB6 |
| EPHX1 |
| EPHX2 |

| |
|---|
| EPM2A |
| EPM2AIP1 |
| EPN1 |
| EPN2 |
| EPN3 |
| EPS15 |
| EPS15L1 |
| EPS8L2 |
| EPS8L3 |
| EPSTI1 |
| ERAL1 |
| ERAP1 |
| ERAS |
| ERBB3 |
| ERBB4 |
| ERCC1 |
| ERCC2 |
| ERCC3 |
| ERCC4 |
| ERCC5 |
| ERCC6L |
| ERCC8 |
| ERF |
| ERGIC1 |
| ERGIC2 |
| ERGIC3 |
| ERH |
| ERI2 |
| ERI3 |
| ERLEC1 |
| ERLIN1 |
| ERLIN2 |
| ERMP1 |
| ERN1 |
| ERP27 |
| ERP29 |
| ERP44 |
| ERRFI1 |
| ESD |
| ESF1 |
| ESPL1 |
| ESPNL |
| ESR1 |
| ESR2 |
| ESRP1 |

| |
|---|
| ESRP2 |
| ESRRA |
| ESRRG |
| ESYT1 |
| ETAA1 |
| ETF1 |
| ETFA |
| ETFB |
| ETFDH |
| ETHE1 |
| ETNK1 |
| ETNK2 |
| ETS1 |
| ETS2 |
| ETV2 |
| ETV3 |
| ETV6 |
| EVC |
| EVI5 |
| EVI5L |
| EVL |
| EVPL |
| EVX1 |
| EWSR1 |
| EXD2 |
| EXD3 |
| EXO1 |
| EXOC1 |
| EXOC2 |
| EXOC3 |
| EXOC3L2 |
| EXOC4 |
| EXOC5 |
| EXOC6 |
| EXOC6B |
| EXOC7 |
| EXOC8 |
| EXOG |
| EXOSC1 |
| EXOSC10 |
| EXOSC2 |
| EXOSC3 |
| EXOSC4 |
| EXOSC5 |
| EXOSC8 |

| |
|---|
| EXOSC9 |
| EXPH5 |
| EXT1 |
| EXT2 |
| EXTL1 |
| EXTL2 |
| EXTL3 |
| EZH1 |
| EZH2 |
| F11R |
| F12 |
| F2RL1 |
| F2RL3 |
| F3 |
| F5 |
| FA2H |
| FAAH |
| FAAP24 |
| FABP3 |
| FABP4 |
| FABP5 |
| FABP7 |
| FADD |
| FADS1 |
| FADS2 |
| FADS3 |
| FAF1 |
| FAF2 |
| FAH |
| FAHD1 |
| FAHD2A |
| FAIM |
| FAM110A |
| FAM110B |
| FAM114A1 |
| FAM117A |
| FAM118B |
| FAM120A |
| FAM120B |
| FAM149B1 |
| FAM151B |
| FAM155A |
| FAM177A1 |
| FAM193B |
| FAM20B |

| |
|---|
| FAM32A |
| FAM50A |
| FAM53C |
| FAM72A |
| FAM76A |
| FAM83D |
| FAM8A1 |
| FAM92A1 |
| FANCA |
| FANCB |
| FANCD2 |
| FANCE |
| FANCF |
| FANCG |
| FANCI |
| FANCL |
| FANCM |
| FAR1 |
| FAR2 |
| FARS2 |
| FARSA |
| FARSB |
| FAS |
| FASTK |
| FASTKD1 |
| FASTKD2 |
| FASTKD3 |
| FASTKD5 |
| FAT1 |
| FAT2 |
| FAT4 |
| FAU |
| FBF1 |
| FBLIM1 |
| FBLN1 |
| FBLN2 |
| FBLN5 |
| FBN1 |
| FBP1 |
| FBP2 |
| FBRSL1 |
| FBXL12 |
| FBXL14 |
| FBXL15 |
| FBXL18 |

| |
|---|
| FBXL19 |
| FBXL2 |
| FBXL20 |
| FBXL22 |
| FBXL4 |
| FBXL5 |
| FBXL7 |
| FBXL8 |
| FBXO10 |
| FBXO11 |
| FBXO15 |
| FBXO17 |
| FBXO21 |
| FBXO22 |
| FBXO24 |
| FBXO25 |
| FBXO28 |
| FBXO3 |
| FBXO31 |
| FBXO32 |
| FBXO33 |
| FBXO34 |
| FBXO38 |
| FBXO4 |
| FBXO42 |
| FBXO43 |
| FBXO45 |
| FBXO5 |
| FBXO6 |
| FBXO7 |
| FBXO8 |
| FBXO9 |
| FBXW11 |
| FBXW12 |
| FBXW2 |
| FBXW4 |
| FBXW5 |
| FBXW9 |
| FCGBP |
| FCGRT |
| FCHO1 |
| FCHO2 |
| FCHSD1 |
| FCHSD2 |
| FCRL3 |

| |
|---|
| FDFT1 |
| FDPS |
| FDX1 |
| FDXR |
| FECH |
| FEM1A |
| FEM1B |
| FEM1C |
| FEN1 |
| FER |
| FER1L6 |
| FERMT2 |
| FES |
| FGD1 |
| FGD2 |
| FGD3 |
| FGD4 |
| FGD5 |
| FGD6 |
| FGF1 |
| FGF13 |
| FGF2 |
| FGFBP1 |
| FGFR1OP2 |
| FGFR2 |
| FGFR3 |
| FGGY |
| FGL2 |
| FGR |
| FH |
| FHDC1 |
| FHL2 |
| FHL3 |
| FHOD1 |
| FHOD3 |
| FIBP |
| FICD |
| FIG4 |
| FIGN |
| FIGNL1 |
| FILIP1 |
| FILIP1L |
| FIP1L1 |
| FITM2 |
| FIZ1 |

| |
|---|
| FKBP10 |
| FKBP11 |
| FKBP15 |
| FKBP3 |
| FKBP5 |
| FKBP7 |
| FKBP8 |
| FKBP9 |
| FKBPL |
| FKRP |
| FKTN |
| FLAD1 |
| FLCN |
| FLI1 |
| FLII |
| FLNA |
| FLNB |
| FLOT1 |
| FLOT2 |
| FLRT1 |
| FLT3 |
| FLVCR1 |
| FLVCR2 |
| FLYWCH1 |
| FLYWCH2 |
| FMN1 |
| FMNL1 |
| FMNL3 |
| FMO5 |
| FMR1 |
| FN3K |
| FN3KRP |
| FNBP1 |
| FNBP1L |
| FNDC1 |
| FNDC3A |
| FNDC3B |
| FNDC4 |
| FNIP1 |
| FNIP2 |
| FNTA |
| FNTB |
| FOLR1 |
| FOLR2 |
| FOS |

| |
|---|
| FOSB |
| FOSL2 |
| FOXA1 |
| FOXA3 |
| FOXC2 |
| FOXG1 |
| FOXH1 |
| FOXJ3 |
| FOXK2 |
| FOXL2 |
| FOXM1 |
| FOXN2 |
| FOXO1 |
| FOXO3 |
| FOXO4 |
| FOXP1 |
| FOXP4 |
| FOXRED1 |
| FOXRED2 |
| FOXS1 |
| FPGS |
| FPGT |
| FRAT1 |
| FRAT2 |
| FRG1 |
| FRK |
| FRMD4B |
| FRMD6 |
| FRMD8 |
| FRMPD1 |
| FRRS1 |
| FRS2 |
| FRS3 |
| FRY |
| FRYL |
| FSCN1 |
| FSD1L |
| FSD2 |
| FST |
| FSTL1 |
| FSTL3 |
| FTH1 |
| FTO |
| FTSJ1 |
| FTSJ3 |

| |
|---|
| FUBP1 |
| FUBP3 |
| FUCA1 |
| FUCA2 |
| FUNDC2 |
| FURIN |
| FUS |
| FUT1 |
| FUT2 |
| FUT4 |
| FUZ |
| FXR1 |
| FXR2 |
| FXYD3 |
| FXYD5 |
| FXYD6 |
| FYCO1 |
| FYTTD1 |
| FZD1 |
| FZD2 |
| FZD3 |
| FZD4 |
| FZD5 |
| FZD7 |
| FZD8 |
| FZD9 |
| FZR1 |
| G0S2 |
| G2E3 |
| G3BP1 |
| G3BP2 |
| G6PC3 |
| G6PD |
| GAA |
| GAB1 |
| GAB2 |
| GABARAP |
| GABBR1 |
| GABPA |
| GABPB1 |
| GABRA2 |
| GABRA3 |
| GABRG1 |
| GABRR2 |
| GADD45A |

| |
|---|
| GADD45B |
| GADD45G |
| GADD45GIP1 |
| GADL1 |
| GAK |
| GAL3ST1 |
| GAL3ST2 |
| GAL3ST3 |
| GAL3ST4 |
| GALC |
| GALE |
| GALK1 |
| GALK2 |
| GALM |
| GALNS |
| GALNT1 |
| GALNT10 |
| GALNT11 |
| GALNT13 |
| GALNT2 |
| GALNT3 |
| GALNT4 |
| GALNT5 |
| GALNT8 |
| GALR3 |
| GALT |
| GAMT |
| GAN |
| GANAB |
| GANC |
| GAPDHS |
| GAPT |
| GAR1 |
| GARNL3 |
| GAS1 |
| GAS2L1 |
| GAS6 |
| GAS7 |
| GAS8 |
| GATA4 |
| GATAD2A |
| GATAD2B |
| GATC |
| GBA |
| GBA2 |

| |
|---|
| GBA3 |
| GBE1 |
| GBGT1 |
| GBP6 |
| GCA |
| GCAT |
| GCC1 |
| GCC2 |
| GCDH |
| GCH1 |
| GCHFR |
| GCLM |
| GCNT1 |
| GCNT2 |
| GCNT3 |
| GCSH |
| GDA |
| GDAP2 |
| GDE1 |
| GDF1 |
| GDF9 |
| GDI1 |
| GDI2 |
| GDPD3 |
| GDPD5 |
| GEM |
| GEMIN4 |
| GEMIN6 |
| GEMIN7 |
| GEMIN8 |
| GEN1 |
| GFER |
| GFM1 |
| GFM2 |
| GFOD2 |
| GFPT1 |
| GFPT2 |
| GGA1 |
| GGA3 |
| GGCT |
| GGCX |
| GGH |
| GGNBP2 |
| GGPS1 |
| GGT1 |

| |
|---|
| GGT7 |
| GGTA1 |
| GHDC |
| GHITM |
| GHR |
| GIGYF1 |
| GIMAP1 |
| GIMAP4 |
| GIMAP6 |
| GIMAP7 |
| GIMAP8 |
| GIN1 |
| GINS1 |
| GINS3 |
| GINS4 |
| GIPC1 |
| GIPC2 |
| GIT1 |
| GIT2 |
| GJA1 |
| GJA5 |
| GJA8 |
| GJB3 |
| GJC1 |
| GK |
| GK2 |
| GK5 |
| GKAP1 |
| GLB1 |
| GLB1L |
| GLCCI1 |
| GLDC |
| GLDN |
| GLG1 |
| GLI3 |
| GLI4 |
| GLIPR2 |
| GLIS2 |
| GLMN |
| GLO1 |
| GLOD4 |
| GLRX |
| GLRX2 |
| GLRX3 |
| GLRX5 |

| |
|---|
| GLS |
| GLS2 |
| GLT1D1 |
| GLT8D1 |
| GLT8D2 |
| GLUD1 |
| GLUL |
| GLYCTK |
| GLYR1 |
| GM2A |
| GMCL1 |
| GMDS |
| GMEB1 |
| GMEB2 |
| GMFB |
| GMFG |
| GMIP |
| GMNN |
| GMPPA |
| GMPR |
| GMPR2 |
| GMPS |
| GNA11 |
| GNA12 |
| GNA13 |
| GNA14 |
| GNAI1 |
| GNAI2 |
| GNAL |
| GNAQ |
| GNAS |
| GNB1 |
| GNB1L |
| GNB2 |
| GNB4 |
| GNB5 |
| GNE |
| GNG10 |
| GNG11 |
| GNG12 |
| GNG3 |
| GNG5 |
| GNG7 |
| GNL1 |
| GNL2 |

| |
|---|
| GNL3 |
| GNPAT |
| GNPDA1 |
| GNPDA2 |
| GNPNAT1 |
| GNPTAB |
| GNPTG |
| GNS |
| GOLGA2 |
| GOLGA3 |
| GOLGA5 |
| GOLGA7 |
| GOLGB1 |
| GOLIM4 |
| GOLM1 |
| GOLPH3 |
| GOLPH3L |
| GOLT1B |
| GON4L |
| GOPC |
| GORAB |
| GORASP2 |
| GOSR1 |
| GOSR2 |
| GOT1 |
| GOT2 |
| GP1BA |
| GP5 |
| GPA33 |
| GPAA1 |
| GPAM |
| GPATCH1 |
| GPATCH3 |
| GPATCH4 |
| GPATCH8 |
| GPBAR1 |
| GPBP1 |
| GPBP1L1 |
| GPC1 |
| GPC2 |
| GPC4 |
| GPC6 |
| GPCPD1 |
| GPD1 |
| GPD1L |

| |
|---|
| GPD2 |
| GPHN |
| GPI |
| GPKOW |
| GPN1 |
| GPN2 |
| GPN3 |
| GPNMB |
| GPR101 |
| GPR108 |
| GPR132 |
| GPR135 |
| GPR137 |
| GPR137B |
| GPR141 |
| GPR143 |
| GPR146 |
| GPR15 |
| GPR153 |
| GPR155 |
| GPR157 |
| GPR162 |
| GPR173 |
| GPR176 |
| GPR179 |
| GPR18 |
| GPR183 |
| GPR3 |
| GPR31 |
| GPR35 |
| GPR39 |
| GPR4 |
| GPR52 |
| GPR63 |
| GPR65 |
| GPR68 |
| GPR75 |
| GPR84 |
| GPR85 |
| GPR88 |
| GPRASP1 |
| GPRC5A |
| GPRC5B |
| GPRC5C |
| GPRC5D |

| |
|---|
| GPRIN3 |
| GPS1 |
| GPSM1 |
| GPSM2 |
| GPT |
| GPT2 |
| GPX1 |
| GPX3 |
| GPX7 |
| GPX8 |
| GRAMD1A |
| GRAMD1B |
| GRAMD1C |
| GRB2 |
| GRB7 |
| GREB1 |
| GREM1 |
| GRHL1 |
| GRHPR |
| GRID2 |
| GRIK5 |
| GRIN2C |
| GRIN3A |
| GRIN3B |
| GRINA |
| GRIP1 |
| GRIPAP1 |
| GRK1 |
| GRK4 |
| GRK5 |
| GRK6 |
| GRM2 |
| GRM3 |
| GRM8 |
| GRN |
| GRPEL1 |
| GRPEL2 |
| GRSF1 |
| GRTP1 |
| GRWD1 |
| GSDMB |
| GSG1 |
| GSK3A |
| GSPT1 |
| GSS |

| |
|---|
| GSTA1 |
| GSTA4 |
| GSTCD |
| GSTK1 |
| GSTM3 |
| GSTO1 |
| GSTP1 |
| GSTZ1 |
| GSX1 |
| GTDC1 |
| GTF2A1 |
| GTF2A2 |
| GTF2B |
| GTF2E1 |
| GTF2E2 |
| GTF2F1 |
| GTF2F2 |
| GTF2H1 |
| GTF2H2 |
| GTF2H3 |
| GTF2I |
| GTF2IRD1 |
| GTF3A |
| GTF3C1 |
| GTF3C2 |
| GTF3C3 |
| GTF3C4 |
| GTF3C5 |
| GTPBP1 |
| GTPBP2 |
| GTPBP3 |
| GTPBP4 |
| GTPBP8 |
| GTSF1L |
| GUCA2B |
| GUCY2C |
| GUCY2D |
| GUF1 |
| GUK1 |
| GULP1 |
| GUSB |
| GXYLT1 |
| GYG1 |
| GYG2 |
| GYS1 |

| |
|---|
| GZF1 |
| GZMA |
| H6PD |
| HABP4 |
| HACE1 |
| HACL1 |
| HADH |
| HAGH |
| HAGHL |
| HAPLN1 |
| HAPLN3 |
| HARBI1 |
| HAS3 |
| HAT1 |
| HAUS1 |
| HAUS2 |
| HAUS3 |
| HAUS4 |
| HAUS5 |
| HAUS6 |
| HAUS7 |
| HAUS8 |
| HAVCR2 |
| HAX1 |
| HBB |
| HBE1 |
| HBEGF |
| HBP1 |
| HBS1L |
| HCCS |
| HCFC1 |
| HCFC1R1 |
| HCFC2 |
| HCK |
| HCN1 |
| HCRT |
| HDAC1 |
| HDAC10 |
| HDAC11 |
| HDAC2 |
| HDAC3 |
| HDAC5 |
| HDAC6 |
| HDAC8 |
| HDC |

| |
|---|
| HDDC2 |
| HDDC3 |
| HDGF |
| HDHD2 |
| HDHD3 |
| HDLBP |
| HEATR3 |
| HEATR4 |
| HEATR5B |
| HEATR6 |
| HEBP2 |
| HECTD2 |
| HECTD3 |
| HEG1 |
| HELB |
| HELLS |
| HELQ |
| HELZ |
| HEMK1 |
| HEPACAM2 |
| HERC1 |
| HERC2 |
| HERC4 |
| HERC5 |
| HERPUD2 |
| HES4 |
| HES6 |
| HEXA |
| HEXB |
| HEXIM1 |
| HEXIM2 |
| HEY2 |
| HFE |
| HFM1 |
| HGF |
| HGS |
| HHEX |
| HHIPL2 |
| HIBADH |
| HIC2 |
| HIF1A |
| HIF1AN |
| HIGD2A |
| HINFP |
| HINT1 |

| |
|---|
| HINT2 |
| HINT3 |
| HIP1R |
| HIPK1 |
| HIPK3 |
| HIRA |
| HIRIP3 |
| HIVEP1 |
| HIVEP2 |
| HIVEP3 |
| HJURP |
| HK1 |
| HK2 |
| HK3 |
| HKDC1 |
| HLCS |
| HLX |
| HM13 |
| HMBS |
| HMCN1 |
| HMG20A |
| HMG20B |
| HMGA1 |
| HMGB1 |
| HMGB3 |
| HMGCL |
| HMGCR |
| HMGCS1 |
| HMGN1 |
| HMGN2 |
| HMGN3 |
| HMGN4 |
| HMGN5 |
| HMGXB4 |
| HMMR |
| HMOX1 |
| HMOX2 |
| HNF4A |
| HNF4G |
| HNMT |
| HNRNPA0 |
| HNRNPA1 |
| HNRNPA3 |
| HNRNPAB |
| HNRNPC |

| |
|---|
| HNRNPD |
| HNRNPF |
| HNRNPH1 |
| HNRNPH2 |
| HNRNPH3 |
| HNRNPK |
| HNRNPL |
| HNRNPM |
| HNRNPR |
| HNRNPU |
| HNRNPUL1 |
| HNRNPUL2 |
| HOMER1 |
| HOMER2 |
| HOMEZ |
| HOOK1 |
| HOOK2 |
| HOOK3 |
| HOXA10 |
| HOXC11 |
| HOXC4 |
| HOXC5 |
| HOXC6 |
| HOXD11 |
| HOXD4 |
| HP1BP3 |
| HPDL |
| HPS1 |
| HPS3 |
| HPS4 |
| HPS6 |
| HPSE |
| HRH1 |
| HRH2 |
| HS1BP3 |
| HS2ST1 |
| HS3ST1 |
| HS3ST2 |
| HS3ST3A1 |
| HS6ST1 |
| HS6ST3 |
| HSD11B1L |
| HSD11B2 |
| HSD17B1 |
| HSD17B10 |

| |
|---|
| HSD17B14 |
| HSD17B2 |
| HSD17B3 |
| HSD17B4 |
| HSD17B7 |
| HSD17B8 |
| HSD3B7 |
| HSDL1 |
| HSDL2 |
| HSF1 |
| HSF2 |
| HSF2BP |
| HSF4 |
| HSP90AA1 |
| HSP90AB1 |
| HSP90B1 |
| HSPA12B |
| HSPA13 |
| HSPA14 |
| HSPA1A |
| HSPA1L |
| HSPA4L |
| HSPA5 |
| HSPA6 |
| HSPA8 |
| HSPA9 |
| HSPB1 |
| HSPB6 |
| HSPBAP1 |
| HSPBP1 |
| HSPD1 |
| HSPE1 |
| HSPG2 |
| HTATIP2 |
| HTATSF1 |
| HTR1F |
| HTR3A |
| HTR5A |
| HTR7 |
| HTRA1 |
| HTRA2 |
| HUS1 |
| HUWE1 |
| HVCN1 |
| HYAL1 |

| |
|---|
| HYAL2 |
| HYLS1 |
| IAH1 |
| IARS2 |
| IBTK |
| ICAM1 |
| ICAM2 |
| ICAM3 |
| ICOSLG |
| ID1 |
| ID2 |
| ID3 |
| IDE |
| IDH1 |
| IDH2 |
| IDH3A |
| IDH3B |
| IDI1 |
| IDO1 |
| IDS |
| IDUA |
| IER2 |
| IER3 |
| IER3IP1 |
| IER5 |
| IER5L |
| IFFO2 |
| IFI16 |
| IFI27L2 |
| IFI30 |
| IFI35 |
| IFI6 |
| IFIH1 |
| IFIT1 |
| IFIT2 |
| IFIT5 |
| IFITM3 |
| IFNAR1 |
| IFNAR2 |
| IFNGR1 |
| IFNGR2 |
| IFNW1 |
| IFRD1 |
| IFRD2 |
| IFT122 |

| |
|---|
| IFT20 |
| IFT52 |
| IFT57 |
| IFT74 |
| IFT80 |
| IFT81 |
| IFT88 |
| IGBP1 |
| IGDCC4 |
| IGF1R |
| IGF2 |
| IGF2BP2 |
| IGFBP2 |
| IGFBP3 |
| IGFBP4 |
| IGFBP5 |
| IGFBP6 |
| IGFBP7 |
| IGHMBP2 |
| IGSF11 |
| IGSF6 |
| IGSF8 |
| IGSF9 |
| IHH |
| IK |
| IKBKB |
| IKBKE |
| IKBKG |
| IKZF1 |
| IKZF2 |
| IKZF3 |
| IKZF4 |
| IKZF5 |
| IL10 |
| IL10RA |
| IL10RB |
| IL11RA |
| IL13RA1 |
| IL15 |
| IL16 |
| IL17D |
| IL17RA |
| IL17RC |
| IL17RD |
| IL18 |

| |
|---|
| IL18BP |
| IL18RAP |
| IL1A |
| IL1RAP |
| IL1RN |
| IL20RA |
| IL27RA |
| IL2RB |
| IL2RG |
| IL34 |
| IL4R |
| IL6 |
| IL6ST |
| ILDR1 |
| ILF2 |
| ILF3 |
| ILK |
| ILKAP |
| ILVBL |
| IMMT |
| IMP3 |
| IMP4 |
| IMPA1 |
| IMPA2 |
| IMPACT |
| IMPDH1 |
| IMPDH2 |
| INCENP |
| ING1 |
| ING3 |
| ING4 |
| ING5 |
| INHA |
| INHBA |
| INHBB |
| INO80 |
| INO80B |
| INO80C |
| INO80D |
| INO80E |
| INPP1 |
| INPP4A |
| INPP5A |
| INPP5B |
| INPP5D |

| |
|---|
| INPP5E |
| INPP5F |
| INPP5K |
| INPPL1 |
| INSIG1 |
| INSIG2 |
| INSL6 |
| INSR |
| INTS10 |
| INTS12 |
| INTS2 |
| INTS3 |
| INTS5 |
| INTS6 |
| INTS7 |
| INTS8 |
| INTS9 |
| INTU |
| IP6K1 |
| IPMK |
| IPO13 |
| IPO4 |
| IPO7 |
| IPO8 |
| IPP |
| IPPK |
| IQCB1 |
| IQCD |
| IQCE |
| IQCG |
| IQGAP1 |
| IQGAP2 |
| IQGAP3 |
| IQSEC2 |
| IRAK1 |
| IRAK1BP1 |
| IRAK2 |
| IRAK3 |
| IRAK4 |
| IREB2 |
| IRF1 |
| IRF2 |
| IRF2BP2 |
| IRF3 |
| IRF6 |

| |
|---|
| IRF7 |
| IRF8 |
| IRS1 |
| IRX3 |
| IRX5 |
| ISCU |
| ISG20 |
| ISG20L2 |
| ISLR2 |
| ISOC1 |
| IST1 |
| ISY1 |
| ISYNA1 |
| ITFG1 |
| ITFG2 |
| ITGA3 |
| ITGA4 |
| ITGA6 |
| ITGAE |
| ITGAL |
| ITGAM |
| ITGAV |
| ITGB1 |
| ITGB1BP1 |
| ITGB1BP2 |
| ITGB3 |
| ITGB3BP |
| ITGB5 |
| ITGB7 |
| ITGB8 |
| ITGBL1 |
| ITIH1 |
| ITM2B |
| ITM2C |
| ITPA |
| ITPK1 |
| ITPKA |
| ITPKB |
| ITPKC |
| ITPR2 |
| ITPR3 |
| ITPRIP |
| ITPRIPL1 |
| ITPRIPL2 |
| ITSN1 |

| |
|---|
| IVD |
| IVNS1ABP |
| IWS1 |
| JAG1 |
| JAGN1 |
| JAK1 |
| JAKMIP2 |
| JAM3 |
| JARID2 |
| JAZF1 |
| JDP2 |
| JKAMP |
| JMJD1C |
| JMJD4 |
| JMJD6 |
| JMJD7 |
| JMJD8 |
| JMY |
| JOSD1 |
| JRK |
| JRKL |
| JSRP1 |
| JTB |
| JUN |
| JUNB |
| JUND |
| JUP |
| KALRN |
| KANK1 |
| KANK2 |
| KANK3 |
| KAT2B |
| KAT5 |
| KATNA1 |
| KATNAL1 |
| KATNB1 |
| KAZALD1 |
| KBTBD2 |
| KBTBD3 |
| KBTBD6 |
| KBTBD7 |
| KCMF1 |
| KCNA2 |
| KCNA4 |
| KCNA5 |

| |
|---|
| KCNA6 |
| KCNAB1 |
| KCNAB2 |
| KCNC2 |
| KCND2 |
| KCNE2 |
| KCNE3 |
| KCNE4 |
| KCNG2 |
| KCNH2 |
| KCNIP4 |
| KCNJ10 |
| KCNJ14 |
| KCNJ15 |
| KCNJ16 |
| KCNJ2 |
| KCNJ5 |
| KCNK1 |
| KCNK3 |
| KCNK5 |
| KCNN3 |
| KCNQ1 |
| KCNQ4 |
| KCNQ5 |
| KCNRG |
| KCNS3 |
| KCTD1 |
| KCTD10 |
| KCTD11 |
| KCTD12 |
| KCTD15 |
| KCTD18 |
| KCTD20 |
| KCTD21 |
| KCTD3 |
| KCTD5 |
| KCTD7 |
| KCTD9 |
| KDELC1 |
| KDELR1 |
| KDELR2 |
| KDELR3 |
| KDM1A |
| KDM1B |
| KDM2A |

| |
|---|
| KDM2B |
| KDM3A |
| KDM4B |
| KDM4C |
| KDM5A |
| KDM5B |
| KDM5C |
| KDM6A |
| KDSR |
| KEAP1 |
| KHDRBS1 |
| KHNYN |
| KIAA0586 |
| KIAA1549 |
| KIAA2013 |
| KIF11 |
| KIF13A |
| KIF13B |
| KIF14 |
| KIF15 |
| KIF16B |
| KIF18A |
| KIF18B |
| KIF19 |
| KIF1C |
| KIF20A |
| KIF20B |
| KIF22 |
| KIF23 |
| KIF24 |
| KIF2A |
| KIF2C |
| KIF3A |
| KIF3B |
| KIF3C |
| KIF4A |
| KIF7 |
| KIFAP3 |
| KIFC1 |
| KIN |
| KIRREL3 |
| KIT |
| KITLG |
| KLC1 |
| KLC2 |

| |
|---|
| KLC3 |
| KLC4 |
| KLF11 |
| KLF12 |
| KLF13 |
| KLF15 |
| KLF16 |
| KLF4 |
| KLF5 |
| KLF6 |
| KLF7 |
| KLF8 |
| KLHDC1 |
| KLHDC10 |
| KLHDC2 |
| KLHDC4 |
| KLHDC7A |
| KLHDC8B |
| KLHL11 |
| KLHL12 |
| KLHL15 |
| KLHL17 |
| KLHL18 |
| KLHL2 |
| KLHL20 |
| KLHL21 |
| KLHL23 |
| KLHL24 |
| KLHL25 |
| KLHL26 |
| KLHL28 |
| KLHL3 |
| KLHL32 |
| KLHL36 |
| KLHL38 |
| KLHL5 |
| KLHL7 |
| KLHL8 |
| KLHL9 |
| KLK14 |
| KLK15 |
| KPNA3 |
| KPNA4 |
| KPNA5 |
| KPNA6 |

| |
|---|
| KPNB1 |
| KRAS |
| KRBA1 |
| KRBA2 |
| KRCC1 |
| KREMEN1 |
| KRI1 |
| KRIT1 |
| KRR1 |
| KRT18 |
| KRT19 |
| KRT20 |
| KRT72 |
| KRT8 |
| KRTCAP2 |
| KSR1 |
| KTI12 |
| KTN1 |
| L1CAM |
| L2HGDH |
| L3MBTL2 |
| L3MBTL3 |
| LACTB |
| LACTB2 |
| LAD1 |
| LAG3 |
| LAGE3 |
| LAMA1 |
| LAMA5 |
| LAMB1 |
| LAMB2 |
| LAMB3 |
| LAMC1 |
| LAMC3 |
| LAMP1 |
| LAMP2 |
| LANCL2 |
| LAP3 |
| LAPTM4A |
| LAPTM4B |
| LAPTM5 |
| LARP1 |
| LARP4 |
| LARP4B |
| LARP6 |

| |
|---|
| LARP7 |
| LARS2 |
| LAS1L |
| LASP1 |
| LAT |
| LATS2 |
| LBH |
| LBR |
| LBX2 |
| LCK |
| LCLAT1 |
| LCMT1 |
| LCMT2 |
| LCN2 |
| LCOR |
| LCORL |
| LCP1 |
| LCP2 |
| LDB1 |
| LDHA |
| LDHAL6B |
| LDHB |
| LDHD |
| LDLR |
| LDLRAD2 |
| LDLRAD3 |
| LDLRAP1 |
| LDOC1 |
| LEF1 |
| LEMD2 |
| LEMD3 |
| LENG1 |
| LENG8 |
| LENG9 |
| LEO1 |
| LEPROT |
| LEPROTL1 |
| LETM1 |
| LETM2 |
| LETMD1 |
| LFNG |
| LGALS3 |
| LGALS3BP |
| LGALS8 |
| LGALS9 |

| |
|---|
| LGMN |
| LGR4 |
| LGSN |
| LHCGR |
| LHFPL2 |
| LHPP |
| LIAS |
| LIF |
| LIFR |
| LIG1 |
| LIG3 |
| LIG4 |
| LIM2 |
| LIMA1 |
| LIMD1 |
| LIMD2 |
| LIMK1 |
| LIMK2 |
| LIN37 |
| LIN52 |
| LIN54 |
| LIN7B |
| LIN7C |
| LINGO2 |
| LINS1 |
| LIPA |
| LIPC |
| LIPE |
| LIPH |
| LIPK |
| LIPT1 |
| LIPT2 |
| LITAF |
| LIX1L |
| LLGL2 |
| LLPH |
| LMAN1 |
| LMAN2 |
| LMAN2L |
| LMBRD1 |
| LMBRD2 |
| LMCD1 |
| LMF1 |
| LMF2 |
| LMNA |

| |
|---|
| LMNB1 |
| LMNB2 |
| LMO2 |
| LMO4 |
| LMO7 |
| LMTK2 |
| LNPEP |
| LNX2 |
| LONP1 |
| LONP2 |
| LONRF1 |
| LONRF3 |
| LOXL3 |
| LOXL4 |
| LPAR2 |
| LPAR5 |
| LPAR6 |
| LPCAT3 |
| LPCAT4 |
| LPGAT1 |
| LPIN1 |
| LPIN3 |
| LPP |
| LPXN |
| LRBA |
| LRCH1 |
| LRCH2 |
| LRCH3 |
| LRCH4 |
| LRFN3 |
| LRFN4 |
| LRIG1 |
| LRIG2 |
| LRIG3 |
| LRP10 |
| LRP12 |
| LRP4 |
| LRP5 |
| LRP6 |
| LRP8 |
| LRPAP1 |
| LRPPRC |
| LRRC1 |
| LRRC10B |
| LRRC14 |

| |
|---|
| LRRC17 |
| LRRC20 |
| LRRC24 |
| LRRC25 |
| LRRC27 |
| LRRC28 |
| LRRC31 |
| LRRC4 |
| LRRC40 |
| LRRC41 |
| LRRC42 |
| LRRC45 |
| LRRC46 |
| LRRC47 |
| LRRC49 |
| LRRC4B |
| LRRC4C |
| LRRC57 |
| LRRC58 |
| LRRC59 |
| LRRC61 |
| LRRC66 |
| LRRC70 |
| LRRC8A |
| LRRC8D |
| LRRCC1 |
| LRRFIP1 |
| LRRFIP2 |
| LRRK1 |
| LRRK2 |
| LRRN2 |
| LRRN3 |
| LRRTM2 |
| LRSAM1 |
| LRWD1 |
| LSG1 |
| LSM1 |
| LSM10 |
| LSM11 |
| LSM12 |
| LSM14A |
| LSM14B |
| LSM3 |
| LSM4 |
| LSM5 |

| |
|---|
| LSM6 |
| LSM7 |
| LSR |
| LSS |
| LTA4H |
| LTB |
| LTB4R |
| LTBP1 |
| LTBP3 |
| LTBR |
| LTV1 |
| LUC7L |
| LUC7L2 |
| LUC7L3 |
| LUZP1 |
| LXN |
| LY6G5B |
| LY75 |
| LY9 |
| LYAR |
| LYN |
| LYNX1 |
| LYPLA1 |
| LYPLA2 |
| LYRM1 |
| LYRM2 |
| LYRM4 |
| LYRM7 |
| LYSMD1 |
| LYSMD2 |
| LYSMD3 |
| LYST |
| LYZ |
| LZTFL1 |
| LZTR1 |
| LZTS1 |
| LZTS2 |
| M6PR |
| MACROD1 |
| MAD1L1 |
| MAD2L1 |
| MAD2L1BP |
| MAD2L2 |
| MADD |
| MAEA |

| |
|---|
| MAF |
| MAF1 |
| MAFB |
| MAFK |
| MAGEA10 |
| MAGED1 |
| MAGED2 |
| MAGEE2 |
| MAGEL2 |
| MAGI3 |
| MAGIX |
| MAGOH |
| MAGOHB |
| MAK16 |
| MAL2 |
| MAMDC4 |
| MAML2 |
| MAML3 |
| MAMLD1 |
| MAMSTR |
| MAN1A1 |
| MAN1A2 |
| MAN1C1 |
| MAN2A1 |
| MAN2A2 |
| MAN2B1 |
| MAN2C1 |
| MANBA |
| MANBAL |
| MANEA |
| MANEAL |
| MANF |
| MAOA |
| MAOB |
| MAP1B |
| MAP1LC3B |
| MAP2 |
| MAP2K1 |
| MAP2K2 |
| MAP2K3 |
| MAP2K5 |
| MAP2K6 |
| MAP2K7 |
| MAP3K1 |
| MAP3K10 |

| |
|---|
| MAP3K11 |
| MAP3K12 |
| MAP3K14 |
| MAP3K15 |
| MAP3K2 |
| MAP3K3 |
| MAP3K4 |
| MAP3K5 |
| MAP3K6 |
| MAP3K7 |
| MAP3K8 |
| MAP4 |
| MAP4K1 |
| MAP4K2 |
| MAP4K3 |
| MAP4K5 |
| MAP6 |
| MAP7 |
| MAP7D1 |
| MAP7D3 |
| MAP9 |
| MAPK11 |
| MAPK12 |
| MAPK13 |
| MAPK14 |
| MAPK15 |
| MAPK4 |
| MAPK6 |
| MAPK7 |
| MAPK8 |
| MAPK8IP1 |
| MAPK8IP3 |
| MAPKAP1 |
| MAPKAPK3 |
| MAPKBP1 |
| MAPRE2 |
| MARCKSL1 |
| MARK1 |
| MARK2 |
| MARK3 |
| MARK4 |
| MARS2 |
| MARVELD1 |
| MARVELD2 |
| MARVELD3 |

| |
|---|
| MASP2 |
| MAST1 |
| MAST3 |
| MASTL |
| MAT2B |
| MATK |
| MATN4 |
| MATR3 |
| MAVS |
| MAX |
| MBD1 |
| MBD3 |
| MBD4 |
| MBD5 |
| MBIP |
| MBLAC1 |
| MBLAC2 |
| MBNL2 |
| MBNL3 |
| MBOAT2 |
| MBP |
| MBTD1 |
| MBTPS1 |
| MBTPS2 |
| MCAM |
| MCAT |
| MCC |
| MCCC1 |
| MCEE |
| MCF2L |
| MCF2L2 |
| MCFD2 |
| MCL1 |
| MCM10 |
| MCM2 |
| MCM3 |
| MCM3AP |
| MCM4 |
| MCM5 |
| MCM6 |
| MCM7 |
| MCM8 |
| MCM9 |
| MCOLN1 |
| MCPH1 |

| |
|---|
| MCRS1 |
| MCTP1 |
| MCTP2 |
| MCTS1 |
| MDC1 |
| MDFIC |
| MDH2 |
| MDK |
| MDM2 |
| MDM4 |
| MDN1 |
| ME1 |
| ME2 |
| ME3 |
| MEA1 |
| MEAF6 |
| MECOM |
| MECP2 |
| MECR |
| MED1 |
| MED10 |
| MED11 |
| MED12 |
| MED12L |
| MED13 |
| MED13L |
| MED15 |
| MED17 |
| MED18 |
| MED19 |
| MED21 |
| MED22 |
| MED23 |
| MED24 |
| MED25 |
| MED26 |
| MED27 |
| MED28 |
| MED29 |
| MED30 |
| MED31 |
| MED4 |
| MED7 |
| MED9 |
| MEF2A |

| |
|---|
| MEF2C |
| MEFV |
| MEGF9 |
| MEI1 |
| MEIS1 |
| MEIS2 |
| MELK |
| MEN1 |
| MEP1B |
| MEPCE |
| MERTK |
| MESP2 |
| MEST |
| METRN |
| METRNL |
| METTL1 |
| METTL13 |
| METTL14 |
| METTL3 |
| METTL4 |
| METTL5 |
| METTL6 |
| METTL7A |
| METTL7B |
| METTL8 |
| METTL9 |
| MEX3A |
| MEX3B |
| MEX3C |
| MFAP1 |
| MFAP2 |
| MFAP3 |
| MFAP3L |
| MFGE8 |
| MFN1 |
| MFRP |
| MFSD1 |
| MFSD10 |
| MFSD11 |
| MFSD3 |
| MFSD6 |
| MFSD6L |
| MFSD8 |
| MFSD9 |
| MGA |

| |
|---|
| MGAT1 |
| MGAT2 |
| MGAT4A |
| MGAT4B |
| MGAT5 |
| MGMT |
| MGRN1 |
| MGST1 |
| MGST2 |
| MGST3 |
| MIA3 |
| MIB1 |
| MIB2 |
| MICAL1 |
| MICAL3 |
| MICALL1 |
| MICALL2 |
| MICB |
| MID1 |
| MID1IP1 |
| MIDN |
| MIER1 |
| MIER2 |
| MIER3 |
| MIF |
| MIF4GD |
| MIIP |
| MINK1 |
| MINPP1 |
| MIOS |
| MIPEP |
| MIPOL1 |
| MIS12 |
| MIS18A |
| MITD1 |
| MITF |
| MKI67 |
| MKKS |
| MKNK1 |
| MKNK2 |
| MKRN2 |
| MKS1 |
| MKX |
| MLEC |
| MLF1 |

| |
|---|
| MLH1 |
| MLH3 |
| MLKL |
| MLLT1 |
| MLLT3 |
| MLLT6 |
| MLST8 |
| MLXIP |
| MLYCD |
| MMAA |
| MMAB |
| MMACHC |
| MMADHC |
| MMD |
| MMGT1 |
| MMP14 |
| MMP15 |
| MMP17 |
| MMP19 |
| MMP2 |
| MMP25 |
| MMP9 |
| MMS19 |
| MN1 |
| MNAT1 |
| MND1 |
| MNS1 |
| MNT |
| MOCOS |
| MOCS1 |
| MOCS2 |
| MOCS3 |
| MON1A |
| MON1B |
| MORC2 |
| MORC3 |
| MORF4L1 |
| MORF4L2 |
| MORN4 |
| MOSPD1 |
| MOSPD3 |
| MOV10 |
| MPDU1 |
| MPDZ |
| MPEG1 |

| |
|---|
| MPG |
| MPHOSPH10 |
| MPHOSPH6 |
| MPHOSPH8 |
| MPHOSPH9 |
| MPI |
| MPL |
| MPND |
| MPP1 |
| MPP2 |
| MPP7 |
| MPPE1 |
| MPPED2 |
| MPRIP |
| MPST |
| MPZL3 |
| MR1 |
| MRAP |
| MRAP2 |
| MRAS |
| MRC2 |
| MRFAP1 |
| MRI1 |
| MRM1 |
| MRO |
| MRPL1 |
| MRPL10 |
| MRPL11 |
| MRPL12 |
| MRPL13 |
| MRPL14 |
| MRPL15 |
| MRPL16 |
| MRPL17 |
| MRPL18 |
| MRPL19 |
| MRPL20 |
| MRPL21 |
| MRPL22 |
| MRPL23 |
| MRPL24 |
| MRPL27 |
| MRPL28 |
| MRPL3 |
| MRPL30 |

| |
|---|
| MRPL32 |
| MRPL33 |
| MRPL34 |
| MRPL35 |
| MRPL36 |
| MRPL37 |
| MRPL38 |
| MRPL39 |
| MRPL4 |
| MRPL41 |
| MRPL42 |
| MRPL43 |
| MRPL44 |
| MRPL45 |
| MRPL46 |
| MRPL47 |
| MRPL48 |
| MRPL49 |
| MRPL51 |
| MRPL52 |
| MRPL53 |
| MRPL54 |
| MRPL55 |
| MRPL9 |
| MRPS10 |
| MRPS11 |
| MRPS15 |
| MRPS16 |
| MRPS17 |
| MRPS18B |
| MRPS18C |
| MRPS2 |
| MRPS21 |
| MRPS22 |
| MRPS23 |
| MRPS24 |
| MRPS25 |
| MRPS26 |
| MRPS27 |
| MRPS28 |
| MRPS30 |
| MRPS31 |
| MRPS33 |
| MRPS34 |
| MRPS35 |

| |
|---|
| MRPS5 |
| MRPS6 |
| MRPS7 |
| MRPS9 |
| MRS2 |
| MSH2 |
| MSH4 |
| MSH5 |
| MSH6 |
| MSI1 |
| MSI2 |
| MSL1 |
| MSL2 |
| MSL3 |
| MSN |
| MSR1 |
| MSRA |
| MSRB2 |
| MSRB3 |
| MST1 |
| MST1R |
| MSTO1 |
| MT1A |
| MT2A |
| MT3 |
| MTA2 |
| MTA3 |
| MTBP |
| MTCH1 |
| MTCH2 |
| MTCP1 |
| MTDH |
| MTF1 |
| MTF2 |
| MTFMT |
| MTFR1 |
| MTG1 |
| MTHFD1L |
| MTHFD2 |
| MTHFD2L |
| MTHFSD |
| MTIF2 |
| MTIF3 |
| MTM1 |
| MTMR1 |

| |
|---|
| MTMR10 |
| MTMR11 |
| MTMR14 |
| MTMR2 |
| MTMR3 |
| MTMR4 |
| MTMR6 |
| MTMR7 |
| MTMR9 |
| MTO1 |
| MTOR |
| MTPAP |
| MTPN |
| MTR |
| MTRF1 |
| MTRF1L |
| MTRR |
| MTSS1 |
| MTTP |
| MTUS1 |
| MTX1 |
| MTX2 |
| MTX3 |
| MUC1 |
| MUC16 |
| MUC2 |
| MUC20 |
| MUL1 |
| MUS81 |
| MVD |
| MVK |
| MVP |
| MX1 |
| MX2 |
| MXD3 |
| MXD4 |
| MXI1 |
| MXRA5 |
| MXRA7 |
| MXRA8 |
| MYADM |
| MYBBP1A |
| MYBL1 |
| MYBL2 |
| MYBPC1 |

| |
|---|
| MYC |
| MYCBP |
| MYCBP2 |
| MYCBPAP |
| MYCN |
| MYD88 |
| MYEF2 |
| MYH10 |
| MYH11 |
| MYH14 |
| MYH9 |
| MYL12B |
| MYL2 |
| MYL6 |
| MYL6B |
| MYL9 |
| MYLIP |
| MYLK |
| MYLPF |
| MYO10 |
| MYO18A |
| MYO19 |
| MYO1A |
| MYO1B |
| MYO1C |
| MYO1D |
| MYO1E |
| MYO1F |
| MYO1G |
| MYO3B |
| MYO5A |
| MYO5C |
| MYO6 |
| MYO7A |
| MYO7B |
| MYO9A |
| MYO9B |
| MYOC |
| MYOCD |
| MYOF |
| MYOM1 |
| MYOZ1 |
| MYPN |
| MYSM1 |
| MZF1 |

| |
|---|
| N4BP1 |
| N4BP2 |
| N4BP2L1 |
| N6AMT1 |
| NAA10 |
| NAA15 |
| NAA16 |
| NAA20 |
| NAA25 |
| NAA30 |
| NAA35 |
| NAA38 |
| NAA40 |
| NAAA |
| NAALADL1 |
| NAB1 |
| NAB2 |
| NACA |
| NACAD |
| NACC1 |
| NACC2 |
| NADK |
| NAE1 |
| NAGA |
| NAGK |
| NAGLU |
| NAGPA |
| NAIF1 |
| NAIP |
| NAMPT |
| NANOS1 |
| NANP |
| NANS |
| NAP1L1 |
| NAP1L5 |
| NAPA |
| NAPB |
| NAPEPLD |
| NAPG |
| NAPSA |
| NARF |
| NASP |
| NAT1 |
| NAT10 |
| NAT14 |

| |
|---|
| NAT9 |
| NAV1 |
| NAV2 |
| NAV3 |
| NBAS |
| NBEA |
| NBEAL2 |
| NBL1 |
| NBN |
| NBR1 |
| NCAM1 |
| NCAM2 |
| NCAPD2 |
| NCAPD3 |
| NCAPG |
| NCAPG2 |
| NCAPH |
| NCAPH2 |
| NCBP1 |
| NCBP2 |
| NCDN |
| NCEH1 |
| NCF1 |
| NCF2 |
| NCF4 |
| NCK1 |
| NCK2 |
| NCKAP1L |
| NCKAP5 |
| NCKAP5L |
| NCKIPSD |
| NCL |
| NCLN |
| NCOA1 |
| NCOA2 |
| NCOA3 |
| NCOA4 |
| NCOA5 |
| NCOA6 |
| NCOA7 |
| NCOR1 |
| NCS1 |
| NCSTN |
| NDC80 |
| NDE1 |

| |
|---|
| NDEL1 |
| NDFIP2 |
| NDOR1 |
| NDRG1 |
| NDRG2 |
| NDST2 |
| NDUFA10 |
| NDUFA12 |
| NDUFA13 |
| NDUFA2 |
| NDUFA3 |
| NDUFA4 |
| NDUFA6 |
| NDUFA8 |
| NDUFAB1 |
| NDUFAF1 |
| NDUFAF3 |
| NDUFAF4 |
| NDUFB1 |
| NDUFB10 |
| NDUFB11 |
| NDUFB2 |
| NDUFB4 |
| NDUFB5 |
| NDUFB6 |
| NDUFB7 |
| NDUFB8 |
| NDUFB9 |
| NDUFC1 |
| NDUFC2 |
| NDUFS2 |
| NDUFS3 |
| NDUFS5 |
| NDUFS6 |
| NDUFS7 |
| NDUFV1 |
| NDUFV2 |
| NDUFV3 |
| NEBL |
| NECAP1 |
| NECAP2 |
| NEDD4 |
| NEDD8 |
| NEDD9 |
| NEFH |

| |
|---|
| NEFL |
| NEGR1 |
| NEIL1 |
| NEIL2 |
| NEIL3 |
| NEK10 |
| NEK2 |
| NEK3 |
| NEK4 |
| NEK6 |
| NEK7 |
| NEK8 |
| NEK9 |
| NELL2 |
| NENF |
| NES |
| NET1 |
| NEU1 |
| NEU3 |
| NEURL1B |
| NEURL2 |
| NEURL4 |
| NEXN |
| NF1 |
| NFAM1 |
| NFAT5 |
| NFATC1 |
| NFATC3 |
| NFE2 |
| NFE2L1 |
| NFE2L3 |
| NFIC |
| NFIL3 |
| NFIX |
| NFKB1 |
| NFKBIA |
| NFKBIB |
| NFKBID |
| NFKBIE |
| NFKBIL1 |
| NFKBIZ |
| NFS1 |
| NFX1 |
| NFYA |
| NFYB |

| |
|---|
| NFYC |
| NGDN |
| NGLY1 |
| NGRN |
| NHEJ1 |
| NHLRC1 |
| NHLRC2 |
| NHLRC3 |
| NHP2 |
| NHS |
| NHSL2 |
| NICN1 |
| NID1 |
| NID2 |
| NIF3L1 |
| NIN |
| NINJ1 |
| NIP7 |
| NIPA1 |
| NIPA2 |
| NIPAL1 |
| NIPAL3 |
| NIPBL |
| NIPSNAP3A |
| NIT1 |
| NIT2 |
| NKAIN1 |
| NKAP |
| NKAPL |
| NKD1 |
| NKG7 |
| NKIRAS1 |
| NKIRAS2 |
| NKPD1 |
| NKRF |
| NKX1-2 |
| NLE1 |
| NLGN2 |
| NLGN3 |
| NLK |
| NLN |
| NLRC3 |
| NLRC4 |
| NLRP3 |
| NLRP5 |

| |
|---|
| NLRP6 |
| NMB |
| NMD3 |
| NME3 |
| NME6 |
| NME7 |
| NMI |
| NMNAT1 |
| NMRAL1 |
| NMT1 |
| NNAT |
| NNT |
| NOB1 |
| NOC2L |
| NOC3L |
| NOC4L |
| NOD1 |
| NOD2 |
| NODAL |
| NOL10 |
| NOL11 |
| NOL12 |
| NOL6 |
| NOL7 |
| NOL8 |
| NOL9 |
| NONO |
| NOP14 |
| NOP16 |
| NOP2 |
| NOP56 |
| NOP58 |
| NOS1AP |
| NOS2 |
| NOSIP |
| NOTCH1 |
| NOTCH2 |
| NOVA1 |
| NOXA1 |
| NOXO1 |
| NPAS3 |
| NPAT |
| NPB |
| NPC2 |
| NPDC1 |

| |
|---|
| NPEPL1 |
| NPEPPS |
| NPHP1 |
| NPHP3 |
| NPHP4 |
| NPL |
| NPLOC4 |
| NPM1 |
| NPM3 |
| NPNT |
| NPR2 |
| NPR3 |
| NPTN |
| NPTX1 |
| NPY1R |
| NQO1 |
| NQO2 |
| NR1D2 |
| NR1H2 |
| NR1H4 |
| NR1I2 |
| NR2C2 |
| NR2C2AP |
| NR2E1 |
| NR2F1 |
| NR2F2 |
| NR3C2 |
| NR4A1 |
| NR4A2 |
| NR4A3 |
| NR5A1 |
| NR5A2 |
| NR6A1 |
| NRAP |
| NRBF2 |
| NRBP1 |
| NRBP2 |
| NRF1 |
| NRIP1 |
| NRIP3 |
| NRM |
| NRP1 |
| NRSN2 |
| NRXN1 |
| NSA2 |

| |
|---|
| NSD1 |
| NSDHL |
| NSF |
| NSFL1C |
| NSL1 |
| NSMAF |
| NSMCE1 |
| NSMCE2 |
| NSMCE4A |
| NSUN3 |
| NSUN4 |
| NSUN5 |
| NSUN6 |
| NSUN7 |
| NT5C |
| NT5C2 |
| NT5DC1 |
| NT5DC3 |
| NT5E |
| NT5M |
| NTAN1 |
| NTHL1 |
| NTN3 |
| NTN4 |
| NTN5 |
| NTRK1 |
| NUAK1 |
| NUAK2 |
| NUB1 |
| NUBP1 |
| NUBP2 |
| NUBPL |
| NUCB1 |
| NUCB2 |
| NUCKS1 |
| NUDC |
| NUDCD1 |
| NUDCD2 |
| NUDCD3 |
| NUDT1 |
| NUDT12 |
| NUDT13 |
| NUDT14 |
| NUDT15 |
| NUDT16 |

| |
|---|
| NUDT16L1 |
| NUDT17 |
| NUDT18 |
| NUDT2 |
| NUDT21 |
| NUDT22 |
| NUDT3 |
| NUDT5 |
| NUDT6 |
| NUDT7 |
| NUDT9 |
| NUF2 |
| NUFIP1 |
| NUFIP2 |
| NUMA1 |
| NUMB |
| NUMBL |
| NUP133 |
| NUP153 |
| NUP155 |
| NUP160 |
| NUP188 |
| NUP205 |
| NUP210 |
| NUP214 |
| NUP35 |
| NUP43 |
| NUP50 |
| NUP54 |
| NUP62 |
| NUP85 |
| NUP88 |
| NUP93 |
| NUP98 |
| NUPR1 |
| NUS1 |
| NUSAP1 |
| NVL |
| NXF1 |
| NXPH1 |
| NXT1 |
| NXT2 |
| NYNRIN |
| OAF |
| OASL |

| |
|---|
| OAT |
| OAZ1 |
| OBSL1 |
| OCEL1 |
| OCIAD1 |
| OCIAD2 |
| OCLN |
| OCRL |
| ODC1 |
| ODF2 |
| ODF2L |
| OFD1 |
| OGFOD1 |
| OGFR |
| OGFRL1 |
| OGG1 |
| OIP5 |
| OIT3 |
| OLA1 |
| OLFM1 |
| OLFM4 |
| OLFML1 |
| OLFML2A |
| OLFML2B |
| OLIG1 |
| OLR1 |
| OMA1 |
| OMD |
| ONECUT1 |
| OPA1 |
| OPA3 |
| OPCML |
| OPHN1 |
| OPRK1 |
| OPTN |
| ORAI1 |
| ORAI3 |
| ORMDL2 |
| ORMDL3 |
| OS9 |
| OSBP |
| OSBP2 |
| OSBPL10 |
| OSBPL11 |
| OSBPL2 |

| |
|---|
| OSBPL3 |
| OSBPL5 |
| OSBPL8 |
| OSBPL9 |
| OSCP1 |
| OSGEP |
| OSGEPL1 |
| OSGIN1 |
| OSM |
| OSR2 |
| OSTC |
| OSTF1 |
| OSTM1 |
| OTOP2 |
| OTUB1 |
| OTUD1 |
| OTUD3 |
| OTUD4 |
| OTUD5 |
| OTUD6B |
| OVCA2 |
| OVGP1 |
| OVOL1 |
| OXA1L |
| OXCT1 |
| OXNAD1 |
| OXR1 |
| OXSM |
| OXSR1 |
| P2RX4 |
| P2RX6 |
| P2RY1 |
| P2RY10 |
| P2RY2 |
| P2RY4 |
| P4HA1 |
| P4HA2 |
| P4HA3 |
| P4HB |
| P4HTM |
| PA2G4 |
| PAAF1 |
| PABPC1 |
| PABPC4 |
| PABPC5 |

| |
|---|
| PABPN1 |
| PACRGL |
| PACSIN2 |
| PACSIN3 |
| PADI1 |
| PADI2 |
| PAF1 |
| PAFAH1B1 |
| PAFAH1B2 |
| PAFAH1B3 |
| PAFAH2 |
| PAG1 |
| PAIP1 |
| PAIP2 |
| PAK1 |
| PAK1IP1 |
| PAK3 |
| PAK4 |
| PALB2 |
| PALLD |
| PALMD |
| PAM |
| PAN2 |
| PAN3 |
| PANK3 |
| PANK4 |
| PANX1 |
| PANX2 |
| PAOX |
| PAPLN |
| PAPOLA |
| PAPOLG |
| PAPSS1 |
| PAPSS2 |
| PAQR3 |
| PAQR4 |
| PAQR5 |
| PAQR6 |
| PAQR8 |
| PARD3 |
| PARD6A |
| PARD6B |
| PARD6G |
| PARG |
| PARK7 |

| |
|---|
| PARL |
| PARN |
| PARP1 |
| PARP11 |
| PARP12 |
| PARP14 |
| PARP16 |
| PARP6 |
| PARP8 |
| PARP9 |
| PARS2 |
| PARVA |
| PARVG |
| PASK |
| PATL1 |
| PATZ1 |
| PAWR |
| PAXIP1 |
| PBK |
| PBLD |
| PBX1 |
| PBX2 |
| PBX3 |
| PBX4 |
| PBXIP1 |
| PC |
| PCBD1 |
| PCBP1 |
| PCBP2 |
| PCBP4 |
| PCCA |
| PCCB |
| PCDH1 |
| PCDH10 |
| PCDH18 |
| PCDH7 |
| PCDH9 |
| PCDHA3 |
| PCDHA4 |
| PCDHB11 |
| PCDHB14 |
| PCDHB15 |
| PCDHB4 |
| PCDHB6 |
| PCDHB7 |

| |
|---|
| PCDHGA5 |
| PCF11 |
| PCGF1 |
| PCGF2 |
| PCGF3 |
| PCGF5 |
| PCGF6 |
| PCID2 |
| PCIF1 |
| PCK1 |
| PCK2 |
| PCM1 |
| PCMT1 |
| PCMTD1 |
| PCMTD2 |
| PCNA |
| PCNP |
| PCOLCE |
| PCOLCE2 |
| PCP4L1 |
| PCSK1N |
| PCSK4 |
| PCSK5 |
| PCSK6 |
| PCSK7 |
| PCTP |
| PCYOX1 |
| PCYOX1L |
| PCYT1A |
| PCYT1B |
| PCYT2 |
| PDAP1 |
| PDCD1 |
| PDCD10 |
| PDCD11 |
| PDCD1LG2 |
| PDCD2 |
| PDCD2L |
| PDCD4 |
| PDCD5 |
| PDCD6 |
| PDCD6IP |
| PDCD7 |
| PDCL |
| PDCL3 |

| |
|---|
| PDE10A |
| PDE11A |
| PDE12 |
| PDE2A |
| PDE4B |
| PDE4C |
| PDE4D |
| PDE4DIP |
| PDE6D |
| PDE7B |
| PDE8A |
| PDE9A |
| PDGFA |
| PDGFB |
| PDGFC |
| PDGFD |
| PDGFRB |
| PDHA1 |
| PDHB |
| PDHX |
| PDIA2 |
| PDIA3 |
| PDIA4 |
| PDIA5 |
| PDIA6 |
| PDIK1L |
| PDK1 |
| PDK3 |
| PDK4 |
| PDLIM1 |
| PDLIM2 |
| PDLIM4 |
| PDLIM5 |
| PDLIM7 |
| PDPK1 |
| PDRG1 |
| PDS5A |
| PDS5B |
| PDSS1 |
| PDXDC1 |
| PDXK |
| PDXP |
| PDZD11 |
| PDZD2 |
| PDZD4 |

| |
|---|
| PDZD8 |
| PDZK1 |
| PDZRN3 |
| PDZRN4 |
| PEBP1 |
| PECAM1 |
| PECR |
| PEG10 |
| PEG3 |
| PELI3 |
| PELO |
| PELP1 |
| PEMT |
| PENK |
| PER1 |
| PER2 |
| PER3 |
| PERP |
| PEX1 |
| PEX10 |
| PEX11A |
| PEX11B |
| PEX11G |
| PEX12 |
| PEX13 |
| PEX14 |
| PEX16 |
| PEX19 |
| PEX2 |
| PEX3 |
| PEX5 |
| PEX6 |
| PEX7 |
| PF4 |
| PFAS |
| PFDN1 |
| PFDN2 |
| PFDN4 |
| PFDN5 |
| PFDN6 |
| PFKFB2 |
| PFKFB3 |
| PFKFB4 |
| PFKL |
| PFKM |

| |
|---|
| PFKP |
| PFN1 |
| PFN2 |
| PGAM1 |
| PGAM2 |
| PGAM5 |
| PGAP1 |
| PGAP2 |
| PGAP3 |
| PGBD1 |
| PGF |
| PGGT1B |
| PGK1 |
| PGLS |
| PGM1 |
| PGM2 |
| PGM2L1 |
| PGM3 |
| PGM5 |
| PGP |
| PGPEP1 |
| PGR |
| PGRMC1 |
| PGRMC2 |
| PGS1 |
| PHACTR1 |
| PHACTR2 |
| PHACTR4 |
| PHAX |
| PHB2 |
| PHC1 |
| PHC2 |
| PHC3 |
| PHF1 |
| PHF11 |
| PHF12 |
| PHF13 |
| PHF14 |
| PHF19 |
| PHF2 |
| PHF20 |
| PHF21A |
| PHF23 |
| PHF3 |
| PHF5A |

| |
|---|
| PHF6 |
| PHF8 |
| PHGDH |
| PHIP |
| PHKA2 |
| PHKB |
| PHKG2 |
| PHLDA1 |
| PHLDA2 |
| PHLDB1 |
| PHLDB2 |
| PHLDB3 |
| PHLPP2 |
| PHOSPHO2 |
| PHYH |
| PI16 |
| PI4K2A |
| PI4K2B |
| PI4KA |
| PI4KB |
| PIAS1 |
| PIAS2 |
| PIAS3 |
| PIAS4 |
| PICALM |
| PICK1 |
| PIF1 |
| PIGA |
| PIGC |
| PIGF |
| PIGH |
| PIGK |
| PIGL |
| PIGM |
| PIGN |
| PIGO |
| PIGP |
| PIGR |
| PIGS |
| PIGT |
| PIGU |
| PIGV |
| PIGY |
| PIH1D1 |
| PIH1D2 |

| |
|---|
| PIK3C2A |
| PIK3C2B |
| PIK3C3 |
| PIK3CA |
| PIK3CB |
| PIK3CD |
| PIK3CG |
| PIK3IP1 |
| PIK3R1 |
| PIK3R2 |
| PIK3R3 |
| PIK3R4 |
| PIK3R5 |
| PIK3R6 |
| PIKFYVE |
| PIM1 |
| PIM2 |
| PIN1 |
| PIN4 |
| PINK1 |
| PINX1 |
| PIP4K2A |
| PIP4K2B |
| PIP5K1A |
| PIP5K1B |
| PIP5K1C |
| PIR |
| PISD |
| PITPNA |
| PITPNM1 |
| PITPNM2 |
| PITRM1 |
| PITX3 |
| PJA1 |
| PKD2 |
| PKD2L2 |
| PKDCC |
| PKHD1 |
| PKIA |
| PKIG |
| PKMYT1 |
| PKN1 |
| PKN2 |
| PKN3 |
| PKNOX1 |

| |
|---|
| PKP1 |
| PKP2 |
| PKP3 |
| PKP4 |
| PLA2G12A |
| PLA2G15 |
| PLA2G6 |
| PLA2G7 |
| PLAA |
| PLAC8 |
| PLAGL1 |
| PLAGL2 |
| PLAU |
| PLAUR |
| PLB1 |
| PLBD1 |
| PLBD2 |
| PLCB2 |
| PLCB3 |
| PLCB4 |
| PLCD1 |
| PLCD3 |
| PLCE1 |
| PLCG1 |
| PLCH2 |
| PLCL2 |
| PLCXD1 |
| PLCXD2 |
| PLD2 |
| PLD3 |
| PLEK |
| PLEK2 |
| PLEKHA1 |
| PLEKHA2 |
| PLEKHA3 |
| PLEKHA4 |
| PLEKHA5 |
| PLEKHA6 |
| PLEKHA7 |
| PLEKHA8 |
| PLEKHB1 |
| PLEKHF1 |
| PLEKHF2 |
| PLEKHG1 |
| PLEKHG2 |

| |
|---|
| PLEKHG3 |
| PLEKHG5 |
| PLEKHG7 |
| PLEKHH1 |
| PLEKHH3 |
| PLEKHJ1 |
| PLEKHM2 |
| PLEKHM3 |
| PLEKHN1 |
| PLEKHO1 |
| PLEKHO2 |
| PLIN1 |
| PLIN2 |
| PLIN3 |
| PLK1 |
| PLK2 |
| PLK3 |
| PLK4 |
| PLOD1 |
| PLOD2 |
| PLOD3 |
| PLP2 |
| PLRG1 |
| PLS3 |
| PLSCR4 |
| PLSCR5 |
| PLTP |
| PLVAP |
| PLXDC1 |
| PLXNA4 |
| PLXNB2 |
| PLXNB3 |
| PLXNC1 |
| PLXND1 |
| PM20D2 |
| PMF1 |
| PML |
| PMM1 |
| PMM2 |
| PMP22 |
| PMPCA |
| PMPCB |
| PMS1 |
| PMVK |
| PNKD |

| |
|---|
| PNKP |
| PNMA1 |
| PNN |
| PNO1 |
| PNPLA2 |
| PNPLA3 |
| PNPLA4 |
| PNPLA6 |
| PNPLA8 |
| PNPO |
| PNPT1 |
| PNRC1 |
| PNRC2 |
| POFUT1 |
| POFUT2 |
| POGK |
| POGZ |
| POLA1 |
| POLA2 |
| POLB |
| POLD1 |
| POLD2 |
| POLD3 |
| POLD4 |
| POLDIP2 |
| POLDIP3 |
| POLE |
| POLE2 |
| POLE3 |
| POLE4 |
| POLG |
| POLG2 |
| POLH |
| POLI |
| POLK |
| POLL |
| POLM |
| POLQ |
| POLR1C |
| POLR1D |
| POLR1E |
| POLR2A |
| POLR2B |
| POLR2C |
| POLR2D |

| |
|---|
| POLR2E |
| POLR2F |
| POLR2G |
| POLR2H |
| POLR2I |
| POLR2J |
| POLR2L |
| POLR3A |
| POLR3B |
| POLR3C |
| POLR3E |
| POLR3F |
| POLR3GL |
| POLR3H |
| POLR3K |
| POLRMT |
| POM121L2 |
| POMC |
| POMGNT1 |
| POMP |
| POMT1 |
| POMT2 |
| PON2 |
| PON3 |
| POP1 |
| POP5 |
| POP7 |
| POR |
| PORCN |
| POSTN |
| POT1 |
| POU2AF1 |
| POU2F1 |
| POU2F3 |
| POU5F1 |
| POU6F1 |
| PPA1 |
| PPA2 |
| PPARA |
| PPARD |
| PPARG |
| PPAT |
| PPCDC |
| PPCS |
| PPDPF |

| |
|---|
| PPFIA1 |
| PPFIA4 |
| PPFIBP1 |
| PPFIBP2 |
| PPHLN1 |
| PPIA |
| PPIB |
| PPIC |
| PPID |
| PPIE |
| PPIF |
| PPIG |
| PPIH |
| PPIL1 |
| PPIL2 |
| PPIL3 |
| PPIL4 |
| PPIL6 |
| PPIP5K1 |
| PPIP5K2 |
| PPL |
| PPM1A |
| PPM1B |
| PPM1D |
| PPM1F |
| PPM1G |
| PPM1K |
| PPM1M |
| PPOX |
| PPP1CA |
| PPP1CC |
| PPP1R10 |
| PPP1R11 |
| PPP1R12A |
| PPP1R12C |
| PPP1R13B |
| PPP1R13L |
| PPP1R14A |
| PPP1R14C |
| PPP1R14D |
| PPP1R15A |
| PPP1R15B |
| PPP1R2 |
| PPP1R3A |
| PPP1R3B |

| |
|---|
| PPP1R3D |
| PPP1R3F |
| PPP1R7 |
| PPP1R8 |
| PPP1R9A |
| PPP2CA |
| PPP2CB |
| PPP2R1A |
| PPP2R1B |
| PPP2R2A |
| PPP2R3A |
| PPP2R3C |
| PPP2R5A |
| PPP2R5B |
| PPP2R5C |
| PPP2R5D |
| PPP2R5E |
| PPP3CA |
| PPP3CB |
| PPP3R1 |
| PPP3R2 |
| PPP4C |
| PPP4R1 |
| PPP4R2 |
| PPP4R4 |
| PPP5C |
| PPP6C |
| PPRC1 |
| PPT1 |
| PPT2 |
| PPTC7 |
| PPWD1 |
| PQBP1 |
| PQLC2 |
| PRAF2 |
| PRAM1 |
| PRAME |
| PRC1 |
| PRCC |
| PRCP |
| PRDM1 |
| PRDM10 |
| PRDM15 |
| PRDM4 |
| PRDM5 |

| |
|---|
| PRDM7 |
| PRDX2 |
| PRDX3 |
| PRDX4 |
| PRDX5 |
| PRDX6 |
| PREB |
| PRELID1 |
| PREP |
| PREPL |
| PREX1 |
| PRICKLE1 |
| PRICKLE3 |
| PRICKLE4 |
| PRIM1 |
| PRIM2 |
| PRKAA1 |
| PRKAA2 |
| PRKAB1 |
| PRKACA |
| PRKACB |
| PRKAG1 |
| PRKAG2 |
| PRKAR1B |
| PRKAR2A |
| PRKCD |
| PRKCE |
| PRKCI |
| PRKCSH |
| PRKCZ |
| PRKD1 |
| PRKD2 |
| PRKD3 |
| PRKDC |
| PRKRA |
| PRKRIP1 |
| PRMT1 |
| PRMT2 |
| PRMT3 |
| PRMT5 |
| PRMT6 |
| PRMT7 |
| PRODH |
| PROM1 |
| PROS1 |

| |
|---|
| PRPF18 |
| PRPF19 |
| PRPF3 |
| PRPF31 |
| PRPF38A |
| PRPF39 |
| PRPF40B |
| PRPF4B |
| PRPF6 |
| PRPF8 |
| PRPS1 |
| PRPS2 |
| PRPSAP1 |
| PRPSAP2 |
| PRR11 |
| PRR12 |
| PRR14 |
| PRR15 |
| PRR15L |
| PRR5 |
| PRR7 |
| PRRC1 |
| PRRG2 |
| PRSS35 |
| PRUNE2 |
| PSAP |
| PSAT1 |
| PSD3 |
| PSD4 |
| PSEN1 |
| PSEN2 |
| PSENEN |
| PSIP1 |
| PSKH1 |
| PSMA1 |
| PSMA2 |
| PSMA3 |
| PSMA5 |
| PSMA6 |
| PSMA7 |
| PSMB1 |
| PSMB10 |
| PSMB11 |
| PSMB2 |
| PSMB3 |

| |
|---|
| PSMB4 |
| PSMB5 |
| PSMB7 |
| PSMB9 |
| PSMC1 |
| PSMC2 |
| PSMC3 |
| PSMC3IP |
| PSMC5 |
| PSMC6 |
| PSMD1 |
| PSMD10 |
| PSMD11 |
| PSMD12 |
| PSMD14 |
| PSMD2 |
| PSMD3 |
| PSMD4 |
| PSMD5 |
| PSMD6 |
| PSMD7 |
| PSMD8 |
| PSMD9 |
| PSME1 |
| PSME2 |
| PSME4 |
| PSMF1 |
| PSMG2 |
| PSMG3 |
| PSMG4 |
| PSPC1 |
| PSPH |
| PSRC1 |
| PSTK |
| PSTPIP1 |
| PTAFR |
| PTAR1 |
| PTBP1 |
| PTCD2 |
| PTCD3 |
| PTCH1 |
| PTCH2 |
| PTCRA |
| PTDSS1 |
| PTEN |

| |
|---|
| PTER |
| PTGDS |
| PTGER1 |
| PTGER2 |
| PTGER3 |
| PTGER4 |
| PTGES2 |
| PTGES3 |
| PTGFR |
| PTGFRN |
| PTGS1 |
| PTGS2 |
| PTH |
| PTH2 |
| PTHLH |
| PTK2 |
| PTK6 |
| PTK7 |
| PTMA |
| PTN |
| PTP4A1 |
| PTP4A2 |
| PTP4A3 |
| PTPDC1 |
| PTPMT1 |
| PTPN1 |
| PTPN11 |
| PTPN12 |
| PTPN18 |
| PTPN2 |
| PTPN21 |
| PTPN23 |
| PTPN3 |
| PTPN6 |
| PTPN9 |
| PTPRA |
| PTPRB |
| PTPRC |
| PTPRD |
| PTPRE |
| PTPRF |
| PTPRH |
| PTPRK |
| PTPRM |
| PTPRN2 |

| |
|---|
| PTPRS |
| PTS |
| PTTG1 |
| PTX3 |
| PUF60 |
| PUM1 |
| PUM2 |
| PURA |
| PURG |
| PUS1 |
| PUS3 |
| PUS7 |
| PUS7L |
| PUSL1 |
| PVR |
| PWP1 |
| PWP2 |
| PWWP2A |
| PWWP2B |
| PXDN |
| PXK |
| PXMP2 |
| PXMP4 |
| PXN |
| PYCARD |
| PYCR1 |
| PYGB |
| PYGL |
| PYGM |
| PYGO1 |
| PYROXD1 |
| QDPR |
| QKI |
| QPCT |
| QPCTL |
| QRFPR |
| QRICH1 |
| QRSL1 |
| QSOX1 |
| QSOX2 |
| QTRT1 |
| R3HCC1 |
| R3HDM2 |
| RAB10 |
| RAB11A |

| |
|---|
| RAB11B |
| RAB11FIP1 |
| RAB11FIP2 |
| RAB11FIP4 |
| RAB11FIP5 |
| RAB12 |
| RAB13 |
| RAB14 |
| RAB15 |
| RAB18 |
| RAB19 |
| RAB1A |
| RAB1B |
| RAB20 |
| RAB21 |
| RAB22A |
| RAB23 |
| RAB24 |
| RAB25 |
| RAB27A |
| RAB27B |
| RAB28 |
| RAB2A |
| RAB2B |
| RAB30 |
| RAB31 |
| RAB32 |
| RAB33B |
| RAB34 |
| RAB35 |
| RAB36 |
| RAB37 |
| RAB38 |
| RAB3B |
| RAB3D |
| RAB3GAP1 |
| RAB3GAP2 |
| RAB3IL1 |
| RAB40B |
| RAB40C |
| RAB4A |
| RAB4B |
| RAB5A |
| RAB5C |
| RAB6A |

| |
|---|
| RAB8A |
| RAB8B |
| RAB9A |
| RAB9B |
| RABEP1 |
| RABEP2 |
| RABEPK |
| RABGAP1L |
| RABGGTA |
| RABIF |
| RABL3 |
| RAC1 |
| RAC2 |
| RAC3 |
| RACGAP1 |
| RAD1 |
| RAD17 |
| RAD18 |
| RAD21 |
| RAD23A |
| RAD23B |
| RAD50 |
| RAD51 |
| RAD51AP1 |
| RAD51C |
| RAD52 |
| RAD54B |
| RAD9A |
| RAE1 |
| RAI1 |
| RAI2 |
| RALA |
| RALB |
| RALBP1 |
| RALGAPA1 |
| RALGAPA2 |
| RALGDS |
| RALGPS1 |
| RALGPS2 |
| RALY |
| RALYL |
| RANBP1 |
| RANBP10 |
| RANBP3 |
| RANBP6 |

| |
|---|
| RANBP9 |
| RANGRF |
| RAP1A |
| RAP1B |
| RAP1GAP |
| RAP2A |
| RAP2B |
| RAP2C |
| RAPGEF1 |
| RAPGEF2 |
| RAPGEF3 |
| RAPGEF4 |
| RAPGEF6 |
| RAPGEFL1 |
| RAPSN |
| RARA |
| RARG |
| RARRES1 |
| RARRES2 |
| RARS2 |
| RASA1 |
| RASA2 |
| RASA3 |
| RASAL2 |
| RASAL3 |
| RASD1 |
| RASEF |
| RASGEF1B |
| RASGEF1C |
| RASGRF1 |
| RASGRP2 |
| RASGRP3 |
| RASGRP4 |
| RASIP1 |
| RASL10B |
| RASL11A |
| RASL11B |
| RASSF1 |
| RASSF2 |
| RASSF3 |
| RASSF5 |
| RASSF6 |
| RASSF7 |
| RASSF8 |
| RASSF9 |

| |
|---|
| RAVER1 |
| RAVER2 |
| RB1 |
| RB1CC1 |
| RBBP4 |
| RBBP5 |
| RBBP6 |
| RBBP7 |
| RBBP8 |
| RBBP9 |
| RBCK1 |
| RBKS |
| RBL2 |
| RBM10 |
| RBM12B |
| RBM14 |
| RBM15 |
| RBM15B |
| RBM17 |
| RBM18 |
| RBM22 |
| RBM25 |
| RBM27 |
| RBM28 |
| RBM3 |
| RBM33 |
| RBM34 |
| RBM41 |
| RBM42 |
| RBM43 |
| RBM45 |
| RBM47 |
| RBM4B |
| RBM5 |
| RBM6 |
| RBM7 |
| RBM8A |
| RBMS1 |
| RBMS3 |
| RBMX |
| RBP2 |
| RBP7 |
| RBPMS2 |
| RBX1 |
| RC3H1 |

| |
|---|
| RCAN3 |
| RCBTB1 |
| RCBTB2 |
| RCC2 |
| RCCD1 |
| RCE1 |
| RCHY1 |
| RCL1 |
| RCN1 |
| RCN2 |
| RCN3 |
| RCOR1 |
| RCOR2 |
| RCOR3 |
| RCSD1 |
| RDH10 |
| RDH11 |
| RDH13 |
| RDH14 |
| RDM1 |
| RECK |
| RECQL |
| RECQL4 |
| RECQL5 |
| REEP1 |
| REEP2 |
| REEP4 |
| REEP5 |
| REEP6 |
| REL |
| RELA |
| RELB |
| RELL1 |
| RELL2 |
| RELN |
| RELT |
| REM1 |
| REM2 |
| REN |
| RENBP |
| REP15 |
| REPIN1 |
| REPS1 |
| REPS2 |
| RER1 |

| |
|---|
| RERE |
| RERG |
| RETSAT |
| REV1 |
| REV3L |
| REXO1 |
| REXO2 |
| RFC2 |
| RFC3 |
| RFC4 |
| RFC5 |
| RFESD |
| RFFL |
| RFK |
| RFNG |
| RFT1 |
| RFTN1 |
| RFTN2 |
| RFWD3 |
| RFX2 |
| RFX7 |
| RFXANK |
| RFXAP |
| RGL1 |
| RGL2 |
| RGMA |
| RGMB |
| RGS1 |
| RGS10 |
| RGS11 |
| RGS12 |
| RGS18 |
| RGS19 |
| RGS2 |
| RGS3 |
| RGS4 |
| RGS5 |
| RGS9BP |
| RHBDD1 |
| RHBDD2 |
| RHBDD3 |
| RHBDF2 |
| RHBDL2 |
| RHEB |
| RHOB |

| |
|---|
| RHOBTB1 |
| RHOBTB3 |
| RHOD |
| RHOF |
| RHOG |
| RHOH |
| RHOQ |
| RHOT1 |
| RHOT2 |
| RHOU |
| RHPN2 |
| RIBC1 |
| RIC3 |
| RIC8A |
| RIC8B |
| RICTOR |
| RIF1 |
| RILPL1 |
| RILPL2 |
| RIMKLA |
| RIMKLB |
| RIMS1 |
| RIMS2 |
| RIN1 |
| RIN3 |
| RING1 |
| RINT1 |
| RIOK1 |
| RIOK2 |
| RIOK3 |
| RIPK1 |
| RIPK2 |
| RIT1 |
| RLF |
| RMI2 |
| RMND5A |
| RMND5B |
| RNASE13 |
| RNASE4 |
| RNASE6 |
| RNASEH1 |
| RNASEH2A |
| RNASEH2B |
| RNASEH2C |
| RNASEK |

| |
|---|
| RNASET2 |
| RND2 |
| RND3 |
| RNF10 |
| RNF103 |
| RNF11 |
| RNF111 |
| RNF112 |
| RNF113A |
| RNF114 |
| RNF115 |
| RNF123 |
| RNF126 |
| RNF128 |
| RNF13 |
| RNF135 |
| RNF139 |
| RNF14 |
| RNF141 |
| RNF145 |
| RNF148 |
| RNF149 |
| RNF166 |
| RNF167 |
| RNF168 |
| RNF169 |
| RNF17 |
| RNF170 |
| RNF180 |
| RNF181 |
| RNF182 |
| RNF185 |
| RNF187 |
| RNF19B |
| RNF2 |
| RNF20 |
| RNF213 |
| RNF214 |
| RNF215 |
| RNF216 |
| RNF219 |
| RNF220 |
| RNF222 |
| RNF24 |
| RNF25 |

| |
|---|
| RNF26 |
| RNF31 |
| RNF32 |
| RNF4 |
| RNF40 |
| RNF41 |
| RNF44 |
| RNF5 |
| RNF6 |
| RNF7 |
| RNFT1 |
| RNFT2 |
| RNGTT |
| RNH1 |
| RNLS |
| RNMT |
| RNPC3 |
| RNPEP |
| RNPEPL1 |
| RNPS1 |
| ROBO1 |
| ROCK1 |
| ROCK2 |
| ROGDI |
| ROM1 |
| ROMO1 |
| RP2 |
| RP9 |
| RPA1 |
| RPA2 |
| RPA3 |
| RPAIN |
| RPAP1 |
| RPAP3 |
| RPE |
| RPF1 |
| RPF2 |
| RPGR |
| RPGRIP1L |
| RPH3AL |
| RPIA |
| RPL10A |
| RPL11 |
| RPL12 |
| RPL13 |

| |
|---|
| RPL13A |
| RPL14 |
| RPL15 |
| RPL18 |
| RPL18A |
| RPL21 |
| RPL22L1 |
| RPL23 |
| RPL23A |
| RPL24 |
| RPL26 |
| RPL27 |
| RPL27A |
| RPL28 |
| RPL29 |
| RPL3 |
| RPL31 |
| RPL32 |
| RPL34 |
| RPL35 |
| RPL36 |
| RPL36A |
| RPL37 |
| RPL38 |
| RPL4 |
| RPL5 |
| RPL6 |
| RPL7 |
| RPL7A |
| RPL7L1 |
| RPL8 |
| RPL9 |
| RPLP1 |
| RPLP2 |
| RPN1 |
| RPN2 |
| RPP21 |
| RPP25 |
| RPP30 |
| RPP38 |
| RPP40 |
| RPRD1A |
| RPRD2 |
| RPRM |
| RPS10 |

| |
|---|
| RPS12 |
| RPS13 |
| RPS14 |
| RPS15 |
| RPS15A |
| RPS16 |
| RPS17 |
| RPS18 |
| RPS19BP1 |
| RPS2 |
| RPS21 |
| RPS24 |
| RPS25 |
| RPS26 |
| RPS27 |
| RPS27A |
| RPS27L |
| RPS3 |
| RPS3A |
| RPS5 |
| RPS6 |
| RPS6KA1 |
| RPS6KA2 |
| RPS6KA3 |
| RPS6KA4 |
| RPS6KA5 |
| RPS6KB1 |
| RPS6KB2 |
| RPS6KC1 |
| RPS7 |
| RPS8 |
| RPS9 |
| RPSA |
| RPTOR |
| RPUSD1 |
| RPUSD2 |
| RPUSD4 |
| RRAD |
| RRAGA |
| RRAGB |
| RRAGC |
| RRAGD |
| RRAS |
| RRAS2 |
| RRM1 |

| |
|---|
| RRM2 |
| RRM2B |
| RRN3 |
| RRP1 |
| RRP12 |
| RRP15 |
| RRP1B |
| RRP7A |
| RRP8 |
| RRP9 |
| RRS1 |
| RSAD1 |
| RSAD2 |
| RSBN1 |
| RSBN1L |
| RSF1 |
| RSL1D1 |
| RSPH3 |
| RSPH4A |
| RSPH6A |
| RSPRY1 |
| RSRC2 |
| RTF1 |
| RTN1 |
| RTN2 |
| RTN3 |
| RTN4 |
| RTN4IP1 |
| RTTN |
| RUFY1 |
| RUFY2 |
| RUFY3 |
| RUNDC1 |
| RUNX1 |
| RUNX3 |
| RUSC1 |
| RUSC2 |
| RUVBL1 |
| RUVBL2 |
| RWDD1 |
| RWDD2A |
| RWDD2B |
| RWDD3 |
| RXFP3 |
| RXFP4 |

| |
|---|
| RXRA |
| RXRB |
| RYBP |
| RYK |
| RYR2 |
| RYR3 |
| S100A10 |
| S100A12 |
| S100A13 |
| S100A16 |
| S100A2 |
| S100A4 |
| S100A6 |
| S100A9 |
| S100PBP |
| S1PR1 |
| S1PR2 |
| S1PR4 |
| SAA4 |
| SAAL1 |
| SACM1L |
| SACS |
| SAE1 |
| SAFB |
| SALL2 |
| SAMD10 |
| SAMD11 |
| SAMD4A |
| SAMD4B |
| SAMD5 |
| SAMD8 |
| SAMD9 |
| SAMHD1 |
| SAMM50 |
| SAP18 |
| SAP30 |
| SAP30BP |
| SAP30L |
| SAR1A |
| SAR1B |
| SARDH |
| SARG |
| SARM1 |
| SARNP |
| SARS2 |

| |
|---|
| SART1 |
| SASH1 |
| SASH3 |
| SASS6 |
| SAT1 |
| SAT2 |
| SAV1 |
| SBDS |
| SBF1 |
| SBF2 |
| SBSN |
| SCAF1 |
| SCAI |
| SCAMP1 |
| SCAMP2 |
| SCAMP3 |
| SCAMP4 |
| SCAND1 |
| SCARA3 |
| SCARB2 |
| SCARF1 |
| SCARF2 |
| SCCPDH |
| SCD5 |
| SCFD1 |
| SCFD2 |
| SCG5 |
| SCGB1A1 |
| SCHIP1 |
| SCIN |
| SCLT1 |
| SCLY |
| SCMH1 |
| SCML1 |
| SCML2 |
| SCN2A |
| SCN9A |
| SCNM1 |
| SCNN1A |
| SCNN1B |
| SCO1 |
| SCO2 |
| SCOC |
| SCP2 |
| SCPEP1 |

| |
|---|
| SCRN1 |
| SCRN2 |
| SCRN3 |
| SCUBE1 |
| SCUBE3 |
| SCYL1 |
| SCYL3 |
| SDAD1 |
| SDC1 |
| SDC2 |
| SDC3 |
| SDC4 |
| SDCBP |
| SDCCAG8 |
| SDF2 |
| SDF2L1 |
| SDF4 |
| SDHAF1 |
| SDHB |
| SDHC |
| SDHD |
| SDK1 |
| SDR39U1 |
| SDR42E1 |
| SEC11A |
| SEC11C |
| SEC13 |
| SEC14L1 |
| SEC14L2 |
| SEC16A |
| SEC16B |
| SEC22A |
| SEC22B |
| SEC22C |
| SEC23A |
| SEC23B |
| SEC23IP |
| SEC24A |
| SEC24B |
| SEC24C |
| SEC24D |
| SEC31A |
| SEC31B |
| SEC61A2 |
| SEC61B |

| |
|---|
| SEC61G |
| SEC62 |
| SEC63 |
| SECISBP2 |
| SECTM1 |
| SEH1L |
| SEL1L |
| SEL1L3 |
| SELE |
| SELENBP1 |
| SELP |
| SELPLG |
| SEMA3B |
| SEMA3F |
| SEMA3G |
| SEMA4B |
| SEMA4C |
| SEMA4D |
| SEMA4F |
| SEMA4G |
| SEMA5A |
| SEMA6A |
| SEMA6B |
| SEMA6C |
| SEMA6D |
| SEMA7A |
| SENP1 |
| SENP2 |
| SENP3 |
| SENP6 |
| SENP7 |
| SEPHS1 |
| SEPHS2 |
| SERAC1 |
| SERBP1 |
| SERGEF |
| SERINC1 |
| SERINC2 |
| SERINC3 |
| SERINC4 |
| SERINC5 |
| SERP1 |
| SERPINA5 |
| SERPINB1 |
| SERPINB2 |

| |
|---|
| SERPINB5 |
| SERPINB6 |
| SERPINB7 |
| SERPINB9 |
| SERPINE1 |
| SERPINE2 |
| SERPINH1 |
| SERPINI1 |
| SERTAD2 |
| SERTAD3 |
| SESN3 |
| SET |
| SETD1A |
| SETD1B |
| SETD2 |
| SETD4 |
| SETD5 |
| SETD6 |
| SETD7 |
| SETMAR |
| SETX |
| SF1 |
| SF3A1 |
| SF3A2 |
| SF3A3 |
| SF3B1 |
| SF3B2 |
| SF3B3 |
| SF3B4 |
| SF3B5 |
| SFI1 |
| SFN |
| SFPQ |
| SFRP1 |
| SFRP4 |
| SFT2D1 |
| SFT2D2 |
| SFT2D3 |
| SFXN1 |
| SFXN2 |
| SFXN4 |
| SFXN5 |
| SGCE |
| SGK1 |
| SGK3 |

| |
|---|
| SGMS2 |
| SGPL1 |
| SGPP1 |
| SGSM1 |
| SGSM3 |
| SGTA |
| SH2B1 |
| SH2B2 |
| SH2B3 |
| SH2D3C |
| SH2D4A |
| SH2D6 |
| SH3BGRL |
| SH3BGRL2 |
| SH3BGRL3 |
| SH3BP2 |
| SH3BP4 |
| SH3BP5 |
| SH3BP5L |
| SH3D19 |
| SH3GL1 |
| SH3GL2 |
| SH3GLB1 |
| SH3GLB2 |
| SH3KBP1 |
| SH3PXD2B |
| SH3RF1 |
| SH3RF2 |
| SH3TC2 |
| SH3YL1 |
| SHARPIN |
| SHB |
| SHC1 |
| SHCBP1 |
| SHD |
| SHF |
| SHISA4 |
| SHISA5 |
| SHISA8 |
| SHKBP1 |
| SHMT1 |
| SHMT2 |
| SHOC2 |
| SHPK |
| SHQ1 |

| |
|---|
| SHROOM1 |
| SHROOM2 |
| SHROOM3 |
| SHROOM4 |
| SIAE |
| SIAH1 |
| SIDT2 |
| SIGLEC1 |
| SIGLEC10 |
| SIGLEC14 |
| SIGLEC8 |
| SIGMAR1 |
| SIK2 |
| SIL1 |
| SIPA1 |
| SIPA1L1 |
| SIPA1L2 |
| SIRPA |
| SIRPB1 |
| SIRT1 |
| SIRT2 |
| SIRT3 |
| SIRT5 |
| SIRT6 |
| SIRT7 |
| SIT1 |
| SIVA1 |
| SIX5 |
| SKA3 |
| SKI |
| SKIL |
| SKP1 |
| SKP2 |
| SLA |
| SLA2 |
| SLAIN1 |
| SLAIN2 |
| SLAMF7 |
| SLAMF8 |
| SLAMF9 |
| SLBP |
| SLC10A3 |
| SLC10A4 |
| SLC10A5 |
| SLC11A1 |

| |
|---|
| SLC11A2 |
| SLC12A2 |
| SLC12A5 |
| SLC12A9 |
| SLC13A2 |
| SLC13A4 |
| SLC15A1 |
| SLC15A3 |
| SLC15A4 |
| SLC16A1 |
| SLC16A11 |
| SLC16A12 |
| SLC16A14 |
| SLC16A3 |
| SLC16A5 |
| SLC16A6 |
| SLC16A7 |
| SLC17A5 |
| SLC19A1 |
| SLC19A2 |
| SLC19A3 |
| SLC1A1 |
| SLC1A4 |
| SLC1A5 |
| SLC1A6 |
| SLC1A7 |
| SLC20A1 |
| SLC20A2 |
| SLC22A15 |
| SLC22A18 |
| SLC22A23 |
| SLC22A5 |
| SLC22A7 |
| SLC23A1 |
| SLC23A2 |
| SLC25A1 |
| SLC25A11 |
| SLC25A12 |
| SLC25A13 |
| SLC25A15 |
| SLC25A16 |
| SLC25A17 |
| SLC25A19 |
| SLC25A22 |
| SLC25A24 |

| |
|---|
| SLC25A25 |
| SLC25A26 |
| SLC25A27 |
| SLC25A28 |
| SLC25A29 |
| SLC25A30 |
| SLC25A35 |
| SLC25A36 |
| SLC25A37 |
| SLC25A38 |
| SLC25A39 |
| SLC25A4 |
| SLC25A42 |
| SLC25A46 |
| SLC25A5 |
| SLC26A1 |
| SLC26A11 |
| SLC26A2 |
| SLC26A3 |
| SLC26A4 |
| SLC26A9 |
| SLC27A1 |
| SLC27A3 |
| SLC27A4 |
| SLC27A5 |
| SLC27A6 |
| SLC28A3 |
| SLC29A2 |
| SLC29A3 |
| SLC29A4 |
| SLC2A1 |
| SLC2A10 |
| SLC2A13 |
| SLC2A2 |
| SLC2A3 |
| SLC2A6 |
| SLC2A8 |
| SLC30A1 |
| SLC30A4 |
| SLC30A5 |
| SLC30A6 |
| SLC30A7 |
| SLC30A9 |
| SLC31A1 |
| SLC31A2 |

| |
|---|
| SLC33A1 |
| SLC34A1 |
| SLC34A2 |
| SLC35A1 |
| SLC35A2 |
| SLC35A4 |
| SLC35B1 |
| SLC35B2 |
| SLC35B3 |
| SLC35B4 |
| SLC35C1 |
| SLC35C2 |
| SLC35D2 |
| SLC35E1 |
| SLC35E4 |
| SLC35F5 |
| SLC36A1 |
| SLC36A4 |
| SLC37A3 |
| SLC37A4 |
| SLC38A10 |
| SLC38A2 |
| SLC38A4 |
| SLC38A5 |
| SLC38A6 |
| SLC38A7 |
| SLC39A1 |
| SLC39A10 |
| SLC39A11 |
| SLC39A14 |
| SLC39A3 |
| SLC39A4 |
| SLC39A5 |
| SLC39A7 |
| SLC39A8 |
| SLC39A9 |
| SLC3A2 |
| SLC40A1 |
| SLC41A1 |
| SLC41A2 |
| SLC41A3 |
| SLC43A1 |
| SLC43A2 |
| SLC44A1 |
| SLC44A2 |

| |
|---|
| SLC44A3 |
| SLC44A4 |
| SLC45A1 |
| SLC45A2 |
| SLC45A4 |
| SLC46A1 |
| SLC47A1 |
| SLC47A2 |
| SLC48A1 |
| SLC4A10 |
| SLC4A1AP |
| SLC4A2 |
| SLC4A3 |
| SLC5A12 |
| SLC5A3 |
| SLC5A6 |
| Slc6a12 |
| SLC6A15 |
| SLC6A6 |
| SLC6A8 |
| SLC7A11 |
| SLC7A2 |
| SLC7A3 |
| SLC7A4 |
| SLC7A5 |
| SLC7A6 |
| SLC7A6OS |
| SLC7A7 |
| SLC7A8 |
| SLC8A1 |
| SLC8A2 |
| SLC9A1 |
| SLC9A3R1 |
| SLC9A3R2 |
| SLC9A6 |
| SLC9A7 |
| SLC9A9 |
| SLCO2B1 |
| SLCO3A1 |
| SLCO4C1 |
| SLFN11 |
| SLIRP |
| SLITRK1 |
| SLITRK2 |
| SLITRK6 |

| |
|---|
| SLK |
| SLMAP |
| SLTM |
| SLU7 |
| SMAD1 |
| SMAD2 |
| SMAD3 |
| SMAD4 |
| SMAD6 |
| SMAD7 |
| SMAGP |
| SMAP |
| SMAP1 |
| SMAP2 |
| SMARCA1 |
| SMARCA4 |
| SMARCA5 |
| SMARCC1 |
| SMARCC2 |
| SMARCD2 |
| SMARCD3 |
| SMARCE1 |
| SMC1A |
| SMC1B |
| SMC2 |
| SMC3 |
| SMC4 |
| SMC5 |
| SMC6 |
| SMCHD1 |
| SMG5 |
| SMG6 |
| SMG7 |
| SMG9 |
| SMN1 |
| SMNDC1 |
| SMO |
| SMOX |
| SMPD1 |
| SMPD2 |
| SMPD4 |
| SMPDL3A |
| SMPDL3B |
| SMS |
| SMTN |

| |
|---|
| SMU1 |
| SMUG1 |
| SMURF1 |
| SMURF2 |
| SMYD2 |
| SMYD3 |
| SMYD4 |
| SMYD5 |
| SNAI1 |
| SNAP23 |
| SNAP29 |
| SNAP47 |
| SNAPC1 |
| SNAPC3 |
| SNAPC4 |
| SNAPC5 |
| SNAPIN |
| SND1 |
| SNF8 |
| SNIP1 |
| SNN |
| SNPH |
| SNRK |
| SNRNP200 |
| SNRNP25 |
| SNRNP27 |
| SNRNP35 |
| SNRNP40 |
| SNRNP48 |
| SNRNP70 |
| SNRPA1 |
| SNRPB |
| SNRPB2 |
| SNRPC |
| SNRPD1 |
| SNRPD3 |
| SNRPE |
| SNRPF |
| SNRPG |
| SNRPN |
| SNTA1 |
| SNTB1 |
| SNTB2 |
| SNTG2 |
| SNUPN |

| |
|---|
| SNW1 |
| SNX1 |
| SNX10 |
| SNX11 |
| SNX13 |
| SNX14 |
| SNX15 |
| SNX16 |
| SNX17 |
| SNX18 |
| SNX19 |
| SNX2 |
| SNX20 |
| SNX21 |
| SNX24 |
| SNX25 |
| SNX27 |
| SNX29 |
| SNX3 |
| SNX30 |
| SNX5 |
| SNX6 |
| SNX7 |
| SNX8 |
| SNX9 |
| SOAT1 |
| SOBP |
| SOCS1 |
| SOCS2 |
| SOCS3 |
| SOCS4 |
| SOCS5 |
| SOCS6 |
| SOD1 |
| SOD2 |
| SOD3 |
| SON |
| SORBS2 |
| SORBS3 |
| SORD |
| SORL1 |
| SOS1 |
| SOS2 |
| SOSTDC1 |
| SOX13 |

| |
|---|
| SOX4 |
| SOX5 |
| SP3 |
| SPA17 |
| SPAG1 |
| SPAG4 |
| SPAG5 |
| SPAG7 |
| SPAG8 |
| SPAG9 |
| SPARC |
| SPAST |
| SPATA13 |
| SPATA18 |
| SPATA2 |
| SPATA20 |
| SPATA24 |
| SPATA2L |
| SPATA4 |
| SPATA5L1 |
| SPATA6 |
| SPATA7 |
| SPC24 |
| SPC25 |
| SPCS1 |
| SPCS2 |
| SPCS3 |
| SPDEF |
| SPDYA |
| SPEN |
| SPG11 |
| SPG7 |
| SPHK1 |
| SPHK2 |
| SPHKAP |
| SPI1 |
| SPIN1 |
| SPIN4 |
| SPINT1 |
| SPINT2 |
| SPIRE1 |
| SPNS1 |
| SPOCK1 |
| SPOCK2 |
| SPON1 |

| |
|---|
| SPON2 |
| SPOP |
| SPOPL |
| SPPL2B |
| SPR |
| SPRED2 |
| SPRY2 |
| SPRY3 |
| SPRY4 |
| SPSB1 |
| SPSB3 |
| SPSB4 |
| SPTAN1 |
| SPTBN1 |
| SPTLC1 |
| SPTLC2 |
| SPTY2D1 |
| SQLE |
| SQSTM1 |
| SRA1 |
| SRBD1 |
| SRC |
| SRD5A3 |
| SREBF1 |
| SREBF2 |
| SRF |
| SRFBP1 |
| SRGAP1 |
| SRGAP2 |
| SRGN |
| SRI |
| SRM |
| SRP14 |
| SRP19 |
| SRP54 |
| SRP68 |
| SRP72 |
| SRP9 |
| SRPK1 |
| SRPK2 |
| SRPK3 |
| SRR |
| SRRD |
| SRRM1 |
| SRRM2 |

| |
|---|
| SRRT |
| SS18 |
| SS18L1 |
| SSB |
| SSBP1 |
| SSBP2 |
| SSBP3 |
| SSBP4 |
| SSH1 |
| SSH2 |
| SSH3 |
| SSNA1 |
| SSPN |
| SSR1 |
| SSR2 |
| SSR3 |
| SSR4 |
| SSRP1 |
| SSU72 |
| SSX2IP |
| ST13 |
| ST14 |
| ST3GAL2 |
| ST3GAL3 |
| ST3GAL4 |
| ST3GAL5 |
| ST3GAL6 |
| ST6GAL1 |
| ST6GAL2 |
| ST6GALNAC1 |
| ST6GALNAC2 |
| ST6GALNAC6 |
| ST7 |
| ST8SIA4 |
| ST8SIA5 |
| STAB1 |
| STAC |
| STAG1 |
| STAM |
| STAM2 |
| STAMBP |
| STAP2 |
| STAR |
| STARD10 |
| STARD13 |

| |
|---|
| STARD3 |
| STARD3NL |
| STARD4 |
| STARD6 |
| STARD7 |
| STARD8 |
| STARD9 |
| STAT1 |
| STAT2 |
| STAT3 |
| STAT5A |
| STAU1 |
| STAU2 |
| STBD1 |
| STC1 |
| STEAP1 |
| STEAP3 |
| STIL |
| STIM1 |
| STIM2 |
| STIP1 |
| STK10 |
| STK11 |
| STK11IP |
| STK16 |
| STK17A |
| STK17B |
| STK19 |
| STK24 |
| STK25 |
| STK3 |
| STK32A |
| STK33 |
| STK35 |
| STK36 |
| STK38 |
| STK38L |
| STK39 |
| STMN1 |
| STOM |
| STOML1 |
| STOML2 |
| STON1 |
| STON2 |
| STOX1 |

| |
|---|
| STRA6 |
| STRADB |
| STRBP |
| STRN |
| STRN3 |
| STRN4 |
| STS |
| STT3A |
| STUB1 |
| STX10 |
| STX11 |
| STX12 |
| STX16 |
| STX17 |
| STX18 |
| STX19 |
| STX1A |
| STX1B |
| STX2 |
| STX4 |
| STX5 |
| STX6 |
| STX7 |
| STX8 |
| STXBP3 |
| STXBP4 |
| STXBP5 |
| STXBP5L |
| STYX |
| SUB1 |
| SUCLA2 |
| SUCLG1 |
| SUCLG2 |
| SUFU |
| SUGT1 |
| SULT1A1 |
| SULT1B1 |
| SUMF1 |
| SUMF2 |
| SUMO1 |
| SUMO3 |
| SUN2 |
| SUOX |
| SUPT16H |
| SUPT3H |

| |
|---|
| SUPT4H1 |
| SUPT5H |
| SUPT6H |
| SUPT7L |
| SUPV3L1 |
| SURF1 |
| SURF2 |
| SURF4 |
| SUSD1 |
| SUSD3 |
| SUV39H1 |
| SUV39H2 |
| SUZ12 |
| SV2C |
| SVIL |
| SWAP70 |
| SYAP1 |
| SYCE2 |
| SYDE1 |
| SYDE2 |
| SYF2 |
| SYK |
| SYN1 |
| SYNC |
| SYNCRIP |
| SYNE1 |
| SYNE2 |
| SYNGAP1 |
| SYNGR1 |
| SYNGR3 |
| SYNJ1 |
| SYNJ2 |
| SYNJ2BP |
| SYNM |
| SYNPO2 |
| SYNRG |
| SYP |
| SYPL1 |
| SYS1 |
| SYT1 |
| SYT10 |
| SYT11 |
| SYT16 |
| SYT17 |
| SYT7 |

| |
|---|
| SYT9 |
| SYTL1 |
| SYTL2 |
| SYTL3 |
| SYTL4 |
| SYTL5 |
| SYVN1 |
| TAC1 |
| TACC2 |
| TACC3 |
| TACR3 |
| TADA1 |
| TADA2A |
| TADA2B |
| TADA3 |
| TAF10 |
| TAF11 |
| TAF12 |
| TAF13 |
| TAF15 |
| TAF1B |
| TAF1C |
| TAF1D |
| TAF4 |
| TAF4B |
| TAF5 |
| TAF5L |
| TAF6 |
| TAF6L |
| TAF7 |
| TAF8 |
| TAF9B |
| TAGAP |
| TAGLN2 |
| TAL2 |
| TALDO1 |
| TAOK1 |
| TAOK2 |
| TAP1 |
| TAP2 |
| TAPBPL |
| TAPT1 |
| TARBP1 |
| TARBP2 |
| TARDBP |

| |
|---|
| TARS2 |
| TAS1R2 |
| TASP1 |
| TATDN1 |
| TATDN2 |
| TATDN3 |
| TAX1BP1 |
| TAX1BP3 |
| TBC1D1 |
| TBC1D10A |
| TBC1D10B |
| TBC1D10C |
| TBC1D12 |
| TBC1D13 |
| TBC1D14 |
| TBC1D15 |
| TBC1D16 |
| TBC1D17 |
| TBC1D19 |
| TBC1D2 |
| TBC1D20 |
| TBC1D22A |
| TBC1D22B |
| TBC1D24 |
| TBC1D2B |
| TBC1D30 |
| TBC1D4 |
| TBC1D7 |
| TBC1D8 |
| TBC1D8B |
| TBC1D9 |
| TBC1D9B |
| TBCA |
| TBCB |
| TBCC |
| TBCD |
| TBK1 |
| TBKBP1 |
| TBL1X |
| TBL1XR1 |
| TBL2 |
| TBL3 |
| TBP |
| TBPL1 |
| TBRG1 |

| |
|---|
| TBRG4 |
| TBXAS1 |
| TCAP |
| TCEA1 |
| TCEA2 |
| TCEA3 |
| TCEAL1 |
| TCEAL4 |
| TCEAL8 |
| TCERG1 |
| TCF12 |
| TCF20 |
| TCF23 |
| TCF25 |
| TCF7 |
| TCF7L1 |
| TCF7L2 |
| TCHP |
| TCN1 |
| TCN2 |
| TCOF1 |
| TCP1 |
| TCP11 |
| TCP11L2 |
| TCTA |
| TCTN1 |
| TDG |
| TDO2 |
| TDP1 |
| TDRD12 |
| TDRD3 |
| TDRD7 |
| TEAD1 |
| TEAD3 |
| TEC |
| TECPR1 |
| TECPR2 |
| TECR |
| TEK |
| TELO2 |
| TERF1 |
| TERF2 |
| TERF2IP |
| TERT |
| TES |

| |
|---|
| TESK1 |
| TESK2 |
| TET2 |
| TET3 |
| TEX14 |
| TEX2 |
| TEX261 |
| TEX264 |
| TF |
| TFAM |
| TFAP4 |
| TFB1M |
| TFB2M |
| TFCP2 |
| TFDP1 |
| TFE3 |
| TFEB |
| TFG |
| TFIP11 |
| TFPI2 |
| TFPT |
| TFRC |
| TGDS |
| TGFB1 |
| TGFB1I1 |
| TGFBR2 |
| TGFBRAP1 |
| TGIF1 |
| TGIF2 |
| TGM2 |
| TGM3 |
| TGS1 |
| THAP1 |
| THAP11 |
| THAP3 |
| THAP4 |
| THAP5 |
| THAP6 |
| THAP9 |
| THBS1 |
| THBS3 |
| THBS4 |
| THG1L |
| THNSL1 |
| THNSL2 |

| |
|---|
| THOC1 |
| THOC2 |
| THOC3 |
| THOC5 |
| THOC6 |
| THOC7 |
| THOP1 |
| THRA |
| THRAP3 |
| THSD4 |
| THSD7A |
| THTPA |
| THUMPD1 |
| THUMPD2 |
| THUMPD3 |
| THYN1 |
| TIA1 |
| TIAL1 |
| TIAM1 |
| TICAM2 |
| TIFA |
| TIGAR |
| TIMELESS |
| TIMM10 |
| TIMM13 |
| TIMM17A |
| TIMM17B |
| TIMM23 |
| TIMM44 |
| TIMM50 |
| TIMM8B |
| TIMM9 |
| TIMP1 |
| TIMP2 |
| TIMP4 |
| TINAGL1 |
| TINF2 |
| TIPARP |
| TIPIN |
| TIPRL |
| TJAP1 |
| TJP1 |
| TJP2 |
| TJP3 |
| TK1 |

| |
|---|
| TK2 |
| TKT |
| TKTL2 |
| TLCD1 |
| TLE1 |
| TLE2 |
| TLE3 |
| TLE4 |
| TLK1 |
| TLK2 |
| TLN1 |
| TLR1 |
| TLR10 |
| TLR2 |
| TLR3 |
| TLR4 |
| TLR5 |
| TLR6 |
| TLR7 |
| TLR8 |
| TM2D1 |
| TM2D2 |
| TM2D3 |
| TM4SF1 |
| TM4SF20 |
| TM6SF2 |
| TM7SF2 |
| TM7SF3 |
| TM9SF1 |
| TM9SF2 |
| TM9SF4 |
| TMBIM1 |
| TMBIM4 |
| TMBIM6 |
| TMC4 |
| TMC5 |
| TMC6 |
| TMC8 |
| TMCC1 |
| TMCC2 |
| TMCC3 |
| TMCO1 |
| TMCO6 |
| TMED1 |
| TMED10 |

| |
|---|
| TMED2 |
| TMED3 |
| TMED4 |
| TMED5 |
| TMED7 |
| TMED9 |
| TMEM101 |
| TMEM104 |
| TMEM106A |
| TMEM106C |
| TMEM108 |
| TMEM109 |
| TMEM11 |
| TMEM115 |
| TMEM117 |
| TMEM120A |
| TMEM120B |
| TMEM125 |
| TMEM126B |
| TMEM127 |
| TMEM128 |
| TMEM129 |
| TMEM131 |
| TMEM132A |
| TMEM134 |
| TMEM135 |
| TMEM138 |
| TMEM139 |
| TMEM140 |
| TMEM141 |
| TMEM143 |
| TMEM144 |
| TMEM147 |
| TMEM14A |
| TMEM14C |
| TMEM150A |
| TMEM150B |
| TMEM151B |
| TMEM160 |
| TMEM161A |
| TMEM161B |
| TMEM164 |
| TMEM165 |
| TMEM167B |
| TMEM17 |

| |
|---|
| TMEM170A |
| TMEM170B |
| TMEM171 |
| TMEM174 |
| TMEM175 |
| TMEM177 |
| TMEM179B |
| TMEM18 |
| TMEM183A |
| TMEM184A |
| TMEM184C |
| TMEM185A |
| TMEM186 |
| TMEM187 |
| TMEM19 |
| TMEM192 |
| TMEM198 |
| TMEM199 |
| TMEM201 |
| TMEM208 |
| TMEM209 |
| TMEM219 |
| TMEM220 |
| TMEM222 |
| TMEM223 |
| TMEM229B |
| TMEM231 |
| TMEM25 |
| TMEM26 |
| TMEM30A |
| TMEM30B |
| TMEM33 |
| TMEM37 |
| TMEM38A |
| TMEM38B |
| TMEM39A |
| TMEM39B |
| TMEM41A |
| TMEM41B |
| TMEM43 |
| TMEM45A |
| TMEM45B |
| TMEM47 |
| TMEM50A |
| TMEM50B |

| |
|---|
| TMEM51 |
| TMEM52 |
| TMEM53 |
| TMEM54 |
| TMEM59 |
| TMEM59L |
| TMEM60 |
| TMEM61 |
| TMEM62 |
| TMEM64 |
| TMEM65 |
| TMEM67 |
| TMEM68 |
| TMEM70 |
| TMEM81 |
| TMEM86A |
| TMEM87A |
| TMEM87B |
| TMEM88 |
| TMEM89 |
| TMEM8B |
| TMEM9 |
| TMEM92 |
| TMEM97 |
| TMEM98 |
| TMEM9B |
| TMF1 |
| TMLHE |
| TMOD3 |
| TMPO |
| TMPPE |
| TMPRSS2 |
| TMPRSS9 |
| TMSB4X |
| TMTC1 |
| TMTC3 |
| TMTC4 |
| TMUB1 |
| TMUB2 |
| TMX1 |
| TMX2 |
| TMX4 |
| TNC |
| TNF |
| TNFAIP1 |

| |
|---|
| TNFAIP2 |
| TNFAIP3 |
| TNFAIP8 |
| TNFAIP8L2 |
| TNFRSF10A |
| TNFRSF11A |
| TNFRSF12A |
| TNFRSF1A |
| TNFRSF1B |
| TNFRSF21 |
| TNFRSF6B |
| TNFRSF8 |
| TNFSF9 |
| TNIK |
| TNIP2 |
| TNK1 |
| TNKS |
| TNKS1BP1 |
| TNNC1 |
| TNNT1 |
| TNPO2 |
| TNPO3 |
| TNRC18 |
| TNRC6A |
| TNRC6B |
| TNS3 |
| TNXB |
| TOB2 |
| TOE1 |
| TOLLIP |
| TOM1 |
| TOM1L1 |
| TOM1L2 |
| TOMM20 |
| TOMM22 |
| TOMM34 |
| TOMM40 |
| TOMM40L |
| TOMM6 |
| TOMM7 |
| TOP1 |
| TOP2A |
| TOP2B |
| TOP3B |
| TOPBP1 |

| |
|---|
| TOPORS |
| TOR1A |
| TOR1AIP1 |
| TOR1AIP2 |
| TOR1B |
| TOR2A |
| TOR3A |
| TOX2 |
| TOX4 |
| TP53 |
| TP53BP1 |
| TP53BP2 |
| TP53I13 |
| TP53I3 |
| TP53INP1 |
| TP53RK |
| TP53TG5 |
| TPBG |
| TPCN2 |
| TPD52 |
| TPD52L2 |
| TPK1 |
| TPM1 |
| TPM2 |
| TPM3 |
| TPM4 |
| TPMT |
| TPP1 |
| TPP2 |
| TPPP |
| TPR |
| TPRA1 |
| TPST1 |
| TPST2 |
| TPT1 |
| TPX2 |
| TRA2A |
| TRA2B |
| TRADD |
| TRAF1 |
| TRAF2 |
| TRAF3 |
| TRAF3IP2 |
| TRAF3IP3 |
| TRAF4 |

| |
|---|
| TRAF5 |
| TRAF6 |
| TRAF7 |
| TRAFD1 |
| TRAIP |
| TRAK1 |
| TRAK2 |
| TRAM1L1 |
| TRAM2 |
| TRANK1 |
| TRAP1 |
| TRAPPC1 |
| TRAPPC10 |
| TRAPPC3 |
| TRAPPC4 |
| TRAPPC5 |
| TRAPPC6B |
| TRAPPC9 |
| TRAT1 |
| TRDMT1 |
| TREH |
| TREM1 |
| TREM2 |
| TREX1 |
| TRIAP1 |
| TRIB1 |
| TRIB2 |
| TRIB3 |
| TRIM11 |
| TRIM14 |
| TRIM15 |
| TRIM2 |
| TRIM21 |
| TRIM23 |
| TRIM24 |
| TRIM25 |
| TRIM26 |
| TRIM27 |
| TRIM28 |
| TRIM3 |
| TRIM31 |
| TRIM32 |
| TRIM33 |
| TRIM35 |
| TRIM36 |

| |
|---|
| TRIM38 |
| TRIM41 |
| TRIM45 |
| TRIM47 |
| TRIM50 |
| TRIM56 |
| TRIM59 |
| TRIM6 |
| TRIM62 |
| TRIM65 |
| TRIM66 |
| TRIM68 |
| TRIM7 |
| TRIM8 |
| TRIM9 |
| TRIML1 |
| TRIO |
| TRIP10 |
| TRIP11 |
| TRIP12 |
| TRIP13 |
| TRIP4 |
| TRIT1 |
| TRMT1 |
| TRMT112 |
| TRMT12 |
| TRMT5 |
| TRMT6 |
| TRMT61A |
| TRMT61B |
| TRNAU1AP |
| TRNT1 |
| TROAP |
| TRPC1 |
| TRPC4 |
| TRPC4AP |
| TRPM2 |
| TRPS1 |
| TRPT1 |
| TRPV1 |
| TRPV2 |
| TRRAP |
| TRUB2 |
| TSC2 |
| TSC22D1 |

| |
|---|
| TSC22D3 |
| TSC22D4 |
| TSEN15 |
| TSFM |
| TSG101 |
| TSGA10 |
| TSHR |
| TSHZ1 |
| TSHZ3 |
| TSKS |
| TSKU |
| TSLP |
| TSN |
| TSNARE1 |
| TSNAX |
| TSPAN1 |
| TSPAN14 |
| TSPAN15 |
| TSPAN17 |
| TSPAN3 |
| TSPAN31 |
| TSPAN5 |
| TSPAN6 |
| TSPAN7 |
| TSPAN8 |
| TSPO |
| TSPYL1 |
| TSPYL2 |
| TSPYL4 |
| TSPYL5 |
| TSR1 |
| TSR2 |
| TSSC4 |
| TSSK1B |
| TSSK3 |
| TST |
| TSTD1 |
| TSTD2 |
| TTBK2 |
| TTC1 |
| TTC12 |
| TTC13 |
| TTC14 |
| TTC17 |
| TTC19 |

| |
|---|
| TTC21B |
| TTC22 |
| TTC23 |
| TTC26 |
| TTC27 |
| TTC28 |
| TTC30A |
| TTC32 |
| TTC33 |
| TTC38 |
| TTC39A |
| TTC39B |
| TTC39C |
| TTC4 |
| TTC5 |
| TTC7A |
| TTC7B |
| TTC8 |
| TTC9C |
| TTF1 |
| TTL |
| TTLL1 |
| TTLL11 |
| TTLL12 |
| TTLL3 |
| TTLL5 |
| TTLL7 |
| TTN |
| TTPAL |
| TTYH2 |
| TUB |
| TUBA1B |
| TUBA1C |
| TUBA4A |
| TUBA8 |
| TUBB2B |
| TUBB3 |
| TUBD1 |
| TUBE1 |
| TUBG1 |
| TUBG2 |
| TUBGCP2 |
| TUBGCP3 |
| TUBGCP4 |
| TUBGCP5 |

| |
|---|
| TUBGCP6 |
| TUFM |
| TULP3 |
| TUSC1 |
| TUSC3 |
| TUT1 |
| TWF1 |
| TWF2 |
| TWIST1 |
| TWSG1 |
| TXLNA |
| TXN |
| TXN2 |
| TXNDC11 |
| TXNDC15 |
| TXNDC16 |
| TXNDC17 |
| TXNDC5 |
| TXNDC9 |
| TXNIP |
| TXNL1 |
| TXNL4A |
| TXNL4B |
| TXNRD1 |
| TXNRD3 |
| TYMS |
| TYRO3 |
| TYW1 |
| TYW3 |
| U2AF2 |
| UACA |
| UAP1L1 |
| UBA1 |
| UBA2 |
| UBA3 |
| UBA5 |
| UBA52 |
| UBA7 |
| UBAC2 |
| UBAP2 |
| UBAP2L |
| UBASH3B |
| UBC |
| UBD |
| UBE2A |

| |
|---|
| UBE2B |
| UBE2C |
| UBE2D1 |
| UBE2D2 |
| UBE2D4 |
| UBE2E1 |
| UBE2E2 |
| UBE2E3 |
| UBE2F |
| UBE2G2 |
| UBE2H |
| UBE2J1 |
| UBE2J2 |
| UBE2K |
| UBE2L3 |
| UBE2L6 |
| UBE2M |
| UBE2N |
| UBE2O |
| UBE2Q1 |
| UBE2Q2 |
| UBE2R2 |
| UBE2S |
| UBE2T |
| UBE2V1 |
| UBE2W |
| UBE2Z |
| UBE3B |
| UBE3C |
| UBE4A |
| UBE4B |
| UBFD1 |
| UBIAD1 |
| UBL3 |
| UBL4A |
| UBL5 |
| UBL7 |
| UBLCP1 |
| UBN2 |
| UBQLN2 |
| UBQLN4 |
| UBR1 |
| UBR2 |
| UBR3 |
| UBR4 |

| |
|---|
| UBR5 |
| UBR7 |
| UBTD1 |
| UBTD2 |
| UBTF |
| UBXN1 |
| UBXN2A |
| UBXN2B |
| UBXN4 |
| UBXN6 |
| UBXN7 |
| UBXN8 |
| UCHL1 |
| UCHL3 |
| UCHL5 |
| UCK1 |
| UCK2 |
| UCKL1 |
| UCN2 |
| UCP1 |
| UCP2 |
| UEVLD |
| UFM1 |
| UFSP1 |
| UGCG |
| UGDH |
| UGGT2 |
| UGT1A1 |
| UHRF1 |
| UHRF1BP1L |
| UHRF2 |
| UIMC1 |
| ULK1 |
| ULK2 |
| ULK3 |
| UMPS |
| UNC119 |
| UNC119B |
| UNC13B |
| UNC13D |
| UNC45A |
| UNC45B |
| UNC50 |
| UNC5B |
| UNC5C |

| |
|---|
| UNC5CL |
| UNC5D |
| UNC93B1 |
| UNG |
| UNK |
| UPB1 |
| UPF1 |
| UPF2 |
| UPF3A |
| UPF3B |
| UPK1B |
| UPK3A |
| UPP1 |
| UPRT |
| UQCR10 |
| UQCRC1 |
| UQCRC2 |
| UQCRH |
| UQCRQ |
| URB1 |
| URB2 |
| URGCP |
| URM1 |
| UROD |
| UROS |
| USE1 |
| USF1 |
| USF2 |
| USH1C |
| USO1 |
| USP1 |
| USP10 |
| USP11 |
| USP14 |
| USP16 |
| USP18 |
| USP19 |
| USP2 |
| USP20 |
| USP21 |
| USP22 |
| USP24 |
| USP25 |
| USP26 |
| USP27X |

| |
|---|
| USP28 |
| USP30 |
| USP33 |
| USP34 |
| USP37 |
| USP38 |
| USP39 |
| USP4 |
| USP40 |
| USP42 |
| USP44 |
| USP45 |
| USP46 |
| USP48 |
| USP49 |
| USP5 |
| USP54 |
| USP7 |
| USP9X |
| USPL1 |
| UST |
| UTP14A |
| UTP15 |
| UTP18 |
| UTP23 |
| UTP3 |
| UTP6 |
| UVRAG |
| UXS1 |
| UXT |
| VAC14 |
| VAMP3 |
| VAMP4 |
| VAMP5 |
| VAMP7 |
| VAMP8 |
| VANGL1 |
| VANGL2 |
| VAPA |
| VAPB |
| VASH1 |
| VASN |
| VASP |
| VAT1 |
| VAV1 |

| |
|---|
| VBP1 |
| VCAM1 |
| VCAN |
| VDAC1 |
| VDAC2 |
| VDR |
| VEGFA |
| VEGFB |
| VEGFC |
| VEZF1 |
| VEZT |
| VGF |
| VGLL4 |
| VHL |
| VIL1 |
| VILL |
| VIM |
| VKORC1 |
| VKORC1L1 |
| VMA21 |
| VPS11 |
| VPS13A |
| VPS13B |
| VPS13C |
| VPS13D |
| VPS16 |
| VPS18 |
| VPS25 |
| VPS26B |
| VPS29 |
| VPS33A |
| VPS33B |
| VPS35 |
| VPS36 |
| VPS37A |
| VPS37D |
| VPS39 |
| VPS41 |
| VPS45 |
| VPS4A |
| VPS4B |
| VPS52 |
| VPS53 |
| VPS54 |
| VPS72 |

| |
|---|
| VPS8 |
| VRK1 |
| VSIG10 |
| VSIG10L |
| VSX2 |
| VTA1 |
| VTI1A |
| VTI1B |
| VWA1 |
| VWA5A |
| VWA5B2 |
| VWF |
| WAC |
| WARS2 |
| WASF1 |
| WASF2 |
| WASF3 |
| WASL |
| WBP1 |
| WBP11 |
| WBP2 |
| WDFY1 |
| WDFY2 |
| WDFY4 |
| WDHD1 |
| WDR1 |
| WDR11 |
| WDR12 |
| WDR13 |
| WDR18 |
| WDR20 |
| WDR24 |
| WDR25 |
| WDR26 |
| WDR27 |
| WDR3 |
| WDR36 |
| WDR37 |
| WDR4 |
| WDR41 |
| WDR44 |
| WDR45 |
| WDR46 |
| WDR48 |
| WDR5 |

| |
|---|
| WDR53 |
| WDR54 |
| WDR55 |
| WDR59 |
| WDR6 |
| WDR61 |
| WDR62 |
| WDR7 |
| WDR70 |
| WDR72 |
| WDR74 |
| WDR75 |
| WDR76 |
| WDR77 |
| WDR81 |
| WDR83 |
| WDR86 |
| WDR91 |
| WDTC1 |
| WEE1 |
| WFDC8 |
| WFIKKN1 |
| WHAMM |
| WIF1 |
| WIPF3 |
| WIPI1 |
| WIZ |
| WNK1 |
| WNK2 |
| WNT10B |
| WNT5A |
| WNT6 |
| WNT9A |
| WRNIP1 |
| WSB2 |
| WT1 |
| WTAP |
| WWC1 |
| WWC2 |
| WWOX |
| WWP1 |
| WWP2 |
| WWTR1 |
| XAB2 |
| XAF1 |

| |
|---|
| XDH |
| XIAP |
| XK |
| XKR4 |
| XKR8 |
| XPA |
| XPC |
| XPNPEP1 |
| XPNPEP3 |
| XPO1 |
| XPO4 |
| XPO5 |
| XPO6 |
| XPO7 |
| XPOT |
| XPR1 |
| XRCC1 |
| XRCC3 |
| XRCC4 |
| XRCC5 |
| XRCC6 |
| XRN1 |
| XRN2 |
| XYLT2 |
| YAF2 |
| YARS2 |
| YBX1 |
| YBX2 |
| YEATS2 |
| YEATS4 |
| YES1 |
| YIF1A |
| YIPF1 |
| YIPF2 |
| YIPF3 |
| YIPF4 |
| YIPF5 |
| YIPF6 |
| YKT6 |
| YLPM1 |
| YME1L1 |
| YPEL1 |
| YPEL3 |
| YRDC |
| YTHDC1 |

| |
|---|
| YTHDC2 |
| YTHDF1 |
| YTHDF2 |
| YTHDF3 |
| YWHAB |
| YWHAE |
| YWHAG |
| YWHAH |
| YWHAQ |
| YWHAZ |
| YY1 |
| ZADH2 |
| ZAP70 |
| ZAR1 |
| ZAR1L |
| ZBED1 |
| ZBED2 |
| ZBED3 |
| ZBED4 |
| ZBED5 |
| ZBP1 |
| ZBTB1 |
| ZBTB11 |
| ZBTB16 |
| ZBTB2 |
| ZBTB20 |
| ZBTB22 |
| ZBTB24 |
| ZBTB25 |
| ZBTB33 |
| ZBTB34 |
| ZBTB37 |
| ZBTB38 |
| ZBTB39 |
| ZBTB4 |
| ZBTB40 |
| ZBTB43 |
| ZBTB44 |
| ZBTB45 |
| ZBTB48 |
| ZBTB49 |
| ZBTB5 |
| ZBTB6 |
| ZBTB7A |
| ZBTB7B |

| |
|---|
| ZBTB7C |
| ZBTB8A |
| ZBTB8OS |
| ZBTB9 |
| ZC3H12A |
| ZC3H12C |
| ZC3H13 |
| ZC3H14 |
| ZC3H15 |
| ZC3H18 |
| ZC3H3 |
| ZC3H4 |
| ZC3H6 |
| ZC3H7A |
| ZC3H7B |
| ZC3H8 |
| ZC3HAV1 |
| ZC3HC1 |
| ZC4H2 |
| ZCCHC10 |
| ZCCHC14 |
| ZCCHC17 |
| ZCCHC3 |
| ZCCHC4 |
| ZCCHC7 |
| ZCCHC8 |
| ZCRB1 |
| ZCWPW1 |
| ZCWPW2 |
| ZDHHC1 |
| ZDHHC13 |
| ZDHHC14 |
| ZDHHC15 |
| ZDHHC16 |
| ZDHHC17 |
| ZDHHC2 |
| ZDHHC20 |
| ZDHHC23 |
| ZDHHC24 |
| ZDHHC4 |
| ZDHHC5 |
| ZDHHC7 |
| ZDHHC9 |
| ZEB1 |
| ZEB2 |

| |
|---|
| ZER1 |
| ZFAND1 |
| ZFAND2A |
| ZFAND2B |
| ZFAND3 |
| ZFAND5 |
| ZFAND6 |
| ZFAT |
| ZFHX2 |
| ZFP1 |
| ZFP2 |
| ZFP3 |
| ZFP36L1 |
| ZFP36L2 |
| ZFP64 |
| ZFP91 |
| ZFPL1 |
| ZFPM1 |
| ZFR |
| ZFR2 |
| ZFX |
| ZFYVE1 |
| ZFYVE16 |
| ZFYVE19 |
| ZFYVE21 |
| ZFYVE26 |
| ZFYVE27 |
| ZFYVE9 |
| ZG16 |
| ZGPAT |
| ZHX2 |
| ZHX3 |
| ZKSCAN1 |
| ZKSCAN2 |
| ZKSCAN4 |
| ZKSCAN5 |
| ZMAT1 |
| ZMAT2 |
| ZMAT3 |
| ZMAT5 |
| ZMIZ1 |
| ZMIZ2 |
| ZMPSTE24 |
| ZMYM1 |
| ZMYM2 |

| |
|---|
| ZMYM3 |
| ZMYM4 |
| ZMYM5 |
| ZMYND10 |
| ZMYND11 |
| ZMYND12 |
| ZMYND8 |
| ZNF12 |
| ZNF131 |
| ZNF133 |
| ZNF134 |
| ZNF135 |
| ZNF140 |
| ZNF142 |
| ZNF143 |
| ZNF148 |
| ZNF165 |
| ZNF169 |
| ZNF174 |
| ZNF175 |
| ZNF180 |
| ZNF182 |
| ZNF184 |
| ZNF19 |
| ZNF197 |
| ZNF2 |
| ZNF205 |
| ZNF207 |
| ZNF208 |
| ZNF211 |
| ZNF212 |
| ZNF217 |
| ZNF219 |
| ZNF22 |
| ZNF227 |
| ZNF236 |
| ZNF239 |
| ZNF24 |
| ZNF248 |
| ZNF25 |
| ZNF250 |
| ZNF26 |
| ZNF260 |
| ZNF263 |
| ZNF268 |

| |
|---|
| ZNF274 |
| ZNF275 |
| ZNF276 |
| ZNF277 |
| ZNF280A |
| ZNF280B |
| ZNF281 |
| ZNF282 |
| ZNF283 |
| ZNF286A |
| ZNF287 |
| ZNF292 |
| ZNF296 |
| ZNF3 |
| ZNF300 |
| ZNF304 |
| ZNF317 |
| ZNF318 |
| ZNF319 |
| ZNF32 |
| ZNF326 |
| ZNF329 |
| ZNF330 |
| ZNF331 |
| ZNF333 |
| ZNF34 |
| ZNF341 |
| ZNF346 |
| ZNF35 |
| ZNF350 |
| ZNF354A |
| ZNF358 |
| ZNF362 |
| ZNF365 |
| ZNF367 |
| ZNF37A |
| ZNF382 |
| ZNF383 |
| ZNF384 |
| ZNF385A |
| ZNF385B |
| ZNF391 |
| ZNF394 |
| ZNF395 |
| ZNF397 |

| |
|---|
| ZNF398 |
| ZNF404 |
| ZNF407 |
| ZNF408 |
| ZNF410 |
| ZNF414 |
| ZNF420 |
| ZNF423 |
| ZNF428 |
| ZNF432 |
| ZNF436 |
| ZNF438 |
| ZNF445 |
| ZNF446 |
| ZNF449 |
| ZNF451 |
| ZNF462 |
| ZNF467 |
| ZNF470 |
| ZNF48 |
| ZNF483 |
| ZNF484 |
| ZNF496 |
| ZNF503 |
| ZNF507 |
| ZNF511 |
| ZNF512 |
| ZNF513 |
| ZNF516 |
| ZNF518A |
| ZNF518B |
| ZNF521 |
| ZNF524 |
| ZNF526 |
| ZNF536 |
| ZNF548 |
| ZNF554 |
| ZNF555 |
| ZNF564 |
| ZNF565 |
| ZNF566 |
| ZNF567 |
| ZNF568 |
| ZNF570 |
| ZNF574 |

| |
|---|
| ZNF575 |
| ZNF576 |
| ZNF579 |
| ZNF581 |
| ZNF583 |
| ZNF584 |
| ZNF593 |
| ZNF596 |
| ZNF599 |
| ZNF608 |
| ZNF609 |
| ZNF613 |
| ZNF614 |
| ZNF618 |
| ZNF619 |
| ZNF622 |
| ZNF623 |
| ZNF628 |
| ZNF629 |
| ZNF638 |
| ZNF641 |
| ZNF644 |
| ZNF646 |
| ZNF652 |
| ZNF653 |
| ZNF655 |
| ZNF664 |
| ZNF667 |
| ZNF668 |
| ZNF672 |
| ZNF674 |
| ZNF677 |
| ZNF684 |
| ZNF687 |
| ZNF688 |
| ZNF689 |
| ZNF691 |
| ZNF692 |
| ZNF696 |
| ZNF697 |
| ZNF703 |
| ZNF704 |
| ZNF710 |
| ZNF711 |
| ZNF713 |

| |
|---|
| ZNF74 |
| ZNF740 |
| ZNF746 |
| ZNF750 |
| ZNF75A |
| ZNF75D |
| ZNF768 |
| ZNF771 |
| ZNF772 |
| ZNF777 |
| ZNF782 |
| ZNF783 |
| ZNF784 |
| ZNF786 |
| ZNF787 |
| ZNF789 |
| ZNF79 |
| ZNF8 |
| ZNF800 |
| ZNF81 |
| ZNF813 |
| ZNF821 |
| ZNF827 |
| ZNF829 |
| ZNF831 |
| ZNF84 |
| ZNF853 |
| ZNFX1 |
| ZNHIT1 |
| ZNHIT2 |
| ZNHIT3 |
| ZNRF1 |
| ZRANB1 |
| ZRANB2 |
| ZRANB3 |
| ZSCAN12 |
| ZSCAN16 |
| ZSCAN2 |
| ZSCAN20 |
| ZSCAN21 |
| ZSCAN23 |
| ZSCAN29 |
| ZSCAN4 |
| ZSWIM1 |
| ZSWIM3 |

| |
|---|
| ZSWIM4 |
| ZSWIM7 |
| ZW10 |
| ZWILCH |
| ZWINT |
| ZXDC |
| ZYG11B |
| ZYX |
| ZZEF1 |
| ZZZ3 |

**Supplemental List 4**

| Gene Name |
|---|
| A2M |
| AAAS |
| AACS |
| AADAC |
| AAK1 |
| AAMP |
| AARSD1 |
| AASDH |
| AASDHPPT |
| AASS |
| AATF |
| AATK |
| ABCA1 |
| ABCA12 |
| ABCA2 |
| ABCA3 |
| ABCA4 |
| ABCA5 |
| ABCA7 |
| ABCB1 |
| ABCB10 |
| ABCB6 |
| ABCB7 |
| ABCB8 |
| ABCB9 |
| ABCC1 |
| ABCC10 |
| ABCC11 |
| ABCC3 |
| ABCC4 |
| ABCD1 |
| ABCD3 |
| ABCD4 |
| ABCE1 |
| ABCF2 |
| ABCF3 |
| ABCG1 |
| ABHD1 |
| ABHD10 |
| ABHD11 |
| ABHD12 |
| ABHD13 |
| ABHD2 |

| |
|---|
| ABHD3 |
| ABHD4 |
| ABHD5 |
| ABHD6 |
| ABHD8 |
| ABI1 |
| ABI3BP |
| ABL2 |
| ABR |
| ABT1 |
| ABTB1 |
| ABTB2 |
| ACAA1 |
| ACAA2 |
| ACACA |
| ACACB |
| ACAD10 |
| ACAD11 |
| ACAD8 |
| ACADS |
| ACADSB |
| ACADVL |
| ACAP1 |
| ACAP2 |
| ACAP3 |
| ACAT1 |
| ACAT2 |
| ACBD3 |
| ACBD4 |
| ACBD5 |
| ACBD6 |
| ACCS |
| ACD |
| ACER3 |
| ACHE |
| ACIN1 |
| ACLY |
| ACOT11 |
| ACOT12 |
| ACOT13 |
| ACOT2 |
| ACOT4 |
| ACOT7 |
| ACOT8 |
| ACOT9 |

| |
|---|
| ACOX1 |
| ACOX3 |
| ACP1 |
| ACP2 |
| ACP6 |
| ACRBP |
| ACSBG1 |
| ACSF2 |
| ACSF3 |
| ACSL1 |
| ACSL3 |
| ACSL6 |
| ACSM3 |
| ACSS1 |
| ACSS2 |
| ACTA1 |
| ACTA2 |
| ACTB |
| ACTBL2 |
| ACTG1 |
| ACTL6A |
| ACTL7A |
| ACTN1 |
| ACTN4 |
| ACTR10 |
| ACTR1B |
| ACTR2 |
| ACTR3 |
| ACTR6 |
| ACTR8 |
| ACVR1 |
| ACVR1B |
| ACVR2A |
| ACVR2B |
| ACY1 |
| ACYP1 |
| ACYP2 |
| ADA |
| ADAL |
| ADAM10 |
| ADAM15 |
| ADAM17 |
| ADAM19 |
| ADAM20 |
| ADAM21 |

| |
|---|
| ADAM22 |
| ADAM32 |
| ADAM8 |
| ADAM9 |
| ADAMTS1 |
| ADAMTS12 |
| ADAMTS13 |
| ADAMTS2 |
| ADAMTS3 |
| ADAMTS5 |
| ADAMTS6 |
| ADAMTS8 |
| ADAMTS9 |
| ADAMTSL4 |
| ADAP1 |
| ADAP2 |
| ADARB1 |
| ADAT1 |
| ADAT2 |
| ADCK1 |
| ADCK2 |
| ADCK5 |
| ADCY1 |
| ADCY3 |
| ADCY4 |
| ADCY6 |
| ADCY7 |
| ADCY9 |
| ADH5 |
| ADHFE1 |
| ADI1 |
| ADIPOR1 |
| ADIPOR2 |
| ADM |
| ADNP2 |
| ADORA1 |
| ADORA2A |
| ADORA2B |
| ADPGK |
| ADPRHL1 |
| ADRA1D |
| ADRA2A |
| ADRM1 |
| ADSL |
| AEBP1 |

| |
|---|
| AEBP2 |
| AFAP1 |
| AFF3 |
| AFF4 |
| AFG3L2 |
| AFMID |
| AFTPH |
| AGAP3 |
| AGBL3 |
| AGBL5 |
| AGFG1 |
| AGFG2 |
| AGGF1 |
| AGK |
| AGL |
| AGMAT |
| AGPAT1 |
| AGPAT2 |
| AGPAT3 |
| AGPAT5 |
| AGPS |
| AGRN |
| AGTPBP1 |
| AGTR2 |
| AHCTF1 |
| AHCY |
| AHCYL1 |
| AHCYL2 |
| AHDC1 |
| AHI1 |
| AHNAK |
| AHNAK2 |
| AHR |
| AHSA1 |
| AIDA |
| AIF1L |
| AIFM1 |
| AIMP1 |
| AIMP2 |
| AIP |
| AK1 |
| AK2 |
| AK6 |
| AKAP1 |
| AKAP11 |

| |
|---|
| AKAP12 |
| AKAP13 |
| AKAP5 |
| AKAP6 |
| AKAP7 |
| AKAP8 |
| AKAP8L |
| AKIP1 |
| AKIRIN1 |
| AKIRIN2 |
| AKNA |
| AKR1A1 |
| AKR1B1 |
| AKT1 |
| AKT1S1 |
| AKT2 |
| AKT3 |
| AKTIP |
| ALAS1 |
| ALCAM |
| ALDH16A1 |
| ALDH18A1 |
| ALDH1A1 |
| ALDH1B1 |
| ALDH1L2 |
| ALDH2 |
| ALDH4A1 |
| ALDH5A1 |
| ALDH6A1 |
| ALDH7A1 |
| ALDH9A1 |
| ALDOA |
| ALG1 |
| ALG11 |
| ALG12 |
| ALG13 |
| ALG14 |
| ALG2 |
| ALG3 |
| ALG5 |
| ALG6 |
| ALG8 |
| ALKBH1 |
| ALKBH2 |
| ALKBH3 |

| |
|---|
| ALKBH4 |
| ALKBH5 |
| ALKBH7 |
| ALMS1 |
| ALOX15 |
| ALPK1 |
| ALPL |
| ALS2 |
| AMACR |
| AMBRA1 |
| AMD1 |
| AMDHD1 |
| AMFR |
| AMH |
| AMIGO1 |
| AMIGO2 |
| AMIGO3 |
| AMN |
| AMN1 |
| AMOT |
| AMOTL1 |
| AMOTL2 |
| AMPD2 |
| AMPD3 |
| AMZ2 |
| ANAPC1 |
| ANAPC10 |
| ANAPC11 |
| ANAPC13 |
| ANAPC7 |
| ANG |
| ANGEL1 |
| ANGEL2 |
| ANGPT2 |
| ANGPTL1 |
| ANGPTL5 |
| ANGPTL7 |
| ANK1 |
| ANK2 |
| ANK3 |
| ANKFY1 |
| ANKH |
| ANKIB1 |
| ANKLE1 |
| ANKLE2 |

| |
|---|
| ANKMY2 |
| ANKRA2 |
| ANKRD10 |
| ANKRD11 |
| ANKRD12 |
| ANKRD13A |
| ANKRD13C |
| ANKRD13D |
| ANKRD16 |
| ANKRD17 |
| ANKRD22 |
| ANKRD26 |
| ANKRD27 |
| ANKRD28 |
| ANKRD29 |
| ANKRD31 |
| ANKRD33B |
| ANKRD34A |
| ANKRD37 |
| ANKRD39 |
| ANKRD40 |
| ANKRD46 |
| ANKRD49 |
| ANKRD50 |
| ANKRD52 |
| ANKRD9 |
| ANKS3 |
| ANKS4B |
| ANKS6 |
| ANKZF1 |
| ANLN |
| ANO10 |
| ANO6 |
| ANO8 |
| ANO9 |
| ANP32A |
| ANP32B |
| ANP32E |
| ANTXR1 |
| ANTXR2 |
| ANXA1 |
| ANXA11 |
| ANXA13 |
| ANXA2 |
| ANXA3 |

| |
|---|
| ANXA5 |
| AOAH |
| AOC3 |
| AOX1 |
| AP1AR |
| AP1B1 |
| AP1G1 |
| AP1G2 |
| AP1M1 |
| AP1S1 |
| AP1S2 |
| AP2A1 |
| AP2A2 |
| AP2M1 |
| AP2S1 |
| AP3B1 |
| AP3D1 |
| AP3M1 |
| AP3S1 |
| AP3S2 |
| AP4B1 |
| AP4E1 |
| AP4M1 |
| AP4S1 |
| APAF1 |
| APBB1 |
| APBB2 |
| APBB3 |
| APC |
| APEH |
| APEX1 |
| APEX2 |
| APH1A |
| APH1B |
| API5 |
| APIP |
| APLF |
| APLP2 |
| APMAP |
| APOA1 |
| APOD |
| APOE |
| APOL3 |
| APOLD1 |
| APOM |

| |
|---|
| APOO |
| APP |
| APPBP2 |
| APPL1 |
| APPL2 |
| APRT |
| APTX |
| AQP11 |
| AQP8 |
| AQR |
| AR |
| ARAF |
| ARAP1 |
| ARAP2 |
| ARC |
| ARF1 |
| ARF4 |
| ARF5 |
| ARF6 |
| ARFGAP2 |
| ARFGAP3 |
| ARFGEF1 |
| ARFGEF2 |
| ARFIP1 |
| ARFIP2 |
| ARFRP1 |
| ARG2 |
| ARGLU1 |
| ARHGAP1 |
| ARHGAP10 |
| ARHGAP11A |
| ARHGAP17 |
| ARHGAP18 |
| ARHGAP19 |
| ARHGAP20 |
| ARHGAP21 |
| ARHGAP22 |
| ARHGAP23 |
| ARHGAP26 |
| ARHGAP29 |
| ARHGAP30 |
| ARHGAP4 |
| ARHGAP5 |
| ARHGAP9 |
| ARHGDIA |

| |
|---|
| ARHGDIB |
| ARHGDIG |
| ARHGEF1 |
| ARHGEF10 |
| ARHGEF10L |
| ARHGEF11 |
| ARHGEF12 |
| ARHGEF16 |
| ARHGEF18 |
| ARHGEF19 |
| ARHGEF2 |
| ARHGEF3 |
| ARHGEF4 |
| ARHGEF6 |
| ARHGEF7 |
| ARHGEF9 |
| ARID1A |
| ARID1B |
| ARID2 |
| ARID3A |
| ARID3B |
| ARID4A |
| ARID5A |
| ARID5B |
| ARIH2 |
| ARL1 |
| ARL10 |
| ARL11 |
| ARL13B |
| ARL15 |
| ARL2 |
| ARL2BP |
| ARL3 |
| ARL4A |
| ARL4C |
| ARL4D |
| ARL5A |
| ARL5B |
| ARL6 |
| ARL6IP1 |
| ARL6IP5 |
| ARL6IP6 |
| ARL8B |
| ARMC1 |
| ARMC5 |

| |
|---|
| ARMC6 |
| ARMC7 |
| ARMC8 |
| ARMC9 |
| ARMCX1 |
| ARNT2 |
| ARNTL |
| ARPC1B |
| ARPC3 |
| ARPC4 |
| ARPC5 |
| ARPC5L |
| ARPP21 |
| ARRB1 |
| ARRDC1 |
| ARRDC2 |
| ARRDC3 |
| ARRDC4 |
| ARSA |
| ARSG |
| ARSK |
| ART4 |
| AS3MT |
| ASAH1 |
| ASAP1 |
| ASAP2 |
| ASAP3 |
| ASB1 |
| ASB13 |
| ASB16 |
| ASB3 |
| ASB4 |
| ASB6 |
| ASB7 |
| ASB8 |
| ASCC1 |
| ASCC2 |
| ASCC3 |
| ASCL1 |
| ASF1A |
| ASF1B |
| ASH1L |
| ASL |
| ASMTL |
| ASNS |

| |
|---|
| ASNSD1 |
| ASPG |
| ASPH |
| ASPHD2 |
| ASPM |
| ASPSCR1 |
| ASRGL1 |
| ASTE1 |
| ASXL1 |
| ASXL2 |
| ASXL3 |
| ATAD1 |
| ATAD2 |
| ATAD5 |
| ATE1 |
| ATF2 |
| ATF3 |
| ATF4 |
| ATF5 |
| ATF7 |
| ATF7IP |
| ATF7IP2 |
| ATG10 |
| ATG101 |
| ATG12 |
| ATG13 |
| ATG16L1 |
| ATG16L2 |
| ATG2A |
| ATG2B |
| ATG3 |
| ATG4A |
| ATG4B |
| ATG4C |
| ATG5 |
| ATG7 |
| ATG9A |
| ATIC |
| ATL3 |
| ATM |
| ATMIN |
| ATN1 |
| ATOH7 |
| ATOX1 |
| ATP10D |

| |
|---|
| ATP11A |
| ATP11B |
| ATP11C |
| ATP12A |
| ATP13A1 |
| ATP13A2 |
| ATP1A1 |
| ATP1A3 |
| ATP1B1 |
| ATP1B2 |
| ATP1B3 |
| ATP2A1 |
| ATP2A2 |
| ATP2A3 |
| ATP2B4 |
| ATP2C1 |
| ATP4A |
| ATP4B |
| ATP6AP1 |
| ATP6AP2 |
| ATP6V0A1 |
| ATP6V0A2 |
| ATP6V0A4 |
| ATP6V0B |
| ATP6V0C |
| ATP6V0D1 |
| ATP6V0D2 |
| ATP6V0E1 |
| ATP6V1A |
| ATP6V1B2 |
| ATP6V1D |
| ATP6V1E1 |
| ATP6V1F |
| ATP6V1G1 |
| ATP6V1G2 |
| ATP6V1H |
| ATP7A |
| ATP7B |
| ATP8A1 |
| ATP8A2 |
| ATP8B1 |
| ATP8B2 |
| ATP8B3 |
| ATP9B |
| ATPAF1 |

| |
|---|
| ATPAF2 |
| ATR |
| ATRIP |
| ATRN |
| ATRNL1 |
| ATRX |
| ATXN1L |
| ATXN2 |
| ATXN2L |
| ATXN3 |
| ATXN7 |
| ATXN7L1 |
| ATXN7L2 |
| ATXN7L3 |
| AUH |
| AUP1 |
| AURKA |
| AURKAIP1 |
| AURKC |
| AUTS2 |
| AVEN |
| AVL9 |
| AVPI1 |
| AXIN1 |
| AXIN2 |
| AXL |
| AZI2 |
| AZIN1 |
| B2M |
| B3GALNT2 |
| B3GALT2 |
| B3GALT4 |
| B3GALT6 |
| B3GAT2 |
| B3GAT3 |
| B3GNT2 |
| B3GNT3 |
| B3GNT4 |
| B3GNT5 |
| B3GNT7 |
| B3GNT8 |
| B3GNT9 |
| B3GNTL1 |
| B4GALNT1 |
| B4GALNT2 |

| |
|---|
| B4GALNT3 |
| B4GALT1 |
| B4GALT2 |
| B4GALT3 |
| B4GALT4 |
| B4GALT5 |
| B4GALT7 |
| B9D1 |
| B9D2 |
| BAALC |
| BACE1 |
| BACE2 |
| BACH2 |
| BAD |
| BAG1 |
| BAG2 |
| BAG3 |
| BAG4 |
| BAG5 |
| BAHCC1 |
| BAHD1 |
| BAIAP2 |
| BAIAP2L2 |
| BAK1 |
| BAMBI |
| BANF1 |
| BANP |
| BARD1 |
| BASP1 |
| BATF2 |
| BAX |
| BAZ1B |
| BAZ2A |
| BAZ2B |
| BBS12 |
| BBS2 |
| BBS4 |
| BBS5 |
| BBS7 |
| BBS9 |
| BBX |
| BCAM |
| BCAP29 |
| BCAP31 |
| BCAR1 |

| |
|---|
| BCAR3 |
| BCAS1 |
| BCAS2 |
| BCAS3 |
| BCAS4 |
| BCAT2 |
| BCCIP |
| BCDIN3D |
| BCHE |
| BCKDHA |
| BCKDHB |
| BCKDK |
| BCL10 |
| BCL11A |
| BCL2 |
| BCL2L1 |
| BCL2L11 |
| BCL2L12 |
| BCL2L15 |
| BCL3 |
| BCL6 |
| BCL6B |
| BCL7B |
| BCL7C |
| BCL9 |
| BCLAF1 |
| BCOR |
| BCORL1 |
| BCR |
| BCS1L |
| BDH2 |
| BDP1 |
| BECN1 |
| BEND3 |
| BEND5 |
| BEND6 |
| BEND7 |
| BEST1 |
| BEST2 |
| BEST4 |
| BET1L |
| BEX2 |
| BEX4 |
| BFAR |
| BFSP1 |

| |
|---|
| BHLHB9 |
| BHLHE41 |
| BICD1 |
| BICD2 |
| BID |
| BIN1 |
| BIN3 |
| BIRC3 |
| BIRC5 |
| BIRC6 |
| BLCAP |
| BLM |
| BLMH |
| BLOC1S1 |
| BLOC1S2 |
| BLVRA |
| BLVRB |
| BLZF1 |
| BMF |
| BMP1 |
| BMP2 |
| BMP3 |
| BMP6 |
| BMPR1A |
| BMPR2 |
| BMS1 |
| BNC2 |
| BNIP1 |
| BNIP3 |
| BNIP3L |
| BOK |
| BOLA1 |
| BOLA3 |
| BOP1 |
| BPGM |
| BPNT1 |
| BPTF |
| BRAF |
| BRCA1 |
| BRCC3 |
| BRD2 |
| BRD3 |
| BRD4 |
| BRD7 |
| BRD8 |

| |
|---|
| BRD9 |
| BRDT |
| BRI3 |
| BRI3BP |
| BRIP1 |
| BRIX1 |
| BRMS1L |
| BRPF1 |
| BRPF3 |
| BRSK1 |
| BRWD1 |
| BRWD3 |
| BSCL2 |
| BSDC1 |
| BSG |
| BSPRY |
| BST1 |
| BST2 |
| BTAF1 |
| BTBD1 |
| BTBD10 |
| BTBD3 |
| BTBD6 |
| BTBD7 |
| BTC |
| BTD |
| BTF3 |
| BTF3L4 |
| BTG1 |
| BTG2 |
| BTG3 |
| BTK |
| BTN1A1 |
| BTN3A3 |
| BTNL9 |
| BTRC |
| BUB1 |
| BUB1B |
| BUB3 |
| BUD13 |
| BUD31 |
| BYSL |
| BZW1 |
| BZW2 |
| C1D |

| |
|---|
| C1GALT1 |
| C1GALT1C1 |
| C1orf35 |
| C1orf52 |
| C1QBP |
| C1QC |
| C1QTNF6 |
| C1R |
| C1RL |
| C1S |
| C2 |
| C2CD2 |
| C2CD2L |
| C4A |
| C5AR1 |
| C6orf120 |
| C8G |
| C9orf64 |
| CA12 |
| CA2 |
| CA4 |
| CA5B |
| CA7 |
| CA9 |
| CAB39L |
| CABIN1 |
| CABLES2 |
| CABYR |
| CACHD1 |
| CACNA1G |
| CACNA1H |
| CACNA2D1 |
| CACNB1 |
| CACNB2 |
| CACNB3 |
| CACYBP |
| CAD |
| CADM1 |
| CADM4 |
| CALCOCO1 |
| CALCOCO2 |
| CALD1 |
| CALHM2 |
| CALM1 |
| CALR |

| |
|---|
| CALU |
| CAMK2D |
| CAMK2G |
| CAMK2N1 |
| CAMKK1 |
| CAMKV |
| CAMLG |
| CAMSAP1 |
| CAMTA1 |
| CAND1 |
| CANT1 |
| CANX |
| CAP1 |
| CAPG |
| CAPN1 |
| CAPN10 |
| CAPN2 |
| CAPN6 |
| CAPN7 |
| CAPNS1 |
| CAPRIN2 |
| CAPZA1 |
| CAPZA2 |
| CAPZB |
| CARD6 |
| CARHSP1 |
| CARM1 |
| CARS2 |
| CASC3 |
| CASD1 |
| CASKIN1 |
| CASKIN2 |
| CASP2 |
| CASP3 |
| CASP7 |
| CASP8 |
| CASP8AP2 |
| CASP9 |
| CASQ1 |
| CASZ1 |
| CATSPER2 |
| CAV2 |
| CBFA2T2 |
| CBFB |
| CBL |

| |
|---|
| CBLB |
| CBLC |
| CBLL1 |
| CBR1 |
| CBR4 |
| CBS |
| CBX1 |
| CBX2 |
| CBX3 |
| CBX5 |
| CBX6 |
| CBX7 |
| CBX8 |
| CBY1 |
| CC2D1A |
| CC2D1B |
| CC2D2A |
| CCAR1 |
| CCBE1 |
| CCDC102A |
| CCDC106 |
| CCDC107 |
| CCDC112 |
| CCDC115 |
| CCDC117 |
| CCDC12 |
| CCDC121 |
| CCDC122 |
| CCDC124 |
| CCDC125 |
| CCDC126 |
| CCDC127 |
| CCDC137 |
| CCDC138 |
| CCDC14 |
| CCDC142 |
| CCDC149 |
| CCDC15 |
| CCDC157 |
| CCDC158 |
| CCDC159 |
| CCDC160 |
| CCDC18 |
| CCDC22 |
| CCDC24 |

| |
|---|
| CCDC25 |
| CCDC28A |
| CCDC28B |
| CCDC32 |
| CCDC34 |
| CCDC43 |
| CCDC47 |
| CCDC50 |
| CCDC51 |
| CCDC59 |
| CCDC61 |
| CCDC65 |
| CCDC66 |
| CCDC69 |
| CCDC7 |
| CCDC71 |
| CCDC73 |
| CCDC74B |
| CCDC77 |
| CCDC80 |
| CCDC82 |
| CCDC85A |
| CCDC85B |
| CCDC85C |
| CCDC86 |
| CCDC88A |
| CCDC88B |
| CCDC88C |
| CCDC9 |
| CCDC90B |
| CCDC91 |
| CCDC93 |
| CCDC96 |
| CCDC97 |
| CCHCR1 |
| CCIN |
| CCL2 |
| CCL20 |
| CCL28 |
| CCL5 |
| CCL8 |
| CCM2 |
| CCNA2 |
| CCNB2 |
| CCNC |

| |
|---|
| CCND3 |
| CCNE1 |
| CCNE2 |
| CCNG1 |
| CCNG2 |
| CCNH |
| CCNI |
| CCNJ |
| CCNK |
| CCNL2 |
| CCNT1 |
| CCNT2 |
| CCNY |
| CCNYL1 |
| CCPG1 |
| CCR7 |
| CCS |
| CCT2 |
| CCT3 |
| CCT4 |
| CCT5 |
| CCT6A |
| CCT7 |
| CCT8 |
| CD109 |
| CD14 |
| CD151 |
| CD164 |
| CD180 |
| CD200 |
| CD2AP |
| CD2BP2 |
| CD302 |
| CD36 |
| CD37 |
| CD40 |
| CD44 |
| CD46 |
| CD47 |
| CD55 |
| CD58 |
| CD63 |
| CD68 |
| CD70 |
| CD72 |

| |
|---|
| CD74 |
| CD81 |
| CD82 |
| CD84 |
| CD8A |
| CD9 |
| CD99L2 |
| CDADC1 |
| CDAN1 |
| CDC14A |
| CDC14B |
| CDC16 |
| CDC25B |
| CDC25C |
| CDC26 |
| CDC27 |
| CDC34 |
| CDC37 |
| CDC37L1 |
| CDC40 |
| CDC42 |
| CDC42BPA |
| CDC42BPG |
| CDC42EP2 |
| CDC42EP3 |
| CDC42EP4 |
| CDC42EP5 |
| CDC42SE2 |
| CDC5L |
| CDC6 |
| CDC7 |
| CDC73 |
| CDCA2 |
| CDCA3 |
| CDCA4 |
| CDCA5 |
| CDCA7 |
| CDCA7L |
| CDCA8 |
| CDCP1 |
| CDH10 |
| CDH12 |
| CDH18 |
| CDH2 |
| CDH24 |

| |
|---|
| CDH3 |
| CDH6 |
| CDH8 |
| CDHR2 |
| CDHR5 |
| CDIPT |
| CDK1 |
| CDK10 |
| CDK12 |
| CDK13 |
| CDK16 |
| CDK17 |
| CDK18 |
| CDK19 |
| CDK20 |
| CDK2AP1 |
| CDK2AP2 |
| CDK3 |
| CDK4 |
| CDK5 |
| CDK5R1 |
| CDK5RAP1 |
| CDK5RAP2 |
| CDK6 |
| CDK7 |
| CDK8 |
| CDK9 |
| CDKAL1 |
| CDKL1 |
| CDKL3 |
| CDKL5 |
| CDKN1A |
| CDKN1B |
| CDKN1C |
| CDKN2A |
| CDKN2AIPNL |
| CDKN2B |
| CDKN2C |
| CDKN2D |
| CDKN3 |
| CDO1 |
| CDON |
| CDR2 |
| CDR2L |
| CDS2 |

| |
|---|
| CDV3 |
| CDYL |
| CDYL2 |
| CEBPA |
| CEBPD |
| CEBPG |
| CEBPZ |
| CELSR1 |
| CELSR2 |
| CENPB |
| CENPE |
| CENPF |
| CENPH |
| CENPI |
| CENPJ |
| CENPK |
| CENPM |
| CENPN |
| CENPO |
| CENPP |
| CENPQ |
| CENPT |
| CENPV |
| CEP135 |
| CEP164 |
| CEP250 |
| CEP350 |
| CEP55 |
| CEP57 |
| CEP68 |
| CEP70 |
| CEP72 |
| CEP76 |
| CEP78 |
| CEP97 |
| CERCAM |
| CES1 |
| CETN2 |
| CETN3 |
| CFB |
| CFD |
| CFDP1 |
| CFL1 |
| CFL2 |
| CGN |

| |
|---|
| CGREF1 |
| CGRRF1 |
| CHAC1 |
| CHAC2 |
| CHADL |
| CHAF1A |
| CHAF1B |
| CHCHD3 |
| CHCHD4 |
| CHCHD6 |
| CHCHD7 |
| CHD1L |
| CHD2 |
| CHD3 |
| CHD4 |
| CHD8 |
| CHD9 |
| CHEK1 |
| CHEK2 |
| CHERP |
| CHFR |
| CHGB |
| CHI3L1 |
| CHIC1 |
| CHIC2 |
| CHKA |
| CHL1 |
| CHML |
| CHMP1A |
| CHMP1B |
| CHMP2A |
| CHMP2B |
| CHMP4A |
| CHMP4B |
| CHMP4C |
| CHMP5 |
| CHMP6 |
| CHORDC1 |
| CHPT1 |
| CHRAC1 |
| CHRD |
| CHRDL1 |
| CHRM3 |
| CHRM5 |
| CHST11 |

| |
|---|
| CHST12 |
| CHST13 |
| CHST14 |
| CHST3 |
| CHST7 |
| CHSY1 |
| CHSY3 |
| CHUK |
| CIAO1 |
| CIAPIN1 |
| CIB1 |
| CINP |
| CIRBP |
| CISD1 |
| CISD2 |
| CISD3 |
| CISH |
| CIT |
| CITED2 |
| CITED4 |
| CIZ1 |
| CKAP2 |
| CKAP2L |
| CKAP4 |
| CKAP5 |
| CKB |
| CKMT2 |
| CKS1B |
| CLASP1 |
| CLASP2 |
| CLCA2 |
| CLCN2 |
| CLCN3 |
| CLCN5 |
| CLCN6 |
| CLCN7 |
| CLDN1 |
| CLDN11 |
| CLDN12 |
| CLDN15 |
| CLDN20 |
| CLDN23 |
| CLDN3 |
| CLDN4 |
| CLDN7 |

| |
|---|
| CLDN8 |
| CLEC11A |
| CLEC16A |
| CLEC3A |
| CLEC3B |
| CLGN |
| CLIC1 |
| CLIC4 |
| CLINT1 |
| CLIP1 |
| CLIP2 |
| CLIP3 |
| CLK1 |
| CLK3 |
| CLK4 |
| CLMN |
| CLN3 |
| CLN5 |
| CLN6 |
| CLN8 |
| CLNS1A |
| CLPB |
| CLPP |
| CLPTM1 |
| CLPTM1L |
| CLPX |
| CLSPN |
| CLSTN1 |
| CLSTN2 |
| CLSTN3 |
| CLTA |
| CLTB |
| CLTC |
| CLU |
| CLUAP1 |
| CLYBL |
| CMBL |
| CMC1 |
| CMPK1 |
| CMPK2 |
| CMTM3 |
| CMTM6 |
| CMTM7 |
| CMYA5 |
| CNBP |

| |
|---|
| CNDP2 |
| CNIH4 |
| CNKSR3 |
| CNN1 |
| CNN2 |
| CNN3 |
| CNNM2 |
| CNNM3 |
| CNNM4 |
| CNOT1 |
| CNOT10 |
| CNOT3 |
| CNOT4 |
| CNOT6 |
| CNOT6L |
| CNOT7 |
| CNOT8 |
| CNP |
| CNPY2 |
| CNPY3 |
| CNPY4 |
| CNR2 |
| CNTLN |
| CNTN1 |
| CNTN3 |
| CNTN6 |
| CNTNAP1 |
| CNTROB |
| COASY |
| COBL |
| COBLL1 |
| COCH |
| COG1 |
| COG2 |
| COG3 |
| COG4 |
| COG5 |
| COG6 |
| COG7 |
| COG8 |
| COIL |
| COL15A1 |
| COL17A1 |
| COL1A1 |
| COL21A1 |

| |
|---|
| COL3A1 |
| COL4A1 |
| COL4A2 |
| COL4A3 |
| COL4A4 |
| COL4A5 |
| COL4A6 |
| COL6A1 |
| COL6A2 |
| COL7A1 |
| COL9A2 |
| COLEC12 |
| COMMD1 |
| COMMD10 |
| COMMD2 |
| COMMD4 |
| COMMD5 |
| COMMD6 |
| COMMD7 |
| COMMD8 |
| COMMD9 |
| COMT |
| COMTD1 |
| COPA |
| COPB1 |
| COPB2 |
| COPE |
| COPG2 |
| COPS3 |
| COPS5 |
| COPS6 |
| COPS7A |
| COPS8 |
| COPZ1 |
| COQ10A |
| COQ10B |
| COQ2 |
| COQ3 |
| COQ5 |
| COQ6 |
| COQ7 |
| COQ9 |
| CORIN |
| CORO1A |
| CORO1B |

| |
|---|
| CORO1C |
| CORO2A |
| CORO6 |
| CORO7 |
| COTL1 |
| COX10 |
| COX11 |
| COX15 |
| COX16 |
| COX18 |
| COX19 |
| COX5A |
| COX5B |
| COX6A1 |
| COX6B1 |
| COX6C |
| COX7A1 |
| COX7A2 |
| COX7A2L |
| COX8A |
| CPD |
| CPEB1 |
| CPEB2 |
| CPEB4 |
| CPNE3 |
| CPOX |
| CPPED1 |
| CPSF1 |
| CPSF2 |
| CPSF3 |
| CPSF4 |
| CPT1A |
| CPT1B |
| CPT1C |
| CPT2 |
| CPVL |
| CPXM1 |
| CPZ |
| CRADD |
| CRAT |
| CRBN |
| CRCP |
| CREB1 |
| CREB3 |
| CREB3L1 |

| |
|---|
| CREB3L2 |
| CREBL2 |
| CREBZF |
| CREG1 |
| CRELD1 |
| CRELD2 |
| CREM |
| CRIM1 |
| CRIP1 |
| CRIP3 |
| CRIPT |
| CRK |
| CRLF3 |
| CRLS1 |
| CRNKL1 |
| CROCC |
| CRTAP |
| CRTC3 |
| CRY2 |
| CRYAB |
| CRYBG3 |
| CRYM |
| CRYZ |
| CRYZL1 |
| CS |
| CSDE1 |
| CSF1 |
| CSF1R |
| CSGALNACT1 |
| CSGALNACT2 |
| CSK |
| CSMD3 |
| CSNK1A1 |
| CSNK1D |
| CSNK1E |
| CSNK1G1 |
| CSNK1G2 |
| CSNK1G3 |
| CSNK2B |
| CSPG4 |
| CSPG5 |
| CSPP1 |
| CSRNP1 |
| CSRNP2 |
| CSRP1 |

| |
|---|
| CSRP2 |
| CST3 |
| CST6 |
| CST7 |
| CSTF1 |
| CSTF2 |
| CSTF2T |
| CSTF3 |
| CSTL1 |
| CTBP1 |
| CTBP2 |
| CTCF |
| CTDSP1 |
| CTDSP2 |
| CTF1 |
| CTH |
| CTHRC1 |
| CTNNA1 |
| CTNNA2 |
| CTNNAL1 |
| CTNNB1 |
| CTNNBL1 |
| CTNND1 |
| CTR9 |
| CTSA |
| CTSB |
| CTSC |
| CTSD |
| CTSH |
| CTSK |
| CTSO |
| CTSZ |
| CTTN |
| CTTNBP2NL |
| CTU1 |
| CTU2 |
| CTXN1 |
| CUEDC2 |
| CUL1 |
| CUL2 |
| CUL3 |
| CUL4B |
| CUL5 |
| CUL7 |
| CUTA |

| |
|---|
| CUTC |
| CUX1 |
| CWC15 |
| CWC22 |
| CWC25 |
| CWC27 |
| CWF19L1 |
| CWF19L2 |
| CX3CR1 |
| CXADR |
| CXCL10 |
| CXCL16 |
| CXCR4 |
| CYB561D2 |
| CYB5B |
| CYB5R1 |
| CYB5R3 |
| CYC1 |
| CYCS |
| CYFIP2 |
| CYHR1 |
| CYLD |
| CYP19A1 |
| CYP1B1 |
| CYP20A1 |
| CYP27A1 |
| CYP27B1 |
| CYP2J2 |
| CYP2U1 |
| CYP4B1 |
| CYP4F3 |
| CYP4V2 |
| CYP8B1 |
| CYSLTR1 |
| CYTH2 |
| CYTH3 |
| D2HGDH |
| DAAM2 |
| DAB2 |
| DAB2IP |
| DACH1 |
| DAD1 |
| DAG1 |
| DAGLA |
| DAGLB |

| |
|---|
| DALRD3 |
| DAP |
| DAPK1 |
| DAPK3 |
| DAXX |
| DAZAP1 |
| DAZAP2 |
| DBF4 |
| DBI |
| DBN1 |
| DBNDD1 |
| DBNL |
| DBP |
| DBR1 |
| DBT |
| DCAF10 |
| DCAF11 |
| DCAF12 |
| DCAF15 |
| DCAF16 |
| DCAF17 |
| DCAF4 |
| DCAF5 |
| DCAF6 |
| DCAF7 |
| DCAF8 |
| DCAKD |
| DCBLD2 |
| DCHS1 |
| DCK |
| DCLRE1A |
| DCLRE1C |
| DCP1A |
| DCP1B |
| DCP2 |
| DCPS |
| DCTD |
| DCTN1 |
| DCTN2 |
| DCTN3 |
| DCTN5 |
| DCTN6 |
| DCTPP1 |
| DCUN1D1 |
| DCUN1D3 |

| |
|---|
| DCUN1D5 |
| DCXR |
| DDA1 |
| DDAH2 |
| DDB1 |
| DDB2 |
| DDHD1 |
| DDHD2 |
| DDI2 |
| DDIT3 |
| DDIT4 |
| DDN |
| DDOST |
| DDR1 |
| DDR2 |
| DDRGK1 |
| DDT |
| DDX1 |
| DDX10 |
| DDX17 |
| DDX18 |
| DDX19A |
| DDX19B |
| DDX20 |
| DDX21 |
| DDX23 |
| DDX24 |
| DDX27 |
| DDX28 |
| DDX31 |
| DDX3X |
| DDX41 |
| DDX46 |
| DDX47 |
| DDX49 |
| DDX5 |
| DDX50 |
| DDX52 |
| DDX54 |
| DDX55 |
| DDX56 |
| DDX58 |
| DDX59 |
| DDX6 |
| DEAF1 |

| |
|---|
| DECR1 |
| DECR2 |
| DEDD |
| DEDD2 |
| DEF8 |
| DEGS1 |
| DEGS2 |
| DENND1A |
| DENND1B |
| DENND1C |
| DENND2A |
| DENND2C |
| DENND2D |
| DENND3 |
| DENND4A |
| DENND4B |
| DENND4C |
| DENND5A |
| DENND5B |
| DEPDC1 |
| DEPDC1B |
| DEPDC5 |
| DEPDC7 |
| DERL1 |
| DERL2 |
| DERL3 |
| DET1 |
| DEXI |
| DFFA |
| DFFB |
| DGAT1 |
| DGAT2 |
| DGCR2 |
| DGCR8 |
| DGKA |
| DGKE |
| DGKQ |
| DGKZ |
| DGUOK |
| DHCR24 |
| DHCR7 |
| DHDDS |
| DHDH |
| DHFR |
| DHH |

| |
|---|
| DHODH |
| DHPS |
| DHRS1 |
| DHRS11 |
| DHRS13 |
| DHRS4 |
| DHRS7 |
| DHRS7B |
| DHRSX |
| DHTKD1 |
| DHX15 |
| DHX16 |
| DHX29 |
| DHX30 |
| DHX32 |
| DHX33 |
| DHX34 |
| DHX35 |
| DHX36 |
| DHX37 |
| DHX38 |
| DHX40 |
| DHX57 |
| DHX9 |
| DIABLO |
| DIAPH2 |
| DIAPH3 |
| DICER1 |
| DIDO1 |
| DIP2A |
| DIP2B |
| DIP2C |
| DIRAS1 |
| DIRAS2 |
| DIRAS3 |
| DIS3 |
| DIS3L |
| DIS3L2 |
| DIXDC1 |
| DKC1 |
| DKK3 |
| DKKL1 |
| DLAT |
| DLC1 |
| DLD |

| |
|---|
| DLEC1 |
| DLG1 |
| DLG4 |
| DLG5 |
| DLGAP1 |
| DLGAP5 |
| DLL1 |
| DLL3 |
| DLX5 |
| DMAP1 |
| DMD |
| DMKN |
| DMPK |
| DMTF1 |
| DMWD |
| DMXL1 |
| DMXL2 |
| DNA2 |
| DNAH1 |
| DNAH2 |
| DNAH7 |
| DNAJA2 |
| DNAJA3 |
| DNAJA4 |
| DNAJB1 |
| DNAJB12 |
| DNAJB14 |
| DNAJB2 |
| DNAJB4 |
| DNAJB6 |
| DNAJB7 |
| DNAJB9 |
| DNAJC10 |
| DNAJC11 |
| DNAJC12 |
| DNAJC13 |
| DNAJC14 |
| DNAJC16 |
| DNAJC17 |
| DNAJC18 |
| DNAJC19 |
| DNAJC2 |
| DNAJC21 |
| DNAJC24 |
| DNAJC25 |

| |
|---|
| DNAJC27 |
| DNAJC3 |
| DNAJC30 |
| DNAJC4 |
| DNAJC5 |
| DNAJC7 |
| DNAJC8 |
| DNAJC9 |
| DNAL1 |
| DNAL4 |
| DNASE1L1 |
| DNASE2 |
| DNHD1 |
| DNM1 |
| DNM1L |
| DNM2 |
| DNMBP |
| DNMT1 |
| DNMT3A |
| DNMT3B |
| DNPEP |
| DNTTIP1 |
| DNTTIP2 |
| DOC2B |
| DOCK1 |
| DOCK11 |
| DOCK3 |
| DOCK4 |
| DOCK5 |
| DOCK6 |
| DOCK7 |
| DOCK8 |
| DOCK9 |
| DOHH |
| DOK1 |
| DOK3 |
| DOK4 |
| DOK7 |
| DOLK |
| DOLPP1 |
| DPCD |
| DPF2 |
| DPF3 |
| DPH2 |
| DPH3 |

| |
|---|
| DPH5 |
| DPM1 |
| DPM2 |
| DPM3 |
| DPP3 |
| DPP7 |
| DPY19L1 |
| DPY19L3 |
| DPY19L4 |
| DPY30 |
| DPYD |
| DPYSL2 |
| DPYSL3 |
| DPYSL5 |
| DQX1 |
| DR1 |
| DRAM2 |
| DRAP1 |
| DRG1 |
| DRG2 |
| DRP2 |
| DSC2 |
| DSCC1 |
| DSE |
| DSG2 |
| DSP |
| DST |
| DSTN |
| DSTYK |
| DTD1 |
| DTNB |
| DTNBP1 |
| DTWD1 |
| DTWD2 |
| DTX1 |
| DTX2 |
| DTX3 |
| DTX3L |
| DTYMK |
| DUOX1 |
| DUS1L |
| DUS3L |
| DUS4L |
| DUSP1 |
| DUSP10 |

| |
|---|
| DUSP11 |
| DUSP12 |
| DUSP14 |
| DUSP16 |
| DUSP18 |
| DUSP19 |
| DUSP23 |
| DUSP3 |
| DUSP5 |
| DUSP6 |
| DUSP7 |
| DUSP9 |
| DUT |
| DVL1 |
| DVL2 |
| DYM |
| DYNC1I2 |
| DYNC1LI1 |
| DYNC2LI1 |
| DYNLL1 |
| DYNLL2 |
| DYNLRB1 |
| DYNLT1 |
| DYNLT3 |
| DYRK1A |
| DYRK1B |
| DYRK3 |
| DZIP1 |
| DZIP3 |
| E2F1 |
| E2F2 |
| E2F3 |
| E2F4 |
| E2F5 |
| E2F6 |
| E4F1 |
| EAF1 |
| EAF2 |
| EAPP |
| EBAG9 |
| EBNA1BP2 |
| EBP |
| EBPL |
| ECD |
| ECE1 |

| |
|---|
| ECH1 |
| ECHDC1 |
| ECHDC3 |
| ECHS1 |
| ECSIT |
| ECT2 |
| EDC3 |
| EDC4 |
| EDEM1 |
| EDEM2 |
| EDEM3 |
| EDF1 |
| EDIL3 |
| EDN1 |
| EDNRA |
| EEA1 |
| EED |
| EEF1A1 |
| EEF1D |
| EEF1E1 |
| EEF1G |
| EEF2 |
| EEF2K |
| EEFSEC |
| EEPD1 |
| EFCAB2 |
| EFCAB6 |
| EFCAB7 |
| EFEMP2 |
| EFHD2 |
| EFNA1 |
| EFNA2 |
| EFNA5 |
| EFNB1 |
| EFNB2 |
| EFR3A |
| EFTUD2 |
| EGFL7 |
| EGFL8 |
| EGLN1 |
| EGLN2 |
| EGLN3 |
| EGR1 |
| EGR3 |
| EHBP1L1 |

| |
|---|
| EHD1 |
| EHD2 |
| EHD4 |
| EHF |
| EHHADH |
| EHMT1 |
| EHMT2 |
| EI24 |
| EID1 |
| EID2 |
| EIF1 |
| EIF1B |
| EIF2AK1 |
| EIF2AK2 |
| EIF2AK3 |
| EIF2AK4 |
| EIF2B1 |
| EIF2B2 |
| EIF2B4 |
| EIF2B5 |
| EIF2S1 |
| EIF2S2 |
| EIF2S3 |
| EIF3B |
| EIF3D |
| EIF3E |
| EIF3F |
| EIF3G |
| EIF3I |
| EIF3J |
| EIF3K |
| EIF3L |
| EIF3M |
| EIF4A1 |
| EIF4A2 |
| EIF4A3 |
| EIF4B |
| EIF4E |
| EIF4E2 |
| EIF4E3 |
| EIF4EBP1 |
| EIF4EBP2 |
| EIF4ENIF1 |
| EIF4G2 |
| EIF4G3 |

| |
|---|
| EIF4H |
| EIF5 |
| EIF5A2 |
| EIF5B |
| ELAC1 |
| ELAC2 |
| ELF1 |
| ELF2 |
| ELF3 |
| ELF4 |
| ELK1 |
| ELK3 |
| ELK4 |
| ELL |
| ELL2 |
| ELMO1 |
| ELMO2 |
| ELMO3 |
| ELMOD1 |
| ELMOD2 |
| ELMOD3 |
| ELOVL1 |
| ELOVL5 |
| ELOVL6 |
| ELP2 |
| ELP3 |
| ELP4 |
| EMD |
| EME1 |
| EME2 |
| EMG1 |
| EMID1 |
| EMILIN1 |
| EMILIN2 |
| EML1 |
| EML2 |
| EML3 |
| EML4 |
| EML6 |
| EMP2 |
| EMP3 |
| ENAH |
| ENC1 |
| ENDOD1 |
| ENDOG |

| |
|---|
| ENHO |
| ENO3 |
| ENOPH1 |
| ENOSF1 |
| ENOX1 |
| ENOX2 |
| ENPP1 |
| ENPP4 |
| ENSA |
| ENTPD1 |
| ENTPD2 |
| ENTPD4 |
| ENTPD5 |
| ENTPD6 |
| ENTPD7 |
| ENTPD8 |
| ENY2 |
| EP400 |
| EPAS1 |
| EPB41 |
| EPB41L2 |
| EPB41L3 |
| EPB41L4B |
| EPB41L5 |
| EPC1 |
| EPC2 |
| EPCAM |
| EPDR1 |
| EPHA1 |
| EPHA2 |
| EPHA4 |
| EPHB1 |
| EPHB4 |
| EPHB6 |
| EPHX1 |
| EPHX2 |
| EPM2A |
| EPM2AIP1 |
| EPN1 |
| EPN2 |
| EPN3 |
| EPS15 |
| EPS15L1 |
| EPS8L2 |
| ERAL1 |

| |
|---|
| ERAP1 |
| ERAS |
| ERBB3 |
| ERBB4 |
| ERCC1 |
| ERCC2 |
| ERCC3 |
| ERCC4 |
| ERCC5 |
| ERCC6L |
| ERCC8 |
| ERF |
| ERGIC1 |
| ERGIC2 |
| ERGIC3 |
| ERH |
| ERI2 |
| ERI3 |
| ERLEC1 |
| ERLIN1 |
| ERLIN2 |
| ERMP1 |
| ERN1 |
| ERP29 |
| ERP44 |
| ERRFI1 |
| ESD |
| ESF1 |
| ESPL1 |
| ESR2 |
| ESRP1 |
| ESRP2 |
| ESRRA |
| ESRRG |
| ESYT1 |
| ETAA1 |
| ETF1 |
| ETFA |
| ETFB |
| ETFDH |
| ETHE1 |
| ETNK1 |
| ETNK2 |
| ETS1 |
| ETS2 |

| |
|---|
| ETV2 |
| ETV3 |
| ETV6 |
| EVC |
| EVI5 |
| EVI5L |
| EVL |
| EVPL |
| EWSR1 |
| EXD2 |
| EXD3 |
| EXO1 |
| EXOC1 |
| EXOC2 |
| EXOC3 |
| EXOC4 |
| EXOC5 |
| EXOC6 |
| EXOC6B |
| EXOC7 |
| EXOC8 |
| EXOG |
| EXOSC1 |
| EXOSC10 |
| EXOSC2 |
| EXOSC3 |
| EXOSC4 |
| EXOSC5 |
| EXOSC8 |
| EXOSC9 |
| EXPH5 |
| EXT1 |
| EXT2 |
| EXTL1 |
| EXTL2 |
| EXTL3 |
| EZH1 |
| EZH2 |
| F11R |
| F12 |
| F2RL1 |
| F3 |
| F5 |
| FA2H |
| FAAH |

| |
|---|
| FAAP24 |
| FABP3 |
| FABP4 |
| FABP5 |
| FABP7 |
| FADD |
| FADS1 |
| FADS2 |
| FADS3 |
| FAF1 |
| FAF2 |
| FAH |
| FAHD1 |
| FAHD2A |
| FAIM |
| FAM110A |
| FAM110B |
| FAM114A1 |
| FAM117A |
| FAM118B |
| FAM120A |
| FAM120B |
| FAM149B1 |
| FAM151B |
| FAM155A |
| FAM177A1 |
| FAM193B |
| FAM20B |
| FAM32A |
| FAM50A |
| FAM53C |
| FAM72A |
| FAM76A |
| FAM83D |
| FAM8A1 |
| FANCA |
| FANCB |
| FANCD2 |
| FANCE |
| FANCF |
| FANCG |
| FANCI |
| FANCL |
| FANCM |
| FAR1 |

| |
|---|
| FAR2 |
| FARS2 |
| FARSA |
| FARSB |
| FAS |
| FASTK |
| FASTKD1 |
| FASTKD2 |
| FASTKD3 |
| FASTKD5 |
| FAT1 |
| FAT2 |
| FAT4 |
| FAU |
| FBF1 |
| FBLIM1 |
| FBLN1 |
| FBLN2 |
| FBLN5 |
| FBN1 |
| FBP1 |
| FBP2 |
| FBRSL1 |
| FBXL12 |
| FBXL14 |
| FBXL15 |
| FBXL18 |
| FBXL19 |
| FBXL2 |
| FBXL20 |
| FBXL22 |
| FBXL4 |
| FBXL5 |
| FBXL7 |
| FBXL8 |
| FBXO10 |
| FBXO11 |
| FBXO15 |
| FBXO17 |
| FBXO21 |
| FBXO22 |
| FBXO24 |
| FBXO25 |
| FBXO28 |
| FBXO3 |

| |
|---|
| FBXO31 |
| FBXO32 |
| FBXO33 |
| FBXO34 |
| FBXO38 |
| FBXO4 |
| FBXO42 |
| FBXO43 |
| FBXO45 |
| FBXO5 |
| FBXO6 |
| FBXO7 |
| FBXO8 |
| FBXO9 |
| FBXW11 |
| FBXW12 |
| FBXW2 |
| FBXW4 |
| FBXW5 |
| FBXW9 |
| FCGBP |
| FCGRT |
| FCHO1 |
| FCHO2 |
| FCHSD1 |
| FCHSD2 |
| FDFT1 |
| FDPS |
| FDX1 |
| FDXR |
| FECH |
| FEM1A |
| FEM1B |
| FEM1C |
| FEN1 |
| FER |
| FERMT2 |
| FES |
| FGD1 |
| FGD2 |
| FGD3 |
| FGD4 |
| FGD5 |
| FGD6 |
| FGF1 |

| |
|---|
| FGF13 |
| FGF2 |
| FGFBP1 |
| FGFR1OP2 |
| FGFR2 |
| FGFR3 |
| FGGY |
| FGL2 |
| FH |
| FHDC1 |
| FHL2 |
| FHL3 |
| FHOD1 |
| FHOD3 |
| FIBP |
| FICD |
| FIG4 |
| FIGN |
| FIGNL1 |
| FILIP1 |
| FILIP1L |
| FIP1L1 |
| FITM2 |
| FIZ1 |
| FKBP10 |
| FKBP11 |
| FKBP15 |
| FKBP3 |
| FKBP5 |
| FKBP7 |
| FKBP8 |
| FKBP9 |
| FKBPL |
| FKRP |
| FKTN |
| FLAD1 |
| FLCN |
| FLII |
| FLNA |
| FLNB |
| FLOT1 |
| FLOT2 |
| FLRT1 |
| FLVCR1 |
| FLVCR2 |

| |
|---|
| FLYWCH1 |
| FLYWCH2 |
| FMN1 |
| FMNL1 |
| FMNL3 |
| FMO5 |
| FMR1 |
| FN3K |
| FN3KRP |
| FNBP1 |
| FNBP1L |
| FNDC1 |
| FNDC3A |
| FNDC3B |
| FNDC4 |
| FNIP1 |
| FNIP2 |
| FNTA |
| FNTB |
| FOLR1 |
| FOS |
| FOSB |
| FOSL2 |
| FOXA1 |
| FOXA3 |
| FOXG1 |
| FOXH1 |
| FOXJ3 |
| FOXK2 |
| FOXL2 |
| FOXM1 |
| FOXN2 |
| FOXO1 |
| FOXO3 |
| FOXO4 |
| FOXP1 |
| FOXP4 |
| FOXRED1 |
| FOXRED2 |
| FOXS1 |
| FPGS |
| FPGT |
| FRAT1 |
| FRAT2 |
| FRG1 |

| |
|---|
| FRK |
| FRMD4B |
| FRMD6 |
| FRMD8 |
| FRMPD1 |
| FRRS1 |
| FRS2 |
| FRS3 |
| FRY |
| FRYL |
| FSCN1 |
| FSD1L |
| FSD2 |
| FST |
| FSTL1 |
| FSTL3 |
| FTH1 |
| FTO |
| FTSJ1 |
| FTSJ3 |
| FUBP1 |
| FUBP3 |
| FUCA1 |
| FUCA2 |
| FUNDC2 |
| FURIN |
| FUS |
| FUT1 |
| FUT2 |
| FUZ |
| FXR1 |
| FXR2 |
| FXYD3 |
| FXYD5 |
| FYCO1 |
| FYTTD1 |
| FZD1 |
| FZD2 |
| FZD3 |
| FZD4 |
| FZD5 |
| FZD7 |
| FZD8 |
| FZD9 |
| FZR1 |

| |
|---|
| G0S2 |
| G2E3 |
| G3BP1 |
| G3BP2 |
| G6PC3 |
| G6PD |
| GAA |
| GAB1 |
| GAB2 |
| GABARAP |
| GABBR1 |
| GABPA |
| GABPB1 |
| GABRA2 |
| GABRA3 |
| GABRG1 |
| GABRR2 |
| GADD45A |
| GADD45B |
| GADD45GIP1 |
| GADL1 |
| GAK |
| GAL3ST1 |
| GAL3ST2 |
| GAL3ST3 |
| GAL3ST4 |
| GALC |
| GALE |
| GALK1 |
| GALK2 |
| GALM |
| GALNS |
| GALNT1 |
| GALNT10 |
| GALNT11 |
| GALNT13 |
| GALNT2 |
| GALNT4 |
| GALNT5 |
| GALR3 |
| GALT |
| GAMT |
| GAN |
| GANAB |
| GANC |

| |
|---|
| GAPDHS |
| GAR1 |
| GARNL3 |
| GAS1 |
| GAS2L1 |
| GAS6 |
| GAS7 |
| GAS8 |
| GATAD2A |
| GATAD2B |
| GATC |
| GBA |
| GBA2 |
| GBA3 |
| GBE1 |
| GBP6 |
| GCA |
| GCAT |
| GCC1 |
| GCC2 |
| GCDH |
| GCH1 |
| GCHFR |
| GCLM |
| GCNT1 |
| GCNT3 |
| GCSH |
| GDA |
| GDAP2 |
| GDE1 |
| GDF9 |
| GDI1 |
| GDI2 |
| GDPD3 |
| GDPD5 |
| GEM |
| GEMIN4 |
| GEMIN6 |
| GEMIN7 |
| GEMIN8 |
| GEN1 |
| GFER |
| GFM1 |
| GFM2 |
| GFOD2 |

| |
|---|
| GFPT1 |
| GFPT2 |
| GGA1 |
| GGA3 |
| GGCT |
| GGCX |
| GGH |
| GGNBP2 |
| GGPS1 |
| GGT1 |
| GGT7 |
| GHDC |
| GHITM |
| GHR |
| GIGYF1 |
| GIMAP1 |
| GIN1 |
| GINS1 |
| GINS3 |
| GINS4 |
| GIPC1 |
| GIT1 |
| GIT2 |
| GJA1 |
| GJA5 |
| GJB3 |
| GJC1 |
| GK |
| GK5 |
| GKAP1 |
| GLB1 |
| GLB1L |
| GLCCI1 |
| GLDC |
| GLDN |
| GLG1 |
| GLI4 |
| GLIPR2 |
| GLIS2 |
| GLMN |
| GLO1 |
| GLOD4 |
| GLRX |
| GLRX2 |
| GLRX3 |

| |
|---|
| GLRX5 |
| GLS |
| GLS2 |
| GLT8D1 |
| GLT8D2 |
| GLUD1 |
| GLUL |
| GLYCTK |
| GLYR1 |
| GM2A |
| GMCL1 |
| GMDS |
| GMEB1 |
| GMEB2 |
| GMFB |
| GMFG |
| GMIP |
| GMNN |
| GMPPA |
| GMPR |
| GMPR2 |
| GMPS |
| GNA11 |
| GNA12 |
| GNA13 |
| GNA14 |
| GNAI1 |
| GNAI2 |
| GNAL |
| GNAQ |
| GNAS |
| GNB1 |
| GNB1L |
| GNB2 |
| GNB4 |
| GNB5 |
| GNE |
| GNG10 |
| GNG11 |
| GNG12 |
| GNG3 |
| GNG5 |
| GNG7 |
| GNL1 |
| GNL2 |

| |
|---|
| GNL3 |
| GNPAT |
| GNPDA1 |
| GNPDA2 |
| GNPNAT1 |
| GNPTAB |
| GNPTG |
| GNS |
| GOLGA2 |
| GOLGA3 |
| GOLGA5 |
| GOLGA7 |
| GOLGB1 |
| GOLIM4 |
| GOLM1 |
| GOLPH3 |
| GOLPH3L |
| GOLT1B |
| GON4L |
| GOPC |
| GORAB |
| GORASP2 |
| GOSR1 |
| GOSR2 |
| GOT1 |
| GOT2 |
| GP1BA |
| GPAA1 |
| GPAM |
| GPATCH1 |
| GPATCH3 |
| GPATCH4 |
| GPATCH8 |
| GPBAR1 |
| GPBP1 |
| GPBP1L1 |
| GPC1 |
| GPC2 |
| GPC4 |
| GPC6 |
| GPCPD1 |
| GPD1 |
| GPD1L |
| GPD2 |
| GPHN |

| |
|---|
| GPI |
| GPKOW |
| GPN1 |
| GPN2 |
| GPN3 |
| GPNMB |
| GPR101 |
| GPR108 |
| GPR135 |
| GPR137 |
| GPR137B |
| GPR141 |
| GPR143 |
| GPR146 |
| GPR15 |
| GPR153 |
| GPR155 |
| GPR157 |
| GPR162 |
| GPR173 |
| GPR176 |
| GPR179 |
| GPR18 |
| GPR3 |
| GPR35 |
| GPR4 |
| GPR63 |
| GPR65 |
| GPR68 |
| GPR75 |
| GPR84 |
| GPRASP1 |
| GPRC5A |
| GPRC5B |
| GPRC5C |
| GPRC5D |
| GPRIN3 |
| GPS1 |
| GPSM1 |
| GPSM2 |
| GPT |
| GPT2 |
| GPX1 |
| GPX3 |
| GPX7 |

| |
|---|
| GPX8 |
| GRAMD1A |
| GRAMD1C |
| GRB2 |
| GRB7 |
| GREB1 |
| GREM1 |
| GRHL1 |
| GRHPR |
| GRID2 |
| GRIK5 |
| GRIN2C |
| GRIN3A |
| GRIN3B |
| GRINA |
| GRIP1 |
| GRIPAP1 |
| GRK4 |
| GRK5 |
| GRK6 |
| GRM3 |
| GRN |
| GRPEL1 |
| GRPEL2 |
| GRSF1 |
| GRTP1 |
| GRWD1 |
| GSDMB |
| GSG1 |
| GSK3A |
| GSPT1 |
| GSS |
| GSTA4 |
| GSTCD |
| GSTK1 |
| GSTM3 |
| GSTO1 |
| GSTP1 |
| GSTZ1 |
| GTDC1 |
| GTF2A1 |
| GTF2A2 |
| GTF2B |
| GTF2E1 |
| GTF2E2 |

| |
|---|
| GTF2F1 |
| GTF2F2 |
| GTF2H1 |
| GTF2H2 |
| GTF2H3 |
| GTF2I |
| GTF2IRD1 |
| GTF3A |
| GTF3C1 |
| GTF3C2 |
| GTF3C3 |
| GTF3C4 |
| GTF3C5 |
| GTPBP1 |
| GTPBP2 |
| GTPBP3 |
| GTPBP4 |
| GTPBP8 |
| GUCA2B |
| GUF1 |
| GUK1 |
| GULP1 |
| GUSB |
| GXYLT1 |
| GYG1 |
| GYG2 |
| GYS1 |
| GZF1 |
| H6PD |
| HABP4 |
| HACE1 |
| HACL1 |
| HADH |
| HAGH |
| HAGHL |
| HAPLN1 |
| HAPLN3 |
| HARBI1 |
| HAS3 |
| HAT1 |
| HAUS1 |
| HAUS2 |
| HAUS3 |
| HAUS4 |
| HAUS5 |

| |
|---|
| HAUS6 |
| HAUS7 |
| HAUS8 |
| HAVCR2 |
| HAX1 |
| HBE1 |
| HBEGF |
| HBP1 |
| HBS1L |
| HCCS |
| HCFC1 |
| HCFC1R1 |
| HCFC2 |
| HCN1 |
| HCRT |
| HDAC1 |
| HDAC10 |
| HDAC11 |
| HDAC2 |
| HDAC3 |
| HDAC5 |
| HDAC6 |
| HDAC8 |
| HDC |
| HDDC2 |
| HDDC3 |
| HDGF |
| HDHD2 |
| HDHD3 |
| HDLBP |
| HEATR3 |
| HEATR4 |
| HEATR5B |
| HEATR6 |
| HEBP2 |
| HECTD2 |
| HECTD3 |
| HEG1 |
| HELB |
| HELLS |
| HELQ |
| HELZ |
| HEMK1 |
| HERC1 |
| HERC2 |

| |
|---|
| HERC4 |
| HERC5 |
| HERPUD2 |
| HES4 |
| HES6 |
| HEXA |
| HEXB |
| HEXIM1 |
| HEXIM2 |
| HEY2 |
| HFE |
| HFM1 |
| HGF |
| HGS |
| HHEX |
| HHIPL2 |
| HIBADH |
| HIC2 |
| HIF1A |
| HIF1AN |
| HIGD2A |
| HINFP |
| HINT1 |
| HINT2 |
| HINT3 |
| HIP1R |
| HIPK1 |
| HIPK3 |
| HIRA |
| HIRIP3 |
| HIVEP1 |
| HIVEP2 |
| HIVEP3 |
| HJURP |
| HK1 |
| HK2 |
| HK3 |
| HLCS |
| HLX |
| HM13 |
| HMBS |
| HMCN1 |
| HMG20A |
| HMG20B |
| HMGA1 |

| |
|---|
| HMGB1 |
| HMGB3 |
| HMGCL |
| HMGCR |
| HMGCS1 |
| HMGN1 |
| HMGN2 |
| HMGN3 |
| HMGN4 |
| HMGN5 |
| HMGXB4 |
| HMMR |
| HMOX1 |
| HMOX2 |
| HNRNPA0 |
| HNRNPA1 |
| HNRNPA3 |
| HNRNPAB |
| HNRNPC |
| HNRNPD |
| HNRNPF |
| HNRNPH1 |
| HNRNPH2 |
| HNRNPH3 |
| HNRNPK |
| HNRNPL |
| HNRNPM |
| HNRNPR |
| HNRNPU |
| HNRNPUL1 |
| HNRNPUL2 |
| HOMER1 |
| HOMER2 |
| HOMEZ |
| HOOK1 |
| HOOK2 |
| HOOK3 |
| HOXA10 |
| HOXC4 |
| HOXC5 |
| HOXC6 |
| HP1BP3 |
| HPDL |
| HPS1 |
| HPS3 |

| |
|---|
| HPS4 |
| HPS6 |
| HPSE |
| HRH1 |
| HS1BP3 |
| HS2ST1 |
| HS3ST1 |
| HS3ST2 |
| HS3ST3A1 |
| HS6ST1 |
| HS6ST3 |
| HSD11B1L |
| HSD11B2 |
| HSD17B1 |
| HSD17B10 |
| HSD17B14 |
| HSD17B3 |
| HSD17B4 |
| HSD17B7 |
| HSD17B8 |
| HSD3B7 |
| HSDL1 |
| HSDL2 |
| HSF1 |
| HSF2 |
| HSF2BP |
| HSF4 |
| HSP90AA1 |
| HSP90AB1 |
| HSP90B1 |
| HSPA12B |
| HSPA13 |
| HSPA14 |
| HSPA1A |
| HSPA1L |
| HSPA4L |
| HSPA5 |
| HSPA6 |
| HSPA8 |
| HSPA9 |
| HSPB1 |
| HSPB6 |
| HSPBAP1 |
| HSPBP1 |
| HSPD1 |

| |
|---|
| HSPE1 |
| HSPG2 |
| HTATIP2 |
| HTATSF1 |
| HTR1F |
| HTR7 |
| HTRA1 |
| HTRA2 |
| HUS1 |
| HUWE1 |
| HVCN1 |
| HYAL1 |
| HYAL2 |
| HYLS1 |
| IAH1 |
| IARS2 |
| IBTK |
| ICAM1 |
| ICOSLG |
| ID1 |
| ID2 |
| ID3 |
| IDE |
| IDH1 |
| IDH2 |
| IDH3A |
| IDH3B |
| IDI1 |
| IDO1 |
| IDS |
| IDUA |
| IER2 |
| IER3 |
| IER3IP1 |
| IER5 |
| IER5L |
| IFFO2 |
| IFI16 |
| IFI27L2 |
| IFI30 |
| IFI35 |
| IFI6 |
| IFIH1 |
| IFIT1 |
| IFIT2 |

| |
|---|
| IFIT5 |
| IFITM3 |
| IFNAR1 |
| IFNAR2 |
| IFNGR1 |
| IFNGR2 |
| IFRD1 |
| IFRD2 |
| IFT122 |
| IFT20 |
| IFT52 |
| IFT57 |
| IFT74 |
| IFT80 |
| IFT81 |
| IFT88 |
| IGBP1 |
| IGDCC4 |
| IGF1R |
| IGF2 |
| IGF2BP2 |
| IGFBP2 |
| IGFBP3 |
| IGFBP4 |
| IGFBP5 |
| IGFBP6 |
| IGFBP7 |
| IGHMBP2 |
| IGSF11 |
| IGSF6 |
| IGSF8 |
| IGSF9 |
| IHH |
| IK |
| IKBKB |
| IKBKE |
| IKBKG |
| IKZF2 |
| IKZF3 |
| IKZF4 |
| IKZF5 |
| IL10 |
| IL10RA |
| IL10RB |
| IL11RA |

| |
|---|
| IL13RA1 |
| IL15 |
| IL16 |
| IL17D |
| IL17RA |
| IL17RC |
| IL17RD |
| IL18 |
| IL18BP |
| IL1A |
| IL1RAP |
| IL1RN |
| IL27RA |
| IL2RB |
| IL34 |
| IL4R |
| IL6ST |
| ILF2 |
| ILF3 |
| ILK |
| ILKAP |
| ILVBL |
| IMMT |
| IMP3 |
| IMP4 |
| IMPA1 |
| IMPA2 |
| IMPACT |
| IMPDH1 |
| IMPDH2 |
| INCENP |
| ING1 |
| ING3 |
| ING4 |
| ING5 |
| INHA |
| INHBA |
| INHBB |
| INO80 |
| INO80B |
| INO80C |
| INO80D |
| INO80E |
| INPP1 |
| INPP4A |

| |
|---|
| INPP5A |
| INPP5B |
| INPP5D |
| INPP5E |
| INPP5F |
| INPP5K |
| INPPL1 |
| INSIG1 |
| INSIG2 |
| INSR |
| INTS10 |
| INTS12 |
| INTS2 |
| INTS3 |
| INTS5 |
| INTS6 |
| INTS7 |
| INTS8 |
| INTS9 |
| INTU |
| IP6K1 |
| IPMK |
| IPO13 |
| IPO4 |
| IPO7 |
| IPO8 |
| IPP |
| IPPK |
| IQCB1 |
| IQCD |
| IQCE |
| IQCG |
| IQGAP1 |
| IQGAP2 |
| IQGAP3 |
| IQSEC2 |
| IRAK1 |
| IRAK1BP1 |
| IRAK2 |
| IRAK3 |
| IRAK4 |
| IREB2 |
| IRF1 |
| IRF2 |
| IRF2BP2 |

| |
|---|
| IRF3 |
| IRF7 |
| IRS1 |
| IRX3 |
| IRX5 |
| ISCU |
| ISG20 |
| ISG20L2 |
| ISOC1 |
| IST1 |
| ISY1 |
| ISYNA1 |
| ITFG1 |
| ITFG2 |
| ITGA3 |
| ITGA4 |
| ITGA6 |
| ITGAE |
| ITGAL |
| ITGAV |
| ITGB1 |
| ITGB1BP1 |
| ITGB1BP2 |
| ITGB3 |
| ITGB3BP |
| ITGB5 |
| ITGB7 |
| ITGB8 |
| ITGBL1 |
| ITM2B |
| ITM2C |
| ITPA |
| ITPK1 |
| ITPKA |
| ITPKB |
| ITPKC |
| ITPR2 |
| ITPR3 |
| ITPRIP |
| ITPRIPL2 |
| ITSN1 |
| IVD |
| IVNS1ABP |
| IWS1 |
| JAG1 |

| |
|---|
| JAGN1 |
| JAK1 |
| JAKMIP2 |
| JAM3 |
| JARID2 |
| JAZF1 |
| JDP2 |
| JKAMP |
| JMJD1C |
| JMJD4 |
| JMJD6 |
| JMJD7 |
| JMJD8 |
| JMY |
| JOSD1 |
| JRK |
| JRKL |
| JSRP1 |
| JTB |
| JUN |
| JUNB |
| JUND |
| JUP |
| KALRN |
| KANK1 |
| KANK2 |
| KANK3 |
| KAT2B |
| KAT5 |
| KATNA1 |
| KATNAL1 |
| KATNB1 |
| KAZALD1 |
| KBTBD2 |
| KBTBD3 |
| KBTBD6 |
| KBTBD7 |
| KCMF1 |
| KCNA4 |
| KCNAB1 |
| KCNAB2 |
| KCNC2 |
| KCND2 |
| KCNE2 |
| KCNE3 |

| |
|---|
| KCNE4 |
| KCNG2 |
| KCNH2 |
| KCNIP4 |
| KCNJ10 |
| KCNJ14 |
| KCNJ15 |
| KCNJ2 |
| KCNJ5 |
| KCNK1 |
| KCNK3 |
| KCNK5 |
| KCNN3 |
| KCNQ1 |
| KCNQ4 |
| KCNQ5 |
| KCNRG |
| KCNS3 |
| KCTD1 |
| KCTD10 |
| KCTD11 |
| KCTD12 |
| KCTD15 |
| KCTD18 |
| KCTD20 |
| KCTD21 |
| KCTD3 |
| KCTD5 |
| KCTD7 |
| KCTD9 |
| KDELR1 |
| KDELR2 |
| KDELR3 |
| KDM1A |
| KDM1B |
| KDM2A |
| KDM2B |
| KDM3A |
| KDM4B |
| KDM4C |
| KDM5A |
| KDM5B |
| KDM5C |
| KDM6A |
| KDSR |

| |
|---|
| KEAP1 |
| KHDRBS1 |
| KHNYN |
| KIAA0586 |
| KIAA1549 |
| KIAA2013 |
| KIF11 |
| KIF13A |
| KIF13B |
| KIF14 |
| KIF15 |
| KIF16B |
| KIF18A |
| KIF18B |
| KIF1C |
| KIF20A |
| KIF20B |
| KIF22 |
| KIF23 |
| KIF24 |
| KIF2A |
| KIF2C |
| KIF3A |
| KIF3B |
| KIF3C |
| KIF4A |
| KIF7 |
| KIFAP3 |
| KIFC1 |
| KIN |
| KIRREL3 |
| KIT |
| KITLG |
| KLC1 |
| KLC2 |
| KLC3 |
| KLC4 |
| KLF11 |
| KLF12 |
| KLF13 |
| KLF15 |
| KLF16 |
| KLF4 |
| KLF5 |
| KLF6 |

| |
|---|
| KLF7 |
| KLF8 |
| KLHDC1 |
| KLHDC10 |
| KLHDC2 |
| KLHDC4 |
| KLHDC8B |
| KLHL11 |
| KLHL12 |
| KLHL15 |
| KLHL17 |
| KLHL18 |
| KLHL2 |
| KLHL20 |
| KLHL21 |
| KLHL23 |
| KLHL24 |
| KLHL25 |
| KLHL26 |
| KLHL28 |
| KLHL3 |
| KLHL36 |
| KLHL5 |
| KLHL7 |
| KLHL8 |
| KLHL9 |
| KLK14 |
| KLK15 |
| KPNA3 |
| KPNA4 |
| KPNA5 |
| KPNA6 |
| KPNB1 |
| KRAS |
| KRBA1 |
| KRBA2 |
| KRCC1 |
| KRI1 |
| KRIT1 |
| KRR1 |
| KRT18 |
| KRT19 |
| KRT20 |
| KRT8 |
| KRTCAP2 |

| |
|---|
| KSR1 |
| KTI12 |
| KTN1 |
| L1CAM |
| L2HGDH |
| L3MBTL2 |
| L3MBTL3 |
| LACTB |
| LACTB2 |
| LAD1 |
| LAG3 |
| LAGE3 |
| LAMA1 |
| LAMA5 |
| LAMB1 |
| LAMB2 |
| LAMB3 |
| LAMC1 |
| LAMC3 |
| LAMP1 |
| LAMP2 |
| LANCL2 |
| LAP3 |
| LAPTM4A |
| LAPTM4B |
| LAPTM5 |
| LARP1 |
| LARP4 |
| LARP4B |
| LARP6 |
| LARP7 |
| LARS2 |
| LAS1L |
| LASP1 |
| LAT |
| LATS2 |
| LBH |
| LBR |
| LBX2 |
| LCLAT1 |
| LCMT1 |
| LCMT2 |
| LCN2 |
| LCOR |
| LCORL |

| |
|---|
| LCP1 |
| LDB1 |
| LDHA |
| LDHB |
| LDHD |
| LDLR |
| LDLRAD2 |
| LDLRAD3 |
| LDLRAP1 |
| LDOC1 |
| LEMD2 |
| LEMD3 |
| LENG1 |
| LENG8 |
| LENG9 |
| LEO1 |
| LEPROT |
| LEPROTL1 |
| LETM1 |
| LETM2 |
| LETMD1 |
| LFNG |
| LGALS3 |
| LGALS3BP |
| LGALS8 |
| LGALS9 |
| LGMN |
| LGR4 |
| LGSN |
| LHFPL2 |
| LHPP |
| LIAS |
| LIF |
| LIFR |
| LIG1 |
| LIG3 |
| LIG4 |
| LIMA1 |
| LIMD1 |
| LIMD2 |
| LIMK1 |
| LIMK2 |
| LIN37 |
| LIN52 |
| LIN54 |

| |
|---|
| LIN7B |
| LIN7C |
| LINGO2 |
| LINS1 |
| LIPA |
| LIPC |
| LIPE |
| LIPH |
| LIPK |
| LIPT1 |
| LIPT2 |
| LITAF |
| LIX1L |
| LLGL2 |
| LLPH |
| LMAN1 |
| LMAN2 |
| LMAN2L |
| LMBRD1 |
| LMBRD2 |
| LMCD1 |
| LMF1 |
| LMF2 |
| LMNA |
| LMNB1 |
| LMNB2 |
| LMO4 |
| LMO7 |
| LMTK2 |
| LNPEP |
| LNX2 |
| LONP1 |
| LONP2 |
| LONRF1 |
| LOXL3 |
| LOXL4 |
| LPAR2 |
| LPAR6 |
| LPCAT3 |
| LPCAT4 |
| LPGAT1 |
| LPIN1 |
| LPIN3 |
| LPP |
| LPXN |

| |
|---|
| LRBA |
| LRCH2 |
| LRCH3 |
| LRCH4 |
| LRFN3 |
| LRFN4 |
| LRIG1 |
| LRIG2 |
| LRIG3 |
| LRP10 |
| LRP12 |
| LRP4 |
| LRP5 |
| LRP6 |
| LRP8 |
| LRPAP1 |
| LRPPRC |
| LRRC1 |
| LRRC10B |
| LRRC14 |
| LRRC17 |
| LRRC20 |
| LRRC24 |
| LRRC25 |
| LRRC27 |
| LRRC28 |
| LRRC31 |
| LRRC4 |
| LRRC40 |
| LRRC41 |
| LRRC42 |
| LRRC45 |
| LRRC46 |
| LRRC47 |
| LRRC49 |
| LRRC4B |
| LRRC4C |
| LRRC57 |
| LRRC58 |
| LRRC59 |
| LRRC66 |
| LRRC8A |
| LRRC8D |
| LRRCC1 |
| LRRFIP1 |

| |
|---|
| LRRFIP2 |
| LRRK1 |
| LRRK2 |
| LRRN2 |
| LRRN3 |
| LRRTM2 |
| LRSAM1 |
| LRWD1 |
| LSG1 |
| LSM1 |
| LSM10 |
| LSM11 |
| LSM12 |
| LSM14A |
| LSM14B |
| LSM3 |
| LSM4 |
| LSM5 |
| LSM6 |
| LSM7 |
| LSR |
| LSS |
| LTA4H |
| LTB |
| LTB4R |
| LTBP1 |
| LTBP3 |
| LTBR |
| LTV1 |
| LUC7L |
| LUC7L2 |
| LUC7L3 |
| LUZP1 |
| LXN |
| LY6G5B |
| LY75 |
| LY9 |
| LYAR |
| LYN |
| LYNX1 |
| LYPLA1 |
| LYPLA2 |
| LYRM1 |
| LYRM2 |
| LYRM4 |

| |
|---|
| LYRM7 |
| LYSMD1 |
| LYSMD2 |
| LYSMD3 |
| LYST |
| LYZ |
| LZTFL1 |
| LZTR1 |
| LZTS1 |
| LZTS2 |
| M6PR |
| MACROD1 |
| MAD1L1 |
| MAD2L1 |
| MAD2L1BP |
| MAD2L2 |
| MADD |
| MAEA |
| MAF |
| MAF1 |
| MAFB |
| MAFK |
| MAGEA10 |
| MAGED1 |
| MAGED2 |
| MAGEL2 |
| MAGI3 |
| MAGIX |
| MAGOH |
| MAGOHB |
| MAK16 |
| MAL2 |
| MAMDC4 |
| MAML2 |
| MAML3 |
| MAMLD1 |
| MAMSTR |
| MAN1A1 |
| MAN1A2 |
| MAN1C1 |
| MAN2A1 |
| MAN2A2 |
| MAN2B1 |
| MAN2C1 |
| MANBA |

| |
|---|
| MANBAL |
| MANEA |
| MANEAL |
| MANF |
| MAOA |
| MAP1B |
| MAP1LC3B |
| MAP2 |
| MAP2K1 |
| MAP2K2 |
| MAP2K3 |
| MAP2K5 |
| MAP2K6 |
| MAP2K7 |
| MAP3K1 |
| MAP3K10 |
| MAP3K11 |
| MAP3K12 |
| MAP3K14 |
| MAP3K15 |
| MAP3K2 |
| MAP3K3 |
| MAP3K4 |
| MAP3K5 |
| MAP3K6 |
| MAP3K7 |
| MAP3K8 |
| MAP4 |
| MAP4K1 |
| MAP4K2 |
| MAP4K3 |
| MAP4K5 |
| MAP7 |
| MAP7D1 |
| MAP7D3 |
| MAP9 |
| MAPK11 |
| MAPK12 |
| MAPK13 |
| MAPK14 |
| MAPK15 |
| MAPK4 |
| MAPK6 |
| MAPK7 |
| MAPK8 |

| |
|---|
| MAPK8IP1 |
| MAPK8IP3 |
| MAPKAP1 |
| MAPKAPK3 |
| MAPKBP1 |
| MAPRE2 |
| MARCKSL1 |
| MARK1 |
| MARK2 |
| MARK3 |
| MARK4 |
| MARS2 |
| MARVELD1 |
| MARVELD2 |
| MARVELD3 |
| MASP2 |
| MAST1 |
| MAST3 |
| MASTL |
| MAT2B |
| MATK |
| MATN4 |
| MATR3 |
| MAVS |
| MAX |
| MBD1 |
| MBD3 |
| MBD4 |
| MBD5 |
| MBIP |
| MBLAC1 |
| MBLAC2 |
| MBNL2 |
| MBNL3 |
| MBOAT2 |
| MBTD1 |
| MBTPS1 |
| MBTPS2 |
| MCAM |
| MCAT |
| MCC |
| MCCC1 |
| MCEE |
| MCF2L |
| MCF2L2 |

| |
|---|
| MCFD2 |
| MCL1 |
| MCM10 |
| MCM2 |
| MCM3 |
| MCM3AP |
| MCM4 |
| MCM5 |
| MCM6 |
| MCM7 |
| MCM8 |
| MCM9 |
| MCOLN1 |
| MCPH1 |
| MCRS1 |
| MCTP1 |
| MCTS1 |
| MDC1 |
| MDFIC |
| MDH2 |
| MDK |
| MDM2 |
| MDM4 |
| MDN1 |
| ME1 |
| ME2 |
| ME3 |
| MEA1 |
| MEAF6 |
| MECOM |
| MECP2 |
| MECR |
| MED1 |
| MED10 |
| MED11 |
| MED12 |
| MED12L |
| MED13 |
| MED13L |
| MED15 |
| MED17 |
| MED18 |
| MED19 |
| MED21 |
| MED22 |

| |
|---|
| MED23 |
| MED24 |
| MED25 |
| MED26 |
| MED27 |
| MED28 |
| MED29 |
| MED30 |
| MED31 |
| MED4 |
| MED7 |
| MED9 |
| MEF2A |
| MEF2C |
| MEFV |
| MEGF9 |
| MEI1 |
| MEIS1 |
| MEIS2 |
| MELK |
| MEN1 |
| MEP1B |
| MEPCE |
| MERTK |
| MESP2 |
| MEST |
| METRN |
| METRNL |
| METTL1 |
| METTL14 |
| METTL3 |
| METTL4 |
| METTL5 |
| METTL6 |
| METTL7A |
| METTL7B |
| METTL8 |
| METTL9 |
| MEX3A |
| MEX3B |
| MEX3C |
| MFAP1 |
| MFAP2 |
| MFAP3 |
| MFAP3L |

| |
|---|
| MFGE8 |
| MFN1 |
| MFSD1 |
| MFSD10 |
| MFSD11 |
| MFSD3 |
| MFSD6 |
| MFSD8 |
| MFSD9 |
| MGA |
| MGAT1 |
| MGAT2 |
| MGAT4A |
| MGAT4B |
| MGAT5 |
| MGMT |
| MGRN1 |
| MGST1 |
| MGST2 |
| MGST3 |
| MIA3 |
| MIB1 |
| MIB2 |
| MICAL1 |
| MICAL3 |
| MICALL1 |
| MICALL2 |
| MICB |
| MID1 |
| MID1IP1 |
| MIDN |
| MIER1 |
| MIER2 |
| MIER3 |
| MIF |
| MIF4GD |
| MIIP |
| MINK1 |
| MINPP1 |
| MIOS |
| MIPEP |
| MIPOL1 |
| MIS12 |
| MIS18A |
| MITD1 |

| |
|---|
| MITF |
| MKI67 |
| MKKS |
| MKNK1 |
| MKNK2 |
| MKRN2 |
| MKS1 |
| MKX |
| MLEC |
| MLF1 |
| MLH1 |
| MLH3 |
| MLKL |
| MLLT1 |
| MLLT3 |
| MLLT6 |
| MLST8 |
| MLXIP |
| MLYCD |
| MMAA |
| MMAB |
| MMACHC |
| MMADHC |
| MMD |
| MMGT1 |
| MMP14 |
| MMP15 |
| MMP17 |
| MMP19 |
| MMP2 |
| MMP25 |
| MMP9 |
| MMS19 |
| MN1 |
| MNAT1 |
| MND1 |
| MNS1 |
| MNT |
| MOCOS |
| MOCS1 |
| MOCS2 |
| MOCS3 |
| MON1A |
| MON1B |
| MORC2 |

| |
|---|
| MORC3 |
| MORF4L1 |
| MORF4L2 |
| MORN4 |
| MOSPD1 |
| MOSPD3 |
| MOV10 |
| MPDU1 |
| MPDZ |
| MPG |
| MPHOSPH10 |
| MPHOSPH6 |
| MPHOSPH8 |
| MPHOSPH9 |
| MPI |
| MPL |
| MPND |
| MPP1 |
| MPP2 |
| MPP7 |
| MPPE1 |
| MPPED2 |
| MPRIP |
| MPST |
| MPZL3 |
| MR1 |
| MRAP |
| MRAP2 |
| MRAS |
| MRC2 |
| MRFAP1 |
| MRI1 |
| MRM1 |
| MRO |
| MRPL1 |
| MRPL10 |
| MRPL11 |
| MRPL12 |
| MRPL13 |
| MRPL14 |
| MRPL15 |
| MRPL16 |
| MRPL17 |
| MRPL18 |
| MRPL19 |

| |
|---|
| MRPL20 |
| MRPL21 |
| MRPL22 |
| MRPL23 |
| MRPL24 |
| MRPL27 |
| MRPL28 |
| MRPL3 |
| MRPL30 |
| MRPL32 |
| MRPL33 |
| MRPL34 |
| MRPL35 |
| MRPL36 |
| MRPL37 |
| MRPL38 |
| MRPL39 |
| MRPL4 |
| MRPL41 |
| MRPL42 |
| MRPL43 |
| MRPL44 |
| MRPL45 |
| MRPL46 |
| MRPL47 |
| MRPL48 |
| MRPL49 |
| MRPL51 |
| MRPL52 |
| MRPL53 |
| MRPL54 |
| MRPL55 |
| MRPL9 |
| MRPS10 |
| MRPS11 |
| MRPS15 |
| MRPS16 |
| MRPS17 |
| MRPS18B |
| MRPS18C |
| MRPS2 |
| MRPS21 |
| MRPS22 |
| MRPS23 |
| MRPS24 |

| |
|---|
| MRPS25 |
| MRPS26 |
| MRPS27 |
| MRPS28 |
| MRPS30 |
| MRPS31 |
| MRPS33 |
| MRPS34 |
| MRPS35 |
| MRPS5 |
| MRPS6 |
| MRPS7 |
| MRPS9 |
| MRS2 |
| MSH2 |
| MSH4 |
| MSH5 |
| MSH6 |
| MSI1 |
| MSI2 |
| MSL1 |
| MSL2 |
| MSL3 |
| MSN |
| MSR1 |
| MSRA |
| MSRB2 |
| MSRB3 |
| MST1 |
| MST1R |
| MSTO1 |
| MT1A |
| MT2A |
| MT3 |
| MTA2 |
| MTA3 |
| MTBP |
| MTCH1 |
| MTCH2 |
| MTCP1 |
| MTDH |
| MTF1 |
| MTF2 |
| MTFMT |
| MTFR1 |

| |
|---|
| MTG1 |
| MTHFD1L |
| MTHFD2 |
| MTHFD2L |
| MTHFSD |
| MTIF2 |
| MTIF3 |
| MTM1 |
| MTMR1 |
| MTMR10 |
| MTMR11 |
| MTMR14 |
| MTMR2 |
| MTMR3 |
| MTMR4 |
| MTMR6 |
| MTMR7 |
| MTMR9 |
| MTO1 |
| MTOR |
| MTPAP |
| MTPN |
| MTR |
| MTRF1 |
| MTRF1L |
| MTRR |
| MTSS1 |
| MTTP |
| MTUS1 |
| MTX1 |
| MTX2 |
| MTX3 |
| MUC1 |
| MUC20 |
| MUL1 |
| MUS81 |
| MVD |
| MVK |
| MVP |
| MX1 |
| MXD3 |
| MXD4 |
| MXI1 |
| MXRA7 |
| MXRA8 |

| |
|---|
| MYADM |
| MYBBP1A |
| MYBL1 |
| MYBL2 |
| MYBPC1 |
| MYC |
| MYCBP |
| MYCBP2 |
| MYCBPAP |
| MYCN |
| MYD88 |
| MYEF2 |
| MYH10 |
| MYH11 |
| MYH14 |
| MYH9 |
| MYL12B |
| MYL6 |
| MYL6B |
| MYL9 |
| MYLIP |
| MYLK |
| MYLPF |
| MYO10 |
| MYO18A |
| MYO19 |
| MYO1A |
| MYO1B |
| MYO1C |
| MYO1D |
| MYO1E |
| MYO1F |
| MYO1G |
| MYO3B |
| MYO5A |
| MYO5C |
| MYO6 |
| MYO7A |
| MYO9A |
| MYO9B |
| MYOC |
| MYOCD |
| MYOF |
| MYOM1 |
| MYOZ1 |

| |
|---|
| MYPN |
| MYSM1 |
| MZF1 |
| N4BP1 |
| N4BP2 |
| N4BP2L1 |
| N6AMT1 |
| NAA10 |
| NAA15 |
| NAA16 |
| NAA20 |
| NAA25 |
| NAA30 |
| NAA35 |
| NAA38 |
| NAA40 |
| NAAA |
| NAALADL1 |
| NAB1 |
| NAB2 |
| NACA |
| NACAD |
| NACC1 |
| NACC2 |
| NADK |
| NAE1 |
| NAGA |
| NAGK |
| NAGLU |
| NAGPA |
| NAIF1 |
| NAIP |
| NAMPT |
| NANOS1 |
| NANP |
| NANS |
| NAP1L1 |
| NAP1L5 |
| NAPA |
| NAPB |
| NAPEPLD |
| NAPG |
| NAPSA |
| NARF |
| NASP |

| |
|---|
| NAT1 |
| NAT10 |
| NAT14 |
| NAT9 |
| NAV1 |
| NAV2 |
| NAV3 |
| NBAS |
| NBEA |
| NBEAL2 |
| NBL1 |
| NBN |
| NBR1 |
| NCAM1 |
| NCAM2 |
| NCAPD2 |
| NCAPD3 |
| NCAPG |
| NCAPG2 |
| NCAPH |
| NCAPH2 |
| NCBP1 |
| NCBP2 |
| NCDN |
| NCEH1 |
| NCF2 |
| NCK1 |
| NCK2 |
| NCKAP1L |
| NCKAP5 |
| NCKAP5L |
| NCKIPSD |
| NCL |
| NCLN |
| NCOA1 |
| NCOA2 |
| NCOA3 |
| NCOA4 |
| NCOA5 |
| NCOA6 |
| NCOA7 |
| NCOR1 |
| NCS1 |
| NCSTN |
| NDC80 |

| |
|---|
| NDE1 |
| NDEL1 |
| NDFIP2 |
| NDOR1 |
| NDRG1 |
| NDRG2 |
| NDST2 |
| NDUFA10 |
| NDUFA12 |
| NDUFA13 |
| NDUFA2 |
| NDUFA3 |
| NDUFA4 |
| NDUFA6 |
| NDUFA8 |
| NDUFAB1 |
| NDUFAF1 |
| NDUFAF3 |
| NDUFAF4 |
| NDUFB1 |
| NDUFB10 |
| NDUFB11 |
| NDUFB2 |
| NDUFB4 |
| NDUFB5 |
| NDUFB6 |
| NDUFB7 |
| NDUFB8 |
| NDUFB9 |
| NDUFC1 |
| NDUFC2 |
| NDUFS2 |
| NDUFS3 |
| NDUFS5 |
| NDUFS6 |
| NDUFS7 |
| NDUFV1 |
| NDUFV2 |
| NDUFV3 |
| NEBL |
| NECAP1 |
| NECAP2 |
| NEDD4 |
| NEDD8 |
| NEDD9 |

| |
|---|
| NEFH |
| NEFL |
| NEGR1 |
| NEIL1 |
| NEIL2 |
| NEIL3 |
| NEK10 |
| NEK2 |
| NEK3 |
| NEK4 |
| NEK6 |
| NEK7 |
| NEK8 |
| NEK9 |
| NELL2 |
| NENF |
| NES |
| NET1 |
| NEU1 |
| NEU3 |
| NEURL1B |
| NEURL2 |
| NEURL4 |
| NEXN |
| NF1 |
| NFAM1 |
| NFAT5 |
| NFATC1 |
| NFATC3 |
| NFE2 |
| NFE2L1 |
| NFE2L3 |
| NFIC |
| NFIL3 |
| NFIX |
| NFKB1 |
| NFKBIA |
| NFKBIB |
| NFKBID |
| NFKBIE |
| NFKBIL1 |
| NFKBIZ |
| NFS1 |
| NFX1 |
| NFYA |

| |
|---|
| NFYB |
| NFYC |
| NGDN |
| NGLY1 |
| NGRN |
| NHEJ1 |
| NHLRC1 |
| NHLRC2 |
| NHLRC3 |
| NHP2 |
| NHSL2 |
| NICN1 |
| NID1 |
| NID2 |
| NIF3L1 |
| NIN |
| NINJ1 |
| NIP7 |
| NIPA1 |
| NIPA2 |
| NIPAL1 |
| NIPAL3 |
| NIPBL |
| NIPSNAP3A |
| NIT1 |
| NIT2 |
| NKAIN1 |
| NKAP |
| NKD1 |
| NKG7 |
| NKIRAS1 |
| NKIRAS2 |
| NKPD1 |
| NKRF |
| NKX1-2 |
| NLE1 |
| NLGN2 |
| NLGN3 |
| NLK |
| NLN |
| NLRC3 |
| NLRC4 |
| NLRP6 |
| NMB |
| NMD3 |

| |
|---|
| NME3 |
| NME6 |
| NME7 |
| NMI |
| NMNAT1 |
| NMRAL1 |
| NMT1 |
| NNAT |
| NNT |
| NOB1 |
| NOC2L |
| NOC3L |
| NOC4L |
| NOD1 |
| NODAL |
| NOL10 |
| NOL11 |
| NOL12 |
| NOL6 |
| NOL7 |
| NOL8 |
| NOL9 |
| NONO |
| NOP14 |
| NOP16 |
| NOP2 |
| NOP56 |
| NOP58 |
| NOS1AP |
| NOSIP |
| NOTCH1 |
| NOTCH2 |
| NOVA1 |
| NOXA1 |
| NPAS3 |
| NPAT |
| NPB |
| NPC2 |
| NPDC1 |
| NPEPL1 |
| NPEPPS |
| NPHP1 |
| NPHP3 |
| NPHP4 |
| NPL |

| |
|---|
| NPLOC4 |
| NPM1 |
| NPM3 |
| NPNT |
| NPR2 |
| NPR3 |
| NPTN |
| NPTX1 |
| NPY1R |
| NQO1 |
| NQO2 |
| NR1D2 |
| NR1H2 |
| NR2C2 |
| NR2C2AP |
| NR2F1 |
| NR2F2 |
| NR3C2 |
| NR4A1 |
| NR4A2 |
| NR4A3 |
| NR5A2 |
| NR6A1 |
| NRBF2 |
| NRBP1 |
| NRBP2 |
| NRF1 |
| NRIP1 |
| NRIP3 |
| NRM |
| NRP1 |
| NRSN2 |
| NRXN1 |
| NSA2 |
| NSD1 |
| NSDHL |
| NSF |
| NSFL1C |
| NSL1 |
| NSMAF |
| NSMCE1 |
| NSMCE2 |
| NSMCE4A |
| NSUN3 |
| NSUN4 |

| |
|---|
| NSUN5 |
| NSUN6 |
| NT5C |
| NT5C2 |
| NT5DC1 |
| NT5DC3 |
| NT5E |
| NT5M |
| NTAN1 |
| NTHL1 |
| NTN3 |
| NTN4 |
| NTN5 |
| NTRK1 |
| NUAK1 |
| NUAK2 |
| NUB1 |
| NUBP1 |
| NUBP2 |
| NUBPL |
| NUCB1 |
| NUCB2 |
| NUCKS1 |
| NUDC |
| NUDCD1 |
| NUDCD2 |
| NUDCD3 |
| NUDT1 |
| NUDT12 |
| NUDT13 |
| NUDT14 |
| NUDT15 |
| NUDT16 |
| NUDT16L1 |
| NUDT17 |
| NUDT18 |
| NUDT2 |
| NUDT21 |
| NUDT22 |
| NUDT3 |
| NUDT5 |
| NUDT6 |
| NUDT7 |
| NUDT9 |
| NUF2 |

| |
|---|
| NUFIP1 |
| NUFIP2 |
| NUMA1 |
| NUMB |
| NUMBL |
| NUP133 |
| NUP153 |
| NUP155 |
| NUP160 |
| NUP188 |
| NUP205 |
| NUP210 |
| NUP214 |
| NUP35 |
| NUP43 |
| NUP50 |
| NUP54 |
| NUP62 |
| NUP85 |
| NUP88 |
| NUP93 |
| NUP98 |
| NUPR1 |
| NUS1 |
| NUSAP1 |
| NVL |
| NXF1 |
| NXPH1 |
| NXT1 |
| NXT2 |
| NYNRIN |
| OAF |
| OASL |
| OAT |
| OAZ1 |
| OBSL1 |
| OCEL1 |
| OCIAD1 |
| OCIAD2 |
| OCLN |
| OCRL |
| ODC1 |
| ODF2 |
| ODF2L |
| OFD1 |

| |
|---|
| OGFOD1 |
| OGFR |
| OGFRL1 |
| OGG1 |
| OIP5 |
| OIT3 |
| OLA1 |
| OLFM1 |
| OLFM4 |
| OLFML1 |
| OLFML2A |
| OLFML2B |
| OLR1 |
| OMA1 |
| ONECUT1 |
| OPA1 |
| OPA3 |
| OPHN1 |
| OPRK1 |
| OPTN |
| ORAI1 |
| ORAI3 |
| ORMDL2 |
| ORMDL3 |
| OS9 |
| OSBP |
| OSBP2 |
| OSBPL10 |
| OSBPL11 |
| OSBPL2 |
| OSBPL3 |
| OSBPL5 |
| OSBPL8 |
| OSBPL9 |
| OSCP1 |
| OSGEP |
| OSGEPL1 |
| OSGIN1 |
| OSR2 |
| OSTC |
| OSTF1 |
| OSTM1 |
| OTUB1 |
| OTUD1 |
| OTUD3 |

| |
|---|
| OTUD4 |
| OTUD5 |
| OTUD6B |
| OVGP1 |
| OVOL1 |
| OXA1L |
| OXCT1 |
| OXNAD1 |
| OXR1 |
| OXSM |
| OXSR1 |
| P2RX4 |
| P2RY1 |
| P2RY4 |
| P4HA1 |
| P4HA2 |
| P4HA3 |
| P4HB |
| P4HTM |
| PA2G4 |
| PAAF1 |
| PABPC1 |
| PABPC4 |
| PABPC5 |
| PABPN1 |
| PACRGL |
| PACSIN2 |
| PACSIN3 |
| PADI1 |
| PADI2 |
| PAF1 |
| PAFAH1B1 |
| PAFAH1B2 |
| PAFAH1B3 |
| PAFAH2 |
| PAG1 |
| PAIP1 |
| PAIP2 |
| PAK1 |
| PAK1IP1 |
| PAK4 |
| PALB2 |
| PALLD |
| PALMD |
| PAM |

| |
|---|
| PAN2 |
| PAN3 |
| PANK3 |
| PANK4 |
| PANX1 |
| PANX2 |
| PAOX |
| PAPLN |
| PAPOLA |
| PAPOLG |
| PAPSS1 |
| PAPSS2 |
| PAQR3 |
| PAQR4 |
| PAQR5 |
| PAQR6 |
| PAQR8 |
| PARD3 |
| PARD6A |
| PARD6B |
| PARD6G |
| PARG |
| PARK7 |
| PARL |
| PARN |
| PARP1 |
| PARP11 |
| PARP12 |
| PARP14 |
| PARP16 |
| PARP6 |
| PARP9 |
| PARS2 |
| PARVA |
| PASK |
| PATL1 |
| PATZ1 |
| PAWR |
| PAXIP1 |
| PBK |
| PBLD |
| PBX1 |
| PBX2 |
| PBX3 |
| PBX4 |

| |
|---|
| PBXIP1 |
| PC |
| PCBD1 |
| PCBP1 |
| PCBP2 |
| PCBP4 |
| PCCA |
| PCCB |
| PCDH1 |
| PCDH10 |
| PCDH18 |
| PCDH7 |
| PCDH9 |
| PCDHA3 |
| PCDHA4 |
| PCDHB11 |
| PCDHB14 |
| PCDHB15 |
| PCDHB4 |
| PCDHB6 |
| PCDHB7 |
| PCDHGA5 |
| PCF11 |
| PCGF1 |
| PCGF2 |
| PCGF3 |
| PCGF5 |
| PCGF6 |
| PCID2 |
| PCIF1 |
| PCK2 |
| PCM1 |
| PCMT1 |
| PCMTD1 |
| PCMTD2 |
| PCNA |
| PCNP |
| PCOLCE |
| PCOLCE2 |
| PCP4L1 |
| PCSK1N |
| PCSK4 |
| PCSK5 |
| PCSK6 |
| PCSK7 |

| |
|---|
| PCTP |
| PCYOX1 |
| PCYOX1L |
| PCYT1A |
| PCYT2 |
| PDAP1 |
| PDCD10 |
| PDCD11 |
| PDCD2 |
| PDCD2L |
| PDCD4 |
| PDCD5 |
| PDCD6 |
| PDCD6IP |
| PDCD7 |
| PDCL |
| PDCL3 |
| PDE12 |
| PDE2A |
| PDE4B |
| PDE4C |
| PDE4D |
| PDE4DIP |
| PDE6D |
| PDE7B |
| PDE8A |
| PDE9A |
| PDGFA |
| PDGFB |
| PDGFC |
| PDGFD |
| PDGFRB |
| PDHA1 |
| PDHB |
| PDHX |
| PDIA2 |
| PDIA3 |
| PDIA4 |
| PDIA5 |
| PDIA6 |
| PDIK1L |
| PDK1 |
| PDK3 |
| PDK4 |
| PDLIM1 |

| |
|---|
| PDLIM2 |
| PDLIM4 |
| PDLIM5 |
| PDLIM7 |
| PDPK1 |
| PDRG1 |
| PDS5A |
| PDS5B |
| PDSS1 |
| PDXDC1 |
| PDXK |
| PDXP |
| PDZD11 |
| PDZD2 |
| PDZD4 |
| PDZD8 |
| PDZK1 |
| PDZRN3 |
| PDZRN4 |
| PEBP1 |
| PECR |
| PEG10 |
| PEG3 |
| PELI3 |
| PELO |
| PELP1 |
| PEMT |
| PENK |
| PER1 |
| PER2 |
| PER3 |
| PERP |
| PEX1 |
| PEX10 |
| PEX11A |
| PEX11B |
| PEX11G |
| PEX12 |
| PEX13 |
| PEX14 |
| PEX16 |
| PEX19 |
| PEX2 |
| PEX3 |
| PEX5 |

| |
|---|
| PEX6 |
| PEX7 |
| PF4 |
| PFAS |
| PFDN1 |
| PFDN2 |
| PFDN4 |
| PFDN5 |
| PFDN6 |
| PFKFB2 |
| PFKFB3 |
| PFKFB4 |
| PFKL |
| PFKM |
| PFKP |
| PFN1 |
| PFN2 |
| PGAM1 |
| PGAM5 |
| PGAP1 |
| PGAP2 |
| PGAP3 |
| PGBD1 |
| PGF |
| PGGT1B |
| PGK1 |
| PGLS |
| PGM1 |
| PGM2 |
| PGM2L1 |
| PGM3 |
| PGM5 |
| PGP |
| PGPEP1 |
| PGR |
| PGRMC1 |
| PGRMC2 |
| PGS1 |
| PHACTR1 |
| PHACTR2 |
| PHACTR4 |
| PHAX |
| PHB2 |
| PHC1 |
| PHC2 |

| |
|---|
| PHC3 |
| PHF1 |
| PHF11 |
| PHF12 |
| PHF13 |
| PHF14 |
| PHF19 |
| PHF2 |
| PHF20 |
| PHF21A |
| PHF23 |
| PHF3 |
| PHF5A |
| PHF6 |
| PHF8 |
| PHGDH |
| PHIP |
| PHKA2 |
| PHKB |
| PHKG2 |
| PHLDA1 |
| PHLDA2 |
| PHLDB1 |
| PHLDB2 |
| PHLDB3 |
| PHLPP2 |
| PHOSPHO2 |
| PHYH |
| PI16 |
| PI4K2A |
| PI4K2B |
| PI4KA |
| PI4KB |
| PIAS1 |
| PIAS2 |
| PIAS3 |
| PIAS4 |
| PICALM |
| PICK1 |
| PIF1 |
| PIGA |
| PIGC |
| PIGF |
| PIGH |
| PIGK |

| |
|---|
| PIGL |
| PIGM |
| PIGN |
| PIGO |
| PIGP |
| PIGR |
| PIGS |
| PIGT |
| PIGU |
| PIGV |
| PIH1D1 |
| PIH1D2 |
| PIK3C2A |
| PIK3C2B |
| PIK3C3 |
| PIK3CA |
| PIK3CB |
| PIK3CD |
| PIK3IP1 |
| PIK3R1 |
| PIK3R2 |
| PIK3R3 |
| PIK3R4 |
| PIK3R5 |
| PIKFYVE |
| PIM1 |
| PIM2 |
| PIN1 |
| PIN4 |
| PINK1 |
| PINX1 |
| PIP4K2A |
| PIP4K2B |
| PIP5K1A |
| PIP5K1B |
| PIP5K1C |
| PIR |
| PISD |
| PITPNA |
| PITPNM1 |
| PITPNM2 |
| PITRM1 |
| PITX3 |
| PJA1 |
| PKD2 |

| |
|---|
| PKD2L2 |
| PKDCC |
| PKIA |
| PKIG |
| PKMYT1 |
| PKN1 |
| PKN2 |
| PKN3 |
| PKNOX1 |
| PKP1 |
| PKP2 |
| PKP3 |
| PKP4 |
| PLA2G12A |
| PLA2G15 |
| PLA2G6 |
| PLA2G7 |
| PLAA |
| PLAC8 |
| PLAGL1 |
| PLAGL2 |
| PLAU |
| PLAUR |
| PLBD1 |
| PLBD2 |
| PLCB3 |
| PLCB4 |
| PLCD1 |
| PLCD3 |
| PLCE1 |
| PLCG1 |
| PLCH2 |
| PLCL2 |
| PLCXD1 |
| PLCXD2 |
| PLD2 |
| PLD3 |
| PLEK |
| PLEK2 |
| PLEKHA1 |
| PLEKHA2 |
| PLEKHA3 |
| PLEKHA4 |
| PLEKHA5 |
| PLEKHA7 |

| |
|---|
| PLEKHA8 |
| PLEKHB1 |
| PLEKHF1 |
| PLEKHF2 |
| PLEKHG1 |
| PLEKHG2 |
| PLEKHG3 |
| PLEKHG5 |
| PLEKHH1 |
| PLEKHH3 |
| PLEKHJ1 |
| PLEKHM2 |
| PLEKHM3 |
| PLEKHN1 |
| PLEKHO1 |
| PLEKHO2 |
| PLIN1 |
| PLIN2 |
| PLIN3 |
| PLK1 |
| PLK2 |
| PLK3 |
| PLK4 |
| PLOD1 |
| PLOD2 |
| PLOD3 |
| PLP2 |
| PLRG1 |
| PLS3 |
| PLSCR4 |
| PLSCR5 |
| PLTP |
| PLXDC1 |
| PLXNB2 |
| PLXNB3 |
| PLXNC1 |
| PLXND1 |
| PM20D2 |
| PMF1 |
| PML |
| PMM1 |
| PMM2 |
| PMP22 |
| PMPCA |
| PMPCB |

| |
|---|
| PMS1 |
| PMVK |
| PNKD |
| PNKP |
| PNMA1 |
| PNN |
| PNO1 |
| PNPLA2 |
| PNPLA3 |
| PNPLA4 |
| PNPLA6 |
| PNPLA8 |
| PNPO |
| PNPT1 |
| PNRC1 |
| PNRC2 |
| POFUT1 |
| POFUT2 |
| POGK |
| POGZ |
| POLA1 |
| POLA2 |
| POLB |
| POLD1 |
| POLD2 |
| POLD3 |
| POLD4 |
| POLDIP2 |
| POLDIP3 |
| POLE |
| POLE2 |
| POLE3 |
| POLE4 |
| POLG |
| POLG2 |
| POLH |
| POLI |
| POLK |
| POLL |
| POLM |
| POLQ |
| POLR1C |
| POLR1D |
| POLR1E |
| POLR2A |

| |
|---|
| POLR2B |
| POLR2C |
| POLR2D |
| POLR2E |
| POLR2F |
| POLR2G |
| POLR2H |
| POLR2I |
| POLR2J |
| POLR2L |
| POLR3A |
| POLR3B |
| POLR3C |
| POLR3E |
| POLR3F |
| POLR3GL |
| POLR3H |
| POLR3K |
| POLRMT |
| POM121L2 |
| POMC |
| POMGNT1 |
| POMP |
| POMT1 |
| POMT2 |
| PON2 |
| POP1 |
| POP5 |
| POP7 |
| POR |
| PORCN |
| POSTN |
| POT1 |
| POU2AF1 |
| POU2F1 |
| POU5F1 |
| POU6F1 |
| PPA1 |
| PPA2 |
| PPARA |
| PPARD |
| PPARG |
| PPAT |
| PPCDC |
| PPCS |

| |
|---|
| PPDPF |
| PPFIA1 |
| PPFIA4 |
| PPFIBP1 |
| PPFIBP2 |
| PPHLN1 |
| PPIA |
| PPIB |
| PPIC |
| PPID |
| PPIE |
| PPIF |
| PPIG |
| PPIH |
| PPIL1 |
| PPIL2 |
| PPIL3 |
| PPIL4 |
| PPIL6 |
| PPIP5K1 |
| PPIP5K2 |
| PPL |
| PPM1A |
| PPM1B |
| PPM1D |
| PPM1F |
| PPM1G |
| PPM1K |
| PPM1M |
| PPOX |
| PPP1CA |
| PPP1CC |
| PPP1R10 |
| PPP1R11 |
| PPP1R12A |
| PPP1R12C |
| PPP1R13B |
| PPP1R13L |
| PPP1R14A |
| PPP1R14C |
| PPP1R14D |
| PPP1R15A |
| PPP1R15B |
| PPP1R2 |
| PPP1R3A |

| |
|---|
| PPP1R3B |
| PPP1R3D |
| PPP1R3F |
| PPP1R7 |
| PPP1R8 |
| PPP1R9A |
| PPP2CA |
| PPP2CB |
| PPP2R1A |
| PPP2R1B |
| PPP2R2A |
| PPP2R3A |
| PPP2R3C |
| PPP2R5A |
| PPP2R5B |
| PPP2R5C |
| PPP2R5D |
| PPP2R5E |
| PPP3CA |
| PPP3CB |
| PPP3R1 |
| PPP4C |
| PPP4R1 |
| PPP4R2 |
| PPP4R4 |
| PPP5C |
| PPP6C |
| PPRC1 |
| PPT1 |
| PPT2 |
| PPTC7 |
| PPWD1 |
| PQBP1 |
| PRAF2 |
| PRAM1 |
| PRAME |
| PRC1 |
| PRCC |
| PRCP |
| PRDM1 |
| PRDM10 |
| PRDM15 |
| PRDM4 |
| PRDM5 |
| PRDM7 |

| |
|---|
| PRDX2 |
| PRDX3 |
| PRDX4 |
| PRDX5 |
| PRDX6 |
| PREB |
| PRELID1 |
| PREP |
| PREPL |
| PREX1 |
| PRICKLE1 |
| PRICKLE3 |
| PRICKLE4 |
| PRIM1 |
| PRIM2 |
| PRKAA1 |
| PRKAA2 |
| PRKAB1 |
| PRKACA |
| PRKACB |
| PRKAG1 |
| PRKAG2 |
| PRKAR1B |
| PRKAR2A |
| PRKCD |
| PRKCE |
| PRKCI |
| PRKCSH |
| PRKCZ |
| PRKD1 |
| PRKD2 |
| PRKD3 |
| PRKDC |
| PRKRA |
| PRKRIP1 |
| PRMT1 |
| PRMT2 |
| PRMT3 |
| PRMT5 |
| PRMT6 |
| PRMT7 |
| PRODH |
| PROS1 |
| PRPF18 |
| PRPF19 |

| |
|---|
| PRPF3 |
| PRPF31 |
| PRPF38A |
| PRPF39 |
| PRPF40B |
| PRPF4B |
| PRPF6 |
| PRPF8 |
| PRPS1 |
| PRPS2 |
| PRPSAP1 |
| PRPSAP2 |
| PRR11 |
| PRR12 |
| PRR14 |
| PRR15 |
| PRR15L |
| PRR5 |
| PRR7 |
| PRRC1 |
| PRRG2 |
| PRSS35 |
| PRUNE2 |
| PSAP |
| PSAT1 |
| PSD3 |
| PSD4 |
| PSEN1 |
| PSEN2 |
| PSENEN |
| PSIP1 |
| PSKH1 |
| PSMA1 |
| PSMA2 |
| PSMA3 |
| PSMA5 |
| PSMA6 |
| PSMA7 |
| PSMB1 |
| PSMB10 |
| PSMB2 |
| PSMB3 |
| PSMB4 |
| PSMB5 |
| PSMB7 |

| |
|---|
| PSMB9 |
| PSMC1 |
| PSMC2 |
| PSMC3 |
| PSMC3IP |
| PSMC5 |
| PSMC6 |
| PSMD1 |
| PSMD10 |
| PSMD11 |
| PSMD12 |
| PSMD14 |
| PSMD2 |
| PSMD3 |
| PSMD4 |
| PSMD5 |
| PSMD6 |
| PSMD7 |
| PSMD8 |
| PSMD9 |
| PSME1 |
| PSME2 |
| PSME4 |
| PSMF1 |
| PSMG2 |
| PSMG3 |
| PSMG4 |
| PSPC1 |
| PSPH |
| PSRC1 |
| PSTK |
| PTAFR |
| PTAR1 |
| PTBP1 |
| PTCD2 |
| PTCD3 |
| PTCH1 |
| PTCH2 |
| PTDSS1 |
| PTEN |
| PTER |
| PTGDS |
| PTGER1 |
| PTGER2 |
| PTGER3 |

| |
|---|
| PTGER4 |
| PTGES2 |
| PTGES3 |
| PTGFR |
| PTGFRN |
| PTGS2 |
| PTH2 |
| PTHLH |
| PTK2 |
| PTK6 |
| PTK7 |
| PTMA |
| PTN |
| PTP4A1 |
| PTP4A2 |
| PTP4A3 |
| PTPDC1 |
| PTPMT1 |
| PTPN1 |
| PTPN11 |
| PTPN12 |
| PTPN18 |
| PTPN2 |
| PTPN21 |
| PTPN23 |
| PTPN3 |
| PTPN6 |
| PTPN9 |
| PTPRA |
| PTPRB |
| PTPRC |
| PTPRD |
| PTPRE |
| PTPRF |
| PTPRH |
| PTPRK |
| PTPRM |
| PTPRS |
| PTS |
| PTTG1 |
| PTX3 |
| PUF60 |
| PUM1 |
| PUM2 |
| PURA |

| |
|---|
| PURG |
| PUS1 |
| PUS3 |
| PUS7 |
| PUS7L |
| PUSL1 |
| PVR |
| PWP1 |
| PWP2 |
| PWWP2A |
| PWWP2B |
| PXK |
| PXMP2 |
| PXMP4 |
| PXN |
| PYCR1 |
| PYGB |
| PYGL |
| PYGM |
| PYGO1 |
| PYROXD1 |
| QDPR |
| QKI |
| QPCTL |
| QRFPR |
| QRICH1 |
| QRSL1 |
| QSOX1 |
| QSOX2 |
| QTRT1 |
| R3HCC1 |
| R3HDM2 |
| RAB10 |
| RAB11A |
| RAB11B |
| RAB11FIP1 |
| RAB11FIP2 |
| RAB11FIP4 |
| RAB11FIP5 |
| RAB12 |
| RAB13 |
| RAB14 |
| RAB15 |
| RAB18 |
| RAB1A |

| |
|---|
| RAB1B |
| RAB20 |
| RAB21 |
| RAB22A |
| RAB23 |
| RAB24 |
| RAB25 |
| RAB27A |
| RAB27B |
| RAB28 |
| RAB2A |
| RAB2B |
| RAB30 |
| RAB31 |
| RAB32 |
| RAB33B |
| RAB34 |
| RAB35 |
| RAB36 |
| RAB37 |
| RAB38 |
| RAB3B |
| RAB3D |
| RAB3GAP1 |
| RAB3GAP2 |
| RAB3IL1 |
| RAB40B |
| RAB40C |
| RAB4A |
| RAB4B |
| RAB5A |
| RAB5C |
| RAB6A |
| RAB8A |
| RAB8B |
| RAB9A |
| RABEP1 |
| RABEP2 |
| RABEPK |
| RABGAP1L |
| RABGGTA |
| RABIF |
| RABL3 |
| RAC1 |
| RAC2 |

| |
|---|
| RAC3 |
| RACGAP1 |
| RAD1 |
| RAD17 |
| RAD18 |
| RAD21 |
| RAD23A |
| RAD23B |
| RAD50 |
| RAD51 |
| RAD51AP1 |
| RAD51C |
| RAD52 |
| RAD54B |
| RAD9A |
| RAE1 |
| RAI1 |
| RAI2 |
| RALA |
| RALB |
| RALBP1 |
| RALGAPA1 |
| RALGAPA2 |
| RALGDS |
| RALGPS1 |
| RALGPS2 |
| RALY |
| RALYL |
| RANBP1 |
| RANBP10 |
| RANBP3 |
| RANBP6 |
| RANBP9 |
| RANGRF |
| RAP1A |
| RAP1B |
| RAP1GAP |
| RAP2A |
| RAP2B |
| RAP2C |
| RAPGEF1 |
| RAPGEF2 |
| RAPGEF3 |
| RAPGEF4 |
| RAPGEF6 |

| |
|---|
| RAPGEFL1 |
| RAPSN |
| RARA |
| RARG |
| RARRES1 |
| RARS2 |
| RASA1 |
| RASA2 |
| RASA3 |
| RASAL2 |
| RASAL3 |
| RASD1 |
| RASEF |
| RASGEF1B |
| RASGEF1C |
| RASGRP2 |
| RASGRP3 |
| RASGRP4 |
| RASIP1 |
| RASL10B |
| RASL11A |
| RASL11B |
| RASSF1 |
| RASSF2 |
| RASSF3 |
| RASSF5 |
| RASSF6 |
| RASSF7 |
| RASSF8 |
| RASSF9 |
| RAVER1 |
| RAVER2 |
| RB1 |
| RB1CC1 |
| RBBP4 |
| RBBP5 |
| RBBP6 |
| RBBP7 |
| RBBP8 |
| RBBP9 |
| RBCK1 |
| RBKS |
| RBL2 |
| RBM10 |
| RBM12B |

| |
|---|
| RBM14 |
| RBM15 |
| RBM15B |
| RBM17 |
| RBM18 |
| RBM22 |
| RBM25 |
| RBM27 |
| RBM28 |
| RBM3 |
| RBM33 |
| RBM34 |
| RBM41 |
| RBM42 |
| RBM43 |
| RBM45 |
| RBM47 |
| RBM4B |
| RBM5 |
| RBM6 |
| RBM7 |
| RBM8A |
| RBMS1 |
| RBMS3 |
| RBMX |
| RBP2 |
| RBP7 |
| RBPMS2 |
| RBX1 |
| RC3H1 |
| RCAN3 |
| RCBTB1 |
| RCBTB2 |
| RCC2 |
| RCCD1 |
| RCE1 |
| RCHY1 |
| RCL1 |
| RCN1 |
| RCN2 |
| RCN3 |
| RCOR1 |
| RCOR2 |
| RCOR3 |
| RCSD1 |

| |
|---|
| RDH10 |
| RDH11 |
| RDH13 |
| RDH14 |
| RDM1 |
| RECK |
| RECQL |
| RECQL4 |
| RECQL5 |
| REEP1 |
| REEP2 |
| REEP4 |
| REEP5 |
| REEP6 |
| REL |
| RELA |
| RELB |
| RELL1 |
| RELL2 |
| RELN |
| RELT |
| REM2 |
| RENBP |
| REP15 |
| REPIN1 |
| REPS1 |
| REPS2 |
| RER1 |
| RERE |
| RERG |
| RETSAT |
| REV1 |
| REV3L |
| REXO1 |
| REXO2 |
| RFC2 |
| RFC3 |
| RFC4 |
| RFC5 |
| RFESD |
| RFFL |
| RFK |
| RFNG |
| RFT1 |
| RFTN1 |

| |
|---|
| RFTN2 |
| RFWD3 |
| RFX2 |
| RFX7 |
| RFXANK |
| RFXAP |
| RGL1 |
| RGL2 |
| RGMA |
| RGMB |
| RGS10 |
| RGS11 |
| RGS12 |
| RGS18 |
| RGS19 |
| RGS2 |
| RGS4 |
| RGS5 |
| RGS9BP |
| RHBDD1 |
| RHBDD2 |
| RHBDD3 |
| RHBDF2 |
| RHBDL2 |
| RHEB |
| RHOB |
| RHOBTB1 |
| RHOBTB3 |
| RHOD |
| RHOF |
| RHOG |
| RHOQ |
| RHOT1 |
| RHOT2 |
| RHOU |
| RHPN2 |
| RIBC1 |
| RIC3 |
| RIC8A |
| RIC8B |
| RICTOR |
| RIF1 |
| RILPL1 |
| RILPL2 |
| RIMKLA |

| |
|---|
| RIMKLB |
| RIMS1 |
| RIMS2 |
| RIN1 |
| RING1 |
| RINT1 |
| RIOK1 |
| RIOK2 |
| RIOK3 |
| RIPK1 |
| RIPK2 |
| RIT1 |
| RLF |
| RMI2 |
| RMND5A |
| RMND5B |
| RNASE4 |
| RNASEH1 |
| RNASEH2A |
| RNASEH2B |
| RNASEH2C |
| RNASEK |
| RNASET2 |
| RND2 |
| RND3 |
| RNF10 |
| RNF103 |
| RNF11 |
| RNF111 |
| RNF112 |
| RNF113A |
| RNF114 |
| RNF115 |
| RNF123 |
| RNF126 |
| RNF13 |
| RNF135 |
| RNF139 |
| RNF14 |
| RNF141 |
| RNF145 |
| RNF148 |
| RNF149 |
| RNF166 |
| RNF167 |

| |
|---|
| RNF168 |
| RNF169 |
| RNF170 |
| RNF181 |
| RNF182 |
| RNF185 |
| RNF187 |
| RNF19B |
| RNF2 |
| RNF20 |
| RNF213 |
| RNF214 |
| RNF215 |
| RNF216 |
| RNF220 |
| RNF222 |
| RNF24 |
| RNF25 |
| RNF26 |
| RNF31 |
| RNF32 |
| RNF4 |
| RNF40 |
| RNF41 |
| RNF44 |
| RNF5 |
| RNF6 |
| RNF7 |
| RNFT1 |
| RNFT2 |
| RNGTT |
| RNH1 |
| RNLS |
| RNMT |
| RNPC3 |
| RNPEP |
| RNPEPL1 |
| RNPS1 |
| ROBO1 |
| ROCK1 |
| ROCK2 |
| ROGDI |
| ROM1 |
| ROMO1 |
| RP2 |

| |
|---|
| RP9 |
| RPA1 |
| RPA2 |
| RPA3 |
| RPAIN |
| RPAP1 |
| RPAP3 |
| RPE |
| RPF1 |
| RPF2 |
| RPGR |
| RPGRIP1L |
| RPH3AL |
| RPIA |
| RPL10A |
| RPL11 |
| RPL12 |
| RPL13 |
| RPL13A |
| RPL14 |
| RPL15 |
| RPL18 |
| RPL18A |
| RPL21 |
| RPL22L1 |
| RPL23 |
| RPL23A |
| RPL24 |
| RPL26 |
| RPL27 |
| RPL27A |
| RPL28 |
| RPL29 |
| RPL3 |
| RPL31 |
| RPL32 |
| RPL34 |
| RPL35 |
| RPL36 |
| RPL36A |
| RPL37 |
| RPL38 |
| RPL4 |
| RPL5 |
| RPL6 |

| |
|---|
| RPL7 |
| RPL7A |
| RPL7L1 |
| RPL8 |
| RPL9 |
| RPLP1 |
| RPLP2 |
| RPN1 |
| RPN2 |
| RPP21 |
| RPP25 |
| RPP30 |
| RPP38 |
| RPP40 |
| RPRD1A |
| RPRD2 |
| RPRM |
| RPS10 |
| RPS12 |
| RPS13 |
| RPS14 |
| RPS15 |
| RPS15A |
| RPS16 |
| RPS17 |
| RPS18 |
| RPS19BP1 |
| RPS2 |
| RPS21 |
| RPS24 |
| RPS25 |
| RPS26 |
| RPS27 |
| RPS27A |
| RPS27L |
| RPS3 |
| RPS3A |
| RPS5 |
| RPS6 |
| RPS6KA1 |
| RPS6KA2 |
| RPS6KA3 |
| RPS6KA4 |
| RPS6KA5 |
| RPS6KB1 |

| |
|---|
| RPS6KB2 |
| RPS6KC1 |
| RPS7 |
| RPS8 |
| RPS9 |
| RPSA |
| RPTOR |
| RPUSD1 |
| RPUSD2 |
| RPUSD4 |
| RRAD |
| RRAGA |
| RRAGB |
| RRAGC |
| RRAGD |
| RRAS |
| RRAS2 |
| RRM1 |
| RRM2 |
| RRM2B |
| RRN3 |
| RRP1 |
| RRP12 |
| RRP15 |
| RRP1B |
| RRP7A |
| RRP8 |
| RRP9 |
| RRS1 |
| RSAD1 |
| RSAD2 |
| RSBN1 |
| RSBN1L |
| RSF1 |
| RSL1D1 |
| RSPH3 |
| RSPH4A |
| RSPRY1 |
| RSRC2 |
| RTF1 |
| RTN1 |
| RTN2 |
| RTN3 |
| RTN4 |
| RTN4IP1 |

| |
|---|
| RTTN |
| RUFY1 |
| RUFY2 |
| RUFY3 |
| RUNDC1 |
| RUNX1 |
| RUNX3 |
| RUSC1 |
| RUSC2 |
| RUVBL1 |
| RUVBL2 |
| RWDD1 |
| RWDD2A |
| RWDD2B |
| RWDD3 |
| RXRA |
| RXRB |
| RYBP |
| RYK |
| RYR2 |
| RYR3 |
| S100A10 |
| S100A13 |
| S100A16 |
| S100A2 |
| S100A4 |
| S100A6 |
| S100A9 |
| S100PBP |
| S1PR1 |
| S1PR2 |
| SAA4 |
| SAAL1 |
| SACM1L |
| SACS |
| SAE1 |
| SAFB |
| SALL2 |
| SAMD10 |
| SAMD11 |
| SAMD4A |
| SAMD4B |
| SAMD5 |
| SAMD8 |
| SAMD9 |

| |
|---|
| SAMHD1 |
| SAMM50 |
| SAP18 |
| SAP30 |
| SAP30BP |
| SAP30L |
| SAR1A |
| SAR1B |
| SARDH |
| SARM1 |
| SARNP |
| SARS2 |
| SART1 |
| SASH1 |
| SASS6 |
| SAT1 |
| SAT2 |
| SAV1 |
| SBDS |
| SBF1 |
| SBF2 |
| SBSN |
| SCAF1 |
| SCAI |
| SCAMP1 |
| SCAMP2 |
| SCAMP3 |
| SCAMP4 |
| SCAND1 |
| SCARA3 |
| SCARB2 |
| SCARF1 |
| SCARF2 |
| SCCPDH |
| SCD5 |
| SCFD1 |
| SCFD2 |
| SCG5 |
| SCIN |
| SCLT1 |
| SCLY |
| SCMH1 |
| SCML1 |
| SCML2 |
| SCN2A |

| |
|---|
| SCN9A |
| SCNM1 |
| SCNN1A |
| SCNN1B |
| SCO1 |
| SCOC |
| SCP2 |
| SCPEP1 |
| SCRN1 |
| SCRN2 |
| SCRN3 |
| SCUBE1 |
| SCUBE3 |
| SCYL1 |
| SCYL3 |
| SDAD1 |
| SDC1 |
| SDC2 |
| SDC3 |
| SDC4 |
| SDCBP |
| SDCCAG8 |
| SDF2 |
| SDF2L1 |
| SDF4 |
| SDHAF1 |
| SDHB |
| SDHC |
| SDHD |
| SDR39U1 |
| SEC11A |
| SEC11C |
| SEC13 |
| SEC14L1 |
| SEC14L2 |
| SEC16A |
| SEC22A |
| SEC22B |
| SEC22C |
| SEC23A |
| SEC23B |
| SEC23IP |
| SEC24A |
| SEC24B |
| SEC24C |

| |
|---|
| SEC24D |
| SEC31A |
| SEC31B |
| SEC61A2 |
| SEC61B |
| SEC61G |
| SEC62 |
| SEC63 |
| SECISBP2 |
| SECTM1 |
| SEH1L |
| SEL1L |
| SEL1L3 |
| SELE |
| SELENBP1 |
| SELPLG |
| SEMA3B |
| SEMA3F |
| SEMA3G |
| SEMA4B |
| SEMA4C |
| SEMA4D |
| SEMA4F |
| SEMA4G |
| SEMA5A |
| SEMA6A |
| SEMA6B |
| SEMA6C |
| SEMA6D |
| SEMA7A |
| SENP1 |
| SENP2 |
| SENP3 |
| SENP6 |
| SENP7 |
| SEPHS1 |
| SEPHS2 |
| SERAC1 |
| SERBP1 |
| SERGEF |
| SERINC1 |
| SERINC2 |
| SERINC3 |
| SERINC5 |
| SERP1 |

| |
|---|
| SERPINA5 |
| SERPINB1 |
| SERPINB2 |
| SERPINB6 |
| SERPINB7 |
| SERPINE1 |
| SERPINE2 |
| SERPINH1 |
| SERPINI1 |
| SERTAD2 |
| SERTAD3 |
| SESN3 |
| SET |
| SETD1A |
| SETD1B |
| SETD2 |
| SETD4 |
| SETD5 |
| SETD6 |
| SETD7 |
| SETMAR |
| SETX |
| SF1 |
| SF3A1 |
| SF3A2 |
| SF3A3 |
| SF3B1 |
| SF3B2 |
| SF3B3 |
| SF3B4 |
| SF3B5 |
| SFI1 |
| SFN |
| SFPQ |
| SFRP1 |
| SFRP4 |
| SFT2D1 |
| SFT2D2 |
| SFXN1 |
| SFXN2 |
| SFXN4 |
| SFXN5 |
| SGCE |
| SGK1 |
| SGK3 |

| |
|---|
| SGMS2 |
| SGPL1 |
| SGPP1 |
| SGSM1 |
| SGSM3 |
| SGTA |
| SH2B1 |
| SH2B2 |
| SH2B3 |
| SH2D3C |
| SH2D4A |
| SH2D6 |
| SH3BGRL |
| SH3BGRL2 |
| SH3BGRL3 |
| SH3BP2 |
| SH3BP4 |
| SH3BP5 |
| SH3BP5L |
| SH3D19 |
| SH3GL1 |
| SH3GL2 |
| SH3GLB1 |
| SH3GLB2 |
| SH3KBP1 |
| SH3PXD2B |
| SH3RF1 |
| SH3TC2 |
| SH3YL1 |
| SHARPIN |
| SHB |
| SHC1 |
| SHCBP1 |
| SHD |
| SHF |
| SHISA4 |
| SHISA5 |
| SHISA8 |
| SHKBP1 |
| SHMT1 |
| SHMT2 |
| SHOC2 |
| SHPK |
| SHQ1 |
| SHROOM1 |

| |
|---|
| SHROOM2 |
| SHROOM3 |
| SIAE |
| SIAH1 |
| SIDT2 |
| SIGLEC10 |
| SIGLEC8 |
| SIGMAR1 |
| SIK2 |
| SIL1 |
| SIPA1 |
| SIPA1L1 |
| SIPA1L2 |
| SIRPA |
| SIRT1 |
| SIRT2 |
| SIRT3 |
| SIRT5 |
| SIRT6 |
| SIRT7 |
| SIT1 |
| SIVA1 |
| SIX5 |
| SKA3 |
| SKI |
| SKIL |
| SKP1 |
| SKP2 |
| SLA2 |
| SLAIN1 |
| SLAIN2 |
| SLAMF7 |
| SLBP |
| SLC10A3 |
| SLC10A4 |
| SLC10A5 |
| SLC11A1 |
| SLC11A2 |
| SLC12A2 |
| SLC12A5 |
| SLC12A9 |
| SLC13A4 |
| SLC15A1 |
| SLC15A3 |
| SLC15A4 |

| |
|---|
| SLC16A1 |
| SLC16A11 |
| SLC16A12 |
| SLC16A14 |
| SLC16A3 |
| SLC16A5 |
| SLC16A6 |
| SLC16A7 |
| SLC17A5 |
| SLC19A1 |
| SLC19A2 |
| SLC1A1 |
| SLC1A4 |
| SLC1A5 |
| SLC20A1 |
| SLC20A2 |
| SLC22A18 |
| SLC22A23 |
| SLC22A5 |
| SLC23A1 |
| SLC23A2 |
| SLC25A1 |
| SLC25A11 |
| SLC25A12 |
| SLC25A13 |
| SLC25A15 |
| SLC25A16 |
| SLC25A17 |
| SLC25A19 |
| SLC25A22 |
| SLC25A24 |
| SLC25A25 |
| SLC25A26 |
| SLC25A27 |
| SLC25A28 |
| SLC25A29 |
| SLC25A30 |
| SLC25A35 |
| SLC25A36 |
| SLC25A37 |
| SLC25A38 |
| SLC25A39 |
| SLC25A4 |
| SLC25A42 |
| SLC25A46 |

| |
|---|
| SLC25A5 |
| SLC26A1 |
| SLC26A11 |
| SLC26A2 |
| SLC26A3 |
| SLC26A4 |
| SLC27A1 |
| SLC27A3 |
| SLC27A4 |
| SLC27A5 |
| SLC27A6 |
| SLC29A2 |
| SLC29A3 |
| SLC29A4 |
| SLC2A1 |
| SLC2A13 |
| SLC2A3 |
| SLC2A6 |
| SLC2A8 |
| SLC30A1 |
| SLC30A4 |
| SLC30A5 |
| SLC30A6 |
| SLC30A7 |
| SLC30A9 |
| SLC31A1 |
| SLC31A2 |
| SLC33A1 |
| SLC34A2 |
| SLC35A1 |
| SLC35A2 |
| SLC35A4 |
| SLC35B1 |
| SLC35B2 |
| SLC35B3 |
| SLC35B4 |
| SLC35C1 |
| SLC35C2 |
| SLC35D2 |
| SLC35E1 |
| SLC35E4 |
| SLC35F5 |
| SLC36A1 |
| SLC36A4 |
| SLC37A3 |

| |
|---|
| SLC37A4 |
| SLC38A10 |
| SLC38A2 |
| SLC38A4 |
| SLC38A5 |
| SLC38A6 |
| SLC38A7 |
| SLC39A1 |
| SLC39A10 |
| SLC39A11 |
| SLC39A14 |
| SLC39A3 |
| SLC39A4 |
| SLC39A5 |
| SLC39A7 |
| SLC39A8 |
| SLC39A9 |
| SLC3A2 |
| SLC40A1 |
| SLC41A1 |
| SLC41A2 |
| SLC41A3 |
| SLC43A1 |
| SLC43A2 |
| SLC44A1 |
| SLC44A2 |
| SLC44A3 |
| SLC44A4 |
| SLC45A1 |
| SLC45A4 |
| SLC46A1 |
| SLC47A1 |
| SLC47A2 |
| SLC48A1 |
| SLC4A1AP |
| SLC4A2 |
| SLC4A3 |
| SLC5A12 |
| SLC5A3 |
| SLC5A6 |
| Slc6a12 |
| SLC6A15 |
| SLC6A6 |
| SLC6A8 |
| SLC7A11 |

| |
|---|
| SLC7A5 |
| SLC7A6 |
| SLC7A6OS |
| SLC7A7 |
| SLC8A1 |
| SLC8A2 |
| SLC9A1 |
| SLC9A3R1 |
| SLC9A3R2 |
| SLC9A6 |
| SLC9A7 |
| SLCO2B1 |
| SLCO3A1 |
| SLCO4C1 |
| SLIRP |
| SLITRK1 |
| SLITRK6 |
| SLK |
| SLMAP |
| SLTM |
| SLU7 |
| SMAD1 |
| SMAD2 |
| SMAD3 |
| SMAD4 |
| SMAD6 |
| SMAD7 |
| SMAGP |
| SMAP1 |
| SMAP2 |
| SMARCA1 |
| SMARCA4 |
| SMARCA5 |
| SMARCC1 |
| SMARCC2 |
| SMARCD2 |
| SMARCD3 |
| SMARCE1 |
| SMC1A |
| SMC1B |
| SMC2 |
| SMC3 |
| SMC4 |
| SMC5 |
| SMC6 |

| |
|---|
| SMCHD1 |
| SMG5 |
| SMG6 |
| SMG7 |
| SMG9 |
| SMN1 |
| SMNDC1 |
| SMO |
| SMOX |
| SMPD1 |
| SMPD2 |
| SMPD4 |
| SMPDL3A |
| SMPDL3B |
| SMS |
| SMTN |
| SMU1 |
| SMUG1 |
| SMURF1 |
| SMURF2 |
| SMYD2 |
| SMYD3 |
| SMYD4 |
| SMYD5 |
| SNAI1 |
| SNAP23 |
| SNAP29 |
| SNAP47 |
| SNAPC1 |
| SNAPC3 |
| SNAPC4 |
| SNAPC5 |
| SNAPIN |
| SND1 |
| SNF8 |
| SNIP1 |
| SNN |
| SNPH |
| SNRK |
| SNRNP200 |
| SNRNP25 |
| SNRNP27 |
| SNRNP35 |
| SNRNP40 |
| SNRNP48 |

| |
|---|
| SNRNP70 |
| SNRPA1 |
| SNRPB |
| SNRPB2 |
| SNRPC |
| SNRPD1 |
| SNRPD3 |
| SNRPE |
| SNRPF |
| SNRPG |
| SNRPN |
| SNTA1 |
| SNTB1 |
| SNTB2 |
| SNUPN |
| SNW1 |
| SNX1 |
| SNX10 |
| SNX11 |
| SNX13 |
| SNX14 |
| SNX15 |
| SNX16 |
| SNX17 |
| SNX18 |
| SNX19 |
| SNX2 |
| SNX21 |
| SNX24 |
| SNX25 |
| SNX27 |
| SNX29 |
| SNX3 |
| SNX30 |
| SNX5 |
| SNX6 |
| SNX7 |
| SNX8 |
| SNX9 |
| SOAT1 |
| SOBP |
| SOCS1 |
| SOCS2 |
| SOCS3 |
| SOCS4 |

| |
|---|
| SOCS5 |
| SOCS6 |
| SOD1 |
| SOD2 |
| SOD3 |
| SON |
| SORBS3 |
| SORD |
| SORL1 |
| SOS1 |
| SOS2 |
| SOSTDC1 |
| SOX13 |
| SOX4 |
| SOX5 |
| SP3 |
| SPA17 |
| SPAG1 |
| SPAG4 |
| SPAG5 |
| SPAG7 |
| SPAG8 |
| SPAG9 |
| SPARC |
| SPAST |
| SPATA13 |
| SPATA18 |
| SPATA2 |
| SPATA20 |
| SPATA24 |
| SPATA2L |
| SPATA5L1 |
| SPATA6 |
| SPATA7 |
| SPC24 |
| SPC25 |
| SPCS1 |
| SPCS2 |
| SPCS3 |
| SPDEF |
| SPDYA |
| SPEN |
| SPG11 |
| SPG7 |
| SPHK1 |

| |
|---|
| SPHK2 |
| SPI1 |
| SPIN1 |
| SPIN4 |
| SPINT1 |
| SPINT2 |
| SPIRE1 |
| SPNS1 |
| SPOCK1 |
| SPOCK2 |
| SPON1 |
| SPON2 |
| SPOP |
| SPOPL |
| SPPL2B |
| SPR |
| SPRED2 |
| SPRY2 |
| SPRY3 |
| SPRY4 |
| SPSB1 |
| SPSB3 |
| SPSB4 |
| SPTAN1 |
| SPTBN1 |
| SPTLC1 |
| SPTLC2 |
| SPTY2D1 |
| SQLE |
| SQSTM1 |
| SRA1 |
| SRBD1 |
| SRC |
| SRD5A3 |
| SREBF1 |
| SREBF2 |
| SRF |
| SRFBP1 |
| SRGAP1 |
| SRGAP2 |
| SRGN |
| SRI |
| SRM |
| SRP14 |
| SRP19 |

| |
|---|
| SRP54 |
| SRP68 |
| SRP72 |
| SRP9 |
| SRPK1 |
| SRPK2 |
| SRPK3 |
| SRR |
| SRRD |
| SRRM1 |
| SRRM2 |
| SRRT |
| SS18 |
| SS18L1 |
| SSB |
| SSBP1 |
| SSBP2 |
| SSBP3 |
| SSBP4 |
| SSH1 |
| SSH2 |
| SSH3 |
| SSNA1 |
| SSR1 |
| SSR2 |
| SSR3 |
| SSR4 |
| SSRP1 |
| SSU72 |
| SSX2IP |
| ST13 |
| ST14 |
| ST3GAL2 |
| ST3GAL3 |
| ST3GAL4 |
| ST3GAL5 |
| ST3GAL6 |
| ST6GAL1 |
| ST6GAL2 |
| ST6GALNAC2 |
| ST6GALNAC6 |
| ST7 |
| ST8SIA4 |
| ST8SIA5 |
| STAB1 |

| |
|---|
| STAC |
| STAG1 |
| STAM |
| STAM2 |
| STAMBP |
| STAP2 |
| STAR |
| STARD10 |
| STARD13 |
| STARD3 |
| STARD3NL |
| STARD4 |
| STARD6 |
| STARD7 |
| STARD8 |
| STARD9 |
| STAT1 |
| STAT2 |
| STAT3 |
| STAT5A |
| STAU1 |
| STAU2 |
| STBD1 |
| STC1 |
| STEAP1 |
| STEAP3 |
| STIL |
| STIM1 |
| STIM2 |
| STIP1 |
| STK10 |
| STK11 |
| STK11IP |
| STK16 |
| STK17A |
| STK17B |
| STK19 |
| STK24 |
| STK25 |
| STK3 |
| STK33 |
| STK35 |
| STK36 |
| STK38 |
| STK38L |

| |
|---|
| STK39 |
| STMN1 |
| STOM |
| STOML1 |
| STOML2 |
| STON1 |
| STON2 |
| STOX1 |
| STRADB |
| STRBP |
| STRN |
| STRN3 |
| STRN4 |
| STS |
| STT3A |
| STUB1 |
| STX10 |
| STX11 |
| STX12 |
| STX16 |
| STX17 |
| STX18 |
| STX19 |
| STX1A |
| STX1B |
| STX2 |
| STX4 |
| STX5 |
| STX6 |
| STX7 |
| STX8 |
| STXBP3 |
| STXBP4 |
| STXBP5 |
| STXBP5L |
| STYX |
| SUB1 |
| SUCLA2 |
| SUCLG1 |
| SUCLG2 |
| SUFU |
| SUGT1 |
| SULT1A1 |
| SULT1B1 |
| SUMF1 |

| |
|---|
| SUMF2 |
| SUMO1 |
| SUMO3 |
| SUN2 |
| SUOX |
| SUPT16H |
| SUPT3H |
| SUPT4H1 |
| SUPT5H |
| SUPT6H |
| SUPT7L |
| SUPV3L1 |
| SURF1 |
| SURF2 |
| SURF4 |
| SUSD1 |
| SUSD3 |
| SUV39H1 |
| SUV39H2 |
| SUZ12 |
| SV2C |
| SVIL |
| SWAP70 |
| SYAP1 |
| SYCE2 |
| SYDE1 |
| SYDE2 |
| SYF2 |
| SYK |
| SYN1 |
| SYNC |
| SYNCRIP |
| SYNE1 |
| SYNE2 |
| SYNGAP1 |
| SYNGR1 |
| SYNGR3 |
| SYNJ1 |
| SYNJ2 |
| SYNJ2BP |
| SYNM |
| SYNPO2 |
| SYNRG |
| SYP |
| SYPL1 |

| |
|---|
| SYS1 |
| SYT1 |
| SYT10 |
| SYT11 |
| SYT17 |
| SYT7 |
| SYT9 |
| SYTL1 |
| SYTL2 |
| SYTL3 |
| SYTL4 |
| SYTL5 |
| SYVN1 |
| TAC1 |
| TACC2 |
| TACC3 |
| TACR3 |
| TADA1 |
| TADA2A |
| TADA2B |
| TADA3 |
| TAF10 |
| TAF11 |
| TAF12 |
| TAF13 |
| TAF15 |
| TAF1B |
| TAF1C |
| TAF1D |
| TAF4 |
| TAF4B |
| TAF5 |
| TAF5L |
| TAF6 |
| TAF6L |
| TAF7 |
| TAF8 |
| TAF9B |
| TAGAP |
| TAGLN2 |
| TAL2 |
| TALDO1 |
| TAOK1 |
| TAOK2 |
| TAP1 |

| |
|---|
| TAP2 |
| TAPBPL |
| TAPT1 |
| TARBP1 |
| TARBP2 |
| TARDBP |
| TARS2 |

**Supplemental List 5**

| Atretic Follicles | Growing Follicles | Plateau Follicles |
|---|---|---|
| AASDH | AAAS | ABCA6 |
| AASS | AATF | ABHD12 |
| ABCA5 | ABCB10 | ACBD3 |
| ABCB1 | ABCD4 | ACBD5 |
| ABCC1 | ABHD5 | ACOT8 |
| ACAA2 | ABHD6 | ACTB |
| ACAD11 | ACAD8 | ACTR2 |
| ACAT1 | ACBD5 | ACYP1 |
| ACOT11 | ACOT7 | ADAM10 |
| ACOX3 | ACOX3 | ADAM21 |
| ACSF2 | ACP1 | ADAT2 |
| ACSS1 | ACSL3 | AGFG1 |
| ACTR6 | ACTA1 | AIMP2 |
| ACVR2A | ACTA2 | AK2 |
| ADI1 | ACTN1 | AKAP11 |
| ADPGK | ACTR6 | AKIRIN1 |
| AFTPH | ACVR2A | AKR1B1 |
| AGBL3 | ACY1 | AKTIP |
| AGL | ADAMTS6 | ALKBH5 |
| AGPAT5 | ADCY6 | ALKBH7 |
| AGTPBP1 | ADCY7 | AMOTL1 |
| AHCTF1 | ADSL | AMZ2 |
| AKIRIN2 | AGFG1 | ANAPC11 |
| AKT2 | AGL | ANKRA2 |
| ALDH18A1 | AGPAT5 | ANKRD12 |
| ALDH1A1 | AGPS | AP2S1 |
| ALDH2 | AGRN | AP3M1 |
| ALDH4A1 | AHCYL2 | APH1A |
| ALDH9A1 | AHI1 | APIP |
| ALKBH2 | AIMP2 | APOO |
| ALMS1 | AK2 | APRT |
| ALPK1 | AK6 | ARF4 |
| AMDHD1 | AKAP1 | ARF5 |
| AMHR2 | AKR1A1 | ARG2 |
| AMN1 | AKR1B1 | ARHGAP6 |
| AMY2A | ALCAM | ARHGDIA |
| ANGEL2 | ALDOA | ARL2BP |
| ANGPTL2 | ALKBH2 | ARL5A |
| ANK3 | ALOX5 | ARL6IP1 |
| ANKH | AMD1 | ARPC3 |
| ANKLE2 | AMIGO2 | ARPC4 |
| ANKMY2 | AMN1 | ASAP2 |
| ANKRD17 | AMZ2 | ATP2B4 |

| | | |
|---|---|---|
| ANKRD22 | ANAPC11 | ATP6V0A1 |
| ANKRD46 | ANKRD10 | ATP6V0B |
| ANO10 | ANKRD12 | ATP6V0E1 |
| ANTXR2 | ANKRD17 | ATP6V1D |
| ANXA5 | ANKRD27 | ATP6V1G1 |
| AOX1 | ANKRD55 | ATXN7L1 |
| AP2M1 | ANP32A | AUP1 |
| AP3M1 | ANP32E | AZI2 |
| AP4B1 | ANXA11 | B3GNT2 |
| APEH | ANXA6 | B9D1 |
| APOA1 | AP1S2 | BBX |
| AQR | AP4B1 | BCAP31 |
| ARFGAP3 | APBB1IP | BCL10 |
| ARFGEF2 | APBB3 | BCR |
| ARFIP1 | APEH | BIN3 |
| ARFIP2 | APOA1 | BOLA1 |
| ARHGAP26 | APOO | BRMS1L |
| ARHGEF10 | AQP11 | BTC |
| ARHGEF3 | AQR | CABLES2 |
| ARHGEF6 | ARF4 | CAMLG |
| ARHGEF9 | ARF5 | CAPNS1 |
| ARID2 | ARFGEF2 | CAPZA2 |
| ARL6 | ARFIP1 | CBR4 |
| ARL6IP1 | ARFRP1 | CBX6 |
| ARMCX1 | ARHGAP10 | CCDC124 |
| ARSA | ARHGAP20 | CCDC28B |
| ARSK | ARHGAP21 | CCDC82 |
| ASB1 | ARHGAP23 | CCNE2 |
| ASB15 | ARHGAP4 | CCNG1 |
| ASH1L | ARHGDIA | CCNG2 |
| ASXL1 | ARHGEF2 | CCNH |
| ATAD1 | ARHGEF3 | CCNI |
| ATF2 | ARHGEF6 | CCT3 |
| ATG12 | ARHGEF9 | CD300LB |
| ATG13 | ARID1A | CD47 |
| ATP10D | ARID3A | CDC42 |
| ATP13A1 | ARL4A | CDC42SE2 |
| ATP6V0D1 | ARL4C | CDCA7L |
| AVL9 | ARL6 | CDIPT |
| B2M | ARMC5 | CDK20 |
| B3GAT3 | ARPC4 | CDK6 |
| B3GNT2 | ARPC5L | CDON |
| B4GALT3 | ARRB1 | CDS2 |
| B4GALT4 | ASAP2 | CENPI |
| BAZ1B | ASB1 | CENPJ |

| | | |
|---|---|---|
| BBS2 | ASB6 | CENPM |
| BBS5 | ASCL4 | CENPV |
| BBS7 | ASF1A | CFDP2 |
| BCAP29 | ASF1B | CHADL |
| BCAR3 | ASH1L | CHCHD3 |
| BCAS4 | ASPH | CHMP1A |
| BCKDHA | ASPM | CHMP2A |
| BCL2L11 | ASXL1 | CHMP5 |
| BDH2 | ATAD1 | CHSY1 |
| BEND6 | ATF2 | CIB1 |
| BEND7 | ATF3 | CINP |
| BET1L | ATG16L1 | CMBL |
| BGN | ATG2B | CMTM6 |
| BIRC3 | ATG3 | CNBP |
| BIRC6 | ATG9A | CNIH4 |
| BLMH | ATL3 | CNN3 |
| BLVRB | ATP1A3 | CNOT7 |
| BMPR1A | ATP6V0A2 | CNTLN |
| BMPR2 | ATP6V0D1 | COMMD10 |
| BNIP3L | ATP6V0E1 | COMMD5 |
| BRPF3 | ATP6V1A | COMMD8 |
| BSG | ATP6V1E1 | COPE |
| BTBD1 | ATP6V1F | COPS8 |
| BZW1 | ATP6V1G2 | COQ10B |
| C1QB | ATP9B | COQ3 |
| C4A | ATXN2 | COX7A2L |
| CAB39L | ATXN2L | CRELD2 |
| CACYBP | ATXN7L3 | CSPP1 |
| CAPN7 | AUP1 | CUL1 |
| CARD6 | AURKA | DAP |
| CASC3 | AVL9 | DBR1 |
| CASP9 | B4GALT1 | DCAF12 |
| CASZ1 | B4GALT5 | DCLRE1A |
| CBR1 | BAG2 | DCTN3 |
| CC2D1A | BAHD1 | DDOST |
| CC2D1B | BAZ1B | DDX41 |
| CC2D2A | BAZ2A | DHRS1 |
| CCDC125 | BBS5 | DHRSX |
| CCDC28A | BBS7 | DIABLO |
| CCDC32 | BCAS4 | DKKL1 |
| CCDC43 | BCL2 | DNAJC15 |
| CCDC50 | BCL7B | DNAJC19 |
| CCDC66 | BCL9 | DNAJC3 |
| CCDC82 | BCOR | DNAL1 |
| CCDC90B | BFSP1 | DOCK9 |

| | | |
|---|---|---|
| CCL5 | BGN | DPM3 |
| CCNI | BIN2 | DPY19L3 |
| CCNK | BIRC5 | DTD1 |
| CCNT1 | BIRC6 | DTYMK |
| CCPG1 | BOLA1 | DUSP23 |
| CD2BP2 | BOP1 | DYNLRB1 |
| CD36 | BRD7 | E2F2 |
| CD46 | BRDT | EBAG9 |
| CD53 | BRIP1 | EDF1 |
| CD68 | BRIX1 | EFCAB2 |
| CD96 | BRMS1L | EID1 |
| CD99L2 | BUB1 | EIF3I |
| CDC37L1 | BUB1B | EIF3K |
| CDC40 | BZW1 | EIF4H |
| CDC7 | BZW2 | ELOVL5 |
| CDH12 | C1orf35 | ELOVL6 |
| CDK10 | C2CD2L | EML4 |
| CDK13 | C6orf120 | EMP2 |
| CDV3 | CA2 | EMX2 |
| CDYL2 | CAB39L | ENOX1 |
| CEP350 | CABLES2 | ENPEP |
| CGRRF1 | CACNA1G | ENSA |
| CHD1L | CACYBP | ERCC1 |
| CHD2 | CALM1 | ERCC8 |
| CHD8 | CAMTA1 | ERGIC3 |
| CHIC1 | CAPN7 | ETFB |
| CHKA | CAPZA2 | ETV2 |
| CHMP2B | CARD6 | EXOSC4 |
| CHMP4C | CASC3 | EZH2 |
| CHMP6 | CASD1 | FADD |
| CHST12 | CASP1 | FAM110A |
| CHUK | CASP8 | FAM32A |
| CIAO1 | CC2D1B | FAM8A1 |
| CITED2 | CCDC125 | FARSA |
| CLASP2 | CCDC159 | FAU |
| CLDN12 | CCDC25 | FBXO8 |
| CLEC1A | CCDC66 | FGD5 |
| CLGN | CCDC85B | FKBP5 |
| CLIP3 | CCL2 | FNBP1L |
| CLN3 | CCNA2 | FOXL2 |
| CLSTN1 | CCNB2 | FUNDC2 |
| CLYBL | CCND2 | FUT1 |
| CNBP | CCNE1 | GCSH |
| CNKSR3 | CCNE2 | GK |
| CNNM3 | CCNG2 | GK2 |

| | | |
|---|---|---|
| CNPY4 | CCNK | GMPS |
| CNTLN | CCNYL1 | GNAQ |
| COG2 | CCT2 | GNAS |
| COG3 | CCT3 | GNB2 |
| COG6 | CCT8 | GNPDA1 |
| COG7 | CD2AP | GOLPH3 |
| COG8 | CD3E | GPR68 |
| COIL | CD40 | GPT2 |
| COL4A5 | CD46 | GSK3A |
| COLEC11 | CD53 | GSTA1 |
| COMMD2 | CD68 | GSTA4 |
| COPG2 | CD74 | GSTK1 |
| COQ10A | CDC25B | GSTM3 |
| COQ6 | CDC25C | GTF2H2 |
| COQ9 | CDC34 | HABP4 |
| COX15 | CDC40 | HAGH |
| COX7A1 | CDC42 | HAX1 |
| CREBZF | CDC42EP3 | HDAC8 |
| CREG1 | CDC6 | HELQ |
| CRIM1 | CDCA3 | HIBADH |
| CRIPT | CDCA7 | HINT1 |
| CRYZL1 | CDCA7L | HMGN1 |
| CSDE1 | CDCA8 | HMOX1 |
| CSGALNACT2 | CDH3 | HNRNPA0 |
| CSN2 | CDIPT | HSD17B7 |
| CTCF | CDK1 | HSPB1 |
| CTNNA2 | CDK13 | HTRA2 |
| CTNNAL1 | CDK16 | IFI27L2 |
| CTNND1 | CDK2AP1 | IFT20 |
| CTSC | CDK4 | IGSF9 |
| CTSD | CDK6 | IL10RB |
| CTSO | CDK8 | IL20RA |
| CTSZ | CDKN2D | ILK |
| CTU1 | CDKN3 | ING4 |
| CUL7 | CDON | IPO8 |
| CWC25 | CDS2 | IQCG |
| CWF19L1 | CDV3 | ITGB1BP1 |
| CYP27A1 | CDYL | ITPA |
| CYP2U1 | CDYL2 | ITPRIPL1 |
| CYP4V2 | CENPE | KBTBD3 |
| D2HGDH | CENPF | KCTD1 |
| DACH1 | CENPH | KDM5C |
| DAGLB | CENPI | KEAP1 |
| DAPK3 | CENPJ | KLF15 |
| DBNL | CENPK | KLHL8 |

| | | |
|---|---|---|
| DCAF15 | CENPM | KRAS |
| DCAF5 | CENPN | KRBA2 |
| DCHS1 | CENPP | KRT20 |
| DCLRE1A | CEP164 | LAPTM4A |
| DCP1A | CEP350 | LBR |
| DCPS | CFDP2 | LEMD2 |
| DCTD | CFL2 | LEPROTL1 |
| DDHD1 | CHADL | LIN52 |
| DDHD2 | CHCHD3 | LMF2 |
| DDO | CHD2 | LMO4 |
| DDR1 | CHD9 | LMTK2 |
| DDX17 | CHEK1 | LNX2 |
| DDX47 | CHERP | LRIG1 |
| DDX5 | CHFR | LRRC28 |
| DDX50 | CHKA | LRRC45 |
| DDX58 | CHMP2A | LRRC8A |
| DDX59 | CHMP4B | LRRN3 |
| DDX6 | CIB1 | LSR |
| DEF8 | CITED4 | MAF1 |
| DENND4B | CKAP2 | MAGOH |
| DENND5B | CKAP2L | MAL2 |
| DFFA | CKAP5 | MAPK4 |
| DHPS | CKS1B | MAPK8IP1 |
| DHTKD1 | CLASP2 | MCC |
| DHX29 | CLEC12A | MCCC1 |
| DHX40 | CLEC14A | MEA1 |
| DHX57 | CLEC1A | MED28 |
| DIDO1 | CLEC3A | MED29 |
| DIP2B | CLIC1 | MED4 |
| DIS3 | CLIP1 | METTL6 |
| DIS3L | CLN8 | METTL9 |
| DLAT | CLNS1A | MFSD6 |
| DMAP1 | CLPX | MIF4GD |
| DMXL1 | CLSTN1 | MMADHC |
| DNAJB2 | CLTC | MMGT1 |
| DNAJC16 | CMPK1 | MOCS2 |
| DNAJC18 | CNBP | MORF4L2 |
| DNAJC24 | CNIH4 | MPDU1 |
| DNAJC8 | CNKSR3 | MPPE1 |
| DPCD | CNN1 | MPST |
| DPH5 | CNN3 | MRAP2 |
| DPYD | CNNM2 | MRPL23 |
| DPYSL2 | CNOT7 | MRPL32 |
| DST | CNP | MRPL36 |
| DSTYK | COG3 | MRPL41 |

| | | |
|---|---|---|
| DTD1 | COL1A1 | MRPL45 |
| DTX3 | COL3A1 | MRPS23 |
| DUS4L | COL4A2 | MRPS34 |
| DUSP12 | COLEC12 | MRPS6 |
| DYNC1I2 | COMMD5 | MSTO1 |
| DYNLT3 | COMMD7 | MT1A |
| DYRK1A | COPE | MT2A |
| DZIP1 | COQ10B | MTCH1 |
| DZIP3 | COQ2 | MTF1 |
| E2F3 | COQ3 | MTHFD2 |
| E2F5 | COQ6 | MTMR10 |
| ECD | CORO1A | MVK |
| ECH1 | CORO1C | MXD4 |
| ECHDC1 | COTL1 | MYL12B |
| ECM2 | COX19 | MYPN |
| EEA1 | COX5A | NBR1 |
| EED | CPEB4 | NCK1 |
| EEF1A1 | CPNE3 | NCS1 |
| EEF1G | CPOX | NDUFA8 |
| EEF2 | CPSF1 | NDUFAB1 |
| EGLN2 | CPSF2 | NDUFB11 |
| EHBP1L1 | CRAT | NDUFB7 |
| EID2 | CREB3L1 | NDUFV2 |
| EIF1 | CREB3L2 | NEFH |
| EIF1B | CREBL2 | NEK10 |
| EIF2B5 | CREBZF | NENF |
| EIF2S3 | CRELD2 | NET1 |
| EIF3E | CS | NINJ1 |
| EIF3F | CSDE1 | NMI |
| EIF3L | CSNK2B | NPY1R |
| EIF4A2 | CSTF3 | NSFL1C |
| EIF4B | CTSC | NSL1 |
| EIF4ENIF1 | CTSD | NT5DC1 |
| EIF5A2 | CTSO | NTHL1 |
| ELAC1 | CTSZ | NUBP1 |
| ELF1 | CUL5 | NUDCD1 |
| ELP2 | CWC22 | NUDT1 |
| ELP3 | CWF19L1 | NUDT21 |
| ENC1 | CWF19L2 | NUDT3 |
| ENPP4 | CXCL2 | NUP160 |
| ENTPD7 | CXCR4 | NXT1 |
| EPHB6 | CYB5B | NXT2 |
| ERLIN2 | CYB5R1 | OAZ1 |
| EXD3 | CYC1 | ORMDL2 |
| EXOC4 | CYP27A1 | OSGIN1 |

| | | |
|---|---|---|
| EXOC7 | DAPK3 | OTUB1 |
| EXOSC10 | DAZAP1 | PAFAH1B3 |
| EXT2 | DBF4 | PAQR3 |
| F11R | DBNL | PARD6B |
| FAF2 | DBR1 | PC |
| FAH | DCAF11 | PCBD1 |
| FAIM | DCAF17 | PCDH7 |
| FAM149B1 | DCAF5 | PCMTD1 |
| FAM151B | DCP1A | PCNA |
| FAM20B | DCPS | PCNP |
| FAM53C | DCTD | PDCD2L |
| FANCM | DCTN1 | PDCL |
| FBXL20 | DCTN5 | PDE9A |
| FBXL4 | DDHD1 | PDZD11 |
| FBXL5 | DDIT4 | PEX11A |
| FBXL7 | DDX10 | PEX16 |
| FBXO32 | DDX21 | PEX2 |
| FBXO38 | DDX24 | PGF |
| FBXO4 | DDX27 | PGLS |
| FBXO9 | DDX3X | PGR |
| FBXW4 | DDX41 | PGRMC1 |
| FECH | DDX46 | PHB2 |
| FEM1A | DDX47 | PHF11 |
| FEM1B | DDX52 | PHOSPHO2 |
| FGGY | DECR2 | PIGL |
| FIP1L1 | DEF8 | PIH1D1 |
| FKBP15 | DENND4B | PIN1 |
| FKRP | DEPDC1 | PIR |
| FLOT2 | DGKE | PKP4 |
| FLVCR1 | DGUOK | PLA2G7 |
| FMO5 | DHRS1 | PLCXD2 |
| FNBP1L | DHX15 | PLEKHA6 |
| FNDC1 | DHX29 | PLEKHJ1 |
| FNDC3B | DIAPH3 | PNPLA4 |
| FPGT | DIDO1 | POLE3 |
| FTO | DIP2B | POLK |
| FUCA1 | DIS3 | POLR1D |
| FXYD3 | DIXDC1 | POLR2E |
| FYCO1 | DKC1 | POLR2G |
| FZD9 | DKK3 | POLR2H |
| FZR1 | DKKL1 | POLR2J |
| G2E3 | DLAT | POMT1 |
| GAA | DLGAP5 | PPARA |
| GAB1 | DMAP1 | PPIB |
| GABBR1 | DMXL1 | PPIE |

| | | |
|---|---|---|
| GALM | DNAJA3 | PPIL3 |
| GALNS | DNAJC13 | PPP1CC |
| GALNT2 | DNAJC27 | PPP1R11 |
| GANAB | DNAJC3 | PPP1R7 |
| GATAD2A | DNAJC8 | PPP6C |
| GBA | DNAL1 | PRDX2 |
| GDAP2 | DND1 | PRKAA1 |
| GEM | DNM1L | PRKAR2A |
| GEMIN6 | DOCK6 | PRMT2 |
| GGA3 | DOK2 | PRMT7 |
| GGPS1 | DPCD | PRPF18 |
| GGT7 | DPF2 | PRRG2 |
| GGTA1 | DPY30 | PSEN1 |
| GKAP1 | DPYSL3 | PSENEN |
| GLB1 | DRG1 | PSMA2 |
| GLI3 | DSG2 | PSMA7 |
| GLOD4 | DTX1 | PSMB7 |
| GLT8D1 | DTX3 | PSMC1 |
| GM2A | DTYMK | PSMC3 |
| GMFB | DUS4L | PSMD11 |
| GNA13 | DUSP1 | PSMG3 |
| GNAL | DUSP10 | PSMG4 |
| GNG7 | DUSP14 | PTAR1 |
| GNL3 | DUSP3 | PTMA |
| GNPAT | DUT | PTPN2 |
| GNS | DYNC1LI1 | PTPRS |
| GOLGA2 | DYNLL1 | PXMP2 |
| GOLGA7 | DYNLL2 | QDPR |
| GOLIM4 | DYNLRB1 | RAB11FIP2 |
| GOLPH3L | DYNLT3 | RAB18 |
| GOPC | DYRK1A | RAB38 |
| GORAB | DZIP1 | RAB5C |
| GORASP2 | DZIP3 | RAB8A |
| GOSR2 | E2F1 | RAC3 |
| GPA33 | E4F1 | RAD21 |
| GPAM | EBNA1BP2 | RAE1 |
| GPBP1 | ECT2 | RALY |
| GPC1 | EDEM3 | RANBP1 |
| GPNMB | EDF1 | RAP2A |
| GPR108 | EEA1 | RARRES2 |
| GPR137B | EED | RBBP4 |
| GPR18 | EFEMP2 | RBBP9 |
| GPR68 | EFHD2 | RBM18 |
| GPRASP1 | EFNA2 | RBM42 |
| GPX3 | EGLN2 | RBMS3 |

| | | |
|---|---|---|
| GPX8 | EHBP1L1 | RBX1 |
| GRIK5 | EHD1 | RCCD1 |
| GRINA | EHMT1 | RCE1 |
| GTDC1 | EI24 | RCN2 |
| GTF2I | EIF2AK2 | RELL1 |
| GTF2IRD1 | EIF2B2 | RER1 |
| GTPBP4 | EIF2B5 | RGMB |
| GUF1 | EIF3I | RNASEH2B |
| GZF1 | EIF4A1 | RNF135 |
| GZMA | EIF4A3 | RNF167 |
| HABP4 | EIF4E3 | RNF187 |
| HACE1 | EIF4G3 | RNF5 |
| HBP1 | EIF5B | RNFT1 |
| HCN1 | ELL | RP2 |
| HEATR5B | ELOVL5 | RPL13 |
| HERC2 | ELOVL6 | RPL13A |
| HLCS | EMILIN1 | RPLP2 |
| HMGCL | EMP2 | RPS26 |
| HMGN3 | EMX2 | RTN3 |
| HMGN4 | ENAH | RYBP |
| HNF4A | ENC1 | S100A2 |
| HNMT | ENPEP | SAMM50 |
| HNRNPH3 | ENPP1 | SAP18 |
| HPS3 | ENY2 | SCAMP1 |
| HS1BP3 | EPB41L2 | SCCPDH |
| HS6ST1 | EPHA2 | SCN2A |
| HSD11B1L | EPHB4 | SDHC |
| HSD17B8 | EPM2A | SEC11A |
| HSF1 | ERN1 | SEC13 |
| HSPBAP1 | ESF1 | SEC61A2 |
| HUWE1 | ESPL1 | SEC61B |
| IARS2 | EXO1 | SEC61G |
| IBTK | EXOC1 | SEC62 |
| IDUA | EXOC5 | SECTM1 |
| IER3 | EXOG | SERPINB6 |
| IER3IP1 | EXOSC4 | SH3BP4 |
| IFNAR1 | EXOSC8 | SHMT2 |
| IFNGR1 | EXTL2 | SIRT2 |
| IFT122 | EZH2 | SKA3 |
| IFT52 | F12 | SKP2 |
| IFT74 | F2RL1 | SLC12A9 |
| IFT88 | FABP4 | SLC25A1 |
| IGBP1 | FABP5 | SLC25A11 |
| IGDCC4 | FADD | SLC25A38 |
| IGFBP7 | FADS1 | SLC30A6 |

| | | |
|---|---|---|
| IK | FADS3 | SLC30A7 |
| IL10RB | FAF2 | SLC35C2 |
| IL13RA1 | FAH | SLC39A14 |
| IL4R | FAM110B | SLC44A2 |
| INPP4A | FAM151B | SNF8 |
| INTS8 | FAM20B | SNRPB |
| IP6K1 | FAM32A | SNRPC |
| IPP | FAM72A | SNRPD3 |
| IQCB1 | FAM83D | SPAST |
| IQGAP1 | FANCD2 | SPATA24 |
| IQGAP2 | FANCE | SPIN1 |
| IREB2 | FANCL | SRGN |
| IRF8 | FAR1 | SRPK2 |
| IST1 | FARSA | SS18 |
| ITFG2 | FARSB | SSR1 |
| ITGB5 | FASTKD1 | SSR2 |
| ITM2B | FAU | SSR3 |
| ITM2C | FBF1 | ST6GALNAC2 |
| IVD | FBLIM1 | STK16 |
| IWS1 | FBLN2 | STK35 |
| JAKMIP2 | FBXL14 | STOM |
| JAM3 | FBXL4 | STX18 |
| JAZF1 | FBXO32 | STX4 |
| JTB | FBXO5 | SUCLG2 |
| KCMF1 | FBXW2 | SUPT4H1 |
| KCNG2 | FCGRT | SUV39H2 |
| KCNIP4 | FDPS | SYNE2 |
| KCNRG | FEM1B | SYT1 |
| KCTD1 | FEN1 | TALDO1 |
| KDM2A | FERMT2 | TAX1BP3 |
| KDM6A | FGD4 | TBC1D1 |
| KIF3B | FGD5 | TBC1D8 |
| KITLG | FGFR1OP2 | TBCA |
| KLF13 | FGL2 | TBCB |
| KLHDC2 | FHL3 | TBCC |
| KLHL11 | FHOD3 | THAP11 |
| KLHL18 | FIGNL1 | THOC3 |
| KLHL2 | FKBP11 | THOC5 |
| KLHL23 | FKBP15 | THOC7 |
| KLHL26 | FLI1 | THUMPD3 |
| KLHL28 | FLNA | TIMM50 |
| KLHL9 | FLNB | TLE3 |
| KPNA4 | FLVCR1 | TMBIM1 |
| KRCC1 | FOS | TMBIM6 |
| KRI1 | FOXA3 | TMED2 |

| | | |
|---|---|---|
| KRT19 | FOXK2 | TMEM128 |
| LAG3 | FSTL3 | TMEM147 |
| LAMA1 | FTO | TMEM160 |
| LAMP1 | FUS | TMEM167B |
| LAP3 | FZR1 | TMEM208 |
| LARP6 | G3BP1 | TMEM209 |
| LARS2 | GABRA2 | TMEM30A |
| LAS1L | GALE | TMEM37 |
| LAT | GALNT4 | TMEM43 |
| LATS2 | GAS8 | TMEM47 |
| LCOR | GATAD2A | TMEM50A |
| LDHB | GBP6 | TMEM50B |
| LDOC1 | GCH1 | TMEM60 |
| LEPROT | GCSH | TMEM87B |
| LGALS9 | GDAP2 | TMEM9B |
| LGMN | GDPD3 | TMPO |
| LIAS | GDPD5 | TMTC4 |
| LIG4 | GFER | TMUB1 |
| LIMA1 | GGA3 | TMX1 |
| LINGO2 | GGCT | TNNC1 |
| LMAN2L | GGNBP2 | TNPO3 |
| LNX2 | GIMAP6 | TOMM22 |
| LONP2 | GIMAP7 | TOR1B |
| LPCAT4 | GINS1 | TPM3 |
| LRBA | GINS4 | TPST1 |
| LRP10 | GJA1 | TRAM2 |
| LRRC10B | GK | TRAPPC5 |
| LRRC42 | GLCCI1 | TRIM24 |
| LRRC47 | GLOD4 | TRPT1 |
| LRRC4C | GLRX3 | TRUB2 |
| LRRC8D | GLT8D2 | TSC22D3 |
| LRRFIP1 | GLUD1 | TSN |
| LRRTM2 | GLUL | TSPAN14 |
| LTB | GMFG | TSPAN31 |
| LTBP3 | GMNN | TSPAN6 |
| LTBR | GNA11 | TTC9C |
| LTV1 | GNAI2 | TUBB2B |
| LUC7L2 | GNAS | TWSG1 |
| LY9 | GNB1 | TXNDC9 |
| LYRM1 | GNG7 | TYW3 |
| LYRM2 | GNL3 | UBAC2 |
| LYSMD1 | GNPDA1 | UBE2M |
| LZTFL1 | GNS | UBE2R2 |
| M6PR | GOLGA2 | UBE2S |
| MAD2L1BP | GOLGB1 | UBE2T |

| | | |
|---|---|---|
| MADD | GOPC | UBL3 |
| MAL2 | GOSR1 | UBQLN4 |
| MAML2 | GP1BA | UBTD2 |
| MAMLD1 | GPATCH4 | UBXN1 |
| MAN1C1 | GPATCH8 | UPK1B |
| MAN2A2 | GPNMB | VAMP3 |
| MAN2B1 | GPR137B | VAMP8 |
| MAN2C1 | GPR162 | VDAC2 |
| MAP3K4 | GPR63 | VEGFB |
| MAP3K7 | GPR85 | VEZF1 |
| MAP4K2 | GPT2 | VPS18 |
| MAP4K3 | GPX8 | VPS25 |
| MAP7 | GRIPAP1 | VPS37A |
| MAPK13 | GRPEL1 | VPS72 |
| MAPK8IP3 | GRWD1 | WAC |
| MAPKAP1 | GSDMB | WBP1 |
| MAPKBP1 | GSK3A | WBP2 |
| MARK1 | GSTK1 | WDFY1 |
| MARK3 | GSTO1 | WDR83 |
| MARK4 | GTF2I | WHAMM |
| MARS2 | GTF2IRD1 | XPO1 |
| MARVELD1 | GTPBP4 | YARS2 |
| MARVELD3 | GZF1 | YEATS4 |
| MBD3 | HABP4 | YIPF6 |
| MBD5 | HACE1 | YTHDF3 |
| MBLAC2 | HAGH | YWHAE |
| MBTPS1 | HAT1 | YWHAZ |
| MBTPS2 | HAUS1 | ZCRB1 |
| MCF2L | HAUS4 | ZDHHC1 |
| MCTS1 | HAUS8 | ZFAND2B |
| MDM2 | HDAC10 | ZFPL1 |
| MEAF6 | HDAC5 | ZGPAT |
| MED12 | HDAC8 | ZMYM1 |
| MED13 | HDDC2 | ZNF148 |
| MED25 | HEATR5B | ZNF391 |
| MED9 | HELLS | ZNF420 |
| MEF2A | HERC2 | ZNF536 |
| MEX3A | HEXIM2 | ZNHIT1 |
| MFAP3 | HEY2 | ZNHIT3 |
| MFN1 | HINFP | ZWINT |
| MFSD10 | HMGB1 | ZYG11B |
| MFSD8 | HMGN3 | |
| MFSD9 | HMGN5 | |
| MGA | HMOX1 | |
| MGRN1 | HNRNPA3 | |

| | |
|---|---|
| MGST3 | HNRNPAB |
| MICALL1 | HNRNPD |
| MIDN | HNRNPF |
| MKKS | HNRNPK |
| MKX | HNRNPL |
| MLLT3 | HNRNPM |
| MLST8 | HNRNPUL2 |
| MMAA | HOMER1 |
| MON1B | HOXC4 |
| MORN4 | HPS3 |
| MOSPD1 | HSDL1 |
| MPP1 | HSF2 |
| MR1 | HSPA12B |
| MRPL1 | HSPB1 |
| MRPL3 | HSPG2 |
| MRPL4 | HTATSF1 |
| MRPL51 | HTR1F |
| MRPL9 | IARS2 |
| MRPS22 | IBTK |
| MRPS9 | ICAM2 |
| MSRB2 | ICOSLG |
| MSRB3 | IER2 |
| MST1 | IER3 |
| MTCP1 | IFNAR1 |
| MTHFD2L | IFNAR2 |
| MTHFSD | IFT20 |
| MTIF3 | IL13RA1 |
| MTM1 | IL1A |
| MTMR2 | IL20RA |
| MTO1 | IL27RA |
| MTTP | ILF2 |
| MXRA8 | ILF3 |
| MYL6B | ILK |
| MYL9 | ILVBL |
| MYLK | INPP5D |
| MYOF | INTS2 |
| MYOZ1 | IPO4 |
| N4BP2L1 | IPO8 |
| NAA16 | IQGAP1 |
| NAA30 | IQGAP3 |
| NAA38 | IRAK2 |
| NAAA | IREB2 |
| NACA | IRS1 |
| NACAD | ISG20 |
| NAP1L5 | ISYNA1 |

| | |
|---|---|
| NAPEPLD | ITPK1 |
| NBL1 | ITPRIPL1 |
| NBN | IVD |
| NCK2 | IWS1 |
| NCOA7 | JAG1 |
| NCOR1 | JAM3 |
| NDEL1 | JARID2 |
| NDOR1 | JMJD7 |
| NDUFC2 | JUN |
| NDUFS5 | KCNJ14 |
| NECAP2 | KCNRG |
| NEIL2 | KCTD10 |
| NEK3 | KCTD11 |
| NEU3 | KCTD12 |
| NFE2L1 | KCTD20 |
| NFE2L3 | KCTD9 |
| NFKBIZ | KDM1B |
| NFS1 | KDM2A |
| NGRN | KHDRBS1 |
| NID1 | KIF11 |
| NIT2 | KIF15 |
| NKAPL | KIF18A |
| NLK | KIF20A |
| NMD3 | KIF20B |
| NME6 | KIF22 |
| NME7 | KIF23 |
| NMNAT1 | KIF2C |
| NOL12 | KIF4A |
| NOL8 | KIF7 |
| NOTCH2 | KIT |
| NPEPPS | KLF4 |
| NPHP1 | KLF6 |
| NR3C2 | KLHL18 |
| NR5A2 | KLHL2 |
| NRIP1 | KLHL21 |
| NRP1 | KLHL32 |
| NSD1 | KPNA4 |
| NT5M | KPNB1 |
| NUB1 | KRAS |
| NUDT12 | KRT18 |
| NUDT13 | KRT19 |
| NUDT3 | KRT8 |
| NUP133 | KTN1 |
| NUP153 | LAG3 |
| NUP88 | LAMC1 |

| | |
|---|---|
| OCIAD1 | LAP3 |
| OFD1 | LAPTM5 |
| OPA1 | LARP6 |
| ORAI3 | LAS1L |
| OSBPL9 | LASP1 |
| OTUD5 | LATS2 |
| PABPN1 | LBR |
| PACRGL | LCK |
| PAM | LCOR |
| PAN3 | LCORL |
| PANK4 | LCP2 |
| PAQR3 | LDOC1 |
| PARP1 | LENG8 |
| PARP11 | LFNG |
| PARP14 | LGALS3 |
| PBXIP1 | LGALS8 |
| PCBD1 | LIMK1 |
| PCCB | LIN7B |
| PCF11 | LIN7C |
| PCGF2 | LIPA |
| PDCD6 | LIPT2 |
| PDCD6IP | LLPH |
| PDCD7 | LMAN2 |
| PDHA1 | LMNA |
| PDHB | LMNB1 |
| PDK1 | LMO2 |
| PDK4 | LNX2 |
| PDRG1 | LPP |
| PDXDC1 | LRP6 |
| PDZRN3 | LRRC1 |
| PEBP1 | LRRC10B |
| PECR | LRRC25 |
| PEG3 | LRRC59 |
| PELO | LRRC66 |
| PEX11B | LSG1 |
| PEX13 | LSM4 |
| PEX5 | LSM5 |
| PFDN5 | LSM7 |
| PFKFB2 | LSR |
| PFKM | LTBP1 |
| PGAP3 | LTV1 |
| PGPEP1 | LUC7L3 |
| PGRMC1 | LXN |
| PGRMC2 | LY9 |
| PHAX | LYAR |

| | |
|---|---|
| PHC3 | LYN |
| PHLDB1 | LYPLA1 |
| PHOSPHO2 | LYZ |
| PHYH | LZTS2 |
| PI4K2A | MAD2L1 |
| PI4K2B | MAD2L1BP |
| PIAS3 | MAFB |
| PICALM | MAN1C1 |
| PIGN | MAN2A2 |
| PIGO | MANEAL |
| PIGP | MANF |
| PIGS | MAP2 |
| PIGT | MAP2K3 |
| PIGY | MAP3K7 |
| PIK3R1 | MAP4K5 |
| PIK3R4 | MAP7 |
| PIN4 | MAP7D1 |
| PINK1 | MAPK14 |
| PIP5K1A | MAPK8 |
| PKD2 | MARK3 |
| PKIA | MARVELD2 |
| PLA2G6 | MARVELD3 |
| PLAA | MAST3 |
| PLBD2 | MBD5 |
| PLCE1 | MBP |
| PLCL2 | MCC |
| PLEKHA5 | MCF2L |
| PLEKHG1 | MCM2 |
| PLSCR5 | MCM3 |
| PMP22 | MCM4 |
| PNPLA3 | MCM5 |
| POGK | MCM6 |
| POLA1 | MDFIC |
| POLDIP2 | MECP2 |
| POLE4 | MED11 |
| POLG | MED27 |
| POLM | MED28 |
| POLR1C | MED9 |
| POLR1E | MEF2A |
| POLR2D | MEF2C |
| POLR3GL | MEIS2 |
| PON2 | MELK |
| PPAT | MEN1 |
| PPHLN1 | MERTK |
| PPM1A | MEST |

| | |
|---|---|
| PPM1K | METRNL |
| PPP1R12C | METTL1 |
| PPP1R15B | METTL3 |
| PPP1R3D | METTL9 |
| PPP2R1B | MEX3C |
| PPP2R5A | MFGE8 |
| PPWD1 | MGAT2 |
| PRCP | MGAT5 |
| PRDM1 | MGST1 |
| PRMT2 | MICALL1 |
| PRODH | MID1 |
| PRPF39 | MIF4GD |
| PRR15 | MIIP |
| PRSS35 | MIS18A |
| PSIP1 | MITD1 |
| PSKH1 | MKKS |
| PSME4 | MKNK2 |
| PTCD2 | MKX |
| PTCH1 | MLH1 |
| PTEN | MMAB |
| PTGFRN | MMD |
| PTP4A3 | MMGT1 |
| PTPN3 | MMP17 |
| PWP1 | MMP19 |
| QSOX1 | MNAT1 |
| R3HDM2 | MND1 |
| RAB11B | MORF4L2 |
| RAB22A | MOSPD1 |
| RAB24 | MPEG1 |
| RAB33B | MPHOSPH8 |
| RAB3GAP1 | MPP1 |
| RAB3GAP2 | MR1 |
| RAB4A | MRAS |
| RAB5A | MRPL1 |
| RAB6A | MRPL14 |
| RABEP1 | MRPL15 |
| RABIF | MRPL17 |
| RAC2 | MRPL36 |
| RAD17 | MRPL38 |
| RALYL | MRPL39 |
| RANBP10 | MRPL4 |
| RANBP9 | MRPL45 |
| RAP2B | MRPL49 |
| RARRES1 | MRPS10 |
| RARS2 | MRPS11 |

| | |
|---|---|
| RASSF1 | MRPS15 |
| RAVER1 | MRPS23 |
| RB1 | MRPS34 |
| RB1CC1 | MRPS35 |
| RBBP9 | MRPS5 |
| RBM12B | MSN |
| RBM17 | MT1A |
| RBM22 | MT2A |
| RBM25 | MTDH |
| RBM5 | MTF2 |
| RBX1 | MTHFD2 |
| RCHY1 | MTMR1 |
| RCL1 | MTMR11 |
| RCN3 | MTMR4 |
| REEP2 | MTMR6 |
| REEP5 | MTRR |
| RELB | MTX3 |
| REPS2 | MVK |
| RERG | MVP |
| REV1 | MXRA7 |
| RFNG | MYL12B |
| RFTN1 | MYL9 |
| RHBDD1 | MYLK |
| RHOQ | MYO1C |
| RIC3 | MYO1D |
| RIC8A | MYO5A |
| RIMKLA | MYO6 |
| RINT1 | MYO9A |
| RIOK2 | MYO9B |
| RIOK3 | NAA10 |
| RIT1 | NAA15 |
| RLF | NAA25 |
| RNF10 | NAAA |
| RNF113A | NACAD |
| RNF114 | NAE1 |
| RNF115 | NAGA |
| RNF13 | NAGK |
| RNF135 | NAIP |
| RNF139 | NANS |
| RNF149 | NAV1 |
| RNF20 | NBN |
| RNF213 | NCAPD2 |
| RNF215 | NCAPG |
| RNF40 | NCAPH |
| RNF7 | NCBP1 |

| | |
|---|---|
| RNLS | NCF2 |
| ROBO1 | NCKAP5L |
| ROGDI | NCKIPSD |
| RPAP3 | NCLN |
| RPF2 | NDC80 |
| RPL10A | NDEL1 |
| RPL12 | NDRG1 |
| RPL15 | NDUFB9 |
| RPL21 | NDUFV1 |
| RPL22L1 | NEK3 |
| RPL23 | NEK8 |
| RPL26 | NET1 |
| RPL27A | NFATC3 |
| RPL3 | NFKB1 |
| RPL31 | NFKBIB |
| RPL32 | NFKBIZ |
| RPL36 | NFS1 |
| RPL36A | NFX1 |
| RPL37 | NFYC |
| RPL4 | NHLRC1 |
| RPL5 | NHLRC3 |
| RPL6 | NID1 |
| RPL7L1 | NIT2 |
| RPL9 | NLK |
| RPRD2 | NMD3 |
| RPS12 | NMI |
| RPS15A | NOC2L |
| RPS18 | NOP14 |
| RPS24 | NOP16 |
| RPS27 | NOP2 |
| RPS3A | NOP56 |
| RPS6KA4 | NOP58 |
| RPS7 | NOS2 |
| RPUSD1 | NOTCH1 |
| RRAGB | NOTCH2 |
| RRP12 | NPHP1 |
| RRP1B | NPHP3 |
| RSF1 | NPLOC4 |
| RTN4 | NR4A1 |
| RTN4IP1 | NR5A2 |
| RTTN | NSMAF |
| RWDD1 | NT5DC1 |
| RWDD2B | NTAN1 |
| S100A4 | NTRK1 |
| SALL2 | NUDCD1 |

| | |
|---|---|
| SCAMP2 | NUDT12 |
| SCARB2 | NUDT21 |
| SCD5 | NUDT9 |
| SCFD1 | NUF2 |
| SCG5 | NUMB |
| SCPEP1 | NUP160 |
| SCYL3 | NUP188 |
| SDR39U1 | NUP214 |
| SEC24A | NUP54 |
| SEC24B | NUP85 |
| SEC63 | NUP88 |
| SEMA4D | NUP93 |
| SEMA4G | NUSAP1 |
| SENP7 | NXT1 |
| SERINC3 | NXT2 |
| SERPINI1 | OCEL1 |
| SETMAR | OIP5 |
| SETX | OLFML1 |
| SF1 | OPA1 |
| SF3A1 | ORAI1 |
| SF3B1 | ORMDL2 |
| SFT2D2 | ORMDL3 |
| SFXN2 | OSBPL3 |
| SGCE | OSBPL9 |
| SGK3 | OSGEPL1 |
| SH3GL1 | OSTC |
| SH3YL1 | OSTM1 |
| SHOC2 | OTUB1 |
| SIAE | OTUD3 |
| SIAH1 | OTUD5 |
| SIK2 | OXNAD1 |
| SIPA1 | OXSR1 |
| SIPA1L1 | PABPC5 |
| SIRT1 | PACRGL |
| SIRT5 | PACSIN3 |
| SLC25A26 | PAFAH1B3 |
| SLC25A28 | PAK4 |
| SLC25A30 | PAM |
| SLC25A35 | PANK3 |
| SLC25A37 | PARP1 |
| SLC25A38 | PATL1 |
| SLC26A11 | PAXIP1 |
| SLC35A1 | PBK |
| SLC38A7 | PBXIP1 |
| SLC39A9 | PC |

| | |
|---|---|
| SLC40A1 | PCCB |
| SLC41A1 | PCGF3 |
| SLC43A2 | PCMTD1 |
| SLC44A3 | PCNA |
| SLC9A3R1 | PCNP |
| SLC9A9 | PDAP1 |
| SLCO2B1 | PDCD2L |
| SLCO3A1 | PDCD6IP |
| SLITRK2 | PDCL |
| SMAP1 | PDE4B |
| SMC6 | PDHA1 |
| SMG6 | PDIA4 |
| SMG7 | PDIA6 |
| SMYD4 | PDK1 |
| SNAP23 | PDLIM1 |
| SNAPIN | PDLIM7 |
| SNX18 | PDS5A |
| SNX21 | PDXP |
| SNX24 | PDZD8 |
| SNX30 | PDZRN3 |
| SNX5 | PERP |
| SNX8 | PEX19 |
| SOAT1 | PEX2 |
| SOD1 | PEX6 |
| SOD2 | PFDN1 |
| SON | PFKM |
| SORBS2 | PFKP |
| SORL1 | PFN1 |
| SPA17 | PGAP3 |
| SPAG9 | PGF |
| SPATA2 | PGGT1B |
| SPATA20 | PGLS |
| SPG11 | PGRMC1 |
| SPG7 | PGS1 |
| SPINT2 | PHAX |
| SPR | PHB2 |
| SPTAN1 | PHGDH |
| SPTBN1 | PI16 |
| SRP72 | PI4K2A |
| SSPN | PIAS3 |
| SSX2IP | PIF1 |
| ST3GAL6 | PIH1D1 |
| ST6GALNAC6 | PIK3C2A |
| STAG1 | PJA1 |
| STAMBP | PKMYT1 |

| | |
|---|---|
| STARD3NL | PLAU |
| STARD9 | PLBD2 |
| STAT2 | PLCB4 |
| STAT3 | PLCD3 |
| STAU1 | PLEK |
| STK36 | PLEKHM2 |
| STK38L | PLK4 |
| STRBP | PLP2 |
| STRN3 | PMM2 |
| STX12 | PMP22 |
| STX16 | PMPCA |
| STX1B | PNKP |
| STX6 | PNO1 |
| STXBP3 | PNPLA4 |
| STYX | PNPLA8 |
| SUB1 | PNPT1 |
| SUMF1 | POGK |
| SUPT6H | POLDIP2 |
| SUPV3L1 | POLE2 |
| SUSD1 | POLE3 |
| SYNE2 | POLM |
| SYNJ2BP | POLR1C |
| SYT9 | POLR1D |
| SYVN1 | POLR2C |
| TADA3 | POLR2E |
| TAF5L | POLR2L |
| TAF7 | POLR3E |
| TARBP1 | POLR3GL |
| TARDBP | POMP |
| TASP1 | POT1 |
| TATDN1 | PPARG |
| TBC1D12 | PPAT |
| TBC1D13 | PPCDC |
| TBC1D17 | PPHLN1 |
| TBC1D19 | PPIA |
| TBC1D20 | PPIB |
| TBC1D9B | PPIG |
| TBL2 | PPIH |
| TBXAS1 | PPIL3 |
| TCAP | PPP1CC |
| TCEA3 | PPP1R2 |
| TCEAL1 | PPP1R8 |
| TCEAL4 | PPP2R5C |
| TCEAL8 | PPRC1 |
| TCF7L2 | PPTC7 |

| | |
|---|---|
| TCOF1 | PRC1 |
| TCP11 | PRCP |
| TCP11L2 | PRDX2 |
| TCTA | PREX1 |
| TDRD3 | PRKAA1 |
| TDRD7 | PRKCD |
| TERF2 | PRMT1 |
| TEX2 | PRPF4B |
| TFAM | PRPF6 |
| TFB2M | PRPSAP2 |
| TFCP2 | PRR12 |
| TFEB | PSMA7 |
| TFIP11 | PSMB1 |
| TGDS | PSMB5 |
| TGIF2 | PSMB9 |
| TGS1 | PSMC1 |
| THAP6 | PSMC6 |
| THOC1 | PSMD10 |
| THRA | PSMD11 |
| THTPA | PSMD2 |
| TIA1 | PSME4 |
| TIAL1 | PSMG2 |
| TK2 | PSPC1 |
| TLE1 | PSPH |
| TLK1 | PTCD3 |
| TLK2 | PTEN |
| TLR3 | PTGER2 |
| TLR6 | PTGES3 |
| TM2D1 | PTK7 |
| TM9SF4 | PTMA |
| TMCO6 | PTPDC1 |
| TMED1 | PTPN11 |
| TMED2 | PTPRS |
| TMED4 | PTTG1 |
| TMEM117 | PUM2 |
| TMEM175 | PUS1 |
| TMEM183A | PUSL1 |
| TMEM184C | PWWP2A |
| TMEM192 | PYCR1 |
| TMEM199 | QRSL1 |
| TMEM229B | QTRT1 |
| TMEM25 | RAB11FIP2 |
| TMEM39A | RAB13 |
| TMEM51 | RAB22A |
| TMEM59 | RAB35 |

| | |
|---|---|
| TMEM87A | RAB3GAP2 |
| TMEM9B | RAB5C |
| TMLHE | RAB6A |
| TNKS | RAB8A |
| TNKS1BP1 | RAB8B |
| TOP2B | RABEP1 |
| TP53RK | RAC1 |
| TPD52 | RAC2 |
| TPD52L2 | RACGAP1 |
| TPK1 | RAD21 |
| TPT1 | RAD23B |
| TRA2A | RAD51AP1 |
| TRAM1L1 | RALBP1 |
| TRANK1 | RALY |
| TRAPPC6B | RANBP1 |
| TRIB2 | RANBP10 |
| TRIM2 | RANBP9 |
| TRIM3 | RAPGEF1 |
| TRIM41 | RARRES1 |
| TRIM62 | RARRES2 |
| TRIM68 | RASA1 |
| TRIP11 | RASA2 |
| TRMT12 | RASD1 |
| TRMT61B | RASSF1 |
| TRPS1 | RASSF5 |
| TSNAX | RAVER1 |
| TSPYL1 | RBBP4 |
| TSPYL4 | RBM17 |
| TTC17 | RBM18 |
| TTC21B | RBM3 |
| TTC5 | RBM43 |
| TTC7B | RBM8A |
| TTLL1 | RBMS1 |
| TUBGCP4 | RCC2 |
| TUSC3 | RCL1 |
| TUT1 | RCOR2 |
| TXNDC15 | REEP2 |
| TXNL4B | REEP4 |
| TYRO3 | REEP6 |
| U2AF2 | REL |
| UACA | RELB |
| UBA5 | RELN |
| UBA7 | REPIN1 |
| UBAC2 | RERE |
| UBAP2 | REXO2 |

| | |
|---|---|
| UBE2A | RFC2 |
| UBE2B | RFC4 |
| UBE2D4 | RFESD |
| UBE2E2 | RFK |
| UBE2E3 | RFNG |
| UBE2V1 | RFXANK |
| UBE3B | RGS1 |
| UBIAD1 | RGS19 |
| UBQLN2 | RHOG |
| UBR3 | RILPL1 |
| UBR4 | RIOK2 |
| UBR5 | RIOK3 |
| UBTD1 | RIT1 |
| UBXN2B | RNASEH2A |
| UBXN8 | RNASEH2B |
| UCK1 | RNF10 |
| ULK3 | RNF135 |
| UNC119 | RNF14 |
| UNC13D | RNF187 |
| URB2 | RNF19B |
| USO1 | RNF20 |
| USP24 | RNF216 |
| USP25 | RNF25 |
| USP33 | RNF40 |
| USP34 | RNF44 |
| USP38 | RNFT1 |
| USP40 | RNGTT |
| USP7 | RNH1 |
| USP9X | ROM1 |
| USPL1 | RPL7L1 |
| UTP14A | RPP40 |
| UTP6 | RPUSD2 |
| VAMP5 | RRAGB |
| VEGFC | RRAGC |
| VEZT | RRM2 |
| VPS13D | RRP12 |
| VPS33A | RRS1 |
| VPS39 | RSL1D1 |
| VPS45 | RSPH6A |
| VPS4A | RSRC2 |
| VPS52 | RUVBL2 |
| VPS8 | RWDD2B |
| VSIG10 | RYBP |
| WDR11 | S100A16 |
| WDR36 | S100A2 |

| | |
|---|---|
| WDR6 | S100A6 |
| WDR81 | S1PR2 |
| XDH | SAMD9 |
| XPO1 | SAP30 |
| XRCC6 | SBDS |
| YARS2 | SCAMP1 |
| ZBED5 | SCARF1 |
| ZBTB38 | SCN9A |
| ZBTB4 | SCO1 |
| ZBTB44 | SCP2 |
| ZC3H13 | SCYL3 |
| ZC3H6 | SDCCAG8 |
| ZCCHC7 | SDF2L1 |
| ZDHHC14 | SEC13 |
| ZDHHC17 | SEC24C |
| ZFAND1 | SEC24D |
| ZFP1 | SEC61A2 |
| ZFP2 | SEC61G |
| ZFX | SEC63 |
| ZFYVE27 | SECTM1 |
| ZKSCAN5 | SEH1L |
| ZMAT1 | SEMA4G |
| ZMIZ2 | SENP2 |
| ZMYM3 | SENP3 |
| ZMYM5 | SENP6 |
| ZNF12 | SENP7 |
| ZNF131 | SERTAD3 |
| ZNF133 | SET |
| ZNF148 | SETMAR |
| ZNF180 | SF3A3 |
| ZNF184 | SF3B1 |
| ZNF19 | SF3B3 |
| ZNF197 | SF3B4 |
| ZNF2 | SFPQ |
| ZNF212 | SFXN2 |
| ZNF219 | SH3BP4 |
| ZNF22 | SH3BP5 |
| ZNF239 | SH3GL1 |
| ZNF281 | SH3GLB2 |
| ZNF292 | SHC1 |
| ZNF3 | SHISA4 |
| ZNF317 | SHMT1 |
| ZNF329 | SIAH1 |
| ZNF354A | SIGMAR1 |
| ZNF358 | SIK2 |

| | |
|---|---|
| ZNF383 | SIRT2 |
| ZNF398 | SIRT5 |
| ZNF407 | SKA3 |
| ZNF408 | SKP2 |
| ZNF48 | SLC10A3 |
| ZNF512 | SLC11A1 |
| ZNF526 | SLC11A2 |
| ZNF565 | SLC15A4 |
| ZNF568 | SLC16A1 |
| ZNF575 | SLC16A11 |
| ZNF576 | SLC16A7 |
| ZNF613 | SLC17A5 |
| ZNF638 | SLC19A1 |
| ZNF644 | SLC20A2 |
| ZNF672 | SLC22A23 |
| ZNF677 | SLC25A17 |
| ZNF687 | SLC25A37 |
| ZNF784 | SLC25A38 |
| ZNF789 | SLC27A3 |
| ZNF813 | SLC27A4 |
| ZNF821 | SLC29A3 |
| ZNFX1 | SLC2A2 |
| ZRANB2 | SLC30A7 |
| ZSCAN16 | SLC35B1 |
| ZXDC | SLC37A3 |
| ZZZ3 | SLC39A10 |
| | SLC39A14 |
| | SLC41A1 |
| | SLC43A1 |
| | SLC45A4 |
| | SLC7A5 |
| | SLC9A6 |
| | SLC9A9 |
| | SLMAP |
| | SMAGP |
| | SMAP |
| | SMAP1 |
| | SMARCA4 |
| | SMC2 |
| | SMC4 |
| | SMG6 |
| | SMG7 |
| | SMPD4 |
| | SMU1 |
| | SNAI1 |

| |
|---|
| SNF8 |
| SNRPB |
| SNRPC |
| SNTB1 |
| SNW1 |
| SNX10 |
| SNX16 |
| SNX18 |
| SNX27 |
| SNX30 |
| SOBP |
| SOCS1 |
| SON |
| SPAG5 |
| SPATA6 |
| SPC24 |
| SPC25 |
| SPCS3 |
| SPI1 |
| SPOP |
| SQSTM1 |
| SREBF2 |
| SRGN |
| SRI |
| SRP72 |
| SRPK1 |
| SRRD |
| SRRM2 |
| SS18 |
| SS18L1 |
| SSBP1 |
| SSBP3 |
| SSR2 |
| SSRP1 |
| SSU72 |
| SSX2IP |
| ST6GALNAC2 |
| STAB1 |
| STAG1 |
| STAMBP |
| STAR |
| STARD3NL |
| STAT2 |
| STAU2 |
| STEAP1 |

| |
|---|
| STIL |
| STIP1 |
| STK16 |
| STK38L |
| STMN1 |
| STOML2 |
| STRN3 |
| STUB1 |
| STX12 |
| STX2 |
| STX4 |
| SUB1 |
| SUCLG2 |
| SUFU |
| SUPT6H |
| SUV39H1 |
| SUV39H2 |
| SUZ12 |
| SYCE2 |
| SYNCRIP |
| SYNRG |
| SYT1 |
| TACC3 |
| TAF15 |
| TARBP1 |
| TARBP2 |
| TARDBP |
| TASP1 |
| TAX1BP3 |
| TBC1D14 |
| TBC1D9 |
| TBCA |
| TBCB |
| TBCC |
| TBK1 |
| TBRG4 |
| TBXAS1 |
| TCEA3 |
| TCERG1 |
| TCF20 |
| TCF25 |
| TCF7 |
| TCOF1 |
| TCP1 |
| TDG |

| |
|---|
| TES |
| TET3 |
| TEX264 |
| TFB2M |
| TFRC |
| TGS1 |
| THOC3 |
| THOP1 |
| THUMPD3 |
| THYN1 |
| TIA1 |
| TIMM10 |
| TIMM17A |
| TINF2 |
| TK2 |
| TLR2 |
| TM4SF1 |
| TM7SF3 |
| TMCC3 |
| TMCO6 |
| TMED7 |
| TMEM120A |
| TMEM128 |
| TMEM143 |
| TMEM144 |
| TMEM147 |
| TMEM14C |
| TMEM150A |
| TMEM150B |
| TMEM165 |
| TMEM170A |
| TMEM192 |
| TMEM201 |
| TMEM209 |
| TMEM231 |
| TMEM51 |
| TMEM60 |
| TMEM65 |
| TMEM67 |
| TMEM87A |
| TMOD3 |
| TMPO |
| TMTC4 |
| TNKS1BP1 |
| TNRC6B |

| |
|---|
| TOMM20 |
| TOMM34 |
| TOP1 |
| TOP2A |
| TOPBP1 |
| TP53BP1 |
| TP53RK |
| TPM2 |
| TPM3 |
| TPM4 |
| TPP2 |
| TPX2 |
| TRAF1 |
| TRAF3IP2 |
| TRAF7 |
| TREM2 |
| TRIM23 |
| TRIM24 |
| TRIM26 |
| TRIO |
| TRIP11 |
| TRIP12 |
| TRMT6 |
| TRMT61B |
| TRUB2 |
| TSKU |
| TSN |
| TSNAX |
| TSPAN6 |
| TSPO |
| TSPYL1 |
| TTC14 |
| TTC39B |
| TTC9C |
| TTLL12 |
| TTLL3 |
| TUBA1C |
| TUBB2B |
| TUBB3 |
| TUBG1 |
| TUT1 |
| TWF2 |
| TYRO3 |
| TYW1 |
| U2AF2 |

| |
|---|
| UACA |
| UBA5 |
| UBAP2L |
| UBC |
| UBD |
| UBE2B |
| UBE2C |
| UBE2E1 |
| UBE2L3 |
| UBE2S |
| UBE2W |
| UBE2Z |
| UBE3B |
| UBE4B |
| UBN2 |
| UBQLN4 |
| UBR5 |
| UBXN2A |
| UCHL1 |
| UCK1 |
| UCK2 |
| UFM1 |
| UGDH |
| UHRF1 |
| UHRF2 |
| UMPS |
| UNC13D |
| UPF1 |
| UPK3A |
| UPP1 |
| UQCRC1 |
| UROD |
| USF2 |
| USO1 |
| USP16 |
| USP24 |
| USP25 |
| USP38 |
| USP40 |
| USP9X |
| USPL1 |
| UTP15 |
| VAPA |
| VCAM1 |
| VGLL4 |

| |
|---|
| VIM |
| VPS18 |
| VPS29 |
| VPS37A |
| VPS45 |
| VPS4A |
| VPS72 |
| VRK1 |
| WASF1 |
| WASF2 |
| WDFY4 |
| WDHD1 |
| WDR11 |
| WDR36 |
| WDR48 |
| WDR59 |
| WDR75 |
| WDR76 |
| WDR77 |
| WDR83 |
| WEE1 |
| WIPI1 |
| WNK1 |
| XDH |
| XKR8 |
| XPO5 |
| YBX1 |
| YEATS4 |
| YRDC |
| YTHDF3 |
| YWHAB |
| YWHAE |
| YWHAG |
| YWHAH |
| ZBTB20 |
| ZBTB4 |
| ZBTB45 |
| ZBTB8OS |
| ZBTB9 |
| ZC3H7B |
| ZCCHC10 |
| ZCCHC8 |
| ZCWPW1 |
| ZFAND2A |
| ZFAND2B |

| |
|---|
| ZFPL1 |
| ZFR |
| ZFYVE21 |
| ZMAT1 |
| ZMIZ2 |
| ZMYM1 |
| ZMYND8 |
| ZNF12 |
| ZNF133 |
| ZNF19 |
| ZNF205 |
| ZNF207 |
| ZNF24 |
| ZNF280B |
| ZNF286A |
| ZNF287 |
| ZNF319 |
| ZNF341 |
| ZNF384 |
| ZNF394 |
| ZNF407 |
| ZNF408 |
| ZNF462 |
| ZNF512 |
| ZNF526 |
| ZNF536 |
| ZNF555 |
| ZNF574 |
| ZNF618 |
| ZNF638 |
| ZNF711 |
| ZNF75D |
| ZNF768 |
| ZNF79 |
| ZNFX1 |
| ZNHIT1 |
| ZNHIT2 |
| ZNHIT3 |
| ZRANB2 |
| ZWILCH |
| ZYG11B |
| ZZZ3 |
